# HY5 enhances *Arabidopsis* tolerance to combined high light and heat stress by coordinating photoprotection and hormone signaling

**DOI:** 10.64898/2025.12.08.693111

**Authors:** Damián Balfagón, Clara Segarra-Medina, David Chávez-Jácome, Tadeu dos Reis de Oliveira, Claudete Santa-Catarina, Vanildo Silveira

## Abstract

Plants are frequently exposed to multiple simultaneous environmental stresses, yet the mechanisms that underlie their tolerance to such combinations remain poorly understood. High light (HL) and heat stress (HS) are two major abiotic factors that commonly co-occur in nature and severely impair photosynthetic performance when combined. The bZIP transcription factor HY5 is a well-known integrator of light and temperature cues, but its role under combined HLHS stress remains largely unexplored. Here, we investigated the role of HY5 in *Arabidopsis* tolerance to HLHS using wild-type (Col-0), HY5-deficient (*hy5-215*), and HY5-overexpressing (HY5OX) lines. Physiological and biochemical analyses revealed that HY5OX plants maintained higher photosynthetic efficiency, lower membrane damage, and improved leaf health under HLHS, while *hy5-215* mutants were hypersensitive. Proteomic profiling showed that HLHS induced distinct HY5-dependent changes in the accumulation of photosynthesis-related proteins, particularly Photosystem II core subunits D1 and D2. NPQ4/PsbS, a key component of non-photochemical quenching (NPQ), was identified as a direct downstream target of HY5, with impaired NPQ activation in *hy5-215* and *npq4-1* mutants correlating with higher Y(NO), lower F_v_/F_m_, and increased oxidative damage. Hormonal profiling further revealed that HY5 is required for proper ABA and JA signaling under HLHS: *hy5-215* mutants failed to induce ABA accumulation and showed disrupted JA signaling despite strong hormone accumulation. Our findings highlight HY5 as a central regulator of tolerance to combined HLHS stress, acting through the transcriptional coordination of photoprotective proteins and hormonal signaling networks. This work provides new insights into how plants integrate environmental and hormonal cues to protect the photosynthetic machinery under multifactorial stress.

## Introduction

Light is one of the primary environmental cues for plant growth and development (Zhang et al., 2024). Plants absorb light energy through chloroplast photosystems to drive photosynthesis, but excessive irradiance progressively saturates photosystem reaction centers. This saturation leaves excess excited energy that can damage the photosynthetic apparatus (particularly membrane components) and thereby diminish photosynthetic capacity (Zhang et al., 2024). Similarly, elevated temperatures (heat stress) adversely affect key photosynthetic processes including carbon fixation (Calvin cycle), electron transport, photochemical reactions, and thylakoid membrane stability (Hu et al., 2020; Zahra et al., 2023). Heat stress triggers overproduction of reactive oxygen species (ROS), which further impairs the photosynthetic machinery by reducing electron transport rates, inactivating Photosystems II (PSII) and I, and degrading essential proteins and pigments (Pospíšil, 2016).

Notably, recent studies have shown that combining high light (HL) and heat stress (HS) aggravates their impacts on plants beyond an additive effect (Balfagón et al., 2019; Balfagón, Zandalinas, et al., 2022; Zhou et al., 2020). Simultaneous exposure to both stresses causes severe physiological and molecular damage that can compromise plant development and survival, even when each stress is at a moderate level that would be tolerated individually (Balfagón et al., 2019). In *Arabidopsis*, for example, the high light and heat combination (HLHS) leads to irreversible injury to PSII (e.g. loss of the D1 protein) and invokes unique stress-response pathways not activated by either stress alone (Balfagón et al., 2019; Balfagón, Zandalinas, et al., 2022). Both high irradiance and heat stress ultimately converge on disrupting photosynthetic proteins and the chloroplast electron transport chain, underscoring the importance of studying their combined effects in order to understand and improve plant resilience under multiple concurrent stresses (Balfagón, Zandalinas, et al., 2022; Nishiyama et al., 2011).

Non-photochemical quenching (NPQ) is a critical photoprotective mechanism that allows plants to safely dissipate excess absorbed light energy as heat (Ruban, 2016). NPQ becomes activated when light absorption exceeds the photosynthetic capacity for energy utilization—either due to high irradiance or when PSII reaction centers are saturated or restricted by environmental constraints such as elevated temperatures (Müller et al., 2001; Qiu et al., 2025). This mechanism is triggered by the generation of a proton gradient (ΔpH) across the thylakoid membrane, which leads to the protonation of the PSII-associated PsbS protein and activation of the enzyme violaxanthin de-epoxidase (VDE). Together, protonated PsbS and zeaxanthin accumulation promote the energy-dependent quenching component (qE), which rapidly converts excess excitation energy into heat, thus preventing damage to the photosynthetic apparatus (Ruban, 2016). The physiological relevance of NPQ has been well documented in model species such as *Arabidopsis thaliana*. Mutants impaired in NPQ function (e.g. *npq1*, *npq4*) display increased susceptibility to photoinhibition and PSII damage under HL compared with wild-type plants, underscoring the essential role of NPQ in mitigating light-induced stress (Müller et al., 2001; Niyogi et al., 2001). By preventing overexcitation and limiting the formation of reactive oxygen species (ROS) within the chloroplast, NPQ serves as a vital component of the plant’s defense strategy against photooxidative stress. Importantly, this mechanism not only contributes to acclimation under HL but also enhances tolerance to other abiotic stressors such as heat, which can exacerbate the imbalance between energy absorption and consumption in the photosynthetic machinery (Foyer, 2018; Takahashi & Badger, 2011).

Elongated Hypocotyl 5 (HY5) is a highly conserved basic leucine zipper (bZIP) transcription factor broadly recognized as a master regulator of light signaling in plants. Acting downstream of multiple photoreceptors, HY5 promotes photomorphogenesis and regulates the expression of approximately one-third of the *Arabidopsis thaliana* genome (F. Wang et al., 2018; Xiao et al., 2022). HY5 orchestrates transcriptional programs that drive the assembly of the photosynthetic machinery, the biosynthesis of light-harvesting pigments (e.g. chlorophylls and carotenoids), and chloroplast development (Toledo-Ortiz et al., 2014). Under HL conditions, HY5 plays a central role in stress acclimation by inducing protective gene expression programs that sustain photosynthetic efficiency. Notably, HY5 directly activates genes such as *PGR5* (Proton Gradient Regulation 5) and *VDE* (Violaxanthin De-epoxidase), thereby enhancing cyclic electron flow and xanthophyll cycle activity. These changes promote NPQ, mitigating photoinhibition and oxidative damage under excess light (Jiang et al., 2020). In addition, HY5 modulates the expression of numerous light-responsive genes to optimize both light capture and photoprotection under fluctuating irradiance. Accordingly, *hy5* loss-of-function mutants exhibit decreased accumulation of photoprotective pigments, reduced NPQ capacity, and heightened sensitivity to photo-oxidative damage (Toledo-Ortiz et al., 2014; Wang et al., 2018) highlighting the essential role of HY5 in orchestrating chlorophyll-binding proteins, antioxidant systems, and PSII repair components to enhance HL tolerance. HY5 also integrates temperature signals during heat stress. Emerging evidence suggests that HY5 acts as a key regulatory node in a heat response pathway that is independent of classical thermomorphogenesis (Xiao et al., 2022). Under acute heat stress, particularly in blue light, HY5 undergoes accelerated COP1-dependent degradation, relieving repression of heat-responsive transcription factors such as *HsfA2* and their downstream effectors (Liu et al., 2025; Yang et al., 2023). This mechanism reveals a dual role for HY5: while it promotes photoprotection under HL, it also fine-tunes thermotolerance by controlling the timely induction of heat shock proteins and other protective elements. Together, these findings establish HY5 as a central integrator of light and temperature cues, modulating gene expression to optimize plant performance under multifactorial environmental stress.

To cope with combined environmental stress, plants activate complex hormonal networks in which abscisic acid (ABA) and jasmonic acid (JA) play central roles. Both hormones accumulate under HL and HS—particularly when both occur simultaneously—through enhanced biosynthesis triggered by osmotic and oxidative cues for ABA, and chloroplast lipid peroxidation for JA (Balfagón et al., 2019; Müller & Munné-Bosch, 2021). ABA acts as a major coordinator of abiotic stress tolerance. Under HL, ABA synthesized in vascular tissues is rapidly transported to distal cells, where it closes stomata to reduce water loss and induces antioxidant defenses such as *APX2* (*ascorbate peroxidase 2*). Chloroplast-derived H₂O₂ amplifies these signals, promoting the dissipation of excess excitation energy and reducing photooxidative damage (Müller & Munné-Bosch, 2021). During heat stress, ABA signaling also contributes to thermotolerance by activating ABA-responsive transcription factors and heat shock proteins (Huang et al., 2016; X. Wang et al., 2017). JA, classically linked to defense and wounding, also mediates acclimation to combined HL and HS stress (HLHS). This dual stress induces strong JA and JA-Ile accumulation and upregulates thousands of JA-dependent genes (Balfagón et al., 2019). JA-deficient mutants display severe susceptibility to HLHS, partly due to the loss of repression control on stress-protective genes such as those regulated by *WRKY48* (Balfagón et al., 2024). Together, ABA and JA integrate with light-signaling networks to fine-tune photosynthetic performance, enhance photoprotection, and strengthen plant tolerance to multifactorial environmental stress.

In this study, we investigated the role of HY5 in *Arabidopsis* tolerance to combined HLHS by integrating physiological, molecular, and biochemical approaches. Using wild-type (Col-0), the HY5-deficient mutant *hy5-215*, and the overexpressor HY5OX, we performed a comparative analysis including proteomics, hormonal profiling, gene expression, chlorophyll fluorescence parameters, and stress tolerance assays. This multifaceted approach aimed to elucidate how HY5 contributes to the orchestration of acclimation responses under this common environmental stress conditions.

## Materials and methods

### Plant Material and Growth Conditions

*Arabidopsis thaliana* wild-type Col (var Columbia-0), *hy5-215* mutant and over expressing line HY5OX (35S::HA-HY5) previously described (Toledo-Ortiz et al., 2014), and *npq4-1* mutant (Li et al., 2000) were used. Plants were grown in peat pellets (Jiffy-7; http://www.jiffygroup.com/) at 23°C under long-day growth conditions (12-h light from 7 AM to 7 PM; 80 μmol m^−2^ sec^−1^/12-h dark from 7 PM to 7 AM).

### Stress Treatments

Individual HL and HS, and HLHS were applied in parallel using 28-day-old *Arabidopsis* plants. HL was applied by exposing 28-d–old plants to 900 μmol m^−2^ s^−1^ at 23°C for 6 h. HS was imposed by transferring 28-d–old plants to 38°C, 80 μmol m^−2^ s^−1^, for 6 h. HLHS was performed by simultaneously subjecting plants to 900 μmol m^−2^ s^−1^ light and 38°C for 6 h. CT plants were maintained at 80 μmol m^−2^ s^−1^, 23°C. After the stress treatments, plants of each treatment were harvested and immediately frozen with N_2_ for use in subsequent analysis. At least five plants of each group were allowed to recover under controlled conditions during 24 h to quantify percentage of healthy leaves (Balfagón, Gómez-Cadenas, et al., 2022). All experiments were carried out at the same time of the light cycle (from 9 am to 3 pm) and were repeated at least three times.

### Malondialdehyde Analysis

Malondialdehyde (MDA) content was quantified using a modified thiobarbituric acid reactive substances (TBARS) assay, based on spectrophotometric detection of MDA-TBA adducts, following (Hodges et al. 1999) with some modifications. Ground frozen rosettes (0.1 g approximately) were homogenized in 2 mL 80 % cold ethanol by sonication for 30 min. Homogenates were centrifuged 12000 g for 10 min and different aliquots of the supernatant were mixed either with 20 % trichloroacetic acid or with a mixture of 20 % trichloroacetic acid and 0.5 % thiobarbituric acid. Both mixtures were incubated in a water bath at 90 °C for 1 h. After that, samples were cooled in an ice bath and centrifuged at 2000 g for 5 min at 4 °C. The absorbance at 440, 534 and 600 nm of the supernatant was read against a blank.

### Photosynthetic Parameters

Y(II), Y(NO), NPQ and F_v_/F_m_ were measured using a portable fluorometer (model no. 110/S FluorPen; PSI, Czech Republic). Y(NO), NPQ and F_v_/F_m_ were measured after a 60 minutes period of dark adaptation following the end of the stress treatments. Photosynthetic measurements were taken for at least 6 plants using two fully expanded young leaves per plant for each stress treatment, and each experiment was repeated at least three times.

Y(NO) and NPQ were recorded with a FluorPen FP 110 using the predefined NPQ1 protocol. After a brief measuring light to obtain minimum fluorescence in darkness (F₀), a short saturating pulse was applied to determine maximum fluorescence in darkness (Fₘ). Following a short dark interval, the sample was exposed to actinic light for 60 s while five saturating pulses were superimposed to probe non-photochemical quenching and effective PSII yield. In NPQ1, the first light-phase pulse occurs at 7 s and subsequent pulses every 12 s; we refer to these light-phase sampling points as L1–L4 and the steady-state point (fifth pulse) as Lss. After switching off the actinic light, dark recovery proceeded for 88 s with three saturating pulses (first at 11 s, then every 26 s), denoted D1–D3. Non-photochemical quenching was computed as NPQ = (Fₘ − Fₘ′)/Fₘ′ (reported as L1–L4, Lss, D1–D3). Y(NO) was derived with the standard “puddle” model using the FluorPen outputs (NPQ and qP) and dark-adapted F₀ and Fₘ: Y(NO) = 1 / (1 + NPQ + qP x (Fₘ / F₀ - 1).

### Protein Extraction and Digestion

Proteins were extracted from lyophilized rosette tissues of *Arabidopsis thaliana* (Col-0, *hy5-215*, and HY5OX) as described by Oliveira et al. (2020), with minor modifications. Three biological replicates, each consisting of pooled tissue from five plants, were analyzed. Proteins were precipitated using the methanol/chloroform method (Nanjo et al., 2012), washed with cold acetone containing DTT, and resuspended in urea/thiourea extraction buffer. Protein concentration was determined using the Bradford assay (Bio-Rad, USA). A total of 100 μg of protein per replicate was digested with trypsin following the filter-aided sample preparation (FASP) procedure described by Wiśniewski et al. (2009), with modifications from Oliveira et al. (2022). The resulting peptides were quantified and stored at −80 °C until LC–MS/MS analysis.

### Bottom-Up Proteomics Analysis

A total of 2 μg of peptides from each replicate was analyzed on a nanoAcquity UPLC M-Class system coupled to a Synapt G2-Si HDMS mass spectrometer (Waters, Manchester, UK) using data-independent acquisition (DIA, HDMSE) mode, as described by Oliveira et al. (2022). Peptides were separated on a reversed-phase C18 column with a linear acetonitrile gradient containing 0.1% formic acid. MS data were acquired with ion-mobility separation at 35,000 FWHM over the 50–2000 *m/z* range. Lock-mass correction with [Glu¹]-fibrinopeptide B was applied during acquisition, and spectra were processed using MassLynx v.4.1 (Waters).

### Data Processing and Quantification

MS/MS spectra were processed using ProteinLynx Global SERVER (PLGS) v.3.02 (Waters) against the *A. thaliana* Araport11 protein database (release 20220914) including common contaminants. Search parameters included one missed cleavage, fixed carbamidomethylation, variable oxidation and phosphorylation, and a ≤1% false discovery rate (FDR). Protein identification required at least two peptides and ≤1% FDR. Only proteins consistently identified across replicates were considered for statistical comparison. Label-free quantification was performed using the TOP3 approach and ISOQuant v.1.7 (Distler et al., 2014). Differentially accumulated proteins were determined using Student’s *t*-test (P < 0.05).

### Abscisic Acid and Jasmonic Acid Analysis

Abscisic acid (ABA) and jasmonic acid (JA) extraction and analysis were carried out as described in (Balfagón et al. 2019). Chromatographic separations were performed on a reversed-phase C18 column (Gravity, 50 × 2.1 mm, 1.6-μm particle size, Luna Omega, Phenomenex, Torrance, CA, USA) inserted on an ultra-performance UPLC system (Xevo TQ-S, Waters Corp., Milford, MA, USA). Hormones were quantified with a triple quadrupole mass spectrometer connected online to the output of the column through an orthogonal Z-spray electrospray ion source. Results were processed using Masslynx v. software, and the phytohormone content was quantified with a standard curve prepared with commercial standards as described in (Balfagón et al., 2019). [^2^H_6_]-ABA and dehydrojasmonic acid (DHJA) were used as internal controls.

### RT-qPCR analysis

Relative expression analysis by RT-qPCR was performed according to (Balfagón et al. 2021) by using a StepOne Real-Time PCR system (Applied Biosystems, Carlsbad, CA, USA) and gene-specific primers (Table S1). The expression levels of all genes were normalized against the expression of *EF1α*.

### Western Blot Analysis

Protein isolation and Western blot procedures were carried out based on previously described methods (Balfagón et al., 2021).

For western blot analysis, 20 μg of protein from each sample were separated using Mini-PROTEAN TGX Stain-Free precast gels (#4568,044, Bio-Rad, USA) with Tris running buffer and transferred to a PVDF membrane following the manufacturer’s protocol (Bio-Rad, USA). Membranes were blocked for 1 hour with 3 % BSA and incubated overnight at 4 °C with primary antibodies against PsbS protein (#AS09 533, Agrisera, Vännäs, Sweden). After washing, membranes were incubated for 1 hour with goat anti-rabbit IgG (H&L) HRP-conjugated secondary antibodies (1:25,000).

Chemiluminescent detection was performed using the ECL2 substrate (Thermo Scientific, USA), and protein bands were visualized with the ImageQuant LAS 500 imaging system (GE Healthcare, Sweden). Stain-Free total protein staining (Bio-Rad, USA) was used as a loading control as described by (Gilda & Gomes 2013). Protein bands were visualized using the ChemiDoc Go System (Bio-Rad, USA).

### Statistical Analysis

All experiments were repeated at least three times. Results are presented as the mean ± SE. Two-way ANOVA followed using a Tukey post hoc test (P < 0.05) or two-tailed Student’s *t*-test (asterisks denote statistical significance at P < 0.05 with respect to CT) were used to discriminate statistical differences. Functional annotations and overrepresentation of GO terms (P < 0.05) were performed using the program STRING (https://string-db.org/; (Szklarczyk et al. 2023) PCA was performed by means of the Soft Independent Modeling of Class Analogy software v. 13.0.3.0, using the log_2_ transformed data and unit variance normalization.

## Results

### HY5 positively influences plant tolerance to combined high light and heat stress

To investigate the role of HY5 in plant acclimation to HLHS, we exposed *Arabidopsis thaliana* wild-type (Col-0), the HY5-deficient mutant (*hy5-215*), and a 35S::HA-HY5 overexpressor line (HY5OX) to HL, HS, and HLHS conditions. Neither HL nor HS alone caused significant leaf damage during the treatment period in any genotype. In contrast, the HLHS combination markedly reduced the percentage of healthy leaves in Col-0 and *hy5-215* plants to 76.7% and 63.9%, respectively, whereas HY5OX plants maintained leaf integrity under all environmental conditions (Figure 1A, B).

**Figure 1.**
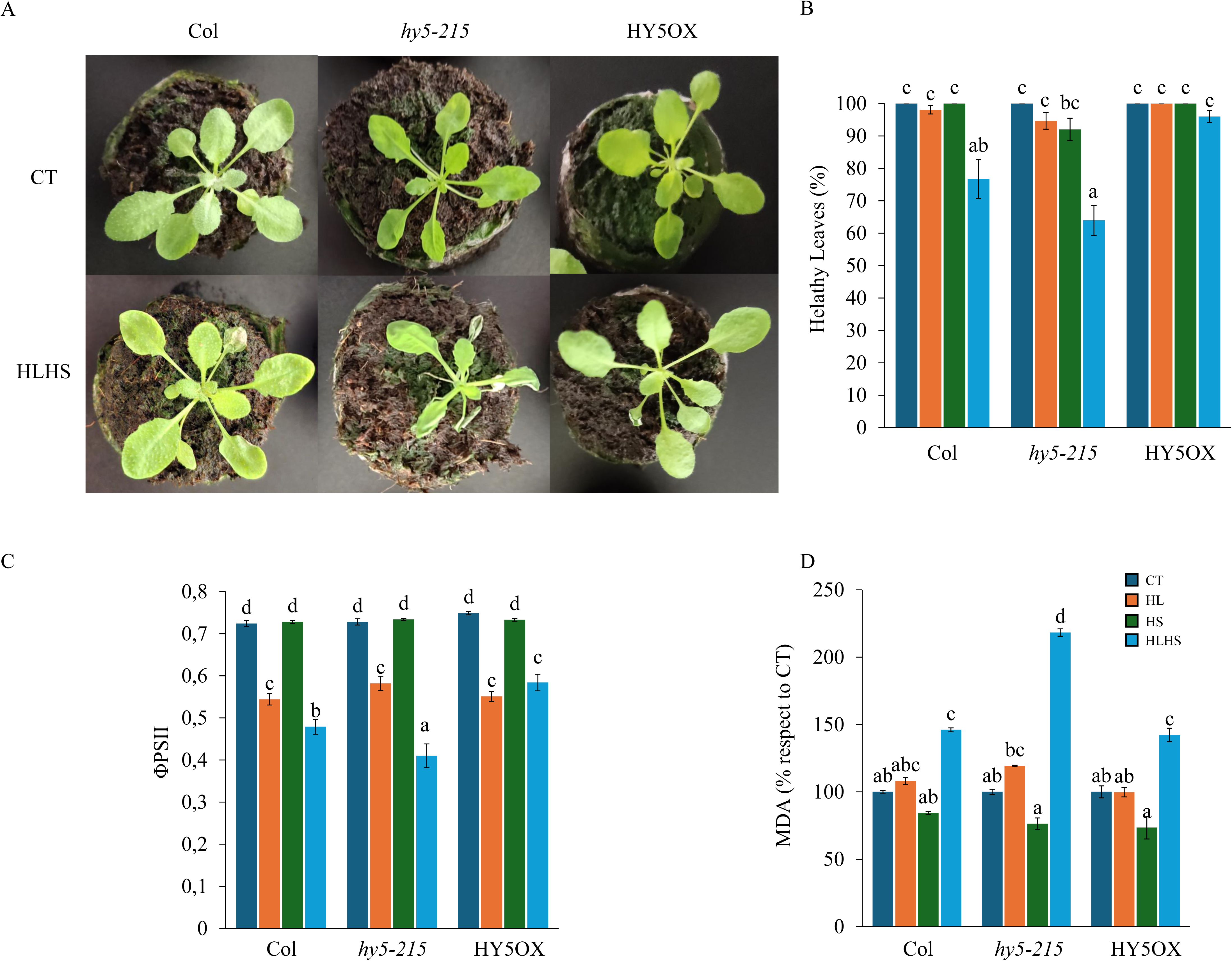
Leaf damage, PSII efficiency, and oxidative stress in Col-0, *hy5-215*, and HY5OX plants exposed to high light (HL), heat stress (HS), and combined HL+HS. (A) Representative images of plants after 24 h of control (CT), high light (HL), heat stress (HS), or combined HLHS treatment. (B) Percentage of healthy leaves after each treatment. (C) Maximum quantum yield of PSII (ΦPSII) measured immediately after stress exposure. (D) Malondialdehyde (MDA) content in leaves as a marker of lipid peroxidation. Error bars represent mean ± standard error (SE) (n = 9).

To assess photosynthetic performance, the quantum yield of PSII (ΦPSII) was measured immediately after stress exposure (Figure 1C). HS alone had no effect on ΦPSII, and HL induced a similar reduction across all genotypes. However, HLHS led to a more pronounced decrease in ΦPSII in *hy5-215* compared to Col-0 and HY5OX. Notably, ΦPSII levels in HY5OX remained comparable under HL and HLHS, suggesting improved resilience, while Col-0 showed an intermediate response.

Oxidative damage to cellular membranes was further evaluated by quantifying malondialdehyde (MDA) levels (Figure 1D). HLHS significantly increased MDA accumulation in all genotypes, but *hy5-215* exhibited a notably higher increase compared to Col-0 and HY5OX.

Taken together, these findings indicate that HY5 plays a positive role in protecting *Arabidopsis* plants against the detrimental effects of combined high light and heat stress, likely by enhancing photoprotection and limiting oxidative damage.

### HY5-dependent proteomic alterations under high light, heat and combined stress conditions

To explore the proteome changes of plants associated with HY5 function HL, HS, and HLHS, we performed a proteomic analysis of Col-0, *hy5-215*, and HY5OX plants using GC-MS (Table S2-S10). Principal component analysis (PCA; Figure 2A) revealed that the primary source of variance (PC1, 22.86%) corresponded to proteomic shifts induced by stress treatments, with HLHS triggering the strongest divergence from control profiles across all genotypes. The second source of variation (PC2, 15.84%) was attributable to the genotypic background, indicating a strong impact of HY5 overexpression or deficiency on the global proteome under both control and stress conditions. Interestingly, HY5OX exhibited a greater proteomic divergence from Col-0 than *hy5-215* (Figure 2A).

**Figure 2.**
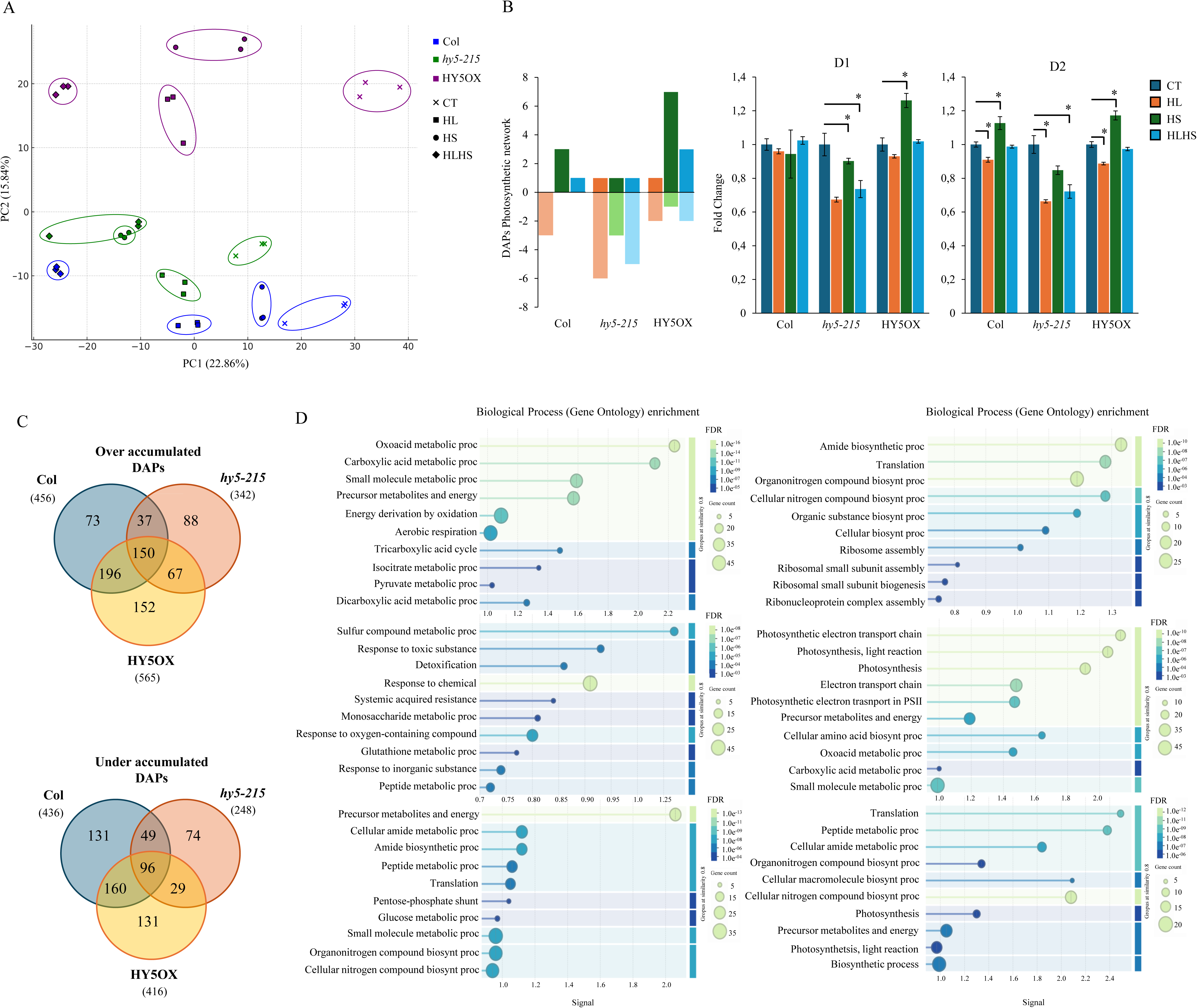
Proteomic responses and photosynthetic protein abundance in Col-0, *hy5-215*, and HY5OX plants under control and stress conditions. (A) Principal component analysis (PCA) of the proteomic profiles in plants subjected to control (CT), high light (HL), heat stress (HS), or combined (HLHS) conditions. (B) Number of photosynthesis-related proteins differentially accumulated (over-accumulated, OA; or under-accumulated, UA) in each genotype and treatment. (C) Venn diagrams showing the number of OA (left) and UA (right) proteins in response to HLHS in Col-0, hy5-215, and HY5OX. (D) Gene ontology enrichment analysis (Biological Process category) of OA proteins (left panel) and UA proteins (right panel) in response to HLHS. Enrichment terms are shown for each genotype: Col-0, *hy5-215*, and HY5OX (top to bottom). Error bars represent mean ± standard error (SE) (n = 3).

Quantification of differentially accumulated proteins (DAPs) under HLHS revealed a clear HY5-dependent pattern. Specifically, 73, 88, and 152 proteins were uniquely over-accumulated (OA) in Col-0, *hy5-215*, and HY5OX, respectively, while 131, 74, and 131 proteins were uniquely under-accumulated (UA) (Figure 3D). Gene ontology (GO) enrichment analysis of these DAPs indicated that HY5 modulates distinct biological processes under stress. In Col-0, OA proteins were mainly associated with metabolic and energy pathways, while UA proteins were enriched in translation and ribosome-related processes. In *hy5-215*, OA proteins were linked to detoxification and response to reactive oxygen species (ROS), whereas UA proteins were enriched in photosynthesis- and electron transport-related processes. HY5OX plants showed a mixture of enriched GO terms, including metabolism, translation, and photosynthesis-related categories, although photosynthetic terms were less prominent than in *hy5-215*.

**Figure 3.**
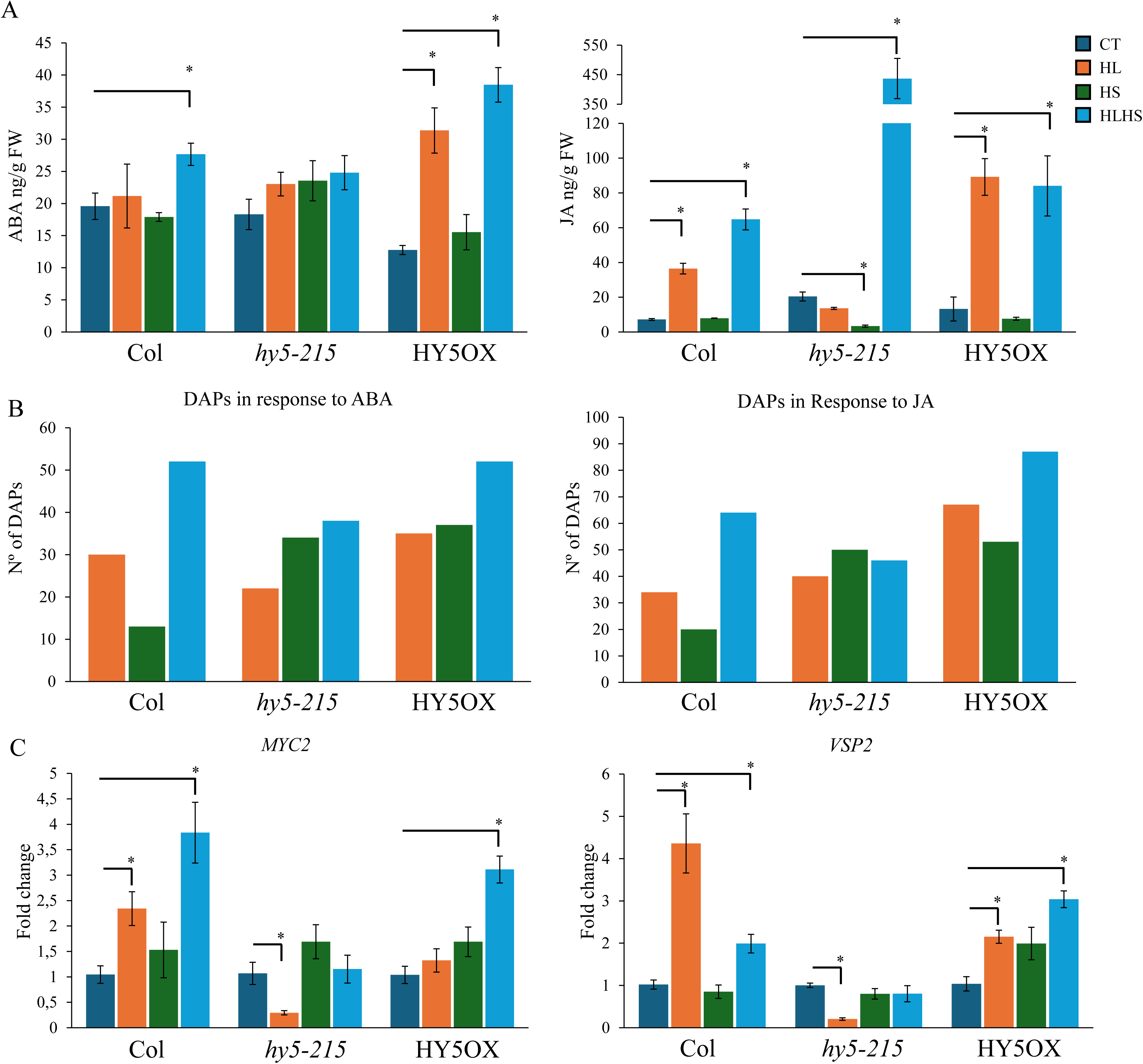
Hormonal profiling, responsive protein accumulation, and gene expression in Col-0, *hy5-215*, and *HY5OX* plants under control and stress conditions. (A) Leaf contents of abscisic acid (ABA) and jasmonic acid (JA) measured in plants subjected to control (CT), high light (HL), heat stress (HS), and combined high light and heat stress (HLHS). (B) Number of differentially accumulated proteins (DAPs) associated with ABA (left) and JA (right) signaling pathways identified in each genotype and treatment. (C) Expression levels of JA-responsive genes (*MYC2* and *VSP2*) measured by qRT-PCR in plants subjected to CT, HL, HS, or HLHS conditions. Error bars represent mean ± standard error (SE) (n = 9).

Given the central role of photosynthetic proteins in acclimation to HL and HLHS, we examined photosynthesis-related DAPs in detail across stress treatments (Figure 2B). *hy5-215* plants exhibited a general trend toward under-accumulation of photosynthetic proteins across all conditions. In Col-0, HL caused under-accumulation of three proteins, while HS and HLHS resulted in over-accumulation of three and one proteins, respectively. HY5OX plants showed over-accumulation of 1, 7, and 3 proteins and under-accumulation of 2, 1, and 2 proteins after HL, HS, and HLHS, respectively.

To further assess the impact on the photosynthetic apparatus, we analyzed the abundance of D1 and D2 core proteins of PSII, key targets of stress-induced photodamage (Figure 2B). HL treatment caused a reduction in D1 protein only in *hy5-215*, and a decrease in D2 across all genotypes. HS led to D1 accumulation in HY5OX and D2 accumulation in Col-0 and HY5OX. Under HLHS, however, only *hy5-215* plants showed a significant reduction in both D1 and D2 protein levels, while Col-0 and HY5OX maintained D1/D2 levels similar to control conditions.

These results demonstrate that HY5 influences the proteome of *Arabidopsis* in response to HL, HS, and especially HLHS, with a marked effect on proteins involved in photosynthesis, detoxification, and stress response. The maintenance of photosynthetic protein abundance in HY5OX plants under combined stress further supports a protective role of HY5 in chloroplast function.

### High light and heat stress combination triggers HY5-dependent ABA and JA signaling

To assess the impact of HY5 on hormonal responses under stress, we analyzed the content of abscisic acid (ABA) and jasmonic acid (JA) in Col-0, *hy5-215*, and HY5OX plants subjected to HL, HS, and HLHS treatments (Figure 3A). In Col-0, ABA levels significantly increased in response to HLHS. HY5OX plants showed elevated ABA accumulation under both HL and HLHS, while *hy5-215* mutants exhibited no significant changes in ABA levels under any stress condition. Consistent with these findings, the number of differentially accumulated proteins (DAPs) associated with ABA signaling was highest under HLHS in all genotypes. However, the number of ABA-related DAPs was greater in Col-0 and HY5OX (52) compared to *hy5-215* (38) (Figure 3B), suggesting an attenuated ABA signaling response in the HY5-deficient background.

JA levels increased in Col-0 and HY5OX plants under HL and HLHS, but remained unchanged under HS. In contrast, *hy5-215* plants exhibited a distinct pattern: JA levels were unchanged under HL, decreased under HS, and strongly increased under HLHS (Figure 3A). However, this accumulation did not correlate with the number of JA-responsive DAPs. While HLHS induced 87 and 64 JA-related DAPs in HY5OX and Col-0, respectively, only 46 were detected in *hy5-215*, despite its elevated JA content (Figure 3B). Additionally, HL and HS induced a higher number of JA-related DAPs in *hy5-215* than in Col-0, but this trend was not maintained under HLHS. To further investigate the discrepancy between JA accumulation and downstream response in *hy5-215*, we analyzed the expression of three JA-responsive genes: *MYC2* and *VSP2* (Figure 3C). Under HLHS, these genes were significantly induced in Col-0 and HY5OX plants, in agreement with JA accumulation. However, no significant upregulation was observed in *hy5-215*, indicating that JA signaling output is compromised in the absence of HY5 despite elevated JA levels.

### NPQ4/PsbS contributes to HLHS tolerance in a HY5-dependent manner

To further explore the proteomic basis of HY5-mediated stress tolerance, we identified proteins that were over-accumulated in Col-0 but not in *hy5-215* under HLHS, and that were also constitutively higher in HY5OX compared to Col-0 under control conditions. This analysis yielded 33 candidate proteins potentially linked to HY5-dependent acclimation (Figure 4A). Gene Ontology enrichment indicated that the most represented term among these proteins was “Photosynthesis,” including six proteins annotated in this category. To validate these findings, we analyzed the expression of the genes encoding the 6 candidate proteins by qRT-PCR under HLHS. Four genes were upregulated in HY5OX, three in Col-0, and only one in *hy5-215*, which displayed downregulation for the remaining genes (Figure 4B), supporting a HY5-dependent transcriptional activation of key protective proteins under combined stress. Among these, NPQ4 (PsbS), a central protein in non-photochemical quenching (NPQ), emerged as a key candidate due to its known role in dissipating excess excitation energy and protecting PSII under HL and HS (Kulasek et al., 2016; Roach & Krieger-Liszkay, 2012). To investigate its role functionally, we analyzed the response of the *npq4-1* mutant to HLHS. *npq4-1* plants exhibited significantly lower PSII efficiency and higher MDA levels compared to Col-0, indicating greater oxidative membrane damage (Figure 4C).

**Figure 4.**
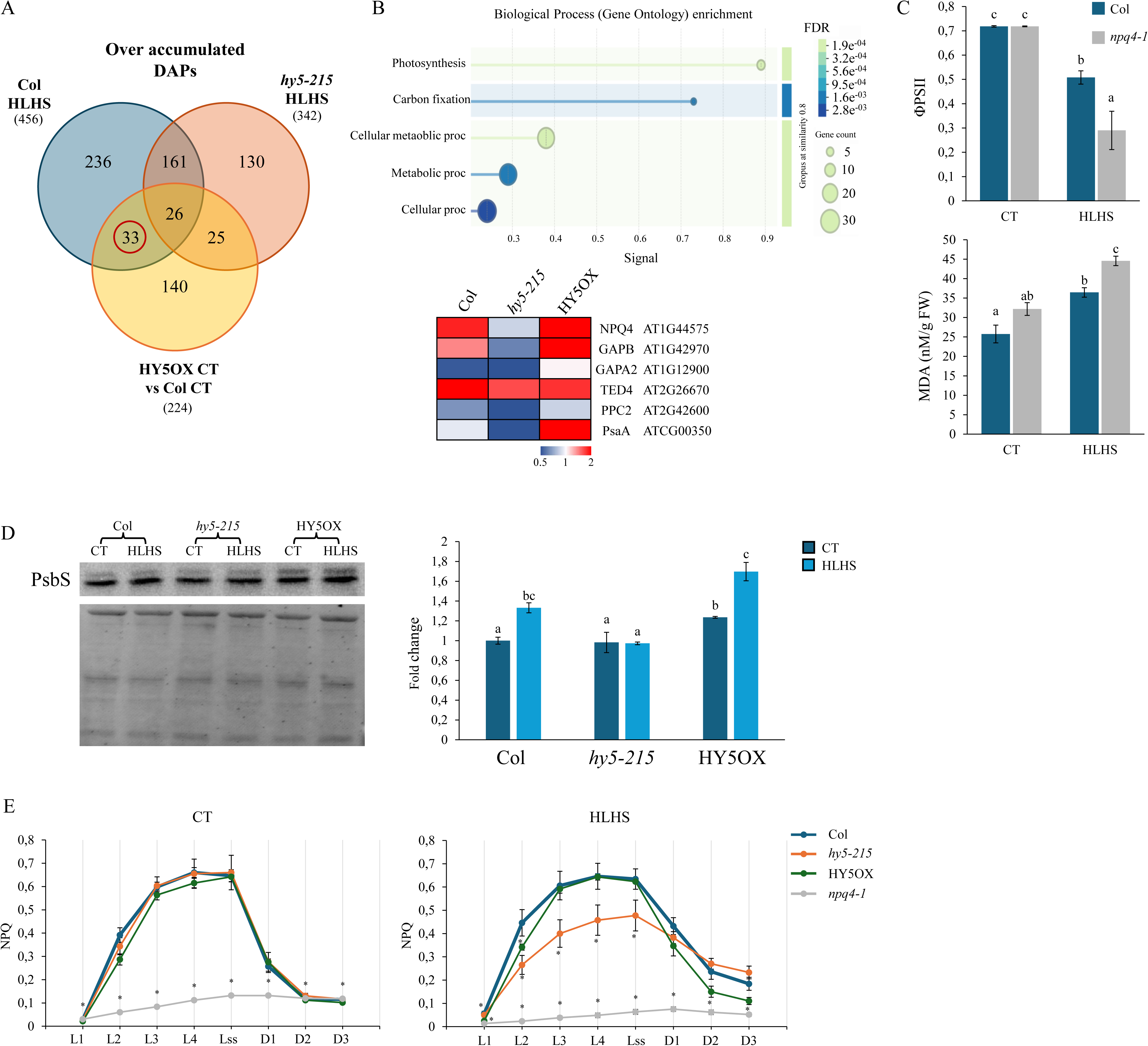
Analysis of NPQ4/PsbS and NPQ-related parameters in Col-0, *hy5-215*, *HY5OX*, and *npq4-1* plants under control and HLHS conditions. (A) Identification of proteins over-accumulated in Col-0 under HLHS, not over-accumulated in *hy5-215*, and constitutively accumulated in HY5OX under control conditions. (B) Top: Gene Ontology (GO) enrichment (Biological Process category) of the 33 proteins identified in panel A. Bottom: Relative expression levels of genes encoding the six proteins annotated under the “Photosynthesis” GO term, measured by qRT-PCR in Col-0, *hy5-215*, and HY5OX under HLHS. (C) PSII quantum yield (ΦPSII) and MDA content in Col-0 and *npq4-1* plants after HLHS stress. (D) Immunoblot detection of PsbS protein and corresponding densitometric quantification in Col-0, *hy5-215*, and HY5OX under CT and HL+HS conditions. (E) NPQ kinetics measured in Col-0, *hy5-215*, HY5OX, and *npq4-1* under CT and HLHS. Error bars represent mean ± standard error (SE) (n = 3).

We next examined NPQ4 transcript and protein levels in Col-0, *hy5-215*, and HY5OX plants subjected to HL, HS, and HLHS. NPQ4 expression was upregulated in Col-0 and HY5OX under HLHS but significantly downregulated in *hy5-215* (Figure 4D and Supplementary Figure S1). Immunoblot analysis confirmed that PsbS protein accumulated under HLHS in Col-0 and HY5OX, but not in *hy5-215*, supporting a role for HY5 in regulating NPQ4 at both transcript and protein levels.

To assess functional consequences, we measured NPQ activation in Col-0, *hy5-215*, HY5OX, and *npq4-1* under control and HLHS conditions. Under control conditions, Col-0, *hy5-215*, and HY5OX displayed similar NPQ induction during actinic light pulses and dark relaxation, while *npq4-1* showed no NPQ activation, as expected due to the absence of NPQ4. After HLHS, NPQ activation was reduced in *hy5-215* compared to Col-0 and HY5OX, particularly during L3, L4 and Lss. HY5OX showed slightly lower NPQ than Col-0 in early pulses but similar values thereafter. As expected, *npq4-1* remained deficient in NPQ under HLHS (Figure 4D). In addition to NPQ, we evaluated the quantum yield of non-regulated energy dissipation [Y(NO)] and maximum PSII efficiency (F_v_/F_m_; Supplementary Figure 2). Under control conditions, all genotypes exhibited similar Y(NO) and F_v_/F_m_ values. However, HLHS triggered significantly higher Y(NO) at time points L2, L3, L4, and Lss in *hy5-215* and *npq4-1*, indicating reduced energy dissipation.

Similarly, the decline in F_v_/F_m_ under HLHS was more pronounced in *hy5-215* and *npq4-1*, reinforcing the role of HY5 and NPQ4 in protecting PSII from photo-oxidative damage under combined stress.

## Discussion

HY5 is a key transcription factor that regulates photoprotective responses by controlling the expression of genes involved in pigment biosynthesis, chloroplast development, and energy dissipation (Gangappa & Botto, 2016; Toledo-Ortiz et al., 2014). Previous studies have shown that HY5 activates genes such as *PGR5* and *VDE*, which are essential for cyclic electron flow and xanthophyll cycle activation under excess light (Jiang et al., 2020). Our proteomic and physiological data now position NPQ4 (PsbS), a key protein in energy dissipation through NPQ, as an additional downstream effector of HY5 during HLHS stress. Beyond its classical role in photomorphogenesis, recent evidence suggests that HY5 also interacts with hormonal signaling pathways to modulate abiotic stress responses (Bian et al., 2022). In particular, ABA and JA have been shown to accumulate under HL and HS and contribute to acclimation by inducing antioxidant defenses and stress-responsive gene expression (Balfagón et al., 2019, 2024; Müller & Munné-Bosch, 2021). Our results suggest that HY5 functions as a central hub integrating light and temperature signals with ABA and JA signaling outputs, thereby coordinating transcriptional and post-transcriptional mechanisms that enhance stress tolerance. Notably, the regulation of NPQ4 (PsbS) and other photosynthesis-related genes and proteins during HLHS stress appears to be HY5-dependent, linking hormonal regulation and transcriptional control to photoprotection and chloroplast stability under HLHS.

The combination of high light and heat stress posed a considerably greater physiological challenge than either stress alone, as shown by the marked reductions in leaf health, PSII efficiency, and increased lipid peroxidation. While HL and HS applied individually had minimal effects on Col-0, *hy5-215*, and HY5OX plants, their combination disproportionately affected Col-0 and was particularly detrimental in the HY5-deficient mutant *hy5-215* (Figure 1). In contrast, HY5OX plants maintained high percentages of healthy leaves, exhibited stable ΦPSII values under HLHS comparable to HL alone, and accumulated less MDA (Figure 1), supporting a role for HY5 in promoting photoprotection and stress tolerance under multifactorial conditions.

This differential tolerance was parallel by major shifts in the proteomic profile. PCA analysis indicated that HLHS was the dominant factor shaping protein accumulation patterns across genotypes, and that HY5OX showed greater divergence from Col-0 than *hy5-215* under both control and stress conditions (Figure 2A). This suggests that HY5 overexpression may drive a basal proteomic reconfiguration that enhances readiness for stress. Notably, GO enrichment analysis revealed that the biological processes most affected by HLHS in *hy5-215* were photosynthesis and electron transport—critical pathways for energy conversion and ROS control (Figure 2D). Proteomic analysis confirmed that *hy5-215* displayed a general under-accumulation of photosynthetic proteins, including the PSII core proteins D1 and D2, under HLHS (Figure 2B). Their depletion likely reflects heightened photodamage and impaired repair capacity in the absence of HY5, which contrasts with their stable accumulation in Col-0 and HY5OX. These findings underscore the essential role of HY5 in maintaining chloroplast function and photosynthetic protein homeostasis under combined stress conditions.

Our results indicate that HY5 may be required for proper activation of ABA and JA signaling pathways under HLHS. In the case of ABA, both Col-0 and HY5OX plants showed a marked increase in ABA content under HLHS, while *hy5-215* mutants failed to accumulate the hormone under any stress condition (Figure 3A). This pattern was mirrored at the proteomic level, where the number of ABA-responsive DAPs was considerably lower in *hy5-215* than in Col-0 or HY5OX, despite exposure to the same stress combination (Figure 3B). These findings suggest that HY5 could be involved not only in modulating ABA-dependent transcriptional responses (as previously suggested by (Chen et al. 2008), but also potentially in facilitating hormone biosynthesis or transport under stress. Given the well-established role of ABA in mediating tolerance to abiotic stresses such as drought, high light, and heat—through stomatal regulation, ROS detoxification, and photoprotective gene expression (Müller & Munné-Bosch, 2021; Sah et al., 2016)—the impaired ABA response in *hy5-215* may contribute to its reduced HLHS tolerance. JA signaling showed a more complex pattern. Strikingly, *hy5-215* plants accumulated extremely high levels of JA under HLHS (Figure 3A), yet this did not translate into increased signaling output: they showed fewer JA-responsive DAPs than Col-0 or HY5OX and failed to induce canonical JA-responsive genes (*MYC2*, *VSP2*; Figures 3B and C). This disconnect suggests that HY5 may be required for efficient transduction of JA signals, potentially through interactions with other transcription factors such as PIFs, MYCs, COI1 or JAZ repressors (Ortigosa et al., 2020; Sun et al., 2025; Yi et al., 2020). One possibility is that, in the absence of HY5, JA signaling becomes decoupled from its canonical downstream pathways. Alternatively, the extreme JA accumulation in *hy5-215* could reflect a shift toward irreversible stress responses such as leaf senescence or programmed cell death, processes in which JA has been implicated under severe stress conditions (Beaugelin et al., 2019; Wasternack & Song, 2016). By contrast, in Col-0 and particularly in HY5OX, JA accumulation under HLHS may act as a signal to activate adaptive responses rather than terminal damage programs. Together, these results suggest that HY5 is not only a regulator of photosynthetic acclimation, but also a critical integrator of hormonal signals under multifactorial stress. However, further studies are needed to elucidate the mechanistic basis of HY5-hormone interactions, especially in terms of direct transcriptional targets, feedback regulation, and interaction with other signaling modules.

Our results provide evidence that HY5 enhances plant tolerance to HLHS by promoting the accumulation and function of NPQ4/PsbS, a central effector of NPQ. NPQ is a fast-acting photoprotective mechanism that dissipates excess excitation energy as heat, preventing overexcitation of PSII and limiting the formation of harmful reactive oxygen species (ROS) under light and thermal stress (Müller et al., 2001; Ruban, 2016). Under HLHS, we observed that HY5OX and Col-0 plants upregulated NPQ4 transcript and protein levels, while *hy5-215* mutants showed significantly reduced expression and failed to accumulate PsbS protein. This pattern was associated with a reduced NPQ capacity in hy5-215, particularly during late actinic light pulses, and with significantly higher Y(NO) values, indicating inefficient energy dissipation and greater excitation pressure on PSII (Kramer et al., 2004). Functionally, these alterations translated into increased oxidative damage in *hy5-215* and *npq4-1* mutants, as shown by elevated MDA levels, greater loss of PSII core proteins D1 and D2, and stronger reductions in F_v_/F_m_ after HLHS. These data support the idea that HY5 contributes to stress tolerance by enabling a PsbS-mediated NPQ response that safeguards PSII integrity. The fact that *npq4-1* phenocopied *hy5-215* in multiple parameters— Y(NO), F_v_/F_m_, and oxidative damage—further underscores the functional relevance of NPQ4 as a downstream effector of HY5. It is well established that PsbS plays a crucial role in rapidly adjusting NPQ in response to changing light environments and protects the thylakoid membrane from photodamage (Li et al., 2000; Roach & Krieger-Liszkay, 2012). Our data now add to this model by placing HY5 as a key upstream regulator of PsbS accumulation under multifactorial stress, linking transcriptional regulation of energy-dissipating proteins with hormonal signaling and proteostasis. The impaired NPQ activation in *hy5-215* likely exacerbates ROS generation and chloroplast dysfunction, contributing to the reduced leaf health and stress performance observed in this genotype.

Overall, our results position HY5 as a central regulator of *Arabidopsis* acclimation to the simultaneous occurrence of high light and heat stress. Through coordinated control of photoprotective mechanisms such as NPQ, regulation of PSII core proteins, and integration with ABA and JA signaling pathways, HY5 contributes to maintaining photosynthetic performance and limiting oxidative damage under multifactorial stress. Notably, we identify NPQ4/PsbS as a key downstream effector of HY5, and provide evidence that HY5 is required for full activation of hormone-responsive gene and protein networks.

---

**Figure S1.**
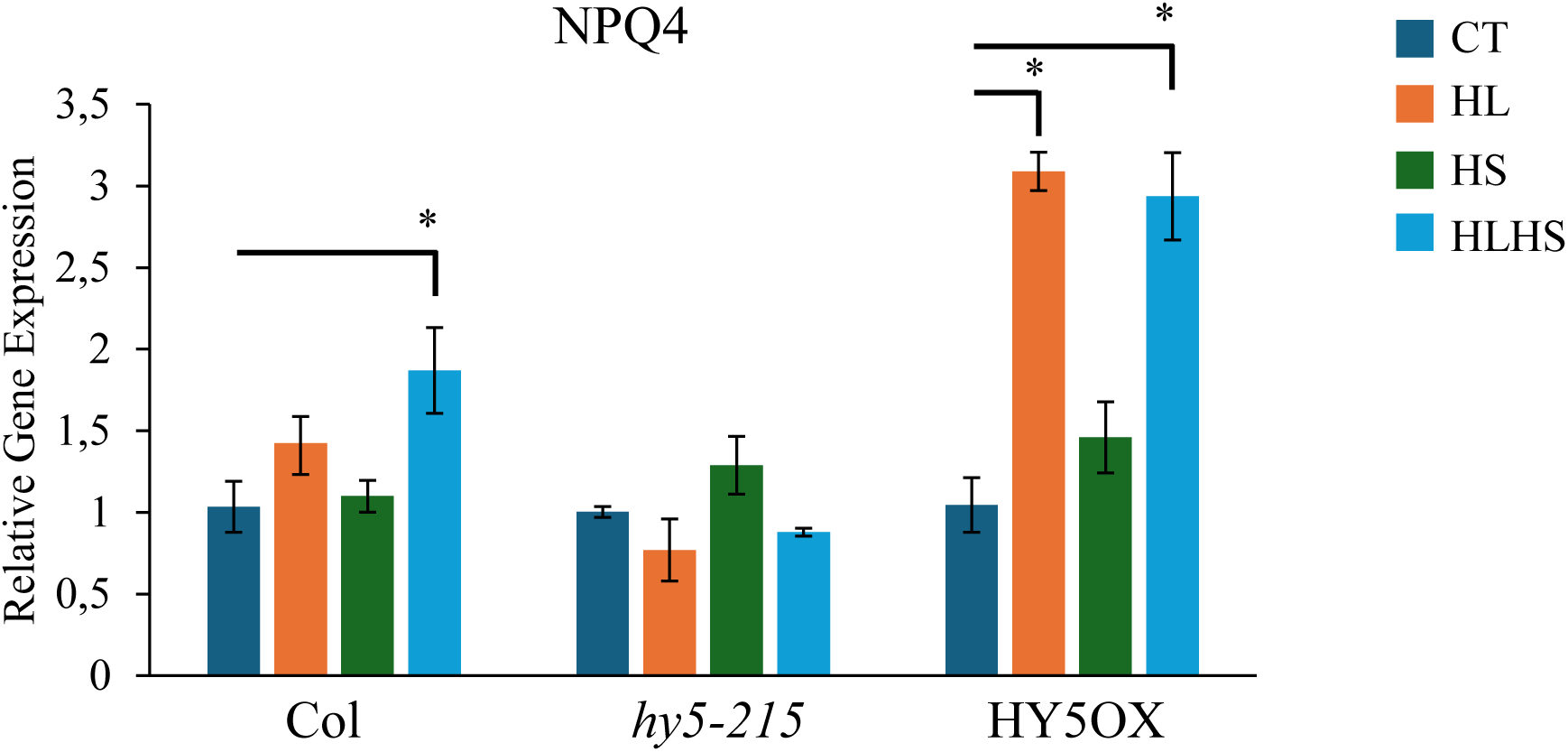
Relative expression levels of the NPQ4 gene in Col-0, *hy5-215*, and HY5OX plants under control (CT), high light (HL), heat stress (HS), and combined high light and heat stress (HLHS) conditions. Expression values were normalized to control levels for each genotype. Error bars represent mean ± standard error (SE) (n = 9).

**Figure S2.**
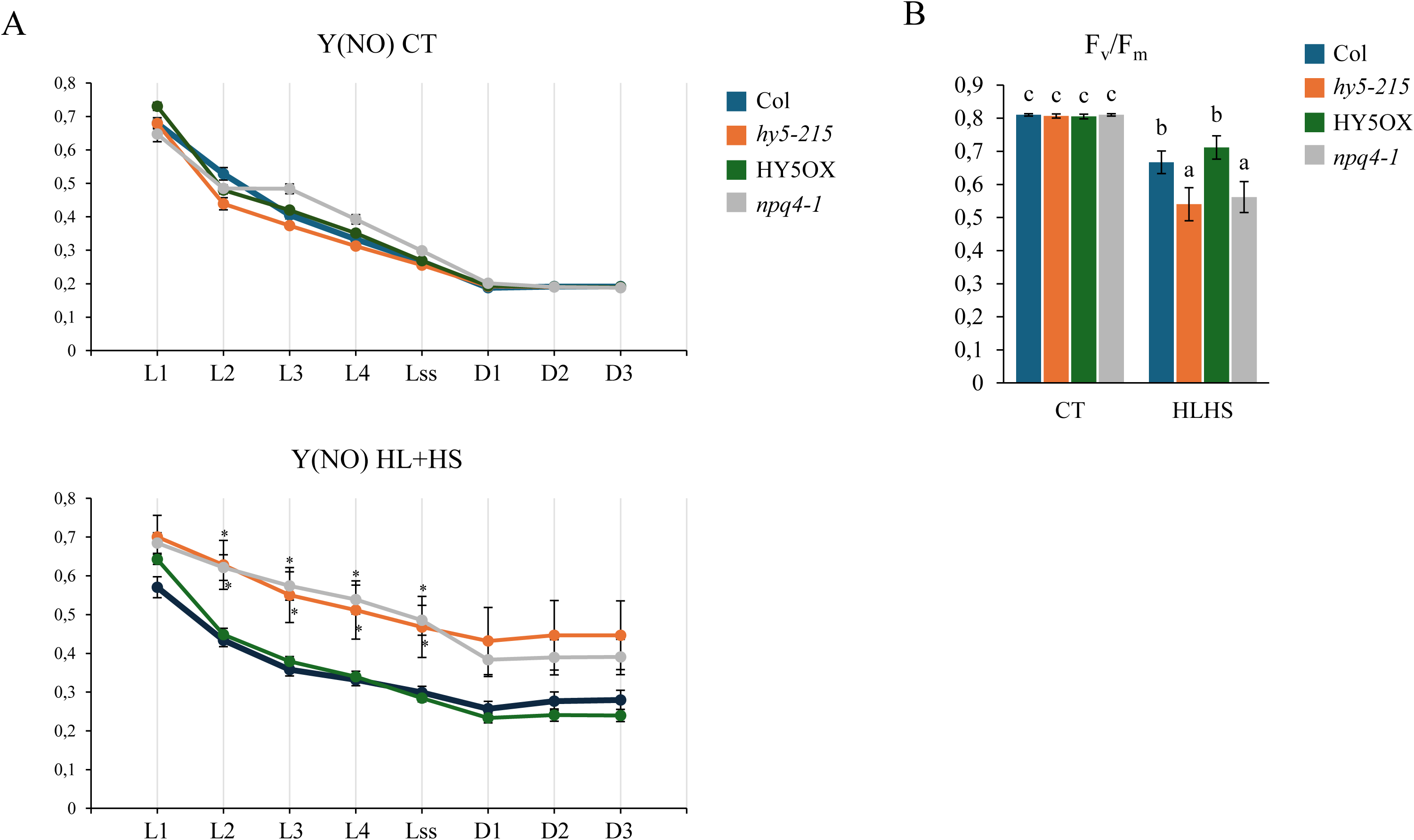
Kinetics of PSII photochemical parameters in Col-0, *hy5-215*, HY5OX, and *npq4-1* plants under CT and HLHS conditions. (A) Non-regulated energy dissipation [Y(NO)] measured over a series of actinic light pulses and dark relaxation points (L1–Lss, D1-D3) in all genotypes under CT and HLHS conditions. (B) Maximum quantum efficiency of PSII (Fv/Fm) in all genotypes under CT and after HLHS treatment. Error bars represent mean ± standard error (SE) (n = 9).

