## Supplementary material for "HY5 enhances *Arabidopsis* tolerance to combined high light and heat stress by coordinating photoprotection and hormone signaling": Table S1

**Supplemental Table S1.** Transcript-specific primers used for relative expression analysis by RT-qPCR.

| Gene | Accession | | Primer |
|---|---|---|---|
| *EF1* α | AT5G60390 | **F** | GAGCCCAAGTTTTTGAAGA |
| | | **R** | CTAACAGCGAAACGTCCCA |
| *WRKY48* | AT5G49520 | **F** | GGTTGCGGAGTGAAGAAGAG |
| | | **R** | GTTGCACCGTGGTCTAGGAT |
| *NPQ4* | AT1G44575 | **F** | ATGCTGCTTACTTCAGGCGT |
| | | **R** | GTTCTTTGAGACCGAGGGCA |
| *GAPB* | AT1G42970 | **F** | CCTGCTCAATGCTCCTCCAA |
| | | **R** | TTGGCTGGTGCAGTGATGAT |
| *GAPA2* | AT1G12900 | **F** | GAGGTGTTGGCATGGTCGTA |
| | | **R** | GCTTTGGCTGCTCCTGTAGA |
| *TED4* | AT2G26670 | **F** | AGTCGCCGTCTTTAGTGGTG |
| | | **R** | AGAAAGTTCGCCGTCCCATT |
| *PPC2* | AT2G42600 | **F** | CAAGGAGCCCCGTTTTGTTG |
| | | **R** | CTTGGGTCACGGATCTGCTT |
| *PsaA* | ATCG00350 | **F** | GCGAGCACCAGTTTGACTTG |
| | | **R** | ACGCCCGCTGAATAGAAACA |
| *MYC2* | AT1G32640 | **F** | GACGCAATCGCTTACATCAA |
| | | **R** | CGAAGAACACGAAGACGACA |
| *VSP2* | AT5G24770 | **F** | CAACTACGCCAACTGCAGAA |
| | | **R** | GGCAAGTCCTTTGGCATAGA |
