## Supplementary material for "HY5 enhances *Arabidopsis* tolerance to combined high light and heat stress by coordinating photoprotection and hormone signaling": Table S2

**Supplemental Table S2. Differentially accumulated proteins compared to control (P < 0.05) in Col leaves subjected to high light stress.**

| Protein ID | Fold Change | p-value | Protein description |
|---|---|---|---|
| AT5G11170.1 | 6,978787475 | 4,253E-08 | UAP56a homolog of human UAP56 a chr5:3553334-3556646 FORWARD LENGTH=427 |
| AT5G14740.2 | 8,968699088 | 4,272E-08 | BETA CA2_ CA2_ DEG12_ CA18 BETA CARBONIC ANHYDRASE 2_ CARBONIC ANHYDRASE 18_ carbonic anhy |
| AT3G62560.1 | 0,399400322 | 1,919E-07 | Sar1d chr3:23137539-23138880 FORWARD LENGTH=193 |
| AT5G61790.1 | 1,342520889 | 2,738E-07 | CNX1_ ATCNX1 calnexin 1 chr5:24827394-24829642 REVERSE LENGTH=530 |
| AT4G02080.1 | 0,606453097 | 1,322E-06 | ATSARA1C_ SAR2_ SAR1C_ ATSAR2_ ASAR1 secretion-associated RAS super family 2 chr4:921554-922547 FORWA |
| AT5G12020.1 | 5,737829824 | 2,26E-06 | HSP17.6II 17.6 kDa class II heat shock protein chr5:3882409-3882876 REVERSE LENGTH=155 |
| AT3G13470.1 | 1,713565674 | 2,337E-06 | CPNB2_ Cpn60beta2 chaperonin-60beta2 chr3:4389685-4392624 FORWARD LENGTH=596 |
| AT4G17090.1 | 1,781155971 | 2,591E-06 | CT-BMY_ BMY8_ AtBAM3_ BAM3 BETA-AMYLASE 8_ BETA-AMYLASE 3_ chloroplast beta-amylase chr4:960526 |
| AT4G13940.1 | 1,283529838 | 2,873E-06 | MEE58_ SAHH1_ EMB1395_ HOG1_ SAH1_ ATSAHH1 EMBRYO DEFECTIVE 1395_ HOMOLOGY-DEPENDENT |
| AT5G12110.1 | 3,950370582 | 4,545E-06 | no symbol available no full name available chr5:3914483-3915732 FORWARD LENGTH=228 |
| AT2G39310.1 | 4,178791474 | 5,048E-06 | JAL22 jacalin-related lectin 22 chr2:16414262-16416323 REVERSE LENGTH=458 |
| AT1G09180.1 | 1,326782499 | 5,37E-06 | SAR1A_ ATSARA1A_ SARA1A_ ATSAR1 SECRETION-ASSOCIATED RAS 1_ secretion-associated RAS super family |
| AT2G33380.1 | 1,881256936 | 5,821E-06 | AtRD20_ PXG3_ CLO-3_ CLO3_ RD20_ AtCLO3 caleosin 3_ Arabidopsis thaliana caleosin 3_ peroxygenase 3_ RESPON |
| AT3G12580.1 | 1,982108533 | 6,382E-06 | HSP70_ ATHSP70_ HSC70-4 ARABIDOPSIS HEAT SHOCK PROTEIN 70_ heat shock protein 70 chr3:3991487-39936 |
| AT1G20260.1 | 1,831269569 | 8,202E-06 | AtVAB3_ VAB3 V-ATPase B subunit 3 chr1:7016971-7020290 FORWARD LENGTH=487 |
| AT4G04020.1 | 3,160543518 | 8,958E-06 | FIB_ PGL35_ FIB1a plastoglobulin 35_ fibrillin 1a_ fibrillin chr4:1932161-1933546 FORWARD LENGTH=318 |
| AT2G39390.1 | 0,296479916 | 9,8E-06 | no symbol available no full name available chr2:16450803-16451762 REVERSE LENGTH=123 |
| AT5G02160.1 | 0,674239913 | 1,616E-05 | FIP FtsH5 Interacting Protein chr5:426392-427024 FORWARD LENGTH=129 |
| AT4G36130.1 | 0,128859286 | 1,864E-05 | no symbol available no full name available chr4:17097613-17098656 FORWARD LENGTH=258 |
| AT3G02560.1 | 0,498102105 | 2,36E-05 | no symbol available no full name available chr3:542341-543168 FORWARD LENGTH=191 |
| AT4G21960.1 | 0,37855313 | 3,504E-05 | PRXR1 chr4:11646613-11648312 REVERSE LENGTH=330 |
| AT5G03300.1 | 2,035029393 | 3,644E-05 | ADK2 adenosine kinase 2 chr5:796573-798997 FORWARD LENGTH=345 |
| AT2G29500.1 | 15,09460288 | 4,126E-05 | HSP17.6B chr2:12633279-12633740 REVERSE LENGTH=153 |
| AT2G35635.1 | 2,409703154 | 4,473E-05 | UBQ7_ RUB2 RELATED TO UBIQUITIN 2_ ubiquitin 7 chr2:14981044-14981943 FORWARD LENGTH=154 |
| AT5G12030.1 | 5,01729245 | 7,089E-05 | AT-HSP17.6A_ HSP17.6A_ HSP17.6 HEAT SHOCK PROTEIN 17.6_ heat shock protein 17.6A chr5:3884214-3884684 R |
| AT2G10940.1 | 1,839152008 | 7,235E-05 | no symbol available no full name available chr2:4311160-4312035 REVERSE LENGTH=291 |
| AT3G16480.1 | 2,084208989 | 7,265E-05 | MPPalpha mitochondrial processing peptidase alpha subunit chr3:5599906-5602716 FORWARD LENGTH=499 |
| AT2G05100.1 | 0,375778286 | 7,878E-05 | LHCB2_ LHCB2.1 LIGHT-HARVESTING CHLOROPHYLL B-BINDING 2_ photosystem II light harvesting complex ge |
| AT4G38510.1 | 0,632405612 | 8,42E-05 | AtVAB2_ VAB2 V-ATPase B subunit 2 chr4:18011155-18014789 REVERSE LENGTH=487 |
| AT1G56330.1 | 1,274923026 | 8,483E-05 | ATSARA1B_ SAR1B_ SAR1_ ATSAR1B_ ATSAR1 SECRETION-ASSOCIATED RAS 1_ ARABIDOPSIS THALIANA |
| AT2G35370.1 | 0,722493747 | 9,404E-05 | GDCH glycine decarboxylase complex H chr2:14891239-14892050 FORWARD LENGTH=165 |
| AT3G49010.4 | 0,151010467 | 0,0001088 | BBC1_ RSU2_ ATBBC1 40S RIBOSOMAL PROTEIN_ breast basic conserved 1 chr3:18166971-18168047 REVERSE L |
| AT5G20290.1 | 0,215780411 | 0,0001093 | no symbol available no full name available chr5:6851695-6853012 REVERSE LENGTH=222 |
| AT5G23860.1 | 0,650532748 | 0,0001156 | TUB8 tubulin beta 8 chr5:8042962-8044528 FORWARD LENGTH=449 |
| AT5G48570.1 | 2,978301462 | 0,0001277 | ROF2_ ATFKBP65_ FKBP65 chr5:19690746-19693656 REVERSE LENGTH=578 |
| AT4G24190.1 | 1,522742637 | 0,0001354 | SHD_ HSP90_ HSP90.7_ AtHsp90-7_ AtHsp90.7 SHEPHERD_ HEAT SHOCK PROTEIN 90.7_ HEAT SHOCK PROTE |

| | | | |
|---|---|---|---|
| AT3G59970.1 | 2,49439194 | 0,0001468 | MTHFR1 methylenetetrahydrofolate reductase 1 chr3:22151303-22153412 FORWARD LENGTH=421 |
| AT5G43010.1 | 0,349198709 | 0,0001531 | RPT4A regulatory particle triple-A ATPase 4A chr5:17248563-17251014 REVERSE LENGTH=399 |
| AT1G12250.1 | 0,589043967 | 0,0001665 | TL20.3 chr1:4159287-4161269 FORWARD LENGTH=280 |
| AT3G45030.1 | 0,390492928 | 0,000179 | no symbol available no full name available chr3:16471606-16472312 REVERSE LENGTH=124 |
| AT2G18450.1 | 1,281002917 | 0,0001803 | SDH1-2 succinate dehydrogenase 1-2 chr2:7997510-8000801 REVERSE LENGTH=632 |
| AT5G49910.1 | 1,27852578 | 0,0001803 | HSC70-7_ cpHsc70-2 chloroplast heat shock protein 70-2_ HEAT SHOCK PROTEIN 70-7 chr5:20303470-20306295 FOR |
| AT1G79040.1 | 0,91447974 | 0,0001808 | PSBR photosystem II subunit R chr1:29736085-29736781 FORWARD LENGTH=140 |
| AT5G52640.1 | 3,19225621 | 0,0001929 | HSP81-1_ ATHSP90.1_ HSP81.1_ AtHsp90-1_ HSP83_ ATHS83_ HSP90.1 HEAT SHOCK PROTEIN 90-1_ heat shock |
| AT1G16470.1 | 0,469537212 | 0,0002027 | PAB1 proteasome subunit PAB1 chr1:5623122-5625439 FORWARD LENGTH=235 |
| AT4G37990.1 | 1,778618719 | 0,0002115 | ELI3-2_ ATCAD8_ ELI3_ CAD-B2 elicitor-activated gene 3-2_ CINNAMYL-ALCOHOL DEHYDROGENASE B2_ ARA |
| AT1G79920.1 | 1,421506079 | 0,0002206 | Hsp70-15_ AtHsp70-15 heat shock protein 70-15 chr1:30059302-30062224 REVERSE LENGTH=736 |
| AT3G15020.1 | 1,242724906 | 0,0002236 | mMDH2 mitochondrial malate dehydrogenase 2 chr3:5056139-5057941 FORWARD LENGTH=341 |
| AT2G29550.1 | 1,365587857 | 0,0002469 | TUB7_ TBB7 tubulin beta-7 chain_ tubulin beta 7 chr2:12644258-12645932 REVERSE LENGTH=449 |
| AT5G17990.1 | 1,449057034 | 0,0002492 | pat1_ TRP1 tryptophan biosynthesis 1_ PHOSPHORIBOSYLANTHRANILATE TRANSFERASE 1 chr5:5957330-595968 |
| AT3G18130.1 | 3,038761382 | 0,0002518 | RACK1C_ RACK1C_AT receptor for activated C kinase 1C chr3:6211109-6212371 REVERSE LENGTH=326 |
| AT1G36280.2 | 0,484324227 | 0,0002613 | no symbol available no full name available chr1:13640600-13642908 FORWARD LENGTH=519 |
| AT3G06720.1 | 1,250404703 | 0,0002644 | IMPA1_ IMPA-1_ AT-IMP_ ATKAP ALPHA_ AIMP ALPHA IMPORTIN ALPHA_ IMPORTIN ALPHA ISOFORM 1_ |
| AT1G07400.1 | 4,968395398 | 0,0002695 | HSP17.8 chr1:2275148-2275621 FORWARD LENGTH=157 |
| AT3G24503.1 | 0,682846651 | 0,0002705 | REF1_ ALDH2C4_ ALDH1A REDUCED EPIDERMAL FLUORESCENCE1_ aldehyde dehydrogenase 2C4_ aldehyde de |
| AT5G59880.2 | 0,696574768 | 0,0002834 | ADF3 actin depolymerizing factor 3 chr5:24120382-24121628 FORWARD LENGTH=124 |
| AT1G03680.1 | 0,721816979 | 0,0003037 | ATHM1_ TRX-M1_ ATM1_ THM1 thioredoxin M-type 1_ THIOREDOXIN M-TYPE 1_ ARABIDOPSIS THIOREDOXI |
| AT2G38540.1 | 2,071978224 | 0,0003193 | ATLTP1_ AtLtpI-4_ LTP1_ LP1 ARABIDOPSIS THALIANA LIPID TRANSFER PROTEIN 1_ lipid transfer protein 1_ I |
| AT1G73600.1 | 2,251129492 | 0,0003204 | DEG26_ NMT_ AtPMT3_ NMT3 Phosphoethanolamine methyltransferase3 chr1:27670825-27673400 FORWARD LENG |
| AT4G18100.1 | 0,179358906 | 0,0003325 | no symbol available no full name available chr4:10035715-10036475 REVERSE LENGTH=133 |
| AT4G21990.1 | 1,656896963 | 0,0003372 | APR3_ PRH26_ PRH-26_ ATAPR3 PAPS REDUCTASE HOMOLOG 26_ APS reductase 3 chr4:11657284-11658973 RE |
| ATCG00830.1 | 0,126304795 | 0,0003511 | RPL2.1 ribosomal protein L2 chrc:84337-85843 REVERSE LENGTH=274 |
| AT1G55060.1 | 1,488359603 | 0,0003846 | UBQ12 ubiquitin 12 chr1:20549533-20550225 FORWARD LENGTH=230 |
| AT2G39590.1 | 1,174733688 | 0,000391 | no symbol available no full name available chr2:16517588-16518247 REVERSE LENGTH=130 |
| AT1G54340.1 | 2,435073985 | 0,0004029 | ICDH isocitrate dehydrogenase chr1:20283520-20286506 FORWARD LENGTH=416 |
| AT1G72810.1 | 2,799708664 | 0,000422 | TSY THREONINE SYNTHASE 2 chr1:27398760-27400393 REVERSE LENGTH=516 |
| AT5G10160.1 | 1,378473571 | 0,0004226 | no symbol available no full name available chr5:3185819-3187159 FORWARD LENGTH=219 |
| AT5G43850.1 | 2,456015277 | 0,0004385 | ATARD4_ ARD4 chr5:17627364-17629122 REVERSE LENGTH=187 |
| AT4G24280.1 | 1,225342807 | 0,0004454 | cpHsc70-1 chloroplast heat shock protein 70-1 chr4:12590094-12593437 FORWARD LENGTH=718 |
| AT4G27440.1 | 0,663173064 | 0,0004607 | PORB protochlorophyllide oxidoreductase B chr4:13725648-13727107 FORWARD LENGTH=401 |
| AT4G14960.1 | 1,952504225 | 0,0004889 | TUA6 Tubulin alpha-6 chr4:8548753-8550319 REVERSE LENGTH=427 |
| AT1G12250.2 | 0,668818078 | 0,0004889 | TL20.3 chr1:4159623-4161269 FORWARD LENGTH=206 |
| AT2G42910.1 | 1,860655428 | 0,0004989 | AtPRS4_ PRS4 phosphoribosyl diphosphate synthase 4 chr2:17856396-17858394 FORWARD LENGTH=337 |
| AT1G43560.1 | 2,703869898 | 0,0005588 | Aty2_ ty2 thioredoxin Y2 chr1:16398359-16399828 REVERSE LENGTH=167 |
| AT3G09440.1 | 1,375268311 | 0,0006658 | no symbol available no full name available chr3:2903434-2905632 REVERSE LENGTH=649 |

| | | | |
|---|---|---|---|
| AT5G25460.1 | 1,424618875 | 0,0006846 | DGR2 DUF642 L-GalL responsive gene 2 chr5:8863430-8865394 FORWARD LENGTH=369 |
| AT3G07090.1 | 2,413287631 | 0,000709 | Desi1 chr3:2243153-2244476 REVERSE LENGTH=265 |
| AT5G52650.1 | 1,596677728 | 0,0007192 | no symbol available no full name available chr5:21355781-21357003 REVERSE LENGTH=179 |
| AT2G21660.1 | 0,650244994 | 0,0007325 | RBGA3_ GR-RBP7_ CCR2_ ATGRP7_ GRP7_ SRBP1 RNA-binding glycine-rich protein A3_ SMALL RNA-BINDING I |
| AT3G04920.2 | 0,270081047 | 0,0007431 | no symbol available no full name available chr3:1360989-1361719 FORWARD LENGTH=112 |
| AT3G13870.1 | 1,704633632 | 0,000764 | GOM8_ RHD3 GOLGI MUTANT 8_ ROOT HAIR DEFECTIVE 3 chr3:4565762-4571109 REVERSE LENGTH=802 |
| AT5G42980.1 | 1,268091637 | 0,0007733 | ATH3_ TRX3_ TRXH3_ ATTRX3_ ATTRXH3 THIOREDOXIN H3_ thioredoxin 3_ thioredoxin H-type 3 chr5:1724277 |
| AT3G26060.1 | 0,868515926 | 0,0007875 | ATPRX Q_ PRXQ peroxiredoxin Q chr3:9524807-9526123 FORWARD LENGTH=216 |
| AT4G16830.2 | 0,705941506 | 0,000801 | AtRGGA chr4:9470979-9472308 FORWARD LENGTH=265 |
| AT1G42970.1 | 1,070489821 | 0,0008072 | GAPB glyceraldehyde-3-phosphate dehydrogenase B subunit chr1:16127552-16129584 FORWARD LENGTH=447 |
| AT5G61780.1 | 1,871536229 | 0,00081 | Tudor2_ AtTudor2_ TSN2 Arabidopsis thaliana TUDOR-SN protein 2_ TUDOR-SN protein 2 chr5:24822012-24826641 F |
| AT3G08590.1 | 1,421291093 | 0,0008275 | iPGAM2 "2_3-biphosphoglycerate-independent phosphoglycerate mutase 2" chr3:2608683-2611237 REVERSE LENGTH= |
| AT1G01100.1 | 1,469542227 | 0,0008396 | RPP1A_ RPP1.1 60S acidic ribosomal protein P1-1_ RPP1 co-orthologous gene 1 chr1:50284-50954 REVERSE LENGTH |
| AT1G62380.1 | 0,475042635 | 0,0008952 | ATACO2_ ACO2 ACC oxidase 2 chr1:23082340-23084068 FORWARD LENGTH=320 |
| AT3G14420.1 | 1,199889695 | 0,0009096 | GOX1 glycolate oxidase 1 chr3:4821804-4823899 FORWARD LENGTH=367 |
| AT1G65350.1 | 1,526691394 | 0,0009186 | UBQ13 ubiquitin 13 chr1:24272518-24277275 REVERSE LENGTH=319 |
| AT3G24170.1 | 1,403085873 | 0,0009358 | ATGR1_ GR1 glutathione-disulfide reductase chr3:8729762-8734115 REVERSE LENGTH=499 |
| AT5G14740.5 | 0,680327357 | 0,0009613 | BETA CA2_ CA2_ DEG12_ CA18 BETA CARBONIC ANHYDRASE 2_ CARBONIC ANHYDRASE 18_ carbonic anhy |
| AT5G35530.1 | 0,610883014 | 0,0009806 | no symbol available no full name available chr5:13710355-13712192 REVERSE LENGTH=248 |
| AT4G37300.1 | 0,740360733 | 0,0010255 | MEE59 maternal effect embryo arrest 59 chr4:17554805-17555498 FORWARD LENGTH=173 |
| AT5G48375.1 | 0,670011028 | 0,00104 | TGG3_ BGLU39 thioglucoside glucosidase 3_ BETA GLUCOSIDASE 39 chr5:19601303-19603883 REVERSE LENGTH |
| AT1G57720.1 | 1,348370408 | 0,0010416 | no symbol available no full name available chr1:21377873-21380114 FORWARD LENGTH=413 |
| AT5G54770.1 | 1,525726427 | 0,0010542 | THI1_ TZ_ THI4 THIAZOLE REQUIRING_ THIAMINE4 chr5:22246634-22247891 FORWARD LENGTH=349 |
| AT5G02500.1 | 1,399350809 | 0,0010881 | AtHsp70-1_ HSP70-1_ HSC70-1_ HSC70_ AT-HSC70-1 ARABIDOPSIS THALIANA HEAT SHOCK COGNATE PROT |
| ATCG00160.1 | 1,234157887 | 0,0010964 | RPS2 ribosomal protein S2 chrc:15013-15723 REVERSE LENGTH=236 |
| AT3G27690.1 | 1,621949527 | 0,001101 | LHCB2.4_ DEG13_ LHCB2_ LHCB2.3 LIGHT-HARVESTING CHLOROPHYLL B-BINDING 2_ photosystem II light ha |
| AT3G53430.1 | 1,487247788 | 0,0011351 | no symbol available no full name available chr3:19809895-19810395 REVERSE LENGTH=166 |
| AT4G04910.1 | 3,436403281 | 0,0011746 | NSF N-ethylmaleimide sensitive factor chr4:2489696-2495666 REVERSE LENGTH=742 |
| AT1G36240.1 | 0,309515871 | 0,0011777 | RPL30A chr1:13614890-13616233 FORWARD LENGTH=112 |
| AT1G65980.1 | 1,205908982 | 0,0011811 | TPX1 thioredoxin-dependent peroxidase 1 chr1:24559524-24560753 REVERSE LENGTH=162 |
| AT5G66120.2 | 1,504428772 | 0,0011951 | no symbol available no full name available chr5:26431516-26433649 REVERSE LENGTH=442 |
| AT4G17520.1 | 0,66412341 | 0,0012064 | HLN HYALURONAN/mRNA BINDING FAMILY PROTEIN chr4:9771496-9773313 FORWARD LENGTH=360 |
| AT5G44020.1 | 2,024753852 | 0,0012301 | no symbol available no full name available chr5:17712433-17714046 FORWARD LENGTH=272 |
| AT5G37640.1 | 1,571256703 | 0,0012541 | UBQ9 ubiquitin 9 chr5:14952782-14953750 REVERSE LENGTH=322 |
| AT1G77490.2 | 0,791919698 | 0,0012823 | TAPX thylakoidal ascorbate peroxidase chr1:29117688-29120649 FORWARD LENGTH=421 |
| AT4G28080.1 | 1,383257414 | 0,0013197 | REC2 REDUCED CHLOROPLAST COVERAGE 2 chr4:13948993-13957840 REVERSE LENGTH=1819 |
| AT3G46430.1 | 1,565950091 | 0,0013237 | AtMtATP6 chr3:17087687-17088497 FORWARD LENGTH=55 |
| AT3G25800.1 | 1,310793331 | 0,0013284 | PP2AA2_ PDF1_ PR 65 protein phosphatase 2A subunit A2 chr3:9422822-9425783 REVERSE LENGTH=587 |
| ATCG00660.1 | 0,409595649 | 0,0013355 | RPL20 ribosomal protein L20 chrc:68512-68865 REVERSE LENGTH=117 |

| | | | |
|---|---|---|---|
| AT2G41100.5 | 0,3317177 | 0,001343 | TCH3_ CML12_ ATCAL4 ARABIDOPSIS THALIANA CALMODULIN LIKE 4_ calmodulin-like 12_ TOUCH 3 chr2:1 |
| AT1G48600.1 | 2,045052894 | 0,0014105 | AtPMT2_ AtPMEAMT_ PMEAMT phosphoethanolamine N-methyltransferase_ Phosphoethanolamine methyltransferase2 |
| AT5G01410.1 | 1,429487664 | 0,0014481 | PDX1_ ATPDX1.3_ ATPDX1_ PDX1.3_ RSR4 REDUCED SUGAR RESPONSE 4_ PYRIDOXINE BIOSYNTHESIS 1. |
| AT2G38230.1 | 1,255439661 | 0,0014538 | ATPDX1.1_ PDX1.1 pyridoxine biosynthesis 1.1_ ARABIDOPSIS THALIANA PYRIDOXINE BIOSYNTHESIS 1.1 chr2 |
| AT4G22240.1 | 1,368850924 | 0,0014694 | FBN1b fibrillin 1b chr4:11766090-11767227 REVERSE LENGTH=310 |
| AT2G45740.1 | 1,258148466 | 0,0015039 | PEX11D peroxin 11D chr2:18839865-18841102 FORWARD LENGTH=236 |
| AT5G65010.1 | 1,48492823 | 0,0015399 | ASN2 asparagine synthetase 2 chr5:25969224-25972278 FORWARD LENGTH=578 |
| AT1G03130.1 | 0,776466135 | 0,0015404 | PSAD-2 photosystem I subunit D-2 chr1:753528-754142 REVERSE LENGTH=204 |
| AT5G20160.1 | 2,045626006 | 0,0015633 | no symbol available no full name available chr5:6804075-6805102 REVERSE LENGTH=128 |
| AT2G29440.1 | 2,617000694 | 0,0015676 | GST24_ ATGSTU6_ GSTU6 glutathione S-transferase tau 6_ GLUTATHIONE S-TRANSFERASE 24 chr2:12620159-126 |
| AT1G78630.1 | 0,198451235 | 0,0015753 | emb1473 embryo defective 1473 chr1:29575997-29577406 FORWARD LENGTH=241 |
| AT1G07890.1 | 1,212059195 | 0,0015786 | MEE6_ ATAPX01_ ATAPX1_ CS1_ APX1 ascorbate peroxidase 1_ maternal effect embryo arrest 6 chr1:2438005-24394 |
| AT1G54270.2 | 0,705253705 | 0,0016436 | EIF4A-2 eif4a-2 chr1:20260495-20262018 FORWARD LENGTH=407 |
| AT5G13490.1 | 1,567324612 | 0,0016902 | AAC2 ADP/ATP carrier 2 chr5:4336034-4337379 FORWARD LENGTH=385 |
| AT1G72150.1 | 0,706957272 | 0,0018023 | PATL1 PATELLIN 1 chr1:27148558-27150652 FORWARD LENGTH=573 |
| AT4G17170.1 | 1,145874394 | 0,0018242 | RAB2A_ AT-RAB2_ ATRAB2A_ ATRABB1C_ RAB-B1B_ ATRAB-B1B_ RABB1C ARABIDOPSIS RAB GTPASE H |
| AT2G28000.1 | 1,354989079 | 0,0018511 | ARC2_ CH-CPN60A_ SLP_ CPN60A_ Cpn60alpha1_ CPNA1 SCHLEPPERLESS_ chaperonin-60alpha1_ CHLOROPLA |
| AT1G30230.1 | 0,447239385 | 0,0018902 | EF1Bb_ eEF-1Bb1 eukaryotic elongation factor 1B beta 1_ elongation factor 1B beta chr1:10639286-10640515 FORWAR |
| AT3G09790.1 | 0,409178054 | 0,0018958 | UBQ8 ubiquitin 8 chr3:3004111-3006006 REVERSE LENGTH=631 |
| AT2G30490.1 | 1,276225901 | 0,0019037 | REF3_ CYP73A5_ C4H_ ATC4H CINNAMATE 4-HYDROXYLASE_ REDUCED EPRDERMAL FLUORESCENCE 3_ |
| AT5G01530.1 | 0,852684858 | 0,0019828 | LHCB4.1 light harvesting complex photosystem II chr5:209084-210243 FORWARD LENGTH=290 |
| AT2G24270.1 | 1,382779117 | 0,0020089 | ALDH11A3 aldehyde dehydrogenase 11A3 chr2:10327325-10329601 REVERSE LENGTH=496 |
| AT2G31790.1 | 1,475556748 | 0,0020211 | no symbol available no full name available chr2:13518269-13520167 FORWARD LENGTH=457 |
| AT1G05010.1 | 0,524688503 | 0,0020799 | EFE_ ACO4_ EAT1 ethylene forming enzyme_ ethylene-forming enzyme chr1:1431419-1432695 REVERSE LENGTH=32 |
| AT4G12420.1 | 0,870364372 | 0,0020923 | SKU5 chr4:7349941-7352868 REVERSE LENGTH=587 |
| AT1G60950.1 | 0,382451436 | 0,002116 | ATFD2_ FD2_ FED A FERREDOXIN 2 chr1:22444565-22445011 FORWARD LENGTH=148 |
| AT5G46290.3 | 1,326818978 | 0,0021466 | KASI_ KAS1 3-ketoacyl-acyl carrier protein synthase I_ KETOACYL-ACP SYNTHASE 1 chr5:18774439-18776629 REV |
| AT4G22670.1 | 1,133411674 | 0,0021485 | AtHip1_ HIP1_ TPR11 HSP70-interacting protein 1_ tetratricopeptide repeat 11 chr4:11918236-11920671 FORWARD LE |
| AT1G77510.1 | 1,503200289 | 0,0021761 | ATPDI6_ PDIL1-2_ ATPDIL1-2_ PDI6 PDI-like 1-2_ PROTEIN DISULFIDE ISOMERASE 6 chr1:29126742-29129433 |
| AT3G09350.1 | 2,939047319 | 0,002183 | Fes1A Fes1A chr3:2871216-2873109 FORWARD LENGTH=363 |
| AT1G50670.1 | 1,433296691 | 0,0021863 | OTU2 ovarian tumor domain (OTU)-containing DUB (deubiquitilating enzyme) 2 chr1:18775086-18776552 REVERSE LE |
| AT5G15970.1 | 13,14805115 | 0,0022583 | AtCor6.6_ KIN2_ COR6.6 COLD-RESPONSIVE 6.6 chr5:5211966-5212441 FORWARD LENGTH=66 |
| AT1G11840.1 | 1,633770816 | 0,0022942 | AtGLYI3_ GLX1_ ATGLX1 glyoxalaseI 3_ glyoxalase I homolog chr1:3996045-3997518 FORWARD LENGTH=283 |
| AT2G47470.1 | 1,202027619 | 0,0023056 | UNE5_ MEE30_ ATPDI11_ PDI11_ ATPDIL2-1 PROTEIN DISULFIDE ISOMERASE 11_ PDI-LIKE 2-1_ ARABIDOP |
| AT5G06290.1 | 1,186373499 | 0,0023173 | 2-Cys Prx B_ 2CPB 2-cysteine peroxiredoxin B_ 2-CYS PEROXIREDOXIN B chr5:1919380-1921211 FORWARD LENG |
| AT1G55490.1 | 1,075574055 | 0,0023727 | Cpn60beta1_ LEN1_ CPNB1_ CPN60B chaperonin-60beta1_ LESION INITIATION 1_ chaperonin 60 beta chr1:207157 |
| AT5G08570.1 | 1,360297782 | 0,0024261 | no symbol available no full name available chr5:2778433-2780300 FORWARD LENGTH=510 |
| AT1G47250.1 | 1,487633342 | 0,0025017 | PAF2 20S proteasome alpha subunit F2 chr1:17319220-17320900 FORWARD LENGTH=277 |
| ATCG00120.1 | 1,107361975 | 0,002561 | ATPA ATP synthase subunit alpha chrc:9938-11461 REVERSE LENGTH=507 |

| | | | |
|---|---|---|---|
| AT4G36250.1 | 1,242610297 | 0,0026882 | ALDH3F1 aldehyde dehydrogenase 3F1 chr4:17151029-17153381 FORWARD LENGTH=484 |
| AT4G27560.1 | 1,607155835 | 0,0026996 | UGT79B2 chr4:13760114-13761481 REVERSE LENGTH=455 |
| AT5G15450.1 | 2,046425932 | 0,0027666 | APG6_ AtCLPB3_ CLPB3_ CLPB-P casein lytic proteinase B3_ CASEIN LYTIC PROTEINASE B-P_ ALBINO AND PA |
| AT2G20560.1 | 5,940191297 | 0,0027689 | DNAJ DNAJ protein chr2:8848353-8849815 REVERSE LENGTH=337 |
| AT1G74040.1 | 1,914371449 | 0,0028332 | IPMS2_ IMS1_ MAML-3 SOPROPYLMALATE SYNTHASE 2_ 2-isopropylmalate synthase 1 chr1:27842258-27845566 |
| AT5G64140.1 | 0,739122294 | 0,0028546 | RPS28 ribosomal protein S28 chr5:25667529-25667723 REVERSE LENGTH=64 |
| AT1G64520.1 | 1,154397369 | 0,0029557 | RPN12a regulatory particle non-ATPase 12A chr1:23956459-23958120 FORWARD LENGTH=267 |
| AT2G21870.1 | 0,783876651 | 0,0030439 | MGP1_ PHI1 PHOSPHITE-INSENSITIVE 1_ MALE GAMETOPHYTE DEFECTIVE 1 chr2:9320456-9322618 REVERS |
| AT3G01390.1 | 1,913793867 | 0,0030851 | AVMA10_ VMA10 vacuolar membrane ATPase 10 chr3:150265-150922 REVERSE LENGTH=110 |
| ATCG00810.1 | 0,171076083 | 0,0031447 | RPL22 ribosomal protein L22 chrc:83467-83949 REVERSE LENGTH=160 |
| AT4G09720.1 | 1,584931572 | 0,0031831 | RABG3A_ ATRABG3A RAB GTPase homolog G3A chr4:6133101-6134959 FORWARD LENGTH=206 |
| AT5G11420.1 | 0,697145143 | 0,0032407 | no symbol available no full name available chr5:3644655-3646991 FORWARD LENGTH=366 |
| AT2G29450.1 | 1,405386137 | 0,0032699 | AT103-1A_ ATGSTU1_ ATGSTU5_ GSTU5 glutathione S-transferase tau 5_ ARABIDOPSIS THALIANA GLUTATHIO |
| AT5G24780.1 | 1,196902455 | 0,003292 | VSP1_ ATVSP1 vegetative storage protein 1 chr5:8507783-8508889 REVERSE LENGTH=270 |
| AT5G55480.1 | 1,517673096 | 0,0032931 | GPDL1_ GDPDL4_ SVL1 SHV3-like 1_ Glycerophosphodiester phosphodiesterase (GDPD) like 4_ glycerophosphodiester |
| AT4G37910.1 | 1,456101178 | 0,0033075 | mtHsc70-1 mitochondrial heat shock protein 70-1 chr4:17825368-17828099 REVERSE LENGTH=682 |
| AT4G10320.1 | 1,478148371 | 0,0033139 | no symbol available no full name available chr4:6397526-6404509 REVERSE LENGTH=1190 |
| AT3G49110.1 | 0,811162669 | 0,0033211 | ATPCA_ PRX33_ ATPRX33_ PRXCA peroxidase CA_ PEROXIDASE CA_ PEROXIDASE 33 chr3:18200713-18202891 |
| AT1G74970.1 | 1,211880324 | 0,0033285 | TWN3_ SOT8_ RPS9_ PRPS9 ribosomal protein S9 chr1:28157761-28159202 REVERSE LENGTH=208 |
| AT3G26450.1 | 1,295771536 | 0,0033749 | no symbol available no full name available chr3:9681593-9683299 REVERSE LENGTH=152 |
| AT3G18780.1 | 1,539946693 | 0,0033955 | LSR2_ ACT2_ ENL2_ DER1_ FIZ2 FRIZZY AND KINKED SHOOTS 2_ LIGHT STRESS-REGULATED 2_ DEFORMI |
| AT1G16890.1 | 1,88438944 | 0,003444 | UBC13B_ AtUBC36_ UBC36 ubiquitin-conjugating enzyme 36_ UBIQUITIN CONJUGATING ENZYME 13B chr1:5776 |
| AT5G63400.1 | 0,835028054 | 0,0036013 | ADK1 adenylate kinase 1 chr5:25393274-25394817 REVERSE LENGTH=246 |
| AT3G08940.1 | 0,783496252 | 0,0036187 | LHCB4.2 light harvesting complex photosystem II chr3:2717717-2718400 FORWARD LENGTH=227 |
| AT3G46060.1 | 1,361233819 | 0,0036843 | ARA3_ RAB8A_ RABE1c_ ARA-3_ ATRAB8A_ ATRABE1C RAB GTPase homolog 8A chr3:16917908-16919740 FOR |
| AT5G63890.1 | 1,470950405 | 0,0037019 | HISN8_ ATHDH_ HDH histidinol dehydrogenase_ HISTIDINE BIOSYNTHESIS 8 chr5:25565600-25567879 REVERSE |
| AT1G22450.1 | 0,697898393 | 0,0037333 | ATCOX6B2_ COX6B cytochrome C oxidase 6B_ CYTOCHROME C OXIDASE 6B2 chr1:7925447-7926918 FORWARD |
| AT2G28950.1 | 0,829066974 | 0,0037452 | ATEXP6_ ATHEXP ALPHA 1.8_ ATEXPA6_ EXPA6 expansin A6_ ARABIDOPSIS THALIANA TEXPANSIN 6 chr2:1 |
| AT4G20890.1 | 0,657968508 | 0,0037469 | TUB9 tubulin beta-9 chain chr4:11182218-11183840 FORWARD LENGTH=444 |
| AT2G01140.1 | 0,774249816 | 0,0038023 | FBA3_ AtFBA3_ PDE345 PIGMENT DEFECTIVE 345_ fructose-bisphosphate aldolase 3 chr2:95006-96491 REVERSE I |
| AT2G42520.1 | 1,172482511 | 0,0038133 | RH37 RNA Helicase 37 chr2:17705382-17708744 FORWARD LENGTH=633 |
| AT5G23140.1 | 0,811779832 | 0,0038839 | CLPP2_ NCLPP7 nuclear-encoded CLP protease P7 chr5:7783811-7784826 FORWARD LENGTH=241 |
| AT1G47260.1 | 0,871016853 | 0,0039716 | APFI_ GAMMA CA2 gamma carbonic anhydrase 2 chr1:17321384-17323347 REVERSE LENGTH=278 |
| AT3G46520.1 | 1,478509072 | 0,0039718 | ACT12 actin-12 chr3:17128567-17129981 FORWARD LENGTH=377 |
| AT3G44890.1 | 0,779929161 | 0,0040702 | RPL9 ribosomal protein L9 chr3:16386505-16387963 FORWARD LENGTH=197 |
| AT1G45000.1 | 1,331575475 | 0,0041408 | RPT4b chr1:17009220-17011607 FORWARD LENGTH=399 |
| AT5G55280.1 | 1,380800719 | 0,0041798 | CPFTSZ_ FTSZ1-1_ ATFTSZ1-1_ FtsZ1 CHLOROPLAST FTSZ_ homolog of bacterial cytokinesis Z-ring protein FTSZ 1 |
| AT2G32060.1 | 0,921937562 | 0,0041966 | no symbol available no full name available chr2:13639228-13640104 REVERSE LENGTH=144 |
| AT3G25230.1 | 1,715230972 | 0,0042168 | FKBP62_ ROF1_ ATFKBP62 rotamase FKBP 1_ FK506 BINDING PROTEIN 62 chr3:9188257-9191137 FORWARD LE |

| | | | |
|---|---|---|---|
| AT4G08900.1 | 0,603046248 | 0,0042215 | ARGAH1 arginine amidohydrolase 1 chr4:5703499-5705180 FORWARD LENGTH=342 |
| AT1G75780.1 | 0,67662842 | 0,0043154 | TUB1 tubulin beta-1 chain chr1:28451378-28453602 REVERSE LENGTH=447 |
| AT3G14415.1 | 1,161332285 | 0,004316 | GOX2 glycolate oxidase 2 chr3:4818667-4820748 FORWARD LENGTH=367 |
| AT4G35830.1 | 1,268181664 | 0,0044009 | ACO1 aconitase 1 chr4:16973007-16977949 REVERSE LENGTH=898 |
| AT3G44300.1 | 1,508777306 | 0,0044824 | AtNIT2_ NIT2 nitrilase 2 chr3:15983351-15985172 FORWARD LENGTH=339 |
| AT4G35250.1 | 1,082989578 | 0,0045717 | HCF244 high chlorophyll fluorescence phenotype 244 chr4:16771401-16773269 REVERSE LENGTH=395 |
| AT5G44340.1 | 1,385720822 | 0,0046565 | TUB4 tubulin beta chain 4 chr5:17859442-17860994 REVERSE LENGTH=444 |
| AT5G04140.1 | 0,550703178 | 0,0048163 | GLUS_ FD-GOGAT_ GLS1_ GLU1 FERREDOXIN-DEPENDENT GLUTAMATE SYNTHASE 1_ glutamate synthase 1_ |
| AT3G63540.1 | 0,786801367 | 0,0048667 | no symbol available no full name available chr3:23459372-23459803 REVERSE LENGTH=143 |
| AT1G45145.1 | 0,206801333 | 0,0048817 | TRX-h5_ ATTRX5_ LIV1_ ATH5_ TRX5 thioredoxin H-type 5_ THIOREDOXIN H-TYPE 5_ LOCUS OF INSENSITIV |
| AT1G74470.1 | 0,831325633 | 0,0049587 | no symbol available no full name available chr1:27991248-27992845 FORWARD LENGTH=467 |
| AT3G03710.1 | 1,271441339 | 0,0050113 | PNP_ RIF10_ PDE326 PIGMENT DEFECTIVE 326_ POLYNUCLEOTIDE PHOSPHORYLASE_ resistant to inhibition [illegible] |
| AT1G56070.1 | 1,207279993 | 0,0050265 | LOS1 LOW EXPRESSION OF OSMOTICALLY RESPONSIVE GENES 1 chr1:20968245-20971077 REVERSE LENGT |
| AT2G33040.1 | 1,152581881 | 0,0052423 | ATP3 gamma subunit of Mt ATP synthase chr2:14018978-14021047 REVERSE LENGTH=325 |
| AT2G36160.1 | 1,165743908 | 0,0052947 | no symbol available no full name available chr2:15169925-15171159 FORWARD LENGTH=150 |
| AT5G59720.1 | 2,494580736 | 0,0052994 | HSP18.2 heat shock protein 18.2 chr5:24062632-24063117 FORWARD LENGTH=161 |
| AT2G26740.1 | 1,403167633 | 0,0053438 | ATSEH_ SEH soluble epoxide hydrolase chr2:11393148-11394257 REVERSE LENGTH=321 |
| AT4G34450.1 | 1,265124884 | 0,0054748 | gamma2-COP gamma2 Coat Protein chr4:16471956-16476795 FORWARD LENGTH=886 |
| AT2G21130.1 | 1,734102103 | 0,0054842 | no symbol available no full name available chr2:9055619-9056143 REVERSE LENGTH=174 |
| AT5G11670.1 | 1,238291764 | 0,0054906 | NADP-ME2_ ATNADP-ME2 NADP-malic enzyme 2_ Arabidopsis thaliana NADP-malic enzyme 2 chr5:3754456-375804 |
| AT2G35840.1 | 1,930617587 | 0,0054912 | no symbol available no full name available chr2:15053952-15055776 FORWARD LENGTH=422 |
| AT5G52470.1 | 1,766956888 | 0,0054913 | ATFIB1_ FBR1_ FIB1_ SKIP7_ ATFBR1 fibrillarin 1_ FIBRILLARIN 1_ SKP1/ASK1-INTERACTING PROTEIN chr5: |
| AT3G19710.1 | 1,755433464 | 0,0054986 | BCAT4 branched-chain aminotransferase4 chr3:6847202-6849429 REVERSE LENGTH=354 |
| AT5G09590.1 | 1,428331811 | 0,0055857 | MTHSC70-2_ HSC70-5 mitochondrial HSO70 2_ HEAT SHOCK COGNATE chr5:2975721-2978508 FORWARD LENG |
| AT1G62740.1 | 0,797424373 | 0,0056227 | Hop2 Hop2 chr1:23231026-23233380 FORWARD LENGTH=571 |
| AT4G20260.1 | 0,788361687 | 0,0056299 | ATPCAP1_ PCAP1_ MDP25 ARABIDOPSIS THALIANA PLASMA-MEMBRANE ASSOCIATED CATION-BINDING |
| AT5G26780.1 | 0,888145863 | 0,0057216 | SHM2 serine hydroxymethyltransferase 2 chr5:9418299-9421725 FORWARD LENGTH=517 |
| AT5G56030.1 | 1,681917538 | 0,0057322 | HSP81-2_ HSP90.2_ AtHsp90.2_ ERD8_ HSP81.2 EARLY-RESPONSIVE TO DEHYDRATION 8_ heat shock protein 81 |
| AT1G48860.1 | 1,452775043 | 0,0058438 | EPSPS 5-enolpyruvylshikimate-3-phosphate synthase chr1:18068892-18071331 REVERSE LENGTH=521 |
| AT5G51440.1 | 2,27269579 | 0,0059112 | HSP23.5 chr5:20891242-20892013 FORWARD LENGTH=210 |
| AT3G63190.1 | 0,817033719 | 0,005933 | cpRRF_ HFP108_ AtcpRRF_ RRF chloroplast ribosome recycling factor_ Arabidopsis thaliana chloroplast ribosome recycl |
| AT2G37040.1 | 1,889050057 | 0,0059994 | PAL1_ ATPAL1 PHE ammonia lyase 1 chr2:15557602-15560237 REVERSE LENGTH=725 |
| AT5G54270.1 | 0,906739946 | 0,0060776 | LHCB3_ LHCB31 light-harvesting chlorophyll B-binding protein 3 chr5:22038424-22039383 FORWARD LENGTH=265 |
| AT2G43560.1 | 0,80921865 | 0,0061409 | no symbol available no full name available chr2:18073995-18075385 REVERSE LENGTH=223 |
| ATCG00780.1 | 0,843151999 | 0,0061451 | RPL14 ribosomal protein L14 chrc:80696-81064 REVERSE LENGTH=122 |
| AT4G11600.1 | 1,138687999 | 0,006197 | GPXL6_ PHGPX_ LSC803_ ATGPX6_ GPX6 glutathione peroxidase 6 chr4:7010021-7011330 REVERSE LENGTH=232 |
| AT2G22240.1 | 3,098441366 | 0,0062007 | MIPS2_ ATIPS2_ ATMIPS2 myo-inositol-1-phosphate synthase 2_ INOSITOL 3-PHOSPHATE SYNTHASE 2_ MYO-IN |
| AT1G55670.1 | 0,714044291 | 0,0062354 | PSAG photosystem I subunit G chr1:20802874-20803356 REVERSE LENGTH=160 |
| AT5G46800.1 | 1,144456459 | 0,0062689 | BOU A BOUT DE SOUFFLE chr5:18988779-18989810 REVERSE LENGTH=300 |

| | | | |
|---|---|---|---|
| AT1G67430.2 | 0,575373188 | 0,0064174 | no symbol available no full name available chr1:25262209-25263627 FORWARD LENGTH=131 |
| AT4G11820.1 | 1,461802218 | 0,0064552 | MVA1_ FKP1_ HMGS FLAKY POLLEN 1_ HYDROXYMETHYLGLUTARYL-COA SYNTHASE chr4:7109124-71112 |
| AT1G50200.1 | 1,172015048 | 0,0065132 | ALATS_ ACD Alanyl-tRNA synthetase chr1:18591429-18598311 REVERSE LENGTH=1003 |
| AT4G37980.1 | 1,197845422 | 0,006765 | ELI3-1_ CHR_ ELI3_ ATCAD7_ CAD7 elicitor-activated gene 3-1_ CINNAMALDEHYDE AND HEXENAL REDUCTA |
| AT1G20440.1 | 0,747178284 | 0,0067687 | AtCOR47_ RD17_ COR47 cold-regulated 47 chr1:7084722-7085664 REVERSE LENGTH=265 |
| AT5G45390.1 | 1,186785153 | 0,0067958 | CLPP4_ NCLPP4 NUCLEAR-ENCODED CLP PROTEASE P4_ CLP protease P4 chr5:18396351-18397586 FORWARD |
| AT5G12040.1 | 1,372143743 | 0,0068188 | no symbol available no full name available chr5:3885162-3887772 FORWARD LENGTH=369 |
| AT3G49120.1 | 0,907606451 | 0,006848 | PRX34_ PRXCB_ ATPCB_ AtPRX34_ PERX34_ ATPERX34 ARABIDOPSIS THALIANA PEROXIDASE CB_ PERO |
| AT2G06850.1 | 1,154039165 | 0,0068577 | EXT_ XTH4_ EXGT-A1 ENDOXYLOGLUCAN TRANSFERASE_ endoxyloglucan transferase A1_ xyloglucan endotrans |
| AT3G16400.1 | 1,238303059 | 0,0069209 | NSP1_ ATNSP1_ ATMLP-470 nitrile specifier protein 1_ NITRILE SPECIFIER PROTEIN 1_ MYROSINASE-BINDING |
| AT5G47870.1 | 1,953272222 | 0,0069773 | RAD52-2_ ODB2_ RAD52-2B radiation sensitive 52-2_ Organellar DNA-Binding protein 2 chr5:19384555-19385808 RE |
| AT4G29010.1 | 1,260844765 | 0,0069782 | AIM1 ABNORMAL INFLORESCENCE MERISTEM chr4:14297312-14302016 REVERSE LENGTH=721 |
| AT3G49680.2 | 1,334278194 | 0,0071251 | BCAT3_ ATBCAT-3 branched-chain aminotransferase 3 chr3:18422768-18425473 FORWARD LENGTH=411 |
| AT5G14590.1 | 1,25814456 | 0,0072346 | no symbol available no full name available chr5:4703533-4706627 REVERSE LENGTH=485 |
| AT5G66510.1 | 0,832724744 | 0,0072698 | GAMMA CA3 gamma carbonic anhydrase 3 chr5:26550016-26551496 REVERSE LENGTH=258 |
| AT1G76180.1 | 0,596036324 | 0,0074507 | ERD14 EARLY RESPONSE TO DEHYDRATION 14 chr1:28587013-28587657 REVERSE LENGTH=185 |
| AT4G39520.1 | 1,807422241 | 0,0074903 | Drg1-1 chr4:18371329-18374000 REVERSE LENGTH=369 |
| AT5G53560.1 | 0,665990442 | 0,0075522 | ATB5-A_ CB5-E_ ATCB5-E_ B5 #2 ARABIDOPSIS CYTOCHROME B5 ISOFORM E_ cytochrome B5 isoform E chr5: |
| AT1G30530.1 | 1,282842907 | 0,0075595 | UGT78D1 UDP-glucosyl transferase 78D1 chr1:10814917-10816374 FORWARD LENGTH=453 |
| AT1G66270.2 | 0,348396892 | 0,0076177 | BGLU21 chr1:24700110-24702995 REVERSE LENGTH=522 |
| AT1G41830.1 | 0,206528734 | 0,0076441 | SKS6 SKU5-similar 6_ SKU5 SIMILAR 6 chr1:15603892-15607802 REVERSE LENGTH=542 |
| AT1G64200.1 | 1,332390565 | 0,0077506 | VHA-E3 vacuolar H+-ATPase subunit E isoform 3 chr1:23828537-23830002 REVERSE LENGTH=237 |
| AT3G06860.1 | 1,264831796 | 0,0079225 | ATMFP2_ MFP2 MULTIFUNCTIONAL PROTEIN 2_ multifunctional protein 2 chr3:2161926-2166009 FORWARD LE |
| AT3G55800.1 | 1,319374144 | 0,0079981 | SBPASE sedoheptulose-bisphosphatase chr3:20709640-20711421 FORWARD LENGTH=393 |
| AT1G56450.1 | 1,127973728 | 0,0080009 | MUD1_ PBG1 20S proteasome beta subunit G1 chr1:21141970-21144186 FORWARD LENGTH=246 |
| AT5G07350.1 | 1,586649349 | 0,0080575 | Tudor1_ TSN1_ AtTudor1 TUDOR-SN protein 1_ Arabidopsis thaliana TUDOR-SN protein 1 chr5:2320344-2324892 RE |
| AT4G26300.4 | 1,366656819 | 0,0082775 | emb1027 embryo defective 1027 chr4:13308400-13312204 REVERSE LENGTH=590 |
| AT3G25760.1 | 1,159975943 | 0,0082905 | AOC1_ ERD12 early-responsive to dehydration 12_ allene oxide cyclase 1 chr3:9403972-9405105 FORWARD LENGTH= |
| AT3G13930.1 | 1,128732908 | 0,0083534 | mtE2-2 mitochondrial pyruvate dehydrogenase subunit 2-2 chr3:4596240-4600143 FORWARD LENGTH=539 |
| AT2G34430.1 | 0,779838894 | 0,0083998 | DEG11_ LHCB1.4_ LHB1B1 light-harvesting chlorophyll-protein complex II subunit B1_ LIGHT-HARVESTING CHLO |
| AT5G60670.1 | 0,764915427 | 0,0084016 | RPL12C Ribosomal Protein Like 12C chr5:24381066-24381566 REVERSE LENGTH=166 |
| AT2G27720.1 | 0,710479477 | 0,0084252 | no symbol available no full name available chr2:11818696-11819370 FORWARD LENGTH=115 |
| AT2G39470.2 | 0,778708669 | 0,0084445 | PnsL1_ PPL2 Photosynthetic NDH subcomplex L 1_ PsbP-like protein 2 chr2:16476335-16477653 FORWARD LENGTH= |
| AT1G18080.1 | 1,344454342 | 0,0086021 | RACK1A_AT_ AtRACK1_ SAC53_ ATARCA_ RACK1A RECEPTOR FOR ACTIVATED C KINASE 1 A_ Suppressor |
| AT1G43170.1 | 0,17301551 | 0,0087117 | emb2207_ RP1_ ARP1_ RPL3A embryo defective 2207_ ribosomal protein 1 chr1:16266992-16268631 FORWARD LEN |
| AT5G26570.1 | 1,279242344 | 0,008822 | ATGWD3_ OK1_ PWD PHOSPHOGLUCAN WATER DIKINASE chr5:9261580-9267526 FORWARD LENGTH=1196 |
| AT1G11750.1 | 1,495472305 | 0,0090875 | NCLPP1_ CLPP6_ NCLPP6 NUCLEAR-ENCODED CLPP 1_ CLP protease proteolytic subunit 6 chr1:3967609-3969535 |
| AT5G19140.2 | 1,424252432 | 0,0091066 | ATAILP1_ AILP1 chr5:6423398-6425785 FORWARD LENGTH=222 |
| AT3G62410.1 | 0,454660291 | 0,0091086 | CP12_ CP12-2 CP12 DOMAIN-CONTAINING PROTEIN 1_ CP12 domain-containing protein 2 chr3:23091006-2309140 |

| | | | |
|---|---|---|---|
| AT5G24770.1 | 1,251481654 | 0,0091807 | ATVSP2_ VSP2 vegetative storage protein 2 chr5:8500713-8501844 REVERSE LENGTH=265 |
| AT5G62690.1 | 1,085974166 | 0,0092627 | TUB2 tubulin beta chain 2 chr5:25181560-25183501 FORWARD LENGTH=450 |
| AT2G22230.1 | 1,175439453 | 0,0093711 | no symbol available no full name available chr2:9450042-9451427 FORWARD LENGTH=220 |
| AT5G19510.1 | 0,754721116 | 0,0095452 | no symbol available no full name available chr5:6581854-6583137 REVERSE LENGTH=224 |
| AT3G46230.1 | 1,429659929 | 0,0096204 | HSP17.4_ ATHSP17.4 heat shock protein 17.4_ ARABIDOPSIS THALIANA HEAT SHOCK PROTEIN 17.4 chr3:16984 |
| AT5G16400.1 | 0,731023474 | 0,0096918 | TRXF2_ ATF2 thioredoxin F2 chr5:5363905-5365249 REVERSE LENGTH=185 |
| AT3G58610.1 | 1,344290152 | 0,0096986 | no symbol available no full name available chr3:21671561-21674639 FORWARD LENGTH=591 |
| AT4G34230.1 | 1,530336225 | 0,0097737 | CAD-5_ ATCAD5_ CAD5 cinnamyl alcohol dehydrogenase 5 chr4:16386898-16388666 REVERSE LENGTH=357 |
| AT3G23990.1 | 1,537001347 | 0,0099069 | HSP60_ HSP60-3B heat shock protein 60_ HEAT SHOCK PROTEIN 60-3B chr3:8669013-8672278 FORWARD LENGT |
| AT4G01150.1 | 0,679519323 | 0,0100237 | CURT1A CURVATURE THYLAKOID 1A chr4:493692-494668 FORWARD LENGTH=164 |
| AT1G20450.1 | 0,697083034 | 0,0100445 | LTI45_ ERD10_ LTI29 EARLY RESPONSIVE TO DEHYDRATION 10_ LOW TEMPERATURE INDUCED 45_ LOW |
| AT3G63140.1 | 0,872074271 | 0,0103351 | CSP41A chloroplast stem-loop binding protein of 41 kDa chr3:23327006-23328620 REVERSE LENGTH=406 |
| AT1G74310.1 | 1,33129516 | 0,0103394 | ATHSP101_ HSP101_ HOT1 heat shock protein 101 chr1:27936715-27939862 REVERSE LENGTH=911 |
| AT3G13580.1 | 0,462695869 | 0,0103823 | no symbol available no full name available chr3:4433809-4435109 FORWARD LENGTH=244 |
| AT3G49720.1 | 0,46492318 | 0,0104193 | CGR2 chr3:18440192-18441655 REVERSE LENGTH=261 |
| AT4G30690.2 | 0,193947846 | 0,0105056 | AtINFC-4_ SVR9L_ AtIF3- 4 SVR9-LIKE1_ Initiation factor 3-4 chr4:14960742-14962328 FORWARD LENGTH=253 |
| AT5G41520.1 | 1,434043744 | 0,0106239 | RPS10B ribosomal protein S10e B chr5:16609377-16610583 REVERSE LENGTH=180 |
| AT1G04710.1 | 1,742630947 | 0,0106651 | KAT1_ PKT4 3-KETO-ACYL-COA THIOLASE 1_ peroxisomal 3-ketoacyl-CoA thiolase 4 chr1:1321941-1324556 FORW |
| AT2G19900.1 | 1,289763191 | 0,0107641 | ATNADP-ME1_ NADP-ME1 NADP-malic enzyme 1_ Arabidopsis thaliana NADP-malic enzyme 1 chr2:8592106-859540 |
| AT4G30530.1 | 1,248318658 | 0,0107911 | GGP1 gamma-glutamyl peptidase 1 chr4:14920605-14922286 FORWARD LENGTH=250 |
| AT3G17240.1 | 0,818142842 | 0,0107976 | mtLPD2 lipoamide dehydrogenase 2 chr3:5890278-5892166 REVERSE LENGTH=507 |
| AT5G27640.1 | 1,338385357 | 0,010848 | ATTIF3B1_ TIF3B1_ EIF3B_ ATEIF3B-1_ EIF3B-1 ARABIDOPSIS THALIANA TRANSLATION INITIATION FACT |
| AT2G37190.1 | 0,780954302 | 0,0108711 | no symbol available no full name available chr2:15619559-15620059 REVERSE LENGTH=166 |
| AT5G67030.2 | 1,2139844 | 0,0108949 | LOS6_ NPQ2_ ZEP_ ABA1_ ATABA1_ ATZEP_ IBS3 ABA DEFICIENT 1_ IMPAIRED IN BABA-INDUCED STERIL |
| AT1G80380.4 | 1,223514272 | 0,0109426 | GLYK glycerate kinase chr1:30217332-30219784 FORWARD LENGTH=450 |
| AT1G58270.1 | 0,810871266 | 0,0110308 | ZW9 chr1:21612394-21614089 REVERSE LENGTH=396 |
| AT3G17390.1 | 0,878530944 | 0,0110718 | SAMS3_ AtSAMS3_ MAT4_ MTO3 METHIONINE ADENOSYLTRANSFERASE 4_ S-ADENOSYLMETHIONINE SY |
| AT1G23740.1 | 1,524160553 | 0,0111682 | AOR alkenal/one oxidoreductase chr1:8398245-8399656 REVERSE LENGTH=386 |
| AT1G19580.1 | 2,321629871 | 0,0112395 | GAMMA CA1 gamma carbonic anhydrase 1 chr1:6774937-6777092 FORWARD LENGTH=275 |
| AT1G78900.1 | 1,227904597 | 0,0113804 | VHA-A vacuolar ATP synthase subunit A chr1:29660463-29664575 FORWARD LENGTH=623 |
| AT2G45960.2 | 1,155914299 | 0,0115045 | TMP-A_ ATHH2_ PIP1;2_ PIP1B TRANSMEMBRANE PROTEIN A_ plasma membrane intrinsic protein 1B_ NAMED |
| AT3G63460.2 | 1,19350579 | 0,0115198 | SEC31B chr3:23431009-23437241 REVERSE LENGTH=1102 |
| AT5G13120.1 | 0,802976943 | 0,0115299 | CYP20-2_ Pnsl5_ ATCYP20-2 cyclophilin 20-2_ Photosynthetic NDH subcomplex L 5_ ARABIDOPSIS THALIANA CY |
| AT2G41090.1 | 0,382997863 | 0,0115335 | CML10 calmodulin like 10 chr2:17135823-17136618 FORWARD LENGTH=191 |
| AT1G04170.1 | 1,256738336 | 0,0115703 | EIF2 GAMMA eukaryotic translation initiation factor 2 gamma subunit chr1:1097423-1099702 FORWARD LENGTH=46 |
| AT4G27520.1 | 0,900799247 | 0,011658 | ENODL2_ AtENODL2 early nodulin-like protein 2 chr4:13750668-13751819 REVERSE LENGTH=349 |
| AT1G79550.1 | 1,503495324 | 0,011697 | PGKc_ PGK_ PGK3 phosphoglycerate kinase_ phosphoglycerate kinase 3 chr1:29924347-29926295 REVERSE LENGTH |
| AT3G48870.1 | 1,160859566 | 0,0117675 | ATCLPC_ HSP93-III_ ClpC2_ ATHSP93-III ClpC2 chr3:18122363-18126008 REVERSE LENGTH=952 |
| AT3G52990.1 | 1,23527623 | 0,0117914 | no symbol available no full name available chr3:19649046-19652237 FORWARD LENGTH=527 |

| | | | |
|---|---|---|---|
| AT5G49460.1 | 1,493529892 | 0,0118981 | ACLB-2 ATP citrate lyase subunit B 2 chr5:20055048-20058195 FORWARD LENGTH=608 |
| AT5G10920.1 | 1,525378504 | 0,0119901 | no symbol available no full name available chr5:3441805-3443892 FORWARD LENGTH=517 |
| AT1G61520.1 | 0,901964081 | 0,0120429 | LHCA3 photosystem I light harvesting complex gene 3 chr1:22700152-22701149 FORWARD LENGTH=273 |
| AT1G24020.1 | 1,295161246 | 0,0120623 | MLP423 MLP-like protein 423 chr1:8500653-8501458 REVERSE LENGTH=155 |
| AT3G18080.1 | 1,30469608 | 0,0120666 | BGLU44 B-S glucosidase 44 chr3:6191586-6194124 FORWARD LENGTH=512 |
| AT1G34430.1 | 1,241887963 | 0,012251 | EMB3003 embryo defective 3003 chr1:12588027-12590084 REVERSE LENGTH=465 |
| AT5G09650.1 | 0,744873147 | 0,0123233 | PPa6_ AtPPa6 pyrophosphorylase 6 chr5:2991331-2993117 REVERSE LENGTH=300 |
| AT4G33680.1 | 1,122709561 | 0,0123309 | AGD2 ABERRANT GROWTH AND DEATH 2_ ARF-GAP domain 2 chr4:16171847-16174630 REVERSE LENGTH=4[illegible] |
| AT3G02360.1 | 1,15106955 | 0,0123574 | PGD2 6-phosphogluconate dehydrogenase 2 chr3:482498-483958 FORWARD LENGTH=486 |
| AT4G35000.1 | 1,238578532 | 0,0123857 | APX3 ascorbate peroxidase 3 chr4:16665007-16667541 REVERSE LENGTH=287 |
| AT2G20610.1 | 1,268941408 | 0,0124326 | RTY1_ RTY_ HLS3_ ALF1_ SUR1 ABERRANT LATERAL ROOT FORMATION 1_ SUPERROOT 1_ ROOTY 1_ RO[illegible] |
| AT3G56940.1 | 0,789443316 | 0,012552 | CRD1_ ACSF_ CHL27 COPPER RESPONSE DEFECT 1 chr3:21076594-21078269 FORWARD LENGTH=409 |
| AT5G26830.1 | 1,334615551 | 0,0125846 | no symbol available no full name available chr5:9437351-9441568 FORWARD LENGTH=709 |
| AT5G11880.1 | 1,219140327 | 0,0126544 | DAPDC2 meso-diaminopimelate decarboxylase 2 chr5:3827806-3829942 REVERSE LENGTH=489 |
| AT2G01290.1 | 0,638727379 | 0,0126681 | RPI2 ribose-5-phosphate isomerase 2 chr2:149192-149989 REVERSE LENGTH=265 |
| ATCG00270.1 | 0,909737967 | 0,0127197 | PSBD photosystem II reaction center protein D chrc:32711-33772 FORWARD LENGTH=353 |
| AT2G21390.1 | 1,100763091 | 0,0128091 | no symbol available no full name available chr2:9152428-9156577 FORWARD LENGTH=1218 |
| AT3G10060.1 | 0,899387611 | 0,0130011 | no symbol available no full name available chr3:3102291-3103801 FORWARD LENGTH=230 |
| AT2G24200.1 | 1,196348919 | 0,0130298 | LAP1_ atLAP1 leucyl aminopeptidase 1 chr2:10287017-10289450 REVERSE LENGTH=520 |
| AT5G35590.1 | 1,30668614 | 0,0131004 | PAA1 proteasome alpha subunit A1 chr5:13765417-13767768 REVERSE LENGTH=246 |
| AT1G22700.3 | 0,692027583 | 0,0132463 | PYG7 chr1:8028323-8029289 REVERSE LENGTH=211 |
| AT5G11450.1 | 0,721296657 | 0,0134763 | PPD5 PsbP domain protein 5 chr5:3654475-3656357 FORWARD LENGTH=297 |
| AT3G28270.1 | 2,711453146 | 0,0134951 | AFL1 At14a-Like1 chr3:10538725-10539849 FORWARD LENGTH=374 |
| AT5G14260.1 | 1,417545646 | 0,0136245 | SAFE1 SAFEGUARD1 chr5:4601139-4603873 FORWARD LENGTH=514 |
| AT1G04480.1 | 0,909285994 | 0,0137322 | no symbol available no full name available chr1:1216110-1217257 FORWARD LENGTH=140 |
| AT1G72730.1 | 1,221928999 | 0,0138335 | no symbol available no full name available chr1:27378040-27379593 REVERSE LENGTH=414 |
| AT3G52500.1 | 0,89411172 | 0,0139435 | no symbol available no full name available chr3:19465644-19467053 REVERSE LENGTH=469 |
| AT1G79530.1 | 1,405000865 | 0,0139499 | GAPCP-1 glyceraldehyde-3-phosphate dehydrogenase of plastid 1 chr1:29916232-29919088 REVERSE LENGTH=422 |
| AT5G13650.1 | 1,172525048 | 0,0140239 | SVR3 SUPPRESSOR OF VARIEGATION 3 chr5:4397821-4402364 FORWARD LENGTH=675 |
| AT1G52030.1 | 1,545611835 | 0,0141268 | F-ATMBP_ MBP1.2_ MBP2 myrosinase-binding protein 2 chr1:19346090-19348282 REVERSE LENGTH=642 |
| AT4G12800.1 | 0,851101558 | 0,0142632 | PSAL photosystem I subunit l chr4:7521469-7522493 FORWARD LENGTH=219 |
| AT3G13460.2 | 1,336704511 | 0,0143034 | ECT2 evolutionarily conserved C-terminal region 2 chr3:4385274-4388220 REVERSE LENGTH=664 |
| AT5G61170.1 | 0,816663815 | 0,0143539 | no symbol available no full name available chr5:24611158-24612202 FORWARD LENGTH=143 |
| AT1G56410.1 | 0,809124014 | 0,0143817 | HSP70T-1_ ERD2 HEAT SHOCK PROTEIN 70T-1_ EARLY-RESPONSIVE TO DEHYDRATION 2 chr1:21117147-211[illegible] |
| AT5G45280.1 | 0,326840813 | 0,0144662 | PAE11 pectin acetylesterase 11 chr5:18346862-18349432 FORWARD LENGTH=370 |
| AT2G21410.1 | 1,791325765 | 0,0145269 | VHA-A2 vacuolar proton ATPase A2 chr2:9162703-9168141 FORWARD LENGTH=821 |
| AT5G10540.1 | 0,704758556 | 0,0146908 | TOP2 thimet metalloendopeptidase 2 chr5:3328119-3332462 FORWARD LENGTH=701 |
| AT4G30910.1 | 1,159462016 | 0,0146956 | no symbol available no full name available chr4:15042621-15045248 REVERSE LENGTH=581 |
| ATCG00130.1 | 1,179524122 | 0,0148309 | ATPF chrc:11529-12798 REVERSE LENGTH=184 |

| | | | |
|---|---|---|---|
| AT1G01100.3 | 1,230335589 | 0,0149023 | RPP1A_ RPP1.1 60S acidic ribosomal protein P1-1_ RPP1 co-orthologous gene 1 chr1:50284-50954 REVERSE LENGTH= |
| AT5G60640.2 | 0,742125771 | 0,014927 | PDI2_ ATPDIL1-4_ ATPDI2_ PDIL1-4 PROTEIN DISULFIDE ISOMERASE 2_ ARABIDOPSIS THALIANA PROTEIN |
| AT2G05710.1 | 1,474959664 | 0,0149716 | ACO3 aconitase 3 chr2:2141591-2146350 FORWARD LENGTH=990 |
| AT4G09040.1 | 0,710517193 | 0,0150724 | CP33C chr4:5795075-5797315 REVERSE LENGTH=304 |
| AT3G07110.1 | 0,460311798 | 0,0157102 | no symbol available no full name available chr3:2252092-2253332 FORWARD LENGTH=206 |
| AT1G09640.1 | 1,228841137 | 0,0159662 | no symbol available no full name available chr1:3120162-3122152 FORWARD LENGTH=414 |
| AT5G15530.1 | 1,710367011 | 0,0161178 | BCCP2_ CAC1-B biotin carboxyl carrier protein 2 chr5:5038955-5040437 FORWARD LENGTH=255 |
| AT3G56130.1 | 0,587585384 | 0,0161582 | BADC1_ BLP3 biotin/lipoyl attachment domain containing 1_ BCCP-Like Protein 3 chr3:20826852-20829007 FORWARD |
| AT5G08670.1 | 1,124582823 | 0,016402 | no symbol available no full name available chr5:2818395-2821149 REVERSE LENGTH=556 |
| AT4G39730.1 | 1,197127531 | 0,0174559 | ATPLAT1_ PLAT1 PLAT domain protein 1_ "Polycystin_ Lipoxygenase_ Alpha-toxin and Triacylglycerol lipase 1" chr4:1 |
| AT5G55220.1 | 0,887005989 | 0,0175921 | HP65b_ TIG1 chr5:22397677-22400678 FORWARD LENGTH=547 |
| AT3G55610.1 | 3,05341998 | 0,0175998 | P5CS2 delta 1-pyrroline-5-carboxylate synthase 2 chr3:20624278-20628989 REVERSE LENGTH=726 |
| AT3G58510.1 | 1,168506386 | 0,0176521 | RH11 RNA Helicase 11 chr3:21640608-21643464 FORWARD LENGTH=612 |
| AT2G22780.1 | 1,654277801 | 0,0178044 | PMDH1 peroxisomal NAD-malate dehydrogenase 1 chr2:9689995-9691923 REVERSE LENGTH=354 |
| AT3G13920.4 | 1,313921424 | 0,0179081 | TIF4A1_ EIF4A1_ RH4 eukaryotic translation initiation factor 4A1 chr3:4592635-4594128 REVERSE LENGTH=407 |
| AT3G60820.1 | 1,150136473 | 0,0181399 | PBF1 chr3:22472038-22473809 REVERSE LENGTH=223 |
| AT5G58290.1 | 0,631300726 | 0,018208 | RPT3 regulatory particle triple-A ATPase 3 chr5:23569155-23571116 FORWARD LENGTH=408 |
| AT2G13360.1 | 1,1574658 | 0,0184152 | SGAT_ AGT_ AGT1 ALANINE:GLYOXYLATE AMINOTRANSFERASE 1_ alanine:glyoxylate aminotransferase_ L-ser |
| AT1G12410.1 | 1,135218788 | 0,0185402 | EMB3146_ CLPR2_ CLP2_ NCLPP2 EMBRYO DEFECTIVE 3146_ CLP protease proteolytic subunit 2_ NUCLEAR-EN |
| AT2G18960.1 | 1,124864472 | 0,0185712 | HA1_ OST2_ AHA1_ PMA H(+)-ATPase 1_ PLASMA MEMBRANE PROTON ATPASE_ OPEN STOMATA 2 chr2:82 |
| AT3G07770.1 | 1,327816737 | 0,0186141 | Hsp89.1_ AtHsp90.6_ AtHsp90-6 HEAT SHOCK PROTEIN 90.6_ HEAT SHOCK PROTEIN 89.1_ HEAT SHOCK PRO |
| AT2G24020.1 | 1,309233614 | 0,0187369 | STIC2 Suppressor of TIC40 2 chr2:10217869-10219269 REVERSE LENGTH=182 |
| AT3G22960.1 | 1,206433689 | 0,0188557 | PKP1_ PKP-ALPHA PLASTIDIAL PYRUVATE KINASE 1 chr3:8139369-8141771 FORWARD LENGTH=596 |
| AT2G22170.1 | 1,503455527 | 0,0188767 | PLAT2 PLAT domain protein 2 chr2:9427010-9427742 REVERSE LENGTH=183 |
| AT3G28290.1 | 1,110316678 | 0,0189207 | AT14A chr3:10547873-10549030 FORWARD LENGTH=385 |
| AT2G20260.1 | 0,892755445 | 0,0190046 | PSAE-2 photosystem I subunit E-2 chr2:8736780-8737644 FORWARD LENGTH=145 |
| AT5G42740.3 | 1,201770698 | 0,019028 | no symbol available no full name available chr5:17136269-17140622 FORWARD LENGTH=528 |
| AT5G28840.1 | 1,177136051 | 0,019065 | GME GDP-D-mannose 3 chr5:10862472-10864024 REVERSE LENGTH=377 |
| AT3G11510.1 | 0,672225645 | 0,0193022 | no symbol available no full name available chr3:3623757-3624866 REVERSE LENGTH=150 |
| AT3G47650.1 | 1,392212885 | 0,0193914 | BSD2 BUNDLE SHEATH DEFECTIVE 2 chr3:17569576-17570262 FORWARD LENGTH=136 |
| AT1G54220.1 | 0,719139186 | 0,0194017 | mtE2-3 mitochondrial pyruvate dehydrogenase subunit 2-3 chr1:20246460-20250208 REVERSE LENGTH=539 |
| AT5G63680.1 | 0,338017844 | 0,0195298 | no symbol available no full name available chr5:25490507-25492530 FORWARD LENGTH=510 |
| AT2G42600.1 | 1,151364786 | 0,0198295 | ATPPC2_ PPC2 phosphoenolpyruvate carboxylase 2 chr2:17734541-17738679 REVERSE LENGTH=963 |
| AT2G36250.4 | 1,125533873 | 0,019867 | FTSZ2-1_ ATFTSZ2-1 chr2:15197661-15199689 REVERSE LENGTH=397 |
| AT3G57490.1 | 0,592218205 | 0,0201081 | no symbol available no full name available chr3:21279824-21280887 REVERSE LENGTH=276 |
| AT5G22800.1 | 1,21827686 | 0,0204382 | EMB263_ EMB1030_ EMB86 EMBRYO DEFECTIVE 263_ EMBRYO DEFECTIVE 1030_ EMBRYO DEFECTIVE 86 |
| AT1G74920.2 | 1,620289931 | 0,0206929 | ALDH10A8 aldehyde dehydrogenase 10A8 chr1:28139175-28142573 REVERSE LENGTH=496 |
| AT4G00660.1 | 1,568347232 | 0,0207736 | RH8_ ATRH8 RNAhelicase-like 8 chr4:274638-277438 FORWARD LENGTH=505 |
| AT4G26970.1 | 1,304625261 | 0,020916 | ACO2 aconitase 2 chr4:13543077-13548427 FORWARD LENGTH=995 |

| | | | |
|---|---|---|---|
| AT3G04870.1 | 1,278441342 | 0,021156 | PDE181_ ZDS_ SPC1 SPONTANEOUS CELL DEATH 1_ PIGMENT DEFECTIVE EMBRYO 181_ zeta-carotene desatu |
| AT2G02930.1 | 1,098529063 | 0,0211618 | ATGSTF3_ GST16_ GSTF3 GLUTATHIONE S-TRANSFERASE 16_ glutathione S-transferase F3 chr2:851348-852106 F |
| AT1G51760.1 | 0,731566985 | 0,0211678 | IAR3_ JR3 IAA-ALANINE RESISTANT 3_ JASMONIC ACID RESPONSIVE 3 chr1:19199562-19201424 FORWARD L |
| AT1G21750.1 | 1,174017448 | 0,0214195 | PDIL1-1_ ATPDI5_ PDI5_ ATPDIL1-1 PDI-like 1-1_ ARABIDOPSIS THALIANA PROTEIN DISULFIDE ISOMERASI |
| AT3G11830.1 | 1,281161789 | 0,0218207 | CCT7 Chaperonin containing T-complex polypeptide-1 subunit 7 chr3:3732734-3736156 FORWARD LENGTH=557 |
| AT1G02500.1 | 0,906198918 | 0,021842 | METK1_ SAM-1_ AtSAM1_ SAM1_ MAT1 S-adenosylmethionine synthetase 1_ S-ADENOSYLMETHIONINE SYNTHI |
| AT4G01800.1 | 1,306979468 | 0,0221488 | SECA1_ AGY1_ AtcpSecA Arabidopsis thaliana chloroplast SecA_ Albino or Glassy Yellow 1 chr4:770926-776131 REVE |
| AT5G25880.3 | 0,828581001 | 0,0223166 | ATNADP-ME3_ NADP-ME3 NADP-malic enzyme 3_ Arabidopsis thaliana NADP-malic enzyme 3 chr5:9024549-902749 |
| AT1G23730.1 | 0,61034958 | 0,0223718 | ATBCA3_ BCA3 beta carbonic anhydrase 3_ BETA CARBONIC ANHYDRASE 3 chr1:8395965-8398014 FORWARD LI |
| AT2G43460.1 | 0,658940453 | 0,0224273 | no symbol available no full name available chr2:18046285-18047292 REVERSE LENGTH=69 |
| AT3G58990.1 | 1,628059584 | 0,0224601 | IPMI SSU3_ IPMI1 isopropylmalate isomerase 1 chr3:21797524-21798285 REVERSE LENGTH=253 |
| AT2G04700.1 | 0,787089453 | 0,0227984 | INAP1_ FTRB Imbalanced NADP Status 1_ ferredoxin/thioredoxin reductase catayltic subunit chr2:1646961-1648345 FOR |
| AT2G47400.1 | 0,808771423 | 0,023307 | CP12_ CP12-1 CP12 DOMAIN-CONTAINING PROTEIN 1_ CP12 domain-containing protein 1 chr2:19446889-1944726 |
| AT2G28900.1 | 1,262955488 | 0,0234101 | OEP16_ OEP16-1_ ATOEP16-L_ ATOEP16-1 outer plastid envelope protein 16-1_ OUTER PLASTID ENVELOPE PRO |
| AT1G74090.1 | 0,461238486 | 0,0235337 | ATST5B_ SOT18_ ATSOT18 DESULFO-GLUCOSINOLATE SULFOTRANSFERASE 18_ ARABIDOPSIS SULFOTR |
| AT5G52310.1 | 0,503652697 | 0,0235711 | COR78_ LTI140_ RD29A_ LTI78 RESPONSIVE TO DESICCATION 29A_ LOW-TEMPERATURE-INDUCED 78_ CO |
| AT4G09320.1 | 1,077028554 | 0,0235894 | ATNDK1_ NDPK1_ NDK1 nucleoside diphosphate kinase 1 chr4:5923484-5924366 FORWARD LENGTH=149 |
| AT4G08870.1 | 1,121097849 | 0,0236883 | ARGAH2 arginine amidohydrolase 2 chr4:5646654-5648693 REVERSE LENGTH=344 |
| AT2G30870.1 | 1,160917143 | 0,0238092 | GSTF10_ ATGSTF10_ ERD13_ ATGSTF4 glutathione S-transferase PHI 10_ ARABIDOPSIS THALIANA GLUTATHIO |
| AT2G02010.1 | 0,295922146 | 0,0240293 | GAD4 glutamate decarboxylase 4 chr2:474375-476495 REVERSE LENGTH=493 |
| AT5G15090.1 | 1,156289637 | 0,0240931 | VDAC3_ AtVDAC-3_ ATVDAC3 ARABIDOPSIS THALIANA VOLTAGE DEPENDENT ANION CHANNEL 3_ voltag |
| AT2G34480.1 | 0,112907842 | 0,0241367 | L18aB_ RPL18aB chr2:14532916-14534161 REVERSE LENGTH=178 |
| AT2G23350.1 | 1,270182584 | 0,0242845 | PABP4_ PAB4 POLY(A) BINDING PROTEIN 4_ poly(A) binding protein 4 chr2:9943209-9946041 FORWARD LENGT |
| AT5G44070.2 | 0,816347382 | 0,0244402 | CAD1_ ARA8_ PCS1_ ATPCS1 PHYTOCHELATIN SYNTHASE 1_ CADMIUM SENSITIVE 1_ ARABIDOPSIS THAI |
| AT3G16050.1 | 3,560771684 | 0,024488 | ATPDX1.2_ A37_ PDX1.2 pyridoxine biosynthesis 1.2_ ARABIDOPSIS THALIANA PYRIDOXINE BIOSYNTHESIS 1. |
| AT2G41220.1 | 0,772865944 | 0,024782 | GLU2 glutamate synthase 2 chr2:17177934-17188388 FORWARD LENGTH=1629 |
| AT3G08030.2 | 0,919131862 | 0,0253307 | AthA2-1 chr3:2564517-2565819 FORWARD LENGTH=323 |
| AT3G53990.1 | 0,876425603 | 0,0254972 | AtUSP_ USP17 Universal stress protein chr3:19989658-19991019 REVERSE LENGTH=160 |
| AT1G24510.3 | 1,373209234 | 0,0259029 | CCT5 Chaperonin containing T-complex polypeptide-1 subunit 5 chr1:8685504-8687802 REVERSE LENGTH=482 |
| AT4G33220.2 | 1,745100755 | 0,0259453 | ATPME44_ PME44 pectin methylesterase 44_ A. THALIANA PECTIN METHYLESTERASE 44 chr4:16024445-160261 |
| AT5G15650.1 | 0,850266431 | 0,0259585 | MUR5_ ATRGP2_ RGP2 MURUS 5_ REVERSIBLY GLYCOSYLATED POLYPEPTIDE 2_ reversibly glycosylated poly |
| AT1G48420.1 | 1,279026548 | 0,0261002 | ACD1_ DCD_ ATACD1_ D-CDES_ AtDCD 1-AMINOCYCLOPROPANE-1-CARBOXYLIC ACID DEAMINASE 1_ A. |
| AT1G12310.1 | 0,913359603 | 0,0262595 | no symbol available no full name available chr1:4187500-4187946 REVERSE LENGTH=148 |
| AT2G46820.1 | 0,684711768 | 0,0262781 | PTAC8_ PSI-P_ PSAP_ CURT1B_ TMP14 CURVATURE THYLAKOID 1B_ PLASTID TRANSCRIPTIONALLY ACT |
| AT5G09660.1 | 1,181604065 | 0,0263391 | PMDH2 peroxisomal NAD-malate dehydrogenase 2 chr5:2993645-2995551 REVERSE LENGTH=354 |
| AT3G04120.1 | 1,108538242 | 0,026419 | GAPC1_ GAPC_ GAPC-1 GLYCERALDEHYDE-3-PHOSPHATE DEHYDROGENASE C SUBUNIT_ glyceraldehyde-3 |
| AT3G02080.1 | 1,542372089 | 0,026526 | no symbol available no full name available chr3:364138-365161 REVERSE LENGTH=143 |
| AT5G47190.1 | 1,469050285 | 0,0272286 | PRPL19 plastid ribosomal proteins of the 50S subunit 19 chr5:19164432-19166064 REVERSE LENGTH=229 |
| AT5G62350.1 | 0,555009955 | 0,0273128 | no symbol available no full name available chr5:25037504-25038112 FORWARD LENGTH=202 |

| | | | |
|---|---|---|---|
| AT5G16990.1 | 0,773397791 | 0,0274221 | no symbol available no full name available chr5:5581831-5583849 REVERSE LENGTH=343 |
| AT1G53240.1 | 1,211115105 | 0,0277263 | mMDH1 mitochondrial malate dehydrogenase 1 chr1:19854966-19856802 REVERSE LENGTH=341 |
| AT4G18440.1 | 1,062808328 | 0,0278301 | no symbol available no full name available chr4:10186385-10188832 REVERSE LENGTH=536 |
| AT1G29900.1 | 1,301827473 | 0,0279194 | CARB_ VEN3 carbamoyl phosphate synthetase B_ VENOSA 3 chr1:10468164-10471976 FORWARD LENGTH=1187 |
| AT1G77590.2 | 1,473177491 | 0,0279457 | LACS9 long chain acyl-CoA synthetase 9 chr1:29148501-29151234 REVERSE LENGTH=545 |
| AT1G27400.1 | 1,308926844 | 0,0282465 | no symbol available no full name available chr1:9515230-9516725 FORWARD LENGTH=176 |
| AT1G66240.1 | 1,264379034 | 0,0282467 | AtHMP14_ ATX1_ ATATX1 HEAVY METAL ASSOCIATED PROTEIN 14_ homolog of anti-oxidant 1 chr1:24686445-[illegible] |
| AT5G65270.1 | 1,261951708 | 0,0283124 | AtRABA4a_ RABA4a RAB GTPase homolog A4A chr5:26083437-26084550 FORWARD LENGTH=226 |
| AT2G37660.1 | 0,928874006 | 0,0284551 | no symbol available no full name available chr2:15795481-15796977 REVERSE LENGTH=325 |
| AT5G57350.3 | 0,394578756 | 0,0285077 | ATAHA3_ HA3_ AHA3 H(+)-ATPase 3_ ARABIDOPSIS THALIANA ARABIDOPSIS H(+)-ATPASE chr5:23231208-2[illegible] |
| AT4G33010.1 | 1,113025433 | 0,0285724 | GLDP1_ AtGLDP1 glycine decarboxylase P-protein 1 chr4:15926852-15931150 REVERSE LENGTH=1037 |
| AT4G37925.1 | 0,836043998 | 0,0286261 | NDH-M_ NdhM subunit NDH-M of NAD(P)H:plastoquinone dehydrogenase complex_ NADH dehydrogenase-like comple[illegible] |
| AT2G26340.1 | 0,822020942 | 0,0288664 | no symbol available no full name available chr2:11215254-11216480 FORWARD LENGTH=253 |
| AT4G34180.1 | 0,499471993 | 0,0290184 | CYCLASE1 CYCLASE1 chr4:16370060-16371383 REVERSE LENGTH=255 |
| AT5G37510.1 | 1,39771862 | 0,0292106 | EMB1467_ CI76 embryo defective 1467 chr5:14897490-14900352 FORWARD LENGTH=745 |
| AT4G14040.1 | 1,278561942 | 0,0293215 | EDA38_ SBP2 selenium-binding protein 2_ EMBRYO SAC DEVELOPMENT ARREST 38 chr4:8100691-8102828 REVE[illegible] |
| AT1G07320.3 | 0,691372783 | 0,0294289 | RPL4_ PRPL4_ EMB2784 plastid ribosomal protein L4_ ribosomal protein L4_ EMBRYO DEFECTIVE 2784 chr1:22491[illegible] |
| AT1G27950.1 | 1,356047851 | 0,0296057 | LTPG1 glycosylphosphatidylinositol-anchored lipid protein transfer 1 chr1:9740740-9741991 FORWARD LENGTH=193 |
| AT1G48630.1 | 0,888949326 | 0,0297748 | RACK1B_ RACK1B_AT receptor for activated C kinase 1B chr1:17981977-17983268 REVERSE LENGTH=326 |
| AT1G63660.2 | 1,791441101 | 0,0301084 | no symbol available no full name available chr1:23604874-23607080 REVERSE LENGTH=434 |
| AT5G01600.1 | 1,564087199 | 0,0302644 | FER1_ ATFER1 ferretin 1_ ARABIDOPSIS THALIANA FERRETIN 1 chr5:228149-229594 REVERSE LENGTH=255 |
| AT3G02730.1 | 0,892537594 | 0,0306383 | TRXF1_ ATF1 thioredoxin F-type 1 chr3:588570-589591 REVERSE LENGTH=178 |
| AT1G64740.1 | 0,921715325 | 0,0312057 | TUA1 alpha-1 tubulin chr1:24050114-24052296 FORWARD LENGTH=450 |
| AT2G37270.1 | 1,087508616 | 0,0313279 | RPS5B_ ATRPS5B ribosomal protein 5B chr2:15647883-15649042 REVERSE LENGTH=207 |
| AT5G40950.1 | 0,809404626 | 0,0314422 | PRPL27_ RPL27 ribosomal protein large subunit 27 chr5:16410866-16411845 FORWARD LENGTH=198 |
| AT2G30110.1 | 0,776870625 | 0,0314492 | MOS5_ ATUBA1_ UBA1 MODIFIER OF SNC1 5_ ubiquitin-activating enzyme 1 chr2:12852632-12857369 REVERSE L[illegible] |
| AT3G20390.1 | 1,131852027 | 0,0315008 | RidA Reactive Intermediate Deaminase A chr3:7110227-7111695 REVERSE LENGTH=187 |
| AT3G16390.1 | 0,86272862 | 0,0315771 | NSP3 nitrile specifier protein 3 chr3:5562602-5564356 FORWARD LENGTH=467 |
| AT4G24770.1 | 1,222738594 | 0,031616 | ATRBP33_ ATRBP31_ CP31A_ RBP31_ CP33a_ CP31 "ARABIDOPSIS THALIANA RNA BINDING PROTEIN_ APP[illegible] |
| AT1G06000.1 | 1,689393143 | 0,0316926 | UGT89C1 chr1:1820495-1821802 REVERSE LENGTH=435 |
| AT3G16470.1 | 1,072941965 | 0,0319476 | JAL35_ AtJAC1_ JR1 jacalin-related lectin 35_ JACALIN-LECTIN LIKE 1_ JASMONATE RESPONSIVE 1 chr3:55960[illegible] |
| AT1G50250.1 | 0,53187555 | 0,0319789 | FTSH1 FTSH protease 1 chr1:18614398-18616930 REVERSE LENGTH=716 |
| AT5G56500.1 | 0,675381487 | 0,0321126 | CPNB3_ Cpn60beta3 chaperonin-60beta3 chr5:22874058-22876966 FORWARD LENGTH=597 |
| AT5G13850.1 | 0,85319988 | 0,0321281 | NACA3 nascent polypeptide-associated complex subunit alpha-like protein 3 chr5:4471361-4472676 FORWARD LENGT[illegible] |
| AT1G31180.1 | 1,100804983 | 0,0324501 | ATIMD3_ IMDH3_ IMD3_ IPMDH1 ISOPROPYLMALATE DEHYDROGENASE 1_ isopropylmalate dehydrogenase 3_ |
| AT3G47470.1 | 0,862179982 | 0,0326739 | LHCA4_ CAB4 light-harvesting chlorophyll-protein complex I subunit A4 chr3:17493622-17494773 REVERSE LENGTH[illegible] |
| AT3G48990.1 | 1,289893204 | 0,0331032 | AAE3 ACYL-ACTIVATING ENZYME 3 chr3:18159031-18161294 REVERSE LENGTH=514 |
| AT1G04410.1 | 1,177892216 | 0,0336862 | c-NAD-MDH1 cytosolic-NAD-dependent malate dehydrogenase 1 chr1:1189418-1191267 REVERSE LENGTH=332 |
| AT3G57560.1 | 1,127987832 | 0,0339712 | NAGK N-acetyl-l-glutamate kinase chr3:21311164-21312207 REVERSE LENGTH=347 |

| | | | |
|---|---|---|---|
| ATCG00680.1 | 0,911449706 | 0,0339896 | PSBB photosystem II reaction center protein B chrc:72371-73897 FORWARD LENGTH=508 |
| AT3G08940.2 | 0,913843852 | 0,0340156 | LHCB4.2 light harvesting complex photosystem II chr3:2717717-2718665 FORWARD LENGTH=287 |
| AT5G48810.1 | 0,657812732 | 0,0340847 | CB5-D_ ATB5-B_ ATCB5-D_ B5 #3_ CYTB5-B cytochrome B5 isoform D_ ARABIDOPSIS CYTOCHROME B5 ISOF( |
| AT1G27450.1 | 1,162049839 | 0,0342356 | APT1_ ATAPT1 ARABIDOPSIS THALIANA ADENINE PHOSPHORIBOSYLTRANSFERASE 1_ adenine phosphoribo |
| AT5G17920.1 | 1,159290598 | 0,0343236 | ATCIMS_ METS1_ ATMETS_ ATMS1 COBALAMIN-INDEPENDENT METHIONINE SYNTHASE_ methionine synth |
| AT5G60360.2 | 0,4319806 | 0,0347571 | AALP_ SAG2_ ALP aleurain-like protease_ SENESCENCE ASSOCIATED GENE2 chr5:24280044-24282152 FORWARI |
| AT5G09900.1 | 1,246524686 | 0,03482 | EMB2107_ MSA_ RPN5A MARIPOSA_ EMBRYO DEFECTIVE 2107_ REGULATORY PARTICLE NON-ATPASE SU |
| AT2G20630.1 | 1,11178911 | 0,0348253 | PIA1 PP2C induced by AVRRPM1 chr2:8897826-8899648 REVERSE LENGTH=279 |
| AT1G54040.2 | 0,449335569 | 0,0349276 | TASTY_ ESR_ ESP epithiospecifier protein_ EPITHIOSPECIFYING SENESCENCE REGULATOR chr1:20170995-2017 |
| AT1G56340.1 | 1,23926442 | 0,0351224 | AtCRT1a_ CRT1_ CRT1a calreticulin 1a_ calreticulin 1 chr1:21090059-21092630 REVERSE LENGTH=425 |
| AT3G12290.1 | 1,245963667 | 0,0352076 | MTHFD1 methylenetetrahydrofolate dehydrogenase/methenyltetrahydrofolate cyclohydrolase chr3:3919591-3921326 FOR\ |
| AT5G48580.1 | 0,771447245 | 0,0352829 | FKBP15-2 FK506- and rapamycin-binding protein 15 kD-2 chr5:19696156-19697304 REVERSE LENGTH=163 |
| AT5G24300.1 | 1,103760709 | 0,0353712 | SS1_ ATSS1 STARCH SYNTHASE 1_ starch synthase 1 chr5:8266934-8270860 FORWARD LENGTH=652 |
| AT4G15545.1 | 0,47325484 | 0,0355027 | NAIP1 NAI2-interacting protein 1 chr4:8875932-8877567 FORWARD LENGTH=337 |
| ATCG00740.1 | 0,899591136 | 0,0357308 | RPOA RNA polymerase subunit alpha chrc:77901-78890 REVERSE LENGTH=329 |
| AT1G59870.1 | 0,919897926 | 0,0359171 | ABCG36_ ATABCG36_ PEN3_ PDR8_ ATPDR8 Arabidopsis thaliana ATP-binding cassette G36_ PLEIOTROPIC DRU( |
| AT5G14660.1 | 0,675732536 | 0,0360543 | ATDEF2_ DEF2_ PDF1B peptide deformylase 1B chr5:4727129-4728671 REVERSE LENGTH=273 |
| AT5G47520.1 | 1,128975353 | 0,036513 | RABA5a_ AtRABA5a RAB GTPase homolog A5A chr5:19277596-19278366 REVERSE LENGTH=221 |
| AT5G20010.1 | 1,15715462 | 0,0366306 | RAN-1_ ATRAN1_ RAN1 RAS-RELATED NUCLEAR PROTEIN_ ARABIDOPSIS THALIANA RAS-RELATED NUCI |
| AT5G20920.2 | 0,749942831 | 0,0367403 | EIF2 BETA_ EMB1401_ eIF-2bs embryo defective 1401_ eukaryotic translation initiation factor 2 beta subunit chr5:70949 |
| AT1G09310.1 | 0,720741738 | 0,0370545 | SVB2_ SVBL SVB-like chr1:3009109-3009648 FORWARD LENGTH=179 |
| AT1G70310.1 | 1,197248717 | 0,0374031 | SPDS2 spermidine synthase 2 chr1:26485497-26487352 REVERSE LENGTH=340 |
| AT1G23820.1 | 1,176657787 | 0,037573 | SPDS1 spermidine synthase 1 chr1:8420410-8422724 FORWARD LENGTH=334 |
| AT4G16660.1 | 1,510646618 | 0,0378015 | HSP70 heat shock protein 70 chr4:9377225-9381232 FORWARD LENGTH=867 |
| AT5G27470.1 | 1,190781212 | 0,0380431 | no symbol available no full name available chr5:9695087-9697154 FORWARD LENGTH=451 |
| AT5G52920.1 | 1,343285753 | 0,0380673 | PKP-BETA1_ PKP2_ PKP1 plastidic pyruvate kinase beta subunit 1_ PLASTIDIAL PYRUVATE KINASE 1_ PLASTIDIA |
| AT3G25920.1 | 0,835563514 | 0,0383307 | RPL15 ribosomal protein L15 chr3:9491268-9492558 REVERSE LENGTH=277 |
| AT5G28500.1 | 1,126245925 | 0,0383768 | no symbol available no full name available chr5:10477810-10479114 FORWARD LENGTH=434 |
| AT5G50950.3 | 0,830318741 | 0,0387009 | FUM2 FUMARASE 2 chr5:20731191-20733636 FORWARD LENGTH=317 |
| AT1G66410.1 | 0,548884136 | 0,0388617 | CAM4_ ACAM-4 calmodulin 4_ CALMODULIN 4 chr1:24774431-24775785 REVERSE LENGTH=149 |
| AT3G14390.1 | 1,263222525 | 0,0389213 | DAPDC1 meso-diaminopimelate decarboxylase 1 chr3:4806771-4808954 FORWARD LENGTH=484 |
| AT3G59970.3 | 1,10493458 | 0,0390973 | MTHFR1 methylenetetrahydrofolate reductase 1 chr3:22151303-22154323 FORWARD LENGTH=592 |
| AT2G44160.1 | 1,266064366 | 0,0394232 | MTHFR2 methylenetetrahydrofolate reductase 2 chr2:18262301-18265185 FORWARD LENGTH=594 |
| AT1G50480.1 | 1,170725857 | 0,0394544 | THFS 10-formyltetrahydrofolate synthetase chr1:18702064-18704687 FORWARD LENGTH=634 |
| AT1G68560.1 | 0,83122894 | 0,0395376 | XYL1_ GH31_ TRG1_ ATXYL1_ AXY3 altered xyloglucan 3_ ALPHA-XYLOSIDASE 1_ alpha-xylosidase 1_ thermoinh |
| AT3G49870.1 | 1,180697335 | 0,0395764 | ARLA1C_ ARL8a_ ATARLA1C ADP-ribosylation factor-like A1C_ ADP-ribosylation factor-like 8a chr3:18492674-18494 |
| AT1G70890.1 | 1,256192925 | 0,0395765 | MLP43 MLP-like protein 43_ major latex protein like 43 chr1:26725912-26726489 REVERSE LENGTH=158 |
| AT4G35100.1 | 0,350738226 | 0,0396418 | PIP3A_ SIMIP_ PIP3_ PIP2;7 plasma membrane intrinsic protein 3_ PLASMA MEMBRANE INTRINSIC PROTEIN 3A_ |
| AT4G09670.1 | 1,252259251 | 0,0398013 | no symbol available no full name available chr4:6107382-6109049 REVERSE LENGTH=362 |

| | | | |
|---|---|---|---|
| AT5G43940.1 | 1,053396502 | 0,0399263 | ADH2_ HOT5_ ATGSNOR1_ PAR2_ GSNOR PARAQUAT RESISTANT 2_ S-NITROSOGLUTATHIONE REDUCTAS |
| AT2G40290.1 | 0,84981588 | 0,039997 | no symbol available no full name available chr2:16829030-16830889 REVERSE LENGTH=344 |
| AT2G21620.1 | 0,797727872 | 0,0401684 | RD2 RESPONSIVE TO DESICCATION 2 chr2:9248749-9249986 FORWARD LENGTH=187 |
| AT4G30610.1 | 0,707492438 | 0,0402795 | BRS1_ SCPL24 BRI1 SUPPRESSOR 1_ SERINE CARBOXYPEPTIDASE 24 PRECURSOR chr4:14944219-14948391 F |
| AT1G10840.1 | 1,360231009 | 0,0408854 | TIF3H1 translation initiation factor 3 subunit H1 chr1:3607885-3610299 REVERSE LENGTH=337 |
| AT1G22740.1 | 0,424486242 | 0,0413197 | RABG3B_ RAB7_ ATRABG3B_ RAB75 RAB GTPase homolog G3B chr1:8049247-8050494 FORWARD LENGTH=20 |
| AT3G52150.1 | 0,864833982 | 0,0413201 | PSRP2 plastid-speciﬁc ribosomal protein 2 chr3:19342074-19343090 FORWARD LENGTH=253 |
| AT5G47840.2 | 0,771747717 | 0,0414074 | AMK2 adenosine monophosphate kinase chr5:19375488-19378058 FORWARD LENGTH=269 |
| AT5G50850.1 | 1,169954884 | 0,0415241 | MAB1 MACCI-BOU chr5:20689671-20692976 FORWARD LENGTH=363 |
| AT2G22990.1 | 0,513474689 | 0,0416949 | SNG1_ SCPL8 sinapoylglucose 1_ SERINE CARBOXYPEPTIDASE-LIKE 8 chr2:9786393-9789925 FORWARD LENG |
| AT3G48690.1 | 6,987287415 | 0,0417831 | ATCXE12_ CXE12 ARABIDOPSIS THALIANA CARBOXYESTERASE 12 chr3:18037186-18038160 REVERSE LENG |
| AT5G36700.1 | 1,116279934 | 0,0418867 | ATPGLP1_ PGLP1 2-phosphoglycolate phosphatase 1 chr5:14421929-14424430 REVERSE LENGTH=362 |
| AT5G59890.2 | 0,761543226 | 0,0420003 | ATADF4_ ADF4 actin depolymerizing factor 4 chr5:24123107-24123596 FORWARD LENGTH=132 |
| AT3G15356.1 | 0,871559676 | 0,0420792 | no symbol available no full name available chr3:5174603-5175418 REVERSE LENGTH=271 |
| AT3G63490.1 | 0,922967817 | 0,0421618 | EMB3126_ PRPL1 proline-rich protein-like 1_ plastid ribosomal protein L1_ EMBRYO DEFECTIVE 3126 chr3:2344426 |
| AT5G48900.1 | 0,811625779 | 0,0422549 | no symbol available no full name available chr5:19825240-19828909 FORWARD LENGTH=417 |
| AT3G10350.1 | 0,819929717 | 0,0425883 | AtGET3b_ GET3b Guided Entry of Tail-anchored proteins 3b chr3:3208310-3210678 FORWARD LENGTH=411 |
| AT3G12915.2 | 0,796953348 | 0,0426611 | no symbol available no full name available chr3:4112834-4115708 FORWARD LENGTH=788 |
| AT1G09430.1 | 1,294853291 | 0,0430316 | ACLA-3 ATP-citrate lyase A-3 chr1:3042135-3044978 FORWARD LENGTH=424 |
| AT1G79230.3 | 1,183924736 | 0,0432842 | ATMST1_ STR1_ MST1_ ATRDH1_ ST1 ARABIDOPSIS THALIANA RHODANESE HOMOLOGUE 1_ mercaptopyru |
| AT3G09640.1 | 0,228411212 | 0,0435663 | AtAPX2_ APX1B_ APX2 ASCORBATE PEROXIDASE 1B_ ascorbate peroxidase 2 chr3:2956301-2958163 FORWARD |
| AT3G52730.1 | 0,782097942 | 0,0437714 | no symbol available no full name available chr3:19543146-19544167 REVERSE LENGTH=72 |
| AT2G43750.1 | 1,100477505 | 0,0442093 | OASB_ CPACS1_ ACS1_ ATCS-B O-acetylserine (thiol) lyase B_ ARABIDOPSIS THALIANA CYSTEIN SYNTHASE- |
| AT5G19820.1 | 1,382503438 | 0,0443737 | IMB3_ KETCH1_ EMB2734 EMBRYO DEFECTIVE 2734_ (karyopherin enabling the transport of the cytoplasmic HYL1 |
| AT1G56110.1 | 1,304282592 | 0,0444375 | NOP56 homolog of nucleolar protein NOP56 chr1:20984544-20986893 REVERSE LENGTH=522 |
| AT1G29880.1 | 1,233994106 | 0,0445769 | no symbol available no full name available chr1:10459662-10462781 REVERSE LENGTH=729 |
| AT5G54900.1 | 1,64580647 | 0,0446802 | RBP45A_ ATRBP45A RNA-binding protein 45A chr5:22295412-22298126 FORWARD LENGTH=387 |
| AT3G63170.1 | 0,685417957 | 0,0447253 | FAP1_ AtFAP1 fatty-acid-binding protein 1 chr3:23334675-23335993 FORWARD LENGTH=279 |
| AT1G28290.2 | 0,962347288 | 0,0451126 | AGP31 arabinogalactan protein 31 chr1:9889331-9890843 REVERSE LENGTH=315 |
| AT3G16460.1 | 1,380504624 | 0,0451333 | JAL34 jacalin-related lectin 34 chr3:5593029-5595522 FORWARD LENGTH=705 |
| AT5G40770.1 | 1,2114561 | 0,0452586 | ATPHB3_ PHB3_ EER3 prohibitin 3 chr5:16315589-16316621 REVERSE LENGTH=277 |
| AT4G15210.3 | 1,388927224 | 0,0453514 | AT-BETA-AMY_ ATBETA-AMY_ BMY1_ RAM1_ BAM5 REDUCED BETA AMYLASE 1_ beta-amylase 5_ ARABID |
| AT1G05190.1 | 0,885190157 | 0,045359 | RPL6_ EMB2394 embryo defective 2394 chr1:1502515-1503738 REVERSE LENGTH=223 |
| AT3G29360.1 | 1,282327827 | 0,045534 | UGD2 UDP-glucose dehydrogenase 2 chr3:11267375-11268817 REVERSE LENGTH=480 |
| AT4G31990.1 | 1,266915346 | 0,04558 | AAT3_ ATAAT1_ ASP5 aspartate aminotransferase 5_ ASPARTATE AMINOTRANSFERASE DEFICIENT 3 chr4:1547 |
| AT2G04390.1 | 0,891998392 | 0,045756 | dS17 chr2:1527911-1528336 FORWARD LENGTH=141 |
| AT5G08380.1 | 0,880387375 | 0,0458503 | AtAGAL1_ AGAL1 alpha-galactosidase 1 chr5:2694851-2697616 REVERSE LENGTH=410 |
| AT5G06870.1 | 0,945173725 | 0,0461646 | PGIP2_ ATPGIP2 ARABIDOPSIS POLYGALACTURONASE INHIBITING PROTEIN 2_ polygalacturonase inhibiting p |
| AT3G15730.1 | 1,296649625 | 0,0462582 | PLD_ PLDALPHA1 phospholipase D alpha 1 chr3:5330835-5333474 FORWARD LENGTH=810 |

| | | | |
|---|---|---|---|
| AT1G71500.1 | 0,649593671 | 0,046266 | PSB33_ LIL8 PhotoSystem B protein 33_ Light-harvesting-like 8 chr1:26936084-26937331 FORWARD LENGTH=287 |
| AT2G43030.1 | 0,749249572 | 0,0463177 | PRPL3 plastid ribosomal proteins of the 50S subunit chr2:17894898-17895713 FORWARD LENGTH=271 |
| AT3G27300.1 | 1,409839069 | 0,0465392 | G6PD5 glucose-6-phosphate dehydrogenase 5 chr3:10083318-10086288 REVERSE LENGTH=516 |
| AT5G03630.1 | 1,447135791 | 0,0470295 | MDAR2 chr5:922378-924616 REVERSE LENGTH=435 |
| AT5G47930.1 | 0,552438587 | 0,0471572 | no symbol available no full name available chr5:19406423-19407329 REVERSE LENGTH=84 |
| AT4G28750.1 | 0,87597013 | 0,0473961 | PSAE-1 PSA E1 KNOCKOUT chr4:14202951-14203888 REVERSE LENGTH=143 |
| ATCG01060.1 | 1,107887703 | 0,0479861 | PSAC chrc:117318-117563 REVERSE LENGTH=81 |
| AT5G35790.1 | 1,167639561 | 0,0480103 | G6PD1 glucose-6-phosphate dehydrogenase 1 chr5:13956879-13959686 REVERSE LENGTH=576 |
| AT4G25630.1 | 0,817107427 | 0,0481521 | ATFIB2_ FIB2 fibrillarin 2 chr4:13074239-13076205 FORWARD LENGTH=320 |
| AT2G30200.1 | 1,244844864 | 0,0481762 | EMB3147_ MCAT_ MCAMT EMBRYO DEFECTIVE 3147_ malonyl CoA-ACP malonyltransferase chr2:12883162-1288 |
| AT5G54640.1 | 1,32816807 | 0,0487981 | HTA1_ RAT5_ ATHTA1 histone H2A 1_ RESISTANT TO AGROBACTERIUM TRANSFORMATION 5 chr5:22196540 |
| AT3G26070.1 | 0,32363795 | 0,0489912 | FBN3a FIBRILLIN3a chr3:9526904-9528199 FORWARD LENGTH=242 |
| AT3G32980.1 | 1,103754497 | 0,0490531 | PRX32 Peroxidase 32 chr3:13526404-13529949 REVERSE LENGTH=352 |
| AT3G02530.1 | 1,154555774 | 0,0495045 | CCT6-2 Chaperonin containing T-complex polypeptide-1 subunit 6-2 chr3:528806-532457 REVERSE LENGTH=535 |
| AT2G44120.1 | 1,081391128 | 0,0497647 | no symbol available no full name available chr2:18249227-18250402 REVERSE LENGTH=242 |

GENE SILENCING 1_ MATERNAL EFFECT EMBRYO ARREST 58_ S-ADENOSYL-L-HOMOCYSTEIN HYDROLASE 1 chr4:8054931-8056676 FORWARD LENGT[illegible]

ABIDOPSIS THALIANA CINNAMYL-ALCOHOL DEHYDROGENASE 8_ ELICITOR-ACTIVATED GENE 3 chr4:17855964-17857388 FORWARD LENGTH=359

?ROTEIN 1_ GLYCINE RICH PROTEIN 7_ "cold_ circadian rhythm_ and rna binding 2"_ GLYCINE-RICH RNA-BINDING PROTEIN 7 chr2:9265477-9266316 REVERS

:EIN 70-1_ heat shock cognate protein 70-1_ HEAT SHOCK COGNATE PROTEIN 70_ HEAT SHOCK PROTEIN 70-1 chr5:554055-556334 REVERSE LENGTH=651

ST CHAPERONIN 60ALPHA_ ACCUMULATION AND REPLICATION OF CHLOROPLASTS 2_ chaperonin-60alpha chr2:11926603-11929184 FORWARD LENGTH=

SIS THALIANA PROTEIN DISULFIDE ISOMERASE 11_ UNFERTILIZED EMBRYO SAC 5_ MATERNAL EFFECT EMBRYO ARREST 30 chr2:19481503-19483683

-1_ ARABIDOPSIS THALIANA HOMOLOG OF BACTERIAL CYTOKINESIS Z-RING PROTEIN FTSZ 1-1 chr5:22420740-22422527 REVERSE LENGTH=433

PROTEIN 1_ microtubule-destabilizing protein 25_ plasma-membrane associated cation-binding protein 1 chr4:10941593-10943227 FORWARD LENGTH=225

ing factor_ "ribosome recycling factor_ chloroplast precursor"_ HIGH CHLOROPHYLL FLUORESCENCE AND PALE GREEN MUTANT 108 chr3:23342861-23344640 R

OR 3B1_ EUKARYOTIC TRANSLATION INITIATION FACTOR 3B1_ EUKARYOTIC TRANSLATION INITIATION FACTOR 3B_ translation initiation factor 3B1 ch

ITY 3_ NON-PHOTOCHEMICAL QUENCHING 2_ ARABIDOPSIS THALIANA ZEAXANTHIN EPOXIDASE_ LOW EXPRESSION OF OSMOTIC STRESS-RESPON

THALIANA 1-AMINOCYCLOPROPANE-1-CARBOXYLIC ACID DEAMINASE 1_ D-cysteine desulfhydrase chr1:17896767-17898803 REVERSE LENGTH=401

G RESISTANCE 8_ ARABIDOPSIS PLEIOTROPIC DRUG RESISTANCE 8_ PENETRATION 3_ ATP-binding cassette G36 chr1:22034661-22039844 FORWARD LENG

B_ CHLOROPLAST O-ACETYLSERINE SULFHYDRYLASE 1_ ARABIDOPSIS CYSTEINE SYNTHASE 1 chr2:18129604-18132322 REVERSE LENGTH=392

SIVE GENES 6_ ARABIDOPSIS THALIANA ABA DEFICIENT 1_ ZEAXANTHIN EPOXIDASE chr5:26754026-26757090 REVERSE LENGTH=610
