## Supplementary material for "HY5 enhances *Arabidopsis* tolerance to combined high light and heat stress by coordinating photoprotection and hormone signaling": Table S4

**Supplemental Table S4. Differentially accumulated proteins compared to control (P < 0.05) in Col leaves subjected to subjected to combined conditions of high ligh**

| Protein ID | Fold Change | p-value | Protein description |
|---|---|---|---|
| AT3G53420.1 | 1,380077349 | 6,26689E-08 | PIP2;1_ AtPIP2;1_ PIP2A_ PIP2 PLASMA MEMBRANE INTRINSIC PROTEIN 2_ PLASMA MEMBRANI |
| AT5G48570.1 | 20,89196334 | 1,71503E-07 | ROF2_ ATFKBP65_ FKBP65 chr5:19690746-19693656 REVERSE LENGTH=578 |
| AT5G12020.1 | 23,41882993 | 2,93873E-07 | HSP17.6II 17.6 kDa class II heat shock protein chr5:3882409-3882876 REVERSE LENGTH=155 |
| AT1G09180.1 | 1,361791149 | 3,48739E-07 | SAR1A_ ATSARA1A_ SARA1A_ ATSAR1 SECRETION-ASSOCIATED RAS 1_ secretion-associated RAS |
| AT5G52640.1 | 19,03320316 | 3,67688E-07 | HSP81-1_ ATHSP90.1_ HSP81.1_ AtHsp90-1_ HSP83_ ATHS83_ HSP90.1 HEAT SHOCK PROTEIN 90-1_ |
| AT4G36130.1 | 0,036298069 | 4,091E-07 | no symbol available no full name available chr4:17097613-17098656 FORWARD LENGTH=258 |
| AT3G08030.2 | 0,524956857 | 4,11146E-07 | AthA2-1 chr3:2564517-2565819 FORWARD LENGTH=323 |
| AT1G74310.1 | 33,9008089 | 4,21006E-07 | ATHSP101_ HSP101_ HOT1 heat shock protein 101 chr1:27936715-27939862 REVERSE LENGTH=911 |
| AT5G12030.1 | 32,28489869 | 5,10304E-07 | AT-HSP17.6A_ HSP17.6A_ HSP17.6 HEAT SHOCK PROTEIN 17.6_ heat shock protein 17.6A chr5:388421 |
| AT4G17090.1 | 1,747842098 | 8,89821E-07 | CT-BMY_ BMY8_ AtBAM3_ BAM3 BETA-AMYLASE 8_ BETA-AMYLASE 3_ chloroplast beta-amylase c |
| AT3G13470.1 | 1,761457882 | 9,8923E-07 | CPNB2_ Cpn60beta2 chaperonin-60beta2 chr3:4389685-4392624 FORWARD LENGTH=596 |
| AT5G03300.1 | 1,901386532 | 1,03402E-06 | ADK2 adenosine kinase 2 chr5:796573-798997 FORWARD LENGTH=345 |
| AT5G51440.1 | 17,45098039 | 1,10409E-06 | HSP23.5 chr5:20891242-20892013 FORWARD LENGTH=210 |
| AT5G59720.1 | 93,00350026 | 1,60096E-06 | HSP18.2 heat shock protein 18.2 chr5:24062632-24063117 FORWARD LENGTH=161 |
| AT3G12580.1 | 8,067081023 | 1,60839E-06 | HSP70_ ATHSP70_ HSC70-4 ARABIDOPSIS HEAT SHOCK PROTEIN 70_ heat shock protein 70 chr3:399 |
| AT5G15450.1 | 3,501427572 | 1,66908E-06 | APG6_ AtCLPB3_ CLPB3_ CLPB-P casein lytic proteinase B3_ CASEIN LYTIC PROTEINASE B-P_ ALBI |
| AT1G16030.1 | 3,913959074 | 1,92637E-06 | Hsp70b heat shock protein 70B chr1:5502386-5504326 REVERSE LENGTH=646 |
| AT4G24190.1 | 1,934032516 | 1,96366E-06 | SHD_ HSP90_ HSP90.7_ AtHsp90-7_ AtHsp90.7 SHEPHERD_ HEAT SHOCK PROTEIN 90.7_ HEAT SHC |
| AT5G05010.1 | 1,956194258 | 2,05362E-06 | no symbol available no full name available chr5:1477137-1479872 FORWARD LENGTH=527 |
| AT1G07400.1 | 38,74972818 | 2,0606E-06 | HSP17.8 chr1:2275148-2275621 FORWARD LENGTH=157 |
| AT2G29500.1 | 43,72532014 | 2,15119E-06 | HSP17.6B chr2:12633279-12633740 REVERSE LENGTH=153 |
| AT1G11840.1 | 3,518221282 | 2,47103E-06 | AtGLYI3_ GLX1_ ATGLX1 glyoxalaseI 3_ glyoxalase I homolog chr1:3996045-3997518 FORWARD LENG |
| AT4G04020.1 | 6,808109833 | 2,6801E-06 | FIB_ PGL35_ FIB1a plastoglobulin 35_ fibrillin 1a_ fibrillin chr4:1932161-1933546 FORWARD LENGTH=3 |
| AT3G09440.1 | 2,186778171 | 2,74235E-06 | no symbol available no full name available chr3:2903434-2905632 REVERSE LENGTH=649 |
| AT2G35370.1 | 0,546850475 | 3,09547E-06 | GDCH glycine decarboxylase complex H chr2:14891239-14892050 FORWARD LENGTH=165 |
| AT5G09590.1 | 3,381562988 | 3,31079E-06 | MTHSC70-2_ HSC70-5 mitochondrial HSO70 2_ HEAT SHOCK COGNATE chr5:2975721-2978508 FORW |
| AT3G07090.1 | 8,519058732 | 4,39514E-06 | Desi1 chr3:2243153-2244476 REVERSE LENGTH=265 |
| AT5G12110.1 | 6,292718784 | 4,64674E-06 | no symbol available no full name available chr5:3914483-3915732 FORWARD LENGTH=228 |
| AT1G59860.1 | 15,70652096 | 4,85981E-06 | HSP17.6A chr1:22031474-22031941 FORWARD LENGTH=155 |
| ATCG00830.1 | 0,042686261 | 5,70821E-06 | RPL2.1 ribosomal protein L2 chrc:84337-85843 REVERSE LENGTH=274 |
| AT3G09350.1 | 21,35123155 | 6,13277E-06 | Fes1A Fes1A chr3:2871216-2873109 FORWARD LENGTH=363 |
| AT3G46230.1 | 7,623725616 | 6,51967E-06 | HSP17.4_ ATHSP17.4 heat shock protein 17.4_ ARABIDOPSIS THALIANA HEAT SHOCK PROTEIN 17.4 |
| AT5G44020.1 | 4,510824074 | 6,67739E-06 | no symbol available no full name available chr5:17712433-17714046 FORWARD LENGTH=272 |
| AT4G21960.1 | 0,154212433 | 6,7092E-06 | PRXR1 chr4:11646613-11648312 REVERSE LENGTH=330 |
| AT4G02080.1 | 0,645004043 | 7,9749E-06 | ATSARA1C_ SAR2_ SAR1C_ ATSAR2_ ASAR1 secretion-associated RAS super family 2 chr4:921554-9225 |
| AT2G35635.1 | 14,0770221 | 8,77064E-06 | UBQ7_ RUB2 RELATED TO UBIQUITIN 2_ ubiquitin 7 chr2:14981044-14981943 FORWARD LENGTH=1 |

| | | | |
|---|---|---|---|
| AT2G20560.1 | 38,03532703 | 9,13016E-06 | DNAJ DNAJ protein chr2:8848353-8849815 REVERSE LENGTH=337 |
| AT4G25630.1 | 0,505674875 | 9,13976E-06 | ATFIB2_ FIB2 fibrillarin 2 chr4:13074239-13076205 FORWARD LENGTH=320 |
| AT5G15520.1 | 0,416178876 | 9,85104E-06 | no symbol available no full name available chr5:5037242-5038136 REVERSE LENGTH=143 |
| AT1G01100.1 | 2,338692991 | 1,07271E-05 | RPP1A_ RPP1.1 60S acidic ribosomal protein P1-1_ RPP1 co-orthologous gene 1 chr1:50284-50954 REVERS |
| AT1G79920.1 | 2,195807244 | 1,13334E-05 | Hsp70-15_ AtHsp70-15 heat shock protein 70-15 chr1:30059302-30062224 REVERSE LENGTH=736 |
| AT5G01410.1 | 1,72052406 | 1,13847E-05 | PDX1_ ATPDX1.3_ ATPDX1_ PDX1.3_ RSR4 REDUCED SUGAR RESPONSE 4_ PYRIDOXINE BIOSYN |
| AT3G52380.1 | 0,379330463 | 1,19931E-05 | PDE322_ CP33 PIGMENT DEFECTIVE 322_ chloroplast RNA-binding protein 33 chr3:19421619-19422855 |
| AT4G25200.1 | 8,648726249 | 1,2028E-05 | ATHSP23.6-MITO_ HSP23.6-MITO mitochondrion-localized small heat shock protein 23.6 chr4:12917089-12 |
| AT3G26060.1 | 0,723885367 | 1,23169E-05 | ATPRX Q_ PRXQ peroxiredoxin Q chr3:9524807-9526123 FORWARD LENGTH=216 |
| AT3G16050.1 | 56,04362159 | 1,31634E-05 | ATPDX1.2_ A37_ PDX1.2 pyridoxine biosynthesis 1.2_ ARABIDOPSIS THALIANA PYRIDOXINE BIOSY |
| AT2G39310.1 | 5,262859 | 1,349E-05 | JAL22 jacalin-related lectin 22 chr2:16414262-16416323 REVERSE LENGTH=458 |
| AT2G42600.1 | 1,362174903 | 1,42453E-05 | ATPPC2_ PPC2 phosphoenolpyruvate carboxylase 2 chr2:17734541-17738679 REVERSE LENGTH=963 |
| AT1G36280.2 | 0,298327602 | 1,49532E-05 | no symbol available no full name available chr1:13640600-13642908 FORWARD LENGTH=519 |
| AT2G21660.1 | 0,277735147 | 1,54914E-05 | RBGA3_ GR-RBP7_ CCR2_ ATGRP7_ GRP7_ SRBP1 RNA-binding glycine-rich protein A3_ SMALL RNA- |
| AT3G53990.1 | 0,307713163 | 1,55593E-05 | AtUSP_ USP17 Universal stress protein chr3:19989658-19991019 REVERSE LENGTH=160 |
| AT5G10160.1 | 1,361203997 | 1,58546E-05 | no symbol available no full name available chr5:3185819-3187159 FORWARD LENGTH=219 |
| AT3G57260.1 | 0,179496352 | 1,63255E-05 | AtBG2_ PR2_ GNS2_ AtPR2_ BG2_ PR-2_ BGL2 "beta-1_3-glucanase 2"_ PATHOGENESIS-RELATED PF |
| AT5G15970.1 | 11,54508807 | 1,69357E-05 | AtCor6.6_ KIN2_ COR6.6 COLD-RESPONSIVE 6.6 chr5:5211966-5212441 FORWARD LENGTH=66 |
| AT3G11130.1 | 2,00670555 | 1,72564E-05 | AtCHC1_ CHC1_ HAS1 clathrin heavy chain 1_ hot ABA deficiency suppressor 1 chr3:3482575-3491667 REV |
| AT2G18450.1 | 1,680759496 | 1,76562E-05 | SDH1-2 succinate dehydrogenase 1-2 chr2:7997510-8000801 REVERSE LENGTH=632 |
| AT5G65430.2 | 1,408882674 | 1,85795E-05 | 14-3-3KAPPA_ GRF8_ AtMIN10_ GF14 KAPPA general regulatory factor 8_ 14-3-3 PROTEIN G-BOX FAC |
| AT3G25230.1 | 4,435309428 | 1,96548E-05 | FKBP62_ ROF1_ ATFKBP62 rotamase FKBP 1_ FK506 BINDING PROTEIN 62 chr3:9188257-9191137 FO |
| AT5G20290.1 | 0,171454956 | 1,99765E-05 | no symbol available no full name available chr5:6851695-6853012 REVERSE LENGTH=222 |
| AT1G56070.1 | 1,592286684 | 2,05659E-05 | LOS1 LOW EXPRESSION OF OSMOTICALLY RESPONSIVE GENES 1 chr1:20968245-20971077 REVER |
| AT3G49120.1 | 0,582386774 | 2,1436E-05 | PRX34_ PRXCB_ ATPCB_ AtPRX34_ PERX34_ ATPERX34 ARABIDOPSIS THALIANA PEROXIDASE ( |
| AT1G57720.1 | 1,672786206 | 2,19802E-05 | no symbol available no full name available chr1:21377873-21380114 FORWARD LENGTH=413 |
| AT1G04710.1 | 1,956383137 | 2,45201E-05 | KAT1_ PKT4 3-KETO-ACYL-COA THIOLASE 1_ peroxisomal 3-ketoacyl-CoA thiolase 4 chr1:1321941-132 |
| AT5G06870.1 | 0,661856943 | 2,46463E-05 | PGIP2_ ATPGIP2 ARABIDOPSIS POLYGALACTURONASE INHIBITING PROTEIN 2_ polygalacturonase |
| AT2G37170.1 | 2,077137294 | 2,73737E-05 | PIP2;2_ PIP2B plasma membrane intrinsic protein 2_ PLASMA MEMBRANE INTRINSIC PROTEIN 2;2 chr |
| AT1G56330.1 | 1,660145528 | 2,82379E-05 | ATSARA1B_ SAR1B_ SAR1_ ATSAR1B_ ATSAR1 SECRETION-ASSOCIATED RAS 1_ ARABIDOPSIS |
| AT5G11170.1 | 1,897221702 | 2,90385E-05 | UAP56a homolog of human UAP56 a chr5:3553334-3556646 FORWARD LENGTH=427 |
| AT1G03680.1 | 0,562461392 | 2,9595E-05 | ATHM1_ TRX-M1_ ATM1_ THM1 thioredoxin M-type 1_ THIOREDOXIN M-TYPE 1_ ARABIDOPSIS TH |
| AT4G27440.1 | 0,293329039 | 3,02476E-05 | PORB protochlorophyllide oxidoreductase B chr4:13725648-13727107 FORWARD LENGTH=401 |
| AT1G50670.1 | 2,402906966 | 3,03471E-05 | OTU2 ovarian tumor domain (OTU)-containing DUB (deubiquitilating enzyme) 2 chr1:18775086-18776552 RI |
| AT4G13430.1 | 0,680407646 | 3,55798E-05 | IIL1_ ATLEUC1 isopropyl malate isomerase large subunit 1 chr4:7804194-7807789 REVERSE LENGTH=50 |
| AT1G09640.1 | 1,750727758 | 3,59359E-05 | no symbol available no full name available chr1:3120162-3122152 FORWARD LENGTH=414 |
| AT2G28000.1 | 1,627136707 | 3,91371E-05 | ARC2_ CH-CPN60A_ SLP_ CPN60A_ Cpn60alpha1_ CPNA1 SCHLEPPERLESS_ chaperonin-60alpha1_ CH |
| AT5G23860.1 | 2,742066346 | 3,93464E-05 | TUB8 tubulin beta 8 chr5:8042962-8044528 FORWARD LENGTH=449 |
| AT5G13650.1 | 0,372332158 | 3,97912E-05 | SVR3 SUPPRESSOR OF VARIEGATION 3 chr5:4397821-4402364 FORWARD LENGTH=675 |

| | | | |
|---|---|---|---|
| AT5G26780.1 | 0,655174773 | 4,26621E-05 | SHM2 serine hydroxymethyltransferase 2 chr5:9418299-9421725 FORWARD LENGTH=517 |
| AT4G22240.1 | 1,756025502 | 4,32329E-05 | FBN1b fibrillin 1b chr4:11766090-11767227 REVERSE LENGTH=310 |
| AT3G49010.4 | 0,063071894 | 4,52956E-05 | BBC1_ RSU2_ ATBBC1 40S RIBOSOMAL PROTEIN_ breast basic conserved 1 chr3:18166971-18168047 R |
| AT1G47260.1 | 0,615533221 | 4,67983E-05 | APFI_ GAMMA CA2 gamma carbonic anhydrase 2 chr1:17321384-17323347 REVERSE LENGTH=278 |
| AT1G16470.1 | 0,528479669 | 4,80805E-05 | PAB1 proteasome subunit PAB1 chr1:5623122-5625439 FORWARD LENGTH=235 |
| AT2G37220.1 | 0,324280349 | 4,83655E-05 | no symbol available no full name available chr2:15634980-15636331 REVERSE LENGTH=289 |
| AT5G43010.1 | 0,229972267 | 4,86589E-05 | RPT4A regulatory particle triple-A ATPase 4A chr5:17248563-17251014 REVERSE LENGTH=399 |
| AT1G20260.1 | 1,754885822 | 5,04854E-05 | AtVAB3_ VAB3 V-ATPase B subunit 3 chr1:7016971-7020290 FORWARD LENGTH=487 |
| AT5G52470.1 | 2,254718169 | 5,11274E-05 | ATFIB1_ FBR1_ FIB1_ SKIP7_ ATFBR1 fibrillarin 1_ FIBRILLARIN 1_ SKP1/ASK1-INTERACTING PRC |
| AT4G34450.1 | 2,057892524 | 5,60719E-05 | gamma2-COP gamma2 Coat Protein chr4:16471956-16476795 FORWARD LENGTH=886 |
| AT5G54770.1 | 1,54353283 | 5,61917E-05 | THI1_ TZ_ THI4 THIAZOLE REQUIRING_ THIAMINE4 chr5:22246634-22247891 FORWARD LENGTH= |
| AT3G08900.1 | 0,057689042 | 5,91702E-05 | RGP_ RGP3 reversibly glycosylated polypeptide 3 chr3:2708347-2709714 REVERSE LENGTH=362 |
| AT2G33380.1 | 1,895706128 | 5,92245E-05 | AtRD20_ PXG3_ CLO-3_ CLO3_ RD20_ AtCLO3 caleosin 3_ Arabidopsis thaliana caleosin 3_ peroxygenase |
| AT3G24503.1 | 0,563628049 | 6,35387E-05 | REF1_ ALDH2C4_ ALDH1A REDUCED EPIDERMAL FLUORESCENCE1_ aldehyde dehydrogenase 2C4_ |
| AT3G02530.1 | 1,371140252 | 6,39943E-05 | CCT6-2 Chaperonin containing T-complex polypeptide-1 subunit 6-2 chr3:528806-532457 REVERSE LENGT |
| AT5G37670.1 | 17,08919166 | 6,56879E-05 | HSP15.7 chr5:14969035-14969448 FORWARD LENGTH=137 |
| AT3G46520.1 | 2,29540762 | 7,08441E-05 | ACT12 actin-12 chr3:17128567-17129981 FORWARD LENGTH=377 |
| AT2G42910.1 | 3,893129971 | 7,67823E-05 | AtPRS4_ PRS4 phosphoribosyl diphosphate synthase 4 chr2:17856396-17858394 FORWARD LENGTH=337 |
| AT5G48375.1 | 0,372074986 | 8,99223E-05 | TGG3_ BGLU39 thioglucoside glucosidase 3_ BETA GLUCOSIDASE 39 chr5:19601303-19603883 REVERS |
| AT5G56030.1 | 2,388207422 | 9,33668E-05 | HSP81-2_ HSP90.2_ AtHsp90.2_ ERD8_ HSP81.2 EARLY-RESPONSIVE TO DEHYDRATION 8_ heat sho |
| AT4G18100.1 | 0,093243171 | 9,38533E-05 | no symbol available no full name available chr4:10035715-10036475 REVERSE LENGTH=133 |
| AT1G54340.1 | 2,851740108 | 0,000102822 | ICDH isocitrate dehydrogenase chr1:20283520-20286506 FORWARD LENGTH=416 |
| AT3G12145.1 | 0,467504854 | 0,000107635 | FLR1_ FTM4 FLOR1_ FLORAL TRANSITION AT THE MERISTEM4 chr3:3874764-3876075 REVERSE L |
| AT4G35830.1 | 1,54698514 | 0,00010864 | ACO1 aconitase 1 chr4:16973007-16977949 REVERSE LENGTH=898 |
| AT1G04480.1 | 0,692243112 | 0,000111282 | no symbol available no full name available chr1:1216110-1217257 FORWARD LENGTH=140 |
| AT4G37300.1 | 0,67267241 | 0,000114866 | MEE59 maternal effect embryo arrest 59 chr4:17554805-17555498 FORWARD LENGTH=173 |
| AT5G06290.1 | 1,288144614 | 0,000117854 | 2-Cys Prx B_ 2CPB 2-cysteine peroxiredoxin B_ 2-CYS PEROXIREDOXIN B chr5:1919380-1921211 FORW |
| AT1G03130.1 | 0,555278712 | 0,000118869 | PSAD-2 photosystem I subunit D-2 chr1:753528-754142 REVERSE LENGTH=204 |
| AT3G63460.2 | 1,69496285 | 0,000119252 | SEC31B chr3:23431009-23437241 REVERSE LENGTH=1102 |
| AT3G45030.1 | 0,340453969 | 0,000122888 | no symbol available no full name available chr3:16471606-16472312 REVERSE LENGTH=124 |
| AT4G10320.1 | 1,939527267 | 0,000123963 | no symbol available no full name available chr4:6397526-6404509 REVERSE LENGTH=1190 |
| AT2G37270.1 | 0,638790712 | 0,000124822 | RPS5B_ ATRPS5B ribosomal protein 5B chr2:15647883-15649042 REVERSE LENGTH=207 |
| AT1G58270.1 | 0,567505868 | 0,000125711 | ZW9 chr1:21612394-21614089 REVERSE LENGTH=396 |
| AT3G04920.2 | 0,123569168 | 0,000125915 | no symbol available no full name available chr3:1360989-1361719 FORWARD LENGTH=112 |
| AT5G46290.2 | 0,900507692 | 0,000126264 | KASI_ KAS1 3-ketoacyl-acyl carrier protein synthase I_ KETOACYL-ACP SYNTHASE 1 chr5:18774439-18[illegible] |
| AT5G56000.1 | 0,84773883 | 0,000126703 | Hsp81.4_ AtHsp90.4 HEAT SHOCK PROTEIN 90.4_ HEAT SHOCK PROTEIN 81.4 chr5:22677602-226800 |
| AT2G30490.1 | 1,451682696 | 0,000128507 | REF3_ CYP73A5_ C4H_ ATC4H CINNAMATE 4-HYDROXYLASE_ REDUCED EPRDERMAL FLUORES |
| AT4G27520.1 | 0,664180172 | 0,000128837 | ENODL2_ AtENODL2 early nodulin-like protein 2 chr4:13750668-13751819 REVERSE LENGTH=349 |
| AT1G22700.3 | 0,573461633 | 0,000130034 | PYG7 chr1:8028323-8029289 REVERSE LENGTH=211 |

| | | | |
|---|---|---|---|
| AT3G57290.1 | 1,640211153 | 0,000130072 | EIF3E_ ATINT6_ TIF3E1_ INT6_ INT-6_ ATEIF3E-1 eukaryotic translation initiation factor 3E chr3:2119678 |
| AT5G02160.1 | 0,636363636 | 0,000133093 | FIP FtsH5 Interacting Protein chr5:426392-427024 FORWARD LENGTH=129 |
| AT3G25800.1 | 1,465215151 | 0,000134289 | PP2AA2_ PDF1_ PR 65 protein phosphatase 2A subunit A2 chr3:9422822-9425783 REVERSE LENGTH=587 |
| AT2G25450.1 | 0,48487893 | 0,000143258 | GSL-OH glucosinolate hydroxylase chr2:10830286-10831563 REVERSE LENGTH=359 |
| AT5G24780.1 | 0,698199917 | 0,000146418 | VSP1_ ATVSP1 vegetative storage protein 1 chr5:8507783-8508889 REVERSE LENGTH=270 |
| AT4G16830.2 | 0,588044725 | 0,000146907 | AtRGGA chr4:9470979-9472308 FORWARD LENGTH=265 |
| AT2G10940.1 | 1,950321624 | 0,000148891 | no symbol available no full name available chr2:4311160-4312035 REVERSE LENGTH=291 |
| AT2G32060.1 | 0,751892142 | 0,000150618 | no symbol available no full name available chr2:13639228-13640104 REVERSE LENGTH=144 |
| AT1G13320.4 | 1,717673117 | 0,000154073 | PP2AA3 protein phosphatase 2A subunit A3 chr1:4563692-4566451 REVERSE LENGTH=412 |
| AT1G47250.1 | 1,5054611 | 0,000159257 | PAF2 20S proteasome alpha subunit F2 chr1:17319220-17320900 FORWARD LENGTH=277 |
| AT2G39390.1 | 0,134259297 | 0,000160052 | no symbol available no full name available chr2:16450803-16451762 REVERSE LENGTH=123 |
| AT5G64140.1 | 0,565470824 | 0,000163029 | RPS28 ribosomal protein S28 chr5:25667529-25667723 REVERSE LENGTH=64 |
| AT2G18960.1 | 1,582391058 | 0,000169275 | HA1_ OST2_ AHA1_ PMA H(+)-ATPase 1_ PLASMA MEMBRANE PROTON ATPASE_ OPEN STOMAT |
| AT5G45280.2 | 0,79449946 | 0,000180423 | PAE11 pectin acetylesterase 11 chr5:18346862-18349488 FORWARD LENGTH=391 |
| AT5G35590.1 | 0,238021672 | 0,000183908 | PAA1 proteasome alpha subunit A1 chr5:13765417-13767768 REVERSE LENGTH=246 |
| AT5G65010.1 | 1,795705243 | 0,0001842 | ASN2 asparagine synthetase 2 chr5:25969224-25972278 FORWARD LENGTH=578 |
| AT3G46060.1 | 1,944931322 | 0,000187458 | ARA3_ RAB8A_ RABE1c_ ARA-3_ ATRAB8A_ ATRABE1C RAB GTPase homolog 8A chr3:16917908-169 |
| AT1G78830.1 | 0,727358676 | 0,00018749 | MNB1 chr1:29637141-29638508 REVERSE LENGTH=455 |
| AT4G38740.1 | 0,725568548 | 0,000187841 | ROC1 rotamase CYP 1 chr4:18083620-18084138 REVERSE LENGTH=172 |
| AT1G48600.1 | 2,646521265 | 0,000191598 | AtPMT2_ AtPMEAMT_ PMEAMT phosphoethanolamine N-methyltransferase_ Phosphoethanolamine methylt |
| AT1G25490.1 | 1,480018271 | 0,000197589 | EER1_ ATB BETA BETA_ RCN1_ REGA ROOTS CURL IN NPA_ ENHANCED ETHYLENE RESPONSE |
| AT2G24200.1 | 1,441479463 | 0,000200012 | LAP1_ atLAP1 leucyl aminopeptidase 1 chr2:10287017-10289450 REVERSE LENGTH=520 |
| AT1G18070.1 | 1,716009639 | 0,000203671 | EF-1alpha chr1:6214236-6218211 REVERSE LENGTH=532 |
| AT5G63400.1 | 0,657958674 | 0,000207635 | ADK1 adenylate kinase 1 chr5:25393274-25394817 REVERSE LENGTH=246 |
| AT3G19760.1 | 2,924065122 | 0,000210448 | EIF4A-III_ RH2 eukaryotic initiation factor 4A-III chr3:6863790-6866242 FORWARD LENGTH=408 |
| AT1G23740.1 | 1,670417362 | 0,000218612 | AOR alkenal/one oxidoreductase chr1:8398245-8399656 REVERSE LENGTH=386 |
| AT5G66760.1 | 0,754616786 | 0,00021897 | SDH1-1 succinate dehydrogenase 1-1 chr5:26653776-26657224 FORWARD LENGTH=634 |
| AT4G29130.1 | 1,549400711 | 0,000219634 | GIN2_ HXK1_ ATHXK1 GLUCOSE INSENSITIVE 2_ hexokinase 1_ ARABIDOPSIS THALIANA HEXOK |
| AT2G06850.1 | 1,505088327 | 0,000221034 | EXT_ XTH4_ EXGT-A1 ENDOXYLOGLUCAN TRANSFERASE_ endoxyloglucan transferase A1_ xylogluc |
| AT5G64040.1 | 0,863565012 | 0,000221514 | PSAN chr5:25628724-25629409 REVERSE LENGTH=171 |
| AT3G08590.1 | 1,877124081 | 0,00022343 | iPGAM2 "2_3-biphosphoglycerate-independent phosphoglycerate mutase 2" chr3:2608683-2611237 REVERSI |
| AT2G29550.1 | 1,653230743 | 0,000225908 | TUB7_ TBB7 tubulin beta-7 chain_ tubulin beta 7 chr2:12644258-12645932 REVERSE LENGTH=449 |
| AT4G26970.1 | 1,516041159 | 0,000235854 | ACO2 aconitase 2 chr4:13543077-13548427 FORWARD LENGTH=995 |
| AT5G27640.1 | 1,88311791 | 0,000248897 | ATTIF3B1_ TIF3B1_ EIF3B_ ATEIF3B-1_ EIF3B-1 ARABIDOPSIS THALIANA TRANSLATION INITIAT |
| AT5G49460.1 | 2,01879836 | 0,000251453 | ACLB-2 ATP citrate lyase subunit B 2 chr5:20055048-20058195 FORWARD LENGTH=608 |
| AT4G38510.1 | 0,749700179 | 0,000254534 | AtVAB2_ VAB2 V-ATPase B subunit 2 chr4:18011155-18014789 REVERSE LENGTH=487 |
| AT5G19510.1 | 0,532564173 | 0,000254945 | no symbol available no full name available chr5:6581854-6583137 REVERSE LENGTH=224 |
| AT3G44890.1 | 0,640777299 | 0,00025707 | RPL9 ribosomal protein L9 chr3:16386505-16387963 FORWARD LENGTH=197 |
| AT3G09840.1 | 1,315298279 | 0,000263125 | ATCDC48_ CDC48_ AtCDC48A_ CDC48A cell division cycle 48 chr3:3019494-3022832 FORWARD LENG |

| | | | |
|---|---|---|---|
| AT2G16600.1 | 0,569385178 | 0,000269051 | AtCYP19-1_ ROC3_ CYP19 rotamase CYP 3_ cyclophilin 19 chr2:7200862-7201383 FORWARD LENGTH= |
| AT1G07890.1 | 1,155759504 | 0,000269651 | MEE6_ ATAPX01_ ATAPX1_ CS1_ APX1 ascorbate peroxidase 1_ maternal effect embryo arrest 6 chr1:243 |
| AT5G54810.1 | 0,730809585 | 0,000270988 | TRP2_ ATTSB1_ TSB1_ TRPB tryptophan synthase beta-subunit 1_ TRYPTOPHAN BIOSYNTHESIS B_ T |
| AT5G10920.1 | 1,727067771 | 0,000280394 | no symbol available no full name available chr5:3441805-3443892 FORWARD LENGTH=517 |
| AT5G59880.2 | 0,609915997 | 0,000285798 | ADF3 actin depolymerizing factor 3 chr5:24120382-24121628 FORWARD LENGTH=124 |
| AT5G67030.2 | 1,377089749 | 0,000293981 | LOS6_ NPQ2_ ZEP_ ABA1_ ATABA1_ ATZEP_ IBS3 ABA DEFICIENT 1_ IMPAIRED IN BABA-INDUC |
| AT1G50200.1 | 1,296450122 | 0,000295803 | ALATS_ ACD Alanyl-tRNA synthetase chr1:18591429-18598311 REVERSE LENGTH=1003 |
| AT5G05600.1 | 0,534108244 | 0,000296409 | JAO2_ JOX2 JASMONATE-INDUCED OXYGENASE2_ Jasmonic Acid Oxidase 2 chr5:1672266-1674602 F |
| AT3G52500.1 | 0,750605949 | 0,000300451 | no symbol available no full name available chr3:19465644-19467053 REVERSE LENGTH=469 |
| AT2G26670.1 | 1,471461633 | 0,00030109 | ATHO1_ HY1_ HY6_ HO1_ GUN2_ TED4 REVERSAL OF THE DET PHENOTYPE 4_ ARABIDOPSIS T |
| AT5G08570.1 | 1,970646547 | 0,000304114 | no symbol available no full name available chr5:2778433-2780300 FORWARD LENGTH=510 |
| AT1G79550.1 | 2,00902792 | 0,000308745 | PGKc_ PGK_ PGK3 phosphoglycerate kinase_ phosphoglycerate kinase 3 chr1:29924347-29926295 REVERS |
| AT1G02920.1 | 0,602608802 | 0,0003169 | ATGSTF8_ GSTF7_ ATGSTF7_ ATGST11_ GST11 GLUTATHIONE S-TRANSFERASE 11_ glutathione S- |
| AT5G23140.1 | 0,715200284 | 0,000320853 | CLPP2_ NCLPP7 nuclear-encoded CLP protease P7 chr5:7783811-7784826 FORWARD LENGTH=241 |
| AT5G66530.1 | 0,61811063 | 0,000327901 | no symbol available no full name available chr5:26553821-26555575 REVERSE LENGTH=307 |
| AT3G04770.1 | 1,524581241 | 0,00033359 | RPSAb 40s ribosomal protein SA B chr3:1309465-1310846 REVERSE LENGTH=332 |
| AT5G61790.1 | 1,360766313 | 0,00033642 | CNX1_ ATCNX1 calnexin 1 chr5:24827394-24829642 REVERSE LENGTH=530 |
| AT3G27690.1 | 1,519622293 | 0,00033654 | LHCB2.4_ DEG13_ LHCB2_ LHCB2.3 LIGHT-HARVESTING CHLOROPHYLL B-BINDING 2_ photosyst |
| AT3G13920.4 | 1,684144035 | 0,000338268 | TIF4A1_ EIF4A1_ RH4 eukaryotic translation initiation factor 4A1 chr3:4592635-4594128 REVERSE LENG |
| AT1G15690.1 | 1,709307974 | 0,000339603 | AtVHP1;1_ AtAVP1_ ATAVP3_ AVP-3_ AVP1_ FUGU5_ VHP1 ARABIDOPSIS THALIANA V-PPASE 3_ |
| AT3G58990.1 | 2,760152144 | 0,000342342 | IPMI SSU3_ IPMI1 isopropylmalate isomerase 1 chr3:21797524-21798285 REVERSE LENGTH=253 |
| AT1G75780.1 | 0,566068052 | 0,000343099 | TUB1 tubulin beta-1 chain chr1:28451378-28453602 REVERSE LENGTH=447 |
| AT2G28190.1 | 0,596763993 | 0,00034339 | CSD2_ CZSOD2_ SOD2_ AtSOD2 COPPER/ZINC SUPEROXIDE DISMUTASE 2_ superoxide dismutase 2_ |
| AT5G07350.1 | 1,63574407 | 0,000343619 | Tudor1_ TSN1_ AtTudor1 TUDOR-SN protein 1_ Arabidopsis thaliana TUDOR-SN protein 1 chr5:2320344-2 |
| AT1G45145.1 | 0,156633486 | 0,000364163 | TRX-h5_ ATTRX5_ LIV1_ ATH5_ TRX5 thioredoxin H-type 5_ THIOREDOXIN H-TYPE 5_ LOCUS OF I |
| AT1G77940.1 | 0,470526796 | 0,000372489 | RPL30B chr1:29304116-29305288 REVERSE LENGTH=112 |
| AT5G54190.1 | 0,443244767 | 0,000387883 | PORA protochlorophyllide oxidoreductase A chr5:21991183-21992773 REVERSE LENGTH=405 |
| AT3G26650.1 | 1,196513422 | 0,000392067 | GAPA_ GAPA1_ GAPA-1 glyceraldehyde 3-phosphate dehydrogenase A subunit_ GLYCERALDEHYDE 3-P |
| AT3G16520.3 | 1,71712234 | 0,00039858 | UGT88A1 UDP-glucosyl transferase 88A1 chr3:5619355-5620833 REVERSE LENGTH=462 |
| AT1G65350.1 | 1,821722089 | 0,000405248 | UBQ13 ubiquitin 13 chr1:24272518-24277275 REVERSE LENGTH=319 |
| AT4G17520.1 | 0,562341814 | 0,000408304 | HLN HYALURONAN/mRNA BINDING FAMILY PROTEIN chr4:9771496-9773313 FORWARD LENGTH |
| AT1G54220.1 | 0,439393677 | 0,000417181 | mtE2-3 mitochondrial pyruvate dehydrogenase subunit 2-3 chr1:20246460-20250208 REVERSE LENGTH=53 |
| AT1G42970.1 | 1,10545736 | 0,000420375 | GAPB glyceraldehyde-3-phosphate dehydrogenase B subunit chr1:16127552-16129584 FORWARD LENGTH |
| AT1G22450.1 | 0,438162745 | 0,000435861 | ATCOX6B2_ COX6B cytochrome C oxidase 6B_ CYTOCHROME C OXIDASE 6B2 chr1:7925447-7926918 |
| AT1G79930.1 | 1,592920638 | 0,000441647 | HSP91_ AtHsp70-14 heat shock protein 91 chr1:30063781-30067067 REVERSE LENGTH=831 |
| AT3G07390.1 | 0,677600218 | 0,000455206 | AIR12 Auxin-Induced in Root cultures 12 chr3:2365452-2366273 FORWARD LENGTH=273 |
| AT2G29450.1 | 1,819543195 | 0,000456872 | AT103-1A_ ATGSTU1_ ATGSTU5_ GSTU5 glutathione S-transferase tau 5_ ARABIDOPSIS THALIANA G |
| AT1G55490.1 | 1,18643695 | 0,000459104 | Cpn60beta1_ LEN1_ CPNB1_ CPN60B chaperonin-60beta1_ LESION INITIATION 1_ chaperonin 60 beta c |
| AT5G11450.1 | 0,303753249 | 0,00046166 | PPD5 PsbP domain protein 5 chr5:3654475-3656357 FORWARD LENGTH=297 |

| | | | |
|---|---|---|---|
| AT5G55220.1 | 0,641952285 | 0,000463622 | HP65b_ TIG1 chr5:22397677-22400678 FORWARD LENGTH=547 |
| AT3G13870.1 | 2,218413911 | 0,000472463 | GOM8_ RHD3 GOLGI MUTANT 8_ ROOT HAIR DEFECTIVE 3 chr3:4565762-4571109 REVERSE LENG |
| AT4G37925.1 | 0,618812143 | 0,000482311 | NDH-M_ NdhM subunit NDH-M of NAD(P)H:plastoquinone dehydrogenase complex_ NADH dehydrogenase |
| AT3G03710.1 | 1,395801831 | 0,000483461 | PNP_ RIF10_ PDE326 PIGMENT DEFECTIVE 326_ POLYNUCLEOTIDE PHOSPHORYLASE_ resistant t |
| AT4G33220.2 | 2,67211412 | 0,000486451 | ATPME44_ PME44 pectin methylesterase 44_ A. THALIANA PECTIN METHYLESTERASE 44 chr4:16024 |
| ATCG00660.1 | 0,269636213 | 0,0004894 | RPL20 ribosomal protein L20 chrc:68512-68865 REVERSE LENGTH=117 |
| AT1G67430.2 | 0,159562682 | 0,000495869 | no symbol available no full name available chr1:25262209-25263627 FORWARD LENGTH=131 |
| AT2G21530.1 | 0,422989806 | 0,000499981 | no symbol available no full name available chr2:9219372-9220464 FORWARD LENGTH=209 |
| AT4G28080.1 | 1,872610931 | 0,000500045 | REC2 REDUCED CHLOROPLAST COVERAGE 2 chr4:13948993-13957840 REVERSE LENGTH=1819 |
| AT2G40290.1 | 0,830689532 | 0,000537401 | no symbol available no full name available chr2:16829030-16830889 REVERSE LENGTH=344 |
| AT5G35630.1 | 0,896927613 | 0,000574775 | GLN2_ GS2_ ATGSL1 GLUTAMINE SYNTHETASE 2_ glutamine synthetase 2_ GLUTAMINE SYNTHET |
| AT1G76080.1 | 0,474309473 | 0,000579893 | CDSP32_ TRXL1_ ATCDSP32 ARABIDOPSIS THALIANA CHLOROPLASTIC DROUGHT-INDUCED ST |
| AT2G06050.1 | 0,583921436 | 0,000594248 | OPR3_ DDE1_ AtOPR3 DELAYED DEHISCENCE 1_ oxophytodienoate-reductase 3 chr2:2359240-2361971 |
| AT5G26000.1 | 0,827117377 | 0,000595661 | TGG1_ AtTGG1_ BGLU38 thioglucoside glucohydrolase 1_ BETA GLUCOSIDASE 38 chr5:9079678-90823 |
| AT1G11580.1 | 0,639448391 | 0,000600295 | PME18_ PMEPCRA_ ATPMEPCRA methylesterase PCR A_ PECTIN METHYLESTERASE 18 chr1:388873 |
| AT2G33040.1 | 1,405691769 | 0,000627876 | ATP3 gamma subunit of Mt ATP synthase chr2:14018978-14021047 REVERSE LENGTH=325 |
| AT2G42590.1 | 1,379143282 | 0,000630113 | GF14 MU_ GRF9_ GRF14 general regulatory factor 9 chr2:17732118-17733775 REVERSE LENGTH=263 |
| AT1G19570.1 | 0,814370108 | 0,000636568 | ATDHAR1_ DHAR1_ DHAR5 DEHYDROASCORBATE REDUCTASE 5_ dehydroascorbate reductase chr1 |
| AT5G13850.1 | 0,576238952 | 0,000639425 | NACA3 nascent polypeptide-associated complex subunit alpha-like protein 3 chr5:4471361-4472676 FORWA |
| AT5G52650.1 | 1,63050682 | 0,000653419 | no symbol available no full name available chr5:21355781-21357003 REVERSE LENGTH=179 |
| AT3G63170.1 | 0,633601901 | 0,000655117 | FAP1_ AtFAP1 fatty-acid-binding protein 1 chr3:23334675-23335993 FORWARD LENGTH=279 |
| AT3G18080.1 | 1,538061875 | 0,000663877 | BGLU44 B-S glucosidase 44 chr3:6191586-6194124 FORWARD LENGTH=512 |
| AT2G36460.1 | 1,400028103 | 0,000668103 | FBA6 fructose-bisphosphate aldolase 6 chr2:15296929-15298387 REVERSE LENGTH=358 |
| AT5G58710.1 | 0,709585778 | 0,00066943 | ROC7 rotamase CYP 7 chr5:23717840-23719495 FORWARD LENGTH=204 |
| AT3G03960.1 | 1,652215979 | 0,0006699 | CCT8 Chaperonin containing T-complex polypeptide-1 subunit 8 chr3:1024432-1027604 FORWARD LENGT |
| AT2G21410.1 | 4,39496295 | 0,000681214 | VHA-A2 vacuolar proton ATPase A2 chr2:9162703-9168141 FORWARD LENGTH=821 |
| AT1G05010.1 | 0,411974583 | 0,000686836 | EFE_ ACO4_ EAT1 ethylene forming enzyme_ ethylene-forming enzyme chr1:1431419-1432695 REVERSE L |
| AT4G37800.1 | 0,760380168 | 0,000696261 | XTH7 xyloglucan endotransglucosylase/hydrolase 7 chr4:17775703-17777372 REVERSE LENGTH=293 |
| AT3G18490.1 | 0,58840025 | 0,000703808 | ASPG1 ASPARTIC PROTEASE IN GUARD CELL 1 chr3:6349090-6350592 REVERSE LENGTH=500 |
| AT3G23990.1 | 1,90032189 | 0,000706218 | HSP60_ HSP60-3B heat shock protein 60_ HEAT SHOCK PROTEIN 60-3B chr3:8669013-8672278 FORWA |
| AT3G15730.1 | 1,967318467 | 0,000707328 | PLD_ PLDALPHA1 phospholipase D alpha 1 chr3:5330835-5333474 FORWARD LENGTH=810 |
| AT1G22300.1 | 1,303634544 | 0,000709301 | 14-3-3EPSILON_ GRF10_ GF14 EPSILON general regulatory factor 10_ 14-3-3 PROTEIN G-BOX FACTOR |
| AT2G28950.1 | 0,654523654 | 0,00072149 | ATEXP6_ ATHEXP ALPHA 1.8_ ATEXPA6_ EXPA6 expansin A6_ ARABIDOPSIS THALIANA TEXPAN |
| AT1G20450.1 | 1,38784931 | 0,000725188 | LTI45_ ERD10_ LTI29 EARLY RESPONSIVE TO DEHYDRATION 10_ LOW TEMPERATURE INDUCE |
| AT5G62350.1 | 0,168883826 | 0,000737448 | no symbol available no full name available chr5:25037504-25038112 FORWARD LENGTH=202 |
| AT5G37640.1 | 1,846139631 | 0,000762799 | UBQ9 ubiquitin 9 chr5:14952782-14953750 REVERSE LENGTH=322 |
| AT5G58070.1 | 2,416050091 | 0,000767844 | TIL_ ATTIL TEMPERATURE-INDUCED LIPOCALIN_ temperature-induced lipocalin chr5:23500512-2350 |
| AT2G25140.1 | 2,630785585 | 0,000779481 | HSP98.7_ CLPB-M_ CLPB4 HEAT SHOCK PROTEIN 98.7_ CASEIN LYTIC PROTEINASE B-M_ casein |
| AT1G13440.2 | 1,886419759 | 0,000807512 | GAPC2_ GAPC-2 glyceraldehyde-3-phosphate dehydrogenase C2_ GLYCERALDEHYDE-3-PHOSPHATE D |

| | | | |
|---|---|---|---|
| AT3G16420.1 | 0,631633015 | 0,000818545 | PBPI_ JAL30_ PBP1 PYK10-binding protein 1_ JACALIN-RELATED LECTIN 30 chr3:5579560-5580674 F[illegible] |
| AT1G12240.1 | 0,325150092 | 0,000820194 | AtVI2_ VAC-INV_ ATBETAFRUCT4_ VI2_ FRUCT4_ AtFRUCT4_ VIN2 VACUOLAR INVERTASE_ va[illegible] |
| AT5G57290.2 | 0,782096242 | 0,000830806 | P3B_ AtP3B ribosomal P3 protein B chr5:23207089-23207835 REVERSE LENGTH=89 |
| AT5G08380.1 | 0,754193765 | 0,000834711 | AtAGAL1_ AGAL1 alpha-galactosidase 1 chr5:2694851-2697616 REVERSE LENGTH=410 |
| AT1G72730.1 | 1,37719872 | 0,000856136 | no symbol available no full name available chr1:27378040-27379593 REVERSE LENGTH=414 |
| AT4G01800.1 | 1,729883722 | 0,000868495 | SECA1_ AGY1_ AtcpSecA Arabidopsis thaliana chloroplast SecA_ Albino or Glassy Yellow 1 chr4:770926-7[illegible] |
| AT2G40010.1 | 0,565203877 | 0,000869317 | no symbol available no full name available chr2:16708578-16710448 REVERSE LENGTH=317 |
| AT3G23570.1 | 0,75835641 | 0,000905341 | no symbol available no full name available chr3:8458052-8459608 REVERSE LENGTH=239 |
| AT4G33090.1 | 1,452076579 | 0,000920327 | APM1_ ATAPM1 AMINOPEPTIDASE M1_ aminopeptidase M1 chr4:15965915-15970418 REVERSE LEN[illegible] |
| AT1G47128.1 | 0,607875622 | 0,000926914 | RD21_ RD21A responsive to dehydration 21A_ responsive to dehydration 21 chr1:17283139-17285609 REVE[illegible] |
| AT3G09640.1 | 4,386706843 | 0,000930041 | AtAPX2_ APX1B_ APX2 ASCORBATE PEROXIDASE 1B_ ascorbate peroxidase 2 chr3:2956301-2958163 [illegible] |
| AT1G30360.1 | 1,379908704 | 0,000948823 | ERD4_ OSCA3.1 early-responsive to dehydration 4 chr1:10715892-10718799 FORWARD LENGTH=724 |
| AT3G16470.1 | 0,749923772 | 0,000950998 | JAL35_ AtJAC1_ JR1 jacalin-related lectin 35_ JACALIN-LECTIN LIKE 1_ JASMONATE RESPONSIVE 1 |
| ATCG00780.1 | 0,726506231 | 0,000977946 | RPL14 ribosomal protein L14 chrc:80696-81064 REVERSE LENGTH=122 |
| AT4G13940.1 | 1,371808054 | 0,000986364 | MEE58_ SAHH1_ EMB1395_ HOG1_ SAH1_ ATSAHH1 EMBRYO DEFECTIVE 1395_ HOMOLOGY-DE[illegible] |
| AT1G74470.1 | 0,750647417 | 0,001023142 | no symbol available no full name available chr1:27991248-27992845 FORWARD LENGTH=467 |
| AT1G65260.1 | 2,595970137 | 0,001033141 | VIPP1_ PTAC4_ IM30 VESICLE-INDUCING PROTEIN IN PLASTIDS 1_ plastid transcriptionally active 4 [illegible] |
| AT1G18080.1 | 1,548160961 | 0,001066085 | RACK1A_AT_ AtRACK1_ SAC53_ ATARCA_ RACK1A RECEPTOR FOR ACTIVATED C KINASE 1 A_ |
| AT3G56070.1 | 0,474395624 | 0,001071457 | ROC2 rotamase cyclophilin 2 chr3:20806987-20807517 REVERSE LENGTH=176 |
| AT2G26080.1 | 0,836359977 | 0,001072159 | GLDP2_ AtGLDP2 glycine decarboxylase P-protein 2 chr2:11109330-11113786 REVERSE LENGTH=1044 |
| AT2G36250.4 | 0,776568455 | 0,001085694 | FTSZ2-1_ ATFTSZ2-1 chr2:15197661-15199689 REVERSE LENGTH=397 |
| AT5G20010.1 | 1,298167034 | 0,001095773 | RAN-1_ ATRAN1_ RAN1 RAS-RELATED NUCLEAR PROTEIN_ ARABIDOPSIS THALIANA RAS-REL[illegible] |
| AT2G35840.1 | 2,283082363 | 0,001098671 | no symbol available no full name available chr2:15053952-15055776 FORWARD LENGTH=422 |
| AT1G36240.1 | 0,324373627 | 0,001105588 | RPL30A chr1:13614890-13616233 FORWARD LENGTH=112 |
| AT2G14720.1 | 0,74876572 | 0,001106303 | MTV2_ BP80-2;1_ MTV4_ VSR4_ VSR2;1 vacuolar sorting receptor 4_ VACUOLAR SORTING RECEPTO[illegible] |
| AT1G45000.1 | 1,397116022 | 0,001114778 | RPT4b chr1:17009220-17011607 FORWARD LENGTH=399 |
| AT4G29410.1 | 0,527869873 | 0,001121262 | no symbol available no full name available chr4:14468439-14469964 REVERSE LENGTH=143 |
| AT3G18190.1 | 1,668045765 | 0,001127795 | CCT4 Chaperonin containing T-complex polypeptide-1 subunit 4 chr3:6232226-6233836 FORWARD LENGT[illegible] |
| AT5G27770.1 | 0,732459279 | 0,001132453 | no symbol available no full name available chr5:9836166-9837113 FORWARD LENGTH=124 |
| AT3G63410.1 | 0,704862495 | 0,001157413 | E37_ VTE3_ IEP37_ APG1 INNER ENVELOPE PROTEIN 37_ VITAMIN E DEFECTIVE 3_ ALBINO OR |
| AT2G36580.1 | 1,447853576 | 0,001164815 | no symbol available no full name available chr2:15339253-15342781 FORWARD LENGTH=527 |
| AT3G25770.1 | 0,727195685 | 0,001188676 | AOC2 allene oxide cyclase 2 chr3:9406975-9407839 FORWARD LENGTH=253 |
| AT3G02730.1 | 0,727032767 | 0,001209973 | TRXF1_ ATF1 thioredoxin F-type 1 chr3:588570-589591 REVERSE LENGTH=178 |
| AT5G67360.1 | 0,592118731 | 0,001210105 | ARA12_ SBT1.7 Subtilisin-like Serine protease 1.7 chr5:26872192-26874465 REVERSE LENGTH=757 |
| AT5G20890.1 | 1,665669644 | 0,001212136 | CCT2 Chaperonin containing T-complex polypeptide-1 subunit 2 chr5:7087020-7089906 REVERSE LENGTH |
| AT1G77490.2 | 0,710622363 | 0,00121479 | TAPX thylakoidal ascorbate peroxidase chr1:29117688-29120649 FORWARD LENGTH=421 |
| AT4G39260.1 | 0,685742175 | 0,001216047 | RBGA6_ CCR1_ ATGRP8_ GR-RBP8_ GRP8 "cold_ circadian rhythm_ and RNA binding 1"_ glycine-rich R[illegible] |
| AT1G03600.1 | 0,219865697 | 0,001218251 | PSB27 chr1:898916-899440 FORWARD LENGTH=174 |
| AT2G43560.1 | 0,732581052 | 0,001219385 | no symbol available no full name available chr2:18073995-18075385 REVERSE LENGTH=223 |

| | | | |
|---|---|---|---|
| AT3G06860.1 | 1,37658123 | 0,001228432 | ATMFP2_ MFP2 MULTIFUNCTIONAL PROTEIN 2_ multifunctional protein 2 chr3:2161926-2166009 FOR |
| AT3G02880.1 | 1,62309008 | 0,001274839 | KIN7 Kinase 7 chr3:634819-636982 FORWARD LENGTH=627 |
| AT2G47390.1 | 0,766685833 | 0,001341899 | CGEP chloroplast glutamyl peptidase chr2:19442278-19446253 REVERSE LENGTH=961 |
| AT3G10350.1 | 0,710405461 | 0,001342968 | AtGET3b_ GET3b Guided Entry of Tail-anchored proteins 3b chr3:3208310-3210678 FORWARD LENGTH= |
| AT5G45390.1 | 1,309902583 | 0,001354668 | CLPP4_ NCLPP4 NUCLEAR-ENCODED CLP PROTEASE P4_ CLP protease P4 chr5:18396351-18397586 I |
| AT3G52580.1 | 0,814496575 | 0,0013594 | no symbol available no full name available chr3:19503324-19504701 FORWARD LENGTH=150 |
| AT2G43100.1 | 0,496822515 | 0,001360695 | IPMI2_ IPMI SSU2_ ATLEUD1 isopropylmalate isomerase 2_ isopropylmalate isomerase small sub-unit 2 chr |
| AT5G62690.1 | 1,264665436 | 0,001377244 | TUB2 tubulin beta chain 2 chr5:25181560-25183501 FORWARD LENGTH=450 |
| AT5G25880.3 | 0,617071588 | 0,00139435 | ATNADP-ME3_ NADP-ME3 NADP-malic enzyme 3_ Arabidopsis thaliana NADP-malic enzyme 3 chr5:9024 |
| AT5G09650.1 | 0,535559447 | 0,001398465 | PPa6_ AtPPa6 pyrophosphorylase 6 chr5:2991331-2993117 REVERSE LENGTH=300 |
| AT1G04530.1 | 1,665623177 | 0,00141188 | TPR4 tetratricopeptide repeat 4 chr1:1234456-1235895 REVERSE LENGTH=310 |
| AT4G32470.1 | 0,607145011 | 0,001414404 | no symbol available no full name available chr4:15669641-15671095 REVERSE LENGTH=122 |
| AT3G11830.1 | 1,589409618 | 0,001438234 | CCT7 Chaperonin containing T-complex polypeptide-1 subunit 7 chr3:3732734-3736156 FORWARD LENGT |
| AT2G09990.1 | 0,448224658 | 0,001441654 | no symbol available no full name available chr2:3781442-3781882 FORWARD LENGTH=146 |
| AT4G09720.1 | 1,669237946 | 0,001451426 | RABG3A_ ATRABG3A RAB GTPase homolog G3A chr4:6133101-6134959 FORWARD LENGTH=206 |
| AT1G09100.1 | 1,183612486 | 0,001453116 | RPT5B 26S proteasome AAA-ATPase subunit RPT5B chr1:2936675-2939258 REVERSE LENGTH=423 |
| AT2G22780.1 | 1,598401685 | 0,001457761 | PMDH1 peroxisomal NAD-malate dehydrogenase 1 chr2:9689995-9691923 REVERSE LENGTH=354 |
| ATCG00810.1 | 0,390120854 | 0,001459129 | RPL22 ribosomal protein L22 chrc:83467-83949 REVERSE LENGTH=160 |
| ATCG00120.1 | 1,19824517 | 0,001463197 | ATPA ATP synthase subunit alpha chrc:9938-11461 REVERSE LENGTH=507 |
| AT2G21870.1 | 0,659961265 | 0,001463984 | MGP1_ PHI1 PHOSPHITE-INSENSITIVE 1_ MALE GAMETOPHYTE DEFECTIVE 1 chr2:9320456-9322 |
| AT2G42210.1 | 0,864369349 | 0,001486121 | OEP16-3_ ATOEP16-3 chr2:17590642-17591591 FORWARD LENGTH=159 |
| AT3G25860.1 | 1,292877768 | 0,001497781 | LTA2_ PLE2 PLASTID E2 SUBUNIT OF PYRUVATE DECARBOXYLASE chr3:9460632-9462585 FORW |
| AT3G62030.1 | 0,850605859 | 0,001507658 | ROC4_ CYP20-3 rotamase CYP 4_ cyclophilin 20-3 chr3:22973708-22975139 FORWARD LENGTH=260 |
| AT5G40770.1 | 1,441663021 | 0,001509539 | ATPHB3_ PHB3_ EER3 prohibitin 3 chr5:16315589-16316621 REVERSE LENGTH=277 |
| AT5G55730.1 | 1,209666644 | 0,001546132 | FLA1 FASCICLIN-like arabinogalactan 1 chr5:22558375-22560392 REVERSE LENGTH=424 |
| AT1G31230.1 | 1,70022618 | 0,001552602 | AK-HSDH I_ AK-HSDH ASPARTATE KINASE-HOMOSERINE DEHYDROGENASE_ aspartate kinase-ho |
| AT3G15356.1 | 0,665100136 | 0,001562705 | no symbol available no full name available chr3:5174603-5175418 REVERSE LENGTH=271 |
| AT5G16710.1 | 0,911556668 | 0,00156763 | DHAR3 dehydroascorbate reductase 1 chr5:5483312-5484926 FORWARD LENGTH=258 |
| AT4G00660.1 | 1,989155838 | 0,001587539 | RH8_ ATRH8 RNAhelicase-like 8 chr4:274638-277438 FORWARD LENGTH=505 |
| AT5G42980.1 | 1,331253891 | 0,001593971 | ATH3_ TRX3_ TRXH3_ ATTRX3_ ATTRXH3 THIOREDOXIN H3_ thioredoxin 3_ thioredoxin H-type 3 ch |
| AT3G04720.1 | 0,762720038 | 0,001620256 | HEL_ PR-4_ AtPR4_ PR4 HEVEIN-LIKE_ pathogenesis-related 4 chr3:1285691-1286531 REVERSE LENGT |
| AT4G21990.1 | 1,449994032 | 0,001627866 | APR3_ PRH26_ PRH-26_ ATAPR3 PAPS REDUCTASE HOMOLOG 26_ APS reductase 3 chr4:11657284-1 |
| AT1G56410.1 | 0,642565296 | 0,001634229 | HSP70T-1_ ERD2 HEAT SHOCK PROTEIN 70T-1_ EARLY-RESPONSIVE TO DEHYDRATION 2 chr1:2 |
| AT4G31180.1 | 1,893866418 | 0,001658076 | IBI1 impaired in BABA-induced disease immunity 1 chr4:15156696-15159362 FORWARD LENGTH=558 |
| AT3G08530.1 | 1,788043804 | 0,001663262 | AtCHC2_ CHC2 clathrin heavy chain 2 chr3:2587171-2595411 REVERSE LENGTH=1703 |
| AT5G16050.1 | 1,875551163 | 0,00167638 | GRF5_ GF14 UPSILON general regulatory factor 5 chr5:5244008-5245402 REVERSE LENGTH=268 |
| AT3G59970.1 | 1,782198827 | 0,001713833 | MTHFR1 methylenetetrahydrofolate reductase 1 chr3:22151303-22153412 FORWARD LENGTH=421 |
| AT2G27860.1 | 1,279955093 | 0,001714621 | AXS1 UDP-D-apiose/UDP-D-xylose synthase 1 chr2:11864684-11866843 REVERSE LENGTH=389 |
| AT4G37910.1 | 1,402149332 | 0,001717983 | mtHsc70-1 mitochondrial heat shock protein 70-1 chr4:17825368-17828099 REVERSE LENGTH=682 |

| | | | |
|---|---|---|---|
| AT2G29440.1 | 2,083121005 | 0,001722379 | GST24_ ATGSTU6_ GSTU6 glutathione S-transferase tau 6_ GLUTATHIONE S-TRANSFERASE 24 chr2:1[illegible] |
| AT3G16390.1 | 0,75509489 | 0,001726428 | NSP3 nitrile specifier protein 3 chr3:5562602-5564356 FORWARD LENGTH=467 |
| AT2G39470.2 | 0,696380997 | 0,001726926 | PnsL1_ PPL2 Photosynthetic NDH subcomplex L 1_ PsbP-like protein 2 chr2:16476335-16477653 FORWAR[illegible] |
| AT1G48860.1 | 1,748629796 | 0,001727293 | EPSPS 5-enolpyruvylshikimate-3-phosphate synthase chr1:18068892-18071331 REVERSE LENGTH=521 |
| AT5G19820.1 | 1,908629945 | 0,001736049 | IMB3_ KETCH1_ EMB2734 EMBRYO DEFECTIVE 2734_ (karyopherin enabling the transport of the cytopl[illegible] |
| AT1G09210.1 | 0,813188122 | 0,001757999 | CRT2_ AtCRT1b_ CRT1b calreticulin 1b_ CALRETICULIN 2 chr1:2973217-2976655 REVERSE LENGTH[illegible] |
| AT1G60950.1 | 0,42807449 | 0,001765357 | ATFD2_ FD2_ FED A FERREDOXIN 2 chr1:22444565-22445011 FORWARD LENGTH=148 |
| AT4G24280.1 | 1,324782573 | 0,001778222 | cpHsc70-1 chloroplast heat shock protein 70-1 chr4:12590094-12593437 FORWARD LENGTH=718 |
| AT2G21390.1 | 1,372735854 | 0,001809934 | no symbol available no full name available chr2:9152428-9156577 FORWARD LENGTH=1218 |
| AT1G78630.1 | 0,089644198 | 0,001820385 | emb1473 embryo defective 1473 chr1:29575997-29577406 FORWARD LENGTH=241 |
| AT1G70310.1 | 1,29798254 | 0,001822639 | SPDS2 spermidine synthase 2 chr1:26485497-26487352 REVERSE LENGTH=340 |
| AT3G02520.1 | 1,488099164 | 0,001826793 | GRF7_ GF14 NU general regulatory factor 7 chr3:526800-527915 REVERSE LENGTH=265 |
| AT3G60820.1 | 1,274670594 | 0,001833778 | PBF1 chr3:22472038-22473809 REVERSE LENGTH=223 |
| AT5G17330.1 | 0,764864715 | 0,001836139 | GAD_ AtGAD1_ GAD1 GLUTAMATE DECARBOXYLASE 1_ glutamate decarboxylase chr5:5711141-571[illegible] |
| AT4G35250.1 | 1,134028962 | 0,001871801 | HCF244 high chlorophyll fluorescence phenotype 244 chr4:16771401-16773269 REVERSE LENGTH=395 |
| AT1G35580.3 | 1,403989015 | 0,001877918 | CINV1_ A/N-InvG_ NIN2 cytosolic invertase 1_ alkaline/neutral invertase G_ neutral invertase 2 chr1:131231[illegible] |
| AT4G14160.3 | 1,625440877 | 0,001914328 | AtSEC23F chr4:8167574-8172266 FORWARD LENGTH=620 |
| AT5G13490.1 | 1,616498753 | 0,00193008 | AAC2 ADP/ATP carrier 2 chr5:4336034-4337379 FORWARD LENGTH=385 |
| AT4G39890.1 | 2,03554351 | 0,001949085 | AtRABH1c_ RABH1c RAB GTPase homolog H1C chr4:18506112-18507459 FORWARD LENGTH=214 |
| AT2G26540.3 | 0,80541028 | 0,00195139 | DUF3_ ATDUF3_ HEMD_ ATUROS_ UROS ARABIDOPSIS THALIANA UROPORPHYRINOGEN III SY[illegible] |
| AT1G78900.1 | 1,347042798 | 0,001980542 | VHA-A vacuolar ATP synthase subunit A chr1:29660463-29664575 FORWARD LENGTH=623 |
| AT3G12290.1 | 1,561039064 | 0,00198067 | MTHFD1 methylenetetrahydrofolate dehydrogenase/methenyltetrahydrofolate cyclohydrolase chr3:3919591-39[illegible] |
| AT1G02500.1 | 0,789384338 | 0,001995172 | METK1_ SAM-1_ AtSAM1_ SAM1_ MAT1 S-adenosylmethionine synthetase 1_ S-ADENOSYLMETHIONI[illegible] |
| AT5G61780.1 | 1,635569623 | 0,002000212 | Tudor2_ AtTudor2_ TSN2 Arabidopsis thaliana TUDOR-SN protein 2_ TUDOR-SN protein 2 chr5:24822012-[illegible] |
| AT3G52730.1 | 1,278501414 | 0,002004258 | no symbol available no full name available chr3:19543146-19544167 REVERSE LENGTH=72 |
| AT1G51760.1 | 0,542248766 | 0,002027495 | IAR3_ JR3 IAA-ALANINE RESISTANT 3_ JASMONIC ACID RESPONSIVE 3 chr1:19199562-19201424 F[illegible] |
| AT5G11670.1 | 1,226822076 | 0,002029096 | NADP-ME2_ ATNADP-ME2 NADP-malic enzyme 2_ Arabidopsis thaliana NADP-malic enzyme 2 chr5:3754[illegible] |
| AT3G13750.1 | 0,47858426 | 0,002044501 | BGAL1 beta galactosidase 1_ beta-galactosidase 1 chr3:4511192-4515756 FORWARD LENGTH=847 |
| AT5G44340.1 | 1,647933196 | 0,002057884 | TUB4 tubulin beta chain 4 chr5:17859442-17860994 REVERSE LENGTH=444 |
| AT2G24270.1 | 1,379304933 | 0,002060461 | ALDH11A3 aldehyde dehydrogenase 11A3 chr2:10327325-10329601 REVERSE LENGTH=496 |
| AT1G04820.1 | 1,24023745 | 0,002093465 | TUA4_ TOR2 TORTIFOLIA 2_ tubulin alpha-4 chain chr1:1356421-1358266 REVERSE LENGTH=450 |
| AT4G18480.1 | 0,608615517 | 0,002107059 | CHL11_ CH-42_ LOST1_ CH42_ CHLI-1_ CHLI1 low temperature with open-stomata 1_ CHLORINA 42 chr[illegible] |
| AT2G19940.1 | 1,468872435 | 0,002107676 | no symbol available no full name available chr2:8613203-8615649 FORWARD LENGTH=389 |
| AT3G25920.1 | 0,64179757 | 0,002169436 | RPL15 ribosomal protein L15 chr3:9491268-9492558 REVERSE LENGTH=277 |
| AT3G18060.1 | 1,751313864 | 0,002175909 | no symbol available no full name available chr3:6183880-6186788 FORWARD LENGTH=609 |
| AT3G52880.1 | 1,38084604 | 0,002176631 | ATMDAR1_ MDAR1 monodehydroascorbate reductase 1 chr3:19601477-19604366 REVERSE LENGTH=43[illegible] |
| AT2G15620.1 | 0,775165703 | 0,002181879 | NIR_ ATHNIR_ NIR1 ARABIDOPSIS THALIANA NITRITE REDUCTASE_ nitrite reductase 1_ NITRITE [illegible] |
| AT3G46430.1 | 1,75453338 | 0,002216455 | AtMtATP6 chr3:17087687-17088497 FORWARD LENGTH=55 |
| AT1G02280.1 | 1,513956876 | 0,002220383 | TOC33_ PPI1_ ATTOC33 PLASTID PROTEIN IMPORT 1_ translocon at the outer envelope membrane of ch[illegible] |

| | | | |
|---|---|---|---|
| AT1G34430.1 | 1,431451187 | 0,002230262 | EMB3003 embryo defective 3003 chr1:12588027-12590084 REVERSE LENGTH=465 |
| AT1G76450.1 | 0,384234845 | 0,002237808 | no symbol available no full name available chr1:28684618-28686109 FORWARD LENGTH=247 |
| AT5G48180.1 | 1,472890958 | 0,002285678 | NSP5_ AtNSP5 nitrile specifier protein 5 chr5:19541283-19542358 REVERSE LENGTH=326 |
| AT4G12420.1 | 0,821778603 | 0,002313914 | SKU5 chr4:7349941-7352868 REVERSE LENGTH=587 |
| AT1G29900.1 | 1,619285981 | 0,002318508 | CARB_ VEN3 carbamoyl phosphate synthetase B_ VENOSA 3 chr1:10468164-10471976 FORWARD LENG |
| ATCG00820.1 | 0,523141371 | 0,002319683 | RPS19 ribosomal protein S19 chrc:84005-84283 REVERSE LENGTH=92 |
| AT1G09430.1 | 1,445460275 | 0,002361845 | ACLA-3 ATP-citrate lyase A-3 chr1:3042135-3044978 FORWARD LENGTH=424 |
| AT5G13120.1 | 0,723939658 | 0,002406001 | CYP20-2_ Pnsl5_ ATCYP20-2 cyclophilin 20-2_ Photosynthetic NDH subcomplex L 5_ ARABIDOPSIS THA |
| AT5G03350.1 | 0,66144245 | 0,002443413 | SAI-LLP1 SA-induced legume lectin-like protein 1 chr5:815804-816628 REVERSE LENGTH=274 |
| AT1G55210.1 | 0,8025715 | 0,002471888 | no symbol available no full name available chr1:20598057-20598620 REVERSE LENGTH=187 |
| AT2G01140.1 | 0,73818062 | 0,002481557 | FBA3_ AtFBA3_ PDE345 PIGMENT DEFECTIVE 345_ fructose-bisphosphate aldolase 3 chr2:95006-96491 |
| AT3G44300.1 | 1,592573695 | 0,002484538 | AtNIT2_ NIT2 nitrilase 2 chr3:15983351-15985172 FORWARD LENGTH=339 |
| AT2G05100.1 | 0,743617177 | 0,002493718 | LHCB2_ LHCB2.1 LIGHT-HARVESTING CHLOROPHYLL B-BINDING 2_ photosystem II light harvesting |
| AT1G12310.1 | 0,786515067 | 0,002521421 | no symbol available no full name available chr1:4187500-4187946 REVERSE LENGTH=148 |
| AT5G22800.1 | 1,321031558 | 0,002608276 | EMB263_ EMB1030_ EMB86 EMBRYO DEFECTIVE 263_ EMBRYO DEFECTIVE 1030_ EMBRYO DEF |
| AT2G04390.1 | 0,713040736 | 0,00260972 | dS17 chr2:1527911-1528336 FORWARD LENGTH=141 |
| AT3G06720.1 | 1,181911174 | 0,002611575 | IMPA1_ IMPA-1_ AT-IMP_ ATKAP ALPHA_ AIMP ALPHA IMPORTIN ALPHA_ IMPORTIN ALPHA IS |
| AT3G17390.1 | 0,814993073 | 0,002624622 | SAMS3_ AtSAMS3_ MAT4_ MTO3 METHIONINE ADENOSYLTRANSFERASE 4_ S-ADENOSYLMETH |
| AT1G28290.2 | 0,637196231 | 0,002683463 | AGP31 arabinogalactan protein 31 chr1:9889331-9890843 REVERSE LENGTH=315 |
| AT4G25130.1 | 0,727053717 | 0,002701854 | MSRA4_ PMSR4 peptide met sulfoxide reductase 4_ methionine sulfoxide reductase A4 chr4:12898802-12899 |
| AT1G24510.3 | 1,730308327 | 0,002736323 | CCT5 Chaperonin containing T-complex polypeptide-1 subunit 5 chr1:8685504-8687802 REVERSE LENGTH |
| AT4G39520.1 | 1,666176582 | 0,002802078 | Drg1-1 chr4:18371329-18374000 REVERSE LENGTH=369 |
| AT2G33070.1 | 0,565416372 | 0,002816415 | ATNSP2_ NSP2 NITRILE-SPECIFIER PROTEIN 2_ nitrile specifier protein 2 chr2:14029350-14030934 REV |
| AT1G64520.1 | 1,428724381 | 0,002825009 | RPN12a regulatory particle non-ATPase 12A chr1:23956459-23958120 FORWARD LENGTH=267 |
| AT3G63490.1 | 0,861249736 | 0,00284519 | EMB3126_ PRPL1 proline-rich protein-like 1_ plastid ribosomal protein L1_ EMBRYO DEFECTIVE 3126 ch |
| AT3G07110.1 | 0,239723561 | 0,002879061 | no symbol available no full name available chr3:2252092-2253332 FORWARD LENGTH=206 |
| AT2G38230.1 | 1,221611717 | 0,002892878 | ATPDX1.1_ PDX1.1 pyridoxine biosynthesis 1.1_ ARABIDOPSIS THALIANA PYRIDOXINE BIOSYNTHE |
| AT1G27450.1 | 1,323041374 | 0,002933539 | APT1_ ATAPT1 ARABIDOPSIS THALIANA ADENINE PHOSPHORIBOSYLTRANSFERASE 1_ adenine |
| AT3G12050.2 | 2,567242604 | 0,002934224 | no symbol available no full name available chr3:3839289-3841303 FORWARD LENGTH=321 |
| AT5G54900.1 | 0,699416942 | 0,002954726 | RBP45A_ ATRBP45A RNA-binding protein 45A chr5:22295412-22298126 FORWARD LENGTH=387 |
| AT5G13410.1 | 0,690905769 | 0,002959961 | no symbol available no full name available chr5:4299830-4301706 REVERSE LENGTH=256 |
| AT4G20890.1 | 0,623041708 | 0,002963302 | TUB9 tubulin beta-9 chain chr4:11182218-11183840 FORWARD LENGTH=444 |
| AT2G37660.1 | 0,830426258 | 0,002993708 | no symbol available no full name available chr2:15795481-15796977 REVERSE LENGTH=325 |
| AT1G74920.2 | 2,024396102 | 0,003030646 | ALDH10A8 aldehyde dehydrogenase 10A8 chr1:28139175-28142573 REVERSE LENGTH=496 |
| AT2G30860.1 | 0,792939459 | 0,003033886 | GSTF9_ ATGSTF9_ ATGSTF7_ GLUTTR glutathione S-transferase PHI 9 chr2:13139132-13140057 FORW |
| AT4G35000.1 | 1,451861877 | 0,003034319 | APX3 ascorbate peroxidase 3 chr4:16665007-16667541 REVERSE LENGTH=287 |
| AT5G17310.2 | 1,32833632 | 0,003036524 | AtUGP2_ UGP2 UDP-GLUCOSE PYROPHOSPHORYLASE 2_ UDP-glucose pyrophosphorylase 2 chr5:569 |
| AT5G16400.1 | 0,680599722 | 0,003085839 | TRXF2_ ATF2 thioredoxin F2 chr5:5363905-5365249 REVERSE LENGTH=185 |
| AT2G39330.1 | 0,60334081 | 0,003127031 | JAL23 jacalin-related lectin 23 chr2:16419787-16421573 REVERSE LENGTH=459 |

| | | | |
|---|---|---|---|
| AT5G40950.1 | 0,569200117 | 0,003146656 | PRPL27_ RPL27 ribosomal protein large subunit 27 chr5:16410866-16411845 FORWARD LENGTH=198 |
| ATCG00740.1 | 0,803957397 | 0,003159372 | RPOA RNA polymerase subunit alpha chrc:77901-78890 REVERSE LENGTH=329 |
| AT4G17470.1 | 0,565703424 | 0,003184884 | CRSH Ca2+-activated RelA-spot homolog chr4:9742922-9744468 REVERSE LENGTH=308 |
| AT5G48880.1 | 0,729881601 | 0,003187402 | PKT2_ PKT1_ KAT5 3-KETO-ACYL-COENZYME A THIOLASE 5_ peroxisomal 3-keto-acyl-CoA thiolase |
| AT1G78380.1 | 1,456015906 | 0,003193313 | GSTU19_ GST8_ ATGSTU19 A. THALIANA GLUTATHIONE S-TRANSFERASE TAU 19_ glutathione S-[illegible] |
| AT3G24170.1 | 1,406739402 | 0,003210073 | ATGR1_ GR1 glutathione-disulfide reductase chr3:8729762-8734115 REVERSE LENGTH=499 |
| AT3G12260.1 | 0,641110533 | 0,003255368 | NDUFA6_ B14 chr3:3909252-3910337 REVERSE LENGTH=133 |
| AT1G11750.1 | 1,558094467 | 0,003273407 | NCLPP1_ CLPP6_ NCLPP6 NUCLEAR-ENCODED CLPP 1_ CLP protease proteolytic subunit 6 chr1:39676[illegible] |
| AT1G14250.1 | 0,409413455 | 0,003277551 | no symbol available no full name available chr1:4868675-4871203 FORWARD LENGTH=488 |
| AT3G29360.1 | 1,510851414 | 0,003301342 | UGD2 UDP-glucose dehydrogenase 2 chr3:11267375-11268817 REVERSE LENGTH=480 |
| AT1G29880.1 | 1,365847428 | 0,003301725 | no symbol available no full name available chr1:10459662-10462781 REVERSE LENGTH=729 |
| AT1G74040.1 | 1,978292637 | 0,003302545 | IPMS2_ IMS1_ MAML-3 SOPROPYLMALATE SYNTHASE 2_ 2-isopropylmalate synthase 1 chr1:2784225[illegible] |
| AT1G76180.1 | 0,529713325 | 0,003326804 | ERD14 EARLY RESPONSE TO DEHYDRATION 14 chr1:28587013-28587657 REVERSE LENGTH=185 |
| AT4G11820.1 | 1,489989782 | 0,003403642 | MVA1_ FKP1_ HMGS FLAKY POLLEN 1_ HYDROXYMETHYLGLUTARYL-COA SYNTHASE chr4:71[illegible] |
| AT5G28540.1 | 1,495836019 | 0,003418231 | BIP1 chr5:10540665-10543274 REVERSE LENGTH=669 |
| ATCG00905.1 | 0,481260486 | 0,00342037 | RPS12_ RPS12C RIBOSOMAL PROTEIN S12_ ribosomal protein S12C chrc:97999-98793 REVERSE LEN[illegible] |
| AT3G63190.1 | 0,790910668 | 0,003443968 | cpRRF_ HFP108_ AtcpRRF_ RRF chloroplast ribosome recycling factor_ Arabidopsis thaliana chloroplast rib[illegible] |
| AT1G12900.1 | 1,112656438 | 0,00348187 | GAPA-2 glyceraldehyde 3-phosphate dehydrogenase A subunit 2 chr1:4392634-4394283 REVERSE LENGTH |
| AT1G35680.1 | 0,856075428 | 0,003518713 | RPL21C_ ASD chloroplast ribosomal protein L21_ ATPase-in-Seed-Development chr1:13208777-13210246 F[illegible] |
| AT4G31500.1 | 0,558113758 | 0,003519012 | RNT1_ RED1_ SUR2_ ATR4_ CYP83B1 RED ELONGATED 1_ SUPERROOT 2_ ALTERED TRYPTOPH[illegible] |
| AT1G64740.1 | 0,830303734 | 0,003532398 | TUA1 alpha-1 tubulin chr1:24050114-24052296 FORWARD LENGTH=450 |
| AT1G19670.1 | 0,632494491 | 0,003651746 | CLH1_ ATCLH1_ CORI1_ ATHCOR1 chlorophyllase 1_ CORONATINE-INDUCED PROTEIN 1 chr1:6803[illegible] |
| AT1G32200.1 | 0,765450792 | 0,003666348 | ACT1_ ATS1 ACYLTRANSFERASE 1 chr1:11602223-11605001 REVERSE LENGTH=459 |
| AT5G23540.1 | 1,107260467 | 0,003675843 | no symbol available no full name available chr5:7937772-7939339 FORWARD LENGTH=308 |
| AT4G24930.1 | 0,630776849 | 0,003803847 | no symbol available no full name available chr4:12821496-12822389 REVERSE LENGTH=225 |
| AT1G09620.1 | 1,479617316 | 0,003835362 | no symbol available no full name available chr1:3113077-3116455 REVERSE LENGTH=1091 |
| AT2G20260.1 | 0,821910791 | 0,003866322 | PSAE-2 photosystem I subunit E-2 chr2:8736780-8737644 FORWARD LENGTH=145 |
| AT2G43460.1 | 0,434798845 | 0,003877514 | no symbol available no full name available chr2:18046285-18047292 REVERSE LENGTH=69 |
| AT2G25080.1 | 0,760696806 | 0,003891974 | GPX1_ ATGPX1_ GPXL1 GLUTATHIONE PEROXIDASE 1_ glutathione peroxidase 1 chr2:10668134-1066[illegible] |
| AT5G09660.1 | 1,311134495 | 0,003959247 | PMDH2 peroxisomal NAD-malate dehydrogenase 2 chr5:2993645-2995551 REVERSE LENGTH=354 |
| AT4G22485.1 | 0,711185621 | 0,004006683 | no symbol available no full name available chr4:11844506-11846476 REVERSE LENGTH=656 |
| AT1G77590.2 | 2,007675624 | 0,004032792 | LACS9 long chain acyl-CoA synthetase 9 chr1:29148501-29151234 REVERSE LENGTH=545 |
| AT3G21370.1 | 1,854555829 | 0,004033657 | BGLU19 beta glucosidase 19 chr3:7524286-7527579 REVERSE LENGTH=527 |
| AT3G58610.1 | 1,487436241 | 0,004116513 | no symbol available no full name available chr3:21671561-21674639 FORWARD LENGTH=591 |
| AT4G09670.1 | 1,613963961 | 0,004118598 | no symbol available no full name available chr4:6107382-6109049 REVERSE LENGTH=362 |
| AT2G24940.1 | 0,722406233 | 0,004124074 | MAPR2_ AtMAPR2 membrane-associated progesterone binding protein 2 chr2:10609447-10609749 FORWA[illegible] |
| AT1G12250.2 | 0,644953474 | 0,004145802 | TL20.3 chr1:4159623-4161269 FORWARD LENGTH=206 |
| AT5G02500.1 | 1,361320074 | 0,004149063 | AtHsp70-1_ HSP70-1_ HSC70-1_ HSC70_ AT-HSC70-1 ARABIDOPSIS THALIANA HEAT SHOCK COG[illegible] |
| AT3G28220.1 | 0,773089064 | 0,0041918 | no symbol available no full name available chr3:10524420-10526497 FORWARD LENGTH=370 |

| | | | |
|---|---|---|---|
| AT1G04270.2 | 0,581823097 | 0,004206005 | RPS15 cytosolic ribosomal protein S15 chr1:1141852-1142960 REVERSE LENGTH=151 |
| AT4G02510.1 | 1,271597246 | 0,004303775 | TOC86_ ATTOC159_ TOC160_ PPI2_ TOC159 translocon at the outer envelope membrane of chloroplasts 15 |
| AT5G24420.1 | 0,577801268 | 0,004305012 | PGL5 6-phosphogluconolactonase 5 chr5:8336943-8337879 REVERSE LENGTH=252 |
| AT5G11520.1 | 0,872876106 | 0,00432232 | YLS4_ ASP3 aspartate aminotransferase 3_ YELLOW-LEAF-SPECIFIC GENE 4 chr5:3685257-3687721 REV |
| AT3G22960.1 | 1,395049367 | 0,004398268 | PKP1_ PKP-ALPHA PLASTIDIAL PYRUVATE KINASE 1 chr3:8139369-8141771 FORWARD LENGTH= |
| AT4G33680.1 | 1,17490206 | 0,004406101 | AGD2 ABERRANT GROWTH AND DEATH 2_ ARF-GAP domain 2 chr4:16171847-16174630 REVERSE L |
| AT2G07732.1 | 1,369520225 | 0,004425176 | no symbol available no full name available chr2:3468424-3468774 REVERSE LENGTH=116 |
| AT1G70770.1 | 1,592963879 | 0,004459838 | no symbol available no full name available chr1:26688622-26691185 REVERSE LENGTH=610 |
| AT5G28840.1 | 1,262495371 | 0,004461606 | GME GDP-D-mannose 3 chr5:10862472-10864024 REVERSE LENGTH=377 |
| AT1G62740.1 | 0,758677445 | 0,004462454 | Hop2 Hop2 chr1:23231026-23233380 FORWARD LENGTH=571 |
| AT1G78300.1 | 1,724943601 | 0,004523821 | GRF2_ 14-3-3OMEGA_ GF14 OMEGA general regulatory factor 2_ 14-3-3 PROTEIN G-BOX FACTOR14 O |
| AT3G04550.1 | 0,801322021 | 0,004534921 | RAF1 Rubisco accumulation factor 1 chr3:1225961-1227310 FORWARD LENGTH=449 |
| AT3G63540.1 | 0,669862589 | 0,004557736 | no symbol available no full name available chr3:23459372-23459803 REVERSE LENGTH=143 |
| AT2G02930.1 | 1,186677785 | 0,00463791 | ATGSTF3_ GST16_ GSTF3 GLUTATHIONE S-TRANSFERASE 16_ glutathione S-transferase F3 chr2:8513 |
| AT1G74970.1 | 1,150926005 | 0,004643194 | TWN3_ SOT8_ RPS9_ PRPS9 ribosomal protein S9 chr1:28157761-28159202 REVERSE LENGTH=208 |
| AT4G27560.1 | 1,397420914 | 0,004663418 | UGT79B2 chr4:13760114-13761481 REVERSE LENGTH=455 |
| AT5G48580.1 | 0,43830809 | 0,004680934 | FKBP15-2 FK506- and rapamycin-binding protein 15 kD-2 chr5:19696156-19697304 REVERSE LENGTH=1 |
| AT5G19770.1 | 1,272656838 | 0,004725621 | TUA3 tubulin alpha-3 chr5:6682761-6684474 REVERSE LENGTH=450 |
| AT1G49630.1 | 0,609205067 | 0,004755177 | PREP2_ ATPREP2 presequence protease 2 chr1:18368405-18375336 REVERSE LENGTH=1080 |
| AT3G23810.1 | 1,206964652 | 0,004761293 | ATSAHH2_ SAHH2 S-ADENOSYL-L-HOMOCYSTEINE (SAH) HYDROLASE 2_ S-adenosyl-l-homocystei |
| AT1G66270.2 | 0,25846767 | 0,004823284 | BGLU21 chr1:24700110-24702995 REVERSE LENGTH=522 |
| AT4G20260.1 | 0,762818935 | 0,004838547 | ATPCAP1_ PCAP1_ MDP25 ARABIDOPSIS THALIANA PLASMA-MEMBRANE ASSOCIATED CATION |
| AT2G34810.1 | 0,50284369 | 0,004929975 | AtBBE16 chr2:14685292-14686914 FORWARD LENGTH=540 |
| AT2G39590.1 | 1,125293379 | 0,004948826 | no symbol available no full name available chr2:16517588-16518247 REVERSE LENGTH=130 |
| AT2G27720.1 | 0,798347057 | 0,004956831 | no symbol available no full name available chr2:11818696-11819370 FORWARD LENGTH=115 |
| AT5G25980.2 | 0,873627669 | 0,004971915 | BGLU37_ TGG2 BETA GLUCOSIDASE 37_ glucoside glucohydrolase 2 chr5:9072730-9075477 FORWARD |
| AT1G35160.1 | 1,796221573 | 0,004994914 | GRF4_ 14-3-3PHI_ GF14 PHI GENERAL REGULATORY FACTOR 4_ 14-3-3 PROTEIN G-BOX FACTOR |
| AT5G26830.1 | 1,414102231 | 0,005002524 | no symbol available no full name available chr5:9437351-9441568 FORWARD LENGTH=709 |
| AT3G04840.1 | 0,822189143 | 0,005031297 | no symbol available no full name available chr3:1329751-1331418 FORWARD LENGTH=262 |
| AT3G16100.1 | 0,384648988 | 0,005056423 | RABG3c_ ATRAB7D_ ATRABG3C RAB GTPase homolog G3C chr3:5459270-5460556 FORWARD LENG |
| AT1G56050.1 | 0,595949567 | 0,005062975 | EngD-2 chr1:20963793-20966181 FORWARD LENGTH=421 |
| AT3G09790.1 | 0,581940211 | 0,005069613 | UBQ8 ubiquitin 8 chr3:3004111-3006006 REVERSE LENGTH=631 |
| AT3G48110.1 | 1,804446961 | 0,005092608 | EDD1_ EDD EMBRYO-DEFECTIVE-DEVELOPMENT 1 chr3:17763111-17770964 FORWARD LENGTH= |
| AT2G05920.1 | 0,56751981 | 0,005097022 | SBT1.8 subtilase 1.8 chr2:2269831-2272207 REVERSE LENGTH=754 |
| AT3G17020.1 | 1,29947275 | 0,005110402 | no symbol available no full name available chr3:5802728-5804063 REVERSE LENGTH=163 |
| AT4G16143.1 | 1,483952837 | 0,005145061 | IMPA-2 importin alpha isoform 2 chr4:9134450-9137134 REVERSE LENGTH=535 |
| AT1G74260.1 | 1,691693302 | 0,005145082 | PUR4 purine biosynthesis 4 chr1:27923005-27927764 REVERSE LENGTH=1407 |
| AT5G14590.1 | 1,229324599 | 0,005153349 | no symbol available no full name available chr5:4703533-4706627 REVERSE LENGTH=485 |
| AT2G42740.1 | 0,923616053 | 0,005182765 | RPL16A ribosomal protein large subunit 16A chr2:17791794-17792946 FORWARD LENGTH=182 |

| | | | |
|---|---|---|---|
| AT5G20980.1 | 1,213145229 | 0,005212338 | MS3_ ATMS3 methionine synthase 3 chr5:7124397-7128353 REVERSE LENGTH=812 |
| AT2G43030.1 | 0,527940666 | 0,005222443 | PRPL3 plastid ribosomal proteins of the 50S subunit chr2:17894898-17895713 FORWARD LENGTH=271 |
| AT2G04842.1 | 0,696445031 | 0,005228029 | EMB2761 EMBRYO DEFECTIVE 2761 chr2:1698466-1701271 REVERSE LENGTH=650 |
| AT5G16990.1 | 0,647442658 | 0,005269801 | no symbol available no full name available chr5:5581831-5583849 REVERSE LENGTH=343 |
| AT5G14660.1 | 0,514574538 | 0,005303721 | ATDEF2_ DEF2_ PDF1B peptide deformylase 1B chr5:4727129-4728671 REVERSE LENGTH=273 |
| AT1G72150.1 | 0,771043623 | 0,005311926 | PATL1 PATELLIN 1 chr1:27148558-27150652 FORWARD LENGTH=573 |
| AT3G46780.1 | 1,26949589 | 0,00532937 | PTAC16 plastid transcriptionally active 16 chr3:17228766-17231021 FORWARD LENGTH=510 |
| AT1G80380.4 | 1,154063705 | 0,005335035 | GLYK glycerate kinase chr1:30217332-30219784 FORWARD LENGTH=450 |
| AT3G60750.1 | 1,143228738 | 0,005385146 | AtTKL1_ TKL1 transketolase 1 chr3:22454004-22456824 FORWARD LENGTH=741 |
| AT1G10760.1 | 1,217062584 | 0,005426011 | GWD_ GWD1_ SOP1_ SOP_ SEX1 STARCH EXCESS 1 chr1:3581210-3590043 REVERSE LENGTH=139 |
| AT4G27670.1 | 58,7372134 | 0,005426077 | HSP21 heat shock protein 21 chr4:13819048-13819895 REVERSE LENGTH=227 |
| AT3G12110.1 | 1,353071191 | 0,005464165 | ACT11 actin-11 chr3:3858116-3859609 FORWARD LENGTH=377 |
| AT5G13630.1 | 0,75281453 | 0,005465198 | ABAR_ CHLH_ GUN5_ CCH_ CCH1 ABA-BINDING PROTEIN_ H SUBUNIT OF MG-CHELATASE_ GE |
| AT4G24780.1 | 0,638398958 | 0,00547466 | PLL19 chr4:12770631-12772227 REVERSE LENGTH=408 |
| AT5G09510.2 | 0,404997962 | 0,005485927 | no symbol available no full name available chr5:2955698-2956353 REVERSE LENGTH=118 |
| AT3G52990.1 | 1,276542044 | 0,005550273 | no symbol available no full name available chr3:19649046-19652237 FORWARD LENGTH=527 |
| AT3G56240.1 | 0,806783099 | 0,005566086 | AtHMP31_ CCH HEAVY METAL ASSOCIATED PROTEIN 31_ copper chaperone chr3:20863460-2086440 |
| ATCG00750.1 | 0,421065748 | 0,005584315 | RPS11 ribosomal protein S11 chrc:78960-79376 REVERSE LENGTH=138 |
| AT4G08870.1 | 0,809269096 | 0,005647762 | ARGAH2 arginine amidohydrolase 2 chr4:5646654-5648693 REVERSE LENGTH=344 |
| AT5G11420.1 | 1,254368137 | 0,00568489 | no symbol available no full name available chr5:3644655-3646991 FORWARD LENGTH=366 |
| AT5G20080.1 | 0,797317141 | 0,005691049 | no symbol available no full name available chr5:6782708-6786360 FORWARD LENGTH=328 |
| AT1G78570.1 | 0,774602351 | 0,005706126 | RHM1_ ATRHM1_ ROL1 REPRESSOR OF LRX1 1_ rhamnose biosynthesis 1_ ARABIDOPSIS THALIANA |
| AT1G66970.1 | 0,673520116 | 0,005820988 | SVL2_ GDPDL1 SHV3-like 2_ Glycerophosphodiester phosphodiesterase (GDPD) like 1 chr1:24992746-2499 |
| AT3G57490.1 | 0,433437141 | 0,00583067 | no symbol available no full name available chr3:21279824-21280887 REVERSE LENGTH=276 |
| AT1G80560.1 | 0,760056344 | 0,005840159 | ATIMD2_ IMD2 isopropylmalate dehydrogenase 2_ ARABIDOPSIS ISOPROPYLMALATE DEHYDROGEN |
| AT1G55060.1 | 0,800443059 | 0,005871909 | UBQ12 ubiquitin 12 chr1:20549533-20550225 FORWARD LENGTH=230 |
| AT1G03475.1 | 0,792703337 | 0,005872985 | HEMF1_ ATCPO-I_ LIN2 LESION INITIATION 2 chr1:869302-871175 REVERSE LENGTH=386 |
| AT2G04700.1 | 0,602744829 | 0,005887952 | INAP1_ FTRB Imbalanced NADP Status 1_ ferredoxin/thioredoxin reductase catayltic subunit chr2:1646961-1 |
| AT2G41840.1 | 0,627330506 | 0,005893678 | no symbol available no full name available chr2:17460016-17461398 REVERSE LENGTH=285 |
| AT3G45140.1 | 0,849212547 | 0,005906651 | ATLOX2_ LOX2 ARABIODOPSIS THALIANA LIPOXYGENASE 2_ lipoxygenase 2 chr3:16525437-16529 |
| AT1G41880.1 | 0,147568307 | 0,005940267 | no symbol available no full name available chr1:15651585-15652427 REVERSE LENGTH=111 |
| AT5G38480.1 | 1,440156379 | 0,005949132 | GRF3_ RCI1 general regulatory factor 3 chr5:15410277-15411285 FORWARD LENGTH=255 |
| AT3G62410.1 | 0,63606685 | 0,006098385 | CP12_ CP12-2 CP12 DOMAIN-CONTAINING PROTEIN 1_ CP12 domain-containing protein 2 chr3:230910 |
| AT3G57890.1 | 1,385120968 | 0,006120566 | no symbol available no full name available chr3:21438271-21441695 FORWARD LENGTH=573 |
| AT2G29630.1 | 0,475611862 | 0,006152426 | THIC_ PY PYRIMIDINE REQUIRING_ thiaminC chr2:12667395-12669569 FORWARD LENGTH=644 |
| AT5G48300.1 | 1,3307267 | 0,006235916 | ADG1_ APS1 ADP-GLUCOSE PYROPHOSPHORYLASE SMALL SUBUNIT 1_ ADP glucose pyrophospho |
| AT1G43170.1 | 0,102457725 | 0,006426911 | emb2207_ RP1_ ARP1_ RPL3A embryo defective 2207_ ribosomal protein 1 chr1:16266992-16268631 FORW |
| AT2G41100.5 | 0,600143612 | 0,006427807 | TCH3_ CML12_ ATCAL4 ARABIDOPSIS THALIANA CALMODULIN LIKE 4_ calmodulin-like 12_ TOU |
| AT3G12915.2 | 0,641977424 | 0,006454867 | no symbol available no full name available chr3:4112834-4115708 FORWARD LENGTH=788 |

| | | | |
|---|---|---|---|
| AT4G16760.2 | 0,818048383 | 0,006514703 | ATACX1_ ACX1 acyl-CoA oxidase 1 chr4:9424930-9428689 REVERSE LENGTH=651 |
| AT5G53560.1 | 0,555237391 | 0,006516126 | ATB5-A_ CB5-E_ ATCB5-E_ B5 #2 ARABIDOPSIS CYTOCHROME B5 ISOFORM E_ cytochrome B5 isof |
| AT1G20440.1 | 1,22002593 | 0,006539429 | AtCOR47_ RD17_ COR47 cold-regulated 47 chr1:7084722-7085664 REVERSE LENGTH=265 |
| AT4G18040.2 | 0,704902038 | 0,006592319 | EIF4A-2 eif4a-2 chr1:20260495-20262018 FORWARD LENGTH=407 |
| AT3G52150.1 | 0,730490226 | 0,006626378 | PSRP2 plastid-speciﬁc ribosomal protein 2 chr3:19342074-19343090 FORWARD LENGTH=253 |
| AT5G17920.1 | 1,160775219 | 0,006649719 | ATCIMS_ METS1_ ATMETS_ ATMS1 COBALAMIN-INDEPENDENT METHIONINE SYNTHASE_ meth |
| AT5G45280.1 | 0,159492058 | 0,006669191 | PAE11 pectin acetylesterase 11 chr5:18346862-18349432 FORWARD LENGTH=370 |
| AT1G74090.1 | 0,388848302 | 0,006703155 | ATST5B_ SOT18_ ATSOT18 DESULFO-GLUCOSINOLATE SULFOTRANSFERASE 18_ ARABIDOPSIS |
| AT5G48480.1 | 2,097689639 | 0,006733205 | no symbol available no full name available chr5:19644814-19645658 FORWARD LENGTH=166 |
| AT5G54270.1 | 1,065715415 | 0,006802153 | LHCB3_ LHCB31 light-harvesting chlorophyll B-binding protein 3 chr5:22038424-22039383 FORWARD LEI |
| AT5G25460.1 | 0,802588782 | 0,006858738 | DGR2 DUF642 L-GalL responsive gene 2 chr5:8863430-8865394 FORWARD LENGTH=369 |
| AT3G02560.1 | 0,565852315 | 0,007012432 | no symbol available no full name available chr3:542341-543168 FORWARD LENGTH=191 |
| AT1G52570.1 | 0,595928401 | 0,007013223 | PLDALPHA2 phospholipase D alpha 2 chr1:19583940-19586551 REVERSE LENGTH=810 |
| AT1G29670.1 | 0,637487743 | 0,007051721 | GDSL1_ GGL6 chr1:10375843-10377717 FORWARD LENGTH=363 |
| AT2G22990.1 | 0,302186671 | 0,007123487 | SNG1_ SCPL8 sinapoylglucose 1_ SERINE CARBOXYPEPTIDASE-LIKE 8 chr2:9786393-9789925 FORWA |
| AT2G37190.1 | 0,690616159 | 0,007183653 | no symbol available no full name available chr2:15619559-15620059 REVERSE LENGTH=166 |
| AT1G29150.1 | 1,488293585 | 0,007250459 | RPN6_ ATS9 non-ATPase subunit 9_ REGULATORY PARTICLE NON-ATPASE 6 chr1:10181240-1018249 |
| AT5G14260.1 | 1,363838413 | 0,007354853 | SAFE1 SAFEGUARD1 chr5:4601139-4603873 FORWARD LENGTH=514 |
| AT3G26450.1 | 1,276039119 | 0,007378205 | no symbol available no full name available chr3:9681593-9683299 REVERSE LENGTH=152 |
| AT3G32980.1 | 0,803998152 | 0,007478133 | PRX32 Peroxidase 32 chr3:13526404-13529949 REVERSE LENGTH=352 |
| AT1G11650.1 | 1,786956014 | 0,007479446 | RBP45B_ ATRBP45B chr1:3914895-3917301 FORWARD LENGTH=306 |
| AT3G59760.3 | 0,696482814 | 0,007513726 | OASC_ ATCS-C ARABIDOPSIS THALIANA CYSTEINSYNTHASE-C_ O-acetylserine (thiol) lyase isoform |
| AT4G22890.4 | 1,675961987 | 0,007637793 | PGR5-LIKE A chr4:12007157-12009175 FORWARD LENGTH=321 |
| AT4G09040.1 | 0,775130119 | 0,00767691 | CP33C chr4:5795075-5797315 REVERSE LENGTH=304 |
| AT1G04040.1 | 1,994882938 | 0,007678539 | no symbol available no full name available chr1:1042564-1043819 REVERSE LENGTH=271 |
| AT5G56500.1 | 0,529203814 | 0,00778331 | CPNB3_ Cpn60beta3 chaperonin-60beta3 chr5:22874058-22876966 FORWARD LENGTH=597 |
| ATCG00900.1 | 0,450597188 | 0,007807092 | RPS7_ RPS7.1 CHLOROPLAST RIBOSOMAL PROTEIN S7 chrc:97478-97945 REVERSE LENGTH=155 |
| AT1G07770.1 | 0,848232866 | 0,00789727 | RPS15A ribosomal protein S15A chr1:2408413-2409065 REVERSE LENGTH=130 |
| AT4G30190.1 | 1,496995239 | 0,007910252 | HA2_ PMA2_ AtHA2_ AHA2 H(+)-ATPase 2_ PLASMA MEMBRANE PROTON ATPASE 2 chr4:1477082 |
| AT2G44050.1 | 1,360132083 | 0,007948382 | COS1 COI1 SUPPRESSOR1_ coronatine insensitive1 suppressor chr2:18224304-18225917 FORWARD LENG |
| AT1G19580.1 | 2,105070695 | 0,007954749 | GAMMA CA1 gamma carbonic anhydrase 1 chr1:6774937-6777092 FORWARD LENGTH=275 |
| AT5G10450.1 | 1,155646987 | 0,008091015 | 14-3-3lambda_ GRF6_ AFT1 14-3-3 PROTEIN G-BOX FACTOR14 LAMBDA_ G-box regulating factor 6 ch |
| AT3G63140.1 | 0,851511666 | 0,008099145 | CSP41A chloroplast stem-loop binding protein of 41 kDa chr3:23327006-23328620 REVERSE LENGTH=406 |
| AT1G43560.1 | 2,026178086 | 0,008152659 | Aty2_ ty2 thioredoxin Y2 chr1:16398359-16399828 REVERSE LENGTH=167 |
| AT5G07340.1 | 0,660758425 | 0,008233679 | no symbol available no full name available chr5:2317300-2319458 FORWARD LENGTH=532 |
| AT3G62120.3 | 1,340082165 | 0,008243679 | ProRS-Cyt_ AtProRS-Cyt prolyl-tRNA synthetase cytosolic chr3:23001227-23003849 REVERSE LENGTH=5 |
| AT1G08520.1 | 0,833475375 | 0,008254937 | ALB1_ PDE166_ ALB-1V_ CHLD_ V157 PIGMENT DEFECTIVE EMBRYO 166_ ALBINA 1 chr1:269653 |
| AT3G14420.1 | 1,145497054 | 0,008266689 | GOX1 glycolate oxidase 1 chr3:4821804-4823899 FORWARD LENGTH=367 |
| AT2G45740.1 | 1,727657796 | 0,008295207 | PEX11D peroxin 11D chr2:18839865-18841102 FORWARD LENGTH=236 |

| | | | |
|---|---|---|---|
| AT3G43980.1 | 0,377006246 | 0,008314216 | no symbol available no full name available chr3:15778555-15779235 REVERSE LENGTH=56 |
| AT1G53750.1 | 1,21416602 | 0,008338215 | RPT1A regulatory particle triple-A 1A chr1:20065921-20068324 REVERSE LENGTH=426 |
| AT3G05560.1 | 0,812516724 | 0,008348612 | no symbol available no full name available chr3:1614641-1615204 FORWARD LENGTH=124 |
| AT1G72810.1 | 2,173888504 | 0,008377869 | TSY THREONINE SYNTHASE 2 chr1:27398760-27400393 REVERSE LENGTH=516 |
| AT3G19710.1 | 1,335583946 | 0,008379936 | BCAT4 branched-chain aminotransferase4 chr3:6847202-6849429 REVERSE LENGTH=354 |
| AT5G40370.1 | 0,836189365 | 0,008433148 | AtGRXC2_ GRXC2_ GRX370 glutaredoxin C2 chr5:16147826-16149052 REVERSE LENGTH=111 |
| AT2G22240.1 | 9,561788548 | 0,008438963 | MIPS2_ ATIPS2_ ATMIPS2 myo-inositol-1-phosphate synthase 2_ INOSITOL 3-PHOSPHATE SYNTHASE |
| AT1G11430.1 | 0,689545102 | 0,008506573 | RIP9_ MORF9 multiple organellar RNA editing factor 9 chr1:3847273-3848938 FORWARD LENGTH=232 |
| AT2G27710.1 | 0,821128408 | 0,008562934 | no symbol available no full name available chr2:11816929-11817670 FORWARD LENGTH=115 |
| AT5G35360.1 | 0,72025862 | 0,00857859 | CAC2 acetyl Co-enzyme a carboxylase biotin carboxylase subunit chr5:13584300-13588268 FORWARD LEN |
| AT5G60360.2 | 0,134226265 | 0,008611781 | AALP_ SAG2_ ALP aleurain-like protease_ SENESCENCE ASSOCIATED GENE2 chr5:24280044-2428215 |
| AT1G56450.1 | 1,118465086 | 0,008685672 | MUD1_ PBG1 20S proteasome beta subunit G1 chr1:21141970-21144186 FORWARD LENGTH=246 |
| AT1G10840.1 | 1,573872355 | 0,008697309 | TIF3H1 translation initiation factor 3 subunit H1 chr1:3607885-3610299 REVERSE LENGTH=337 |
| AT2G47400.1 | 0,848534748 | 0,008700016 | CP12_ CP12-1 CP12 DOMAIN-CONTAINING PROTEIN 1_ CP12 domain-containing protein 1 chr2:194468 |
| AT4G23900.1 | 0,742866345 | 0,008733735 | no symbol available no full name available chr4:12424505-12426318 FORWARD LENGTH=237 |
| AT3G17810.1 | 0,834053949 | 0,008766735 | PYD1 pyrimidine 1 chr3:6094279-6096289 FORWARD LENGTH=426 |
| ATCG00130.1 | 1,199690466 | 0,0088088 | ATPF chrc:11529-12798 REVERSE LENGTH=184 |
| ATCG00480.1 | 1,068186748 | 0,008843695 | PB_ ATPB_ CF1beta_ AthCF1beta ATP synthase subunit beta chrc:52660-54156 REVERSE LENGTH=498 |
| AT4G30690.2 | 0,151555093 | 0,008922574 | AtINFC-4_ SVR9L_ AtIF3- 4 SVR9-LIKE1_ Initiation factor 3-4 chr4:14960742-14962328 FORWARD LEN |
| AT4G39330.1 | 1,223725816 | 0,008937088 | ATCAD9_ CAD9 cinnamyl alcohol dehydrogenase 9 chr4:18291268-18292772 FORWARD LENGTH=360 |
| AT5G11200.2 | 1,163399362 | 0,008974824 | UAP56b homolog of human UAP56 b chr5:3567389-3570686 FORWARD LENGTH=486 |
| AT5G47870.1 | 1,6637246 | 0,008990784 | RAD52-2_ ODB2_ RAD52-2B radiation sensitive 52-2_ Organellar DNA-Binding protein 2 chr5:19384555-19 |
| AT1G07320.3 | 0,567870566 | 0,009228641 | RPL4_ PRPL4_ EMB2784 plastid ribosomal protein L4_ ribosomal protein L4_ EMBRYO DEFECTIVE 2784 |
| AT5G16510.1 | 1,38128818 | 0,009279546 | RGP5 reversibly glycosylated polypeptide 5_ reversibly glycosylated protein 5 chr5:5393296-5394342 FORW |
| AT2G31610.1 | 1,199055709 | 0,009384063 | no symbol available no full name available chr2:13450384-13451669 FORWARD LENGTH=250 |
| AT4G05180.1 | 0,871228923 | 0,009477486 | PSBQ-2_ PSBQ_ PSII-Q photosystem II subunit Q-2_ PHOTOSYSTEM II SUBUNIT Q chr4:2672093-26731 |
| AT3G18780.1 | 1,633639271 | 0,009532062 | LSR2_ ACT2_ ENL2_ DER1_ FIZ2 FRIZZY AND KINKED SHOOTS 2_ LIGHT STRESS-REGULATED 2 |
| AT3G49680.2 | 1,660158061 | 0,009625666 | BCAT3_ ATBCAT-3 branched-chain aminotransferase 3 chr3:18422768-18425473 FORWARD LENGTH=41 |
| AT5G43850.1 | 3,691194568 | 0,009627542 | ATARD4_ ARD4 chr5:17627364-17629122 REVERSE LENGTH=187 |
| AT2G42690.1 | 1,643291712 | 0,009661993 | AGAP1 ACYLATED GALACTOLIPID- ASSOCIATED PHOSPHOLIPASE 1 chr2:17776356-17777682 REV |
| AT5G15650.1 | 0,846737746 | 0,009680439 | MUR5_ ATRGP2_ RGP2 MURUS 5_ REVERSIBLY GLYCOSYLATED POLYPEPTIDE 2_ reversibly glyc |
| AT5G63570.1 | 1,088960297 | 0,009866887 | GSA1 "glutamate-1-semialdehyde-2_1-aminomutase" chr5:25451957-25453620 FORWARD LENGTH=474 |
| AT1G75040.1 | 0,479991993 | 0,009899126 | PR-5_ PR5 pathogenesis-related gene 5 chr1:28177754-28178731 FORWARD LENGTH=239 |
| AT3G12390.1 | 0,796747286 | 0,009995162 | no symbol available no full name available chr3:3942344-3943595 FORWARD LENGTH=203 |
| AT4G04640.1 | 1,107474053 | 0,010091307 | ATPC1 chr4:2350761-2351882 REVERSE LENGTH=373 |
| AT3G26740.1 | 0,591032466 | 0,01013298 | CCL CCR-like chr3:9827868-9828461 FORWARD LENGTH=141 |
| AT3G48930.1 | 0,28989601 | 0,010262554 | EMB1080 embryo defective 1080 chr3:18141017-18142189 REVERSE LENGTH=160 |
| AT5G07030.1 | 0,834998404 | 0,010322212 | no symbol available no full name available chr5:2183600-2185717 REVERSE LENGTH=455 |
| AT4G36250.1 | 1,201190113 | 0,010347161 | ALDH3F1 aldehyde dehydrogenase 3F1 chr4:17151029-17153381 FORWARD LENGTH=484 |

| | | | |
|---|---|---|---|
| AT1G52400.1 | 0,73068973 | 0,010595859 | BGL1_ ATBG1_ BGLU18 A. THALIANA BETA-GLUCOSIDASE 1_ BETA-GLUCOSIDASE HOMOLOG |
| AT5G16590.1 | 1,435177169 | 0,010645177 | LRR1 Leucine rich repeat protein 1 chr5:5431862-5433921 FORWARD LENGTH=625 |
| AT1G59870.1 | 1,159584977 | 0,010648734 | ABCG36_ ATABCG36_ PEN3_ PDR8_ ATPDR8 Arabidopsis thaliana ATP-binding cassette G36_ PLEIOTR |
| AT5G51110.1 | 0,698487512 | 0,010685856 | ATP1_ SDIRIP1_ RAF2 SDIR1-INTERACTING PROTEIN1_ AtAIRP2 Target Protein 1_ Rubisco Assembly |
| ATCG00350.1 | 1,26245526 | 0,01069574 | PSAA chrc:39605-41857 REVERSE LENGTH=750 |
| AT2G47110.1 | 0,755318191 | 0,010766806 | UBI6_ UBQ6_ RPS27aB UBIQUITIN EXTENSION PROTEIN 6_ ubiquitin 6_ Ribosomal protein S27aB chr: |
| AT1G49240.1 | 1,159367371 | 0,010993686 | ACT8_ FIZ1 FRIZZY AND KINKED SHOOTS_ actin 8 chr1:18216539-18217947 FORWARD LENGTH=37 |
| AT5G20920.2 | 0,646503158 | 0,011145007 | EIF2 BETA_ EMB1401_ eIF-2bs embryo defective 1401_ eukaryotic translation initiation factor 2 beta subunit |
| AT5G62670.1 | 10,08466278 | 0,011198002 | HA11_ AHA11 H(+)-ATPase 11 chr5:25159495-25164957 FORWARD LENGTH=956 |
| AT2G32120.1 | 2,114679878 | 0,011253702 | HSP70T-2 heat-shock protein 70T-2 chr2:13651720-13653411 REVERSE LENGTH=563 |
| AT3G27240.1 | 0,688637888 | 0,011275693 | Cyc1-1 chr3:10056144-10058370 REVERSE LENGTH=307 |
| AT5G35790.1 | 1,215405578 | 0,011404429 | G6PD1 glucose-6-phosphate dehydrogenase 1 chr5:13956879-13959686 REVERSE LENGTH=576 |
| AT3G52300.1 | 0,734825465 | 0,011447638 | ATPQ_ ATPd "ATP synthase D chain_ mitochondrial" chr3:19396689-19398119 FORWARD LENGTH=168 |
| AT5G42740.3 | 1,219753718 | 0,011515334 | no symbol available no full name available chr5:17136269-17140622 FORWARD LENGTH=528 |
| AT3G08580.1 | 1,247096269 | 0,011560479 | AAC1 ADP/ATP carrier 1 chr3:2605706-2607030 REVERSE LENGTH=381 |
| AT3G09200.2 | 1,259936892 | 0,01178331 | no symbol available no full name available chr3:2823364-2825020 REVERSE LENGTH=287 |
| AT5G15530.1 | 1,310733067 | 0,011803755 | BCCP2_ CAC1-B biotin carboxyl carrier protein 2 chr5:5038955-5040437 FORWARD LENGTH=255 |
| AT4G37980.1 | 1,335035604 | 0,01185309 | ELI3-1_ CHR_ ELI3_ ATCAD7_ CAD7 elicitor-activated gene 3-1_ CINNAMALDEHYDE AND HEXENAL |
| AT1G08450.2 | 0,429358547 | 0,011863774 | CRT3_ AtCRT3_ EBS2_ PSL1 A. thaliana calreticulin 3_ EMS-MUTAGENIZED BRI1 SUPPRESSOR 2_ cal |
| AT5G19220.1 | 1,113657435 | 0,012059821 | ADG2_ APL1 ADP glucose pyrophosphorylase large subunit 1_ ADP GLUCOSE PYROPHOSPHORYLASE : |
| AT4G25100.1 | 1,304836428 | 0,012080274 | FSD1_ ATFSD1 ARABIDOPSIS FE SUPEROXIDE DISMUTASE 1_ Fe superoxide dismutase 1 chr4:128846 |
| AT4G39980.1 | 0,709836353 | 0,01214085 | AtDAHP1_ DHS1_ DAHP1 3-DEOXY-D-ARABINO-HEPTULOSONATE-7-PHOSPHATE 1_ 3-deoxy-D-ar |
| AT3G17240.1 | 0,714588869 | 0,012223104 | mtLPD2 lipoamide dehydrogenase 2 chr3:5890278-5892166 REVERSE LENGTH=507 |
| AT1G67700.1 | 0,72926142 | 0,012225525 | HHL1 HYPERSENSITIVE TO HIGH LIGHT 1 chr1:25374295-25375716 FORWARD LENGTH=230 |
| AT3G09630.1 | 0,340008266 | 0,012248186 | SAC56 Suppressor of Acaulis 56 chr3:2953813-2955444 FORWARD LENGTH=406 |
| AT4G28750.1 | 0,862036749 | 0,012264799 | PSAE-1 PSA E1 KNOCKOUT chr4:14202951-14203888 REVERSE LENGTH=143 |
| AT3G08740.1 | 0,376193541 | 0,012392395 | no symbol available no full name available chr3:2654788-2656154 REVERSE LENGTH=236 |
| AT4G37000.1 | 0,780532362 | 0,012447222 | ACD2_ ATRCCR ARABIDOPSIS THALIANA RED CHLOROPHYLL CATABOLITE REDUCTASE_ ACC |
| AT5G15090.1 | 1,169424269 | 0,012540814 | VDAC3_ AtVDAC-3_ ATVDAC3 ARABIDOPSIS THALIANA VOLTAGE DEPENDENT ANION CHANN |
| AT3G13930.1 | 1,149127458 | 0,012587595 | mtE2-2 mitochondrial pyruvate dehydrogenase subunit 2-2 chr3:4596240-4600143 FORWARD LENGTH=539 |
| AT3G16530.1 | 0,580661082 | 0,012638692 | no symbol available no full name available chr3:5624586-5625416 REVERSE LENGTH=276 |
| AT1G09270.1 | 1,203670414 | 0,012763259 | IMPA-4 importin alpha isoform 4 chr1:2994506-2997833 FORWARD LENGTH=538 |
| AT4G23850.1 | 1,387699231 | 0,012775393 | LACS4 long-chain acyl-CoA synthetase 4 chr4:12403720-12408263 REVERSE LENGTH=666 |
| AT1G08200.1 | 0,902665736 | 0,012939405 | AXS2 UDP-D-apiose/UDP-D-xylose synthase 2 chr1:2574259-2576609 REVERSE LENGTH=389 |
| AT3G56940.1 | 0,859239164 | 0,013034437 | CRD1_ ACSF_ CHL27 COPPER RESPONSE DEFECT 1 chr3:21076594-21078269 FORWARD LENGTH=4 |
| AT2G20990.1 | 1,485145326 | 0,013122094 | NTMC2T1.1_ ATSYTA_ SYT1_ NTMC2TYPE1.1_ AtSYT1_ SYTA SYNAPTOTAGMIN 1_ synaptotagmin |
| AT3G49720.1 | 0,433291917 | 0,013217039 | CGR2 chr3:18440192-18441655 REVERSE LENGTH=261 |
| AT4G30270.1 | 0,528839556 | 0,013389233 | MERI5B_ XTH24_ MERI-5_ SEN4 xyloglucan endotransglucosylase/hydrolase 24_ SENESCENCE 4_ MERI |
| AT3G01120.1 | 1,297342267 | 0,013395209 | AtCYS1_ CGS_ AtCGS1_ CGS1_ MTO1 CYSTATHIONINE GAMMA-SYNTHASE 1_ CYSTATHIONINE |

| | | | |
|---|---|---|---|
| AT4G26530.1 | 0,844275069 | 0,013474012 | FBA5_ AtFBA5_ DEG22 fructose-bisphosphate aldolase 5 chr4:13391566-13392937 FORWARD LENGTH= |
| ATCG00650.1 | 0,394751314 | 0,013623453 | RPS18 ribosomal protein S18 chrc:67917-68222 FORWARD LENGTH=101 |
| AT5G03290.1 | 1,456063119 | 0,013708044 | IDH-V isocitrate dehydrogenase V chr5:794043-795939 FORWARD LENGTH=374 |
| AT3G23940.2 | 0,887692425 | 0,013708317 | DHAD Dihydroxyacid dehydratase chr3:8648780-8652323 FORWARD LENGTH=606 |
| AT5G50950.3 | 0,866582073 | 0,013773843 | FUM2 FUMARASE 2 chr5:20731191-20733636 FORWARD LENGTH=317 |
| AT1G65980.1 | 1,173310184 | 0,01381188 | TPX1 thioredoxin-dependent peroxidase 1 chr1:24559524-24560753 REVERSE LENGTH=162 |
| AT1G70890.1 | 1,511890529 | 0,014008374 | MLP43 MLP-like protein 43_ major latex protein like 43 chr1:26725912-26726489 REVERSE LENGTH=158 |
| AT2G30930.1 | 0,610050503 | 0,014186896 | no symbol available no full name available chr2:13162458-13163156 FORWARD LENGTH=164 |
| AT1G23730.1 | 0,533915238 | 0,014196021 | ATBCA3_ BCA3 beta carbonic anhydrase 3_ BETA CARBONIC ANHYDRASE 3 chr1:8395965-8398014 F |
| AT5G26742.1 | 1,212610735 | 0,014196921 | AtRH3_ RH3_ emb1138 embryo defective 1138 chr5:9285540-9288871 REVERSE LENGTH=747 |
| AT5G27670.1 | 0,680546597 | 0,014261292 | HTA7_ h2a.w.7 histone H2A 7 chr5:9792807-9793365 REVERSE LENGTH=150 |
| AT1G54780.1 | 0,705362753 | 0,014406792 | TLP18.3_ AtTLP18.3 thylakoid lumen protein 18.3 chr1:20439533-20440953 FORWARD LENGTH=285 |
| AT3G12780.1 | 1,118595071 | 0,014469863 | PGKp1_ PGK1 phosphoglycerate kinase 1 chr3:4061127-4063140 REVERSE LENGTH=481 |
| AT5G08280.1 | 0,915762688 | 0,014634268 | HEMC_ RUG1 RUGOSA 1_ hydroxymethylbilane synthase chr5:2663763-2665596 REVERSE LENGTH=38 |
| AT3G06050.1 | 0,765452424 | 0,01469009 | PRXIIF_ ATPRXIIF peroxiredoxin IIF_ PEROXIREDOXIN IIF chr3:1826311-1827809 REVERSE LENGTH |
| AT5G56010.1 | 1,356282956 | 0,014890559 | AtHsp90-3_ AtHsp90.3_ Hsp81.3_ HSP81-3 HEAT SHOCK PROTEIN 90-3_ HEAT SHOCK PROTEIN 81. |
| AT4G37990.1 | 1,55882603 | 0,014903663 | ELI3-2_ ATCAD8_ ELI3_ CAD-B2 elicitor-activated gene 3-2_ CINNAMYL-ALCOHOL DEHYDROGENA |
| AT3G19820.1 | 1,527627293 | 0,014915688 | DWF1_ DIM_ DIM1_ CBB1_ EVE1 ENHANCED VERY-LOW-FLUENCE RESPONSES 1_ DIMINUTIA_ |
| AT2G13360.1 | 1,164032239 | 0,014928839 | SGAT_ AGT_ AGT1 ALANINE:GLYOXYLATE AMINOTRANSFERASE 1_ alanine:glyoxylate aminotrans |
| AT1G01470.1 | 2,278337231 | 0,015126456 | LEA14_ LSR3_ AtLEA14_ LEA1 LIGHT STRESS-REGULATED 3_ Arabidopsis thaliana Late Embryogenes |
| AT5G66120.2 | 1,266016825 | 0,015157264 | no symbol available no full name available chr5:26431516-26433649 REVERSE LENGTH=442 |
| AT5G64050.1 | 3,085936375 | 0,015192367 | ATERS_ OVA3_ ERS glutamate tRNA synthetase_ OVULE ABORTION 3 chr5:25630196-25633099 REVER |
| AT3G48730.1 | 1,320727064 | 0,015382566 | GSAM_ GSA2 glutamate-l-semialdehyde aminomutase_ "glutamate-1-semialdehyde 2_1-aminomutase 2" chr3: |
| ATCG00790.1 | 0,319769163 | 0,015409142 | RPL16 ribosomal protein L16 chrc:81189-82652 REVERSE LENGTH=135 |
| AT2G43750.1 | 1,141171328 | 0,015553743 | OASB_ CPACS1_ ACS1_ ATCS-B O-acetylserine (thiol) lyase B_ ARABIDOPSIS THALIANA CYSTEIN S |
| AT1G54040.2 | 0,326346844 | 0,015579024 | TASTY_ ESR_ ESP epithiospecifier protein_ EPITHIOSPECIFYING SENESCENCE REGULATOR chr1:20 |
| AT2G19900.1 | 1,23545655 | 0,015678206 | ATNADP-ME1_ NADP-ME1 NADP-malic enzyme 1_ Arabidopsis thaliana NADP-malic enzyme 1 chr2:8592 |
| AT2G32920.1 | 1,928432789 | 0,015696278 | PDIL2-3_ PDI9_ ATPDIL2-3_ ATPDI9 ARABIDOPSIS THALIANA PROTEIN DISULFIDE ISOMERASE |
| AT5G03340.1 | 1,31529483 | 0,015795127 | AtCDC48C cell division cycle 48C chr5:810091-813133 REVERSE LENGTH=810 |
| AT3G17210.1 | 1,222605288 | 0,015876464 | ATHS1_ HS1 heat stable protein 1_ A. THALIANA HEAT STABLE PROTEIN 1 chr3:5882318-5882896 FO |
| AT2G31570.1 | 0,577194586 | 0,015928414 | ATGPX2_ GPX2_ GPXL2 glutathione peroxidase 2 chr2:13438211-13439775 REVERSE LENGTH=169 |
| AT5G20160.1 | 2,257111795 | 0,015952625 | no symbol available no full name available chr5:6804075-6805102 REVERSE LENGTH=128 |
| AT1G20020.1 | 0,820281546 | 0,016014955 | LFNR2_ FNR2_ ATLFNR2 leaf-type chloroplast-targeted FNR 2_ ferredoxin-NADP(+)-oxidoreductase 2_ LE |
| AT1G73600.1 | 1,368227072 | 0,016042377 | DEG26_ NMT_ AtPMT3_ NMT3 Phosphoethanolamine methyltransferase3 chr1:27670825-27673400 FORW |
| AT3G16480.1 | 1,794743571 | 0,016044411 | MPPalpha mitochondrial processing peptidase alpha subunit chr3:5599906-5602716 FORWARD LENGTH=4 |
| AT3G42050.1 | 1,447391576 | 0,016075429 | VHA-H chr3:14228846-14232228 REVERSE LENGTH=441 |
| AT5G04140.1 | 0,684831526 | 0,016454086 | GLUS_ FD-GOGAT_ GLS1_ GLU1 FERREDOXIN-DEPENDENT GLUTAMATE SYNTHASE 1_ glutamat |
| AT5G41520.1 | 1,404056322 | 0,0164859 | RPS10B ribosomal protein S10e B chr5:16609377-16610583 REVERSE LENGTH=180 |
| AT1G12000.1 | 1,381910648 | 0,01653617 | no symbol available no full name available chr1:4050159-4053727 REVERSE LENGTH=566 |

| | | | |
|---|---|---|---|
| AT2G14260.2 | 1,37230746 | 0,016594706 | PIP_ PAP1 proline iminopeptidase_ prolyl aminopeptidase 1 chr2:6041441-6043475 REVERSE LENGTH=32 |
| AT5G57870.1 | 1,36615607 | 0,016690781 | eIFiso4G1 eukaryotic translation Initiation Factor isoform 4G1 chr5:23439755-23443433 FORWARD LENGT |
| AT4G34670.1 | 0,864726939 | 0,016732448 | no symbol available no full name available chr4:16548724-16550222 FORWARD LENGTH=262 |
| AT4G02520.1 | 0,914255155 | 0,01697864 | ATGSTF2_ GSTF2_ ATPM24.1_ GST2_ ATPM24 glutathione S-transferase PHI 2 chr4:1110673-1111531 R |
| AT1G44575.1 | 1,136394288 | 0,017116887 | CP22_ PSBS_ NPQ4 NONPHOTOCHEMICAL QUENCHING 4_ PHOTOSYSTEM II SUBUNIT S chr1:168 |
| AT4G01310.1 | 0,812514275 | 0,017175825 | PRPL5 plastid ribosomal proteins of the 50S subunit 5 chr4:544166-545480 REVERSE LENGTH=262 |
| AT3G16460.1 | 1,478849922 | 0,017331517 | JAL34 jacalin-related lectin 34 chr3:5593029-5595522 FORWARD LENGTH=705 |
| AT1G62660.1 | 0,682212186 | 0,017439028 | VI1 VACUOLAR INVERTASE 1 chr1:23199949-23203515 FORWARD LENGTH=648 |
| AT3G44320.1 | 1,367426618 | 0,017578186 | NIT3_ AtNIT3 NITRILASE 3_ nitrilase 3 chr3:15993419-15995493 FORWARD LENGTH=346 |
| AT5G66510.1 | 0,831064991 | 0,017585335 | GAMMA CA3 gamma carbonic anhydrase 3 chr5:26550016-26551496 REVERSE LENGTH=258 |
| AT2G42130.2 | 1,150725102 | 0,017636368 | no symbol available no full name available chr2:17566242-17567830 FORWARD LENGTH=269 |
| AT3G22460.1 | 3,008818402 | 0,017672488 | OASA2 O-acetylserine (thiol) lyase (OAS-TL) isoform A2 chr3:7964204-7965751 FORWARD LENGTH=250 |
| AT2G46280.1 | 0,811158681 | 0,017891399 | TIF3I1_ TRIP-1_ TRIP1 TGF-beta receptor interacting protein 1 chr2:19003656-19005393 REVERSE LENG |
| AT4G01050.1 | 1,194686536 | 0,018000297 | TROL thylakoid rhodanese-like chr4:455874-458175 FORWARD LENGTH=466 |
| AT3G48990.1 | 1,333262768 | 0,018058828 | AAE3 ACYL-ACTIVATING ENZYME 3 chr3:18159031-18161294 REVERSE LENGTH=514 |
| AT2G19760.1 | 0,585279563 | 0,018070157 | PFN1_ PRF1 profilin 1_ PROFILIN 1 chr2:8517074-8518067 REVERSE LENGTH=131 |
| AT3G53460.1 | 0,513788834 | 0,018085041 | CP29 chloroplast RNA-binding protein 29 chr3:19819738-19821423 REVERSE LENGTH=342 |
| AT2G33210.2 | 1,116182424 | 0,018226392 | HSP60_ HSP60-2 heat shock protein 60-2 chr2:14075093-14078568 REVERSE LENGTH=580 |
| AT5G57350.3 | 0,31158146 | 0,018482203 | ATAHA3_ HA3_ AHA3 H(+)-ATPase 3_ ARABIDOPSIS THALIANA ARABIDOPSIS H(+)-ATPASE chr5: |
| AT4G27090.1 | 0,52896024 | 0,018644307 | RPL14B chr4:13594104-13595187 REVERSE LENGTH=134 |
| AT2G05840.3 | 1,438653551 | 0,018665602 | PAA2 20S proteasome subunit PAA2 chr2:2234226-2235533 FORWARD LENGTH=205 |
| AT5G06600.2 | 1,241830279 | 0,018788326 | UBP12_ AtUBP12 ubiquitin-specific protease 12 chr5:2019545-2027834 REVERSE LENGTH=1115 |
| AT5G60640.2 | 0,606487569 | 0,019021321 | PDI2_ ATPDIL1-4_ ATPDI2_ PDIL1-4 PROTEIN DISULFIDE ISOMERASE 2_ ARABIDOPSIS THALIAN |
| AT3G55610.1 | 2,780192376 | 0,01903378 | P5CS2 delta 1-pyrroline-5-carboxylate synthase 2 chr3:20624278-20628989 REVERSE LENGTH=726 |
| AT1G61520.1 | 0,904042127 | 0,019049166 | LHCA3 photosystem I light harvesting complex gene 3 chr1:22700152-22701149 FORWARD LENGTH=273 |
| AT2G04030.1 | 1,340882374 | 0,019090429 | CR88_ EMB1956_ AtHsp90.5_ HSP90C_ AtHsp90C_ Hsp88.1_ HSP90.5 HEAT SHOCK PROTEIN 90.5_ E |
| AT2G28815.1 | 1,571869285 | 0,019347138 | no symbol available no full name available chr2:12367001-12368064 REVERSE LENGTH=291 |
| AT2G27530.1 | 0,742981875 | 0,019378575 | PGY1 PIGGYBACK1 chr2:11763443-11764570 REVERSE LENGTH=216 |
| AT4G02840.1 | 0,59275429 | 0,019518168 | SmD1b chr4:1264726-1266253 FORWARD LENGTH=116 |
| AT5G23010.1 | 0,565403146 | 0,019610522 | GSM1_ MAM1_ IMS3 glucosinolate metabolism 1_ 2-ISOPROPYLMALATE SYNTHASE 3_ methylthioalky |
| AT4G23400.1 | 1,302961048 | 0,019612966 | PIP1D_ PIP1;5 plasma membrane intrinsic protein 1;5 chr4:12220792-12222155 FORWARD LENGTH=287 |
| AT1G66410.1 | 0,426664103 | 0,019645672 | CAM4_ ACAM-4 calmodulin 4_ CALMODULIN 4 chr1:24774431-24775785 REVERSE LENGTH=149 |
| AT1G01200.1 | 1,459678324 | 0,019709895 | RABA3_ ATRABA3_ ATRAB-A3 ARABIDOPSIS RAB GTPASE HOMOLOG A3_ RAB GTPase homolog |
| AT2G16360.1 | 1,715519092 | 0,019762698 | no symbol available no full name available chr2:7076713-7077310 REVERSE LENGTH=109 |
| AT4G11600.1 | 1,149836682 | 0,019764465 | GPXL6_ PHGPX_ LSC803_ ATGPX6_ GPX6 glutathione peroxidase 6 chr4:7010021-7011330 REVERSE L |
| AT3G11510.1 | 0,671464095 | 0,019773063 | no symbol available no full name available chr3:3623757-3624866 REVERSE LENGTH=150 |
| AT2G45300.3 | 1,567078293 | 0,019873759 | no symbol available no full name available chr2:18677518-18680118 FORWARD LENGTH=518 |
| AT5G02870.1 | 0,363734694 | 0,01987578 | RPL4 ribosomal large subunit 4 chr5:657830-659526 FORWARD LENGTH=407 |
| AT5G09900.1 | 1,709944083 | 0,019880115 | EMB2107_ MSA_ RPN5A MARIPOSA_ EMBRYO DEFECTIVE 2107_ REGULATORY PARTICLE NON- |

| | | | |
|---|---|---|---|
| AT1G68560.1 | 0,75950726 | 0,019886525 | XYL1_ GH31_ TRG1_ ATXYL1_ AXY3 altered xyloglucan 3_ ALPHA-XYLOSIDASE 1_ alpha-xylosidase |
| AT2G36160.1 | 0,874622573 | 0,019934704 | no symbol available no full name available chr2:15169925-15171159 FORWARD LENGTH=150 |
| AT1G56110.1 | 1,444975374 | 0,020054577 | NOP56 homolog of nucleolar protein NOP56 chr1:20984544-20986893 REVERSE LENGTH=522 |
| AT3G10060.1 | 0,887997217 | 0,020127795 | no symbol available no full name available chr3:3102291-3103801 FORWARD LENGTH=230 |
| AT3G47070.1 | 0,4911978 | 0,020176515 | no symbol available no full name available chr3:17337205-17337507 REVERSE LENGTH=100 |
| AT4G29010.1 | 1,194408477 | 0,020202789 | AIM1 ABNORMAL INFLORESCENCE MERISTEM chr4:14297312-14302016 REVERSE LENGTH=721 |
| AT4G30610.1 | 0,606188327 | 0,020270278 | BRS1_ SCPL24 BRI1 SUPPRESSOR 1_ SERINE CARBOXYPEPTIDASE 24 PRECURSOR chr4:14944219- |
| AT5G52920.1 | 1,420115782 | 0,020282221 | PKP-BETA1_ PKP2_ PKP1 plastidic pyruvate kinase beta subunit 1_ PLASTIDIAL PYRUVATE KINASE 1_ |
| AT2G41220.1 | 0,755702393 | 0,020605639 | GLU2 glutamate synthase 2 chr2:17177934-17188388 FORWARD LENGTH=1629 |
| AT1G18500.1 | 1,478272307 | 0,020708763 | MAML-4_ IPMS1 methylthioalkylmalate synthase-like 4_ ISOPROPYLMALATE SYNTHASE 1 chr1:636934 |
| AT1G29660.1 | 0,764007756 | 0,020812078 | GGL5 chr1:10371955-10373624 FORWARD LENGTH=364 |
| AT2G42130.3 | 1,243689471 | 0,020890364 | no symbol available no full name available chr2:17566389-17567916 FORWARD LENGTH=271 |
| AT5G50920.1 | 1,161728736 | 0,021320082 | DCA1_ CLPC_ ATHSP93-V_ CLPC1_ HSP93-V HEAT SHOCK PROTEIN 93-V_ CLPC homologue 1_ DE- |
| AT2G23350.1 | 1,284579809 | 0,021334916 | PABP4_ PAB4 POLY(A) BINDING PROTEIN 4_ poly(A) binding protein 4 chr2:9943209-9946041 FORWA |
| AT3G14415.1 | 1,170937132 | 0,021438295 | GOX2 glycolate oxidase 2 chr3:4818667-4820748 FORWARD LENGTH=367 |
| AT4G27585.1 | 0,661094049 | 0,021462177 | SLP1_ AtSLP1 stomatin-like protein 1 chr4:13766984-13769832 REVERSE LENGTH=411 |
| AT3G18740.1 | 0,696541658 | 0,021525584 | RPL30C chr3:6453437-6453870 FORWARD LENGTH=112 |
| AT4G24830.1 | 1,449397652 | 0,021528805 | no symbol available no full name available chr4:12793085-12795857 REVERSE LENGTH=494 |
| AT2G03440.1 | 1,215937198 | 0,021753645 | ATNRP1_ NRP1 nodulin-related protein 1 chr2:1039409-1039972 REVERSE LENGTH=187 |
| AT2G26740.1 | 1,538772762 | 0,021908612 | ATSEH_ SEH soluble epoxide hydrolase chr2:11393148-11394257 REVERSE LENGTH=321 |
| AT3G01390.1 | 1,523757581 | 0,021912285 | AVMA10_ VMA10 vacuolar membrane ATPase 10 chr3:150265-150922 REVERSE LENGTH=110 |
| AT4G34230.1 | 1,482683223 | 0,021958682 | CAD-5_ ATCAD5_ CAD5 cinnamyl alcohol dehydrogenase 5 chr4:16386898-16388666 REVERSE LENGTH |
| AT3G16400.1 | 1,128164739 | 0,021994303 | NSP1_ ATNSP1_ ATMLP-470 nitrile specifier protein 1_ NITRILE SPECIFIER PROTEIN 1_ MYROSINAS |
| AT1G11860.1 | 0,840492999 | 0,022194698 | GLDT chr1:4001801-4003245 FORWARD LENGTH=408 |
| AT3G59970.3 | 1,157566352 | 0,022253212 | MTHFR1 methylenetetrahydrofolate reductase 1 chr3:22151303-22154323 FORWARD LENGTH=592 |
| AT3G54890.4 | 0,751151085 | 0,02236486 | LHCA1 photosystem I light harvesting complex gene 1 chr3:20339881-20340922 REVERSE LENGTH=213 |
| AT3G48690.1 | 5,880888605 | 0,022833117 | ATCXE12_ CXE12 ARABIDOPSIS THALIANA CARBOXYESTERASE 12 chr3:18037186-18038160 REVI |
| AT1G62180.1 | 1,592134372 | 0,022864999 | APSR_ PRH43_ PRH_ ATAPR2_ APR2 ADENOSINE-5'-PHOSPHOSULFATE REDUCTASE_ 3'-PHOSPH |
| AT4G35860.2 | 1,463170108 | 0,022897225 | ATRABB1B_ ATGB2_ ATRAB2C_ GB2 GTP-binding 2 chr4:16987118-16988587 REVERSE LENGTH=16 |
| AT4G10480.1 | 0,814338581 | 0,022900818 | no symbol available no full name available chr4:6478089-6479079 REVERSE LENGTH=212 |
| AT4G04910.1 | 5,449641782 | 0,022991074 | NSF N-ethylmaleimide sensitive factor chr4:2489696-2495666 REVERSE LENGTH=742 |
| AT4G24620.1 | 1,296090334 | 0,022991083 | PGI1_ PGI phosphoglucose isomerase 1 chr4:12708972-12712610 REVERSE LENGTH=613 |
| AT1G74910.1 | 1,778496539 | 0,023012556 | KJC1 KONJAC 1 chr1:28135770-28138456 REVERSE LENGTH=415 |
| AT1G24180.1 | 0,775752479 | 0,023034995 | IAR4 IAA-CONJUGATE-RESISTANT 4 chr1:8560777-8563382 REVERSE LENGTH=393 |
| AT2G45470.1 | 0,734633739 | 0,023044481 | AGP8_ FLA8 ARABINOGALACTAN PROTEIN 8_ FASCICLIN-like arabinogalactan protein 8 chr2:187427 |
| AT5G28500.1 | 1,073711501 | 0,023062544 | no symbol available no full name available chr5:10477810-10479114 FORWARD LENGTH=434 |
| AT1G55670.1 | 0,803041075 | 0,023398422 | PSAG photosystem I subunit G chr1:20802874-20803356 REVERSE LENGTH=160 |
| AT3G18890.1 | 1,349073028 | 0,023555594 | Tic62_ AtTic62 translocon at the inner envelope membrane of chloroplasts 62 chr3:6511169-6514729 FORWA |
| AT2G35040.1 | 0,881364921 | 0,023612215 | no symbol available no full name available chr2:14765347-14768269 REVERSE LENGTH=596 |

| | | | |
|---|---|---|---|
| AT3G54050.1 | 1,181853318 | 0,023744384 | HCEF1_ cfbp1 high cyclic electron flow 1 chr3:20016951-20018527 FORWARD LENGTH=417 |
| AT5G37510.1 | 1,337404767 | 0,023929819 | EMB1467_ CI76 embryo defective 1467 chr5:14897490-14900352 FORWARD LENGTH=745 |
| AT1G79210.1 | 1,137043537 | 0,023956243 | no symbol available no full name available chr1:29796286-29798240 REVERSE LENGTH=235 |
| AT4G24770.1 | 1,217168186 | 0,023957972 | ATRBP33_ ATRBP31_ CP31A_ RBP31_ CP33a_ CP31 "ARABIDOPSIS THALIANA RNA BINDING PRO |
| AT1G76010.1 | 0,605310895 | 0,02396805 | ALBA1_ Atalba1_ ALBA4 chr1:28528505-28530488 REVERSE LENGTH=350 |
| AT1G54010.1 | 0,798173332 | 0,023979209 | GLL23 GDSL-like lipase 23 chr1:20158854-20160747 REVERSE LENGTH=386 |
| AT5G03630.1 | 1,559755228 | 0,024080214 | MDAR2 chr5:922378-924616 REVERSE LENGTH=435 |
| AT5G55480.1 | 1,490922016 | 0,024196926 | GPDL1_ GDPDL4_ SVL1 SHV3-like 1_ Glycerophosphodiester phosphodiesterase (GDPD) like 4_ glyceroph |
| AT1G09750.1 | 0,866444022 | 0,024222988 | no symbol available no full name available chr1:3157541-3158960 FORWARD LENGTH=449 |
| AT4G09000.1 | 1,106362811 | 0,024542263 | GRF1_ GF14 CHI GENERAL REGULATORY FACTOR1-G-BOX FACTOR 14-3-3 HOMOLOG ISOFORM |
| AT4G23170.1 | 0,815348148 | 0,024689468 | CRK9_ EP1 CYSTEINE-RICH RLK (RECEPTOR-LIKE PROTEIN KINASE) 9 chr4:12135205-12136002 F( |
| AT1G75350.1 | 0,609329175 | 0,024993018 | emb2184 embryo defective 2184 chr1:28272163-28272687 FORWARD LENGTH=144 |
| AT1G23820.1 | 1,387176772 | 0,025046163 | SPDS1 spermidine synthase 1 chr1:8420410-8422724 FORWARD LENGTH=334 |
| AT1G01320.2 | 1,420646436 | 0,025081863 | REC1_ FLL2 FLOURY ENDOSPERM LIKE 2_ REDUCED CHLOROPLAST COVERAGE chr1:121582-13( |
| AT2G22990.2 | 0,490866969 | 0,025098702 | SNG1_ SCPL8 sinapoylglucose 1_ SERINE CARBOXYPEPTIDASE-LIKE 8 chr2:9787071-9789925 FORW/ |
| AT3G61050.1 | 1,311072056 | 0,025171644 | CLB1_ SYT7_ AtCLB_ NTMC2TYPE4_ NTMC2T4 calcium-dependent lipid-binding protein_ Synaptotagmir |
| AT3G13120.1 | 0,844575763 | 0,025184728 | PRPS10 plastid ribosomal protein of the 30S subunit 10 chr3:4220310-4221526 REVERSE LENGTH=191 |
| AT3G48140.1 | 1,287117227 | 0,025337444 | no symbol available no full name available chr3:17778471-17779299 FORWARD LENGTH=88 |
| AT5G01530.1 | 0,906884793 | 0,025524163 | LHCB4.1 light harvesting complex photosystem II chr5:209084-210243 FORWARD LENGTH=290 |
| AT5G49360.1 | 0,7750893 | 0,02563421 | ATBXL1_ BXL1 beta-xylosidase 1_ BETA-XYLOSIDASE 1 chr5:20012179-20016659 REVERSE LENGTH |
| AT2G28790.1 | 0,561225142 | 0,025732729 | no symbol available no full name available chr2:12354664-12355413 REVERSE LENGTH=249 |
| AT1G03220.1 | 1,534082031 | 0,025805816 | SAP2 secreted aspartic protease 2 chr1:787143-788444 FORWARD LENGTH=433 |
| AT2G45960.2 | 1,169630018 | 0,026015637 | TMP-A_ ATHH2_ PIP1;2_ PIP1B TRANSMEMBRANE PROTEIN A_ plasma membrane intrinsic protein 1B |
| ATCG00770.1 | 0,758070165 | 0,026119971 | RPS8 ribosomal protein S8 chrc:80068-80472 REVERSE LENGTH=134 |
| AT5G56350.1 | 2,427736273 | 0,026256707 | no symbol available no full name available chr5:22820254-22822529 REVERSE LENGTH=498 |
| AT2G47730.1 | 1,267669068 | 0,02636344 | GST6_ GSTF8_ ATGSTF8_ ATGSTF5 glutathione S-transferase phi 8_ Arabidopsis thaliana glutathione S-tra |
| AT1G22780.1 | 0,612323876 | 0,026503024 | RPS18A_ PFL_ PFL1 POINTED FIRST LEAVES_ POINTED FIRST LEAVES 1_ 40S RIBOSOMAL PROT |
| AT2G44610.1 | 1,585907243 | 0,026615549 | ATRAB6A_ RAB6_ RAB6A_ ATRABH1B chr2:18411778-18413883 REVERSE LENGTH=208 |
| AT5G11880.1 | 1,200628148 | 0,026643333 | DAPDC2 meso-diaminopimelate decarboxylase 2 chr5:3827806-3829942 REVERSE LENGTH=489 |
| AT5G39320.1 | 0,800820645 | 0,026661931 | UDG4 UDP-glucose dehydrogenase 4 chr5:15743254-15744696 FORWARD LENGTH=480 |
| AT3G28270.1 | 2,572102761 | 0,026818664 | AFL1 At14a-Like1 chr3:10538725-10539849 FORWARD LENGTH=374 |
| AT4G16660.1 | 1,63198962 | 0,026896457 | HSP70 heat shock protein 70 chr4:9377225-9381232 FORWARD LENGTH=867 |
| AT4G26300.4 | 1,529670113 | 0,026953686 | emb1027 embryo defective 1027 chr4:13308400-13312204 REVERSE LENGTH=590 |
| AT2G30200.1 | 1,298390827 | 0,027241969 | EMB3147_ MCAT_ MCAMT EMBRYO DEFECTIVE 3147_ malonyl CoA-ACP malonyltransferase chr2:128 |
| AT2G33530.1 | 0,689889751 | 0,027312212 | scpl46 serine carboxypeptidase-like 46 chr2:14197866-14200536 REVERSE LENGTH=465 |
| AT2G30950.1 | 2,88298665 | 0,027355281 | FTSH2_ VAR2 VARIEGATED 2 chr2:13174692-13177064 FORWARD LENGTH=695 |
| AT1G08830.1 | 0,31665286 | 0,027373106 | CSD1_ AtSOD1_ SOD1 superoxide dismutase 1_ copper/zinc superoxide dismutase 1 chr1:2827700-2829053 ] |
| AT4G34200.1 | 1,292857543 | 0,02773157 | EDA9_ PGDH1 phosphoglycerate dehydrogenase 1_ embryo sac development arrest 9 chr4:16374041-1637656 |
| AT5G47700.1 | 1,2421331 | 0,028373795 | RPP1C_ RPP1.3 60S acidic ribosomal protein P1-3_ RPP1 co-orthologous gene 3 chr5:19328019-19328724 R] |

| | | | |
|---|---|---|---|
| AT2G45710.1 | 0,860059967 | 0,028401093 | no symbol available no full name available chr2:18831243-18831999 FORWARD LENGTH=84 |
| AT1G11870.6 | 0,409050236 | 0,028505752 | SRS_ OVA7_ ATSRS Seryl-tRNA synthetase_ ovule abortion 7 chr1:4004608-4006556 FORWARD LENGTH |
| AT5G54640.1 | 0,630370762 | 0,028837486 | HTA1_ RAT5_ ATHTA1 histone H2A 1_ RESISTANT TO AGROBACTERIUM TRANSFORMATION 5 ch |
| AT2G01250.1 | 0,737716713 | 0,028845194 | RPL7B chr2:132943-134264 REVERSE LENGTH=242 |
| AT2G19730.1 | 0,540372122 | 0,028957431 | no symbol available no full name available chr2:8511752-8512995 FORWARD LENGTH=143 |
| AT1G50900.1 | 0,815200384 | 0,029166012 | GDC1_ LTD Grana Deficient Chloroplast 1_ LHCP translocation defect chr1:18866272-18867014 FORWARD |
| AT5G23250.1 | 1,340629241 | 0,029255915 | no symbol available no full name available chr5:7830460-7832491 FORWARD LENGTH=341 |
| AT2G38040.1 | 0,847010309 | 0,029588376 | CAC3 acetyl Co-enzyme a carboxylase carboxyltransferase alpha subunit chr2:15917612-15920749 FORWARD |
| AT1G09080.2 | 0,782625164 | 0,029651332 | BIP3 binding protein 3 chr1:2929268-2931804 REVERSE LENGTH=665 |
| AT2G38540.1 | 1,491914809 | 0,029713006 | ATLTP1_ AtLtpI-4_ LTP1_ LP1 ARABIDOPSIS THALIANA LIPID TRANSFER PROTEIN 1_ lipid transfer |
| AT5G17170.1 | 0,660436432 | 0,02984414 | ENH1 enhancer of sos3-1 chr5:5649335-5650975 FORWARD LENGTH=271 |
| AT3G19010.2 | 0,636704901 | 0,03002162 | no symbol available no full name available chr3:6556567-6557862 REVERSE LENGTH=297 |
| AT3G10950.1 | 0,296787331 | 0,030040473 | no symbol available no full name available chr3:3423893-3424566 FORWARD LENGTH=92 |
| AT4G30620.1 | 0,688150228 | 0,030255631 | STCL STIC2 Like chr4:14948724-14950035 REVERSE LENGTH=180 |
| AT3G10670.1 | 1,092709998 | 0,03032217 | ABCI6_ ATNAP7_ NAP7 non-intrinsic ABC protein 7_ ATP-binding cassette I6 chr3:3335325-3337304 REV |
| AT2G44640.1 | 0,905711038 | 0,030376451 | no symbol available no full name available chr2:18417286-18419063 FORWARD LENGTH=451 |
| AT3G20390.1 | 0,879872044 | 0,030428244 | RidA Reactive Intermediate Deaminase A chr3:7110227-7111695 REVERSE LENGTH=187 |
| AT5G19990.1 | 1,169490126 | 0,030483976 | RPT6A_ ATSUG1 regulatory particle triple-A ATPase 6A chr5:6752144-6754918 FORWARD LENGTH=419 |
| AT5G15200.1 | 0,628696685 | 0,030525802 | no symbol available no full name available chr5:4935124-4936334 REVERSE LENGTH=198 |
| AT1G62780.1 | 0,816532181 | 0,030936507 | no symbol available no full name available chr1:23249349-23251066 REVERSE LENGTH=237 |
| AT5G62790.1 | 1,363395407 | 0,031024693 | PDE129_ DXR 1-deoxy-D-xylulose 5-phosphate reductoisomerase_ PIGMENT-DEFECTIVE EMBRYO 129 c |
| AT2G45790.1 | 0,665762241 | 0,031122985 | PMM_ ATPMM phosphomannomutase_ PHOSPHOMANNOMUTASE chr2:18855876-18857753 FORWARD |
| AT2G12550.1 | 0,730911741 | 0,03120899 | NUB1 homolog of human NUB1 chr2:5114881-5118486 FORWARD LENGTH=562 |
| AT2G44120.1 | 0,932130624 | 0,031247 | no symbol available no full name available chr2:18249227-18250402 REVERSE LENGTH=242 |
| AT3G11630.1 | 1,115490442 | 0,031340099 | 2CPA 2-Cys peroxiredoxin A chr3:3672189-3673937 FORWARD LENGTH=266 |
| AT1G08360.1 | 0,863821529 | 0,031371154 | no symbol available no full name available chr1:2636231-2637694 FORWARD LENGTH=216 |
| AT2G34480.1 | 0,167673813 | 0,031401628 | L18aB_ RPL18aB chr2:14532916-14534161 REVERSE LENGTH=178 |
| AT1G09590.1 | 0,135609356 | 0,031853949 | no symbol available no full name available chr1:3106549-3107606 FORWARD LENGTH=164 |
| AT1G58380.1 | 0,838423755 | 0,031988362 | XW6 chr1:21689115-21690085 FORWARD LENGTH=284 |
| AT4G01690.1 | 1,239979344 | 0,032079733 | PPO1_ PPOX_ HEMG1 chr4:729929-732309 FORWARD LENGTH=537 |
| AT3G03250.1 | 1,107033425 | 0,032198316 | AtUGP1_ UGP_ UGP1 UDP-glucose pyrophosphorylase_ UDP-GLUCOSE PYROPHOSPHORYLASE 1 chr3 |
| AT2G40100.1 | 0,526672615 | 0,0322134 | LHCB8_ LHCB4.3 light harvesting complex photosystem II chr2:16745884-16747190 FORWARD LENGTH= |
| AT5G13870.1 | 2,139773401 | 0,032340283 | EXGT-A4_ XTH5 endoxyloglucan transferase A4_ xyloglucan endotransglucosylase/hydrolase 5 chr5:4475089 |
| AT3G26070.1 | 0,216622379 | 0,032341914 | FBN3a FIBRILLIN3a chr3:9526904-9528199 FORWARD LENGTH=242 |
| AT5G39730.1 | 0,589235386 | 0,032352157 | no symbol available no full name available chr5:15901740-15902624 FORWARD LENGTH=172 |
| AT2G27030.1 | 1,205841446 | 0,032573274 | ACAM-2_ CAM5 calmodulin 5 chr2:11532069-11533060 FORWARD LENGTH=149 |
| AT1G78850.1 | 1,790030882 | 0,032765514 | MBL1_ GAL1 apple domain lectin-1_ Mannose Binding Lectin1 chr1:29642072-29643397 REVERSE LENGT |
| AT3G20820.1 | 0,799342734 | 0,032991928 | no symbol available no full name available chr3:7280930-7282027 FORWARD LENGTH=365 |
| AT3G16640.1 | 0,775452687 | 0,033035513 | AtTCTP1_ TCTP1 translationally controlled tumor protein chr3:5669709-5670729 REVERSE LENGTH=168 |

| | | | |
|---|---|---|---|
| AT3G54660.1 | 0,797080266 | 0,033126767 | EMB2360_ MIAO_ ATGR2_ GR2_ GR glutathione reductase_ GRISEA 2 chr3:20230356-20233100 REVERS |
| AT1G77510.1 | 1,330312374 | 0,033145976 | ATPDI6_ PDIL1-2_ ATPDIL1-2_ PDI6 PDI-like 1-2_ PROTEIN DISULFIDE ISOMERASE 6 chr1:2912674 |
| AT1G06430.1 | 0,398052611 | 0,033166241 | FTSH8 FTSH protease 8 chr1:1960214-1962525 REVERSE LENGTH=685 |
| AT5G23820.1 | 0,806416546 | 0,033488815 | ML3 MD2-related lipid recognition 3 chr5:8031386-8032809 FORWARD LENGTH=164 |
| AT1G18540.1 | 0,557550788 | 0,033491443 | no symbol available no full name available chr1:6377448-6378548 REVERSE LENGTH=233 |
| AT5G63980.1 | 0,686157545 | 0,033518984 | ALX8_ SUPO1_ AtFRY1_ HOS2_ ATSAL1_ SAL1_ RON1_ FRY1 HIGH EXPRESSION OF OSMOTICAL |
| AT5G46290.3 | 1,133911955 | 0,033657444 | KASI_ KAS1 3-ketoacyl-acyl carrier protein synthase I_ KETOACYL-ACP SYNTHASE 1 chr5:18774439-18 |
| AT3G24430.1 | 1,3570949 | 0,033786724 | HCF101 HIGH-CHLOROPHYLL-FLUORESCENCE 101 chr3:8868731-8872154 REVERSE LENGTH=532 |
| AT2G25060.1 | 0,669784338 | 0,034126988 | ENODL14_ AtENODL14 early nodulin-like protein 14 chr2:10662308-10662930 FORWARD LENGTH=182 |
| AT1G12410.1 | 1,271561133 | 0,034221222 | EMB3146_ CLPR2_ CLP2_ NCLPP2 EMBRYO DEFECTIVE 3146_ CLP protease proteolytic subunit 2_ NU |
| AT3G15060.1 | 4,766884848 | 0,034281057 | RABA1g_ AtRABA1g RAB GTPase homolog A1G chr3:5069239-5070025 FORWARD LENGTH=217 |
| AT5G66570.1 | 0,884154751 | 0,034437213 | OEE1_ PSBO-1_ MSP-1_ OE33_ OEE33_ PSBO1 PS II OXYGEN-EVOLVING COMPLEX 1_ OXYGEN E |
| AT1G67280.1 | 0,81650451 | 0,034616102 | AtGLYI6 GlyoxalaseI 6 chr1:25188563-25190547 REVERSE LENGTH=350 |
| ATCG00470.1 | 0,636299639 | 0,034761946 | ATPE ATP synthase epsilon chain chrc:52265-52663 REVERSE LENGTH=132 |
| AT5G59890.2 | 0,669220781 | 0,034769781 | ATADF4_ ADF4 actin depolymerizing factor 4 chr5:24123107-24123596 FORWARD LENGTH=132 |
| AT4G22930.1 | 1,520180296 | 0,034969754 | DHOASE_ PYR4 DIHYDROOROTASE_ pyrimidin 4 chr4:12019315-12021200 FORWARD LENGTH=377 |
| AT5G17990.1 | 1,278032184 | 0,035234565 | pat1_ TRP1 tryptophan biosynthesis 1_ PHOSPHORIBOSYLANTHRANILATE TRANSFERASE 1 chr5:595 |
| AT1G09340.1 | 1,129447224 | 0,035482745 | CRB_ CSP41B_ HIP1.3 chloroplast RNA binding_ heteroglycan-interacting protein 1.3_ CHLOROPLAST ST |
| AT1G55260.2 | 0,703952295 | 0,035734468 | LTPG6 glycosylphosphatidylinositol-anchored lipid protein transfer 6 chr1:20614663-20616158 FORWARD L |
| AT3G26520.1 | 1,24534266 | 0,03578427 | GAMMA-TIP2_ SITIP_ TIP2_ TIP1;2 SALT-STRESS INDUCIBLE TONOPLAST INTRINSIC PROTEIN_ |
| AT1G79500.1 | 1,57672724 | 0,035871004 | AtkdsA1_ KDO8PS 3-Deoxy-D-manno-octulosonate 8-phosphate synthase chr1:29903604-29905989 FORWA |
| AT1G70820.1 | 0,84604371 | 0,036280737 | no symbol available no full name available chr1:26705594-26708034 FORWARD LENGTH=615 |
| AT3G48870.1 | 1,139477993 | 0,036354763 | ATCLPC_ HSP93-III_ ClpC2_ ATHSP93-III ClpC2 chr3:18122363-18126008 REVERSE LENGTH=952 |
| AT5G42650.1 | 1,148015556 | 0,036803189 | CYP74A_ AOS_ DDE2 allene oxide synthase_ DELAYED DEHISCENCE 2_ CYTOCHROME P450 74A chr |
| AT5G02960.1 | 0,298876185 | 0,036969777 | no symbol available no full name available chr5:693280-694396 REVERSE LENGTH=142 |
| AT2G29560.1 | 1,687062358 | 0,037078091 | ENOC_ ENO3 cytosolic enolase_ enolase 3 chr2:12646635-12649694 FORWARD LENGTH=475 |
| AT1G63660.2 | 2,471934069 | 0,037083185 | no symbol available no full name available chr1:23604874-23607080 REVERSE LENGTH=434 |
| AT5G11560.1 | 1,819580692 | 0,037121803 | PNET5 chr5:3709734-3713994 REVERSE LENGTH=982 |
| AT5G23120.1 | 0,857284378 | 0,037327475 | HCF136 HIGH CHLOROPHYLL FLUORESCENCE 136 chr5:7778154-7780463 FORWARD LENGTH=403 |
| AT3G49110.1 | 0,896443867 | 0,037527015 | ATPCA_ PRX33_ ATPRX33_ PRXCA peroxidase CA_ PEROXIDASE CA_ PEROXIDASE 33 chr3:182007 |
| AT4G15545.1 | 0,474716707 | 0,037896918 | NAIP1 NAI2-interacting protein 1 chr4:8875932-8877567 FORWARD LENGTH=337 |
| AT5G08670.1 | 1,110504814 | 0,037902107 | no symbol available no full name available chr5:2818395-2821149 REVERSE LENGTH=556 |
| AT5G44500.1 | 0,730316445 | 0,037976007 | no symbol available no full name available chr5:17927505-17928269 FORWARD LENGTH=254 |
| AT3G56190.1 | 1,766942175 | 0,037982116 | ALPHA-SNAP2_ ASNAP alpha-soluble NSF attachment protein 2 chr3:20846119-20848356 REVERSE LEN |
| AT4G03520.1 | 0,782714094 | 0,038023801 | ATHM2_ TRXm2 thioredoxin m2 chr4:1562585-1564055 REVERSE LENGTH=186 |
| AT2G07698.1 | 1,190264697 | 0,039238611 | no symbol available no full name available chr2:3361474-3364028 FORWARD LENGTH=777 |
| AT1G08110.4 | 0,88770391 | 0,03926157 | AtGLYI2_ GLYI2_ GLXI:3 Glyoxalase12_ Glyoxalase I;3 chr1:2535463-2537630 FORWARD LENGTH=23 |
| AT3G56130.1 | 0,705888404 | 0,04018778 | BADC1_ BLP3 biotin/lipoyl attachment domain containing 1_ BCCP-Like Protein 3 chr3:20826852-20829007 |
| AT5G14200.1 | 0,847022677 | 0,040585202 | ATIMD1_ IMD1 isopropylmalate dehydrogenase 1_ ARABIDOPSIS ISOPROPYLMALATE DEHYDROGEN |

| | | | |
|---|---|---|---|
| AT5G47840.2 | 0,65328831 | 0,040693849 | AMK2 adenosine monophosphate kinase chr5:19375488-19378058 FORWARD LENGTH=269 |
| AT1G68010.1 | 1,224612697 | 0,041011101 | HPR_ ATHPR1 hydroxypyruvate reductase chr1:25493418-25495720 FORWARD LENGTH=386 |
| AT1G21440.1 | 0,779536871 | 0,041101376 | no symbol available no full name available chr1:7502325-7504103 REVERSE LENGTH=336 |
| AT5G45750.1 | 3,748495925 | 0,041578499 | RABA1c_ AtRABA1c RAB GTPase homolog A1C chr5:18559318-18560639 FORWARD LENGTH=216 |
| AT1G65930.1 | 1,330105531 | 0,041766006 | cICDH cytosolic NADP+-dependent isocitrate dehydrogenase chr1:24539088-24541861 FORWARD LENGTH |
| AT1G79850.1 | 0,341789717 | 0,041986853 | RPS17_ PDE347_ CS17_ PRPS17 PIGMENT DEFECTIVE 347_ PLASTID RIBOSOMAL SMALL SUBUNI |
| AT3G54400.1 | 0,84489512 | 0,042145739 | no symbol available no full name available chr3:20140291-20142599 REVERSE LENGTH=425 |
| AT1G58080.1 | 0,757607621 | 0,04228955 | ATATP-PRT1_ ATP-PRT1_ HISN1A ATP phosphoribosyl transferase 1 chr1:21504562-21507429 REVERSE |
| AT3G15950.2 | 1,160157514 | 0,042398251 | NAI2 chr3:5397783-5402610 REVERSE LENGTH=734 |
| AT5G58250.1 | 0,785654432 | 0,042929169 | LCAA/YCF54_ EMB3143 low chlorophyll accumulation/hypothetical chloroplast open reading frame 54_ EMI |
| AT5G53480.1 | 1,293109934 | 0,043170213 | AtKPNB1_ KPNB1_ IMB1 homolog of human KPNB1 chr5:21714016-21716709 FORWARD LENGTH=870 |
| AT1G79230.3 | 1,186763466 | 0,04347845 | ATMST1_ STR1_ MST1_ ATRDH1_ ST1 ARABIDOPSIS THALIANA RHODANESE HOMOLOGUE 1_ m |
| AT3G16410.1 | 0,875814133 | 0,043514015 | NSP4 nitrile specifier protein 4 chr3:5572145-5574359 FORWARD LENGTH=619 |
| AT3G53870.1 | 0,880381812 | 0,043700821 | no symbol available no full name available chr3:19951547-19952782 FORWARD LENGTH=249 |
| AT2G22230.1 | 0,877118774 | 0,043760325 | no symbol available no full name available chr2:9450042-9451427 FORWARD LENGTH=220 |
| AT5G27470.1 | 1,180757077 | 0,043922558 | no symbol available no full name available chr5:9695087-9697154 FORWARD LENGTH=451 |
| AT5G27850.1 | 0,457737946 | 0,044413594 | RPL18C chr5:9873169-9874297 FORWARD LENGTH=187 |
| AT4G35100.1 | 0,20152619 | 0,044630379 | PIP3A_ SIMIP_ PIP3_ PIP2;7 plasma membrane intrinsic protein 3_ PLASMA MEMBRANE INTRINSIC PR |
| AT1G34760.2 | 0,714194358 | 0,044902832 | RHS5_ GRF11_ GF14 OMICRON ROOT HAIR SPECIFIC 5_ general regulatory factor 11 chr1:12743981-12 |
| AT1G69740.1 | 1,217173172 | 0,045599177 | HEMB1_ ALAD1 5-aminolevulinic acid dehydratase 1 chr1:26232197-26234713 FORWARD LENGTH=430 |
| AT5G60670.1 | 0,809917144 | 0,045681771 | RPL12C Ribosomal Protein Like 12C chr5:24381066-24381566 REVERSE LENGTH=166 |
| AT1G66580.1 | 0,161836912 | 0,045866577 | SAG24_ RPL10C senescence associated gene 24_ ribosomal protein L10 C chr1:24839208-24840439 FORWA |
| AT5G22440.1 | 1,292346347 | 0,046147677 | no symbol available no full name available chr5:7435328-7436486 REVERSE LENGTH=217 |
| AT3G02780.2 | 0,801431475 | 0,046226379 | IDI2_ IPIAT1_ IPP2 isopentenyl pyrophosphate:dimethylallyl pyrophosphate isomerase 2_ ATISOPENTENYI |
| AT5G48810.1 | 0,683993845 | 0,046558757 | CB5-D_ ATB5-B_ ATCB5-D_ B5 #3_ CYTB5-B cytochrome B5 isoform D_ ARABIDOPSIS CYTOCHROM |
| AT1G06410.1 | 1,361047798 | 0,046569164 | ATTPSA_ TPS7_ ATTPS7 TREHALOSE -6-PHOSPHATASE SYNTHASE S7_ trehalose-phosphatase/syntha |
| AT3G57410.1 | 1,178387024 | 0,046831254 | VLN3_ ATVLN3 villin 3 chr3:21243615-21249809 REVERSE LENGTH=965 |
| AT4G14670.1 | 1,405437046 | 0,0471179 | CLPB2 casein lytic proteinase B2 chr4:8410054-8412557 FORWARD LENGTH=623 |
| AT1G80600.1 | 3,071470283 | 0,047209212 | TUP5_ WIN1 HOPW1-1-interacting 1_ TUMOR PRONE 5 chr1:30298675-30300513 REVERSE LENGTH=4 |
| AT3G25520.2 | 0,785312873 | 0,047491774 | RPL5A_ ATL5_ PGY3_ OLI5 ribosomal protein L5_ RIBOSOMAL PROTEIN L5 A_ PIGGYBACK3_ OLIG |
| AT4G12800.1 | 0,884434579 | 0,04776515 | PSAL photosystem I subunit l chr4:7521469-7522493 FORWARD LENGTH=219 |
| AT5G52520.1 | 1,257947042 | 0,047767479 | ProRS-Org_ OVA6_ PRORS1_ AtProRS-Org PROLYL-TRNA SYNTHETASE 1_ OVULE ABORTION 6_ p |
| AT1G02930.1 | 1,112313804 | 0,047812402 | ATGSTF3_ ATGSTF6_ GST1_ ERD11_ GSTF6_ ATGST1 EARLY RESPONSIVE TO DEHYDRATION 11 |
| AT3G46010.1 | 0,487012624 | 0,048220617 | atadf_ ADF1_ ATADF1 actin depolymerizing factor 1 chr3:16909679-16910678 REVERSE LENGTH=139 |
| AT1G02780.1 | 0,346077116 | 0,048254088 | emb2386 embryo defective 2386 chr1:608120-609391 REVERSE LENGTH=214 |
| AT2G20580.1 | 1,297001801 | 0,048399929 | ATRPN1A_ RPN1A 26S PROTEASOME REGULATORY SUBUNIT S2 1A_ 26S proteasome regulatory sub |
| AT2G17390.1 | 0,810158835 | 0,048746931 | AKR2B ankyrin repeat-containing 2B chr2:7555870-7557743 FORWARD LENGTH=344 |
| AT5G53540.1 | 1,114079393 | 0,048986359 | APP1 chr5:21749561-21751099 REVERSE LENGTH=403 |
| AT4G39800.1 | 1,094119699 | 0,04901504 | ATMIPS1_ ATIPS1_ MIPS1_ MI-1-P SYNTHASE INOSITOL 3-PHOSPHATE SYNTHASE 1_ MYO-INOS |

| | | | |
|---|---|---|---|
| AT3G56650.1 | 0,642739263 | 0,049040466 | PPD6 PsbP-domain protein 6 chr3:20984807-20985913 FORWARD LENGTH=262 |
| AT1G52410.1 | 0,590491336 | 0,049234857 | TSA1_ AtTSA1 TSK-associating protein 1 chr1:19520762-19525361 FORWARD LENGTH=755 |

-BINDING PROTEIN 1_ GLYCINE RICH PROTEIN 7_ "cold_ circadian rhythm_ and rna binding 2"_ GLYCINE-RICH RNA-BINDING PROTEIN 7 chr2:9265477-92663

ILOROPLAST CHAPERONIN 60ALPHA_ ACCUMULATION AND REPLICATION OF CHLOROPLASTS 2_ chaperonin-60alpha chr2:11926603-11929184 FORWARD ck protein 81-2_ heat shock protein 81.2_ HEAT SHOCK PROTEIN 90.2_ HEAT SHOCK PROTEIN 90.2 chr5:22686923-22689433 FORWARD LENGTH=699

TION FACTOR 3B1_ EUKARYOTIC TRANSLATION INITIATION FACTOR 3B1_ EUKARYOTIC TRANSLATION INITIATION FACTOR 3B_ translation initiation fa

:ED STERILITY 3_ NON-PHOTOCHEMICAL QUENCHING 2_ ARABIDOPSIS THALIANA ZEAXANTHIN EPOXIDASE_ LOW EXPRESSION OF OSMOTIC STRES

HALIANA HEME OXYGENASE 1_ HEME OXYGENASE 1_ GENOMES UNCOUPLED 2_ HEME OXYGENASE 6 chr2:11341816-11343394 FORWARD LENGTH=28

:PENDENT GENE SILENCING 1_ MATERNAL EFFECT EMBRYO ARREST 58_ S-ADENOSYL-L-HOMOCYSTEIN HYDROLASE 1 chr4:8054931-8056676 FORWA

osome recycling factor_ "ribosome recycling factor_ chloroplast precursor"_ HIGH CHLOROPHYLL FLUORESCENCE AND PALE GREEN MUTANT 108 chr3:23342861

NATE PROTEIN 70-1_ heat shock cognate protein 70-1_ HEAT SHOCK COGNATE PROTEIN 70_ HEAT SHOCK PROTEIN 70-1 chr5:554055-556334 REVERSE LENG

59_ PLASTID PROTEIN IMPORT 2_ TRANSLOCON AT THE OUTER ENVELOPE MEMBRANE OF CHLOROPLASTS 86_ TRANSLOCON AT THE OUTER ENVEL

N-BINDING PROTEIN 1_ microtubule-destabilizing protein 25_ plasma-membrane associated cation-binding protein 1 chr4:10941593-10943227 FORWARD LENGTH=225

.OPIC DRUG RESISTANCE 8_ ARABIDOPSIS PLEIOTROPIC DRUG RESISTANCE 8_ PENETRATION 3_ ATP-binding cassette G36 chr1:22034661-22039844 FORW

GAMMA-SYNTHASE_ A. thaliana cystathionine gamma-synthetase 1_ METHIONINE OVERACCUMULATION 1 chr3:39234-41865 REVERSE LENGTH=563

SE B2_ ARABIDOPSIS THALIANA CINNAMYL-ALCOHOL DEHYDROGENASE 8_ ELICITOR-ACTIVATED GENE 3 chr4:17855964-17857388 FORWARD LENGT

YNTHASE-B_ CHLOROPLAST O-ACETYLSERINE SULFHYDRYLASE 1_ ARABIDOPSIS CYSTEINE SYNTHASE 1 chr2:18129604-18132322 REVERSE LENGTH=

OADENOSINE-5'-PHOSPHOSULFATE (PAPS) REDUCTASE HOMOLOG 43_ 5'adenylylphosphosulfate reductase 2 chr1:22975794-22977465 REVERSE LENGTH=454

LY RESPONSIVE GENES 2_ suppressors of PIN1 overexpression 1_ ALTERED EXPRESSION OF APX2 8_ FIERY1_ ROTUNDA 1 chr5:25610002-25611802 FORWAR

VOLVING COMPLEX 33 KILODALTON PROTEIN_ OXYGEN EVOLVING ENHANCER PROTEIN 33_ PS II oxygen-evolving complex 1_ MANGANESE-STABILIZI

_ ARABIDOPSIS GLUTATHIONE S-TRANSFERASE 1_ ARABIDOPSIS THALIANA GLUATIONE S-TRANSFERASE F3_ glutathione S-transferase 6_ GLUTATHION

ITOL-1-PHOSPHATE SYNTHASE 1_ myo-inositol-1-phosphate synthase 1_ D-myo-Inositol 3-Phosphate Synthase 1 chr4:18469659-18471893 REVERSE LENGTH=511

SS-RESPONSIVE GENES 6_ ARABIDOPSIS THALIANA ABA DEFICIENT 1_ ZEAXANTHIN EPOXIDASE chr5:26754026-26757090 REVERSE LENGTH=610
