## Supplementary material for "HY5 enhances *Arabidopsis* tolerance to combined high light and heat stress by coordinating photoprotection and hormone signaling": Table S5

**Supplemental Table S5. Differentially accumulated proteins compared to control (P < 0.05) in *hy5-215* leaves subjected to high light stress.**

| Protein ID | Fold Change | p-value | Protein description |
|---|---|---|---|
| AT4G04020.1 | 3,4866997 | 1,56E-07 | FIB_ PGL35_ FIB1a plastoglobulin 35_ fibrillin 1a_ fibrillin chr4:1932161-1933546 FORWARD LENGTH=318 |
| AT5G04140.2 | 0,579945378 | 2,461E-06 | GLUS_ FD-GOGAT_ GLS1_ GLU1 FERREDOXIN-DEPENDENT GLUTAMATE SYNTHASE 1_ glutamate synthase 1_ FE |
| AT5G14740.5 | 2,06083956 | 2,539E-06 | BETA CA2_ CA2_ DEG12_ CA18 BETA CARBONIC ANHYDRASE 2_ CARBONIC ANHYDRASE 18_ carbonic anhydras |
| AT5G54770.1 | 1,798115394 | 5,686E-06 | THI1_ TZ_ THI4 THIAZOLE REQUIRING_ THIAMINE4 chr5:22246634-22247891 FORWARD LENGTH=349 |
| AT5G23860.1 | 0,450767956 | 6,72E-06 | TUB8 tubulin beta 8 chr5:8042962-8044528 FORWARD LENGTH=449 |
| AT5G15520.1 | 3,489137622 | 7,654E-06 | no symbol available no full name available chr5:5037242-5038136 REVERSE LENGTH=143 |
| AT3G04720.1 | 1,147938997 | 8,388E-06 | HEL_ PR-4_ AtPR4_ PR4 HEVEIN-LIKE_ pathogenesis-related 4 chr3:1285691-1286531 REVERSE LENGTH=212 |
| AT2G35635.1 | 4,277924234 | 1,166E-05 | UBQ7_ RUB2 RELATED TO UBIQUITIN 2_ ubiquitin 7 chr2:14981044-14981943 FORWARD LENGTH=154 |
| AT3G08940.2 | 0,683183139 | 1,927E-05 | LHCB4.2 light harvesting complex photosystem II chr3:2717717-2718665 FORWARD LENGTH=287 |
| AT1G24020.1 | 1,459743905 | 1,942E-05 | MLP423 MLP-like protein 423 chr1:8500653-8501458 REVERSE LENGTH=155 |
| AT3G53430.1 | 1,690678328 | 2,239E-05 | no symbol available no full name available chr3:19809895-19810395 REVERSE LENGTH=166 |
| AT1G48600.1 | 1,780581695 | 2,773E-05 | AtPMT2_ AtPMEAMT_ PMEAMT phosphoethanolamine N-methyltransferase_ Phosphoethanolamine methyltransferase2 chr1 |
| AT1G32900.1 | 1,568635624 | 3,621E-05 | GBSS1 granule bound starch synthase 1 chr1:11920582-11923506 REVERSE LENGTH=610 |
| AT3G13470.1 | 1,449286576 | 6,355E-05 | CPNB2_ Cpn60beta2 chaperonin-60beta2 chr3:4389685-4392624 FORWARD LENGTH=596 |
| AT5G01410.1 | 1,537308498 | 6,521E-05 | PDX1_ ATPDX1.3_ ATPDX1_ PDX1.3_ RSR4 REDUCED SUGAR RESPONSE 4_ PYRIDOXINE BIOSYNTHESIS 1.3_ A |
| AT5G19760.1 | 1,266961752 | 7,063E-05 | no symbol available no full name available chr5:6679591-6681845 REVERSE LENGTH=298 |
| AT3G01390.1 | 4,018428783 | 8,031E-05 | AVMA10_ VMA10 vacuolar membrane ATPase 10 chr3:150265-150922 REVERSE LENGTH=110 |
| AT1G73600.1 | 3,279223025 | 8,285E-05 | DEG26_ NMT_ AtPMT3_ NMT3 Phosphoethanolamine methyltransferase3 chr1:27670825-27673400 FORWARD LENGTH= |
| AT2G47470.1 | 1,208674389 | 8,69E-05 | UNE5_ MEE30_ ATPDI11_ PDI11_ ATPDIL2-1 PROTEIN DISULFIDE ISOMERASE 11_ PDI-LIKE 2-1_ ARABIDOPSIS |
| AT1G76160.1 | 2,975954374 | 9,025E-05 | sks5 SKU5 similar 5 chr1:28578211-28581020 REVERSE LENGTH=541 |
| AT5G61170.1 | 0,487089184 | 9,251E-05 | no symbol available no full name available chr5:24611158-24612202 FORWARD LENGTH=143 |
| AT3G25920.1 | 0,849879244 | 9,983E-05 | RPL15 ribosomal protein L15 chr3:9491268-9492558 REVERSE LENGTH=277 |
| AT4G24190.1 | 1,368296923 | 1E-04 | SHD_ HSP90_ HSP90.7_ AtHsp90-7_ AtHsp90.7 SHEPHERD_ HEAT SHOCK PROTEIN 90.7_ HEAT SHOCK PROTEIN 9 |
| AT1G51980.1 | 1,204558813 | 0,0001008 | no symbol available no full name available chr1:19323692-19326771 REVERSE LENGTH=503 |
| AT2G39310.1 | 1,669548378 | 0,0001135 | JAL22 jacalin-related lectin 22 chr2:16414262-16416323 REVERSE LENGTH=458 |
| AT5G26667.2 | 3,794614683 | 0,0001158 | PYR6 chr5:9276659-9278091 FORWARD LENGTH=202 |
| AT2G37190.1 | 0,672120036 | 0,0001264 | no symbol available no full name available chr2:15619559-15620059 REVERSE LENGTH=166 |
| AT5G56500.1 | 0,711671786 | 0,000127 | CPNB3_ Cpn60beta3 chaperonin-60beta3 chr5:22874058-22876966 FORWARD LENGTH=597 |
| AT1G65350.1 | 1,747447702 | 0,0001389 | UBQ13 ubiquitin 13 chr1:24272518-24277275 REVERSE LENGTH=319 |
| AT5G03340.1 | 0,759130339 | 0,0001514 | AtCDC48C cell division cycle 48C chr5:810091-813133 REVERSE LENGTH=810 |
| AT5G13490.1 | 1,58545286 | 0,0001616 | AAC2 ADP/ATP carrier 2 chr5:4336034-4337379 FORWARD LENGTH=385 |
| AT5G61790.1 | 1,263315035 | 0,0001756 | CNX1_ ATCNX1 calnexin 1 chr5:24827394-24829642 REVERSE LENGTH=530 |
| AT3G12110.1 | 2,487990442 | 0,0001788 | ACT11 actin-11 chr3:3858116-3859609 FORWARD LENGTH=377 |
| AT1G79930.1 | 1,264118121 | 0,0001793 | HSP91_ AtHsp70-14 heat shock protein 91 chr1:30063781-30067067 REVERSE LENGTH=831 |
| AT5G09590.1 | 1,435263785 | 0,0001886 | MTHSC70-2_ HSC70-5 mitochondrial HSO70 2_ HEAT SHOCK COGNATE chr5:2975721-2978508 FORWARD LENGTH= |
| AT5G60670.1 | 1,361722053 | 0,0001896 | RPL12C Ribosomal Protein Like 12C chr5:24381066-24381566 REVERSE LENGTH=166 |

| | | | |
|---|---|---|---|
| AT5G43010.1 | 0,558041082 | 0,0001933 | RPT4A regulatory particle triple-A ATPase 4A chr5:17248563-17251014 REVERSE LENGTH=399 |
| AT4G39730.1 | 1,324876523 | 0,0002086 | ATPLAT1_ PLAT1 PLAT domain protein 1_ "Polycystin_ Lipoxygenase_ Alpha-toxin and Triacylglycerol lipase 1" chr4:1843[illegible] |
| AT2G38540.1 | 7,329467661 | 0,0002168 | ATLTP1_ AtLtpI-4_ LTP1_ LP1 ARABIDOPSIS THALIANA LIPID TRANSFER PROTEIN 1_ lipid transfer protein 1_ LIPI[illegible] |
| AT3G08590.1 | 1,528497087 | 0,0002201 | iPGAM2 "2_3-biphosphoglycerate-independent phosphoglycerate mutase 2" chr3:2608683-2611237 REVERSE LENGTH=560 |
| AT4G38510.1 | 1,295177987 | 0,0002493 | AtVAB2_ VAB2 V-ATPase B subunit 2 chr4:18011155-18014789 REVERSE LENGTH=487 |
| AT4G02520.1 | 1,390528726 | 0,0002495 | ATGSTF2_ GSTF2_ ATPM24.1_ GST2_ ATPM24 glutathione S-transferase PHI 2 chr4:1110673-1111531 REVERSE LENG[illegible] |
| AT4G39200.2 | 0,785123125 | 0,0002664 | no symbol available no full name available chr4:18257464-18258464 FORWARD LENGTH=107 |
| AT4G02080.1 | 0,592118988 | 0,0002685 | ATSARA1C_ SAR2_ SAR1C_ ATSAR2_ ASAR1 secretion-associated RAS super family 2 chr4:921554-922547 FORWARD [illegible] |
| AT2G33410.1 | 0,492581385 | 0,0002818 | RBGD2 RNA-binding glycine-rich protein D2 chr2:14156085-14157435 FORWARD LENGTH=404 |
| AT4G01050.1 | 0,755460833 | 0,0003019 | TROL thylakoid rhodanese-like chr4:455874-458175 FORWARD LENGTH=466 |
| AT5G03300.1 | 1,734614656 | 0,0003109 | ADK2 adenosine kinase 2 chr5:796573-798997 FORWARD LENGTH=345 |
| AT2G33380.1 | 3,247194946 | 0,0003116 | AtRD20_ PXG3_ CLO-3_ CLO3_ RD20_ AtCLO3 caleosin 3_ Arabidopsis thaliana caleosin 3_ peroxygenase 3_ RESPONSIV[illegible] |
| AT2G42910.1 | 1,763214188 | 0,0003327 | AtPRS4_ PRS4 phosphoribosyl diphosphate synthase 4 chr2:17856396-17858394 FORWARD LENGTH=337 |
| AT3G23990.1 | 1,39153439 | 0,0003496 | HSP60_ HSP60-3B heat shock protein 60_ HEAT SHOCK PROTEIN 60-3B chr3:8669013-8672278 FORWARD LENGTH=5[illegible] |
| AT2G38230.1 | 1,208700313 | 0,0003719 | ATPDX1.1_ PDX1.1 pyridoxine biosynthesis 1.1_ ARABIDOPSIS THALIANA PYRIDOXINE BIOSYNTHESIS 1.1 chr2:16[illegible] |
| AT1G31330.1 | 0,892999046 | 0,0003947 | PSAF photosystem I subunit F chr1:11215011-11215939 REVERSE LENGTH=221 |
| AT3G16390.1 | 0,831102316 | 0,000395 | NSP3 nitrile specifier protein 3 chr3:5562602-5564356 FORWARD LENGTH=467 |
| AT3G04550.1 | 1,340696312 | 0,0004072 | RAF1 Rubisco accumulation factor 1 chr3:1225961-1227310 FORWARD LENGTH=449 |
| AT5G13630.1 | 0,687740727 | 0,0004629 | ABAR_ CHLH_ GUN5_ CCH_ CCH1 ABA-BINDING PROTEIN_ H SUBUNIT OF MG-CHELATASE_ GENOMES UNCO[illegible] |
| AT5G01600.1 | 13,00394322 | 0,0004772 | FER1_ ATFER1 ferretin 1_ ARABIDOPSIS THALIANA FERRETIN 1 chr5:228149-229594 REVERSE LENGTH=255 |
| AT4G12800.1 | 0,692411429 | 0,0004874 | PSAL photosystem I subunit l chr4:7521469-7522493 FORWARD LENGTH=219 |
| AT1G36280.2 | 0,52522247 | 0,0005125 | no symbol available no full name available chr1:13640600-13642908 FORWARD LENGTH=519 |
| AT1G65980.1 | 1,475439483 | 0,000519 | TPX1 thioredoxin-dependent peroxidase 1 chr1:24559524-24560753 REVERSE LENGTH=162 |
| AT5G66510.1 | 0,347961466 | 0,000522 | GAMMA CA3 gamma carbonic anhydrase 3 chr5:26550016-26551496 REVERSE LENGTH=258 |
| AT2G05100.1 | 0,748878278 | 0,0005572 | LHCB2_ LHCB2.1 LIGHT-HARVESTING CHLOROPHYLL B-BINDING 2_ photosystem II light harvesting complex gene 2.[illegible] |
| AT3G12915.1 | 0,668155395 | 0,0005603 | no symbol available no full name available chr3:4112999-4115708 FORWARD LENGTH=820 |
| AT2G06050.1 | 1,236821727 | 0,0005623 | OPR3_ DDE1_ AtOPR3 DELAYED DEHISCENCE 1_ oxophytodienoate-reductase 3 chr2:2359240-2361971 REVERSE LEN[illegible] |
| AT1G75040.1 | 2,400773796 | 0,0005699 | PR-5_ PR5 pathogenesis-related gene 5 chr1:28177754-28178731 FORWARD LENGTH=239 |
| AT4G33010.1 | 1,271591293 | 0,0005715 | GLDP1_ AtGLDP1 glycine decarboxylase P-protein 1 chr4:15926852-15931150 REVERSE LENGTH=1037 |
| AT5G11170.1 | 1,204207295 | 0,0005726 | UAP56a homolog of human UAP56 a chr5:3553334-3556646 FORWARD LENGTH=427 |
| AT3G19710.1 | 1,633217893 | 0,0006372 | BCAT4 branched-chain aminotransferase4 chr3:6847202-6849429 REVERSE LENGTH=354 |
| AT3G46060.1 | 0,37613852 | 0,0006481 | ARA3_ RAB8A_ RABE1c_ ARA-3_ ATRAB8A_ ATRABE1C RAB GTPase homolog 8A chr3:16917908-16919740 FORWA[illegible] |
| AT5G17770.1 | 1,191350741 | 0,0006871 | CBR_ CBR1_ ATCBR NADH:cytochrome B5 reductase 1_ NADH:CYTOCHROME B5 REDUCTASE 1 chr5:5864543-58664[illegible] |
| AT4G21280.1 | 1,06618535 | 0,0006883 | PSBQ-1_ PSBQA_ PSBQ photosystem II subunit QA_ PHOTOSYSTEM II SUBUNIT Q-1_ PHOTOSYSTEM II SUBUNIT Q[illegible] |
| AT2G29440.1 | 1,381341204 | 0,0007057 | GST24_ ATGSTU6_ GSTU6 glutathione S-transferase tau 6_ GLUTATHIONE S-TRANSFERASE 24 chr2:12620159-126210[illegible] |
| AT5G01530.1 | 0,742875425 | 0,0007206 | LHCB4.1 light harvesting complex photosystem II chr5:209084-210243 FORWARD LENGTH=290 |
| AT1G09310.1 | 0,855004775 | 0,0007258 | SVB2_ SVBL SVB-like chr1:3009109-3009648 FORWARD LENGTH=179 |
| AT1G79550.1 | 1,380915827 | 0,0007261 | PGKc_ PGK_ PGK3 phosphoglycerate kinase_ phosphoglycerate kinase 3 chr1:29924347-29926295 REVERSE LENGTH=401 |
| AT3G44890.1 | 0,714003491 | 0,000739 | RPL9 ribosomal protein L9 chr3:16386505-16387963 FORWARD LENGTH=197 |

| | | | |
|---|---|---|---|
| AT1G31812.1 | 1,58758048 | 0,0007521 | ACBP6_ ACBP_ AtACBP6 acyl-CoA-binding protein 6_ ACYL-COA-BINDING PROTEIN chr1:11411132-11412099 REVEF |
| AT1G74270.1 | 0,739973101 | 0,0007559 | no symbol available no full name available chr1:27928415-27929466 REVERSE LENGTH=112 |
| ATCG00905.1 | 0,751835713 | 0,0007715 | RPS12_ RPS12C RIBOSOMAL PROTEIN S12_ ribosomal protein S12C chrc:97999-98793 REVERSE LENGTH=85 |
| AT5G47700.1 | 1,479221656 | 0,0007855 | RPP1C_ RPP1.3 60S acidic ribosomal protein P1-3_ RPP1 co-orthologous gene 3 chr5:19328019-19328724 REVERSE LENG |
| AT1G30530.1 | 1,209925982 | 0,0007931 | UGT78D1 UDP-glucosyl transferase 78D1 chr1:10814917-10816374 FORWARD LENGTH=453 |
| AT5G17310.2 | 1,37225563 | 0,0008383 | AtUGP2_ UGP2 UDP-GLUCOSE PYROPHOSPHORYLASE 2_ UDP-glucose pyrophosphorylase 2 chr5:5696955-5700845 R |
| AT3G48140.1 | 1,407221258 | 0,0008477 | no symbol available no full name available chr3:17778471-17779299 FORWARD LENGTH=88 |
| AT2G35840.1 | 1,439193124 | 0,000853 | no symbol available no full name available chr2:15053952-15055776 FORWARD LENGTH=422 |
| AT1G75270.1 | 1,636065788 | 0,0008752 | DHAR2 dehydroascorbate reductase 2 chr1:28250255-28251237 REVERSE LENGTH=213 |
| AT3G09440.1 | 1,42096976 | 0,0008906 | no symbol available no full name available chr3:2903434-2905632 REVERSE LENGTH=649 |
| AT4G27440.1 | 0,614196757 | 0,0009018 | PORB protochlorophyllide oxidoreductase B chr4:13725648-13727107 FORWARD LENGTH=401 |
| AT2G14610.1 | 1,600983819 | 0,000943 | PR1_ ATPR1_ AtCAPE9_ PR 1 PATHOGENESIS-RELATED GENE 1_ pathogenesis-related gene 1 chr2:6241944-6242429 F |
| AT5G55070.1 | 1,330938704 | 0,0009494 | E2-OGDH2 chr5:22347637-22350409 FORWARD LENGTH=464 |
| AT1G43560.1 | 2,333283515 | 0,0009574 | Aty2_ ty2 thioredoxin Y2 chr1:16398359-16399828 REVERSE LENGTH=167 |
| AT2G02930.1 | 1,4267578 | 0,0009671 | ATGSTF3_ GST16_ GSTF3 GLUTATHIONE S-TRANSFERASE 16_ glutathione S-transferase F3 chr2:851348-852106 REVI |
| AT3G53990.1 | 0,667566143 | 0,0010298 | AtUSP_ USP17 Universal stress protein chr3:19989658-19991019 REVERSE LENGTH=160 |
| AT3G53230.1 | 1,344612299 | 0,0010498 | AtCDC48B cell division cycle 48B chr3:19723416-19726489 FORWARD LENGTH=815 |
| ATCG00280.1 | 0,608312557 | 0,0011002 | PSBC photosystem II reaction center protein C chrc:33720-35141 FORWARD LENGTH=473 |
| AT3G18740.1 | 0,832460564 | 0,0011305 | RPL30C chr3:6453437-6453870 FORWARD LENGTH=112 |
| AT1G13930.1 | 1,891487235 | 0,0011589 | no symbol available no full name available chr1:4761091-4761558 FORWARD LENGTH=155 |
| AT3G15360.1 | 1,317321757 | 0,0011907 | ATM4_ TRX-M4_ ATHM4 ARABIDOPSIS THIOREDOXIN M-TYPE 4_ thioredoxin M-type 4 chr3:5188448-5189457 FOR |
| AT1G52220.1 | 0,663978986 | 0,0012305 | CURT1C CURVATURE THYLAKOID 1C chr1:19453770-19454605 REVERSE LENGTH=156 |
| ATCG00790.1 | 0,621427625 | 0,001273 | RPL16 ribosomal protein L16 chrc:81189-82652 REVERSE LENGTH=135 |
| AT5G52640.1 | 2,000121319 | 0,0012746 | HSP81-1_ ATHSP90.1_ HSP81.1_ AtHsp90-1_ HSP83_ ATHS83_ HSP90.1 HEAT SHOCK PROTEIN 90-1_ heat shock prote |
| AT4G14040.1 | 1,178488664 | 0,0012956 | EDA38_ SBP2 selenium-binding protein 2_ EMBRYO SAC DEVELOPMENT ARREST 38 chr4:8100691-8102828 REVERSE |
| AT1G53240.1 | 1,292811294 | 0,0013119 | mMDH1 mitochondrial malate dehydrogenase 1 chr1:19854966-19856802 REVERSE LENGTH=341 |
| AT1G58270.1 | 0,74105439 | 0,0013527 | ZW9 chr1:21612394-21614089 REVERSE LENGTH=396 |
| AT5G15970.1 | 5,42982719 | 0,0013839 | AtCor6.6_ KIN2_ COR6.6 COLD-RESPONSIVE 6.6 chr5:5211966-5212441 FORWARD LENGTH=66 |
| AT4G24280.1 | 1,186856613 | 0,0014108 | cpHsc70-1 chloroplast heat shock protein 70-1 chr4:12590094-12593437 FORWARD LENGTH=718 |
| AT1G74470.1 | 0,815161389 | 0,0014184 | no symbol available no full name available chr1:27991248-27992845 FORWARD LENGTH=467 |
| AT2G24020.1 | 1,623540944 | 0,0014373 | STIC2 Suppressor of TIC40 2 chr2:10217869-10219269 REVERSE LENGTH=182 |
| AT5G11670.1 | 1,232313471 | 0,0014431 | NADP-ME2_ ATNADP-ME2 NADP-malic enzyme 2_ Arabidopsis thaliana NADP-malic enzyme 2 chr5:3754456-3758040 FO |
| AT5G27850.1 | 0,711278792 | 0,0015303 | RPL18C chr5:9873169-9874297 FORWARD LENGTH=187 |
| AT4G28440.1 | 1,255751393 | 0,0015313 | no symbol available no full name available chr4:14060054-14060970 FORWARD LENGTH=153 |
| AT1G30360.1 | 0,611313225 | 0,0015371 | ERD4_ OSCA3.1 early-responsive to dehydration 4 chr1:10715892-10718799 FORWARD LENGTH=724 |
| AT3G09790.1 | 0,737107461 | 0,0015762 | UBQ8 ubiquitin 8 chr3:3004111-3006006 REVERSE LENGTH=631 |
| AT5G23140.1 | 0,892620427 | 0,0016056 | CLPP2_ NCLPP7 nuclear-encoded CLP protease P7 chr5:7783811-7784826 FORWARD LENGTH=241 |
| AT4G15210.3 | 1,88049962 | 0,0016475 | AT-BETA-AMY_ ATBETA-AMY_ BMY1_ RAM1_ BAM5 REDUCED BETA AMYLASE 1_ beta-amylase 5_ ARABIDOPS |
| AT2G34430.1 | 0,686683539 | 0,0016744 | DEG11_ LHCB1.4_ LHB1B1 light-harvesting chlorophyll-protein complex II subunit B1_ LIGHT-HARVESTING CHLOROPI |

| | | | |
|---|---|---|---|
| AT1G72150.1 | 0,633512954 | 0,0016892 | PATL1 PATELLIN 1 chr1:27148558-27150652 FORWARD LENGTH=573 |
| AT2G29450.1 | 1,776876216 | 0,0016943 | AT103-1A_ ATGSTU1_ ATGSTU5_ GSTU5 glutathione S-transferase tau 5_ ARABIDOPSIS THALIANA GLUTATHIONE |
| AT3G47470.1 | 0,720752376 | 0,0017183 | LHCA4_ CAB4 light-harvesting chlorophyll-protein complex I subunit A4 chr3:17493622-17494773 REVERSE LENGTH=251 |
| AT3G24170.1 | 1,439297815 | 0,0017265 | ATGR1_ GR1 glutathione-disulfide reductase chr3:8729762-8734115 REVERSE LENGTH=499 |
| AT4G37910.1 | 1,417153816 | 0,0017508 | mtHsc70-1 mitochondrial heat shock protein 70-1 chr4:17825368-17828099 REVERSE LENGTH=682 |
| AT1G48630.1 | 1,186774266 | 0,001759 | RACK1B_ RACK1B_AT receptor for activated C kinase 1B chr1:17981977-17983268 REVERSE LENGTH=326 |
| ATCG00680.1 | 0,617455553 | 0,0017638 | PSBB photosystem II reaction center protein B chrc:72371-73897 FORWARD LENGTH=508 |
| AT1G29660.1 | 0,753349105 | 0,0017927 | GGL5 chr1:10371955-10373624 FORWARD LENGTH=364 |
| AT4G37930.1 | 1,121693802 | 0,0018042 | STM_ SHMT1_ SHM1 SERINE HYDROXYMETHYLTRANSFERASE 1_ serine transhydroxymethyltransferase 1_ SERINE |
| AT5G49910.1 | 1,263046201 | 0,0018731 | HSC70-7_ cpHsc70-2 chloroplast heat shock protein 70-2_ HEAT SHOCK PROTEIN 70-7 chr5:20303470-20306295 FORWA |
| AT1G79040.1 | 0,651962362 | 0,0018738 | PSBR photosystem II subunit R chr1:29736085-29736781 FORWARD LENGTH=140 |
| AT1G11750.1 | 1,27587842 | 0,0018835 | NCLPP1_ CLPP6_ NCLPP6 NUCLEAR-ENCODED CLPP 1_ CLP protease proteolytic subunit 6 chr1:3967609-3969535 FOR |
| AT1G09750.1 | 0,844962596 | 0,001899 | no symbol available no full name available chr1:3157541-3158960 FORWARD LENGTH=449 |
| AT5G17050.1 | 1,397476052 | 0,0019034 | UGT78D2 UDP-glucosyl transferase 78D2 chr5:5607828-5609392 REVERSE LENGTH=460 |
| AT5G24770.1 | 1,203721337 | 0,0019114 | ATVSP2_ VSP2 vegetative storage protein 2 chr5:8500713-8501844 REVERSE LENGTH=265 |
| AT2G47110.1 | 1,280444905 | 0,0019261 | UBI6_ UBQ6_ RPS27aB UBIQUITIN EXTENSION PROTEIN 6_ ubiquitin 6_ Ribosomal protein S27aB chr2:19344701-193 |
| AT2G34480.1 | 0,465290879 | 0,0019661 | L18aB_ RPL18aB chr2:14532916-14534161 REVERSE LENGTH=178 |
| AT4G09010.2 | 0,739870879 | 0,0019692 | APX4_ TL29 ascorbate peroxidase 4_ thylakoid lumen 29 chr4:5777502-5779064 REVERSE LENGTH=284 |
| AT3G47070.1 | 1,870896609 | 0,0020337 | no symbol available no full name available chr3:17337205-17337507 REVERSE LENGTH=100 |
| AT5G28840.1 | 1,129321562 | 0,0020698 | GME GDP-D-mannose 3 chr5:10862472-10864024 REVERSE LENGTH=377 |
| AT2G42540.1 | 1,296030408 | 0,0020901 | COR15_ AtCOR15A_ COR15A cold-regulated 15a chr2:17711241-17711930 REVERSE LENGTH=127 |
| AT5G19940.1 | 0,927673373 | 0,0021816 | FBN6_ PAP8 FIBRILLIN6_ Probable Plastid-Lipid Associated Protein chr5:6739693-6740661 FORWARD LENGTH=239 |
| AT1G05010.1 | 0,696003565 | 0,0022396 | EFE_ ACO4_ EAT1 ethylene forming enzyme_ ethylene-forming enzyme chr1:1431419-1432695 REVERSE LENGTH=323 |
| AT3G48990.1 | 1,217398591 | 0,0022584 | AAE3 ACYL-ACTIVATING ENZYME 3 chr3:18159031-18161294 REVERSE LENGTH=514 |
| AT2G31610.1 | 1,114173842 | 0,0022874 | no symbol available no full name available chr2:13450384-13451669 FORWARD LENGTH=250 |
| AT5G44500.1 | 1,165800149 | 0,002289 | no symbol available no full name available chr5:17927505-17928269 FORWARD LENGTH=254 |
| AT5G19370.1 | 0,737444392 | 0,0023717 | no symbol available no full name available chr5:6524247-6526629 REVERSE LENGTH=299 |
| AT2G18960.1 | 0,802607987 | 0,0023746 | HA1_ OST2_ AHA1_ PMA H(+)-ATPase 1_ PLASMA MEMBRANE PROTON ATPASE_ OPEN STOMATA 2 chr2:822185 |
| AT4G13430.1 | 1,110081924 | 0,0024404 | IIL1_ ATLEUC1 isopropyl malate isomerase large subunit 1 chr4:7804194-7807789 REVERSE LENGTH=509 |
| AT2G36460.1 | 1,162870254 | 0,0024537 | FBA6 fructose-bisphosphate aldolase 6 chr2:15296929-15298387 REVERSE LENGTH=358 |
| AT1G12840.1 | 1,134400271 | 0,0024805 | DET3_ ATVHA-C ARABIDOPSIS THALIANA VACUOLAR ATP SYNTHASE SUBUNIT C_ DE-ETIOLATED 3 chr1:437 |
| ATCG00470.1 | 0,868096521 | 0,0025443 | ATPE ATP synthase epsilon chain chrc:52265-52663 REVERSE LENGTH=132 |
| AT3G01280.1 | 1,108458993 | 0,0025668 | VDAC1_ ATVDAC1 ARABIDOPSIS THALIANA VOLTAGE DEPENDENT ANION CHANNEL 1_ voltage dependent anion |
| AT3G52300.1 | 1,542364442 | 0,0025789 | ATPQ_ ATPd "ATP synthase D chain_ mitochondrial" chr3:19396689-19398119 FORWARD LENGTH=168 |
| AT3G03250.1 | 1,284707903 | 0,002584 | AtUGP1_ UGP_ UGP1 UDP-glucose pyrophosphorylase_ UDP-GLUCOSE PYROPHOSPHORYLASE 1 chr3:749761-754014 |
| AT2G39330.1 | 1,172799778 | 0,0026229 | JAL23 jacalin-related lectin 23 chr2:16419787-16421573 REVERSE LENGTH=459 |
| AT5G04140.1 | 1,124425668 | 0,0026705 | GLUS_ FD-GOGAT_ GLS1_ GLU1 FERREDOXIN-DEPENDENT GLUTAMATE SYNTHASE 1_ glutamate synthase 1_ FE |
| AT1G27400.1 | 0,650229933 | 0,0027231 | no symbol available no full name available chr1:9515230-9516725 FORWARD LENGTH=176 |
| AT1G11840.6 | 1,199224151 | 0,002765 | AtGLYI3_ GLX1_ ATGLX1 glyoxalaseI 3_ glyoxalase I homolog chr1:3995928-3997518 FORWARD LENGTH=322 |

| | | | |
|---|---|---|---|
| AT3G08940.1 | 0,417817678 | 0,0027658 | LHCB4.2 light harvesting complex photosystem II chr3:2717717-2718400 FORWARD LENGTH=227 |
| AT2G31670.1 | 1,216601443 | 0,0028348 | UP3 UP3 chr2:13472699-13473490 REVERSE LENGTH=263 |
| AT1G59900.1 | 1,073501877 | 0,0028349 | AT-E1 ALPHA_ E1 ALPHA_ IAR4L pyruvate dehydrogenase complex E1 alpha subunit_ IAR4-LIKE chr1:22051368-2205366 |
| AT1G18210.1 | 0,614327875 | 0,0028351 | no symbol available no full name available chr1:6268273-6268785 REVERSE LENGTH=170 |
| AT3G59970.1 | 1,308621263 | 0,0028392 | MTHFR1 methylenetetrahydrofolate reductase 1 chr3:22151303-22153412 FORWARD LENGTH=421 |
| AT1G57720.1 | 1,191693944 | 0,0029299 | no symbol available no full name available chr1:21377873-21380114 FORWARD LENGTH=413 |
| AT3G27690.1 | 0,834362889 | 0,0029439 | LHCB2.4_ DEG13_ LHCB2_ LHCB2.3 LIGHT-HARVESTING CHLOROPHYLL B-BINDING 2_ photosystem II light harves |
| AT5G56950.1 | 1,597414317 | 0,002946 | NFA3_ NAP1;3_ NFA03 NUCLEOSOME/CHROMATIN ASSEMBLY FACTOR GROUP A 03_ NUCLEOSOME/CHROMA |
| AT2G21580.2 | 1,164393079 | 0,00295 | no symbol available no full name available chr2:9236629-9237510 FORWARD LENGTH=107 |
| AT5G03940.1 | 0,857672898 | 0,0029534 | SRP54CP_ 54CP_ FFC_ CPSRP54 54 CHLOROPLAST PROTEIN_ chloroplast signal recognition particle 54 kDa subunit_ SI |
| AT1G20630.1 | 1,23322448 | 0,0030035 | CAT1 catalase 1 chr1:7146812-7149609 FORWARD LENGTH=492 |
| AT1G01100.1 | 1,602879733 | 0,0030176 | RPP1A_ RPP1.1 60S acidic ribosomal protein P1-1_ RPP1 co-orthologous gene 1 chr1:50284-50954 REVERSE LENGTH=112 |
| AT4G02770.1 | 0,823827323 | 0,0030182 | PSAD1_ PSAD-1 photosystem I subunit D-1 chr4:1229247-1229873 REVERSE LENGTH=208 |
| AT3G25230.1 | 1,50966351 | 0,0030279 | FKBP62_ ROF1_ ATFKBP62 rotamase FKBP 1_ FK506 BINDING PROTEIN 62 chr3:9188257-9191137 FORWARD LENG |
| AT1G62820.1 | 1,40944844 | 0,0030701 | CML14 chr1:23263822-23264268 REVERSE LENGTH=148 |
| AT3G12580.1 | 1,68489185 | 0,0031049 | HSP70_ ATHSP70_ HSC70-4 ARABIDOPSIS HEAT SHOCK PROTEIN 70_ heat shock protein 70 chr3:3991487-3993689 R |
| AT5G57870.1 | 1,131814103 | 0,0031092 | eIFiso4G1 eukaryotic translation Initiation Factor isoform 4G1 chr5:23439755-23443433 FORWARD LENGTH=780 |
| AT1G04410.1 | 1,211906219 | 0,0032084 | c-NAD-MDH1 cytosolic-NAD-dependent malate dehydrogenase 1 chr1:1189418-1191267 REVERSE LENGTH=332 |
| AT4G21960.1 | 0,498305996 | 0,0032093 | PRXR1 chr4:11646613-11648312 REVERSE LENGTH=330 |
| ATCG00270.1 | 0,663139248 | 0,0032811 | PSBD photosystem II reaction center protein D chrc:32711-33772 FORWARD LENGTH=353 |
| AT3G15020.1 | 0,754848079 | 0,0034105 | mMDH2 mitochondrial malate dehydrogenase 2 chr3:5056139-5057941 FORWARD LENGTH=341 |
| AT5G42790.1 | 1,29879047 | 0,0034326 | ARS5_ PAF1_ ATPSM30 ARSENIC TOLERANCE 5_ proteasome alpha subunit F1 chr5:17159270-17160975 REVERSE LEN |
| AT5G57290.2 | 1,12401194 | 0,0034688 | P3B_ AtP3B ribosomal P3 protein B chr5:23207089-23207835 REVERSE LENGTH=89 |
| AT1G32990.1 | 0,903704326 | 0,00352 | PRPL11 plastid ribosomal protein l11 chr1:11955827-11957139 FORWARD LENGTH=222 |
| AT5G03630.1 | 1,281674313 | 0,0035774 | MDAR2 chr5:922378-924616 REVERSE LENGTH=435 |
| AT3G27830.1 | 1,068605039 | 0,0036191 | RPL12-A_ RPL12 ribosomal protein L12-A_ RIBOSOMAL PROTEIN L12 chr3:10318576-10319151 FORWARD LENGTH= |
| AT5G45280.1 | 0,484041525 | 0,0036505 | PAE11 pectin acetylesterase 11 chr5:18346862-18349432 FORWARD LENGTH=370 |
| AT3G26740.1 | 0,645807806 | 0,0037149 | CCL CCR-like chr3:9827868-9828461 FORWARD LENGTH=141 |
| AT3G20390.1 | 1,131175329 | 0,0037215 | RidA Reactive Intermediate Deaminase A chr3:7110227-7111695 REVERSE LENGTH=187 |
| ATCG00730.1 | 0,72081634 | 0,0037346 | PETD photosynthetic electron transfer D chrc:76481-77672 FORWARD LENGTH=160 |
| AT1G21750.1 | 1,247269132 | 0,0038672 | PDIL1-1_ ATPDI5_ PDI5_ ATPDIL1-1 PDI-like 1-1_ ARABIDOPSIS THALIANA PROTEIN DISULFIDE ISOMERASE 5_ |
| AT4G17090.1 | 2,214129315 | 0,0039249 | CT-BMY_ BMY8_ AtBAM3_ BAM3 BETA-AMYLASE 8_ BETA-AMYLASE 3_ chloroplast beta-amylase chr4:9605266-96 |
| AT3G42050.1 | 0,730554563 | 0,0040095 | VHA-H chr3:14228846-14232228 REVERSE LENGTH=441 |
| AT1G77510.1 | 1,368414425 | 0,0040359 | ATPDI6_ PDIL1-2_ ATPDIL1-2_ PDI6 PDI-like 1-2_ PROTEIN DISULFIDE ISOMERASE 6 chr1:29126742-29129433 FOR |
| AT1G15820.1 | 0,76982017 | 0,0041902 | CP24_ LHCB6 light harvesting complex photosystem II subunit 6 chr1:5446685-5447676 REVERSE LENGTH=258 |
| AT2G26150.1 | 2,22992723 | 0,0042013 | ATHSFA2_ HSFA2 heat shock transcription factor A2 chr2:11135856-11137217 FORWARD LENGTH=345 |
| AT4G21990.1 | 1,810776619 | 0,0042953 | APR3_ PRH26_ PRH-26_ ATAPR3 PAPS REDUCTASE HOMOLOG 26_ APS reductase 3 chr4:11657284-11658973 REVER |
| AT5G65020.1 | 1,191568324 | 0,0044364 | ANNAT2_ AtANN2 annexin 2 chr5:25973915-25975554 FORWARD LENGTH=317 |
| AT4G30530.1 | 1,186406126 | 0,0044623 | GGP1 gamma-glutamyl peptidase 1 chr4:14920605-14922286 FORWARD LENGTH=250 |

| | | | |
|---|---|---|---|
| AT1G58080.1 | 1,407669338 | 0,0045756 | ATATP-PRT1_ ATP-PRT1_ HISN1A ATP phosphoribosyl transferase 1 chr1:21504562-21507429 REVERSE LENGTH=411 |
| AT1G27950.1 | 2,181011907 | 0,0045925 | LTPG1 glycosylphosphatidylinositol-anchored lipid protein transfer 1 chr1:9740740-9741991 FORWARD LENGTH=193 |
| AT5G42020.2 | 1,303614239 | 0,0046 | BIP2_ BIP luminal binding protein chr5:16807697-16810480 REVERSE LENGTH=613 |
| AT1G07320.3 | 0,894049141 | 0,0047478 | RPL4_ PRPL4_ EMB2784 plastid ribosomal protein L4_ ribosomal protein L4_ EMBRYO DEFECTIVE 2784 chr1:2249190-2[illegible] |
| AT3G07770.1 | 1,26731145 | 0,0047605 | Hsp89.1_ AtHsp90.6_ AtHsp90-6 HEAT SHOCK PROTEIN 90.6_ HEAT SHOCK PROTEIN 89.1_ HEAT SHOCK PROTEI[illegible] |
| AT2G05840.3 | 1,35919396 | 0,0048493 | PAA2 20S proteasome subunit PAA2 chr2:2234226-2235533 FORWARD LENGTH=205 |
| AT1G67430.2 | 1,76862232 | 0,0048544 | no symbol available no full name available chr1:25262209-25263627 FORWARD LENGTH=131 |
| AT5G10360.1 | 0,643152418 | 0,0049059 | RPS6B_ EMB3010 Ribosomal protein small subunit 6b_ embryo defective 3010 chr5:3258734-3260142 REVERSE LENGTH=[illegible] |
| AT2G20610.1 | 1,271059039 | 0,0049865 | RTY1_ RTY_ HLS3_ ALF1_ SUR1 ABERRANT LATERAL ROOT FORMATION 1_ SUPERROOT 1_ ROOTY 1_ ROOTY[illegible] |
| AT2G21410.1 | 0,548498013 | 0,005004 | VHA-A2 vacuolar proton ATPase A2 chr2:9162703-9168141 FORWARD LENGTH=821 |
| AT2G21170.1 | 1,10910201 | 0,0050337 | TIM_ PDTPI triosephosphate isomerase_ PLASTID ISOFORM TRIOSE PHOSPHATE ISOMERASE chr2:9071047-9073106 [illegible] |
| AT3G53870.1 | 0,880311089 | 0,0050772 | no symbol available no full name available chr3:19951547-19952782 FORWARD LENGTH=249 |
| AT3G28220.1 | 0,810017458 | 0,0050796 | no symbol available no full name available chr3:10524420-10526497 FORWARD LENGTH=370 |
| AT2G28000.1 | 1,213019232 | 0,0051014 | ARC2_ CH-CPN60A_ SLP_ CPN60A_ Cpn60alpha1_ CPNA1 SCHLEPPERLESS_ chaperonin-60alpha1_ CHLOROPLAST [illegible] |
| AT5G20010.1 | 1,128725964 | 0,0052203 | RAN-1_ ATRAN1_ RAN1 RAS-RELATED NUCLEAR PROTEIN_ ARABIDOPSIS THALIANA RAS-RELATED NUCLEA[illegible] |
| AT5G57350.3 | 0,540259504 | 0,0052257 | ATAHA3_ HA3_ AHA3 H(+)-ATPase 3_ ARABIDOPSIS THALIANA ARABIDOPSIS H(+)-ATPASE chr5:23231208-23234[illegible] |
| AT5G45930.1 | 1,192950292 | 0,005227 | CHLI2_ CHL I2_ CHLI-2 magnesium chelatase i2 chr5:18628095-18629565 FORWARD LENGTH=418 |
| AT2G46820.1 | 0,766789071 | 0,0052964 | PTAC8_ PSI-P_ PSAP_ CURT1B_ TMP14 CURVATURE THYLAKOID 1B_ PLASTID TRANSCRIPTIONALLY ACTIVE [illegible] |
| AT3G57260.1 | 1,462097205 | 0,0053278 | AtBG2_ PR2_ GNS2_ AtPR2_ BG2_ PR-2_ BGL2 "beta-1_3-glucanase 2"_ PATHOGENESIS-RELATED PROTEIN 2_ "&#[illegible] |
| AT3G46970.1 | 1,139223473 | 0,0053336 | ATPHS2_ PHS2 alpha-glucan phosphorylase 2_ Arabidopsis thaliana alpha-glucan phosphorylase 2 chr3:17301625-17306111 [illegible] |
| AT5G13850.1 | 0,837714426 | 0,0054405 | NACA3 nascent polypeptide-associated complex subunit alpha-like protein 3 chr5:4471361-4472676 FORWARD LENGTH=2[illegible] |
| AT1G75780.1 | 0,649091883 | 0,0054623 | TUB1 tubulin beta-1 chain chr1:28451378-28453602 REVERSE LENGTH=447 |
| AT1G23190.1 | 1,257400502 | 0,0054727 | PGM3 phosphoglucomutase 3 chr1:8219946-8224186 FORWARD LENGTH=583 |
| AT3G52880.1 | 1,227356111 | 0,0054937 | ATMDAR1_ MDAR1 monodehydroascorbate reductase 1 chr3:19601477-19604366 REVERSE LENGTH=434 |
| AT3G61470.1 | 0,847858411 | 0,0055106 | LHCA2 photosystem I light harvesting complex gene 2 chr3:22745736-22747032 FORWARD LENGTH=257 |
| AT2G22170.1 | 0,332418142 | 0,0055734 | PLAT2 PLAT domain protein 2 chr2:9427010-9427742 REVERSE LENGTH=183 |
| AT1G15690.1 | 0,696398433 | 0,005619 | AtVHP1;1_ AtAVP1_ ATAVP3_ AVP-3_ AVP1_ FUGU5_ VHP1 ARABIDOPSIS THALIANA V-PPASE 3_ FUGU 5 chr1:5[illegible] |
| AT3G55610.1 | 1,635541089 | 0,0056565 | P5CS2 delta 1-pyrroline-5-carboxylate synthase 2 chr3:20624278-20628989 REVERSE LENGTH=726 |
| AT2G41100.5 | 0,449899605 | 0,0057793 | TCH3_ CML12_ ATCAL4 ARABIDOPSIS THALIANA CALMODULIN LIKE 4_ calmodulin-like 12_ TOUCH 3 chr2:17138[illegible] |
| AT1G80380.4 | 1,052093643 | 0,0059146 | GLYK glycerate kinase chr1:30217332-30219784 FORWARD LENGTH=450 |
| AT1G58380.1 | 1,096930958 | 0,0059385 | XW6 chr1:21689115-21690085 FORWARD LENGTH=284 |
| AT1G66270.2 | 1,812392733 | 0,0059405 | BGLU21 chr1:24700110-24702995 REVERSE LENGTH=522 |
| AT3G02730.1 | 1,16439972 | 0,0059661 | TRXF1_ ATF1 thioredoxin F-type 1 chr3:588570-589591 REVERSE LENGTH=178 |
| AT1G50670.1 | 1,197846117 | 0,0060581 | OTU2 ovarian tumor domain (OTU)-containing DUB (deubiquitilating enzyme) 2 chr1:18775086-18776552 REVERSE LENG[illegible] |
| AT5G54270.1 | 0,764951029 | 0,0061473 | LHCB3_ LHCB31 light-harvesting chlorophyll B-binding protein 3 chr5:22038424-22039383 FORWARD LENGTH=265 |
| AT3G56940.1 | 0,534532984 | 0,0063433 | CRD1_ ACSF_ CHL27 COPPER RESPONSE DEFECT 1 chr3:21076594-21078269 FORWARD LENGTH=409 |
| AT1G02780.1 | 0,616915235 | 0,006561 | emb2386 embryo defective 2386 chr1:608120-609391 REVERSE LENGTH=214 |
| ATCG00350.1 | 0,684099305 | 0,0065928 | PSAA chrc:39605-41857 REVERSE LENGTH=750 |
| AT3G44860.1 | 1,37247093 | 0,0066259 | FAMT farnesoic acid carboxyl-O-methyltransferase chr3:16379689-16380939 FORWARD LENGTH=348 |

| | | | |
|---|---|---|---|
| AT3G09350.1 | 2,182240085 | 0,0066487 | Fes1A Fes1A chr3:2871216-2873109 FORWARD LENGTH=363 |
| AT2G27860.1 | 0,729467473 | 0,0066571 | AXS1 UDP-D-apiose/UDP-D-xylose synthase 1 chr2:11864684-11866843 REVERSE LENGTH=389 |
| AT4G34620.1 | 0,840584373 | 0,0067605 | SSR16 small subunit ribosomal protein 16 chr4:16535084-16536092 REVERSE LENGTH=113 |
| AT4G38740.1 | 1,256147292 | 0,0068518 | ROC1 rotamase CYP 1 chr4:18083620-18084138 REVERSE LENGTH=172 |
| AT1G71500.1 | 0,666237777 | 0,0068947 | PSB33_ LIL8 PhotoSystem B protein 33_ Light-harvesting-like 8 chr1:26936084-26937331 FORWARD LENGTH=287 |
| AT1G55260.2 | 1,745910567 | 0,0069887 | LTPG6 glycosylphosphatidylinositol-anchored lipid protein transfer 6 chr1:20614663-20616158 FORWARD LENGTH=224 |
| AT1G68010.1 | 1,20447348 | 0,0070274 | HPR_ ATHPR1 hydroxypyruvate reductase chr1:25493418-25495720 FORWARD LENGTH=386 |
| AT4G02530.1 | 1,221554779 | 0,0070748 | MPH2 MAINTENANCE OF PHOTOSYSTEM II UNDER HIGH LIGHT 2 chr4:1112335-1114005 REVERSE LENGTH=216 |
| AT4G05530.1 | 1,219364585 | 0,0071718 | SDRA_ IBR1 indole-3-butyric acid response 1_ SHORT-CHAIN DEHYDROGENASE/REDUCTASE A chr4:2816462-281807 |
| AT2G03440.1 | 1,562704857 | 0,0071857 | ATNRP1_ NRP1 nodulin-related protein 1 chr2:1039409-1039972 REVERSE LENGTH=187 |
| AT5G02790.1 | 1,294622963 | 0,0073958 | GSTL3 Glutathione transferase L3 chr5:632877-634858 FORWARD LENGTH=235 |
| AT5G12860.2 | 0,793150603 | 0,0074042 | DiT1 dicarboxylate transporter 1 chr5:4059850-4061919 REVERSE LENGTH=556 |
| AT1G02560.1 | 1,134740735 | 0,0074647 | NCLPP1_ NCLPP5_ CLPP5 NUCLEAR CLPP 5_ NUCLEAR-ENCODED CLPP 1_ nuclear encoded CLP protease 5 chr1:538 |
| AT1G41880.1 | 0,80199827 | 0,0075955 | no symbol available no full name available chr1:15651585-15652427 REVERSE LENGTH=111 |
| AT1G59870.1 | 0,738950285 | 0,0076373 | ABCG36_ ATABCG36_ PEN3_ PDR8_ ATPDR8 Arabidopsis thaliana ATP-binding cassette G36_ PLEIOTROPIC DRUG RE |
| AT5G25880.3 | 1,63366645 | 0,0076852 | ATNADP-ME3_ NADP-ME3 NADP-malic enzyme 3_ Arabidopsis thaliana NADP-malic enzyme 3 chr5:9024549-9027498 FO |
| AT3G09200.2 | 1,426713497 | 0,0078465 | no symbol available no full name available chr3:2823364-2825020 REVERSE LENGTH=287 |
| AT5G19550.1 | 1,463801852 | 0,0079101 | AAT2_ ASP2 ASPARTATE AMINOTRANSFERASE 2_ aspartate aminotransferase 2 chr5:6598201-6601597 FORWARD LE |
| AT1G51760.1 | 1,251457523 | 0,0079245 | IAR3_ JR3 IAA-ALANINE RESISTANT 3_ JASMONIC ACID RESPONSIVE 3 chr1:19199562-19201424 FORWARD LEN |
| AT2G26080.1 | 1,21708465 | 0,0081547 | GLDP2_ AtGLDP2 glycine decarboxylase P-protein 2 chr2:11109330-11113786 REVERSE LENGTH=1044 |
| AT4G36130.1 | 0,258207012 | 0,0081949 | no symbol available no full name available chr4:17097613-17098656 FORWARD LENGTH=258 |
| AT1G55670.1 | 0,623084431 | 0,0082974 | PSAG photosystem I subunit G chr1:20802874-20803356 REVERSE LENGTH=160 |
| ATCG00540.1 | 0,856631879 | 0,0084503 | PETA photosynthetic electron transfer A chrc:61657-62619 FORWARD LENGTH=320 |
| AT4G32260.1 | 1,162608022 | 0,0085074 | PDE334 PIGMENT DEFECTIVE 334 chr4:15573859-15574586 REVERSE LENGTH=219 |
| AT3G11250.1 | 0,59445624 | 0,0086288 | no symbol available no full name available chr3:3521453-3522826 FORWARD LENGTH=323 |
| AT3G01910.1 | 1,353885585 | 0,008685 | AtSO_ AT-SO_ SOX sulfite oxidase chr3:314919-317274 REVERSE LENGTH=393 |
| AT3G12290.1 | 1,270158166 | 0,0087016 | MTHFD1 methylenetetrahydrofolate dehydrogenase/methenyltetrahydrofolate cyclohydrolase chr3:3919591-3921326 FORWAF |
| AT4G11600.1 | 1,225433691 | 0,0087917 | GPXL6_ PHGPX_ LSC803_ ATGPX6_ GPX6 glutathione peroxidase 6 chr4:7010021-7011330 REVERSE LENGTH=232 |
| AT2G40610.1 | 0,461472678 | 0,008813 | ATHEXP ALPHA 1.11_ EXP8_ ATEXPA8_ ATEXP8_ EXPA8 expansin A8 chr2:16949121-16950472 REVERSE LENGTH= |
| ATCG00020.1 | 0,673350023 | 0,0088211 | PSBA photosystem II reaction center protein A chrc:383-1444 REVERSE LENGTH=353 |
| AT1G07890.1 | 1,205772796 | 0,0088984 | MEE6_ ATAPX01_ ATAPX1_ CS1_ APX1 ascorbate peroxidase 1_ maternal effect embryo arrest 6 chr1:2438005-2439435 F( |
| AT5G28540.1 | 1,220826263 | 0,0089402 | BIP1 chr5:10540665-10543274 REVERSE LENGTH=669 |
| AT4G37925.1 | 0,814126159 | 0,0090309 | NDH-M_ NdhM subunit NDH-M of NAD(P)H:plastoquinone dehydrogenase complex_ NADH dehydrogenase-like complex M |
| AT3G10350.1 | 0,794862575 | 0,0090404 | AtGET3b_ GET3b Guided Entry of Tail-anchored proteins 3b chr3:3208310-3210678 FORWARD LENGTH=411 |
| AT3G44300.1 | 1,250844217 | 0,0090618 | AtNIT2_ NIT2 nitrilase 2 chr3:15983351-15985172 FORWARD LENGTH=339 |
| AT1G66280.1 | 0,723323653 | 0,0090848 | BGLU22 chr1:24706759-24709737 REVERSE LENGTH=524 |
| AT1G14250.1 | 0,873570454 | 0,0091456 | no symbol available no full name available chr1:4868675-4871203 FORWARD LENGTH=488 |
| AT5G60360.2 | 0,468545122 | 0,0092626 | AALP_ SAG2_ ALP aleurain-like protease_ SENESCENCE ASSOCIATED GENE2 chr5:24280044-24282152 FORWARD LE |
| AT1G75330.1 | 0,334382347 | 0,009298 | OTC ornithine carbamoyltransferase chr1:28266457-28268383 REVERSE LENGTH=375 |

| | | | |
|---|---|---|---|
| AT1G62180.1 | 1,450820869 | 0,009446 | APSR_ PRH43_ PRH_ ATAPR2_ APR2 ADENOSINE-5'-PHOSPHOSULFATE REDUCTASE_ 3'-PHOSPHOADENOSINE-: |
| AT3G07110.1 | 0,577216524 | 0,0094797 | no symbol available no full name available chr3:2252092-2253332 FORWARD LENGTH=206 |
| ATCG00750.1 | 0,642387888 | 0,0094834 | RPS11 ribosomal protein S11 chrc:78960-79376 REVERSE LENGTH=138 |
| AT3G16420.1 | 0,839774178 | 0,0095191 | PBPI_ JAL30_ PBP1 PYK10-binding protein 1_ JACALIN-RELATED LECTIN 30 chr3:5579560-5580674 FORWARD LENC |
| AT3G02880.1 | 1,625712405 | 0,0095367 | KIN7 Kinase 7 chr3:634819-636982 FORWARD LENGTH=627 |
| AT4G37800.1 | 0,841360515 | 0,0097909 | XTH7 xyloglucan endotransglucosylase/hydrolase 7 chr4:17775703-17777372 REVERSE LENGTH=293 |
| AT4G25200.1 | 0,744935436 | 0,0098105 | ATHSP23.6-MITO_ HSP23.6-MITO mitochondrion-localized small heat shock protein 23.6 chr4:12917089-12917858 FORWA |
| AT5G56030.1 | 1,458980785 | 0,0099084 | HSP81-2_ HSP90.2_ AtHsp90.2_ ERD8_ HSP81.2 EARLY-RESPONSIVE TO DEHYDRATION 8_ heat shock protein 81-2_ |
| AT1G09795.1 | 1,210740877 | 0,0100046 | ATATP-PRT2_ HISN1B_ ATP-PRT2 ATP phosphoribosyl transferase 2 chr1:3173588-3176690 FORWARD LENGTH=413 |
| AT3G54400.1 | 0,905064602 | 0,0101722 | no symbol available no full name available chr3:20140291-20142599 REVERSE LENGTH=425 |
| AT4G09040.1 | 0,706724595 | 0,0102313 | CP33C chr4:5795075-5797315 REVERSE LENGTH=304 |
| AT4G30270.1 | 0,684710969 | 0,0104109 | MERI5B_ XTH24_ MERI-5_ SEN4 xyloglucan endotransglucosylase/hydrolase 24_ SENESCENCE 4_ MERISTEM 5_ meriste |
| AT1G06000.1 | 1,286531762 | 0,0104124 | UGT89C1 chr1:1820495-1821802 REVERSE LENGTH=435 |
| AT3G53610.1 | 0,74196767 | 0,0104633 | RAB8B_ RAB8_ AtRab8B_ ATRAB8_ AtRABE1a RAB GTPase homolog 8 chr3:19876531-19878264 REVERSE LENGTH= |
| AT5G59290.1 | 1,362190292 | 0,010626 | ATUXS3_ UXS3 UDP-glucuronic acid decarboxylase 3 chr5:23915814-23917953 REVERSE LENGTH=342 |
| AT5G64040.1 | 1,383687186 | 0,0106294 | PSAN chr5:25628724-25629409 REVERSE LENGTH=171 |
| AT3G54210.1 | 0,587979655 | 0,0106673 | PRPL17 plastid ribosomal proteins of the 50S subunit 17 chr3:20067672-20068385 REVERSE LENGTH=211 |
| AT1G32060.1 | 1,108135485 | 0,0108637 | PRK phosphoribulokinase chr1:11532668-11534406 FORWARD LENGTH=395 |
| AT4G18440.1 | 1,0956881 | 0,0111027 | no symbol available no full name available chr4:10186385-10188832 REVERSE LENGTH=536 |
| AT3G48560.1 | 1,202490413 | 0,0111989 | CSR1_ IMR1_ TZP5_ ALS_ AHAS ACETOLACTATE SYNTHASE_ TRIAZOLOPYRIMIDINE RESISTANT 5_ IMIDAZO |
| AT4G30910.1 | 1,253854211 | 0,0112109 | no symbol available no full name available chr4:15042621-15045248 REVERSE LENGTH=581 |
| AT1G45000.1 | 1,16839196 | 0,011226 | RPT4b chr1:17009220-17011607 FORWARD LENGTH=399 |
| AT3G46440.1 | 1,960034574 | 0,0113438 | UXS5 UDP-XYL synthase 5 chr3:17089268-17091611 REVERSE LENGTH=341 |
| AT5G37640.1 | 1,198343023 | 0,0115082 | UBQ9 ubiquitin 9 chr5:14952782-14953750 REVERSE LENGTH=322 |
| ATCG00720.1 | 0,642969109 | 0,0115778 | PETB photosynthetic electron transfer B chrc:74841-76292 FORWARD LENGTH=215 |
| ATCG00820.1 | 0,693584141 | 0,0115778 | RPS19 ribosomal protein S19 chrc:84005-84283 REVERSE LENGTH=92 |
| AT3G57560.1 | 1,062326668 | 0,0116298 | NAGK N-acetyl-l-glutamate kinase chr3:21311164-21312207 REVERSE LENGTH=347 |
| AT2G21330.1 | 1,250294394 | 0,011774 | FBA1_ AtFBA1 fructose-bisphosphate aldolase 1 chr2:9128416-9130152 REVERSE LENGTH=399 |
| AT5G54160.1 | 1,228267304 | 0,0118804 | ATOMT1_ OMT1_ OMT3_ AtCOMT_ COMT1 caffeate O-methyltransferase 1_ O-methyltransferase 1_ O-methyltransferase 3 |
| AT4G35090.1 | 1,465099513 | 0,0119698 | CAT2 catalase 2 chr4:16700937-16703215 REVERSE LENGTH=492 |
| AT1G02920.1 | 1,209573603 | 0,0121792 | ATGSTF8_ GSTF7_ ATGSTF7_ ATGST11_ GST11 GLUTATHIONE S-TRANSFERASE 11_ glutathione S-transferase 7_ A |
| AT1G74100.1 | 1,237468047 | 0,012278 | ATSOT16_ ATST5A_ CORI-7_ SOT16 sulfotransferase 16_ CORONATINE INDUCED-7_ SULFOTRANSFERASE 16_ AR |
| AT5G12030.1 | 0,459023176 | 0,0123024 | AT-HSP17.6A_ HSP17.6A_ HSP17.6 HEAT SHOCK PROTEIN 17.6_ heat shock protein 17.6A chr5:3884214-3884684 REVI |
| AT5G14320.1 | 0,85217882 | 0,0123976 | EMB3137 EMBRYO DEFECTIVE 3137 chr5:4617839-4618772 REVERSE LENGTH=169 |
| AT5G40760.1 | 1,273219219 | 0,0125032 | G6PD6 glucose-6-phosphate dehydrogenase 6 chr5:16311284-16314556 FORWARD LENGTH=515 |
| AT4G37990.1 | 2,008391824 | 0,0125106 | ELI3-2_ ATCAD8_ ELI3_ CAD-B2 elicitor-activated gene 3-2_ CINNAMYL-ALCOHOL DEHYDROGENASE B2_ ARABII |
| AT1G79920.1 | 1,202054421 | 0,0125814 | Hsp70-15_ AtHsp70-15 heat shock protein 70-15 chr1:30059302-30062224 REVERSE LENGTH=736 |
| AT3G63460.2 | 1,068414819 | 0,0126123 | SEC31B chr3:23431009-23437241 REVERSE LENGTH=1102 |
| AT4G14030.1 | 1,228321227 | 0,0126533 | SBP1_ AtSBP1 selenium-binding protein 1 chr4:8098121-8100165 REVERSE LENGTH=490 |

| | | | |
|---|---|---|---|
| ATCG00830.1 | 0,326815561 | 0,0127402 | RPL2.1 ribosomal protein L2 chrc:84337-85843 REVERSE LENGTH=274 |
| AT4G11010.1 | 1,283225101 | 0,0127524 | NDPK3 nucleoside diphosphate kinase 3 chr4:6732780-6734298 REVERSE LENGTH=238 |
| AT2G12550.1 | 1,117948569 | 0,0128912 | NUB1 homolog of human NUB1 chr2:5114881-5118486 FORWARD LENGTH=562 |
| AT5G09660.1 | 1,258935506 | 0,0129586 | PMDH2 peroxisomal NAD-malate dehydrogenase 2 chr5:2993645-2995551 REVERSE LENGTH=354 |
| AT3G10950.1 | 0,801989573 | 0,0130369 | no symbol available no full name available chr3:3423893-3424566 FORWARD LENGTH=92 |
| AT4G23850.1 | 1,082826904 | 0,0133776 | LACS4 long-chain acyl-CoA synthetase 4 chr4:12403720-12408263 REVERSE LENGTH=666 |
| AT4G13010.1 | 1,107835774 | 0,0133796 | CeQORH chloroplast envelope Quinone Oxidoreductase Homolog chr4:7600682-7602567 FORWARD LENGTH=329 |
| AT4G24930.1 | 0,821512871 | 0,013608 | no symbol available no full name available chr4:12821496-12822389 REVERSE LENGTH=225 |
| AT3G28710.1 | 0,562445057 | 0,0136812 | VHA-d1 chr3:10773144-10775594 REVERSE LENGTH=351 |
| AT2G31790.1 | 1,261945353 | 0,0136845 | no symbol available no full name available chr2:13518269-13520167 FORWARD LENGTH=457 |
| AT1G48030.1 | 1,144408411 | 0,0137155 | mtLPD1 mitochondrial lipoamide dehydrogenase 1 chr1:17717432-17719141 REVERSE LENGTH=507 |
| AT2G33210.2 | 1,098924502 | 0,0138801 | HSP60_ HSP60-2 heat shock protein 60-2 chr2:14075093-14078568 REVERSE LENGTH=580 |
| AT5G52470.1 | 1,219441469 | 0,0140593 | ATFIB1_ FBR1_ FIB1_ SKIP7_ ATFBR1 fibrillarin 1_ FIBRILLARIN 1_ SKP1/ASK1-INTERACTING PROTEIN chr5:2129 |
| AT2G05710.1 | 1,455988074 | 0,0140789 | ACO3 aconitase 3 chr2:2141591-2146350 FORWARD LENGTH=990 |
| AT3G14420.1 | 1,12147916 | 0,0140911 | GOX1 glycolate oxidase 1 chr3:4821804-4823899 FORWARD LENGTH=367 |
| AT5G64140.1 | 1,211943941 | 0,0140926 | RPS28 ribosomal protein S28 chr5:25667529-25667723 REVERSE LENGTH=64 |
| AT1G23310.1 | 1,198564351 | 0,0142603 | GGT1_ AOAT1_ GGAT1 GLUTAMATE:GLYOXYLATE AMINOTRANSFERASE 1_ glutamate:glyoxylate aminotransferas |
| AT3G49110.1 | 0,628058248 | 0,0142907 | ATPCA_ PRX33_ ATPRX33_ PRXCA peroxidase CA_ PEROXIDASE CA_ PEROXIDASE 33 chr3:18200713-18202891 FO |
| AT3G19820.1 | 0,745720959 | 0,0143961 | DWF1_ DIM_ DIM1_ CBB1_ EVE1 ENHANCED VERY-LOW-FLUENCE RESPONSES 1_ DIMINUTIA_ DIMINUTO 1_ |
| AT5G56000.1 | 0,875186022 | 0,014463 | Hsp81.4_ AtHsp90.4 HEAT SHOCK PROTEIN 90.4_ HEAT SHOCK PROTEIN 81.4 chr5:22677602-22680067 REVERSE L |
| AT1G80560.1 | 0,755528813 | 0,0145355 | ATIMD2_ IMD2 isopropylmalate dehydrogenase 2_ ARABIDOPSIS ISOPROPYLMALATE DEHYDROGENASE 2 chr1:302 |
| AT5G02870.1 | 0,622811647 | 0,0145503 | RPL4 ribosomal large subunit 4 chr5:657830-659526 FORWARD LENGTH=407 |
| AT2G45710.1 | 1,096492932 | 0,0146011 | no symbol available no full name available chr2:18831243-18831999 FORWARD LENGTH=84 |
| AT3G49010.4 | 0,532717093 | 0,0146663 | BBC1_ RSU2_ ATBBC1 40S RIBOSOMAL PROTEIN_ breast basic conserved 1 chr3:18166971-18168047 REVERSE LENG |
| AT3G48420.1 | 1,165193422 | 0,0146693 | no symbol available no full name available chr3:17929743-17931551 FORWARD LENGTH=319 |
| AT1G49240.1 | 0,845865063 | 0,0146741 | ACT8_ FIZ1 FRIZZY AND KINKED SHOOTS_ actin 8 chr1:18216539-18217947 FORWARD LENGTH=377 |
| AT1G66580.1 | 0,623779568 | 0,0149213 | SAG24_ RPL10C senescence associated gene 24_ ribosomal protein L10 C chr1:24839208-24840439 FORWARD LENGTH=2 |
| AT2G32060.1 | 1,196398015 | 0,0152517 | no symbol available no full name available chr2:13639228-13640104 REVERSE LENGTH=144 |
| AT2G29500.1 | 1,985451062 | 0,0152782 | HSP17.6B chr2:12633279-12633740 REVERSE LENGTH=153 |
| AT5G44070.2 | 1,429522846 | 0,0153708 | CAD1_ ARA8_ PCS1_ ATPCS1 PHYTOCHELATIN SYNTHASE 1_ CADMIUM SENSITIVE 1_ ARABIDOPSIS THALIA |
| AT1G08200.1 | 1,205276458 | 0,0156033 | AXS2 UDP-D-apiose/UDP-D-xylose synthase 2 chr1:2574259-2576609 REVERSE LENGTH=389 |
| AT4G16830.2 | 0,920216038 | 0,0157174 | AtRGGA chr4:9470979-9472308 FORWARD LENGTH=265 |
| AT3G03780.1 | 1,161501218 | 0,0157192 | MS2_ ATMS2 methionine synthase 2 chr3:957602-960740 FORWARD LENGTH=765 |
| AT5G50850.1 | 1,148935295 | 0,0158234 | MAB1 MACCI-BOU chr5:20689671-20692976 FORWARD LENGTH=363 |
| AT5G59370.1 | 1,213733959 | 0,016169 | ACT4 actin 4 chr5:23950109-23951586 FORWARD LENGTH=377 |
| AT1G50200.1 | 1,131427804 | 0,0162457 | ALATS_ ACD Alanyl-tRNA synthetase chr1:18591429-18598311 REVERSE LENGTH=1003 |
| AT2G17360.1 | 0,801263332 | 0,0164223 | no symbol available no full name available chr2:7546598-7548138 FORWARD LENGTH=261 |
| AT3G22110.1 | 1,311264337 | 0,0168918 | PAC1 20S proteasome alpha subunit C1 chr3:7792819-7793571 REVERSE LENGTH=250 |
| AT4G25740.1 | 1,595336288 | 0,0170109 | no symbol available no full name available chr4:13107488-13108751 REVERSE LENGTH=177 |

| | | | |
|---|---|---|---|
| ATCG00740.1 | 0,798249394 | 0,0170123 | RPOA RNA polymerase subunit alpha chrc:77901-78890 REVERSE LENGTH=329 |
| AT3G09820.1 | 0,796252052 | 0,0170735 | ATADK1_ ADK1 adenosine kinase 1 chr3:3012122-3014624 FORWARD LENGTH=344 |
| AT4G27520.1 | 0,936131574 | 0,0172543 | ENODL2_ AtENODL2 early nodulin-like protein 2 chr4:13750668-13751819 REVERSE LENGTH=349 |
| AT3G08530.1 | 0,708518691 | 0,0173295 | AtCHC2_ CHC2 clathrin heavy chain 2 chr3:2587171-2595411 REVERSE LENGTH=1703 |
| AT5G66120.2 | 0,746642087 | 0,0173306 | no symbol available no full name available chr5:26431516-26433649 REVERSE LENGTH=442 |
| AT3G10920.1 | 1,145699653 | 0,0173957 | ATMSD1_ MEE33_ MSD1_ AtSOD1 ARABIDOPSIS MANGANESE SUPEROXIDE DISMUTASE 1_ superoxide dismutase |
| AT3G12145.1 | 0,850231509 | 0,0174983 | FLR1_ FTM4 FLOR1_ FLORAL TRANSITION AT THE MERISTEM4 chr3:3874764-3876075 REVERSE LENGTH=325 |
| AT5G20290.1 | 0,549940223 | 0,0174986 | no symbol available no full name available chr5:6851695-6853012 REVERSE LENGTH=222 |
| AT1G29910.1 | 0,865630761 | 0,017581 | CAB3_ LHCB1.2_ AB180 LIGHT HARVESTING CHLOROPHYLL A/B BINDING PROTEIN 1.2_ chlorophyll A/B binding |
| AT5G51110.1 | 1,28759542 | 0,0180527 | ATP1_ SDIRIP1_ RAF2 SDIR1-INTERACTING PROTEIN1_ AtAIRP2 Target Protein 1_ Rubisco Assembly Factor 2 chr5:20 |
| AT2G19900.1 | 1,188068835 | 0,0182623 | ATNADP-ME1_ NADP-ME1 NADP-malic enzyme 1_ Arabidopsis thaliana NADP-malic enzyme 1 chr2:8592106-8595403 RE |
| AT1G22760.1 | 1,44334521 | 0,0184363 | PAB3_ PABP3 poly(A) binding protein 3 chr1:8055599-8058799 FORWARD LENGTH=660 |
| AT5G16050.1 | 1,182705477 | 0,0188021 | GRF5_ GF14 UPSILON general regulatory factor 5 chr5:5244008-5245402 REVERSE LENGTH=268 |
| AT2G20140.1 | 1,159767007 | 0,0188277 | RPT2b regulatory particle AAA-ATPase 2b chr2:8692736-8694837 FORWARD LENGTH=443 |
| AT3G23570.1 | 1,2194817 | 0,0190002 | no symbol available no full name available chr3:8458052-8459608 REVERSE LENGTH=239 |
| AT1G05190.1 | 0,883937126 | 0,0191002 | RPL6_ EMB2394 embryo defective 2394 chr1:1502515-1503738 REVERSE LENGTH=223 |
| AT1G75280.1 | 2,174814571 | 0,0191378 | no symbol available no full name available chr1:28252030-28253355 FORWARD LENGTH=310 |
| AT2G26740.1 | 1,268998062 | 0,0192618 | ATSEH_ SEH soluble epoxide hydrolase chr2:11393148-11394257 REVERSE LENGTH=321 |
| AT3G22630.1 | 2,670583367 | 0,0192854 | PBD1_ PRCGB 20S proteasome beta subunit D1 chr3:8009709-8010774 REVERSE LENGTH=204 |
| AT4G27560.1 | 1,315105775 | 0,0193091 | UGT79B2 chr4:13760114-13761481 REVERSE LENGTH=455 |
| AT4G34670.1 | 1,072105912 | 0,0194273 | no symbol available no full name available chr4:16548724-16550222 FORWARD LENGTH=262 |
| AT5G16620.1 | 1,288858844 | 0,0194934 | PDE120_ TIC40_ ATTIC40 translocon at the inner envelope membrane of chloroplasts 40_ pigment defective embryo 120 chr5 |
| AT5G39740.1 | 0,92236429 | 0,0196555 | RPL5B_ OLI7 ribosomal protein L5 B_ OLIGOCELLULA 7 chr5:15903365-15905185 FORWARD LENGTH=301 |
| AT2G14880.1 | 1,538211299 | 0,0196696 | SWIB2 chr2:6393686-6394841 REVERSE LENGTH=141 |
| AT5G44020.1 | 0,696224231 | 0,019705 | no symbol available no full name available chr5:17712433-17714046 FORWARD LENGTH=272 |
| AT5G20720.1 | 1,139516298 | 0,0197749 | ATCPN21_ CPN20_ CPN21_ CHCPN10_ CPN10 chaperonin 20_ CHLOROPLAST CHAPERONIN 10 chr5:7015015-701635 |
| AT2G39390.1 | 0,542176498 | 0,019934 | no symbol available no full name available chr2:16450803-16451762 REVERSE LENGTH=123 |
| AT2G36880.1 | 0,338550598 | 0,0199344 | MAT3 methionine adenosyltransferase 3 chr2:15479721-15480893 REVERSE LENGTH=390 |
| AT3G13930.1 | 1,235400708 | 0,0200833 | mtE2-2 mitochondrial pyruvate dehydrogenase subunit 2-2 chr3:4596240-4600143 FORWARD LENGTH=539 |
| AT2G37040.1 | 1,236144648 | 0,0201432 | PAL1_ ATPAL1 PHE ammonia lyase 1 chr2:15557602-15560237 REVERSE LENGTH=725 |
| AT1G03130.1 | 1,125438734 | 0,0201506 | PSAD-2 photosystem I subunit D-2 chr1:753528-754142 REVERSE LENGTH=204 |
| AT3G11510.1 | 0,839609842 | 0,0203629 | no symbol available no full name available chr3:3623757-3624866 REVERSE LENGTH=150 |
| AT1G66970.1 | 0,75812553 | 0,0203865 | SVL2_ GDPDL1 SHV3-like 2_ Glycerophosphodiester phosphodiesterase (GDPD) like 1 chr1:24992746-24996005 REVERSE |
| AT1G56340.1 | 1,312765584 | 0,0206623 | AtCRT1a_ CRT1_ CRT1a calreticulin 1a_ calreticulin 1 chr1:21090059-21092630 REVERSE LENGTH=425 |
| AT2G20260.1 | 0,902649163 | 0,020709 | PSAE-2 photosystem I subunit E-2 chr2:8736780-8737644 FORWARD LENGTH=145 |
| AT1G76010.1 | 0,697087223 | 0,020826 | ALBA1_ Atalba1_ ALBA4 chr1:28528505-28530488 REVERSE LENGTH=350 |
| AT3G16410.1 | 1,201168007 | 0,0209211 | NSP4 nitrile specifier protein 4 chr3:5572145-5574359 FORWARD LENGTH=619 |
| AT1G67700.1 | 0,864253349 | 0,02096 | HHL1 HYPERSENSITIVE TO HIGH LIGHT 1 chr1:25374295-25375716 FORWARD LENGTH=230 |
| AT5G07440.3 | 1,60535195 | 0,0212493 | GDH2 glutamate dehydrogenase 2 chr5:2356153-2357546 FORWARD LENGTH=309 |

| | | | |
|---|---|---|---|
| AT5G47210.1 | 1,102018036 | 0,0212752 | no symbol available no full name available chr5:19169222-19171012 REVERSE LENGTH=357 |
| AT4G19410.1 | 1,132161941 | 0,0214507 | PAE7_ AtPAE7 Pectin Acetyesterase 7_ pectin acetylesterase 7 chr4:10582188-10584766 REVERSE LENGTH=391 |
| AT1G10760.1 | 1,224757392 | 0,0216037 | GWD_ GWD1_ SOP1_ SOP_ SEX1 STARCH EXCESS 1 chr1:3581210-3590043 REVERSE LENGTH=1399 |
| AT3G06860.1 | 1,244554306 | 0,0217487 | ATMFP2_ MFP2 MULTIFUNCTIONAL PROTEIN 2_ multifunctional protein 2 chr3:2161926-2166009 FORWARD LENGT |
| AT1G13060.1 | 1,214284667 | 0,0218185 | PBE1 20S proteasome beta subunit E1 chr1:4452641-4454663 FORWARD LENGTH=274 |
| AT3G26450.1 | 1,297763641 | 0,0219086 | no symbol available no full name available chr3:9681593-9683299 REVERSE LENGTH=152 |
| AT4G23900.1 | 0,89680907 | 0,021956 | no symbol available no full name available chr4:12424505-12426318 FORWARD LENGTH=237 |
| AT4G16760.2 | 1,354407989 | 0,0224035 | ATACX1_ ACX1 acyl-CoA oxidase 1 chr4:9424930-9428689 REVERSE LENGTH=651 |
| AT2G01290.1 | 0,861176569 | 0,0224894 | RPI2 ribose-5-phosphate isomerase 2 chr2:149192-149989 REVERSE LENGTH=265 |
| AT3G18000.1 | 1,394264156 | 0,0225955 | PEAMT1_ AtPMT1_ PEAMT_ NMT1_ XPL1_ DPR2 N-METHYLTRANSFERASE 1_ Phosphoethanolamine methyltransfera |
| AT5G62350.1 | 0,589230894 | 0,0229526 | no symbol available no full name available chr5:25037504-25038112 FORWARD LENGTH=202 |
| AT1G03680.1 | 1,380074881 | 0,0229891 | ATHM1_ TRX-M1_ ATM1_ THM1 thioredoxin M-type 1_ THIOREDOXIN M-TYPE 1_ ARABIDOPSIS THIOREDOXIN M |
| AT3G09630.1 | 0,664440344 | 0,0231542 | SAC56 Suppressor of Acaulis 56 chr3:2953813-2955444 FORWARD LENGTH=406 |
| AT1G30230.1 | 1,713400535 | 0,0231997 | EF1Bb_ eEF-1Bb1 eukaryotic elongation factor 1B beta 1_ elongation factor 1B beta chr1:10639286-10640515 FORWARD LE |
| AT3G54640.1 | 0,228732283 | 0,0236563 | TRP3_ TSA1 TRYPTOPHAN-REQUIRING 3_ tryptophan synthase alpha chain chr3:20223331-20225303 REVERSE LENGT |
| AT2G23350.1 | 1,141241829 | 0,0239839 | PABP4_ PAB4 POLY(A) BINDING PROTEIN 4_ poly(A) binding protein 4 chr2:9943209-9946041 FORWARD LENGTH=6 |
| AT1G61520.1 | 0,924733854 | 0,0243526 | LHCA3 photosystem I light harvesting complex gene 3 chr1:22700152-22701149 FORWARD LENGTH=273 |
| AT3G56070.1 | 1,114826311 | 0,0243849 | ROC2 rotamase cyclophilin 2 chr3:20806987-20807517 REVERSE LENGTH=176 |
| AT3G28290.1 | 0,806789509 | 0,0248338 | AT14A chr3:10547873-10549030 FORWARD LENGTH=385 |
| AT3G25770.1 | 0,908168168 | 0,0250188 | AOC2 allene oxide cyclase 2 chr3:9406975-9407839 FORWARD LENGTH=253 |
| AT4G02450.2 | 1,095872816 | 0,0255348 | p23-1 chr4:1073987-1075765 REVERSE LENGTH=240 |
| AT4G18360.1 | 0,796882019 | 0,025661 | GOX3 glycolate oxidase 3 chr4:10146141-10148386 REVERSE LENGTH=368 |
| AT4G22240.1 | 1,244035039 | 0,0256703 | FBN1b fibrillin 1b chr4:11766090-11767227 REVERSE LENGTH=310 |
| AT1G66240.1 | 1,484201988 | 0,0257626 | AtHMP14_ ATX1_ ATATX1 HEAVY METAL ASSOCIATED PROTEIN 14_ homolog of anti-oxidant 1 chr1:24686445-2468 |
| AT1G52230.1 | 0,958466513 | 0,0260004 | PSI-H_ PSAH-2_ PSAH2 photosystem I subunit H2_ PHOTOSYSTEM I SUBUNIT H-2 chr1:19454902-19455508 FORWARI |
| AT1G55210.1 | 1,092473982 | 0,0261297 | no symbol available no full name available chr1:20598057-20598620 REVERSE LENGTH=187 |
| AT1G26480.1 | 0,605700591 | 0,0263587 | GF14 IOTA_ GRF12 general regulatory factor 12 chr1:9156573-9157845 REVERSE LENGTH=268 |
| AT3G14310.1 | 0,890717884 | 0,0265054 | OZS2_ ATPME3_ PME3 pectin methylesterase 3_ OVERLY ZINC SENSITIVE 2 chr3:4772214-4775095 REVERSE LENGT |
| AT4G36250.1 | 1,133858929 | 0,0265487 | ALDH3F1 aldehyde dehydrogenase 3F1 chr4:17151029-17153381 FORWARD LENGTH=484 |
| AT2G30870.1 | 1,110828752 | 0,0269611 | GSTF10_ ATGSTF10_ ERD13_ ATGSTF4 glutathione S-transferase PHI 10_ ARABIDOPSIS THALIANA GLUTATHIONE |
| AT4G38680.1 | 1,843685618 | 0,0270684 | GRP2_ CSP2_ ATCSP2_ CSDP2 glycine rich protein 2_ COLD SHOCK DOMAIN PROTEIN 2_ ARABIDOPSIS THALIANA |
| AT5G28510.1 | 0,874602707 | 0,0271923 | BGLU24 beta glucosidase 24 chr5:10481041-10484022 REVERSE LENGTH=533 |
| AT2G43030.1 | 0,787763048 | 0,0271935 | PRPL3 plastid ribosomal proteins of the 50S subunit chr2:17894898-17895713 FORWARD LENGTH=271 |
| AT3G27300.1 | 1,420635293 | 0,0272851 | G6PD5 glucose-6-phosphate dehydrogenase 5 chr3:10083318-10086288 REVERSE LENGTH=516 |
| AT2G35370.1 | 0,897799103 | 0,0274564 | GDCH glycine decarboxylase complex H chr2:14891239-14892050 FORWARD LENGTH=165 |
| AT5G44340.1 | 1,139481912 | 0,0274632 | TUB4 tubulin beta chain 4 chr5:17859442-17860994 REVERSE LENGTH=444 |
| AT1G64190.1 | 1,264528084 | 0,0275015 | PGD1 6-phosphogluconate dehydrogenase 1 chr1:23825549-23827012 REVERSE LENGTH=487 |
| AT1G20010.1 | 0,930931461 | 0,0275601 | TUB5 tubulin beta-5 chain chr1:6938033-6940481 REVERSE LENGTH=449 |
| AT2G22290.1 | 0,649024052 | 0,0283712 | RAB-H1D_ ATRABH1D_ ATRAB6_ RABH1d_ ATRAB-H1D RAB GTPASE HOMOLOG H1D_ ARABIDOPSIS RAB GTF |

| | | | |
|---|---|---|---|
| ATCG01110.1 | 0,674944682 | 0,0288979 | NDHH NAD(P)H dehydrogenase subunit H chrc:122011-123192 REVERSE LENGTH=393 |
| AT3G23600.1 | 1,246485804 | 0,0290167 | no symbol available no full name available chr3:8473833-8475655 FORWARD LENGTH=239 |
| AT1G78570.1 | 1,170326522 | 0,0294043 | RHM1_ ATRHM1_ ROL1 REPRESSOR OF LRX1 1_ rhamnose biosynthesis 1_ ARABIDOPSIS THALIANA RHAMNOSE E |
| AT1G55060.1 | 0,899425816 | 0,029618 | UBQ12 ubiquitin 12 chr1:20549533-20550225 FORWARD LENGTH=230 |
| AT1G56450.1 | 1,128310655 | 0,0296278 | MUD1_ PBG1 20S proteasome beta subunit G1 chr1:21141970-21144186 FORWARD LENGTH=246 |
| AT5G57850.1 | 0,81527475 | 0,0299451 | ADCL_ DAT1 D-AA specific transaminase D-AAT_ 4-amino-4-deoxychorismate lyase chr5:23435548-23437287 REVERSE L |
| AT5G27470.1 | 1,240332581 | 0,030012 | no symbol available no full name available chr5:9695087-9697154 FORWARD LENGTH=451 |
| AT1G14410.1 | 0,514888275 | 0,0300527 | PTAC1_ ATWHY1_ WHY1 A. THALIANA WHIRLY 1_ WHIRLY 1 chr1:4929352-4930810 REVERSE LENGTH=263 |
| AT5G12020.1 | 1,410509153 | 0,0307001 | HSP17.6II 17.6 kDa class II heat shock protein chr5:3882409-3882876 REVERSE LENGTH=155 |
| AT3G46830.1 | 1,225800085 | 0,0307998 | ATRAB-A2C_ ATRAB11A_ ATRABA2C_ RABA2c_ RAB-A2C RAB GTPASE HOMOLOG A2C_ ARABIDOPSIS RAB G |
| AT1G19580.1 | 1,690302969 | 0,0308909 | GAMMA CA1 gamma carbonic anhydrase 1 chr1:6774937-6777092 FORWARD LENGTH=275 |
| AT2G17630.1 | 0,768340803 | 0,0312705 | PSAT2 phosphoserine aminotransferase 2 chr2:7666637-7667905 FORWARD LENGTH=422 |
| AT1G77490.2 | 0,84260092 | 0,0314422 | TAPX thylakoidal ascorbate peroxidase chr1:29117688-29120649 FORWARD LENGTH=421 |
| AT5G48880.1 | 1,234901306 | 0,0321431 | PKT2_ PKT1_ KAT5 3-KETO-ACYL-COENZYME A THIOLASE 5_ peroxisomal 3-keto-acyl-CoA thiolase 2_ PEROXISOM |
| AT4G39800.1 | 1,637111202 | 0,0326441 | ATMIPS1_ ATIPS1_ MIPS1_ MI-1-P SYNTHASE INOSITOL 3-PHOSPHATE SYNTHASE 1_ MYO-INOSITOL-1-PHOSPI |
| ATCG00900.1 | 0,638528992 | 0,0326788 | RPS7_ RPS7.1 CHLOROPLAST RIBOSOMAL PROTEIN S7 chrc:97478-97945 REVERSE LENGTH=155 |
| AT1G55490.1 | 1,080703777 | 0,0327118 | Cpn60beta1_ LEN1_ CPNB1_ CPN60B chaperonin-60beta1_ LESION INITIATION 1_ chaperonin 60 beta chr1:20715717-20 |
| AT4G35460.1 | 1,288269424 | 0,0327399 | ATNTRB_ NTR1_ NTRB NADPH-DEPENDENT THIOREDOXIN REDUCTASE 1_ NADPH-dependent thioredoxin reducta |
| AT5G53850.1 | 1,281729479 | 0,0327986 | DEP1 DEHYDRATASE-ENOLASE-PHOSPHATASE-COMPLEX 1 chr5:21861617-21864817 REVERSE LENGTH=402 |
| AT2G27730.1 | 1,577472896 | 0,0328138 | no symbol available no full name available chr2:11820056-11820867 REVERSE LENGTH=113 |
| AT2G21060.1 | 1,397487758 | 0,0333735 | GRP2B_ ATCSP4_ ATGRP2B glycine-rich protein 2B_ COLD SHOCK DOMAIN PROTEIN 4 chr2:9036983-9037588 REVE |
| AT5G43940.1 | 1,068787408 | 0,0334663 | ADH2_ HOT5_ ATGSNOR1_ PAR2_ GSNOR PARAQUAT RESISTANT 2_ S-NITROSOGLUTATHIONE REDUCTASE_ |
| AT2G34460.1 | 1,784267369 | 0,0337758 | no symbol available no full name available chr2:14529635-14530732 FORWARD LENGTH=280 |
| AT1G54000.1 | 0,760617762 | 0,0338399 | GLL22 GDSL lipase-like protein 22 chr1:20154548-20156365 REVERSE LENGTH=391 |
| AT1G06400.1 | 1,381294226 | 0,0339303 | ARA2_ ATRABA1A_ ATRAB11E_ ARA-2 ARABIDOPSIS THALIANA RAB GTPASE HOMOLOG A1A chr1:1951089-19 |
| AT3G14415.1 | 1,15890408 | 0,0344372 | GOX2 glycolate oxidase 2 chr3:4818667-4820748 FORWARD LENGTH=367 |
| AT3G57490.1 | 0,786959339 | 0,0348223 | no symbol available no full name available chr3:21279824-21280887 REVERSE LENGTH=276 |
| AT5G59240.1 | 0,447792157 | 0,0348771 | no symbol available no full name available chr5:23902626-23903670 REVERSE LENGTH=210 |
| AT5G25980.2 | 1,062198576 | 0,0351483 | BGLU37_ TGG2 BETA GLUCOSIDASE 37_ glucoside glucohydrolase 2 chr5:9072730-9075477 FORWARD LENGTH=547 |
| AT2G30050.1 | 1,497749376 | 0,0352151 | no symbol available no full name available chr2:12825540-12826448 FORWARD LENGTH=302 |
| AT5G02940.1 | 0,722171946 | 0,0352977 | PEC1 PLASTID ENVELOPE ION CHANNELS 1 chr5:684671-689674 REVERSE LENGTH=813 |
| AT1G06410.1 | 0,750730249 | 0,0354179 | ATTPSA_ TPS7_ ATTPS7 TREHALOSE -6-PHOSPHATASE SYNTHASE S7_ trehalose-phosphatase/synthase 7 chr1:19554 |
| AT2G33800.1 | 0,897144745 | 0,0356043 | SCA1_ RPS5_ EMB3113_ PRPS5 plastid ribosomal protein of the 30S subunit 5_ EMBRYO DEFECTIVE 3113_ ribosomal pr |
| AT1G32080.1 | 0,639541107 | 0,0359595 | AtLrgB_ LrgB_ PLGG_ PLGG1 chr1:11537572-11539756 REVERSE LENGTH=512 |
| AT4G37980.1 | 1,177873158 | 0,036151 | ELI3-1_ CHR_ ELI3_ ATCAD7_ CAD7 elicitor-activated gene 3-1_ CINNAMALDEHYDE AND HEXENAL REDUCTASE_ |
| AT5G13410.1 | 0,801537604 | 0,0361926 | no symbol available no full name available chr5:4299830-4301706 REVERSE LENGTH=256 |
| AT1G62660.1 | 1,202649308 | 0,0362 | VI1 VACUOLAR INVERTASE 1 chr1:23199949-23203515 FORWARD LENGTH=648 |
| AT3G61050.1 | 0,895004893 | 0,0363721 | CLB1_ SYT7_ AtCLB_ NTMC2TYPE4_ NTMC2T4 calcium-dependent lipid-binding protein_ Synaptotagmin 7 chr3:2259748 |
| AT3G26070.1 | 0,681495044 | 0,0364195 | FBN3a FIBRILLIN3a chr3:9526904-9528199 FORWARD LENGTH=242 |

| | | | |
|---|---|---|---|
| AT4G26900.1 | 1,251420367 | 0,0365307 | AT-HF_ HISN4 HIS HF chr4:13515514-13519608 FORWARD LENGTH=592 |
| AT3G46780.1 | 0,902942187 | 0,0365755 | PTAC16 plastid transcriptionally active 16 chr3:17228766-17231021 FORWARD LENGTH=510 |
| AT1G08450.2 | 1,322550809 | 0,0368992 | CRT3_ AtCRT3_ EBS2_ PSL1 A. thaliana calreticulin 3_ EMS-MUTAGENIZED BRI1 SUPPRESSOR 2_ calreticulin 3_ PRI( |
| AT5G20950.1 | 1,156718062 | 0,036983 | BGLC1 chr5:7107609-7110775 REVERSE LENGTH=624 |
| AT5G42980.1 | 1,167715785 | 0,0372719 | ATH3_ TRX3_ TRXH3_ ATTRX3_ ATTRXH3 THIOREDOXIN H3_ thioredoxin 3_ thioredoxin H-type 3 chr5:17242772-17 |
| AT4G22670.1 | 1,089203904 | 0,0374399 | AtHip1_ HIP1_ TPR11 HSP70-interacting protein 1_ tetratricopeptide repeat 11 chr4:11918236-11920671 FORWARD LENG |
| AT1G01620.1 | 0,770553947 | 0,0374629 | TMP-B_ PIP1;3_ PIP1C plasma membrane intrinsic protein 1C_ PLASMA MEMBRANE INTRINSIC PROTEIN 1;3 chr1:225 |
| AT4G11820.1 | 1,277318292 | 0,0378737 | MVA1_ FKP1_ HMGS FLAKY POLLEN 1_ HYDROXYMETHYLGLUTARYL-COA SYNTHASE chr4:7109124-7111213 F |
| AT2G02010.1 | 0,399272468 | 0,037912 | GAD4 glutamate decarboxylase 4 chr2:474375-476495 REVERSE LENGTH=493 |
| AT5G39730.1 | 0,702237749 | 0,0382773 | no symbol available no full name available chr5:15901740-15902624 FORWARD LENGTH=172 |
| AT5G59880.2 | 0,845601554 | 0,0383266 | ADF3 actin depolymerizing factor 3 chr5:24120382-24121628 FORWARD LENGTH=124 |
| AT4G30920.1 | 1,131661756 | 0,0383442 | LAP2_ AtLAP2 leucyl aminopeptidase 2 chr4:15046589-15049304 REVERSE LENGTH=583 |
| AT5G02500.1 | 1,284166533 | 0,0384539 | AtHsp70-1_ HSP70-1_ HSC70-1_ HSC70_ AT-HSC70-1 ARABIDOPSIS THALIANA HEAT SHOCK COGNATE PROTEIN |
| AT3G16480.1 | 0,754327812 | 0,0385156 | MPPalpha mitochondrial processing peptidase alpha subunit chr3:5599906-5602716 FORWARD LENGTH=499 |
| AT5G16590.1 | 0,854860295 | 0,0386306 | LRR1 Leucine rich repeat protein 1 chr5:5431862-5433921 FORWARD LENGTH=625 |
| AT2G44650.1 | 1,363867355 | 0,0386866 | CHL-CPN10_ CPN10 CHLOROPLAST CHAPERONIN 10_ chloroplast chaperonin 10 chr2:18419521-18420510 REVERSE I |
| AT5G40370.1 | 1,2346449 | 0,0389404 | AtGRXC2_ GRXC2_ GRX370 glutaredoxin C2 chr5:16147826-16149052 REVERSE LENGTH=111 |
| AT5G62690.1 | 1,169970316 | 0,0390293 | TUB2 tubulin beta chain 2 chr5:25181560-25183501 FORWARD LENGTH=450 |
| AT5G08280.1 | 0,914335472 | 0,0400062 | HEMC_ RUG1 RUGOSA 1_ hydroxymethylbilane synthase chr5:2663763-2665596 REVERSE LENGTH=382 |
| AT2G45290.1 | 0,761058516 | 0,0400596 | TKL2 transketolase 2 chr2:18672737-18675589 FORWARD LENGTH=741 |
| AT5G54640.1 | 0,653651176 | 0,0403026 | HTA1_ RAT5_ ATHTA1 histone H2A 1_ RESISTANT TO AGROBACTERIUM TRANSFORMATION 5 chr5:22196540-221 |
| AT5G23010.1 | 1,368067775 | 0,0403237 | GSM1_ MAM1_ IMS3 glucosinolate metabolism 1_ 2-ISOPROPYLMALATE SYNTHASE 3_ methylthioalkylmalate synthase |
| AT4G29130.1 | 1,20997632 | 0,0407013 | GIN2_ HXK1_ ATHXK1 GLUCOSE INSENSITIVE 2_ hexokinase 1_ ARABIDOPSIS THALIANA HEXOKINASE 1 chr4:1 |
| AT5G50950.3 | 0,827057873 | 0,0412903 | FUM2 FUMARASE 2 chr5:20731191-20733636 FORWARD LENGTH=317 |
| AT2G06850.1 | 1,132479262 | 0,0413233 | EXT_ XTH4_ EXGT-A1 ENDOXYLOGLUCAN TRANSFERASE_ endoxyloglucan transferase A1_ xyloglucan endotransgluc |
| AT3G62870.1 | 0,79707311 | 0,041721 | no symbol available no full name available chr3:23242862-23244273 REVERSE LENGTH=256 |
| AT1G78380.1 | 1,112969787 | 0,0418375 | GSTU19_ GST8_ ATGSTU19 A. THALIANA GLUTATHIONE S-TRANSFERASE TAU 19_ glutathione S-transferase TAU |
| AT1G78300.1 | 0,927083394 | 0,0421196 | GRF2_ 14-3-3OMEGA_ GF14 OMEGA general regulatory factor 2_ 14-3-3 PROTEIN G-BOX FACTOR14 OMEGA chr1:294 |
| AT2G36620.1 | 0,782933727 | 0,0424908 | RPL24A ribosomal protein L24 chr2:15350548-15351819 REVERSE LENGTH=164 |
| AT1G11860.1 | 1,087343783 | 0,0425155 | GLDT chr1:4001801-4003245 FORWARD LENGTH=408 |
| AT4G39980.1 | 1,121123194 | 0,0438166 | AtDAHP1_ DHS1_ DAHP1 3-DEOXY-D-ARABINO-HEPTULOSONATE-7-PHOSPHATE 1_ 3-deoxy-D-arabino-heptulosor |
| AT1G04710.1 | 1,196935202 | 0,0438574 | KAT1_ PKT4 3-KETO-ACYL-COA THIOLASE 1_ peroxisomal 3-ketoacyl-CoA thiolase 4 chr1:1321941-1324556 FORWAR |
| AT5G14920.1 | 2,401094924 | 0,0441084 | GASA14 A-stimulated in Arabidopsis 14 chr5:4826598-4827761 FORWARD LENGTH=275 |
| AT3G60770.1 | 0,654178133 | 0,044236 | no symbol available no full name available chr3:22460525-22461656 REVERSE LENGTH=151 |
| AT3G51800.3 | 1,256616779 | 0,0446902 | ATG2_ EBP1_ ATEBP1_ G2p A. THALIANA ERBB-3 BINDING PROTEIN 1_ ERBB-3 BINDING PROTEIN 1 chr3:19211 |
| AT1G44575.1 | 0,780192436 | 0,0448754 | CP22_ PSBS_ NPQ4 NONPHOTOCHEMICAL QUENCHING 4_ PHOTOSYSTEM II SUBUNIT S chr1:16871768-16873194 |
| AT5G15090.1 | 1,191326553 | 0,0449062 | VDAC3_ AtVDAC-3_ ATVDAC3 ARABIDOPSIS THALIANA VOLTAGE DEPENDENT ANION CHANNEL 3_ voltage de |
| AT1G74260.1 | 1,193907648 | 0,0452385 | PUR4 purine biosynthesis 4 chr1:27923005-27927764 REVERSE LENGTH=1407 |
| AT3G15356.1 | 1,350630709 | 0,0455342 | no symbol available no full name available chr3:5174603-5175418 REVERSE LENGTH=271 |

| | | | |
|---|---|---|---|
| AT3G18780.1 | 1,336013305 | 0,0459327 | LSR2_ ACT2_ ENL2_ DER1_ FIZ2 FRIZZY AND KINKED SHOOTS 2_ LIGHT STRESS-REGULATED 2_ DEFORMED R |
| AT5G13030.1 | 1,308373897 | 0,0461658 | SELO SELENOPROTEIN O chr5:4133216-4136461 FORWARD LENGTH=633 |
| AT4G35830.1 | 1,172651804 | 0,0462421 | ACO1 aconitase 1 chr4:16973007-16977949 REVERSE LENGTH=898 |
| AT3G17210.1 | 1,133025444 | 0,0467207 | ATHS1_ HS1 heat stable protein 1_ A. THALIANA HEAT STABLE PROTEIN 1 chr3:5882318-5882896 FORWARD LENGT |
| AT3G04940.1 | 1,366144957 | 0,0471738 | ATCYSD1_ CYSD1 CYSTEINE SYNTHASE D1_ cysteine synthase D1 chr3:1365681-1367508 FORWARD LENGTH=324 |
| AT1G18080.1 | 1,203428594 | 0,0472115 | RACK1A_AT_ AtRACK1_ SAC53_ ATARCA_ RACK1A RECEPTOR FOR ACTIVATED C KINASE 1 A_ Suppressor of A |
| AT4G18100.1 | 0,411317038 | 0,0478469 | no symbol available no full name available chr4:10035715-10036475 REVERSE LENGTH=133 |
| AT5G24780.1 | 1,09792355 | 0,0479086 | VSP1_ ATVSP1 vegetative storage protein 1 chr5:8507783-8508889 REVERSE LENGTH=270 |
| AT3G16400.1 | 0,871309292 | 0,0482341 | NSP1_ ATNSP1_ ATMLP-470 nitrile specifier protein 1_ NITRILE SPECIFIER PROTEIN 1_ MYROSINASE-BINDING PRO |
| AT1G50250.1 | 0,896613201 | 0,0482632 | FTSH1 FTSH protease 1 chr1:18614398-18616930 REVERSE LENGTH=716 |
| AT2G31680.1 | 0,736799228 | 0,0488146 | AtRABA5d_ RABA5d RAB GTPase homolog A5D chr2:13473781-13474957 REVERSE LENGTH=219 |
| AT3G14290.1 | 1,168499141 | 0,0489986 | PAE2 20S proteasome alpha subunit E2 chr3:4764364-4766381 FORWARD LENGTH=237 |
| AT2G30970.1 | 1,121755124 | 0,0490736 | ASP1 aspartate aminotransferase 1 chr2:13179012-13181686 FORWARD LENGTH=430 |
| AT2G47730.1 | 1,208416754 | 0,0491286 | GST6_ GSTF8_ ATGSTF8_ ATGSTF5 glutathione S-transferase phi 8_ Arabidopsis thaliana glutathione S-transferase phi 8_ G |
| AT1G10840.1 | 1,242749859 | 0,049311 | TIF3H1 translation initiation factor 3 subunit H1 chr1:3607885-3610299 REVERSE LENGTH=337 |
| AT3G08740.1 | 0,761292912 | 0,0496953 | no symbol available no full name available chr3:2654788-2656154 REVERSE LENGTH=236 |
| AT5G38660.1 | 0,646012194 | 0,0497244 | APE1 ACCLIMATION OF PHOTOSYNTHESIS TO ENVIRONMENT chr5:15473285-15475497 REVERSE LENGTH=286 |

THALIANA PROTEIN DISULFIDE ISOMERASE 11_ UNFERTILIZED EMBRYO SAC 5_ MATERNAL EFFECT EMBRYO ARREST 30 chr2:19481503-19483683 FOR

GNAL RECOGNITION PARTICLE 54 KDA SUBUNIT CHLOROPLAST PROTEIN_ FIFTY-FOUR CHLOROPLAST HOMOLOGUE chr5:1060265-1063257 REVERSE

CHAPERONIN 60ALPHA_ ACCUMULATION AND REPLICATION OF CHLOROPLASTS 2_ chaperonin-60alpha chr2:11926603-11929184 FORWARD LENGTH=586

:SISTANCE 8_ ARABIDOPSIS PLEIOTROPIC DRUG RESISTANCE 8_ PENETRATION 3_ ATP-binding cassette G36 chr1:22034661-22039844 FORWARD LENGTH=

)OPSIS THALIANA CINNAMYL-ALCOHOL DEHYDROGENASE 8_ ELICITOR-ACTIVATED GENE 3 chr4:17855964-17857388 FORWARD LENGTH=359

70-1_ heat shock cognate protein 70-1_ HEAT SHOCK COGNATE PROTEIN 70_ HEAT SHOCK PROTEIN 70-1 chr5:554055-556334 REVERSE LENGTH=651

ꝛWARD LENGTH=361

LENGTH=564
