## Supplementary material for "HY5 enhances *Arabidopsis* tolerance to combined high light and heat stress by coordinating photoprotection and hormone signaling": Table S6

**Supplemental Table S6. Differentially accumulated proteins compared to control (P < 0.05) in *hy5-215* leaves subjected to heat stress.**

| Protein ID | Fold Change | p-value | Protein description |
|---|---|---|---|
| AT2G35635.1 | 6,11137702 | 1,79643E-07 | UBQ7_ RUB2 RELATED TO UBIQUITIN 2_ ubiquitin 7 chr2:14981044-14981943 FORWARD LENGTH=154 |
| AT5G14740.2 | 0,220838142 | 8,35569E-07 | BETA CA2_ CA2_ DEG12_ CA18 BETA CARBONIC ANHYDRASE 2_ CARBONIC ANHYDRASE 18_ carbonic anhydrase |
| AT5G52640.1 | 7,260875257 | 1,69035E-06 | HSP81-1_ ATHSP90.1_ HSP81.1_ AtHsp90-1_ HSP83_ ATHS83_ HSP90.1 HEAT SHOCK PROTEIN 90-1_ heat shock prote |
| AT5G11170.1 | 0,301806741 | 1,99461E-06 | UAP56a homolog of human UAP56 a chr5:3553334-3556646 FORWARD LENGTH=427 |
| AT3G25920.1 | 0,746191264 | 2,601E-06 | RPL15 ribosomal protein L15 chr3:9491268-9492558 REVERSE LENGTH=277 |
| AT2G42910.1 | 2,462021985 | 2,77197E-06 | AtPRS4_ PRS4 phosphoribosyl diphosphate synthase 4 chr2:17856396-17858394 FORWARD LENGTH=337 |
| AT1G65350.1 | 1,979882468 | 3,30494E-06 | UBQ13 ubiquitin 13 chr1:24272518-24277275 REVERSE LENGTH=319 |
| AT5G23860.1 | 3,032925272 | 3,67301E-06 | TUB8 tubulin beta 8 chr5:8042962-8044528 FORWARD LENGTH=449 |
| AT1G65980.1 | 1,48680282 | 5,28666E-06 | TPX1 thioredoxin-dependent peroxidase 1 chr1:24559524-24560753 REVERSE LENGTH=162 |
| AT1G74260.1 | 1,355346412 | 5,61841E-06 | PUR4 purine biosynthesis 4 chr1:27923005-27927764 REVERSE LENGTH=1407 |
| AT5G14740.5 | 1,648932191 | 5,91915E-06 | BETA CA2_ CA2_ DEG12_ CA18 BETA CARBONIC ANHYDRASE 2_ CARBONIC ANHYDRASE 18_ carbonic anhydrase |
| AT2G38540.1 | 8,29194943 | 5,9374E-06 | ATLTP1_ AtLtpI-4_ LTP1_ LP1 ARABIDOPSIS THALIANA LIPID TRANSFER PROTEIN 1_ lipid transfer protein 1_ LIPI |
| AT1G18210.1 | 0,489277919 | 6,25889E-06 | no symbol available no full name available chr1:6268273-6268785 REVERSE LENGTH=170 |
| AT2G29500.1 | 11,12640643 | 6,41697E-06 | HSP17.6B chr2:12633279-12633740 REVERSE LENGTH=153 |
| AT5G03340.1 | 0,609671865 | 6,92138E-06 | AtCDC48C cell division cycle 48C chr5:810091-813133 REVERSE LENGTH=810 |
| AT1G41880.1 | 0,293591078 | 8,83121E-06 | no symbol available no full name available chr1:15651585-15652427 REVERSE LENGTH=111 |
| AT3G53430.1 | 1,612619652 | 1,13657E-05 | no symbol available no full name available chr3:19809895-19810395 REVERSE LENGTH=166 |
| AT5G12030.1 | 4,29731545 | 1,28415E-05 | AT-HSP17.6A_ HSP17.6A_ HSP17.6 HEAT SHOCK PROTEIN 17.6_ heat shock protein 17.6A chr5:3884214-3884684 REVI |
| AT5G09590.1 | 1,798181223 | 1,45161E-05 | MTHSC70-2_ HSC70-5 mitochondrial HSO70 2_ HEAT SHOCK COGNATE chr5:2975721-2978508 FORWARD LENGTH= |
| AT5G48570.1 | 4,057212925 | 1,53127E-05 | ROF2_ ATFKBP65_ FKBP65 chr5:19690746-19693656 REVERSE LENGTH=578 |
| AT5G44020.1 | 1,506394824 | 1,80968E-05 | no symbol available no full name available chr5:17712433-17714046 FORWARD LENGTH=272 |
| AT4G04020.1 | 2,789361093 | 1,84051E-05 | FIB_ PGL35_ FIB1a plastoglobulin 35_ fibrillin 1a_ fibrillin chr4:1932161-1933546 FORWARD LENGTH=318 |
| AT5G27850.1 | 0,542215755 | 1,89295E-05 | RPL18C chr5:9873169-9874297 FORWARD LENGTH=187 |
| AT1G77510.1 | 1,348492852 | 2,18574E-05 | ATPDI6_ PDIL1-2_ ATPDIL1-2_ PDI6 PDI-like 1-2_ PROTEIN DISULFIDE ISOMERASE 6 chr1:29126742-29129433 FOR |
| AT3G13470.1 | 1,288218823 | 2,56171E-05 | CPNB2_ Cpn60beta2 chaperonin-60beta2 chr3:4389685-4392624 FORWARD LENGTH=596 |
| AT4G24190.1 | 1,797743645 | 2,85467E-05 | SHD_ HSP90_ HSP90.7_ AtHsp90-7_ AtHsp90.7 SHEPHERD_ HEAT SHOCK PROTEIN 90.7_ HEAT SHOCK PROTEIN 9 |
| AT3G48930.1 | 0,290369098 | 3,2739E-05 | EMB1080 embryo defective 1080 chr3:18141017-18142189 REVERSE LENGTH=160 |
| AT1G55490.1 | 1,298042541 | 3,30911E-05 | Cpn60beta1_ LEN1_ CPNB1_ CPN60B chaperonin-60beta1_ LESION INITIATION 1_ chaperonin 60 beta chr1:20715717-20' |
| AT1G48630.1 | 0,532166889 | 3,48036E-05 | RACK1B_ RACK1B_AT receptor for activated C kinase 1B chr1:17981977-17983268 REVERSE LENGTH=326 |
| AT2G47470.1 | 1,323557938 | 3,70759E-05 | UNE5_ MEE30_ ATPDI11_ PDI11_ ATPDIL2-1 PROTEIN DISULFIDE ISOMERASE 11_ PDI-LIKE 2-1_ ARABIDOPSIS ' |
| AT3G07110.1 | 0,238161769 | 3,91448E-05 | no symbol available no full name available chr3:2252092-2253332 FORWARD LENGTH=206 |
| AT5G59720.1 | 4,891617686 | 4,45285E-05 | HSP18.2 heat shock protein 18.2 chr5:24062632-24063117 FORWARD LENGTH=161 |
| AT1G74310.1 | 3,603144923 | 4,87937E-05 | ATHSP101_ HSP101_ HOT1 heat shock protein 101 chr1:27936715-27939862 REVERSE LENGTH=911 |
| AT5G43010.1 | 0,51179492 | 5,11735E-05 | RPT4A regulatory particle triple-A ATPase 4A chr5:17248563-17251014 REVERSE LENGTH=399 |
| AT3G12580.1 | 2,373274507 | 5,15225E-05 | HSP70_ ATHSP70_ HSC70-4 ARABIDOPSIS HEAT SHOCK PROTEIN 70_ heat shock protein 70 chr3:3991487-3993689 RI |
| AT4G23850.1 | 1,261944801 | 5,28216E-05 | LACS4 long-chain acyl-CoA synthetase 4 chr4:12403720-12408263 REVERSE LENGTH=666 |

| | | | |
|---|---|---|---|
| AT5G12020.1 | 8,316664006 | 5,78582E-05 | HSP17.6II 17.6 kDa class II heat shock protein chr5:3882409-3882876 REVERSE LENGTH=155 |
| AT3G16390.1 | 0,644921286 | 5,82838E-05 | NSP3 nitrile specifier protein 3 chr3:5562602-5564356 FORWARD LENGTH=467 |
| AT2G47110.1 | 1,390840672 | 5,8284E-05 | UBI6_ UBQ6_ RPS27aB UBIQUITIN EXTENSION PROTEIN 6_ ubiquitin 6_ Ribosomal protein S27aB chr2:1934[illegible] |
| AT5G61790.1 | 1,285036514 | 6,21454E-05 | CNX1_ ATCNX1 calnexin 1 chr5:24827394-24829642 REVERSE LENGTH=530 |
| AT3G18780.1 | 1,766681925 | 6,34978E-05 | LSR2_ ACT2_ ENL2_ DER1_ FIZ2 FRIZZY AND KINKED SHOOTS 2_ LIGHT STRESS-REGULATED 2_ DEFORMED [illegible] |
| AT3G12110.1 | 2,163629224 | 6,72383E-05 | ACT11 actin-11 chr3:3858116-3859609 FORWARD LENGTH=377 |
| AT5G06870.1 | 0,711049213 | 7,00147E-05 | PGIP2_ ATPGIP2 ARABIDOPSIS POLYGALACTURONASE INHIBITING PROTEIN 2_ polygalacturonase inhibiting protei[illegible] |
| AT5G62690.1 | 1,286349879 | 7,58289E-05 | TUB2 tubulin beta chain 2 chr5:25181560-25183501 FORWARD LENGTH=450 |
| AT4G39200.2 | 0,618161337 | 7,71671E-05 | no symbol available no full name available chr4:18257464-18258464 FORWARD LENGTH=107 |
| AT1G64190.1 | 1,648413545 | 7,87691E-05 | PGD1 6-phosphogluconate dehydrogenase 1 chr1:23825549-23827012 REVERSE LENGTH=487 |
| AT2G42600.1 | 1,203355098 | 7,89183E-05 | ATPPC2_ PPC2 phosphoenolpyruvate carboxylase 2 chr2:17734541-17738679 REVERSE LENGTH=963 |
| AT1G11840.6 | 1,252361423 | 8,31223E-05 | AtGLYI3_ GLX1_ ATGLX1 glyoxalaseI 3_ glyoxalase I homolog chr1:3995928-3997518 FORWARD LENGTH=322 |
| AT2G24020.1 | 1,590755068 | 8,41066E-05 | STIC2 Suppressor of TIC40 2 chr2:10217869-10219269 REVERSE LENGTH=182 |
| AT3G01390.1 | 3,29801401 | 8,6236E-05 | AVMA10_ VMA10 vacuolar membrane ATPase 10 chr3:150265-150922 REVERSE LENGTH=110 |
| AT1G55060.1 | 1,348854134 | 8,96148E-05 | UBQ12 ubiquitin 12 chr1:20549533-20550225 FORWARD LENGTH=230 |
| AT1G76180.1 | 0,763067967 | 9,35573E-05 | ERD14 EARLY RESPONSE TO DEHYDRATION 14 chr1:28587013-28587657 REVERSE LENGTH=185 |
| AT5G61170.1 | 0,502982348 | 9,43828E-05 | no symbol available no full name available chr5:24611158-24612202 FORWARD LENGTH=143 |
| AT3G09630.1 | 0,310135794 | 0,000102164 | SAC56 Suppressor of Acaulis 56 chr3:2953813-2955444 FORWARD LENGTH=406 |
| AT3G25230.1 | 2,089753945 | 0,000108365 | FKBP62_ ROF1_ ATFKBP62 rotamase FKBP 1_ FK506 BINDING PROTEIN 62 chr3:9188257-9191137 FORWARD LENG[illegible] |
| AT4G30270.1 | 0,47062903 | 0,000109522 | MERI5B_ XTH24_ MERI-5_ SEN4 xyloglucan endotransglucosylase/hydrolase 24_ SENESCENCE 4_ MERISTEM 5_ merist[illegible] |
| AT1G11750.1 | 1,33335134 | 0,000113348 | NCLPP1_ CLPP6_ NCLPP6 NUCLEAR-ENCODED CLPP 1_ CLP protease proteolytic subunit 6 chr1:3967609-3969535 FOR[illegible] |
| AT3G04720.1 | 1,254850423 | 0,000116193 | HEL_ PR-4_ AtPR4_ PR4 HEVEIN-LIKE_ pathogenesis-related 4 chr3:1285691-1286531 REVERSE LENGTH=212 |
| AT3G46060.1 | 0,302687522 | 0,000129208 | ARA3_ RAB8A_ RABE1c_ ARA-3_ ATRAB8A_ ATRABE1C RAB GTPase homolog 8A chr3:16917908-16919740 FORWA[illegible] |
| AT3G09350.1 | 4,175756095 | 0,0001294 | Fes1A Fes1A chr3:2871216-2873109 FORWARD LENGTH=363 |
| AT3G18130.1 | 1,85746916 | 0,000133333 | RACK1C_ RACK1C_AT receptor for activated C kinase 1C chr3:6211109-6212371 REVERSE LENGTH=326 |
| AT4G38510.1 | 0,684274282 | 0,000137407 | AtVAB2_ VAB2 V-ATPase B subunit 2 chr4:18011155-18014789 REVERSE LENGTH=487 |
| AT2G25450.1 | 0,427519085 | 0,000138715 | GSL-OH glucosinolate hydroxylase chr2:10830286-10831563 REVERSE LENGTH=359 |
| AT5G28060.1 | 0,507175461 | 0,000144494 | RPS24B chr5:10069791-10070792 REVERSE LENGTH=133 |
| ATCG00790.1 | 0,436109078 | 0,000149693 | RPL16 ribosomal protein L16 chrc:81189-82652 REVERSE LENGTH=135 |
| AT1G08360.1 | 0,761091834 | 0,000153005 | no symbol available no full name available chr1:2636231-2637694 FORWARD LENGTH=216 |
| AT2G36620.1 | 0,663529263 | 0,000157246 | RPL24A ribosomal protein L24 chr2:15350548-15351819 REVERSE LENGTH=164 |
| AT1G22780.1 | 0,708575668 | 0,000157436 | RPS18A_ PFL_ PFL1 POINTED FIRST LEAVES_ POINTED FIRST LEAVES 1_ 40S RIBOSOMAL PROTEIN S18 chr1:80[illegible] |
| AT4G21280.1 | 1,079500734 | 0,000157866 | PSBQ-1_ PSBQA_ PSBQ photosystem II subunit QA_ PHOTOSYSTEM II SUBUNIT Q-1_ PHOTOSYSTEM II SUBUNIT Q[illegible] |
| ATCG00750.1 | 0,5005744 | 0,000161308 | RPS11 ribosomal protein S11 chrc:78960-79376 REVERSE LENGTH=138 |
| AT5G15520.1 | 1,947309686 | 0,000163402 | no symbol available no full name available chr5:5037242-5038136 REVERSE LENGTH=143 |
| AT5G37640.1 | 1,351231744 | 0,000164919 | UBQ9 ubiquitin 9 chr5:14952782-14953750 REVERSE LENGTH=322 |
| AT3G62870.1 | 0,456292195 | 0,00016549 | no symbol available no full name available chr3:23242862-23244273 REVERSE LENGTH=256 |
| AT1G59900.1 | 1,19507259 | 0,000166487 | AT-E1 ALPHA_ E1 ALPHA_ IAR4L pyruvate dehydrogenase complex E1 alpha subunit_ IAR4-LIKE chr1:22051368-220536[illegible] |
| AT3G26450.1 | 1,466852707 | 0,000184591 | no symbol available no full name available chr3:9681593-9683299 REVERSE LENGTH=152 |

| | | | |
|---|---|---|---|
| AT5G23140.1 | 0,756230624 | 0,000187301 | CLPP2_ NCLPP7 nuclear-encoded CLP protease P7 chr5:7783811-7784826 FORWARD LENGTH=241 |
| AT1G55210.1 | 0,775462559 | 0,000195382 | no symbol available no full name available chr1:20598057-20598620 REVERSE LENGTH=187 |
| AT5G10360.1 | 0,20761497 | 0,000197664 | RPS6B_ EMB3010 Ribosomal protein small subunit 6b_ embryo defective 3010 chr5:3258734-3260142 REVERSE LENGTH= |
| AT2G30860.1 | 0,833022418 | 0,000202323 | GSTF9_ ATGSTF9_ ATGSTF7_ GLUTTR glutathione S-transferase PHI 9 chr2:13139132-13140057 FORWARD LENGTH=[illegible] |
| AT3G12145.1 | 0,618906374 | 0,00021257 | FLR1_ FTM4 FLOR1_ FLORAL TRANSITION AT THE MERISTEM4 chr3:3874764-3876075 REVERSE LENGTH=325 |
| AT2G29450.1 | 1,837348097 | 0,0002135 | AT103-1A_ ATGSTU1_ ATGSTU5_ GSTU5 glutathione S-transferase tau 5_ ARABIDOPSIS THALIANA GLUTATHIONE [illegible] |
| ATCG00900.1 | 0,442130605 | 0,000214922 | RPS7_ RPS7.1 CHLOROPLAST RIBOSOMAL PROTEIN S7 chrc:97478-97945 REVERSE LENGTH=155 |
| AT1G57720.1 | 1,330758995 | 0,000216567 | no symbol available no full name available chr1:21377873-21380114 FORWARD LENGTH=413 |
| AT2G33210.2 | 1,142243331 | 0,000219293 | HSP60_ HSP60-2 heat shock protein 60-2 chr2:14075093-14078568 REVERSE LENGTH=580 |
| AT5G56500.1 | 0,721426316 | 0,000247782 | CPNB3_ Cpn60beta3 chaperonin-60beta3 chr5:22874058-22876966 FORWARD LENGTH=597 |
| AT4G02520.1 | 1,187930219 | 0,000247827 | ATGSTF2_ GSTF2_ ATPM24.1_ GST2_ ATPM24 glutathione S-transferase PHI 2 chr4:1110673-1111531 REVERSE LENG[illegible] |
| AT1G09750.1 | 0,819979935 | 0,000261745 | no symbol available no full name available chr1:3157541-3158960 FORWARD LENGTH=449 |
| AT5G11670.1 | 1,208041781 | 0,000263289 | NADP-ME2_ ATNADP-ME2 NADP-malic enzyme 2_ Arabidopsis thaliana NADP-malic enzyme 2 chr5:3754456-3758040 FO |
| AT1G79550.1 | 1,476441854 | 0,000271067 | PGKc_ PGK_ PGK3 phosphoglycerate kinase_ phosphoglycerate kinase 3 chr1:29924347-29926295 REVERSE LENGTH=401 |
| AT4G25200.1 | 1,852207929 | 0,000280266 | ATHSP23.6-MITO_ HSP23.6-MITO mitochondrion-localized small heat shock protein 23.6 chr4:12917089-12917858 FORWA |
| AT3G09440.1 | 1,665249898 | 0,000282659 | no symbol available no full name available chr3:2903434-2905632 REVERSE LENGTH=649 |
| AT2G34480.1 | 0,318622682 | 0,000285051 | L18aB_ RPL18aB chr2:14532916-14534161 REVERSE LENGTH=178 |
| AT1G10840.1 | 1,42797165 | 0,000302961 | TIF3H1 translation initiation factor 3 subunit H1 chr1:3607885-3610299 REVERSE LENGTH=337 |
| AT5G13650.1 | 0,705932034 | 0,000306032 | SVR3 SUPPRESSOR OF VARIEGATION 3 chr5:4397821-4402364 FORWARD LENGTH=675 |
| AT5G28540.1 | 1,697946563 | 0,000308326 | BIP1 chr5:10540665-10543274 REVERSE LENGTH=669 |
| AT3G63460.2 | 1,406607504 | 0,000313038 | SEC31B chr3:23431009-23437241 REVERSE LENGTH=1102 |
| AT5G16050.1 | 1,569214876 | 0,000319215 | GRF5_ GF14 UPSILON general regulatory factor 5 chr5:5244008-5245402 REVERSE LENGTH=268 |
| AT2G17360.1 | 0,484333435 | 0,000329277 | no symbol available no full name available chr2:7546598-7548138 FORWARD LENGTH=261 |
| AT1G04270.2 | 0,628914408 | 0,00034052 | RPS15 cytosolic ribosomal protein S15 chr1:1141852-1142960 REVERSE LENGTH=151 |
| AT4G26530.1 | 0,8287062 | 0,000345658 | FBA5_ AtFBA5_ DEG22 fructose-bisphosphate aldolase 5 chr4:13391566-13392937 FORWARD LENGTH=358 |
| AT4G37925.1 | 0,669823258 | 0,000354289 | NDH-M_ NdhM subunit NDH-M of NAD(P)H:plastoquinone dehydrogenase complex_ NADH dehydrogenase-like complex M |
| AT2G28000.1 | 1,402860151 | 0,000361768 | ARC2_ CH-CPN60A_ SLP_ CPN60A_ Cpn60alpha1_ CPNA1 SCHLEPPERLESS_ chaperonin-60alpha1_ CHLOROPLAST [illegible] |
| AT3G10950.1 | 0,473878962 | 0,000364997 | no symbol available no full name available chr3:3423893-3424566 FORWARD LENGTH=92 |
| AT1G12840.1 | 1,271456877 | 0,000366376 | DET3_ ATVHA-C ARABIDOPSIS THALIANA VACUOLAR ATP SYNTHASE SUBUNIT C_ DE-ETIOLATED 3 chr1:437[illegible] |
| AT2G34810.1 | 0,678492714 | 0,000373714 | AtBBE16 chr2:14685292-14686914 FORWARD LENGTH=540 |
| AT3G16470.1 | 0,742042175 | 0,000379423 | JAL35_ AtJAC1_ JR1 jacalin-related lectin 35_ JACALIN-LECTIN LIKE 1_ JASMONATE RESPONSIVE 1 chr3:5596096-5[illegible] |
| AT5G02870.1 | 0,302431064 | 0,000383332 | RPL4 ribosomal large subunit 4 chr5:657830-659526 FORWARD LENGTH=407 |
| AT3G25660.1 | 0,036174789 | 0,000402316 | no symbol available no full name available chr3:9339640-9342044 REVERSE LENGTH=537 |
| AT2G06850.1 | 1,341059517 | 0,000427883 | EXT_ XTH4_ EXGT-A1 ENDOXYLOGLUCAN TRANSFERASE_ endoxyloglucan transferase A1_ xyloglucan endotransgluc |
| AT1G25490.1 | 1,386291616 | 0,000429028 | EER1_ ATB BETA BETA_ RCN1_ REGA ROOTS CURL IN NPA_ ENHANCED ETHYLENE RESPONSE 1 chr1:8951700- |
| AT2G36460.1 | 1,398112707 | 0,000429805 | FBA6 fructose-bisphosphate aldolase 6 chr2:15296929-15298387 REVERSE LENGTH=358 |
| AT2G37190.1 | 0,774359539 | 0,000430256 | no symbol available no full name available chr2:15619559-15620059 REVERSE LENGTH=166 |
| AT4G34450.1 | 1,487429814 | 0,000436886 | gamma2-COP gamma2 Coat Protein chr4:16471956-16476795 FORWARD LENGTH=886 |
| AT5G10920.1 | 1,602598018 | 0,000442565 | no symbol available no full name available chr5:3441805-3443892 FORWARD LENGTH=517 |

| | | | |
|---|---|---|---|
| AT2G21580.2 | 0,714901508 | 0,000469212 | no symbol available no full name available chr2:9236629-9237510 FORWARD LENGTH=107 |
| AT5G10160.1 | 1,319295581 | 0,000479226 | no symbol available no full name available chr5:3185819-3187159 FORWARD LENGTH=219 |
| AT5G39320.1 | 0,527067689 | 0,000523673 | UDG4 UDP-glucose dehydrogenase 4 chr5:15743254-15744696 FORWARD LENGTH=480 |
| AT1G04410.1 | 1,084256769 | 0,000528748 | c-NAD-MDH1 cytosolic-NAD-dependent malate dehydrogenase 1 chr1:1189418-1191267 REVERSE LENGTH=332 |
| AT3G24830.1 | 0,293000793 | 0,000544572 | no symbol available no full name available chr3:9064613-9065871 FORWARD LENGTH=206 |
| AT5G12250.1 | 0,700140905 | 0,000554676 | TUB6 beta-6 tubulin chr5:3961317-3962971 REVERSE LENGTH=449 |
| AT3G12050.2 | 2,2920467 | 0,000565367 | no symbol available no full name available chr3:3839289-3841303 FORWARD LENGTH=321 |
| AT2G34420.1 | 1,444867524 | 0,000567947 | LHCB1.5_ LHB1B2 PHOTOSYSTEM II LIGHT HARVESTING COMPLEX GENE 1.5_ photosystem II light harvesting com |
| AT1G37130.1 | 0,726160007 | 0,00061611 | B29_ ATNR2_ NIA2_ NIA2-1_ NR2_ CHL3_ NR NITRATE REDUCTASE 2_ CHLORATE RESISTANT 3_ ARABIDOPSIS |
| AT1G48600.1 | 1,610405388 | 0,000633597 | AtPMT2_ AtPMEAMT_ PMEAMT phosphoethanolamine N-methyltransferase_ Phosphoethanolamine methyltransferase2 chr1 |
| AT3G52880.1 | 1,385747342 | 0,000636986 | ATMDAR1_ MDAR1 monodehydroascorbate reductase 1 chr3:19601477-19604366 REVERSE LENGTH=434 |
| AT3G25520.2 | 0,666022015 | 0,000640255 | RPL5A_ ATL5_ PGY3_ OLI5 ribosomal protein L5_ RIBOSOMAL PROTEIN L5 A_ PIGGYBACK3_ OLIGOCELLULA 5 c |
| AT5G27470.1 | 1,136688736 | 0,000642877 | no symbol available no full name available chr5:9695087-9697154 FORWARD LENGTH=451 |
| AT4G02080.1 | 0,61991415 | 0,000646136 | ATSARA1C_ SAR2_ SAR1C_ ATSAR2_ ASAR1 secretion-associated RAS super family 2 chr4:921554-922547 FORWARD L |
| AT2G46820.1 | 0,832707464 | 0,000673701 | PTAC8_ PSI-P_ PSAP_ CURT1B_ TMP14 CURVATURE THYLAKOID 1B_ PLASTID TRANSCRIPTIONALLY ACTIVE |
| AT5G40770.1 | 1,237227618 | 0,000682052 | ATPHB3_ PHB3_ EER3 prohibitin 3 chr5:16315589-16316621 REVERSE LENGTH=277 |
| AT1G79230.3 | 1,165021259 | 0,000691332 | ATMST1_ STR1_ MST1_ ATRDH1_ ST1 ARABIDOPSIS THALIANA RHODANESE HOMOLOGUE 1_ mercaptopyruvate |
| AT3G05590.1 | 0,3668428 | 0,000700235 | RPL18 ribosomal protein L18 chr3:1621511-1622775 FORWARD LENGTH=187 |
| AT5G16130.1 | 0,773483475 | 0,000701641 | no symbol available no full name available chr5:5268984-5269912 FORWARD LENGTH=190 |
| AT2G33380.1 | 2,443678161 | 0,000720171 | AtRD20_ PXG3_ CLO-3_ CLO3_ RD20_ AtCLO3 caleosin 3_ Arabidopsis thaliana caleosin 3_ peroxygenase 3_ RESPONSIV |
| AT2G39330.1 | 0,781586479 | 0,000760655 | JAL23 jacalin-related lectin 23 chr2:16419787-16421573 REVERSE LENGTH=459 |
| AT3G54210.1 | 0,297863772 | 0,000761526 | PRPL17 plastid ribosomal proteins of the 50S subunit 17 chr3:20067672-20068385 REVERSE LENGTH=211 |
| AT5G19940.1 | 1,164241447 | 0,000771202 | FBN6_ PAP8 FIBRILLIN6_ Probable Plastid-Lipid Associated Protein chr5:6739693-6740661 FORWARD LENGTH=239 |
| AT5G45390.1 | 1,186823826 | 0,000782104 | CLPP4_ NCLPP4 NUCLEAR-ENCODED CLP PROTEASE P4_ CLP protease P4 chr5:18396351-18397586 FORWARD LEN |
| AT3G26650.1 | 1,076914063 | 0,000809191 | GAPA_ GAPA1_ GAPA-1 glyceraldehyde 3-phosphate dehydrogenase A subunit_ GLYCERALDEHYDE 3-PHOSPHATE DE |
| AT1G76160.1 | 5,233595037 | 0,000822539 | sks5 SKU5 similar 5 chr1:28578211-28581020 REVERSE LENGTH=541 |
| AT5G13490.1 | 1,819674294 | 0,000824053 | AAC2 ADP/ATP carrier 2 chr5:4336034-4337379 FORWARD LENGTH=385 |
| AT3G04920.2 | 0,43544436 | 0,000860354 | no symbol available no full name available chr3:1360989-1361719 FORWARD LENGTH=112 |
| AT5G67360.1 | 0,786543308 | 0,00086295 | ARA12_ SBT1.7 Subtilisin-like Serine protease 1.7 chr5:26872192-26874465 REVERSE LENGTH=757 |
| AT1G73600.1 | 0,570202316 | 0,000863629 | DEG26_ NMT_ AtPMT3_ NMT3 Phosphoethanolamine methyltransferase3 chr1:27670825-27673400 FORWARD LENGTH= |
| AT3G53610.1 | 0,616397254 | 0,000904098 | RAB8B_ RAB8_ AtRab8B_ ATRAB8_ AtRABE1a RAB GTPase homolog 8 chr3:19876531-19878264 REVERSE LENGTH= |
| AT3G46230.1 | 1,409997373 | 0,00090554 | HSP17.4_ ATHSP17.4 heat shock protein 17.4_ ARABIDOPSIS THALIANA HEAT SHOCK PROTEIN 17.4 chr3:16984263- |
| AT1G32990.1 | 0,894748359 | 0,000916274 | PRPL11 plastid ribosomal protein l11 chr1:11955827-11957139 FORWARD LENGTH=222 |
| AT5G13630.1 | 0,675709458 | 0,000916782 | ABAR_ CHLH_ GUN5_ CCH_ CCH1 ABA-BINDING PROTEIN_ H SUBUNIT OF MG-CHELATASE_ GENOMES UNCO |
| AT5G14320.1 | 0,754647067 | 0,000942752 | EMB3137 EMBRYO DEFECTIVE 3137 chr5:4617839-4618772 REVERSE LENGTH=169 |
| AT3G21370.1 | 1,172855647 | 0,000946248 | BGLU19 beta glucosidase 19 chr3:7524286-7527579 REVERSE LENGTH=527 |
| AT2G37170.1 | 1,840406789 | 0,000950305 | PIP2;2_ PIP2B plasma membrane intrinsic protein 2_ PLASMA MEMBRANE INTRINSIC PROTEIN 2;2 chr2:15613624-1561 |
| AT1G66580.1 | 0,369621877 | 0,000956035 | SAG24_ RPL10C senescence associated gene 24_ ribosomal protein L10 C chr1:24839208-24840439 FORWARD LENGTH=2 |
| AT5G59370.1 | 1,458353267 | 0,000964005 | ACT4 actin 4 chr5:23950109-23951586 FORWARD LENGTH=377 |

| | | | |
|---|---|---|---|
| AT3G08590.1 | 1,356208065 | 0,000980027 | iPGAM2 "2_3-biphosphoglycerate-independent phosphoglycerate mutase 2" chr3:2608683-2611237 REVERSE LENGTH=560 |
| AT3G44890.1 | 0,686817792 | 0,000996517 | RPL9 ribosomal protein L9 chr3:16386505-16387963 FORWARD LENGTH=197 |
| AT1G78850.1 | 2,017510937 | 0,001004307 | MBL1_ GAL1 apple domain lectin-1_ Mannose Binding Lectin1 chr1:29642072-29643397 REVERSE LENGTH=441 |
| AT5G48880.1 | 0,713528586 | 0,001007546 | PKT2_ PKT1_ KAT5 3-KETO-ACYL-COENZYME A THIOLASE 5_ peroxisomal 3-keto-acyl-CoA thiolase 2_ PEROXISOM |
| AT5G14660.1 | 0,706047374 | 0,001011101 | ATDEF2_ DEF2_ PDF1B peptide deformylase 1B chr5:4727129-4728671 REVERSE LENGTH=273 |
| AT3G15360.1 | 1,188841957 | 0,001017277 | ATM4_ TRX-M4_ ATHM4 ARABIDOPSIS THIOREDOXIN M-TYPE 4_ thioredoxin M-type 4 chr3:5188448-5189457 FOR |
| AT3G24503.1 | 1,348559497 | 0,001038434 | REF1_ ALDH2C4_ ALDH1A REDUCED EPIDERMAL FLUORESCENCE1_ aldehyde dehydrogenase 2C4_ aldehyde dehydr |
| AT1G12240.1 | 0,823625645 | 0,001048364 | AtVI2_ VAC-INV_ ATBETAFRUCT4_ VI2_ FRUCT4_ AtFRUCT4_ VIN2 VACUOLAR INVERTASE_ vacuolar invertase 2 |
| AT1G05010.1 | 0,371672309 | 0,001071522 | EFE_ ACO4_ EAT1 ethylene forming enzyme_ ethylene-forming enzyme chr1:1431419-1432695 REVERSE LENGTH=323 |
| AT1G52000.1 | 0,680718463 | 0,001074125 | no symbol available no full name available chr1:19333352-19335700 REVERSE LENGTH=730 |
| AT4G12400.2 | 8,327711889 | 0,001077305 | Hop3 Hop3 chr4:7338866-7341239 REVERSE LENGTH=558 |
| AT1G56340.1 | 1,373900128 | 0,00110197 | AtCRT1a_ CRT1_ CRT1a calreticulin 1a_ calreticulin 1 chr1:21090059-21092630 REVERSE LENGTH=425 |
| AT3G11830.1 | 1,207459473 | 0,001117871 | CCT7 Chaperonin containing T-complex polypeptide-1 subunit 7 chr3:3732734-3736156 FORWARD LENGTH=557 |
| AT1G52570.1 | 0,773818413 | 0,001124459 | PLDALPHA2 phospholipase D alpha 2 chr1:19583940-19586551 REVERSE LENGTH=810 |
| AT5G50950.3 | 0,478695843 | 0,001126299 | FUM2 FUMARASE 2 chr5:20731191-20733636 FORWARD LENGTH=317 |
| AT1G62660.1 | 0,70947631 | 0,001143242 | VI1 VACUOLAR INVERTASE 1 chr1:23199949-23203515 FORWARD LENGTH=648 |
| AT5G57350.3 | 0,489368663 | 0,001208638 | ATAHA3_ HA3_ AHA3 H(+)-ATPase 3_ ARABIDOPSIS THALIANA ARABIDOPSIS H(+)-ATPASE chr5:23231208-23234 |
| AT3G19170.1 | 1,086485903 | 0,00123154 | ATZNMP_ ATPREP1_ PREP1 presequence protease 1 chr3:6625578-6631874 REVERSE LENGTH=1080 |
| AT4G08900.1 | 0,398621422 | 0,001243989 | ARGAH1 arginine amidohydrolase 1 chr4:5703499-5705180 FORWARD LENGTH=342 |
| AT5G49460.1 | 1,490470812 | 0,001245886 | ACLB-2 ATP citrate lyase subunit B 2 chr5:20055048-20058195 FORWARD LENGTH=608 |
| AT5G52650.1 | 1,870461817 | 0,001251787 | no symbol available no full name available chr5:21355781-21357003 REVERSE LENGTH=179 |
| AT3G23990.1 | 1,428862756 | 0,001257942 | HSP60_ HSP60-3B heat shock protein 60_ HEAT SHOCK PROTEIN 60-3B chr3:8669013-8672278 FORWARD LENGTH=5 |
| AT4G27520.1 | 0,80767335 | 0,001278784 | ENODL2_ AtENODL2 early nodulin-like protein 2 chr4:13750668-13751819 REVERSE LENGTH=349 |
| AT5G19510.1 | 0,711884211 | 0,001284483 | no symbol available no full name available chr5:6581854-6583137 REVERSE LENGTH=224 |
| AT1G75040.1 | 1,475916038 | 0,001322651 | PR-5_ PR5 pathogenesis-related gene 5 chr1:28177754-28178731 FORWARD LENGTH=239 |
| AT1G73060.1 | 1,589733826 | 0,001330524 | LPA3 Low PSII Accumulation 3 chr1:27479027-27481258 FORWARD LENGTH=358 |
| AT1G02560.1 | 1,182878091 | 0,001336736 | NCLPP1_ NCLPP5_ CLPP5 NUCLEAR CLPP 5_ NUCLEAR-ENCODED CLPP 1_ nuclear encoded CLP protease 5 chr1:538 |
| AT1G71500.1 | 0,666037504 | 0,001363606 | PSB33_ LIL8 PhotoSystem B protein 33_ Light-harvesting-like 8 chr1:26936084-26937331 FORWARD LENGTH=287 |
| AT3G43980.1 | 0,668662648 | 0,001387817 | no symbol available no full name available chr3:15778555-15779235 REVERSE LENGTH=56 |
| AT5G15200.1 | 0,549062866 | 0,001398325 | no symbol available no full name available chr5:4935124-4936334 REVERSE LENGTH=198 |
| AT1G55260.2 | 1,802764687 | 0,001400458 | LTPG6 glycosylphosphatidylinositol-anchored lipid protein transfer 6 chr1:20614663-20616158 FORWARD LENGTH=224 |
| AT4G24280.1 | 1,204489249 | 0,001419077 | cpHsc70-1 chloroplast heat shock protein 70-1 chr4:12590094-12593437 FORWARD LENGTH=718 |
| AT2G14260.2 | 1,435482394 | 0,001428035 | PIP_ PAP1 proline iminopeptidase_ prolyl aminopeptidase 1 chr2:6041441-6043475 REVERSE LENGTH=329 |
| AT3G28220.1 | 0,738081539 | 0,00143363 | no symbol available no full name available chr3:10524420-10526497 FORWARD LENGTH=370 |
| AT5G25980.2 | 0,841310901 | 0,001445853 | BGLU37_ TGG2 BETA GLUCOSIDASE 37_ glucoside glucohydrolase 2 chr5:9072730-9075477 FORWARD LENGTH=547 |
| AT3G26070.1 | 0,445430956 | 0,001447935 | FBN3a FIBRILLIN3a chr3:9526904-9528199 FORWARD LENGTH=242 |
| AT1G02780.1 | 0,587407967 | 0,001493267 | emb2386 embryo defective 2386 chr1:608120-609391 REVERSE LENGTH=214 |
| AT5G39740.1 | 0,708292712 | 0,001502069 | RPL5B_ OLI7 ribosomal protein L5 B_ OLIGOCELLULA 7 chr5:15903365-15905185 FORWARD LENGTH=301 |
| AT5G14920.1 | 2,324989801 | 0,001519022 | GASA14 A-stimulated in Arabidopsis 14 chr5:4826598-4827761 FORWARD LENGTH=275 |

| | | | |
|---|---|---|---|
| AT4G21960.1 | 0,378599759 | 0,001522268 | PRXR1 chr4:11646613-11648312 REVERSE LENGTH=330 |
| AT3G12290.1 | 1,413627617 | 0,001531092 | MTHFD1 methylenetetrahydrofolate dehydrogenase/methenyltetrahydrofolate cyclohydrolase chr3:3919591-3921326 FORWAR |
| AT5G02960.1 | 0,423607046 | 0,001560456 | no symbol available no full name available chr5:693280-694396 REVERSE LENGTH=142 |
| AT2G41220.1 | 0,810607014 | 0,001595109 | GLU2 glutamate synthase 2 chr2:17177934-17188388 FORWARD LENGTH=1629 |
| AT1G58270.1 | 0,789116958 | 0,001597232 | ZW9 chr1:21612394-21614089 REVERSE LENGTH=396 |
| AT1G17290.1 | 1,20251817 | 0,001614433 | AlaAT1 alanine aminotransferase chr1:5922771-5926093 FORWARD LENGTH=543 |
| AT1G56330.1 | 1,502938854 | 0,001646544 | ATSARA1B_ SAR1B_ SAR1_ ATSAR1B_ ATSAR1 SECRETION-ASSOCIATED RAS 1_ ARABIDOPSIS THALIANA SEC |
| AT2G28950.1 | 0,802078471 | 0,001647717 | ATEXP6_ ATHEXP ALPHA 1.8_ ATEXPA6_ EXPA6 expansin A6_ ARABIDOPSIS THALIANA TEXPANSIN 6 chr2:1243 |
| AT1G27400.1 | 0,630108638 | 0,001670687 | no symbol available no full name available chr1:9515230-9516725 FORWARD LENGTH=176 |
| AT1G08830.1 | 0,473805005 | 0,001740814 | CSD1_ AtSOD1_ SOD1 superoxide dismutase 1_ copper/zinc superoxide dismutase 1 chr1:2827700-2829053 FORWARD LEN |
| AT5G03630.1 | 1,247626166 | 0,00181763 | MDAR2 chr5:922378-924616 REVERSE LENGTH=435 |
| AT2G22230.1 | 1,261245485 | 0,00184903 | no symbol available no full name available chr2:9450042-9451427 FORWARD LENGTH=220 |
| AT5G19550.1 | 1,315322706 | 0,001852531 | AAT2_ ASP2 ASPARTATE AMINOTRANSFERASE 2_ aspartate aminotransferase 2 chr5:6598201-6601597 FORWARD LE |
| AT2G05920.1 | 0,556753902 | 0,00186343 | SBT1.8 subtilase 1.8 chr2:2269831-2272207 REVERSE LENGTH=754 |
| AT5G24780.1 | 0,767560332 | 0,001863946 | VSP1_ ATVSP1 vegetative storage protein 1 chr5:8507783-8508889 REVERSE LENGTH=270 |
| AT5G45280.1 | 0,468255955 | 0,0019081 | PAE11 pectin acetylesterase 11 chr5:18346862-18349432 FORWARD LENGTH=370 |
| AT2G21260.1 | 3,086288205 | 0,001950891 | no symbol available no full name available chr2:9105693-9107308 REVERSE LENGTH=309 |
| AT5G46290.3 | 1,525572583 | 0,001994237 | KASI_ KAS1 3-ketoacyl-acyl carrier protein synthase I_ KETOACYL-ACP SYNTHASE 1 chr5:18774439-18776629 REVERS |
| AT1G26570.1 | 0,83173895 | 0,002005814 | UGD1_ ATUGD1 UDP-GLUCOSE DEHYDROGENASE 1_ UDP-glucose dehydrogenase 1 chr1:9182801-9184246 FORWAR |
| AT3G55610.1 | 1,07406566 | 0,002075613 | P5CS2 delta 1-pyrroline-5-carboxylate synthase 2 chr3:20624278-20628989 REVERSE LENGTH=726 |
| AT1G29660.1 | 0,738647092 | 0,002129777 | GGL5 chr1:10371955-10373624 FORWARD LENGTH=364 |
| AT1G05190.1 | 0,835796469 | 0,002200645 | RPL6_ EMB2394 embryo defective 2394 chr1:1502515-1503738 REVERSE LENGTH=223 |
| ATCG00650.1 | 0,540043852 | 0,002243932 | RPS18 ribosomal protein S18 chrc:67917-68222 FORWARD LENGTH=101 |
| AT3G15190.1 | 0,304138783 | 0,002312687 | PRPS20 plastid ribosomal protein S20 chr3:5116216-5117412 FORWARD LENGTH=202 |
| AT5G14780.1 | 0,694635294 | 0,002375031 | FDH_ AtFDH1 formate dehydrogenase chr5:4777043-4779190 FORWARD LENGTH=384 |
| AT2G01250.1 | 0,774637506 | 0,002430057 | RPL7B chr2:132943-134264 REVERSE LENGTH=242 |
| AT1G36240.1 | 1,144272081 | 0,002463339 | RPL30A chr1:13614890-13616233 FORWARD LENGTH=112 |
| AT2G18450.1 | 1,422421032 | 0,002470215 | SDH1-2 succinate dehydrogenase 1-2 chr2:7997510-8000801 REVERSE LENGTH=632 |
| AT1G27950.1 | 1,989780835 | 0,00247067 | LTPG1 glycosylphosphatidylinositol-anchored lipid protein transfer 1 chr1:9740740-9741991 FORWARD LENGTH=193 |
| AT1G79930.1 | 1,370298321 | 0,002544883 | HSP91_ AtHsp70-14 heat shock protein 91 chr1:30063781-30067067 REVERSE LENGTH=831 |
| AT3G54640.1 | 0,273135399 | 0,002548198 | TRP3_ TSA1 TRYPTOPHAN-REQUIRING 3_ tryptophan synthase alpha chain chr3:20223331-20225303 REVERSE LENGT |
| AT1G76080.1 | 1,122772257 | 0,002574642 | CDSP32_ TRXL1_ ATCDSP32 ARABIDOPSIS THALIANA CHLOROPLASTIC DROUGHT-INDUCED STRESS PROTEIN |
| AT1G49760.1 | 0,749371163 | 0,002591975 | PABP8_ PAB8 POLY(A) BINDING PROTEIN 8_ poly(A) binding protein 8 chr1:18416740-18419753 FORWARD LENGTH |
| AT3G60770.1 | 0,329428169 | 0,002606333 | no symbol available no full name available chr3:22460525-22461656 REVERSE LENGTH=151 |
| ATCG01060.1 | 1,14348634 | 0,002633274 | PSAC chrc:117318-117563 REVERSE LENGTH=81 |
| AT1G03630.1 | 1,187655888 | 0,00270117 | POR C_ PORC protochlorophyllide oxidoreductase C chr1:907699-909245 FORWARD LENGTH=401 |
| AT5G66510.1 | 0,553483913 | 0,002714045 | GAMMA CA3 gamma carbonic anhydrase 3 chr5:26550016-26551496 REVERSE LENGTH=258 |
| AT5G27640.1 | 1,369746095 | 0,002714332 | ATTIF3B1_ TIF3B1_ EIF3B_ ATEIF3B-1_ EIF3B-1 ARABIDOPSIS THALIANA TRANSLATION INITIATION FACTOR |
| AT4G09040.1 | 0,63806475 | 0,002722043 | CP33C chr4:5795075-5797315 REVERSE LENGTH=304 |

| | | | |
|---|---|---|---|
| AT3G52580.1 | 0,693889192 | 0,002727343 | no symbol available no full name available chr3:19503324-19504701 FORWARD LENGTH=150 |
| AT3G13750.1 | 0,57260326 | 0,002742417 | BGAL1 beta galactosidase 1_ beta-galactosidase 1 chr3:4511192-4515756 FORWARD LENGTH=847 |
| AT3G12260.1 | 0,741461454 | 0,002743562 | NDUFA6_ B14 chr3:3909252-3910337 REVERSE LENGTH=133 |
| AT1G09270.1 | 1,232179945 | 0,00275647 | IMPA-4 importin alpha isoform 4 chr1:2994506-2997833 FORWARD LENGTH=538 |
| AT5G66120.2 | 0,57601187 | 0,002783622 | no symbol available no full name available chr5:26431516-26433649 REVERSE LENGTH=442 |
| AT5G47190.1 | 1,808246502 | 0,002807361 | PRPL19 plastid ribosomal proteins of the 50S subunit 19 chr5:19164432-19166064 REVERSE LENGTH=229 |
| AT3G10920.1 | 1,241204058 | 0,00282018 | ATMSD1_ MEE33_ MSD1_ AtSOD1 ARABIDOPSIS MANGANESE SUPEROXIDE DISMUTASE 1_ superoxide dismutase |
| AT1G52400.1 | 0,771699674 | 0,002827651 | BGL1_ ATBG1_ BGLU18 A. THALIANA BETA-GLUCOSIDASE 1_ BETA-GLUCOSIDASE HOMOLOG 1_ beta glucosida |
| AT4G12800.1 | 0,844795335 | 0,002834954 | PSAL photosystem I subunit l chr4:7521469-7522493 FORWARD LENGTH=219 |
| AT1G43670.1 | 0,942471695 | 0,002847556 | FBP_ AtcFBP_ cyfbp_ FINS1 "fructose-1_6-bisphosphatase"_ "Arabidopsis thaliana cytosolic fructose-1_6-bisphosphatase"_ F |
| AT5G08670.1 | 1,134354175 | 0,002902763 | no symbol available no full name available chr5:2818395-2821149 REVERSE LENGTH=556 |
| AT1G74040.1 | 1,408762915 | 0,00292955 | IPMS2_ IMS1_ MAML-3 SOPROPYLMALATE SYNTHASE 2_ 2-isopropylmalate synthase 1 chr1:27842258-27845566 FOR |
| AT3G08030.2 | 0,828199413 | 0,002942328 | AthA2-1 chr3:2564517-2565819 FORWARD LENGTH=323 |
| AT4G30910.1 | 1,263806663 | 0,002943407 | no symbol available no full name available chr4:15042621-15045248 REVERSE LENGTH=581 |
| AT5G02790.1 | 1,177456997 | 0,002965553 | GSTL3 Glutathione transferase L3 chr5:632877-634858 FORWARD LENGTH=235 |
| AT2G05710.1 | 1,511435719 | 0,002990781 | ACO3 aconitase 3 chr2:2141591-2146350 FORWARD LENGTH=990 |
| AT1G01320.2 | 1,275590396 | 0,003000066 | REC1_ FLL2 FLOURY ENDOSPERM LIKE 2_ REDUCED CHLOROPLAST COVERAGE chr1:121582-130099 REVERSE |
| AT5G51440.1 | 3,209237183 | 0,003016455 | HSP23.5 chr5:20891242-20892013 FORWARD LENGTH=210 |
| AT2G39390.1 | 0,297939513 | 0,003046596 | no symbol available no full name available chr2:16450803-16451762 REVERSE LENGTH=123 |
| AT3G09790.1 | 0,740612875 | 0,003076995 | UBQ8 ubiquitin 8 chr3:3004111-3006006 REVERSE LENGTH=631 |
| AT5G22800.1 | 1,171751637 | 0,00314655 | EMB263_ EMB1030_ EMB86 EMBRYO DEFECTIVE 263_ EMBRYO DEFECTIVE 1030_ EMBRYO DEFECTIVE 86 chr5 |
| AT1G04480.1 | 0,821338309 | 0,003149842 | no symbol available no full name available chr1:1216110-1217257 FORWARD LENGTH=140 |
| AT2G43030.1 | 0,641502861 | 0,00317864 | PRPL3 plastid ribosomal proteins of the 50S subunit chr2:17894898-17895713 FORWARD LENGTH=271 |
| AT2G31790.1 | 1,250420449 | 0,003226961 | no symbol available no full name available chr2:13518269-13520167 FORWARD LENGTH=457 |
| ATCG00905.1 | 0,546658817 | 0,003235274 | RPS12_ RPS12C RIBOSOMAL PROTEIN S12_ ribosomal protein S12C chrc:97999-98793 REVERSE LENGTH=85 |
| AT4G32260.1 | 1,222638646 | 0,003252554 | PDE334 PIGMENT DEFECTIVE 334 chr4:15573859-15574586 REVERSE LENGTH=219 |
| AT5G15650.1 | 0,877971456 | 0,003257237 | MUR5_ ATRGP2_ RGP2 MURUS 5_ REVERSIBLY GLYCOSYLATED POLYPEPTIDE 2_ reversibly glycosylated polypept |
| AT5G11520.1 | 0,819121033 | 0,003259237 | YLS4_ ASP3 aspartate aminotransferase 3_ YELLOW-LEAF-SPECIFIC GENE 4 chr5:3685257-3687721 REVERSE LENGTH |
| AT3G53990.2 | 1,077083998 | 0,003272175 | AtUSP_ USP17 Universal stress protein chr3:19990558-19991019 REVERSE LENGTH=126 |
| AT1G13930.1 | 1,707277516 | 0,003308172 | no symbol available no full name available chr1:4761091-4761558 FORWARD LENGTH=155 |
| ATCG00830.1 | 0,131449614 | 0,003318156 | RPL2.1 ribosomal protein L2 chrc:84337-85843 REVERSE LENGTH=274 |
| AT3G56070.1 | 0,794242441 | 0,003337434 | ROC2 rotamase cyclophilin 2 chr3:20806987-20807517 REVERSE LENGTH=176 |
| AT2G32920.1 | 2,003666425 | 0,003360719 | PDIL2-3_ PDI9_ ATPDIL2-3_ ATPDI9 ARABIDOPSIS THALIANA PROTEIN DISULFIDE ISOMERASE 9_ PDI-like 2-3_ |
| AT1G22530.1 | 0,816050867 | 0,003412952 | PATL2 PATELLIN 2 chr1:7955773-7958326 REVERSE LENGTH=683 |
| AT2G43910.2 | 0,831400656 | 0,003417943 | ATHOL1_ HOL1 HARMLESS TO OZONE LAYER 1 chr2:18184831-18186951 REVERSE LENGTH=217 |
| AT1G29150.1 | 1,309729603 | 0,003419472 | RPN6_ ATS9 non-ATPase subunit 9_ REGULATORY PARTICLE NON-ATPASE 6 chr1:10181240-10182499 FORWARD L |
| AT3G08940.2 | 0,890163118 | 0,003442791 | LHCB4.2 light harvesting complex photosystem II chr3:2717717-2718665 FORWARD LENGTH=287 |
| AT4G00100.1 | 0,357974388 | 0,003447368 | RPS13_ RPS13A_ PFL2_ ATRPS13A ribosomal protein S13A_ POINTED FIRST LEAF 2 chr4:37172-38123 FORWARD LE |
| AT5G47870.1 | 1,799003368 | 0,003457776 | RAD52-2_ ODB2_ RAD52-2B radiation sensitive 52-2_ Organellar DNA-Binding protein 2 chr5:19384555-19385808 REVERS |

| | | | |
|---|---|---|---|
| AT5G43940.1 | 1,07016253 | 0,003458732 | ADH2_ HOT5_ ATGSNOR1_ PAR2_ GSNOR PARAQUAT RESISTANT 2_ S-NITROSOGLUTATHIONE REDUCTASE_ |
| AT5G40950.1 | 0,838485238 | 0,00347266 | PRPL27_ RPL27 ribosomal protein large subunit 27 chr5:16410866-16411845 FORWARD LENGTH=198 |
| AT3G19710.1 | 1,556448943 | 0,003485605 | BCAT4 branched-chain aminotransferase4 chr3:6847202-6849429 REVERSE LENGTH=354 |
| AT5G27770.1 | 0,857470481 | 0,003496515 | no symbol available no full name available chr5:9836166-9837113 FORWARD LENGTH=124 |
| AT2G41840.1 | 0,827249109 | 0,003526907 | no symbol available no full name available chr2:17460016-17461398 REVERSE LENGTH=285 |
| AT3G20050.1 | 1,263051061 | 0,003549141 | ATTCP-1_ TCP-1_ CCT1 Chaperonin containing T-complex polypeptide-1 subunit 1_ T-complex protein 1 alpha subunit chr3: |
| AT4G18360.1 | 0,8243284 | 0,003587794 | GOX3 glycolate oxidase 3 chr4:10146141-10148386 REVERSE LENGTH=368 |
| AT2G31610.1 | 1,063072889 | 0,003598716 | no symbol available no full name available chr2:13450384-13451669 FORWARD LENGTH=250 |
| AT1G43560.1 | 2,93444926 | 0,003625594 | Aty2_ ty2 thioredoxin Y2 chr1:16398359-16399828 REVERSE LENGTH=167 |
| AT3G02560.1 | 0,77961889 | 0,003638341 | no symbol available no full name available chr3:542341-543168 FORWARD LENGTH=191 |
| AT3G52380.1 | 0,700761305 | 0,003671099 | PDE322_ CP33 PIGMENT DEFECTIVE 322_ chloroplast RNA-binding protein 33 chr3:19421619-19422855 FORWARD LE |
| AT3G16410.1 | 1,300751595 | 0,003693931 | NSP4 nitrile specifier protein 4 chr3:5572145-5574359 FORWARD LENGTH=619 |
| AT2G39310.1 | 2,158040292 | 0,00371178 | JAL22 jacalin-related lectin 22 chr2:16414262-16416323 REVERSE LENGTH=458 |
| AT3G16530.1 | 0,569415615 | 0,003711849 | no symbol available no full name available chr3:5624586-5625416 REVERSE LENGTH=276 |
| AT2G02930.1 | 1,236560283 | 0,003791526 | ATGSTF3_ GST16_ GSTF3 GLUTATHIONE S-TRANSFERASE 16_ glutathione S-transferase F3 chr2:851348-852106 REV |
| AT1G78300.1 | 1,213247864 | 0,003832951 | GRF2_ 14-3-3OMEGA_ GF14 OMEGA general regulatory factor 2_ 14-3-3 PROTEIN G-BOX FACTOR14 OMEGA chr1:294 |
| AT3G46440.1 | 1,742932083 | 0,003861905 | UXS5 UDP-XYL synthase 5 chr3:17089268-17091611 REVERSE LENGTH=341 |
| AT2G27860.1 | 0,698463321 | 0,003870155 | AXS1 UDP-D-apiose/UDP-D-xylose synthase 1 chr2:11864684-11866843 REVERSE LENGTH=389 |
| AT1G14250.1 | 0,575027913 | 0,003889702 | no symbol available no full name available chr1:4868675-4871203 FORWARD LENGTH=488 |
| AT1G51980.1 | 1,102960627 | 0,003927086 | no symbol available no full name available chr1:19323692-19326771 REVERSE LENGTH=503 |
| AT5G20920.2 | 0,846683132 | 0,003965182 | EIF2 BETA_ EMB1401_ eIF-2bs embryo defective 1401_ eukaryotic translation initiation factor 2 beta subunit chr5:7094994-7 |
| AT2G45790.1 | 0,604960005 | 0,003968497 | PMM_ ATPMM phosphomannomutase_ PHOSPHOMANNOMUTASE chr2:18855876-18857753 FORWARD LENGTH=246 |
| AT4G37910.1 | 1,293924293 | 0,003986548 | mtHsc70-1 mitochondrial heat shock protein 70-1 chr4:17825368-17828099 REVERSE LENGTH=682 |
| AT5G37510.1 | 1,166890581 | 0,004082471 | EMB1467_ CI76 embryo defective 1467 chr5:14897490-14900352 FORWARD LENGTH=745 |
| AT3G53990.1 | 0,933141867 | 0,004116678 | AtUSP_ USP17 Universal stress protein chr3:19989658-19991019 REVERSE LENGTH=160 |
| AT3G54400.1 | 0,86346613 | 0,004126602 | no symbol available no full name available chr3:20140291-20142599 REVERSE LENGTH=425 |
| ATCG00470.1 | 0,901279273 | 0,004132376 | ATPE ATP synthase epsilon chain chrc:52265-52663 REVERSE LENGTH=132 |
| AT3G28290.1 | 0,745208404 | 0,004210272 | AT14A chr3:10547873-10549030 FORWARD LENGTH=385 |
| AT3G49010.4 | 0,452836752 | 0,00424208 | BBC1_ RSU2_ ATBBC1 40S RIBOSOMAL PROTEIN_ breast basic conserved 1 chr3:18166971-18168047 REVERSE LENG |
| AT1G79920.1 | 1,224626606 | 0,004303609 | Hsp70-15_ AtHsp70-15 heat shock protein 70-15 chr1:30059302-30062224 REVERSE LENGTH=736 |
| AT1G18500.1 | 1,245634808 | 0,004338702 | MAML-4_ IPMS1 methylthioalkylmalate synthase-like 4_ ISOPROPYLMALATE SYNTHASE 1 chr1:6369347-6372861 FOR |
| AT2G01290.1 | 1,152784602 | 0,004340313 | RPI2 ribose-5-phosphate isomerase 2 chr2:149192-149989 REVERSE LENGTH=265 |
| AT4G37800.1 | 0,776879732 | 0,004346655 | XTH7 xyloglucan endotransglucosylase/hydrolase 7 chr4:17775703-17777372 REVERSE LENGTH=293 |
| AT2G41090.1 | 0,369147133 | 0,004347201 | CML10 calmodulin like 10 chr2:17135823-17136618 FORWARD LENGTH=191 |
| AT4G34670.1 | 0,864255821 | 0,004370771 | no symbol available no full name available chr4:16548724-16550222 FORWARD LENGTH=262 |
| AT5G64040.1 | 1,487994579 | 0,00447701 | PSAN chr5:25628724-25629409 REVERSE LENGTH=171 |
| AT2G29340.2 | 1,11938604 | 0,004552274 | no symbol available no full name available chr2:12597131-12598306 FORWARD LENGTH=262 |
| AT3G03250.1 | 1,096035999 | 0,004680857 | AtUGP1_ UGP_ UGP1 UDP-glucose pyrophosphorylase_ UDP-GLUCOSE PYROPHOSPHORYLASE 1 chr3:749761-754014 |
| AT4G16830.2 | 0,872862759 | 0,00468234 | AtRGGA chr4:9470979-9472308 FORWARD LENGTH=265 |

| | | | |
|---|---|---|---|
| AT3G49910.1 | 0,35873824 | 0,00471469 | no symbol available no full name available chr3:18504311-18504751 FORWARD LENGTH=146 |
| AT3G10350.1 | 0,734423268 | 0,004732312 | AtGET3b_ GET3b Guided Entry of Tail-anchored proteins 3b chr3:3208310-3210678 FORWARD LENGTH=411 |
| AT5G42980.1 | 1,245904431 | 0,004774167 | ATH3_ TRX3_ TRXH3_ ATTRX3_ ATTRXH3 THIOREDOXIN H3_ thioredoxin 3_ thioredoxin H-type 3 chr5:17242772-172 |
| AT5G20010.1 | 1,126876018 | 0,004792689 | RAN-1_ ATRAN1_ RAN1 RAS-RELATED NUCLEAR PROTEIN_ ARABIDOPSIS THALIANA RAS-RELATED NUCLEA |
| AT3G15730.1 | 1,278141839 | 0,004906563 | PLD_ PLDALPHA1 phospholipase D alpha 1 chr3:5330835-5333474 FORWARD LENGTH=810 |
| AT5G60360.2 | 0,419658531 | 0,004909793 | AALP_ SAG2_ ALP aleurain-like protease_ SENESCENCE ASSOCIATED GENE2 chr5:24280044-24282152 FORWARD LE |
| AT1G23730.1 | 0,68828155 | 0,004924857 | ATBCA3_ BCA3 beta carbonic anhydrase 3_ BETA CARBONIC ANHYDRASE 3 chr1:8395965-8398014 FORWARD LENG |
| AT5G19760.1 | 1,167990477 | 0,004927957 | no symbol available no full name available chr5:6679591-6681845 REVERSE LENGTH=298 |
| AT4G17560.1 | 0,724156523 | 0,005086298 | no symbol available no full name available chr4:9780343-9781752 FORWARD LENGTH=225 |
| AT2G38230.1 | 0,911323169 | 0,005250723 | ATPDX1.1_ PDX1.1 pyridoxine biosynthesis 1.1_ ARABIDOPSIS THALIANA PYRIDOXINE BIOSYNTHESIS 1.1 chr2:160 |
| AT1G04710.1 | 1,532852984 | 0,005254008 | KAT1_ PKT4 3-KETO-ACYL-COA THIOLASE 1_ peroxisomal 3-ketoacyl-CoA thiolase 4 chr1:1321941-1324556 FORWAR |
| AT2G33040.1 | 1,327368286 | 0,005258439 | ATP3 gamma subunit of Mt ATP synthase chr2:14018978-14021047 REVERSE LENGTH=325 |
| AT4G29840.1 | 1,123339546 | 0,005341804 | MTO2_ TS THREONINE SYNTHASE_ METHIONINE OVER-ACCUMULATOR 2 chr4:14599434-14601014 REVERSE LE |
| AT1G47128.1 | 0,70200512 | 0,005351095 | RD21_ RD21A responsive to dehydration 21A_ responsive to dehydration 21 chr1:17283139-17285609 REVERSE LENGTH= |
| AT1G07320.3 | 0,820145208 | 0,005479563 | RPL4_ PRPL4_ EMB2784 plastid ribosomal protein L4_ ribosomal protein L4_ EMBRYO DEFECTIVE 2784 chr1:2249190-2 |
| AT1G09640.1 | 1,375659986 | 0,005504648 | no symbol available no full name available chr1:3120162-3122152 FORWARD LENGTH=414 |
| AT5G27670.1 | 0,895650272 | 0,005536912 | HTA7_ h2a.w.7 histone H2A 7 chr5:9792807-9793365 REVERSE LENGTH=150 |
| AT1G64770.1 | 1,3049268 | 0,005539516 | NDF2_ PnsB2_ NDH45 NAD(P)H DEHYDROGENASE SUBUNIT 45_ NDH-dependent cyclic electron flow 1_ Photosyntheti |
| AT3G62530.1 | 0,787933307 | 0,005553365 | no symbol available no full name available chr3:23132219-23133121 FORWARD LENGTH=221 |
| AT3G29360.1 | 1,223867258 | 0,005559907 | UGD2 UDP-glucose dehydrogenase 2 chr3:11267375-11268817 REVERSE LENGTH=480 |
| AT2G42170.1 | 0,598725655 | 0,005726362 | no symbol available no full name available chr2:17578683-17580222 FORWARD LENGTH=329 |
| AT3G45030.1 | 0,860668094 | 0,005755582 | no symbol available no full name available chr3:16471606-16472312 REVERSE LENGTH=124 |
| AT3G47370.1 | 0,860668094 | 0,005755582 | no symbol available no full name available chr3:17453671-17454437 REVERSE LENGTH=122 |
| AT1G36280.2 | 0,723161108 | 0,005783943 | no symbol available no full name available chr1:13640600-13642908 FORWARD LENGTH=519 |
| AT3G08530.1 | 1,244567593 | 0,005798998 | AtCHC2_ CHC2 clathrin heavy chain 2 chr3:2587171-2595411 REVERSE LENGTH=1703 |
| AT3G13920.4 | 1,219242993 | 0,006059439 | TIF4A1_ EIF4A1_ RH4 eukaryotic translation initiation factor 4A1 chr3:4592635-4594128 REVERSE LENGTH=407 |
| AT1G74470.1 | 0,784491794 | 0,006131893 | no symbol available no full name available chr1:27991248-27992845 FORWARD LENGTH=467 |
| AT4G28080.1 | 1,235264293 | 0,006142021 | REC2 REDUCED CHLOROPLAST COVERAGE 2 chr4:13948993-13957840 REVERSE LENGTH=1819 |
| AT5G20630.1 | 0,851135238 | 0,006148981 | GER3_ GLP3A_ ATGER3_ GLP3_ GLP3B ARABIDOPSIS THALIANA GERMIN 3_ germin 3_ GERMIN-LIKE PROTEIN |
| AT1G35580.3 | 1,474944432 | 0,006150438 | CINV1_ A/N-InvG_ NIN2 cytosolic invertase 1_ alkaline/neutral invertase G_ neutral invertase 2 chr1:13123183-13124808 RE |
| AT5G41520.1 | 1,441455559 | 0,006292646 | RPS10B ribosomal protein S10e B chr5:16609377-16610583 REVERSE LENGTH=180 |
| AT2G29550.1 | 1,334344195 | 0,006307518 | TUB7_ TBB7 tubulin beta-7 chain_ tubulin beta 7 chr2:12644258-12645932 REVERSE LENGTH=449 |
| AT1G11840.1 | 1,17535767 | 0,006417181 | AtGLYI3_ GLX1_ ATGLX1 glyoxalaseI 3_ glyoxalase I homolog chr1:3996045-3997518 FORWARD LENGTH=283 |
| AT2G19900.1 | 1,20748244 | 0,006423711 | ATNADP-ME1_ NADP-ME1 NADP-malic enzyme 1_ Arabidopsis thaliana NADP-malic enzyme 1 chr2:8592106-8595403 RE |
| AT2G12550.1 | 0,861033704 | 0,006451034 | NUB1 homolog of human NUB1 chr2:5114881-5118486 FORWARD LENGTH=562 |
| AT5G40760.1 | 1,341221371 | 0,006476449 | G6PD6 glucose-6-phosphate dehydrogenase 6 chr5:16311284-16314556 FORWARD LENGTH=515 |
| AT1G22300.1 | 1,112813911 | 0,006547962 | 14-3-3EPSILON_ GRF10_ GF14 EPSILON general regulatory factor 10_ 14-3-3 PROTEIN G-BOX FACTOR14 EPSILON chr |
| AT4G17090.1 | 1,45307187 | 0,006558799 | CT-BMY_ BMY8_ AtBAM3_ BAM3 BETA-AMYLASE 8_ BETA-AMYLASE 3_ chloroplast beta-amylase chr4:9605266-96 |
| AT1G72150.1 | 0,74192112 | 0,006613802 | PATL1 PATELLIN 1 chr1:27148558-27150652 FORWARD LENGTH=573 |

| | | | |
|---|---|---|---|
| AT5G13850.1 | 0,831783479 | 0,006656115 | NACA3 nascent polypeptide-associated complex subunit alpha-like protein 3 chr5:4471361-4472676 FORWARD LENGTH=20 |
| AT1G77940.1 | 0,353861228 | 0,00667265 | RPL30B chr1:29304116-29305288 REVERSE LENGTH=112 |
| AT1G23820.1 | 1,200800493 | 0,006675926 | SPDS1 spermidine synthase 1 chr1:8420410-8422724 FORWARD LENGTH=334 |
| AT2G20360.1 | 1,24912098 | 0,006698897 | no symbol available no full name available chr2:8786070-8789098 FORWARD LENGTH=402 |
| AT2G44120.1 | 0,877835576 | 0,00670745 | no symbol available no full name available chr2:18249227-18250402 REVERSE LENGTH=242 |
| AT3G06580.1 | 1,2416886 | 0,006715525 | GAL1_ GALK GALACTOSE KINASE 1 chr3:2049141-2051867 REVERSE LENGTH=496 |
| AT5G01410.1 | 1,158026143 | 0,006745792 | PDX1_ ATPDX1.3_ ATPDX1_ PDX1.3_ RSR4 REDUCED SUGAR RESPONSE 4_ PYRIDOXINE BIOSYNTHESIS 1.3_ A |
| AT4G22010.1 | 1,167767333 | 0,006894734 | sks4 SKU5 similar 4 chr4:11663429-11666463 FORWARD LENGTH=541 |
| AT2G37220.1 | 0,765632146 | 0,006949585 | no symbol available no full name available chr2:15634980-15636331 REVERSE LENGTH=289 |
| AT5G60600.1 | 1,156327633 | 0,006988399 | ISPG_ CSB3_ HDS_ GCPE_ CLB4 CONSTITUTIVE SUBTILISIN 3_ 4-hydroxy-3-methylbut-2-enyl diphosphate synthase_ C |
| AT4G18100.1 | 0,173236757 | 0,007031953 | no symbol available no full name available chr4:10035715-10036475 REVERSE LENGTH=133 |
| AT3G09640.1 | 0,296128121 | 0,007062214 | AtAPX2_ APX1B_ APX2 ASCORBATE PEROXIDASE 1B_ ascorbate peroxidase 2 chr3:2956301-2958163 FORWARD LEN |
| AT3G01910.1 | 1,30194834 | 0,007337109 | AtSO_ AT-SO_ SOX sulfite oxidase chr3:314919-317274 REVERSE LENGTH=393 |
| AT3G45140.1 | 0,753435455 | 0,007501252 | ATLOX2_ LOX2 ARABIODOPSIS THALIANA LIPOXYGENASE 2_ lipoxygenase 2 chr3:16525437-16529233 FORWARD |
| AT4G10320.1 | 1,387370256 | 0,007535698 | no symbol available no full name available chr4:6397526-6404509 REVERSE LENGTH=1190 |
| AT5G66570.1 | 0,890383385 | 0,007569382 | OEE1_ PSBO-1_ MSP-1_ OE33_ OEE33_ PSBO1 PS II OXYGEN-EVOLVING COMPLEX 1_ OXYGEN EVOLVING COM |
| AT2G47730.1 | 1,383285162 | 0,007580445 | GST6_ GSTF8_ ATGSTF8_ ATGSTF5 glutathione S-transferase phi 8_ Arabidopsis thaliana glutathione S-transferase phi 8_ G |
| AT1G07770.1 | 0,882683233 | 0,007648041 | RPS15A ribosomal protein S15A chr1:2408413-2409065 REVERSE LENGTH=130 |
| AT5G19370.1 | 0,45182627 | 0,007652276 | no symbol available no full name available chr5:6524247-6526629 REVERSE LENGTH=299 |
| AT5G14040.1 | 1,235742611 | 0,007654554 | MPT3_ PHT3;1 phosphate transporter 3;1_ mitochondrial phosphate transporter 3 chr5:4531059-4532965 REVERSE LENGTH |
| AT5G58070.1 | 1,890690956 | 0,0077024 | TIL_ ATTIL TEMPERATURE-INDUCED LIPOCALIN_ temperature-induced lipocalin chr5:23500512-23501156 REVERSE |
| AT3G53230.1 | 1,197767827 | 0,007749498 | AtCDC48B cell division cycle 48B chr3:19723416-19726489 FORWARD LENGTH=815 |
| AT4G33090.1 | 1,321792741 | 0,007785074 | APM1_ ATAPM1 AMINOPEPTIDASE M1_ aminopeptidase M1 chr4:15965915-15970418 REVERSE LENGTH=879 |
| AT2G18960.1 | 1,143368526 | 0,007792385 | HA1_ OST2_ AHA1_ PMA H(+)-ATPase 1_ PLASMA MEMBRANE PROTON ATPASE_ OPEN STOMATA 2 chr2:822185 |
| AT3G25760.1 | 0,920070073 | 0,007846857 | AOC1_ ERD12 early-responsive to dehydration 12_ allene oxide cyclase 1 chr3:9403972-9405105 FORWARD LENGTH=254 |
| AT3G02880.1 | 1,790790077 | 0,007853443 | KIN7 Kinase 7 chr3:634819-636982 FORWARD LENGTH=627 |
| AT2G47510.1 | 1,255764287 | 0,007858751 | FUM1 fumarase 1 chr2:19498614-19502020 FORWARD LENGTH=492 |
| AT3G58610.1 | 1,201139547 | 0,007996023 | no symbol available no full name available chr3:21671561-21674639 FORWARD LENGTH=591 |
| AT1G76010.1 | 0,645654192 | 0,008001567 | ALBA1_ Atalba1_ ALBA4 chr1:28528505-28530488 REVERSE LENGTH=350 |
| AT2G03440.1 | 1,222713308 | 0,00805634 | ATNRP1_ NRP1 nodulin-related protein 1 chr2:1039409-1039972 REVERSE LENGTH=187 |
| AT5G56030.1 | 1,402010019 | 0,008116533 | HSP81-2_ HSP90.2_ AtHsp90.2_ ERD8_ HSP81.2 EARLY-RESPONSIVE TO DEHYDRATION 8_ heat shock protein 81-2_ |
| AT3G45940.1 | 1,610018875 | 0,008137493 | no symbol available no full name available chr3:16886226-16889171 REVERSE LENGTH=868 |
| AT4G36130.1 | 0,257117007 | 0,008236819 | no symbol available no full name available chr4:17097613-17098656 FORWARD LENGTH=258 |
| AT2G45290.1 | 0,779910141 | 0,008352773 | TKL2 transketolase 2 chr2:18672737-18675589 FORWARD LENGTH=741 |
| AT4G33680.1 | 1,148557084 | 0,00835761 | AGD2 ABERRANT GROWTH AND DEATH 2_ ARF-GAP domain 2 chr4:16171847-16174630 REVERSE LENGTH=461 |
| AT2G40610.1 | 0,50150957 | 0,008511224 | ATHEXP ALPHA 1.11_ EXP8_ ATEXPA8_ ATEXP8_ EXPA8 expansin A8 chr2:16949121-16950472 REVERSE LENGTH= |
| AT1G66270.2 | 1,572432061 | 0,008556657 | BGLU21 chr1:24700110-24702995 REVERSE LENGTH=522 |
| AT4G35450.1 | 1,28609009 | 0,008791923 | AFT_ AKR2_ AKR2A ankyrin repeat-containing protein 2 chr4:16839862-16841759 FORWARD LENGTH=342 |
| AT2G43950.1 | 1,542653306 | 0,008927872 | ATOEP37_ OEP37 chloroplast outer envelope protein 37_ ARABIDOPSIS CHLOROPLAST OUTER ENVELOPE PROTEIN |

| | | | |
|---|---|---|---|
| AT5G03940.1 | 1,073423876 | 0,008931116 | SRP54CP_ 54CP_ FFC_ CPSRP54 54 CHLOROPLAST PROTEIN_ chloroplast signal recognition particle 54 kDa subunit_ SI[illegible] |
| AT2G42590.1 | 1,242762904 | 0,008965422 | GF14 MU_ GRF9_ GRF14 general regulatory factor 9 chr2:17732118-17733775 REVERSE LENGTH=263 |
| AT2G15620.1 | 0,870657288 | 0,009357521 | NIR_ ATHNIR_ NIR1 ARABIDOPSIS THALIANA NITRITE REDUCTASE_ nitrite reductase 1_ NITRITE REDUCTASE c[illegible] |
| AT5G11420.1 | 0,744621535 | 0,009396214 | no symbol available no full name available chr5:3644655-3646991 FORWARD LENGTH=366 |
| AT2G45810.1 | 1,34348023 | 0,009408716 | RH6 RNA Helicase 6 chr2:18859836-18862318 FORWARD LENGTH=528 |
| AT1G62180.1 | 0,580303283 | 0,009455091 | APSR_ PRH43_ PRH_ ATAPR2_ APR2 ADENOSINE-5'-PHOSPHOSULFATE REDUCTASE_ 3'-PHOSPHOADENOSINE-[illegible] |
| AT3G53420.1 | 1,078439829 | 0,009497383 | PIP2;1_ AtPIP2;1_ PIP2A_ PIP2 PLASMA MEMBRANE INTRINSIC PROTEIN 2_ PLASMA MEMBRANE INTRINSIC PR[illegible] |
| AT5G48375.1 | 0,841757482 | 0,009560477 | TGG3_ BGLU39 thioglucoside glucosidase 3_ BETA GLUCOSIDASE 39 chr5:19601303-19603883 REVERSE LENGTH=43[illegible] |
| AT5G12470.1 | 0,699023521 | 0,00971352 | RER4 RETICULATA-RELATED 4 chr5:4044950-4047290 REVERSE LENGTH=386 |
| AT5G54900.1 | 4,670923929 | 0,009806886 | RBP45A_ ATRBP45A RNA-binding protein 45A chr5:22295412-22298126 FORWARD LENGTH=387 |
| AT1G20340.1 | 0,875757511 | 0,010198566 | PETE2_ DRT112 DNA-DAMAGE-REPAIR/TOLERATION PROTEIN 112_ PLASTOCYANIN 2 chr1:7042770-7043273 RE[illegible] |
| AT5G17310.2 | 1,226857461 | 0,010213525 | AtUGP2_ UGP2 UDP-GLUCOSE PYROPHOSPHORYLASE 2_ UDP-glucose pyrophosphorylase 2 chr5:5696955-5700845 R[illegible] |
| AT1G30380.1 | 2,200014614 | 0,010225522 | PSAK photosystem I subunit K chr1:10722325-10723013 FORWARD LENGTH=130 |
| AT3G52750.4 | 1,526022433 | 0,010281798 | FTSZ2-2 chr3:19550926-19552435 REVERSE LENGTH=350 |
| AT5G17990.1 | 1,200964087 | 0,01034188 | pat1_ TRP1 tryptophan biosynthesis 1_ PHOSPHORIBOSYLANTHRANILATE TRANSFERASE 1 chr5:5957330-5959681 F[illegible] |
| AT5G20720.1 | 1,071738685 | 0,010342392 | ATCPN21_ CPN20_ CPN21_ CHCPN10_ CPN10 chaperonin 20_ CHLOROPLAST CHAPERONIN 10 chr5:7015015-701635[illegible] |
| AT3G52500.1 | 0,86718246 | 0,010754666 | no symbol available no full name available chr3:19465644-19467053 REVERSE LENGTH=469 |
| AT3G03960.1 | 1,370934819 | 0,010783173 | CCT8 Chaperonin containing T-complex polypeptide-1 subunit 8 chr3:1024432-1027604 FORWARD LENGTH=549 |
| AT3G19760.1 | 2,842040567 | 0,010794904 | EIF4A-III_ RH2 eukaryotic initiation factor 4A-III chr3:6863790-6866242 FORWARD LENGTH=408 |
| AT1G63660.2 | 1,557084034 | 0,010879945 | no symbol available no full name available chr1:23604874-23607080 REVERSE LENGTH=434 |
| AT1G07790.1 | 0,758771798 | 0,011044722 | HTB1 chr1:2413049-2413495 FORWARD LENGTH=148 |
| AT1G41830.1 | 1,139423845 | 0,011125531 | SKS6 SKU5-similar 6_ SKU5 SIMILAR 6 chr1:15603892-15607802 REVERSE LENGTH=542 |
| AT1G75940.1 | 1,405778601 | 0,011252867 | ATA27_ BGLU20 BETA GLUCOSIDASE 20 chr1:28511198-28514044 FORWARD LENGTH=535 |
| AT3G47070.1 | 2,210520876 | 0,011366086 | no symbol available no full name available chr3:17337205-17337507 REVERSE LENGTH=100 |
| AT1G02500.1 | 0,823528075 | 0,011447761 | METK1_ SAM-1_ AtSAM1_ SAM1_ MAT1 S-adenosylmethionine synthetase 1_ S-ADENOSYLMETHIONINE SYNTHETA[illegible] |
| AT1G58290.1 | 0,394464256 | 0,011452779 | HEMA1_ GluTR_ AtHEMA1 glutamyl-tRNA reductase_ Arabidopsis thaliana hemA 1 chr1:21624028-21626051 REVERSE L[illegible] |
| AT1G02930.1 | 0,82529781 | 0,011549066 | ATGSTF3_ ATGSTF6_ GST1_ ERD11_ GSTF6_ ATGST1 EARLY RESPONSIVE TO DEHYDRATION 11_ ARABIDOPSI[illegible] |
| AT2G41530.1 | 1,162401986 | 0,011583848 | SFGH_ ATSFGH S-formylglutathione hydrolase_ ARABIDOPSIS THALIANA S-FORMYLGLUTATHIONE HYDROLASE [illegible] |
| AT5G37600.1 | 0,829011543 | 0,011662288 | ATGSR1_ ATGLN1;1_ GLN1;1_ GSR 1 ARABIDOPSIS THALIANA GLUTAMINE SYNTHASE CLONE R1_ ARABIDOP[illegible] |
| AT2G23350.1 | 1,179460857 | 0,011696464 | PABP4_ PAB4 POLY(A) BINDING PROTEIN 4_ poly(A) binding protein 4 chr2:9943209-9946041 FORWARD LENGTH=6[illegible] |
| AT1G14320.2 | 0,384661393 | 0,011839894 | RPL10A_ SAC52_ RPL10 SUPPRESSOR OF ACAULIS 52_ ribosomal protein L10_ ribosomal protein L10 A chr1:4888650-4[illegible] |
| AT3G12390.1 | 0,852105981 | 0,011944246 | no symbol available no full name available chr3:3942344-3943595 FORWARD LENGTH=203 |
| AT5G58290.1 | 0,760993964 | 0,012082168 | RPT3 regulatory particle triple-A ATPase 3 chr5:23569155-23571116 FORWARD LENGTH=408 |
| AT5G54810.1 | 0,906363853 | 0,012141137 | TRP2_ ATTSB1_ TSB1_ TRPB tryptophan synthase beta-subunit 1_ TRYPTOPHAN BIOSYNTHESIS B_ TRYPTOPHAN B[illegible] |
| AT2G26740.1 | 1,099165867 | 0,012223202 | ATSEH_ SEH soluble epoxide hydrolase chr2:11393148-11394257 REVERSE LENGTH=321 |
| AT4G08870.1 | 0,757224904 | 0,012573809 | ARGAH2 arginine amidohydrolase 2 chr4:5646654-5648693 REVERSE LENGTH=344 |
| AT5G15970.1 | 5,668398095 | 0,012625752 | AtCor6.6_ KIN2_ COR6.6 COLD-RESPONSIVE 6.6 chr5:5211966-5212441 FORWARD LENGTH=66 |
| AT4G17470.1 | 0,742263757 | 0,012870831 | CRSH Ca2+-activated RelA-spot homolog chr4:9742922-9744468 REVERSE LENGTH=308 |
| AT1G35720.1 | 0,916822896 | 0,013084834 | ANNAT1_ ATOXY5_ OXY5_ ANN1_ AtANN1 annexin 1 chr1:13225304-13226939 FORWARD LENGTH=317 |

| | | | |
|---|---|---|---|
| AT1G32900.1 | 0,905406295 | 0,013123163 | GBSS1 granule bound starch synthase 1 chr1:11920582-11923506 REVERSE LENGTH=610 |
| AT5G62350.1 | 0,507236948 | 0,013124831 | no symbol available no full name available chr5:25037504-25038112 FORWARD LENGTH=202 |
| AT2G06050.1 | 0,890156618 | 0,013184823 | OPR3_ DDE1_ AtOPR3 DELAYED DEHISCENCE 1_ oxophytodienoate-reductase 3 chr2:2359240-2361971 REVERSE LEN |
| AT5G09900.1 | 1,313074643 | 0,013242843 | EMB2107_ MSA_ RPN5A MARIPOSA_ EMBRYO DEFECTIVE 2107_ REGULATORY PARTICLE NON-ATPASE SUBU |
| AT1G49240.1 | 1,065889805 | 0,013368786 | ACT8_ FIZ1 FRIZZY AND KINKED SHOOTS_ actin 8 chr1:18216539-18217947 FORWARD LENGTH=377 |
| AT4G25100.1 | 1,159405974 | 0,013580178 | FSD1_ ATFSD1 ARABIDOPSIS FE SUPEROXIDE DISMUTASE 1_ Fe superoxide dismutase 1 chr4:12884649-12886501 RI |
| ATCG00820.1 | 0,487920072 | 0,013582369 | RPS19 ribosomal protein S19 chrc:84005-84283 REVERSE LENGTH=92 |
| AT5G35630.1 | 0,922363051 | 0,013625123 | GLN2_ GS2_ ATGSL1 GLUTAMINE SYNTHETASE 2_ glutamine synthetase 2_ GLUTAMINE SYNTHETASE LIKE 1 chr |
| AT2G28790.1 | 0,690348482 | 0,013653588 | no symbol available no full name available chr2:12354664-12355413 REVERSE LENGTH=249 |
| AT4G25630.1 | 1,188834492 | 0,01367275 | ATFIB2_ FIB2 fibrillarin 2 chr4:13074239-13076205 FORWARD LENGTH=320 |
| AT1G31812.1 | 1,440982325 | 0,013835453 | ACBP6_ ACBP_ AtACBP6 acyl-CoA-binding protein 6_ ACYL-COA-BINDING PROTEIN chr1:11411132-11412099 REVEF |
| AT3G13930.1 | 1,187512666 | 0,013950427 | mtE2-2 mitochondrial pyruvate dehydrogenase subunit 2-2 chr3:4596240-4600143 FORWARD LENGTH=539 |
| AT2G33070.1 | 1,540574188 | 0,014222274 | ATNSP2_ NSP2 NITRILE-SPECIFIER PROTEIN 2_ nitrile specifier protein 2 chr2:14029350-14030934 REVERSE LENGTH |
| AT4G35000.1 | 1,262554937 | 0,014265458 | APX3 ascorbate peroxidase 3 chr4:16665007-16667541 REVERSE LENGTH=287 |
| AT4G27700.1 | 1,213815977 | 0,014365996 | no symbol available no full name available chr4:13826541-13827673 REVERSE LENGTH=224 |
| AT3G32980.1 | 0,696983697 | 0,01465981 | PRX32 Peroxidase 32 chr3:13526404-13529949 REVERSE LENGTH=352 |
| AT3G46780.1 | 1,098257274 | 0,014698206 | PTAC16 plastid transcriptionally active 16 chr3:17228766-17231021 FORWARD LENGTH=510 |
| AT1G80560.1 | 0,835283906 | 0,014700018 | ATIMD2_ IMD2 isopropylmalate dehydrogenase 2_ ARABIDOPSIS ISOPROPYLMALATE DEHYDROGENASE 2 chr1:302 |
| AT2G28815.1 | 1,922784579 | 0,014701292 | no symbol available no full name available chr2:12367001-12368064 REVERSE LENGTH=291 |
| AT3G16420.1 | 0,816524458 | 0,014728587 | PBPI_ JAL30_ PBP1 PYK10-binding protein 1_ JACALIN-RELATED LECTIN 30 chr3:5579560-5580674 FORWARD LENC |
| AT4G09670.1 | 1,223815187 | 0,014729263 | no symbol available no full name available chr4:6107382-6109049 REVERSE LENGTH=362 |
| AT1G54220.1 | 0,799600656 | 0,014864254 | mtE2-3 mitochondrial pyruvate dehydrogenase subunit 2-3 chr1:20246460-20250208 REVERSE LENGTH=539 |
| AT3G23570.1 | 0,820116394 | 0,015203867 | no symbol available no full name available chr3:8458052-8459608 REVERSE LENGTH=239 |
| AT2G14610.1 | 1,223884933 | 0,0152364 | PR1_ ATPR1_ AtCAPE9_ PR 1 PATHOGENESIS-RELATED GENE 1_ pathogenesis-related gene 1 chr2:6241944-6242429 F |
| AT2G20610.1 | 1,193664092 | 0,015417946 | RTY1_ RTY_ HLS3_ ALF1_ SUR1 ABERRANT LATERAL ROOT FORMATION 1_ SUPERROOT 1_ ROOTY 1_ ROOTY |
| AT1G29670.1 | 0,678893697 | 0,015991467 | GDSL1_ GGL6 chr1:10375843-10377717 FORWARD LENGTH=363 |
| AT1G56110.1 | 1,187455014 | 0,016340434 | NOP56 homolog of nucleolar protein NOP56 chr1:20984544-20986893 REVERSE LENGTH=522 |
| AT5G13030.1 | 1,508328263 | 0,016578755 | SELO SELENOPROTEIN O chr5:4133216-4136461 FORWARD LENGTH=633 |
| AT2G16600.1 | 0,865395604 | 0,016723015 | AtCYP19-1_ ROC3_ CYP19 rotamase CYP 3_ cyclophilin 19 chr2:7200862-7201383 FORWARD LENGTH=173 |
| AT5G36880.4 | 1,250396941 | 0,016764932 | ACS acetyl-CoA synthetase chr5:14535506-14539084 REVERSE LENGTH=610 |
| AT1G09080.2 | 0,757665589 | 0,016791428 | BIP3 binding protein 3 chr1:2929268-2931804 REVERSE LENGTH=665 |
| AT4G27000.1 | 0,693386132 | 0,016837625 | ATRBP45C chr4:13554983-13557763 REVERSE LENGTH=415 |
| AT5G47700.1 | 1,176505943 | 0,016888553 | RPP1C_ RPP1.3 60S acidic ribosomal protein P1-3_ RPP1 co-orthologous gene 3 chr5:19328019-19328724 REVERSE LENG |
| AT2G27710.1 | 0,798282694 | 0,016985127 | no symbol available no full name available chr2:11816929-11817670 FORWARD LENGTH=115 |
| ATCG00680.1 | 0,760385786 | 0,017008346 | PSBB photosystem II reaction center protein B chrc:72371-73897 FORWARD LENGTH=508 |
| AT5G20290.1 | 0,547637579 | 0,017158057 | no symbol available no full name available chr5:6851695-6853012 REVERSE LENGTH=222 |
| AT5G52470.1 | 0,828600472 | 0,017273345 | ATFIB1_ FBR1_ FIB1_ SKIP7_ ATFBR1 fibrillarin 1_ FIBRILLARIN 1_ SKP1/ASK1-INTERACTING PROTEIN chr5:2129 |
| AT1G09010.1 | 1,232824341 | 0,017342094 | no symbol available no full name available chr1:2895259-2899287 REVERSE LENGTH=944 |
| AT4G38740.1 | 0,9043369 | 0,017496277 | ROC1 rotamase CYP 1 chr4:18083620-18084138 REVERSE LENGTH=172 |

| | | | |
|---|---|---|---|
| AT1G13060.1 | 1,154679445 | 0,017514521 | PBE1 20S proteasome beta subunit E1 chr1:4452641-4454663 FORWARD LENGTH=274 |
| AT2G25140.1 | 1,380849396 | 0,017532836 | HSP98.7_ CLPB-M_ CLPB4 HEAT SHOCK PROTEIN 98.7_ CASEIN LYTIC PROTEINASE B-M_ casein lytic proteinase B |
| AT2G14880.1 | 1,608488838 | 0,017546662 | SWIB2 chr2:6393686-6394841 REVERSE LENGTH=141 |
| AT4G09010.2 | 1,103835115 | 0,017844201 | APX4_ TL29 ascorbate peroxidase 4_ thylakoid lumen 29 chr4:5777502-5779064 REVERSE LENGTH=284 |
| AT3G04940.1 | 1,3515451 | 0,017898991 | ATCYSD1_ CYSD1 CYSTEINE SYNTHASE D1_ cysteine synthase D1 chr3:1365681-1367508 FORWARD LENGTH=324 |
| AT3G25770.1 | 0,872336739 | 0,018039421 | AOC2 allene oxide cyclase 2 chr3:9406975-9407839 FORWARD LENGTH=253 |
| AT1G45145.1 | 1,295307119 | 0,018275057 | TRX-h5_ ATTRX5_ LIV1_ ATH5_ TRX5 thioredoxin H-type 5_ THIOREDOXIN H-TYPE 5_ LOCUS OF INSENSITIVITY |
| AT2G28190.1 | 0,737105278 | 0,018646719 | CSD2_ CZSOD2_ SOD2_ AtSOD2 COPPER/ZINC SUPEROXIDE DISMUTASE 2_ superoxide dismutase 2_ copper/zinc sup |
| AT5G03350.1 | 0,816256388 | 0,018674312 | SAI-LLP1 SA-induced legume lectin-like protein 1 chr5:815804-816628 REVERSE LENGTH=274 |
| AT1G07660.1 | 0,791026892 | 0,018717029 | no symbol available no full name available chr1:2369212-2369523 FORWARD LENGTH=103 |
| AT5G24420.1 | 0,513799087 | 0,018828 | PGL5 6-phosphogluconolactonase 5 chr5:8336943-8337879 REVERSE LENGTH=252 |
| AT2G18020.1 | 0,35628874 | 0,018929727 | EMB2296 embryo defective 2296 chr2:7837151-7838160 FORWARD LENGTH=258 |
| AT1G09620.1 | 1,189336316 | 0,019192782 | no symbol available no full name available chr1:3113077-3116455 REVERSE LENGTH=1091 |
| AT1G31330.1 | 0,92951968 | 0,019502753 | PSAF photosystem I subunit F chr1:11215011-11215939 REVERSE LENGTH=221 |
| AT2G33410.1 | 0,762799551 | 0,019573283 | RBGD2 RNA-binding glycine-rich protein D2 chr2:14156085-14157435 FORWARD LENGTH=404 |
| AT2G17390.1 | 0,793765417 | 0,019577629 | AKR2B ankyrin repeat-containing 2B chr2:7555870-7557743 FORWARD LENGTH=344 |
| AT1G33120.1 | 0,800212805 | 0,020007896 | no symbol available no full name available chr1:12010986-12012223 FORWARD LENGTH=194 |
| AT1G66970.1 | 0,853176088 | 0,020098043 | SVL2_ GDPDL1 SHV3-like 2_ Glycerophosphodiester phosphodiesterase (GDPD) like 1 chr1:24992746-24996005 REVERSE |
| AT2G18110.1 | 1,385314884 | 0,020766785 | no symbol available no full name available chr2:7872636-7873713 FORWARD LENGTH=231 |
| AT5G50950.1 | 0,586017301 | 0,020930016 | FUM2 FUMARASE 2 chr5:20729687-20733476 FORWARD LENGTH=510 |
| AT5G65430.2 | 1,099563785 | 0,02093035 | 14-3-3KAPPA_ GRF8_ AtMIN10_ GF14 KAPPA general regulatory factor 8_ 14-3-3 PROTEIN G-BOX FACTOR14 KAPPA |
| AT4G20260.1 | 0,881180413 | 0,020963073 | ATPCAP1_ PCAP1_ MDP25 ARABIDOPSIS THALIANA PLASMA-MEMBRANE ASSOCIATED CATION-BINDING PRO |
| AT5G62790.1 | 1,251152857 | 0,021095244 | PDE129_ DXR 1-deoxy-D-xylulose 5-phosphate reductoisomerase_ PIGMENT-DEFECTIVE EMBRYO 129 chr5:25214358-2 |
| AT4G25740.1 | 1,336494729 | 0,021540564 | no symbol available no full name available chr4:13107488-13108751 REVERSE LENGTH=177 |
| AT2G47400.1 | 0,913024295 | 0,021610226 | CP12_ CP12-1 CP12 DOMAIN-CONTAINING PROTEIN 1_ CP12 domain-containing protein 1 chr2:19446889-19447263 FO |
| AT5G14030.5 | 1,102167304 | 0,021623725 | no symbol available no full name available chr5:4526878-4527917 FORWARD LENGTH=159 |
| AT5G12860.2 | 0,854533798 | 0,021856395 | DiT1 dicarboxylate transporter 1 chr5:4059850-4061919 REVERSE LENGTH=556 |
| AT4G22670.1 | 1,097713785 | 0,021984085 | AtHip1_ HIP1_ TPR11 HSP70-interacting protein 1_ tetratricopeptide repeat 11 chr4:11918236-11920671 FORWARD LENGT |
| AT4G35090.1 | 1,195610393 | 0,022054468 | CAT2 catalase 2 chr4:16700937-16703215 REVERSE LENGTH=492 |
| AT3G06650.1 | 1,177361917 | 0,022179394 | ACLB-1 ATP-citrate lyase B-1 chr3:2079247-2082633 REVERSE LENGTH=608 |
| AT2G21660.1 | 0,747821452 | 0,022183584 | RBGA3_ GR-RBP7_ CCR2_ ATGRP7_ GRP7_ SRBP1 RNA-binding glycine-rich protein A3_ SMALL RNA-BINDING PRO |
| AT4G02530.1 | 1,126212256 | 0,022192452 | MPH2 MAINTENANCE OF PHOTOSYSTEM II UNDER HIGH LIGHT 2 chr4:1112335-1114005 REVERSE LENGTH=216 |
| AT1G19670.1 | 0,83235539 | 0,022387298 | CLH1_ ATCLH1_ CORI1_ ATHCOR1 chlorophyllase 1_ CORONATINE-INDUCED PROTEIN 1 chr1:6803796-6804923 RE |
| AT4G27440.1 | 0,853988736 | 0,022542304 | PORB protochlorophyllide oxidoreductase B chr4:13725648-13727107 FORWARD LENGTH=401 |
| AT3G27690.1 | 0,85250789 | 0,02277387 | LHCB2.4_ DEG13_ LHCB2_ LHCB2.3 LIGHT-HARVESTING CHLOROPHYLL B-BINDING 2_ photosystem II light harves |
| AT2G33800.1 | 0,88325784 | 0,022917082 | SCA1_ RPS5_ EMB3113_ PRPS5 plastid ribosomal protein of the 30S subunit 5_ EMBRYO DEFECTIVE 3113_ ribosomal pr |
| AT5G63980.1 | 0,680196633 | 0,023469612 | ALX8_ SUPO1_ AtFRY1_ HOS2_ ATSAL1_ SAL1_ RON1_ FRY1 HIGH EXPRESSION OF OSMOTICALLY RESPONSIV |
| AT3G44860.1 | 0,922009395 | 0,023819786 | FAMT farnesoic acid carboxyl-O-methyltransferase chr3:16379689-16380939 FORWARD LENGTH=348 |
| AT5G43850.1 | 1,168292928 | 0,023976721 | ATARD4_ ARD4 chr5:17627364-17629122 REVERSE LENGTH=187 |

| | | | |
|---|---|---|---|
| AT3G24170.1 | 1,181866547 | 0,024255441 | ATGR1_ GR1 glutathione-disulfide reductase chr3:8729762-8734115 REVERSE LENGTH=499 |
| AT5G65020.1 | 1,127176656 | 0,024293263 | ANNAT2_ AtANN2 annexin 2 chr5:25973915-25975554 FORWARD LENGTH=317 |
| AT5G13510.1 | 0,134996253 | 0,024343765 | EMB3136 EMBRYO DEFECTIVE 3136 chr5:4341294-4341956 FORWARD LENGTH=220 |
| AT1G67430.2 | 0,829996523 | 0,024480241 | no symbol available no full name available chr1:25262209-25263627 FORWARD LENGTH=131 |
| AT3G46430.1 | 1,458809741 | 0,024571399 | AtMtATP6 chr3:17087687-17088497 FORWARD LENGTH=55 |
| AT1G19570.1 | 0,880166113 | 0,02464296 | ATDHAR1_ DHAR1_ DHAR5 DEHYDROASCORBATE REDUCTASE 5_ dehydroascorbate reductase chr1:6773462-67744 |
| AT2G29630.1 | 0,604714802 | 0,025017343 | THIC_ PY PYRIMIDINE REQUIRING_ thiaminC chr2:12667395-12669569 FORWARD LENGTH=644 |
| ATCG00740.1 | 0,830973747 | 0,025076349 | RPOA RNA polymerase subunit alpha chrc:77901-78890 REVERSE LENGTH=329 |
| AT5G23120.1 | 0,933935761 | 0,025123012 | HCF136 HIGH CHLOROPHYLL FLUORESCENCE 136 chr5:7778154-7780463 FORWARD LENGTH=403 |
| ATCG00280.1 | 0,815159583 | 0,025258858 | PSBC photosystem II reaction center protein C chrc:33720-35141 FORWARD LENGTH=473 |
| AT3G13580.1 | 1,922030734 | 0,025546654 | no symbol available no full name available chr3:4433809-4435109 FORWARD LENGTH=244 |
| AT3G56190.1 | 1,184378074 | 0,025717332 | ALPHA-SNAP2_ ASNAP alpha-soluble NSF attachment protein 2 chr3:20846119-20848356 REVERSE LENGTH=289 |
| AT2G25080.1 | 1,270093102 | 0,025968258 | GPX1_ ATGPX1_ GPXL1 GLUTATHIONE PEROXIDASE 1_ glutathione peroxidase 1 chr2:10668134-10669828 FORWAR |
| AT1G09795.1 | 1,128997408 | 0,026064602 | ATATP-PRT2_ HISN1B_ ATP-PRT2 ATP phosphoribosyl transferase 2 chr1:3173588-3176690 FORWARD LENGTH=413 |
| AT5G59880.2 | 0,800961661 | 0,02611354 | ADF3 actin depolymerizing factor 3 chr5:24120382-24121628 FORWARD LENGTH=124 |
| AT5G45280.2 | 0,811691964 | 0,026274451 | PAE11 pectin acetylesterase 11 chr5:18346862-18349488 FORWARD LENGTH=391 |
| AT1G79040.1 | 0,834983797 | 0,026545311 | PSBR photosystem II subunit R chr1:29736085-29736781 FORWARD LENGTH=140 |
| AT2G26340.1 | 0,805705512 | 0,026565151 | no symbol available no full name available chr2:11215254-11216480 FORWARD LENGTH=253 |
| AT1G75950.1 | 0,401646079 | 0,026588962 | UIP1_ ASK1_ SKP1A_ ATSKP1_ SKP1 ARABIDOPSIS SKP1 HOMOLOGUE 1_ UFO INTERACTING PROTEIN 1_ S pha |
| AT3G55040.1 | 0,875502272 | 0,026906153 | GSTL2 glutathione transferase lambda 2 chr3:20398718-20400305 REVERSE LENGTH=292 |
| AT3G09260.1 | 0,684785788 | 0,026995639 | LEB_ BGLU23_ PYK10_ PSR3.1 LONG ER BODY chr3:2840657-2843730 REVERSE LENGTH=524 |
| AT3G49120.1 | 0,755072366 | 0,027008968 | PRX34_ PRXCB_ ATPCB_ AtPRX34_ PERX34_ ATPERX34 ARABIDOPSIS THALIANA PEROXIDASE CB_ PEROXIDA |
| AT1G03220.1 | 1,266038298 | 0,027111341 | SAP2 secreted aspartic protease 2 chr1:787143-788444 FORWARD LENGTH=433 |
| AT1G60950.1 | 0,606263814 | 0,027118313 | ATFD2_ FD2_ FED A FERREDOXIN 2 chr1:22444565-22445011 FORWARD LENGTH=148 |
| AT4G21210.1 | 0,521189319 | 0,02720727 | RP1_ ATRP1 PPDK regulatory protein chr4:11307002-11308587 FORWARD LENGTH=403 |
| AT3G53900.1 | 0,916918047 | 0,02723832 | PYRR_ UPP PYRIMIDINE R_ uracil phosphoribosyltransferase chr3:19956914-19958699 REVERSE LENGTH=231 |
| AT5G14300.1 | 0,628734587 | 0,027311812 | ATPHB5_ PHB5 prohibitin 5 chr5:4613102-4614023 FORWARD LENGTH=249 |
| AT3G08940.1 | 0,701232082 | 0,027346069 | LHCB4.2 light harvesting complex photosystem II chr3:2717717-2718400 FORWARD LENGTH=227 |
| AT5G23740.1 | 1,285994712 | 0,027373766 | RPS11-BETA ribosomal protein S11-beta chr5:8008251-8009330 REVERSE LENGTH=159 |
| AT1G80380.4 | 1,068013496 | 0,027478091 | GLYK glycerate kinase chr1:30217332-30219784 FORWARD LENGTH=450 |
| AT5G28840.1 | 1,086414623 | 0,027602295 | GME GDP-D-mannose 3 chr5:10862472-10864024 REVERSE LENGTH=377 |
| AT3G16400.1 | 0,855665219 | 0,02789286 | NSP1_ ATNSP1_ ATMLP-470 nitrile specifier protein 1_ NITRILE SPECIFIER PROTEIN 1_ MYROSINASE-BINDING PR |
| AT5G02160.1 | 0,497606208 | 0,028371625 | FIP FtsH5 Interacting Protein chr5:426392-427024 FORWARD LENGTH=129 |
| AT1G45000.1 | 1,151266429 | 0,028405391 | RPT4b chr1:17009220-17011607 FORWARD LENGTH=399 |
| AT1G24510.3 | 1,399900578 | 0,028814333 | CCT5 Chaperonin containing T-complex polypeptide-1 subunit 5 chr1:8685504-8687802 REVERSE LENGTH=482 |
| AT1G52040.1 | 0,462509173 | 0,028817132 | MBP1_ ATMBP myrosinase-binding protein 1 chr1:19350595-19352578 REVERSE LENGTH=462 |
| AT5G42020.2 | 1,193975024 | 0,029100942 | BIP2_ BIP luminal binding protein chr5:16807697-16810480 REVERSE LENGTH=613 |
| AT3G08580.1 | 1,084636147 | 0,029139253 | AAC1 ADP/ATP carrier 1 chr3:2605706-2607030 REVERSE LENGTH=381 |
| AT4G01050.1 | 0,929957274 | 0,029237773 | TROL thylakoid rhodanese-like chr4:455874-458175 FORWARD LENGTH=466 |

| | | | |
|---|---|---|---|
| AT1G32060.1 | 1,109766814 | 0,029749371 | PRK phosphoribulokinase chr1:11532668-11534406 FORWARD LENGTH=395 |
| AT4G24780.1 | 0,638291147 | 0,029848231 | PLL19 chr4:12770631-12772227 REVERSE LENGTH=408 |
| AT3G02230.1 | 0,858435988 | 0,029929675 | RGP1_ ATRGP1 ARABIDOPSIS THALIANA REVERSIBLY GLYCOSYLATED POLYPEPTIDE 1_ reversibly glycosylated |
| AT5G13410.1 | 0,805827298 | 0,03004236 | no symbol available no full name available chr5:4299830-4301706 REVERSE LENGTH=256 |
| AT3G44300.1 | 1,193003503 | 0,030085181 | AtNIT2_ NIT2 nitrilase 2 chr3:15983351-15985172 FORWARD LENGTH=339 |
| AT1G20020.1 | 0,863990188 | 0,030324555 | LFNR2_ FNR2_ ATLFNR2 leaf-type chloroplast-targeted FNR 2_ ferredoxin-NADP(+)-oxidoreductase 2_ LEAF FNR 2 chr1:6 |
| AT2G39770.1 | 0,79822629 | 0,030327711 | CYT1_ EMB101_ GMP1_ VTC1_ SOZ1 GDP-MANNOSE PYROPHOSPHORYLASE 1_ CYTOKINESIS DEFECTIVE 1_ S |
| AT3G18490.1 | 0,835494682 | 0,030362724 | ASPG1 ASPARTIC PROTEASE IN GUARD CELL 1 chr3:6349090-6350592 REVERSE LENGTH=500 |
| AT3G52960.1 | 1,133198431 | 0,030394982 | PrxIIE peroxiredoxin-II-E chr3:19639699-19640403 FORWARD LENGTH=234 |
| AT5G44500.1 | 1,197045586 | 0,030406562 | no symbol available no full name available chr5:17927505-17928269 FORWARD LENGTH=254 |
| AT3G53460.1 | 0,724877356 | 0,030431288 | CP29 chloroplast RNA-binding protein 29 chr3:19819738-19821423 REVERSE LENGTH=342 |
| AT5G49360.1 | 0,629518694 | 0,030441607 | ATBXL1_ BXL1 beta-xylosidase 1_ BETA-XYLOSIDASE 1 chr5:20012179-20016659 REVERSE LENGTH=774 |
| AT3G56340.1 | 0,45165687 | 0,030656132 | RPS26e Ribosomal Protein S26e chr3:20892309-20893343 REVERSE LENGTH=130 |
| AT4G09650.1 | 1,236094177 | 0,030691401 | PDE332_ ATPD PIGMENT DEFECTIVE 332_ ATP synthase delta-subunit gene chr4:6100799-6101503 FORWARD LENGT |
| AT1G53750.1 | 1,140186507 | 0,030700219 | RPT1A regulatory particle triple-A 1A chr1:20065921-20068324 REVERSE LENGTH=426 |
| AT1G12000.1 | 1,519894959 | 0,031039893 | no symbol available no full name available chr1:4050159-4053727 REVERSE LENGTH=566 |
| AT2G27730.1 | 1,576601095 | 0,031158781 | no symbol available no full name available chr2:11820056-11820867 REVERSE LENGTH=113 |
| ATCG01110.1 | 0,68043043 | 0,03143935 | NDHH NAD(P)H dehydrogenase subunit H chrc:122011-123192 REVERSE LENGTH=393 |
| AT1G72730.1 | 1,175513048 | 0,031480982 | no symbol available no full name available chr1:27378040-27379593 REVERSE LENGTH=414 |
| AT1G32200.1 | 0,862401876 | 0,031740797 | ACT1_ ATS1 ACYLTRANSFERASE 1 chr1:11602223-11605001 REVERSE LENGTH=459 |
| AT2G20420.1 | 1,161654939 | 0,031797794 | no symbol available no full name available chr2:8805574-8807858 FORWARD LENGTH=421 |
| AT5G10240.2 | 1,509574229 | 0,03185847 | ASN3 asparagine synthetase 3 chr5:3212934-3216418 REVERSE LENGTH=577 |
| AT2G17840.2 | 0,778755937 | 0,032070677 | ERD7 EARLY-RESPONSIVE TO DEHYDRATION 7 chr2:7756194-7757798 REVERSE LENGTH=394 |
| AT1G26480.1 | 0,500376445 | 0,032097425 | GF14 IOTA_ GRF12 general regulatory factor 12 chr1:9156573-9157845 REVERSE LENGTH=268 |
| AT1G48830.1 | 0,776415728 | 0,032239487 | no symbol available no full name available chr1:18059854-18060935 REVERSE LENGTH=191 |
| AT1G09310.1 | 0,880304512 | 0,032795111 | SVB2_ SVBL SVB-like chr1:3009109-3009648 FORWARD LENGTH=179 |
| AT5G24770.1 | 0,877035509 | 0,032895417 | ATVSP2_ VSP2 vegetative storage protein 2 chr5:8500713-8501844 REVERSE LENGTH=265 |
| AT4G37980.1 | 0,803228905 | 0,033007858 | ELI3-1_ CHR_ ELI3_ ATCAD7_ CAD7 elicitor-activated gene 3-1_ CINNAMALDEHYDE AND HEXENAL REDUCTASE_ |
| AT4G12420.1 | 0,893670919 | 0,033374921 | SKU5 chr4:7349941-7352868 REVERSE LENGTH=587 |
| AT1G04170.1 | 1,623018059 | 0,033765455 | EIF2 GAMMA eukaryotic translation initiation factor 2 gamma subunit chr1:1097423-1099702 FORWARD LENGTH=465 |
| AT4G37990.1 | 0,655417781 | 0,033852396 | ELI3-2_ ATCAD8_ ELI3_ CAD-B2 elicitor-activated gene 3-2_ CINNAMYL-ALCOHOL DEHYDROGENASE B2_ ARABID |
| AT3G26060.1 | 0,910631942 | 0,033885409 | ATPRX Q_ PRXQ peroxiredoxin Q chr3:9524807-9526123 FORWARD LENGTH=216 |
| AT4G15510.3 | 0,700010717 | 0,034020193 | PPD1 PsbP-Domain Protein1 chr4:8861351-8862529 FORWARD LENGTH=205 |
| AT1G23740.1 | 1,265629775 | 0,034626586 | AOR alkenal/one oxidoreductase chr1:8398245-8399656 REVERSE LENGTH=386 |
| AT2G04390.1 | 0,802433635 | 0,03480919 | dS17 chr2:1527911-1528336 FORWARD LENGTH=141 |
| AT4G26300.4 | 1,511445448 | 0,035238094 | emb1027 embryo defective 1027 chr4:13308400-13312204 REVERSE LENGTH=590 |
| AT1G30120.1 | 0,512646836 | 0,03572916 | PDH-E1 BETA pyruvate dehydrogenase E1 beta chr1:10584350-10586477 REVERSE LENGTH=406 |
| AT5G57870.1 | 1,117064109 | 0,035818337 | eIFiso4G1 eukaryotic translation Initiation Factor isoform 4G1 chr5:23439755-23443433 FORWARD LENGTH=780 |
| AT1G72640.4 | 1,169004572 | 0,0360697 | no symbol available no full name available chr1:27346873-27348147 REVERSE LENGTH=203 |

| | | | |
|---|---|---|---|
| AT3G59920.1 | 0,803296868 | 0,036076423 | GDI2_ ATGDI2 RAB GDP dissociation inhibitor 2 chr3:22135157-22138221 FORWARD LENGTH=444 |
| AT5G02500.1 | 1,231956051 | 0,036113576 | AtHsp70-1_ HSP70-1_ HSC70-1_ HSC70_ AT-HSC70-1 ARABIDOPSIS THALIANA HEAT SHOCK COGNATE PROTEIN |
| AT1G09180.1 | 1,08329052 | 0,03655604 | SAR1A_ ATSARA1A_ SARA1A_ ATSAR1 SECRETION-ASSOCIATED RAS 1_ secretion-associated RAS super family 1 ch |
| AT5G50850.1 | 1,135938162 | 0,036647835 | MAB1 MACCI-BOU chr5:20689671-20692976 FORWARD LENGTH=363 |
| AT5G67500.3 | 0,766019358 | 0,036930381 | VDAC2_ ATVDAC2 ARABIDOPSIS THALIANA VOLTAGE DEPENDENT ANION CHANNEL 2_ voltage dependent anion |
| AT3G02780.2 | 0,814821488 | 0,037192832 | IDI2_ IPIAT1_ IPP2 isopentenyl pyrophosphate:dimethylallyl pyrophosphate isomerase 2_ ATISOPENTENYL DIPHOSPHE IS |
| AT1G29900.1 | 1,155857623 | 0,037263263 | CARB_ VEN3 carbamoyl phosphate synthetase B_ VENOSA 3 chr1:10468164-10471976 FORWARD LENGTH=1187 |
| AT4G15545.1 | 0,58377799 | 0,037375876 | NAIP1 NAI2-interacting protein 1 chr4:8875932-8877567 FORWARD LENGTH=337 |
| AT4G03520.1 | 1,121631711 | 0,037445036 | ATHM2_ TRXm2 thioredoxin m2 chr4:1562585-1564055 REVERSE LENGTH=186 |
| AT3G03780.1 | 1,161746928 | 0,037468509 | MS2_ ATMS2 methionine synthase 2 chr3:957602-960740 FORWARD LENGTH=765 |
| AT5G23820.1 | 0,787771678 | 0,037497516 | ML3 MD2-related lipid recognition 3 chr5:8031386-8032809 FORWARD LENGTH=164 |
| AT3G19820.1 | 1,146705805 | 0,037881815 | DWF1_ DIM_ DIM1_ CBB1_ EVE1 ENHANCED VERY-LOW-FLUENCE RESPONSES 1_ DIMINUTIA_ DIMINUTO 1_ C |
| AT1G18540.1 | 0,771081514 | 0,038051905 | no symbol available no full name available chr1:6377448-6378548 REVERSE LENGTH=233 |
| AT5G06600.2 | 1,185738049 | 0,038192931 | UBP12_ AtUBP12 ubiquitin-specific protease 12 chr5:2019545-2027834 REVERSE LENGTH=1115 |
| AT1G35160.1 | 1,108467882 | 0,038197264 | GRF4_ 14-3-3PHI_ GF14 PHI GENERAL REGULATORY FACTOR 4_ 14-3-3 PROTEIN G-BOX FACTOR14 PHI_ GF14 p |
| AT1G29880.1 | 1,341953291 | 0,038260107 | no symbol available no full name available chr1:10459662-10462781 REVERSE LENGTH=729 |
| AT1G56580.1 | 5,707007748 | 0,038269155 | SVB SMALLER WITH VARIABLE BRANCHES chr1:21198402-21198902 REVERSE LENGTH=166 |
| ATCG00660.1 | 0,548763163 | 0,038630446 | RPL20 ribosomal protein L20 chrc:68512-68865 REVERSE LENGTH=117 |
| AT1G56070.1 | 1,133569811 | 0,038767037 | LOS1 LOW EXPRESSION OF OSMOTICALLY RESPONSIVE GENES 1 chr1:20968245-20971077 REVERSE LENGTH=8 |
| AT2G17265.1 | 0,553136563 | 0,038988203 | DRM1_ HSK_ DMR1 DOWNY MILDEW RESISTANT 1_ homoserine kinase chr2:7508606-7509718 FORWARD LENGTH= |
| AT3G48750.1 | 0,772606209 | 0,039312015 | CDC2AAT_ CDKA;1_ CDK2_ CDC2_ CDKA1_ CDC2A cell division control 2 chr3:18072238-18074296 FORWARD LENG |
| AT1G17880.1 | 0,72981188 | 0,039678282 | ATBTF3_ BTF3 basic transcription factor 3 chr1:6152572-6153425 REVERSE LENGTH=165 |
| AT4G12060.1 | 2,051510363 | 0,039974258 | ClpT2 chr4:7228269-7229898 REVERSE LENGTH=241 |
| AT1G07140.1 | 0,873703881 | 0,040631366 | SIRANBP chr1:2192360-2193688 REVERSE LENGTH=228 |
| AT1G60710.1 | 0,80923173 | 0,040970377 | ATB2 chr1:22355073-22356627 REVERSE LENGTH=345 |
| AT3G52300.1 | 1,393683359 | 0,041236695 | ATPQ_ ATPd "ATP synthase D chain_ mitochondrial" chr3:19396689-19398119 FORWARD LENGTH=168 |
| AT3G05560.1 | 0,85269672 | 0,041531881 | no symbol available no full name available chr3:1614641-1615204 FORWARD LENGTH=124 |
| AT1G50200.1 | 1,177873689 | 0,041569916 | ALATS_ ACD Alanyl-tRNA synthetase chr1:18591429-18598311 REVERSE LENGTH=1003 |
| AT5G14200.1 | 0,716620723 | 0,042065061 | ATIMD1_ IMD1 isopropylmalate dehydrogenase 1_ ARABIDOPSIS ISOPROPYLMALATE DEHYDROGENASE 1 chr5:457 |
| AT5G14260.1 | 1,524032667 | 0,042088143 | SAFE1 SAFEGUARD1 chr5:4601139-4603873 FORWARD LENGTH=514 |
| AT1G49970.1 | 0,835632253 | 0,042176116 | ClpR1_ SVR2_ NCLPP5_ CLPR1 NUCLEAR CLPP 5_ CLP protease proteolytic subunit 1_ SUPPRESSOR OF VARIEGATIC |
| AT2G22170.1 | 0,487121749 | 0,042176152 | PLAT2 PLAT domain protein 2 chr2:9427010-9427742 REVERSE LENGTH=183 |
| AT4G23170.1 | 1,301183976 | 0,042329834 | CRK9_ EP1 CYSTEINE-RICH RLK (RECEPTOR-LIKE PROTEIN KINASE) 9 chr4:12135205-12136002 FORWARD LENG |
| AT2G20580.1 | 1,355493002 | 0,043249756 | ATRPN1A_ RPN1A 26S PROTEASOME REGULATORY SUBUNIT S2 1A_ 26S proteasome regulatory subunit S2 1A chr2:8 |
| AT5G60660.1 | 0,623636322 | 0,043467461 | PIP2F_ PIP2;4 plasma membrane intrinsic protein 2;4 chr5:24375673-24376939 REVERSE LENGTH=291 |
| AT5G63570.1 | 1,108505983 | 0,043795876 | GSA1 "glutamate-1-semialdehyde-2_1-aminomutase" chr5:25451957-25453620 FORWARD LENGTH=474 |
| AT1G20630.1 | 1,186907474 | 0,043923847 | CAT1 catalase 1 chr1:7146812-7149609 FORWARD LENGTH=492 |
| AT1G15690.1 | 0,853582991 | 0,043986992 | AtVHP1;1_ AtAVP1_ ATAVP3_ AVP-3_ AVP1_ FUGU5_ VHP1 ARABIDOPSIS THALIANA V-PPASE 3_ FUGU 5 chr1:5 |
| AT5G01530.1 | 0,746866417 | 0,04402834 | LHCB4.1 light harvesting complex photosystem II chr5:209084-210243 FORWARD LENGTH=290 |

| | | | |
|---|---|---|---|
| AT1G20450.1 | 0,909580768 | 0,044339653 | LTI45_ ERD10_ LTI29 EARLY RESPONSIVE TO DEHYDRATION 10_ LOW TEMPERATURE INDUCED 45_ LOW TEM |
| AT2G42220.1 | 0,740291216 | 0,045455156 | no symbol available no full name available chr2:17592105-17593305 FORWARD LENGTH=234 |
| AT2G22240.1 | 2,016936788 | 0,045524595 | MIPS2_ ATIPS2_ ATMIPS2 myo-inositol-1-phosphate synthase 2_ INOSITOL 3-PHOSPHATE SYNTHASE 2_ MYO-INOSI |
| AT5G07350.1 | 1,23524067 | 0,046067315 | Tudor1_ TSN1_ AtTudor1 TUDOR-SN protein 1_ Arabidopsis thaliana TUDOR-SN protein 1 chr5:2320344-2324892 REVER |
| AT1G70310.1 | 1,252293397 | 0,04607895 | SPDS2 spermidine synthase 2 chr1:26485497-26487352 REVERSE LENGTH=340 |
| AT5G09500.1 | 0,692747339 | 0,046172141 | no symbol available no full name available chr5:2954044-2954850 REVERSE LENGTH=150 |
| AT5G02240.1 | 0,822478779 | 0,046205957 | no symbol available no full name available chr5:451502-452984 FORWARD LENGTH=253 |
| AT3G49720.1 | 0,581971263 | 0,046362891 | CGR2 chr3:18440192-18441655 REVERSE LENGTH=261 |
| AT1G75270.1 | 1,19812941 | 0,046596904 | DHAR2 dehydroascorbate reductase 2 chr1:28250255-28251237 REVERSE LENGTH=213 |
| AT1G74270.1 | 0,79878359 | 0,04664112 | no symbol available no full name available chr1:27928415-27929466 REVERSE LENGTH=112 |
| AT1G52030.1 | 1,173033246 | 0,046644134 | F-ATMBP_ MBP1.2_ MBP2 myrosinase-binding protein 2 chr1:19346090-19348282 REVERSE LENGTH=642 |
| AT5G53490.1 | 1,122121678 | 0,0470542 | TL17 Thylakoid lumenal 17.4 kDa protein chr5:21723488-21724621 REVERSE LENGTH=236 |
| AT1G56500.2 | 0,797756963 | 0,047245831 | SOQ1 suppressor of quenching 1 chr1:21159775-21166313 FORWARD LENGTH=878 |
| AT4G34620.1 | 0,880844501 | 0,047360831 | SSR16 small subunit ribosomal protein 16 chr4:16535084-16536092 REVERSE LENGTH=113 |
| AT5G16620.1 | 1,152476809 | 0,047638533 | PDE120_ TIC40_ ATTIC40 translocon at the inner envelope membrane of chloroplasts 40_ pigment defective embryo 120 chr5 |
| AT1G21750.1 | 1,140957676 | 0,047640103 | PDIL1-1_ ATPDI5_ PDI5_ ATPDIL1-1 PDI-like 1-1_ ARABIDOPSIS THALIANA PROTEIN DISULFIDE ISOMERASE 5_ |
| AT2G44610.1 | 1,378995341 | 0,04781963 | ATRAB6A_ RAB6_ RAB6A_ ATRABH1B chr2:18411778-18413883 REVERSE LENGTH=208 |
| AT1G20260.1 | 1,372499766 | 0,047886358 | AtVAB3_ VAB3 V-ATPase B subunit 3 chr1:7016971-7020290 FORWARD LENGTH=487 |
| AT3G63140.1 | 0,905841555 | 0,047967209 | CSP41A chloroplast stem-loop binding protein of 41 kDa chr3:23327006-23328620 REVERSE LENGTH=406 |
| AT1G75330.1 | 0,563024121 | 0,048047653 | OTC ornithine carbamoyltransferase chr1:28266457-28268383 REVERSE LENGTH=375 |
| AT1G74920.2 | 1,292856995 | 0,048080323 | ALDH10A8 aldehyde dehydrogenase 10A8 chr1:28139175-28142573 REVERSE LENGTH=496 |
| AT1G56050.1 | 0,768326124 | 0,048273518 | EngD-2 chr1:20963793-20966181 FORWARD LENGTH=421 |
| ATCG00540.1 | 0,908145348 | 0,048424212 | PETA photosynthetic electron transfer A chrc:61657-62619 FORWARD LENGTH=320 |
| AT4G21990.1 | 0,659590036 | 0,048624046 | APR3_ PRH26_ PRH-26_ ATAPR3 PAPS REDUCTASE HOMOLOG 26_ APS reductase 3 chr4:11657284-11658973 REVER |
| AT1G18080.1 | 1,168048047 | 0,049703014 | RACK1A_AT_ AtRACK1_ SAC53_ ATARCA_ RACK1A RECEPTOR FOR ACTIVATED C KINASE 1 A_ Suppressor of A |

THALIANA PROTEIN DISULFIDE ISOMERASE 11_ UNFERTILIZED EMBRYO SAC 5_ MATERNAL EFFECT EMBRYO ARREST 30 chr2:19481503-19483683 FOR

[illegible]HAPERONIN 60ALPHA_ ACCUMULATION AND REPLICATION OF CHLOROPLASTS 2_ chaperonin-60alpha chr2:11926603-11929184 FORWARD LENGTH=586

3B1_ EUKARYOTIC TRANSLATION INITIATION FACTOR 3B1_ EUKARYOTIC TRANSLATION INITIATION FACTOR 3B_ translation initiation factor 3B1 chr5:9[illegible]

IPLEX 33 KILODALTON PROTEIN_ OXYGEN EVOLVING ENHANCER PROTEIN 33_ PS II oxygen-evolving complex 1_ MANGANESE-STABILIZING PROTEIN 1_

GNAL RECOGNITION PARTICLE 54 KDA SUBUNIT CHLOROPLAST PROTEIN_ FIFTY-FOUR CHLOROPLAST HOMOLOGUE chr5:1060265-1063257 REVERSE

S GLUTATHIONE S-TRANSFERASE 1_ ARABIDOPSIS THALIANA GLUATIONE S-TRANSFERASE F3_ glutathione S-transferase 6_ GLUTATHIONE S-TRANSFEI

TEIN 1_ GLYCINE RICH PROTEIN 7_ "cold_ circadian rhythm_ and rna binding 2"_ GLYCINE-RICH RNA-BINDING PROTEIN 7 chr2:9265477-9266316 REVERSE LI

'E GENES 2_ suppressors of PIN1 overexpression 1_ ALTERED EXPRESSION OF APX2 8_ FIERY1_ ROTUNDA 1 chr5:25610002-25611802 FORWARD LENGTH=35

)OPSIS THALIANA CINNAMYL-ALCOHOL DEHYDROGENASE 8_ ELICITOR-ACTIVATED GENE 3 chr4:17855964-17857388 FORWARD LENGTH=359

70-1_ heat shock cognate protein 70-1_ HEAT SHOCK COGNATE PROTEIN 70_ HEAT SHOCK PROTEIN 70-1 chr5:554055-556334 REVERSE LENGTH=651

_ 33 KDA OXYGEN EVOLVING POLYPEPTIDE 1 chr5:26568744-26570124 FORWARD LENGTH=332
