## Supplementary material for "HY5 enhances *Arabidopsis* tolerance to combined high light and heat stress by coordinating photoprotection and hormone signaling": Table S7

**Supplemental Table S7. Differentially accumulated proteins compared to control (P < 0.05) in *hy5-215* leaves subjected to combined conditions of high light and heat stre**

| Protein ID | Fold Change | p-value | Protein description |
|---|---|---|---|
| AT5G59720.1 | 81,98632274 | 8,69418E-07 | HSP18.2 heat shock protein 18.2 chr5:24062632-24063117 FORWARD LENGTH=161 |
| AT5G52640.1 | 20,44937966 | 1,16299E-06 | HSP81-1_ ATHSP90.1_ HSP81.1_ AtHsp90-1_ HSP83_ ATHS83_ HSP90.1 HEAT SHOCK PROTEIN 90-1_ heat shock |
| AT5G48570.1 | 12,80696628 | 1,42131E-06 | ROF2_ ATFKBP65_ FKBP65 chr5:19690746-19693656 REVERSE LENGTH=578 |
| AT3G12580.1 | 6,340228691 | 1,77978E-06 | HSP70_ ATHSP70_ HSC70-4 ARABIDOPSIS HEAT SHOCK PROTEIN 70_ heat shock protein 70 chr3:3991487-39936 |
| AT2G35635.1 | 23,51011352 | 2,41387E-06 | UBQ7_ RUB2 RELATED TO UBIQUITIN 2_ ubiquitin 7 chr2:14981044-14981943 FORWARD LENGTH=154 |
| AT5G61790.1 | 1,493705533 | 2,45083E-06 | CNX1_ ATCNX1 calnexin 1 chr5:24827394-24829642 REVERSE LENGTH=530 |
| AT5G13650.1 | 0,182193647 | 4,26901E-06 | SVR3 SUPPRESSOR OF VARIEGATION 3 chr5:4397821-4402364 FORWARD LENGTH=675 |
| AT3G09350.1 | 23,6423147 | 5,34835E-06 | Fes1A Fes1A chr3:2871216-2873109 FORWARD LENGTH=363 |
| AT3G09440.1 | 3,15289147 | 5,62629E-06 | no symbol available no full name available chr3:2903434-2905632 REVERSE LENGTH=649 |
| AT1G12240.1 | 0,299395949 | 7,99987E-06 | AtVI2_ VAC-INV_ ATBETAFRUCT4_ VI2_ FRUCT4_ AtFRUCT4_ VIN2 VACUOLAR INVERTASE_ vacuolar invert |
| AT1G07400.1 | 32,97086268 | 9,73639E-06 | HSP17.8 chr1:2275148-2275621 FORWARD LENGTH=157 |
| AT2G39310.1 | 5,699452634 | 1,02201E-05 | JAL22 jacalin-related lectin 22 chr2:16414262-16416323 REVERSE LENGTH=458 |
| AT1G14250.1 | 0,235601156 | 1,16294E-05 | no symbol available no full name available chr1:4868675-4871203 FORWARD LENGTH=488 |
| AT1G78820.1 | 3,180288716 | 1,25067E-05 | no symbol available no full name available chr1:29634401-29635768 REVERSE LENGTH=455 |
| AT1G16030.1 | 4,978515304 | 1,25417E-05 | Hsp70b heat shock protein 70B chr1:5502386-5504326 REVERSE LENGTH=646 |
| AT5G09590.1 | 3,826408751 | 1,31797E-05 | MTHSC70-2_ HSC70-5 mitochondrial HSO70 2_ HEAT SHOCK COGNATE chr5:2975721-2978508 FORWARD LENG |
| AT4G39730.1 | 0,401052617 | 1,3373E-05 | ATPLAT1_ PLAT1 PLAT domain protein 1_ "Polycystin_ Lipoxygenase_ Alpha-toxin and Triacylglycerol lipase 1" chr4:1 |
| AT2G29500.1 | 38,676793 | 1,56088E-05 | HSP17.6B chr2:12633279-12633740 REVERSE LENGTH=153 |
| AT2G37270.1 | 0,678085032 | 2,00457E-05 | RPS5B_ ATRPS5B ribosomal protein 5B chr2:15647883-15649042 REVERSE LENGTH=207 |
| AT3G25230.1 | 3,7562595 | 2,00477E-05 | FKBP62_ ROF1_ ATFKBP62 rotamase FKBP 1_ FK506 BINDING PROTEIN 62 chr3:9188257-9191137 FORWARD LE |
| AT5G14740.5 | 1,879683123 | 2,28949E-05 | BETA CA2_ CA2_ DEG12_ CA18 BETA CARBONIC ANHYDRASE 2_ CARBONIC ANHYDRASE 18_ carbonic anhy |
| AT5G01600.1 | 8,16447599 | 2,61846E-05 | FER1_ ATFER1 ferretin 1_ ARABIDOPSIS THALIANA FERRETIN 1 chr5:228149-229594 REVERSE LENGTH=255 |
| AT5G19940.1 | 0,797815524 | 2,87346E-05 | FBN6_ PAP8 FIBRILLIN6_ Probable Plastid-Lipid Associated Protein chr5:6739693-6740661 FORWARD LENGTH=23 |
| AT4G04020.1 | 8,511186113 | 2,94949E-05 | FIB_ PGL35_ FIB1a plastoglobulin 35_ fibrillin 1a_ fibrillin chr4:1932161-1933546 FORWARD LENGTH=318 |
| AT5G03340.1 | 0,761074242 | 3,34606E-05 | AtCDC48C cell division cycle 48C chr5:810091-813133 REVERSE LENGTH=810 |
| AT4G39980.1 | 0,607018704 | 4,23775E-05 | AtDAHP1_ DHS1_ DAHP1 3-DEOXY-D-ARABINO-HEPTULOSONATE-7-PHOSPHATE 1_ 3-deoxy-D-arabino-heptul |
| AT3G52380.1 | 0,244019372 | 4,49903E-05 | PDE322_ CP33 PIGMENT DEFECTIVE 322_ chloroplast RNA-binding protein 33 chr3:19421619-19422855 FORWARD |
| AT1G58270.1 | 0,429280495 | 4,53045E-05 | ZW9 chr1:21612394-21614089 REVERSE LENGTH=396 |
| AT5G19760.1 | 1,262229554 | 6,01709E-05 | no symbol available no full name available chr5:6679591-6681845 REVERSE LENGTH=298 |
| AT1G59860.1 | 8,527008607 | 6,08839E-05 | HSP17.6A chr1:22031474-22031941 FORWARD LENGTH=155 |
| AT2G05100.1 | 0,490874496 | 6,19012E-05 | LHCB2_ LHCB2.1 LIGHT-HARVESTING CHLOROPHYLL B-BINDING 2_ photosystem II light harvesting complex ge |
| AT1G66430.1 | 2,360167933 | 6,40459E-05 | FRK6_ FRK3 Fructokinase 6_ Fructokinase 3 chr1:24778400-24780393 FORWARD LENGTH=384 |
| AT3G25770.1 | 0,717206752 | 6,62876E-05 | AOC2 allene oxide cyclase 2 chr3:9406975-9407839 FORWARD LENGTH=253 |
| AT4G27670.1 | 35,03062851 | 6,91878E-05 | HSP21 heat shock protein 21 chr4:13819048-13819895 REVERSE LENGTH=227 |
| AT5G28540.1 | 1,715282082 | 6,98914E-05 | BIP1 chr5:10540665-10543274 REVERSE LENGTH=669 |
| AT1G54040.2 | 5,843616316 | 7,073E-05 | TASTY_ ESR_ ESP epithiospecifier protein_ EPITHIOSPECIFYING SENESCENCE REGULATOR chr1:20170995-2017 |

| | | | |
|---|---|---|---|
| AT3G13470.1 | 1,195009605 | 7,18566E-05 | CPNB2_ Cpn60beta2 chaperonin-60beta2 chr3:4389685-4392624 FORWARD LENGTH=596 |
| AT5G01410.1 | 1,805786498 | 7,5777E-05 | PDX1_ ATPDX1.3_ ATPDX1_ PDX1.3_ RSR4 REDUCED SUGAR RESPONSE 4_ PYRIDOXINE BIOSYNTHESIS 1.[illegible] |
| AT4G22240.1 | 1,886332791 | 7,67865E-05 | FBN1b fibrillin 1b chr4:11766090-11767227 REVERSE LENGTH=310 |
| AT4G30270.1 | 0,403857083 | 7,81742E-05 | MERI5B_ XTH24_ MERI-5_ SEN4 xyloglucan endotransglucosylase/hydrolase 24_ SENESCENCE 4_ MERISTEM 5_ m[illegible] |
| AT2G06850.1 | 1,978787918 | 7,89725E-05 | EXT_ XTH4_ EXGT-A1 ENDOXYLOGLUCAN TRANSFERASE_ endoxyloglucan transferase A1_ xyloglucan endotrans[illegible] |
| AT3G23990.1 | 1,590612241 | 7,90484E-05 | HSP60_ HSP60-3B heat shock protein 60_ HEAT SHOCK PROTEIN 60-3B chr3:8669013-8672278 FORWARD LENGT[illegible] |
| AT2G05920.1 | 0,658343731 | 7,95998E-05 | SBT1.8 subtilase 1.8 chr2:2269831-2272207 REVERSE LENGTH=754 |
| AT2G34810.1 | 0,54611263 | 8,72812E-05 | AtBBE16 chr2:14685292-14686914 FORWARD LENGTH=540 |
| AT1G65350.1 | 1,71299846 | 8,85861E-05 | UBQ13 ubiquitin 13 chr1:24272518-24277275 REVERSE LENGTH=319 |
| AT4G27440.1 | 0,307302115 | 0,000101721 | PORB protochlorophyllide oxidoreductase B chr4:13725648-13727107 FORWARD LENGTH=401 |
| AT1G23730.1 | 0,249971632 | 0,000103706 | ATBCA3_ BCA3 beta carbonic anhydrase 3_ BETA CARBONIC ANHYDRASE 3 chr1:8395965-8398014 FORWARD L[illegible] |
| AT2G25450.1 | 0,295693609 | 0,000105883 | GSL-OH glucosinolate hydroxylase chr2:10830286-10831563 REVERSE LENGTH=359 |
| AT3G08940.2 | 0,632984793 | 0,000106629 | LHCB4.2 light harvesting complex photosystem II chr3:2717717-2718665 FORWARD LENGTH=287 |
| AT5G13630.1 | 0,384170026 | 0,00010939 | ABAR_ CHLH_ GUN5_ CCH_ CCH1 ABA-BINDING PROTEIN_ H SUBUNIT OF MG-CHELATASE_ GENOMES UN[illegible] |
| AT1G75040.1 | 4,161723359 | 0,000109773 | PR-5_ PR5 pathogenesis-related gene 5 chr1:28177754-28178731 FORWARD LENGTH=239 |
| AT1G79040.1 | 0,537490581 | 0,000118028 | PSBR photosystem II subunit R chr1:29736085-29736781 FORWARD LENGTH=140 |
| AT5G14920.1 | 2,94403648 | 0,000118579 | GASA14 A-stimulated in Arabidopsis 14 chr5:4826598-4827761 FORWARD LENGTH=275 |
| AT3G23700.1 | 0,703594274 | 0,000119213 | SRRP1 S1 RNA-binding ribosomal protein 1 chr3:8531689-8533742 REVERSE LENGTH=392 |
| AT5G12020.1 | 35,39900955 | 0,00012207 | HSP17.6II 17.6 kDa class II heat shock protein chr5:3882409-3882876 REVERSE LENGTH=155 |
| AT2G36460.1 | 1,653456358 | 0,000123991 | FBA6 fructose-bisphosphate aldolase 6 chr2:15296929-15298387 REVERSE LENGTH=358 |
| AT1G55490.1 | 1,182177545 | 0,000125795 | Cpn60beta1_ LEN1_ CPNB1_ CPN60B chaperonin-60beta1_ LESION INITIATION 1_ chaperonin 60 beta chr1:2071571[illegible] |
| AT1G49630.1 | 0,462066289 | 0,000139216 | PREP2_ ATPREP2 presequence protease 2 chr1:18368405-18375336 REVERSE LENGTH=1080 |
| AT2G47110.1 | 1,57778865 | 0,000149124 | UBI6_ UBQ6_ RPS27aB UBIQUITIN EXTENSION PROTEIN 6_ ubiquitin 6_ Ribosomal protein S27aB chr2:19344701-[illegible] |
| AT3G10350.1 | 0,682223407 | 0,000152962 | AtGET3b_ GET3b Guided Entry of Tail-anchored proteins 3b chr3:3208310-3210678 FORWARD LENGTH=411 |
| AT4G21960.1 | 0,124845259 | 0,000160853 | PRXR1 chr4:11646613-11648312 REVERSE LENGTH=330 |
| AT1G77510.1 | 1,500344561 | 0,000163827 | ATPDI6_ PDIL1-2_ ATPDIL1-2_ PDI6 PDI-like 1-2_ PROTEIN DISULFIDE ISOMERASE 6 chr1:29126742-29129433 [illegible] |
| AT1G47250.1 | 2,287453566 | 0,000164121 | PAF2 20S proteasome alpha subunit F2 chr1:17319220-17320900 FORWARD LENGTH=277 |
| AT1G75940.1 | 1,511093339 | 0,000164751 | ATA27_ BGLU20 BETA GLUCOSIDASE 20 chr1:28511198-28514044 FORWARD LENGTH=535 |
| AT4G25200.1 | 12,84623591 | 0,000165155 | ATHSP23.6-MITO_ HSP23.6-MITO mitochondrion-localized small heat shock protein 23.6 chr4:12917089-12917858 FO[illegible] |
| AT3G16420.1 | 0,559383195 | 0,000165254 | PBPI_ JAL30_ PBP1 PYK10-binding protein 1_ JACALIN-RELATED LECTIN 30 chr3:5579560-5580674 FORWARD L[illegible] |
| AT4G33010.1 | 0,879848656 | 0,00018162 | GLDP1_ AtGLDP1 glycine decarboxylase P-protein 1 chr4:15926852-15931150 REVERSE LENGTH=1037 |
| AT2G37220.1 | 0,326541221 | 0,000186837 | no symbol available no full name available chr2:15634980-15636331 REVERSE LENGTH=289 |
| AT5G51440.1 | 21,64526741 | 0,000200201 | HSP23.5 chr5:20891242-20892013 FORWARD LENGTH=210 |
| AT5G06870.1 | 0,810723878 | 0,000221799 | PGIP2_ ATPGIP2 ARABIDOPSIS POLYGALACTURONASE INHIBITING PROTEIN 2_ polygalacturonase inhibiting p[illegible] |
| AT5G37640.1 | 1,559629045 | 0,000233886 | UBQ9 ubiquitin 9 chr5:14952782-14953750 REVERSE LENGTH=322 |
| AT1G36280.2 | 0,413643152 | 0,000238551 | no symbol available no full name available chr1:13640600-13642908 FORWARD LENGTH=519 |
| AT5G15520.1 | 3,062587028 | 0,000247995 | no symbol available no full name available chr5:5037242-5038136 REVERSE LENGTH=143 |
| AT5G12030.1 | 19,90274687 | 0,000250349 | AT-HSP17.6A_ HSP17.6A_ HSP17.6 HEAT SHOCK PROTEIN 17.6_ heat shock protein 17.6A chr5:3884214-3884684 R[illegible] |
| AT4G35090.1 | 1,658186221 | 0,000255179 | CAT2 catalase 2 chr4:16700937-16703215 REVERSE LENGTH=492 |

| | | | |
|---|---|---|---|
| AT3G28290.1 | 0,489743559 | 0,000261051 | AT14A chr3:10547873-10549030 FORWARD LENGTH=385 |
| AT5G24780.1 | 0,65246613 | 0,000270478 | VSP1_ ATVSP1 vegetative storage protein 1 chr5:8507783-8508889 REVERSE LENGTH=270 |
| AT1G35580.3 | 2,12370202 | 0,000283241 | CINV1_ A/N-InvG_ NIN2 cytosolic invertase 1_ alkaline/neutral invertase G_ neutral invertase 2 chr1:13123183-1312480[illegible] |
| AT4G02520.1 | 1,368625997 | 0,000283596 | ATGSTF2_ GSTF2_ ATPM24.1_ GST2_ ATPM24 glutathione S-transferase PHI 2 chr4:1110673-1111531 REVERSE LE[illegible] |
| AT3G23570.1 | 0,706448875 | 0,000284827 | no symbol available no full name available chr3:8458052-8459608 REVERSE LENGTH=239 |
| AT3G16390.1 | 0,809454595 | 0,000293719 | NSP3 nitrile specifier protein 3 chr3:5562602-5564356 FORWARD LENGTH=467 |
| AT5G42020.2 | 1,613655738 | 0,000314157 | BIP2_ BIP luminal binding protein chr5:16807697-16810480 REVERSE LENGTH=613 |
| AT5G12250.1 | 0,659352267 | 0,000316642 | TUB6 beta-6 tubulin chr5:3961317-3962971 REVERSE LENGTH=449 |
| AT2G33210.2 | 1,23814035 | 0,000318339 | HSP60_ HSP60-2 heat shock protein 60-2 chr2:14075093-14078568 REVERSE LENGTH=580 |
| AT2G15620.1 | 0,705606776 | 0,000325439 | NIR_ ATHNIR_ NIR1 ARABIDOPSIS THALIANA NITRITE REDUCTASE_ nitrite reductase 1_ NITRITE REDUCTAS[illegible] |
| AT5G47190.1 | 1,698233506 | 0,000332246 | PRPL19 plastid ribosomal proteins of the 50S subunit 19 chr5:19164432-19166064 REVERSE LENGTH=229 |
| AT5G25980.2 | 0,694268712 | 0,000333399 | BGLU37_ TGG2 BETA GLUCOSIDASE 37_ glucoside glucohydrolase 2 chr5:9072730-9075477 FORWARD LENGTH=[illegible] |
| AT4G24190.1 | 1,684003252 | 0,000341942 | SHD_ HSP90_ HSP90.7_ AtHsp90-7_ AtHsp90.7 SHEPHERD_ HEAT SHOCK PROTEIN 90.7_ HEAT SHOCK PROTE[illegible] |
| AT3G11510.1 | 1,673695139 | 0,000351951 | no symbol available no full name available chr3:3623757-3624866 REVERSE LENGTH=150 |
| AT5G61170.1 | 0,553411675 | 0,000352618 | no symbol available no full name available chr5:24611158-24612202 FORWARD LENGTH=143 |
| AT1G62660.1 | 0,720772805 | 0,000380221 | VI1 VACUOLAR INVERTASE 1 chr1:23199949-23203515 FORWARD LENGTH=648 |
| AT3G02880.1 | 1,956546608 | 0,000387965 | KIN7 Kinase 7 chr3:634819-636982 FORWARD LENGTH=627 |
| AT1G09180.1 | 1,182453347 | 0,000392483 | SAR1A_ ATSARA1A_ SARA1A_ ATSAR1 SECRETION-ASSOCIATED RAS 1_ secretion-associated RAS super family |
| AT4G02080.1 | 0,630463951 | 0,000399336 | ATSARA1C_ SAR2_ SAR1C_ ATSAR2_ ASAR1 secretion-associated RAS super family 2 chr4:921554-922547 FORWA[illegible] |
| AT3G09840.1 | 1,453907376 | 0,000408525 | ATCDC48_ CDC48_ AtCDC48A_ CDC48A cell division cycle 48 chr3:3019494-3022832 FORWARD LENGTH=809 |
| AT5G04140.2 | 0,84365197 | 0,000414661 | GLUS_ FD-GOGAT_ GLS1_ GLU1 FERREDOXIN-DEPENDENT GLUTAMATE SYNTHASE 1_ glutamate synthase 1_ |
| AT1G74470.1 | 0,685525143 | 0,000417115 | no symbol available no full name available chr1:27991248-27992845 FORWARD LENGTH=467 |
| AT5G14660.1 | 0,604034967 | 0,000417975 | ATDEF2_ DEF2_ PDF1B peptide deformylase 1B chr5:4727129-4728671 REVERSE LENGTH=273 |
| AT3G18780.1 | 1,755117547 | 0,00045357 | LSR2_ ACT2_ ENL2_ DER1_ FIZ2 FRIZZY AND KINKED SHOOTS 2_ LIGHT STRESS-REGULATED 2_ DEFORMI[illegible] |
| AT1G09795.1 | 1,55407148 | 0,00046644 | ATATP-PRT2_ HISN1B_ ATP-PRT2 ATP phosphoribosyl transferase 2 chr1:3173588-3176690 FORWARD LENGTH=4[illegible] |
| AT5G58070.1 | 2,096270177 | 0,000490743 | TIL_ ATTIL TEMPERATURE-INDUCED LIPOCALIN_ temperature-induced lipocalin chr5:23500512-23501156 REVE[illegible] |
| AT2G29450.1 | 2,556369069 | 0,000499558 | AT103-1A_ ATGSTU1_ ATGSTU5_ GSTU5 glutathione S-transferase tau 5_ ARABIDOPSIS THALIANA GLUTATHI[illegible] |
| AT4G23170.1 | 1,549734792 | 0,000505573 | CRK9_ EP1 CYSTEINE-RICH RLK (RECEPTOR-LIKE PROTEIN KINASE) 9 chr4:12135205-12136002 FORWARD L[illegible] |
| AT1G74310.1 | 47,52578499 | 0,000515104 | ATHSP101_ HSP101_ HOT1 heat shock protein 101 chr1:27936715-27939862 REVERSE LENGTH=911 |
| AT5G23820.1 | 0,665159597 | 0,00052798 | ML3 MD2-related lipid recognition 3 chr5:8031386-8032809 FORWARD LENGTH=164 |
| AT1G56050.1 | 0,467065209 | 0,000546073 | EngD-2 chr1:20963793-20966181 FORWARD LENGTH=421 |
| AT4G12400.2 | 21,02563728 | 0,000555758 | Hop3 Hop3 chr4:7338866-7341239 REVERSE LENGTH=558 |
| AT5G66760.1 | 0,52214675 | 0,000560145 | SDH1-1 succinate dehydrogenase 1-1 chr5:26653776-26657224 FORWARD LENGTH=634 |
| AT1G12840.1 | 1,248245755 | 0,000567954 | DET3_ ATVHA-C ARABIDOPSIS THALIANA VACUOLAR ATP SYNTHASE SUBUNIT C_ DE-ETIOLATED 3 chr1[illegible] |
| AT2G18450.1 | 2,512430494 | 0,000569252 | SDH1-2 succinate dehydrogenase 1-2 chr2:7997510-8000801 REVERSE LENGTH=632 |
| AT3G12290.1 | 1,858413128 | 0,00057053 | MTHFD1 methylenetetrahydrofolate dehydrogenase/methenyltetrahydrofolate cyclohydrolase chr3:3919591-3921326 FOR[illegible] |
| AT2G22240.1 | 8,483352849 | 0,000572856 | MIPS2_ ATIPS2_ ATMIPS2 myo-inositol-1-phosphate synthase 2_ INOSITOL 3-PHOSPHATE SYNTHASE 2_ MYO-IN[illegible] |
| AT5G60670.1 | 1,574072766 | 0,00062879 | RPL12C Ribosomal Protein Like 12C chr5:24381066-24381566 REVERSE LENGTH=166 |
| AT4G05530.1 | 1,423283281 | 0,000642092 | SDRA_ IBR1 indole-3-butyric acid response 1_ SHORT-CHAIN DEHYDROGENASE/REDUCTASE A chr4:2816462-28[illegible] |

| | | | |
|---|---|---|---|
| AT3G55610.1 | 1,821994242 | 0,000652925 | P5CS2 delta 1-pyrroline-5-carboxylate synthase 2 chr3:20624278-20628989 REVERSE LENGTH=726 |
| AT5G43850.1 | 3,590936526 | 0,000670228 | ATARD4_ ARD4 chr5:17627364-17629122 REVERSE LENGTH=187 |
| AT2G45810.1 | 1,36067453 | 0,000675045 | RH6 RNA Helicase 6 chr2:18859836-18862318 FORWARD LENGTH=528 |
| AT3G57260.1 | 1,498309889 | 0,000679768 | AtBG2_ PR2_ GNS2_ AtPR2_ BG2_ PR-2_ BGL2 "beta-1_3-glucanase 2"_ PATHOGENESIS-RELATED PROTEIN 2_ ' |
| AT5G66190.1 | 0,817422152 | 0,000708813 | LFNR1_ ATLFNR1_ FNR1 leaf-type chloroplast-targeted FNR 1_ LEAF FNR 1_ ferredoxin-NADP(+)-oxidoreductase 1 cl |
| AT3G19170.1 | 0,600795488 | 0,00071135 | ATZNMP_ ATPREP1_ PREP1 presequence protease 1 chr3:6625578-6631874 REVERSE LENGTH=1080 |
| AT3G09640.1 | 2,521016104 | 0,000716534 | AtAPX2_ APX1B_ APX2 ASCORBATE PEROXIDASE 1B_ ascorbate peroxidase 2 chr3:2956301-2958163 FORWARD |
| AT5G50950.3 | 0,478645809 | 0,000728569 | FUM2 FUMARASE 2 chr5:20731191-20733636 FORWARD LENGTH=317 |
| AT4G37800.1 | 0,62622981 | 0,000733468 | XTH7 xyloglucan endotransglucosylase/hydrolase 7 chr4:17775703-17777372 REVERSE LENGTH=293 |
| AT3G15520.2 | 0,677809372 | 0,000737269 | no symbol available no full name available chr3:5249739-5251861 REVERSE LENGTH=326 |
| AT1G21750.1 | 1,327427649 | 0,000753862 | PDIL1-1_ ATPDI5_ PDI5_ ATPDIL1-1 PDI-like 1-1_ ARABIDOPSIS THALIANA PROTEIN DISULFIDE ISOMERASI |
| AT1G79930.1 | 1,616873973 | 0,00075674 | HSP91_ AtHsp70-14 heat shock protein 91 chr1:30063781-30067067 REVERSE LENGTH=831 |
| AT5G25460.1 | 0,504501997 | 0,000774687 | DGR2 DUF642 L-GalL responsive gene 2 chr5:8863430-8865394 FORWARD LENGTH=369 |
| AT2G21660.1 | 0,497932487 | 0,000781619 | RBGA3_ GR-RBP7_ CCR2_ ATGRP7_ GRP7_ SRBP1 RNA-binding glycine-rich protein A3_ SMALL RNA-BINDING I |
| AT3G09790.1 | 0,61550034 | 0,000783224 | UBQ8 ubiquitin 8 chr3:3004111-3006006 REVERSE LENGTH=631 |
| AT5G04430.1 | 2,449647028 | 0,000819557 | BTR1_ BTR1L_ BTR1S BINDING TO TOMV RNA 1S (SHORT FORM)_ binding to TOMV RNA 1L (long form)_ BINI |
| AT2G14610.1 | 2,166622078 | 0,000835547 | PR1_ ATPR1_ AtCAPE9_ PR 1 PATHOGENESIS-RELATED GENE 1_ pathogenesis-related gene 1 chr2:6241944-62424 |
| AT3G46230.1 | 9,502252155 | 0,000839002 | HSP17.4_ ATHSP17.4 heat shock protein 17.4_ ARABIDOPSIS THALIANA HEAT SHOCK PROTEIN 17.4 chr3:16984 |
| AT2G47470.1 | 1,143559803 | 0,000874421 | UNE5_ MEE30_ ATPDI11_ PDI11_ ATPDIL2-1 PROTEIN DISULFIDE ISOMERASE 11_ PDI-LIKE 2-1_ ARABIDOP |
| AT5G11420.1 | 1,514407272 | 0,000921659 | no symbol available no full name available chr5:3644655-3646991 FORWARD LENGTH=366 |
| AT1G79920.1 | 1,716869816 | 0,000941412 | Hsp70-15_ AtHsp70-15 heat shock protein 70-15 chr1:30059302-30062224 REVERSE LENGTH=736 |
| AT4G09040.1 | 0,515611959 | 0,000964334 | CP33C chr4:5795075-5797315 REVERSE LENGTH=304 |
| AT1G37130.1 | 0,703663751 | 0,000968377 | B29_ ATNR2_ NIA2_ NIA2-1_ NR2_ CHL3_ NR NITRATE REDUCTASE 2_ CHLORATE RESISTANT 3_ ARABIDO |
| AT3G08740.1 | 0,35102077 | 0,001002073 | no symbol available no full name available chr3:2654788-2656154 REVERSE LENGTH=236 |
| AT1G78380.1 | 1,528430054 | 0,001072178 | GSTU19_ GST8_ ATGSTU19 A. THALIANA GLUTATHIONE S-TRANSFERASE TAU 19_ glutathione S-transferase T |
| AT1G22410.1 | 0,31099377 | 0,001086033 | no symbol available no full name available chr1:7912120-7914742 FORWARD LENGTH=527 |
| AT1G56110.1 | 1,523039273 | 0,001114255 | NOP56 homolog of nucleolar protein NOP56 chr1:20984544-20986893 REVERSE LENGTH=522 |
| AT2G18960.1 | 1,222227357 | 0,001138522 | HA1_ OST2_ AHA1_ PMA H(+)-ATPase 1_ PLASMA MEMBRANE PROTON ATPASE_ OPEN STOMATA 2 chr2:82 |
| AT5G24770.1 | 0,613354852 | 0,00115879 | ATVSP2_ VSP2 vegetative storage protein 2 chr5:8500713-8501844 REVERSE LENGTH=265 |
| AT1G13320.4 | 0,560832296 | 0,001167038 | PP2AA3 protein phosphatase 2A subunit A3 chr1:4563692-4566451 REVERSE LENGTH=412 |
| AT1G76160.1 | 3,68018724 | 0,001195138 | sks5 SKU5 similar 5 chr1:28578211-28581020 REVERSE LENGTH=541 |
| AT1G15980.1 | 0,72468182 | 0,001203616 | NDF1_ NDH48_ PnsB1 NAD(P)H DEHYDROGENASE SUBUNIT 48_ Photosynthetic NDH subcomplex B 1_ NDH-dep |
| AT4G38510.1 | 0,72437384 | 0,00121067 | AtVAB2_ VAB2 V-ATPase B subunit 2 chr4:18011155-18014789 REVERSE LENGTH=487 |
| AT1G45145.1 | 1,850141402 | 0,001216096 | TRX-h5_ ATTRX5_ LIV1_ ATH5_ TRX5 thioredoxin H-type 5_ THIOREDOXIN H-TYPE 5_ LOCUS OF INSENSITIV |
| ATCG00820.1 | 0,500654561 | 0,001222103 | RPS19 ribosomal protein S19 chrc:84005-84283 REVERSE LENGTH=92 |
| AT5G10160.1 | 1,40401175 | 0,001247271 | no symbol available no full name available chr5:3185819-3187159 FORWARD LENGTH=219 |
| AT1G20450.1 | 2,344154499 | 0,001295533 | LTI45_ ERD10_ LTI29 EARLY RESPONSIVE TO DEHYDRATION 10_ LOW TEMPERATURE INDUCED 45_ LOW |
| AT5G43010.1 | 0,726438067 | 0,001310835 | RPT4A regulatory particle triple-A ATPase 4A chr5:17248563-17251014 REVERSE LENGTH=399 |
| AT3G46780.1 | 0,806062828 | 0,001313361 | PTAC16 plastid transcriptionally active 16 chr3:17228766-17231021 FORWARD LENGTH=510 |

| | | | |
|---|---|---|---|
| AT1G51980.1 | 1,23454433 | 0,001313365 | no symbol available no full name available chr1:19323692-19326771 REVERSE LENGTH=503 |
| AT4G37925.1 | 0,653713661 | 0,001318854 | NDH-M_ NdhM subunit NDH-M of NAD(P)H:plastoquinone dehydrogenase complex_ NADH dehydrogenase-like comple |
| AT2G47730.1 | 1,70745262 | 0,001331503 | GST6_ GSTF8_ ATGSTF8_ ATGSTF5 glutathione S-transferase phi 8_ Arabidopsis thaliana glutathione S-transferase phi |
| AT3G62530.1 | 0,568366453 | 0,001338641 | no symbol available no full name available chr3:23132219-23133121 FORWARD LENGTH=221 |
| AT5G48880.1 | 0,733254195 | 0,001354828 | PKT2_ PKT1_ KAT5 3-KETO-ACYL-COENZYME A THIOLASE 5_ peroxisomal 3-keto-acyl-CoA thiolase 2_ PEROXI |
| AT3G04720.1 | 1,279279721 | 0,00135706 | HEL_ PR-4_ AtPR4_ PR4 HEVEIN-LIKE_ pathogenesis-related 4 chr3:1285691-1286531 REVERSE LENGTH=212 |
| AT1G30360.1 | 1,233622313 | 0,001372217 | ERD4_ OSCA3.1 early-responsive to dehydration 4 chr1:10715892-10718799 FORWARD LENGTH=724 |
| AT1G65980.1 | 1,431184681 | 0,001403596 | TPX1 thioredoxin-dependent peroxidase 1 chr1:24559524-24560753 REVERSE LENGTH=162 |
| AT5G54500.1 | 1,404119184 | 0,001405071 | FQR1 flavodoxin-like quinone reductase 1 chr5:22124674-22126256 FORWARD LENGTH=204 |
| AT2G33380.1 | 3,593758088 | 0,001405967 | AtRD20_ PXG3_ CLO-3_ CLO3_ RD20_ AtCLO3 caleosin 3_ Arabidopsis thaliana caleosin 3_ peroxygenase 3_ RESPON |
| AT1G72150.1 | 0,589966916 | 0,001406335 | PATL1 PATELLIN 1 chr1:27148558-27150652 FORWARD LENGTH=573 |
| AT5G23860.1 | 0,754786485 | 0,001470497 | TUB8 tubulin beta 8 chr5:8042962-8044528 FORWARD LENGTH=449 |
| AT2G47390.1 | 0,789727729 | 0,001504125 | CGEP chloroplast glutamyl peptidase chr2:19442278-19446253 REVERSE LENGTH=961 |
| AT2G06050.1 | 0,856876438 | 0,001514986 | OPR3_ DDE1_ AtOPR3 DELAYED DEHISCENCE 1_ oxophytodienoate-reductase 3 chr2:2359240-2361971 REVERSE |
| AT1G64190.1 | 1,248986357 | 0,001521571 | PGD1 6-phosphogluconate dehydrogenase 1 chr1:23825549-23827012 REVERSE LENGTH=487 |
| AT3G12110.1 | 1,509596757 | 0,001553391 | ACT11 actin-11 chr3:3858116-3859609 FORWARD LENGTH=377 |
| AT3G62030.1 | 0,849728524 | 0,001556582 | ROC4_ CYP20-3 rotamase CYP 4_ cyclophilin 20-3 chr3:22973708-22975139 FORWARD LENGTH=260 |
| AT3G53460.1 | 0,380253054 | 0,001567945 | CP29 chloroplast RNA-binding protein 29 chr3:19819738-19821423 REVERSE LENGTH=342 |
| AT4G37990.1 | 3,52919581 | 0,001599823 | ELI3-2_ ATCAD8_ ELI3_ CAD-B2 elicitor-activated gene 3-2_ CINNAMYL-ALCOHOL DEHYDROGENASE B2_ ARA |
| AT4G01150.1 | 3,63355693 | 0,001635987 | CURT1A CURVATURE THYLAKOID 1A chr4:493692-494668 FORWARD LENGTH=164 |
| AT1G71500.1 | 0,674143525 | 0,001672346 | PSB33_ LIL8 PhotoSystem B protein 33_ Light-harvesting-like 8 chr1:26936084-26937331 FORWARD LENGTH=287 |
| AT5G43940.1 | 1,123723956 | 0,001672412 | ADH2_ HOT5_ ATGSNOR1_ PAR2_ GSNOR PARAQUAT RESISTANT 2_ S-NITROSOGLUTATHIONE REDUCTAS |
| AT1G24020.1 | 1,159100759 | 0,001678296 | MLP423 MLP-like protein 423 chr1:8500653-8501458 REVERSE LENGTH=155 |
| AT2G30870.1 | 1,227389566 | 0,001682109 | GSTF10_ ATGSTF10_ ERD13_ ATGSTF4 glutathione S-transferase PHI 10_ ARABIDOPSIS THALIANA GLUTATHIO |
| AT4G17090.1 | 1,752279753 | 0,001736349 | CT-BMY_ BMY8_ AtBAM3_ BAM3 BETA-AMYLASE 8_ BETA-AMYLASE 3_ chloroplast beta-amylase chr4:960526 |
| AT5G57850.1 | 0,625280219 | 0,001760525 | ADCL_ DAT1 D-AA specific transaminase D-AAT_ 4-amino-4-deoxychorismate lyase chr5:23435548-23437287 REVERS |
| AT3G08030.2 | 0,71385725 | 0,00179153 | AthA2-1 chr3:2564517-2565819 FORWARD LENGTH=323 |
| AT1G48630.1 | 1,521723615 | 0,001807452 | RACK1B_ RACK1B_AT receptor for activated C kinase 1B chr1:17981977-17983268 REVERSE LENGTH=326 |
| AT1G56330.1 | 1,299977037 | 0,001811525 | ATSARA1B_ SAR1B_ SAR1_ ATSAR1B_ ATSAR1 SECRETION-ASSOCIATED RAS 1_ ARABIDOPSIS THALIANA |
| AT1G07890.1 | 1,295042465 | 0,001817335 | MEE6_ ATAPX01_ ATAPX1_ CS1_ APX1 ascorbate peroxidase 1_ maternal effect embryo arrest 6 chr1:2438005-243943 |
| AT3G29360.1 | 1,223643607 | 0,001835798 | UGD2 UDP-glucose dehydrogenase 2 chr3:11267375-11268817 REVERSE LENGTH=480 |
| AT4G25080.1 | 0,53584177 | 0,001881464 | CHLM magnesium-protoporphyrin IX methyltransferase chr4:12877015-12878128 FORWARD LENGTH=312 |
| AT4G35830.1 | 1,523664515 | 0,001913614 | ACO1 aconitase 1 chr4:16973007-16977949 REVERSE LENGTH=898 |
| AT3G53420.1 | 1,087297373 | 0,00191664 | PIP2;1_ AtPIP2;1_ PIP2A_ PIP2 PLASMA MEMBRANE INTRINSIC PROTEIN 2_ PLASMA MEMBRANE INTRINSI |
| AT4G26530.1 | 0,723614642 | 0,001918331 | FBA5_ AtFBA5_ DEG22 fructose-bisphosphate aldolase 5 chr4:13391566-13392937 FORWARD LENGTH=358 |
| AT3G52880.1 | 1,452697228 | 0,001934882 | ATMDAR1_ MDAR1 monodehydroascorbate reductase 1 chr3:19601477-19604366 REVERSE LENGTH=434 |
| AT3G19760.1 | 4,330202332 | 0,001953831 | EIF4A-III_ RH2 eukaryotic initiation factor 4A-III chr3:6863790-6866242 FORWARD LENGTH=408 |
| AT5G56030.1 | 1,771186449 | 0,001974161 | HSP81-2_ HSP90.2_ AtHsp90.2_ ERD8_ HSP81.2 EARLY-RESPONSIVE TO DEHYDRATION 8_ heat shock protein 81 |
| AT3G08590.1 | 1,492326564 | 0,001982162 | iPGAM2 "2_3-biphosphoglycerate-independent phosphoglycerate mutase 2" chr3:2608683-2611237 REVERSE LENGTH= |

| | | | |
|---|---|---|---|
| AT4G37980.1 | 1,548689008 | 0,002008328 | ELI3-1_ CHR_ ELI3_ ATCAD7_ CAD7 elicitor-activated gene 3-1_ CINNAMALDEHYDE AND HEXENAL REDUCTA |
| AT1G63660.2 | 2,357848958 | 0,002021001 | no symbol available no full name available chr1:23604874-23607080 REVERSE LENGTH=434 |
| AT3G13750.1 | 0,346085077 | 0,002031968 | BGAL1 beta galactosidase 1_ beta-galactosidase 1 chr3:4511192-4515756 FORWARD LENGTH=847 |
| AT1G02560.1 | 1,110052567 | 0,002098686 | NCLPP1_ NCLPP5_ CLPP5 NUCLEAR CLPP 5_ NUCLEAR-ENCODED CLPP 1_ nuclear encoded CLP protease 5 chr1 |
| AT1G05010.1 | 0,497634069 | 0,002121201 | EFE_ ACO4_ EAT1 ethylene forming enzyme_ ethylene-forming enzyme chr1:1431419-1432695 REVERSE LENGTH=32 |
| AT5G65430.2 | 1,829377896 | 0,002151056 | 14-3-3KAPPA_ GRF8_ AtMIN10_ GF14 KAPPA general regulatory factor 8_ 14-3-3 PROTEIN G-BOX FACTOR14 KA |
| AT1G26570.1 | 0,793400821 | 0,002201402 | UGD1_ ATUGD1 UDP-GLUCOSE DEHYDROGENASE 1_ UDP-glucose dehydrogenase 1 chr1:9182801-9184246 FOR |
| AT4G24280.1 | 1,215140012 | 0,002207617 | cpHsc70-1 chloroplast heat shock protein 70-1 chr4:12590094-12593437 FORWARD LENGTH=718 |
| AT3G46060.1 | 0,675207479 | 0,002231197 | ARA3_ RAB8A_ RABE1c_ ARA-3_ ATRAB8A_ ATRABE1C RAB GTPase homolog 8A chr3:16917908-16919740 FOR |
| AT4G30190.1 | 1,331683486 | 0,002237246 | HA2_ PMA2_ AtHA2_ AHA2 H(+)-ATPase 2_ PLASMA MEMBRANE PROTON ATPASE 2 chr4:14770820-14775920 |
| AT3G54640.1 | 0,241895233 | 0,002251492 | TRP3_ TSA1 TRYPTOPHAN-REQUIRING 3_ tryptophan synthase alpha chain chr3:20223331-20225303 REVERSE LE |
| AT1G80560.1 | 0,792029592 | 0,002295226 | ATIMD2_ IMD2 isopropylmalate dehydrogenase 2_ ARABIDOPSIS ISOPROPYLMALATE DEHYDROGENASE 2 chr1 |
| AT1G56580.1 | 4,412154266 | 0,002314704 | SVB SMALLER WITH VARIABLE BRANCHES chr1:21198402-21198902 REVERSE LENGTH=166 |
| AT1G20020.1 | 0,75774957 | 0,002329862 | LFNR2_ FNR2_ ATLFNR2 leaf-type chloroplast-targeted FNR 2_ ferredoxin-NADP(+)-oxidoreductase 2_ LEAF FNR 2 c |
| AT5G23060.1 | 0,543463553 | 0,002362066 | CaS calcium sensing receptor chr5:7736760-7738412 REVERSE LENGTH=387 |
| AT1G01800.1 | 1,332737641 | 0,002372247 | no symbol available no full name available chr1:293396-294888 FORWARD LENGTH=295 |
| AT1G09100.1 | 1,559553422 | 0,002380854 | RPT5B 26S proteasome AAA-ATPase subunit RPT5B chr1:2936675-2939258 REVERSE LENGTH=423 |
| AT1G02930.1 | 1,432341628 | 0,0024406 | ATGSTF3_ ATGSTF6_ GST1_ ERD11_ GSTF6_ ATGST1 EARLY RESPONSIVE TO DEHYDRATION 11_ ARABID |
| AT4G27520.1 | 0,721844446 | 0,002512855 | ENODL2_ AtENODL2 early nodulin-like protein 2 chr4:13750668-13751819 REVERSE LENGTH=349 |
| AT2G16600.1 | 0,757496217 | 0,002569907 | AtCYP19-1_ ROC3_ CYP19 rotamase CYP 3_ cyclophilin 19 chr2:7200862-7201383 FORWARD LENGTH=173 |
| AT2G41220.1 | 0,833874493 | 0,002570924 | GLU2 glutamate synthase 2 chr2:17177934-17188388 FORWARD LENGTH=1629 |
| AT5G08670.1 | 1,226044308 | 0,002585465 | no symbol available no full name available chr5:2818395-2821149 REVERSE LENGTH=556 |
| AT2G44610.1 | 1,248239124 | 0,002595589 | ATRAB6A_ RAB6_ RAB6A_ ATRABH1B chr2:18411778-18413883 REVERSE LENGTH=208 |
| AT3G52960.1 | 1,234738671 | 0,002625505 | PrxIIE peroxiredoxin-II-E chr3:19639699-19640403 FORWARD LENGTH=234 |
| AT2G05840.3 | 1,488383977 | 0,002678558 | PAA2 20S proteasome subunit PAA2 chr2:2234226-2235533 FORWARD LENGTH=205 |
| AT3G55040.1 | 0,728402766 | 0,002732293 | GSTL2 glutathione transferase lambda 2 chr3:20398718-20400305 REVERSE LENGTH=292 |
| AT1G66970.1 | 0,738486622 | 0,002736484 | SVL2_ GDPDL1 SHV3-like 2_ Glycerophosphodiester phosphodiesterase (GDPD) like 1 chr1:24992746-24996005 REVE |
| AT3G12915.2 | 1,317704166 | 0,002800044 | no symbol available no full name available chr3:4112834-4115708 FORWARD LENGTH=788 |
| AT1G29660.1 | 0,733569541 | 0,002886025 | GGL5 chr1:10371955-10373624 FORWARD LENGTH=364 |
| AT1G58290.1 | 0,357880524 | 0,002904856 | HEMA1_ GluTR_ AtHEMA1 glutamyl-tRNA reductase_ Arabidopsis thaliana hemA 1 chr1:21624028-21626051 REVERS |
| AT3G28220.1 | 0,782633029 | 0,00302729 | no symbol available no full name available chr3:10524420-10526497 FORWARD LENGTH=370 |
| AT1G47128.1 | 0,569820952 | 0,003060065 | RD21_ RD21A responsive to dehydration 21A_ responsive to dehydration 21 chr1:17283139-17285609 REVERSE LENG |
| AT4G24780.1 | 0,558188147 | 0,003064142 | PLL19 chr4:12770631-12772227 REVERSE LENGTH=408 |
| ATCG00740.1 | 0,714946863 | 0,003133429 | RPOA RNA polymerase subunit alpha chrc:77901-78890 REVERSE LENGTH=329 |
| AT1G77490.2 | 0,839069359 | 0,003149653 | TAPX thylakoidal ascorbate peroxidase chr1:29117688-29120649 FORWARD LENGTH=421 |
| AT4G34230.1 | 2,061079098 | 0,003182342 | CAD-5_ ATCAD5_ CAD5 cinnamyl alcohol dehydrogenase 5 chr4:16386898-16388666 REVERSE LENGTH=357 |
| AT3G55440.1 | 1,355136185 | 0,003199549 | ATCTIMC_ CYTOTPI_ TPI CYTOSOLIC ISOFORM TRIOSE PHOSPHATE ISOMERASE_ triosephosphate isomerase_ |
| AT1G32200.1 | 0,695430816 | 0,003288399 | ACT1_ ATS1 ACYLTRANSFERASE 1 chr1:11602223-11605001 REVERSE LENGTH=459 |
| AT5G62690.1 | 1,472763015 | 0,003309809 | TUB2 tubulin beta chain 2 chr5:25181560-25183501 FORWARD LENGTH=450 |

| | | | |
|---|---|---|---|
| AT3G54400.1 | 0,812125664 | 0,003316685 | no symbol available no full name available chr3:20140291-20142599 REVERSE LENGTH=425 |
| AT1G02780.1 | 0,538889938 | 0,00333176 | emb2386 embryo defective 2386 chr1:608120-609391 REVERSE LENGTH=214 |
| AT5G62350.1 | 0,361139458 | 0,003340392 | no symbol available no full name available chr5:25037504-25038112 FORWARD LENGTH=202 |
| AT5G45280.1 | 0,402990572 | 0,003348419 | PAE11 pectin acetylesterase 11 chr5:18346862-18349432 FORWARD LENGTH=370 |
| AT1G04710.1 | 1,570662017 | 0,003475817 | KAT1_ PKT4 3-KETO-ACYL-COA THIOLASE 1_ peroxisomal 3-ketoacyl-CoA thiolase 4 chr1:1321941-1324556 FORW |
| AT5G08380.1 | 1,216320688 | 0,003512699 | AtAGAL1_ AGAL1 alpha-galactosidase 1 chr5:2694851-2697616 REVERSE LENGTH=410 |
| AT5G20010.1 | 1,278007281 | 0,003550891 | RAN-1_ ATRAN1_ RAN1 RAS-RELATED NUCLEAR PROTEIN_ ARABIDOPSIS THALIANA RAS-RELATED NUC |
| AT5G40370.1 | 1,687254291 | 0,003553131 | AtGRXC2_ GRXC2_ GRX370 glutaredoxin C2 chr5:16147826-16149052 REVERSE LENGTH=111 |
| AT5G01650.1 | 1,421848136 | 0,003596717 | MDL2 MIF/D-DT-like 2 chr5:242734-244033 REVERSE LENGTH=115 |
| AT1G43560.1 | 1,606012341 | 0,003641946 | Aty2_ ty2 thioredoxin Y2 chr1:16398359-16399828 REVERSE LENGTH=167 |
| AT3G56070.1 | 0,674246214 | 0,003782418 | ROC2 rotamase cyclophilin 2 chr3:20806987-20807517 REVERSE LENGTH=176 |
| AT2G45710.1 | 1,227144898 | 0,003800019 | no symbol available no full name available chr2:18831243-18831999 FORWARD LENGTH=84 |
| AT1G10760.1 | 0,636276102 | 0,003803987 | GWD_ GWD1_ SOP1_ SOP_ SEX1 STARCH EXCESS 1 chr1:3581210-3590043 REVERSE LENGTH=1399 |
| AT2G20610.1 | 1,296057076 | 0,003816036 | RTY1_ RTY_ HLS3_ ALF1_ SUR1 ABERRANT LATERAL ROOT FORMATION 1_ SUPERROOT 1_ ROOTY 1_ RO |
| AT4G14880.1 | 1,191339936 | 0,003855612 | OASA1_ OLD3_ CYTACS1_ ATCYS-3A ONSET OF LEAF DEATH 3_ O-acetylserine (thiol) lyase (OAS-TL) isoform A |
| AT3G19710.1 | 1,370478485 | 0,003873095 | BCAT4 branched-chain aminotransferase4 chr3:6847202-6849429 REVERSE LENGTH=354 |
| AT1G56340.1 | 1,297897102 | 0,003955122 | AtCRT1a_ CRT1_ CRT1a calreticulin 1a_ calreticulin 1 chr1:21090059-21092630 REVERSE LENGTH=425 |
| AT3G12915.1 | 0,700954512 | 0,003970874 | no symbol available no full name available chr3:4112999-4115708 FORWARD LENGTH=820 |
| AT3G09820.1 | 0,690060858 | 0,003976511 | ATADK1_ ADK1 adenosine kinase 1 chr3:3012122-3014624 FORWARD LENGTH=344 |
| AT1G76080.1 | 0,588167634 | 0,003997488 | CDSP32_ TRXL1_ ATCDSP32 ARABIDOPSIS THALIANA CHLOROPLASTIC DROUGHT-INDUCED STRESS PRO |
| AT2G24020.1 | 1,439096586 | 0,004066607 | STIC2 Suppressor of TIC40 2 chr2:10217869-10219269 REVERSE LENGTH=182 |
| AT1G79210.1 | 1,249792737 | 0,004166492 | no symbol available no full name available chr1:29796286-29798240 REVERSE LENGTH=235 |
| AT4G09670.1 | 1,263994748 | 0,004182883 | no symbol available no full name available chr4:6107382-6109049 REVERSE LENGTH=362 |
| AT2G36580.1 | 1,562963882 | 0,004327533 | no symbol available no full name available chr2:15339253-15342781 FORWARD LENGTH=527 |
| AT1G55060.1 | 1,19291607 | 0,004340675 | UBQ12 ubiquitin 12 chr1:20549533-20550225 FORWARD LENGTH=230 |
| AT1G32900.1 | 0,825969813 | 0,004362964 | GBSS1 granule bound starch synthase 1 chr1:11920582-11923506 REVERSE LENGTH=610 |
| AT5G19550.1 | 1,548734861 | 0,004371978 | AAT2_ ASP2 ASPARTATE AMINOTRANSFERASE 2_ aspartate aminotransferase 2 chr5:6598201-6601597 FORWAR |
| AT3G23600.1 | 1,223084463 | 0,004408161 | no symbol available no full name available chr3:8473833-8475655 FORWARD LENGTH=239 |
| AT3G16530.1 | 0,675418363 | 0,004451353 | no symbol available no full name available chr3:5624586-5625416 REVERSE LENGTH=276 |
| AT3G09260.1 | 0,461666778 | 0,004470729 | LEB_ BGLU23_ PYK10_ PSR3.1 LONG ER BODY chr3:2840657-2843730 REVERSE LENGTH=524 |
| AT5G52650.1 | 1,582968965 | 0,004482975 | no symbol available no full name available chr5:21355781-21357003 REVERSE LENGTH=179 |
| AT2G35410.1 | 0,798763482 | 0,004492346 | no symbol available no full name available chr2:14898341-14899590 FORWARD LENGTH=308 |
| AT1G19550.1 | 1,222462782 | 0,004498236 | DHAR dehydroascorbate reductase chr1:6767451-6767983 REVERSE LENGTH=153 |
| AT5G45750.1 | 0,337953402 | 0,00454263 | RABA1c_ AtRABA1c RAB GTPase homolog A1C chr5:18559318-18560639 FORWARD LENGTH=216 |
| AT2G26080.1 | 0,722773008 | 0,004578859 | GLDP2_ AtGLDP2 glycine decarboxylase P-protein 2 chr2:11109330-11113786 REVERSE LENGTH=1044 |
| AT1G53240.1 | 1,194070086 | 0,004582918 | mMDH1 mitochondrial malate dehydrogenase 1 chr1:19854966-19856802 REVERSE LENGTH=341 |
| AT3G20050.1 | 1,235493494 | 0,004601543 | ATTCP-1_ TCP-1_ CCT1 Chaperonin containing T-complex polypeptide-1 subunit 1_ T-complex protein 1 alpha subunit c |
| AT3G12145.1 | 0,758813234 | 0,00463983 | FLR1_ FTM4 FLOR1_ FLORAL TRANSITION AT THE MERISTEM4 chr3:3874764-3876075 REVERSE LENGTH=32 |
| AT1G22760.1 | 1,571852648 | 0,004736739 | PAB3_ PABP3 poly(A) binding protein 3 chr1:8055599-8058799 FORWARD LENGTH=660 |

| | | | |
|---|---|---|---|
| AT2G29630.1 | 0,390335254 | 0,0047996 | THIC_ PY PYRIMIDINE REQUIRING_ thiaminC chr2:12667395-12669569 FORWARD LENGTH=644 |
| AT1G34430.1 | 0,661894759 | 0,004952269 | EMB3003 embryo defective 3003 chr1:12588027-12590084 REVERSE LENGTH=465 |
| AT1G79550.1 | 1,682433223 | 0,004956086 | PGKc_ PGK_ PGK3 phosphoglycerate kinase_ phosphoglycerate kinase 3 chr1:29924347-29926295 REVERSE LENGTH= |
| AT2G42590.1 | 1,204101696 | 0,005009254 | GF14 MU_ GRF9_ GRF14 general regulatory factor 9 chr2:17732118-17733775 REVERSE LENGTH=263 |
| AT3G44890.1 | 0,760822321 | 0,005032152 | RPL9 ribosomal protein L9 chr3:16386505-16387963 FORWARD LENGTH=197 |
| AT2G30950.1 | 1,961623972 | 0,005032458 | FTSH2_ VAR2 VARIEGATED 2 chr2:13174692-13177064 FORWARD LENGTH=695 |
| AT1G13930.1 | 1,684435829 | 0,005032874 | no symbol available no full name available chr1:4761091-4761558 FORWARD LENGTH=155 |
| AT5G28500.1 | 0,895911302 | 0,005035649 | no symbol available no full name available chr5:10477810-10479114 FORWARD LENGTH=434 |
| AT4G14040.1 | 1,288610766 | 0,00506623 | EDA38_ SBP2 selenium-binding protein 2_ EMBRYO SAC DEVELOPMENT ARREST 38 chr4:8100691-8102828 REVE |
| AT1G75270.1 | 1,316805436 | 0,005098659 | DHAR2 dehydroascorbate reductase 2 chr1:28250255-28251237 REVERSE LENGTH=213 |
| AT1G13060.1 | 1,317204884 | 0,005159352 | PBE1 20S proteasome beta subunit E1 chr1:4452641-4454663 FORWARD LENGTH=274 |
| AT1G20440.1 | 2,030070536 | 0,005385442 | AtCOR47_ RD17_ COR47 cold-regulated 47 chr1:7084722-7085664 REVERSE LENGTH=265 |
| AT4G31180.1 | 1,670324509 | 0,005389216 | IBI1 impaired in BABA-induced disease immunity 1 chr4:15156696-15159362 FORWARD LENGTH=558 |
| AT5G66530.1 | 0,788770053 | 0,005402809 | no symbol available no full name available chr5:26553821-26555575 REVERSE LENGTH=307 |
| AT4G17470.1 | 0,529410063 | 0,005553073 | CRSH Ca2+-activated RelA-spot homolog chr4:9742922-9744468 REVERSE LENGTH=308 |
| AT5G16590.1 | 1,175562532 | 0,005579035 | LRR1 Leucine rich repeat protein 1 chr5:5431862-5433921 FORWARD LENGTH=625 |
| AT5G57350.3 | 0,536244738 | 0,005753618 | ATAHA3_ HA3_ AHA3 H(+)-ATPase 3_ ARABIDOPSIS THALIANA ARABIDOPSIS H(+)-ATPASE chr5:23231208-2 |
| AT1G76010.1 | 0,642580994 | 0,005908195 | ALBA1_ Atalba1_ ALBA4 chr1:28528505-28530488 REVERSE LENGTH=350 |
| AT1G19670.1 | 0,726834565 | 0,005940214 | CLH1_ ATCLH1_ CORI1_ ATHCOR1 chlorophyllase 1_ CORONATINE-INDUCED PROTEIN 1 chr1:6803796-680492 |
| AT2G33040.1 | 1,501811471 | 0,005988883 | ATP3 gamma subunit of Mt ATP synthase chr2:14018978-14021047 REVERSE LENGTH=325 |
| AT3G13930.1 | 1,250266823 | 0,006020752 | mtE2-2 mitochondrial pyruvate dehydrogenase subunit 2-2 chr3:4596240-4600143 FORWARD LENGTH=539 |
| AT3G16470.1 | 0,879431949 | 0,006092644 | JAL35_ AtJAC1_ JR1 jacalin-related lectin 35_ JACALIN-LECTIN LIKE 1_ JASMONATE RESPONSIVE 1 chr3:55960 |
| AT3G14290.1 | 1,183897046 | 0,00615065 | PAE2 20S proteasome alpha subunit E2 chr3:4764364-4766381 FORWARD LENGTH=237 |
| AT1G22530.1 | 0,808779192 | 0,006155986 | PATL2 PATELLIN 2 chr1:7955773-7958326 REVERSE LENGTH=683 |
| AT1G27950.1 | 2,277097664 | 0,006168268 | LTPG1 glycosylphosphatidylinositol-anchored lipid protein transfer 1 chr1:9740740-9741991 FORWARD LENGTH=193 |
| AT2G45300.3 | 0,702925477 | 0,006201219 | no symbol available no full name available chr2:18677518-18680118 FORWARD LENGTH=518 |
| AT1G48600.1 | 1,824740691 | 0,006233142 | AtPMT2_ AtPMEAMT_ PMEAMT phosphoethanolamine N-methyltransferase_ Phosphoethanolamine methyltransferase2 |
| AT4G01900.1 | 1,38994581 | 0,006237983 | PII_ GLB1 GLNB1 homolog chr4:821736-823294 FORWARD LENGTH=196 |
| AT1G45000.1 | 1,189071368 | 0,00624333 | RPT4b chr1:17009220-17011607 FORWARD LENGTH=399 |
| AT2G32920.1 | 1,903177423 | 0,006281223 | PDIL2-3_ PDI9_ ATPDIL2-3_ ATPDI9 ARABIDOPSIS THALIANA PROTEIN DISULFIDE ISOMERASE 9_ PDI-like |
| AT1G09310.1 | 0,808378313 | 0,006329755 | SVB2_ SVBL SVB-like chr1:3009109-3009648 FORWARD LENGTH=179 |
| AT2G37170.1 | 1,653349121 | 0,00657443 | PIP2;2_ PIP2B plasma membrane intrinsic protein 2_ PLASMA MEMBRANE INTRINSIC PROTEIN 2;2 chr2:15613624- |
| AT1G52570.1 | 1,169240164 | 0,006643885 | PLDALPHA2 phospholipase D alpha 2 chr1:19583940-19586551 REVERSE LENGTH=810 |
| AT4G13430.1 | 0,772424122 | 0,006717674 | IIL1_ ATLEUC1 isopropyl malate isomerase large subunit 1 chr4:7804194-7807789 REVERSE LENGTH=509 |
| AT1G56450.1 | 1,169983768 | 0,006729095 | MUD1_ PBG1 20S proteasome beta subunit G1 chr1:21141970-21144186 FORWARD LENGTH=246 |
| AT1G13440.2 | 0,747070284 | 0,006770516 | GAPC2_ GAPC-2 glyceraldehyde-3-phosphate dehydrogenase C2_ GLYCERALDEHYDE-3-PHOSPHATE DEHYDROG |
| AT4G37910.1 | 1,254324504 | 0,00690042 | mtHsc70-1 mitochondrial heat shock protein 70-1 chr4:17825368-17828099 REVERSE LENGTH=682 |
| AT1G62750.1 | 0,696334628 | 0,006953282 | ATSCO1/CPEF-G_ ATSCO1_ SCO1 SNOWY COTYLEDON 1 chr1:23233622-23236321 REVERSE LENGTH=783 |
| AT5G26570.1 | 0,770925506 | 0,007050489 | ATGWD3_ OK1_ PWD PHOSPHOGLUCAN WATER DIKINASE chr5:9261580-9267526 FORWARD LENGTH=1196 |

| | | | |
|---|---|---|---|
| AT3G45140.1 | 0,742307619 | 0,007098726 | ATLOX2_ LOX2 ARABIDOPSIS THALIANA LIPOXYGENASE 2_ lipoxygenase 2 chr3:16525437-16529233 FORWA |
| AT4G17520.1 | 0,513296846 | 0,007149417 | HLN HYALURONAN/mRNA BINDING FAMILY PROTEIN chr4:9771496-9773313 FORWARD LENGTH=360 |
| AT3G05530.1 | 1,219984092 | 0,007369216 | RPT5A_ ATS6A.2 regulatory particle triple-A ATPase 5A chr3:1603540-1605993 FORWARD LENGTH=424 |
| AT1G70820.1 | 0,630678858 | 0,007395034 | no symbol available no full name available chr1:26705594-26708034 FORWARD LENGTH=615 |
| AT4G14030.1 | 1,300348411 | 0,0074338 | SBP1_ AtSBP1 selenium-binding protein 1 chr4:8098121-8100165 REVERSE LENGTH=490 |
| AT3G63140.1 | 0,867196792 | 0,007593488 | CSP41A chloroplast stem-loop binding protein of 41 kDa chr3:23327006-23328620 REVERSE LENGTH=406 |
| AT2G20360.1 | 1,2274275 | 0,007593875 | no symbol available no full name available chr2:8786070-8789098 FORWARD LENGTH=402 |
| AT3G01390.1 | 2,811494137 | 0,007697054 | AVMA10_ VMA10 vacuolar membrane ATPase 10 chr3:150265-150922 REVERSE LENGTH=110 |
| AT2G40010.1 | 1,411237969 | 0,007728063 | no symbol available no full name available chr2:16708578-16710448 REVERSE LENGTH=317 |
| AT4G03520.1 | 0,795116631 | 0,007738787 | ATHM2_ TRXm2 thioredoxin m2 chr4:1562585-1564055 REVERSE LENGTH=186 |
| AT1G25490.1 | 1,425430146 | 0,007746267 | EER1_ ATB BETA BETA_ RCN1_ REGA ROOTS CURL IN NPA_ ENHANCED ETHYLENE RESPONSE 1 chr1:8951 |
| ATCG00280.1 | 0,67443049 | 0,007754739 | PSBC photosystem II reaction center protein C chrc:33720-35141 FORWARD LENGTH=473 |
| AT3G56940.1 | 0,570312721 | 0,007851177 | CRD1_ ACSF_ CHL27 COPPER RESPONSE DEFECT 1 chr3:21076594-21078269 FORWARD LENGTH=409 |
| AT1G70410.1 | 1,118366842 | 0,0079417 | BCA4_ ATBCA4_ CA4 beta carbonic anhydrase 4_ BETA CARBONIC ANHYDRASE 4 chr1:26534167-26536505 REVI |
| AT1G80380.4 | 0,798562928 | 0,008055693 | GLYK glycerate kinase chr1:30217332-30219784 FORWARD LENGTH=450 |
| AT5G56010.1 | 1,462847198 | 0,008081558 | AtHsp90-3_ AtHsp90.3_ Hsp81.3_ HSP81-3 HEAT SHOCK PROTEIN 90-3_ HEAT SHOCK PROTEIN 81.3_ heat shock |
| AT1G04410.1 | 1,226740698 | 0,008084595 | c-NAD-MDH1 cytosolic-NAD-dependent malate dehydrogenase 1 chr1:1189418-1191267 REVERSE LENGTH=332 |
| AT3G08580.1 | 1,224441257 | 0,008117801 | AAC1 ADP/ATP carrier 1 chr3:2605706-2607030 REVERSE LENGTH=381 |
| AT5G14200.1 | 0,851441208 | 0,008134375 | ATIMD1_ IMD1 isopropylmalate dehydrogenase 1_ ARABIDOPSIS ISOPROPYLMALATE DEHYDROGENASE 1 chr5 |
| AT4G05180.1 | 0,830975663 | 0,008275205 | PSBQ-2_ PSBQ_ PSII-Q photosystem II subunit Q-2_ PHOTOSYSTEM II SUBUNIT Q chr4:2672093-2673170 REVERS |
| AT5G51110.1 | 0,666203037 | 0,008296493 | ATP1_ SDIRIP1_ RAF2 SDIR1-INTERACTING PROTEIN1_ AtAIRP2 Target Protein 1_ Rubisco Assembly Factor 2 chr |
| AT3G02090.1 | 0,867398119 | 0,008302375 | MPPBETA chr3:365624-368526 FORWARD LENGTH=531 |
| AT5G49460.1 | 1,447239769 | 0,008304573 | ACLB-2 ATP citrate lyase subunit B 2 chr5:20055048-20058195 FORWARD LENGTH=608 |
| AT2G43945.1 | 0,691858832 | 0,00835248 | no symbol available no full name available chr2:18197995-18199988 REVERSE LENGTH=289 |
| AT3G27690.1 | 1,248844012 | 0,008419079 | LHCB2.4_ DEG13_ LHCB2_ LHCB2.3 LIGHT-HARVESTING CHLOROPHYLL B-BINDING 2_ photosystem II light ha |
| AT1G03130.1 | 1,255077387 | 0,008437282 | PSAD-2 photosystem I subunit D-2 chr1:753528-754142 REVERSE LENGTH=204 |
| AT4G33090.1 | 1,356039772 | 0,008764688 | APM1_ ATAPM1 AMINOPEPTIDASE M1_ aminopeptidase M1 chr4:15965915-15970418 REVERSE LENGTH=879 |
| AT4G12730.1 | 0,759105555 | 0,008860275 | FLA2 FASCICLIN-like arabinogalactan 2 chr4:7491598-7492809 REVERSE LENGTH=403 |
| AT1G09080.2 | 0,746948888 | 0,008925369 | BIP3 binding protein 3 chr1:2929268-2931804 REVERSE LENGTH=665 |
| AT1G35160.1 | 1,22036776 | 0,009050703 | GRF4_ 14-3-3PHI_ GF14 PHI GENERAL REGULATORY FACTOR 4_ 14-3-3 PROTEIN G-BOX FACTOR14 PHI_ GF |
| AT4G23850.1 | 1,355061243 | 0,009080125 | LACS4 long-chain acyl-CoA synthetase 4 chr4:12403720-12408263 REVERSE LENGTH=666 |
| AT1G16080.1 | 0,805495322 | 0,009149037 | no symbol available no full name available chr1:5514394-5515761 FORWARD LENGTH=313 |
| AT2G02930.1 | 1,189456224 | 0,009213253 | ATGSTF3_ GST16_ GSTF3 GLUTATHIONE S-TRANSFERASE 16_ glutathione S-transferase F3 chr2:851348-852106 F |
| AT3G27280.1 | 1,900456915 | 0,009239371 | PHB4_ ATPHB4 prohibitin 4 chr3:10076904-10078051 FORWARD LENGTH=279 |
| AT1G54220.1 | 0,670209019 | 0,009279735 | mtE2-3 mitochondrial pyruvate dehydrogenase subunit 2-3 chr1:20246460-20250208 REVERSE LENGTH=539 |
| AT5G54190.1 | 0,610083907 | 0,009383264 | PORA protochlorophyllide oxidoreductase A chr5:21991183-21992773 REVERSE LENGTH=405 |
| AT1G31812.1 | 1,676108927 | 0,009422526 | ACBP6_ ACBP_ AtACBP6 acyl-CoA-binding protein 6_ ACYL-COA-BINDING PROTEIN chr1:11411132-11412099 RE |
| AT2G39800.2 | 0,813707347 | 0,009426476 | P5CS1_ ATP5CS delta1-pyrroline-5-carboxylate synthase 1 chr2:16598516-16601976 REVERSE LENGTH=614 |
| AT2G01520.1 | 3,865542834 | 0,00961394 | MLP328_ ZCE1 (Zusammen-CA)-enhanced 1_ MLP-like protein 328 chr2:235992-236881 FORWARD LENGTH=151 |

| | | | |
|---|---|---|---|
| AT4G18480.1 | 0,681931857 | 0,009634617 | CHL11_ CH-42_ LOST1_ CH42_ CHLI-1_ CHLI1 low temperature with open-stomata 1_ CHLORINA 42 chr4:10201897- |
| AT3G08530.1 | 1,501766699 | 0,009642589 | AtCHC2_ CHC2 clathrin heavy chain 2 chr3:2587171-2595411 REVERSE LENGTH=1703 |
| AT5G60360.2 | 0,205825755 | 0,009659231 | AALP_ SAG2_ ALP aleurain-like protease_ SENESCENCE ASSOCIATED GENE2 chr5:24280044-24282152 FORWARI |
| AT5G11670.1 | 1,330625171 | 0,009735877 | NADP-ME2_ ATNADP-ME2 NADP-malic enzyme 2_ Arabidopsis thaliana NADP-malic enzyme 2 chr5:3754456-375804( |
| AT5G15970.1 | 6,741084807 | 0,009759952 | AtCor6.6_ KIN2_ COR6.6 COLD-RESPONSIVE 6.6 chr5:5211966-5212441 FORWARD LENGTH=66 |
| AT5G42980.1 | 1,24455814 | 0,00978152 | ATH3_ TRX3_ TRXH3_ ATTRX3_ ATTRXH3 THIOREDOXIN H3_ thioredoxin 3_ thioredoxin H-type 3 chr5:1724277 |
| AT1G02920.1 | 1,297919981 | 0,009896007 | ATGSTF8_ GSTF7_ ATGSTF7_ ATGST11_ GST11 GLUTATHIONE S-TRANSFERASE 11_ glutathione S-transferase 7 |
| AT5G47210.1 | 1,166203873 | 0,00998257 | no symbol available no full name available chr5:19169222-19171012 REVERSE LENGTH=357 |
| AT4G26970.1 | 3,721191606 | 0,010029786 | ACO2 aconitase 2 chr4:13543077-13548427 FORWARD LENGTH=995 |
| AT2G26150.1 | 3,858843433 | 0,010227513 | ATHSFA2_ HSFA2 heat shock transcription factor A2 chr2:11135856-11137217 FORWARD LENGTH=345 |
| AT5G54160.1 | 1,214398796 | 0,010250593 | ATOMT1_ OMT1_ OMT3_ AtCOMT_ COMT1 caffeate O-methyltransferase 1_ O-methyltransferase 1_ O-methyltransfer |
| AT1G07410.1 | 0,583826199 | 0,010269978 | ATRAB-A2B_ ATRABA2B_ RABA2b_ RAB-A2B ARABIDOPSIS RAB GTPASE HOMOLOG A2B_ RAB GTPase hon |
| AT5G47870.1 | 1,65497084 | 0,010355597 | RAD52-2_ ODB2_ RAD52-2B radiation sensitive 52-2_ Organellar DNA-Binding protein 2 chr5:19384555-19385808 RE |
| AT4G18440.1 | 0,747343661 | 0,0104695 | no symbol available no full name available chr4:10186385-10188832 REVERSE LENGTH=536 |
| AT4G02530.1 | 1,240091783 | 0,010556625 | MPH2 MAINTENANCE OF PHOTOSYSTEM II UNDER HIGH LIGHT 2 chr4:1112335-1114005 REVERSE LENGTH= |
| AT4G35000.1 | 1,336226617 | 0,010672928 | APX3 ascorbate peroxidase 3 chr4:16665007-16667541 REVERSE LENGTH=287 |
| AT3G26060.1 | 0,811175865 | 0,010787373 | ATPRX Q_ PRXQ peroxiredoxin Q chr3:9524807-9526123 FORWARD LENGTH=216 |
| AT4G12420.1 | 0,82993454 | 0,010861656 | SKU5 chr4:7349941-7352868 REVERSE LENGTH=587 |
| AT2G20140.1 | 1,124963905 | 0,011033436 | RPT2b regulatory particle AAA-ATPase 2b chr2:8692736-8694837 FORWARD LENGTH=443 |
| AT2G42910.1 | 3,381384501 | 0,011044412 | AtPRS4_ PRS4 phosphoribosyl diphosphate synthase 4 chr2:17856396-17858394 FORWARD LENGTH=337 |
| AT1G01470.1 | 2,315504355 | 0,01111179 | LEA14_ LSR3_ AtLEA14_ LEA1 LIGHT STRESS-REGULATED 3_ Arabidopsis thaliana Late Embryogenesis abundant |
| AT3G44300.1 | 1,281549771 | 0,011386058 | AtNIT2_ NIT2 nitrilase 2 chr3:15983351-15985172 FORWARD LENGTH=339 |
| AT1G10840.1 | 1,358397324 | 0,01142513 | TIF3H1 translation initiation factor 3 subunit H1 chr1:3607885-3610299 REVERSE LENGTH=337 |
| AT5G27470.1 | 1,152914755 | 0,011443774 | no symbol available no full name available chr5:9695087-9697154 FORWARD LENGTH=451 |
| AT4G34450.1 | 1,461374328 | 0,011713829 | gamma2-COP gamma2 Coat Protein chr4:16471956-16476795 FORWARD LENGTH=886 |
| AT3G46440.1 | 1,472092626 | 0,012098568 | UXS5 UDP-XYL synthase 5 chr3:17089268-17091611 REVERSE LENGTH=341 |
| AT3G17210.1 | 1,271032317 | 0,012137096 | ATHS1_ HS1 heat stable protein 1_ A. THALIANA HEAT STABLE PROTEIN 1 chr3:5882318-5882896 FORWARD LE |
| AT2G28950.1 | 0,783546812 | 0,012252123 | ATEXP6_ ATHEXP ALPHA 1.8_ ATEXPA6_ EXPA6 expansin A6_ ARABIDOPSIS THALIANA TEXPANSIN 6 chr2:1 |
| ATCG00540.1 | 0,888735092 | 0,012563129 | PETA photosynthetic electron transfer A chrc:61657-62619 FORWARD LENGTH=320 |
| AT5G16050.1 | 1,370949579 | 0,012578607 | GRF5_ GF14 UPSILON general regulatory factor 5 chr5:5244008-5245402 REVERSE LENGTH=268 |
| AT5G67360.1 | 0,803905211 | 0,0126304 | ARA12_ SBT1.7 Subtilisin-like Serine protease 1.7 chr5:26872192-26874465 REVERSE LENGTH=757 |
| AT3G16950.1 | 1,35575773 | 0,012633439 | ptlpd1_ LPD1 lipoamide dehydrogenase 1 chr3:5786761-5790383 REVERSE LENGTH=570 |
| AT5G13870.1 | 1,789524733 | 0,01276947 | EXGT-A4_ XTH5 endoxyloglucan transferase A4_ xyloglucan endotransglucosylase/hydrolase 5 chr5:4475089-4476217 R |
| AT1G19570.1 | 0,852571386 | 0,012873997 | ATDHAR1_ DHAR1_ DHAR5 DEHYDROASCORBATE REDUCTASE 5_ dehydroascorbate reductase chr1:6773462-67 |
| AT1G14810.1 | 1,273721479 | 0,012906884 | no symbol available no full name available chr1:5102684-5104633 REVERSE LENGTH=375 |
| AT1G08520.1 | 0,596423819 | 0,012911613 | ALB1_ PDE166_ ALB-1V_ CHLD_ V157 PIGMENT DEFECTIVE EMBRYO 166_ ALBINA 1 chr1:2696538-2700819 F |
| AT5G58250.1 | 0,718077808 | 0,012925151 | LCAA/YCF54_ EMB3143 low chlorophyll accumulation/hypothetical chloroplast open reading frame 54_ EMBRYO DEFI |
| AT3G16410.1 | 0,789644425 | 0,012971385 | NSP4 nitrile specifier protein 4 chr3:5572145-5574359 FORWARD LENGTH=619 |
| AT5G25450.1 | 1,964059042 | 0,013016386 | no symbol available no full name available chr5:8857036-8857849 FORWARD LENGTH=122 |

| | | | |
|---|---|---|---|
| AT5G46290.3 | 1,319585824 | 0,01316185 | KASI_ KAS1 3-ketoacyl-acyl carrier protein synthase I_ KETOACYL-ACP SYNTHASE 1 chr5:18774439-18776629 REV |
| AT5G20630.1 | 0,889110881 | 0,013528728 | GER3_ GLP3A_ ATGER3_ GLP3_ GLP3B ARABIDOPSIS THALIANA GERMIN 3_ germin 3_ GERMIN-LIKE PROTI |
| AT2G38540.1 | 5,682159121 | 0,013593253 | ATLTP1_ AtLtpI-4_ LTP1_ LP1 ARABIDOPSIS THALIANA LIPID TRANSFER PROTEIN 1_ lipid transfer protein 1_ I |
| ATCG00270.1 | 0,721538778 | 0,013601414 | PSBD photosystem II reaction center protein D chrc:32711-33772 FORWARD LENGTH=353 |
| AT5G35630.1 | 0,917002605 | 0,013702286 | GLN2_ GS2_ ATGSL1 GLUTAMINE SYNTHETASE 2_ glutamine synthetase 2_ GLUTAMINE SYNTHETASE LIKE 1 |
| AT5G17990.1 | 1,296378928 | 0,013707547 | pat1_ TRP1 tryptophan biosynthesis 1_ PHOSPHORIBOSYLANTHRANILATE TRANSFERASE 1 chr5:5957330-595968 |
| AT2G27860.1 | 0,779055719 | 0,013736398 | AXS1 UDP-D-apiose/UDP-D-xylose synthase 1 chr2:11864684-11866843 REVERSE LENGTH=389 |
| AT3G24430.1 | 0,410637977 | 0,013749267 | HCF101 HIGH-CHLOROPHYLL-FLUORESCENCE 101 chr3:8868731-8872154 REVERSE LENGTH=532 |
| AT1G08360.1 | 0,672236678 | 0,013835727 | no symbol available no full name available chr1:2636231-2637694 FORWARD LENGTH=216 |
| AT1G22300.1 | 1,209611867 | 0,013924794 | 14-3-3EPSILON_ GRF10_ GF14 EPSILON general regulatory factor 10_ 14-3-3 PROTEIN G-BOX FACTOR14 EPSILON |
| AT3G26070.1 | 0,521906692 | 0,014212559 | FBN3a FIBRILLIN3a chr3:9526904-9528199 FORWARD LENGTH=242 |
| AT5G56350.1 | 1,620277165 | 0,014245825 | no symbol available no full name available chr5:22820254-22822529 REVERSE LENGTH=498 |
| AT3G61430.1 | 1,255047127 | 0,014407433 | ATPIP1_ PIP1A_ PIP1_ PIP1;1 plasma membrane intrinsic protein 1A_ PLASMA MEMBRANE INTRINSIC PROTEIN 1 |
| AT2G35490.1 | 1,179796585 | 0,014552769 | FBN2 FIBRILLIN2 chr2:14912309-14913797 REVERSE LENGTH=376 |
| ATCG01060.1 | 1,228862737 | 0,01456502 | PSAC chrc:117318-117563 REVERSE LENGTH=81 |
| AT3G15950.2 | 0,69997927 | 0,014618257 | NAI2 chr3:5397783-5402610 REVERSE LENGTH=734 |
| AT5G19510.1 | 0,765392784 | 0,014769956 | no symbol available no full name available chr5:6581854-6583137 REVERSE LENGTH=224 |
| AT1G62180.1 | 1,48024032 | 0,014949897 | APSR_ PRH43_ PRH_ ATAPR2_ APR2 ADENOSINE-5'-PHOSPHOSULFATE REDUCTASE_ 3'-PHOSPHOADENOSI |
| AT4G39520.1 | 1,40861318 | 0,015126019 | Drg1-1 chr4:18371329-18374000 REVERSE LENGTH=369 |
| AT3G08940.1 | 0,618026023 | 0,015131175 | LHCB4.2 light harvesting complex photosystem II chr3:2717717-2718400 FORWARD LENGTH=227 |
| AT2G14260.2 | 1,216812133 | 0,01525512 | PIP_ PAP1 proline iminopeptidase_ prolyl aminopeptidase 1 chr2:6041441-6043475 REVERSE LENGTH=329 |
| AT2G28000.1 | 1,213897096 | 0,015399421 | ARC2_ CH-CPN60A_ SLP_ CPN60A_ Cpn60alpha1_ CPNA1 SCHLEPPERLESS_ chaperonin-60alpha1_ CHLOROPLA |
| AT3G53430.1 | 1,18858266 | 0,015526064 | no symbol available no full name available chr3:19809895-19810395 REVERSE LENGTH=166 |
| AT3G49110.1 | 0,62275995 | 0,015553719 | ATPCA_ PRX33_ ATPRX33_ PRXCA peroxidase CA_ PEROXIDASE CA_ PEROXIDASE 33 chr3:18200713-18202891 |
| AT5G28840.1 | 1,175855527 | 0,015634078 | GME GDP-D-mannose 3 chr5:10862472-10864024 REVERSE LENGTH=377 |
| AT4G11600.1 | 1,327456371 | 0,015882577 | GPXL6_ PHGPX_ LSC803_ ATGPX6_ GPX6 glutathione peroxidase 6 chr4:7010021-7011330 REVERSE LENGTH=232 |
| AT3G26740.1 | 0,697007442 | 0,016160508 | CCL CCR-like chr3:9827868-9828461 FORWARD LENGTH=141 |
| AT5G15090.1 | 1,270741766 | 0,016169835 | VDAC3_ AtVDAC-3_ ATVDAC3 ARABIDOPSIS THALIANA VOLTAGE DEPENDENT ANION CHANNEL 3_ voltag |
| AT1G65260.1 | 3,203343033 | 0,016281232 | VIPP1_ PTAC4_ IM30 VESICLE-INDUCING PROTEIN IN PLASTIDS 1_ plastid transcriptionally active 4 chr1:242363 |
| AT5G50850.1 | 1,17561152 | 0,016424523 | MAB1 MACCI-BOU chr5:20689671-20692976 FORWARD LENGTH=363 |
| AT5G19220.1 | 0,885536027 | 0,016429378 | ADG2_ APL1 ADP glucose pyrophosphorylase large subunit 1_ ADP GLUCOSE PYROPHOSPHORYLASE 2 chr5:6463 |
| AT1G29150.1 | 1,295663922 | 0,016643813 | RPN6_ ATS9 non-ATPase subunit 9_ REGULATORY PARTICLE NON-ATPASE 6 chr1:10181240-10182499 FORWAR |
| AT5G64040.1 | 1,484520656 | 0,01665042 | PSAN chr5:25628724-25629409 REVERSE LENGTH=171 |
| AT3G44110.1 | 1,664171905 | 0,016988549 | ATJ3_ ATJ_ J3 DNAJ homologue 3 chr3:15869115-15871059 REVERSE LENGTH=420 |
| AT5G08280.1 | 0,893533518 | 0,017105094 | HEMC_ RUG1 RUGOSA 1_ hydroxymethylbilane synthase chr5:2663763-2665596 REVERSE LENGTH=382 |
| AT3G63460.2 | 1,320429324 | 0,017263515 | SEC31B chr3:23431009-23437241 REVERSE LENGTH=1102 |
| AT5G41520.1 | 1,504780883 | 0,017411038 | RPS10B ribosomal protein S10e B chr5:16609377-16610583 REVERSE LENGTH=180 |
| AT2G04842.1 | 0,617196259 | 0,017457356 | EMB2761 EMBRYO DEFECTIVE 2761 chr2:1698466-1701271 REVERSE LENGTH=650 |
| AT4G32260.1 | 1,138700274 | 0,017809948 | PDE334 PIGMENT DEFECTIVE 334 chr4:15573859-15574586 REVERSE LENGTH=219 |

| | | | |
|---|---|---|---|
| AT5G02160.1 | 0,4760938 | 0,018240421 | FIP FtsH5 Interacting Protein chr5:426392-427024 FORWARD LENGTH=129 |
| AT1G10630.1 | 1,191313924 | 0,018321888 | ARFA1F_ ATARFA1F ADP-ribosylation factor A1F chr1:3513189-3514230 REVERSE LENGTH=181 |
| AT1G23190.1 | 1,172408662 | 0,018692568 | PGM3 phosphoglucomutase 3 chr1:8219946-8224186 FORWARD LENGTH=583 |
| AT1G56500.2 | 0,742576592 | 0,018761852 | SOQ1 suppressor of quenching 1 chr1:21159775-21166313 FORWARD LENGTH=878 |
| AT1G17880.1 | 0,64332264 | 0,018829325 | ATBTF3_ BTF3 basic transcription factor 3 chr1:6152572-6153425 REVERSE LENGTH=165 |
| AT5G41670.1 | 0,78076919 | 0,01918538 | PGD3 6-phosphogluconate dehydrogenase 3 chr5:16665647-16667110 REVERSE LENGTH=487 |
| AT1G18210.1 | 0,604463784 | 0,019293121 | no symbol available no full name available chr1:6268273-6268785 REVERSE LENGTH=170 |
| AT4G01050.1 | 0,900358415 | 0,019308645 | TROL thylakoid rhodanese-like chr4:455874-458175 FORWARD LENGTH=466 |
| AT3G56190.1 | 1,334128299 | 0,019331801 | ALPHA-SNAP2_ ASNAP alpha-soluble NSF attachment protein 2 chr3:20846119-20848356 REVERSE LENGTH=289 |
| AT4G13940.1 | 1,274928956 | 0,019382395 | MEE58_ SAHH1_ EMB1395_ HOG1_ SAH1_ ATSAHH1 EMBRYO DEFECTIVE 1395_ HOMOLOGY-DEPENDENT [illegible] |
| AT2G27730.1 | 1,667832793 | 0,019551997 | no symbol available no full name available chr2:11820056-11820867 REVERSE LENGTH=113 |
| AT1G75330.1 | 0,61813436 | 0,019561129 | OTC ornithine carbamoyltransferase chr1:28266457-28268383 REVERSE LENGTH=375 |
| AT1G09430.1 | 1,218605609 | 0,019655518 | ACLA-3 ATP-citrate lyase A-3 chr1:3042135-3044978 FORWARD LENGTH=424 |
| ATCG00680.1 | 0,724833601 | 0,019689963 | PSBB photosystem II reaction center protein B chrc:72371-73897 FORWARD LENGTH=508 |
| AT2G21385.2 | 0,772079284 | 0,019813141 | AtCGLD11_ BFA3_ CGLD11 CONSERVED IN THE GREEN LINEAGE AND DIATOMS 11_ biogenesis factors require[illegible] |
| AT1G11750.1 | 1,243330572 | 0,01987073 | NCLPP1_ CLPP6_ NCLPP6 NUCLEAR-ENCODED CLPP 1_ CLP protease proteolytic subunit 6 chr1:3967609-3969535 [illegible] |
| AT3G14310.1 | 0,904789513 | 0,019894748 | OZS2_ ATPME3_ PME3 pectin methylesterase 3_ OVERLY ZINC SENSITIVE 2 chr3:4772214-4775095 REVERSE LEN[illegible] |
| AT1G11840.6 | 1,168083046 | 0,01996788 | AtGLYI3_ GLX1_ ATGLX1 glyoxalaseI 3_ glyoxalase I homolog chr1:3995928-3997518 FORWARD LENGTH=322 |
| AT2G05710.1 | 1,288123774 | 0,020143045 | ACO3 aconitase 3 chr2:2141591-2146350 FORWARD LENGTH=990 |
| AT1G20620.1 | 0,904902463 | 0,02029219 | SEN2_ ROG1_ ATCAT3_ CAT3 SENESCENCE 2_ catalase 3_ REPRESSOR OF GSNOR1 chr1:7143142-7146193 FOR[illegible] |
| AT5G03650.1 | 0,520506468 | 0,020738681 | SBE2.2 starch branching enzyme 2.2 chr5:931924-937470 FORWARD LENGTH=805 |
| AT5G17310.2 | 1,386041807 | 0,020892107 | AtUGP2_ UGP2 UDP-GLUCOSE PYROPHOSPHORYLASE 2_ UDP-glucose pyrophosphorylase 2 chr5:5696955-57008[illegible] |
| AT4G34620.1 | 0,827868394 | 0,020908883 | SSR16 small subunit ribosomal protein 16 chr4:16535084-16536092 REVERSE LENGTH=113 |
| AT2G37620.1 | 1,355811779 | 0,020964941 | ACT1_ AAc1 ARABIDOPSIS ACTIN 1_ actin 1 chr2:15779761-15781241 FORWARD LENGTH=377 |
| AT1G09620.1 | 1,159228721 | 0,021274643 | no symbol available no full name available chr1:3113077-3116455 REVERSE LENGTH=1091 |
| AT5G40760.1 | 1,284326536 | 0,021417396 | G6PD6 glucose-6-phosphate dehydrogenase 6 chr5:16311284-16314556 FORWARD LENGTH=515 |
| AT2G28815.1 | 1,483158758 | 0,021452057 | no symbol available no full name available chr2:12367001-12368064 REVERSE LENGTH=291 |
| AT4G21990.1 | 1,605320168 | 0,021516097 | APR3_ PRH26_ PRH-26_ ATAPR3 PAPS REDUCTASE HOMOLOG 26_ APS reductase 3 chr4:11657284-11658973 RE[illegible] |
| AT2G21410.1 | 1,463000482 | 0,021593703 | VHA-A2 vacuolar proton ATPase A2 chr2:9162703-9168141 FORWARD LENGTH=821 |
| AT5G37600.1 | 1,37107827 | 0,022498552 | ATGSR1_ ATGLN1;1_ GLN1;1_ GSR 1 ARABIDOPSIS THALIANA GLUTAMINE SYNTHASE CLONE R1_ ARABI[illegible] |
| AT1G45201.1 | 0,801758766 | 0,022655917 | TLL1_ ATTLL1 ARABIDOPSIS THALIANA TRIACYLGLYCEROL LIPASE-LIKE 1_ triacylglycerol lipase-like 1 chr1[illegible] |
| AT3G63410.1 | 0,82435061 | 0,022760433 | E37_ VTE3_ IEP37_ APG1 INNER ENVELOPE PROTEIN 37_ VITAMIN E DEFECTIVE 3_ ALBINO OR PALE GRE[illegible] |
| AT5G03300.1 | 1,154273951 | 0,023202687 | ADK2 adenosine kinase 2 chr5:796573-798997 FORWARD LENGTH=345 |
| AT5G63980.1 | 0,726246174 | 0,023405162 | ALX8_ SUPO1_ AtFRY1_ HOS2_ ATSAL1_ SAL1_ RON1_ FRY1 HIGH EXPRESSION OF OSMOTICALLY RESPO[illegible] |
| AT4G38680.1 | 1,94250537 | 0,02343804 | GRP2_ CSP2_ ATCSP2_ CSDP2 glycine rich protein 2_ COLD SHOCK DOMAIN PROTEIN 2_ ARABIDOPSIS THALI[illegible] |
| AT2G37190.1 | 0,868867971 | 0,023589832 | no symbol available no full name available chr2:15619559-15620059 REVERSE LENGTH=166 |
| AT4G01850.1 | 1,215401702 | 0,023696189 | AtSAM2_ SAM-2_ MAT2_ SAM2 S-adenosylmethionine synthetase 2_ S-ADENOSYLMETHIONINE SYNTHETASE 2 [illegible] |
| AT1G52510.1 | 0,875618532 | 0,023756883 | no symbol available no full name available chr1:19563039-19565260 REVERSE LENGTH=380 |
| AT1G65960.2 | 1,359470468 | 0,023759428 | GAD2 glutamate decarboxylase 2 chr1:24552094-24557253 FORWARD LENGTH=494 |

| | | | |
|---|---|---|---|
| AT2G42220.1 | 0,669177456 | 0,023764477 | no symbol available no full name available chr2:17592105-17593305 FORWARD LENGTH=234 |
| AT3G06580.1 | 1,332588864 | 0,023909202 | GAL1_ GALK GALACTOSE KINASE 1 chr3:2049141-2051867 REVERSE LENGTH=496 |
| AT1G55260.2 | 1,633123887 | 0,024165452 | LTPG6 glycosylphosphatidylinositol-anchored lipid protein transfer 6 chr1:20614663-20616158 FORWARD LENGTH=22 |
| AT5G44320.1 | 1,135230965 | 0,024368441 | no symbol available no full name available chr5:17854901-17856667 REVERSE LENGTH=588 |
| AT5G66570.1 | 0,889283273 | 0,024381952 | OEE1_ PSBO-1_ MSP-1_ OE33_ OEE33_ PSBO1 PS II OXYGEN-EVOLVING COMPLEX 1_ OXYGEN EVOLVING [illegible] |
| AT4G37930.1 | 0,940194827 | 0,024577559 | STM_ SHMT1_ SHM1 SERINE HYDROXYMETHYLTRANSFERASE 1_ serine transhydroxymethyltransferase 1_ SERI |
| AT1G77090.1 | 0,804705863 | 0,024801397 | no symbol available no full name available chr1:28960576-28961875 REVERSE LENGTH=260 |
| AT5G60640.2 | 0,78299236 | 0,024991137 | PDI2_ ATPDIL1-4_ ATPDI2_ PDIL1-4 PROTEIN DISULFIDE ISOMERASE 2_ ARABIDOPSIS THALIANA PROTEI |
| AT3G06860.1 | 1,307576156 | 0,024994703 | ATMFP2_ MFP2 MULTIFUNCTIONAL PROTEIN 2_ multifunctional protein 2 chr3:2161926-2166009 FORWARD LE |
| AT4G35460.1 | 1,368882124 | 0,025201204 | ATNTRB_ NTR1_ NTRB NADPH-DEPENDENT THIOREDOXIN REDUCTASE 1_ NADPH-dependent thioredoxin red |
| AT1G23820.1 | 1,203551161 | 0,025382321 | SPDS1 spermidine synthase 1 chr1:8420410-8422724 FORWARD LENGTH=334 |
| AT2G17265.1 | 0,529329088 | 0,02565147 | DRM1_ HSK_ DMR1 DOWNY MILDEW RESISTANT 1_ homoserine kinase chr2:7508606-7509718 FORWARD LEN |
| AT4G20850.1 | 0,850068947 | 0,025792922 | TPP2 tripeptidyl peptidase ii chr4:11160935-11169889 REVERSE LENGTH=1380 |
| AT2G36530.1 | 1,131538628 | 0,025863363 | LOS2_ ENO2 LOW EXPRESSION OF OSMOTICALLY RESPONSIVE GENES 2_ enolase 2 chr2:15321081-15323786 |
| AT1G52400.1 | 0,78146464 | 0,026335294 | BGL1_ ATBG1_ BGLU18 A. THALIANA BETA-GLUCOSIDASE 1_ BETA-GLUCOSIDASE HOMOLOG 1_ beta gluc |
| AT2G43100.1 | 0,58338201 | 0,02655025 | IPMI2_ IPMI SSU2_ ATLEUD1 isopropylmalate isomerase 2_ isopropylmalate isomerase small sub-unit 2 chr2:17920685- |
| AT1G62780.1 | 0,657748178 | 0,026650932 | no symbol available no full name available chr1:23249349-23251066 REVERSE LENGTH=237 |
| AT2G39990.1 | 1,112158583 | 0,026732066 | EIF2_ AteIF3f_ eIF3F Arabidopsis thaliana eukaryotic translation initiation factor 3 subunit F_ eukaryotic translation initia |
| AT3G17240.1 | 0,752499464 | 0,026922172 | mtLPD2 lipoamide dehydrogenase 2 chr3:5890278-5892166 REVERSE LENGTH=507 |
| AT4G26300.4 | 1,504246052 | 0,027342793 | emb1027 embryo defective 1027 chr4:13308400-13312204 REVERSE LENGTH=590 |
| AT3G04600.1 | 1,356735341 | 0,027391066 | no symbol available no full name available chr3:1243152-1245958 FORWARD LENGTH=402 |
| AT5G47700.1 | 1,353261152 | 0,027710333 | RPP1C_ RPP1.3 60S acidic ribosomal protein P1-3_ RPP1 co-orthologous gene 3 chr5:19328019-19328724 REVERSE LE |
| AT5G14040.1 | 1,315758017 | 0,027977222 | MPT3_ PHT3;1 phosphate transporter 3;1_ mitochondrial phosphate transporter 3 chr5:4531059-4532965 REVERSE LEN |
| AT3G24170.1 | 1,412386846 | 0,028188696 | ATGR1_ GR1 glutathione-disulfide reductase chr3:8729762-8734115 REVERSE LENGTH=499 |
| AT3G02520.1 | 1,25758037 | 0,028243254 | GRF7_ GF14 NU general regulatory factor 7 chr3:526800-527915 REVERSE LENGTH=265 |
| ATCG00750.1 | 0,875833991 | 0,028282174 | RPS11 ribosomal protein S11 chrc:78960-79376 REVERSE LENGTH=138 |
| AT1G78300.1 | 1,216465208 | 0,028396826 | GRF2_ 14-3-3OMEGA_ GF14 OMEGA general regulatory factor 2_ 14-3-3 PROTEIN G-BOX FACTOR14 OMEGA chr1 |
| AT1G67700.1 | 0,874966189 | 0,028472727 | HHL1 HYPERSENSITIVE TO HIGH LIGHT 1 chr1:25374295-25375716 FORWARD LENGTH=230 |
| AT5G59290.1 | 1,246629041 | 0,028484294 | ATUXS3_ UXS3 UDP-glucuronic acid decarboxylase 3 chr5:23915814-23917953 REVERSE LENGTH=342 |
| AT4G08870.1 | 0,801420119 | 0,028561053 | ARGAH2 arginine amidohydrolase 2 chr4:5646654-5648693 REVERSE LENGTH=344 |
| AT3G05590.1 | 0,614255286 | 0,028654371 | RPL18 ribosomal protein L18 chr3:1621511-1622775 FORWARD LENGTH=187 |
| AT4G02450.2 | 1,160771021 | 0,029444029 | p23-1 chr4:1073987-1075765 REVERSE LENGTH=240 |
| AT3G03960.1 | 1,268624278 | 0,029935885 | CCT8 Chaperonin containing T-complex polypeptide-1 subunit 8 chr3:1024432-1027604 FORWARD LENGTH=549 |
| AT4G22670.1 | 1,270590108 | 0,030005268 | AtHip1_ HIP1_ TPR11 HSP70-interacting protein 1_ tetratricopeptide repeat 11 chr4:11918236-11920671 FORWARD LE |
| AT5G11450.1 | 0,455109581 | 0,030414983 | PPD5 PsbP domain protein 5 chr5:3654475-3656357 FORWARD LENGTH=297 |
| AT5G23010.1 | 0,576145479 | 0,030635318 | GSM1_ MAM1_ IMS3 glucosinolate metabolism 1_ 2-ISOPROPYLMALATE SYNTHASE 3_ methylthioalkylmalate synt |
| AT4G21280.1 | 1,128734055 | 0,030806107 | PSBQ-1_ PSBQA_ PSBQ photosystem II subunit QA_ PHOTOSYSTEM II SUBUNIT Q-1_ PHOTOSYSTEM II SUBUN |
| AT1G73230.1 | 1,485861214 | 0,031245385 | no symbol available no full name available chr1:27540506-27541364 REVERSE LENGTH=165 |
| AT4G25630.1 | 1,16496054 | 0,031332256 | ATFIB2_ FIB2 fibrillarin 2 chr4:13074239-13076205 FORWARD LENGTH=320 |

| | | | |
|---|---|---|---|
| AT5G10360.1 | 0,548409378 | 0,031363838 | RPS6B_ EMB3010 Ribosomal protein small subunit 6b_ embryo defective 3010 chr5:3258734-3260142 REVERSE LENG |
| AT5G03630.1 | 1,402510439 | 0,031464095 | MDAR2 chr5:922378-924616 REVERSE LENGTH=435 |
| AT3G26450.1 | 1,229713964 | 0,032158243 | no symbol available no full name available chr3:9681593-9683299 REVERSE LENGTH=152 |
| AT3G53990.2 | 1,21139711 | 0,032323742 | AtUSP_ USP17 Universal stress protein chr3:19990558-19991019 REVERSE LENGTH=126 |
| AT1G79230.3 | 1,124337364 | 0,032342134 | ATMST1_ STR1_ MST1_ ATRDH1_ ST1 ARABIDOPSIS THALIANA RHODANESE HOMOLOGUE 1_ mercaptopyru |
| AT5G44500.1 | 1,338354695 | 0,032378258 | no symbol available no full name available chr5:17927505-17928269 FORWARD LENGTH=254 |
| AT5G39740.1 | 0,895853875 | 0,032405306 | RPL5B_ OLI7 ribosomal protein L5 B_ OLIGOCELLULA 7 chr5:15903365-15905185 FORWARD LENGTH=301 |
| AT1G79870.2 | 1,13972299 | 0,032493888 | HPPR2 Hydroxyphenylpyruvate reductase 2 chr1:30044794-30045851 FORWARD LENGTH=294 |
| AT4G25740.1 | 1,297403115 | 0,032608161 | no symbol available no full name available chr4:13107488-13108751 REVERSE LENGTH=177 |
| AT1G49760.1 | 0,863316512 | 0,032671063 | PABP8_ PAB8 POLY(A) BINDING PROTEIN 8_ poly(A) binding protein 8 chr1:18416740-18419753 FORWARD LEN |
| AT4G39990.1 | 1,911978585 | 0,03281555 | ATGB3_ RABA4B_ ATRABA4B_ ATRAB11G GTP-BINDING PROTEIN 3_ RAB GTPase homolog A4B_ ARABIDOP |
| AT5G04140.1 | 0,949156562 | 0,033879591 | GLUS_ FD-GOGAT_ GLS1_ GLU1 FERREDOXIN-DEPENDENT GLUTAMATE SYNTHASE 1_ glutamate synthase 1_ |
| AT5G38480.1 | 1,180652137 | 0,034165253 | GRF3_ RCI1 general regulatory factor 3 chr5:15410277-15411285 FORWARD LENGTH=255 |
| AT5G16510.1 | 1,411815756 | 0,034227863 | RGP5 reversibly glycosylated polypeptide 5_ reversibly glycosylated protein 5 chr5:5393296-5394342 FORWARD LENG |
| ATCG00020.1 | 0,735718445 | 0,034542136 | PSBA photosystem II reaction center protein A chrc:383-1444 REVERSE LENGTH=353 |
| AT2G43090.1 | 0,471661938 | 0,034636996 | IPMI SSU1 isopropylmalate isomerase small subunit 1 chr2:17918957-17919712 FORWARD LENGTH=251 |
| AT5G12040.1 | 1,591049925 | 0,034669969 | no symbol available no full name available chr5:3885162-3887772 FORWARD LENGTH=369 |
| AT1G60710.1 | 0,776774534 | 0,034723532 | ATB2 chr1:22355073-22356627 REVERSE LENGTH=345 |
| AT4G08900.1 | 0,72718168 | 0,034826438 | ARGAH1 arginine amidohydrolase 1 chr4:5703499-5705180 FORWARD LENGTH=342 |
| AT3G12050.2 | 2,589304782 | 0,034870461 | no symbol available no full name available chr3:3839289-3841303 FORWARD LENGTH=321 |
| AT5G06320.1 | 1,200059922 | 0,034888856 | NHL3 NDR1/HIN1-like 3 chr5:1931016-1931711 REVERSE LENGTH=231 |
| AT5G28510.1 | 0,838476024 | 0,035691378 | BGLU24 beta glucosidase 24 chr5:10481041-10484022 REVERSE LENGTH=533 |
| AT1G74970.1 | 1,197904071 | 0,035806297 | TWN3_ SOT8_ RPS9_ PRPS9 ribosomal protein S9 chr1:28157761-28159202 REVERSE LENGTH=208 |
| AT1G09780.1 | 0,70195479 | 0,035828398 | iPGAM1 "2_3-biphosphoglycerate-independent phosphoglycerate mutase 1" chr1:3165550-3167812 REVERSE LENGTH= |
| AT5G24300.1 | 0,819564814 | 0,035958439 | SS1_ ATSS1 STARCH SYNTHASE 1_ starch synthase 1 chr5:8266934-8270860 FORWARD LENGTH=652 |
| AT3G14600.1 | 0,293382563 | 0,036406664 | no symbol available no full name available chr3:4910773-4911933 FORWARD LENGTH=178 |
| AT5G20720.1 | 1,097452475 | 0,036529024 | ATCPN21_ CPN20_ CPN21_ CHCPN10_ CPN10 chaperonin 20_ CHLOROPLAST CHAPERONIN 10 chr5:7015015-70 |
| AT2G39330.1 | 0,888607212 | 0,036565969 | JAL23 jacalin-related lectin 23 chr2:16419787-16421573 REVERSE LENGTH=459 |
| AT5G54810.1 | 0,894824641 | 0,036683484 | TRP2_ ATTSB1_ TSB1_ TRPB tryptophan synthase beta-subunit 1_ TRYPTOPHAN BIOSYNTHESIS B_ TRYPTOPHA |
| AT2G41530.1 | 1,119530947 | 0,03671055 | SFGH_ ATSFGH S-formylglutathione hydrolase_ ARABIDOPSIS THALIANA S-FORMYLGLUTATHIONE HYDROLA |
| AT3G02230.1 | 0,865687144 | 0,036857527 | RGP1_ ATRGP1 ARABIDOPSIS THALIANA REVERSIBLY GLYCOSYLATED POLYPEPTIDE 1_ reversibly glycosyl |
| AT2G31610.1 | 1,065948392 | 0,036895444 | no symbol available no full name available chr2:13450384-13451669 FORWARD LENGTH=250 |
| AT3G60820.1 | 1,187032356 | 0,03698016 | PBF1 chr3:22472038-22473809 REVERSE LENGTH=223 |
| AT1G78830.1 | 0,814367227 | 0,037008529 | MNB1 chr1:29637141-29638508 REVERSE LENGTH=455 |
| AT5G59370.1 | 1,237533546 | 0,037087998 | ACT4 actin 4 chr5:23950109-23951586 FORWARD LENGTH=377 |
| AT2G20580.1 | 1,255557534 | 0,037331345 | ATRPN1A_ RPN1A 26S PROTEASOME REGULATORY SUBUNIT S2 1A_ 26S proteasome regulatory subunit S2 1A c |
| AT1G52000.1 | 1,335081862 | 0,037593753 | no symbol available no full name available chr1:19333352-19335700 REVERSE LENGTH=730 |
| AT1G28290.2 | 1,191083468 | 0,037859913 | AGP31 arabinogalactan protein 31 chr1:9889331-9890843 REVERSE LENGTH=315 |
| AT3G54890.4 | 0,812230366 | 0,038389017 | LHCA1 photosystem I light harvesting complex gene 1 chr3:20339881-20340922 REVERSE LENGTH=213 |

| | | | |
|---|---|---|---|
| AT3G46830.1 | 1,806204362 | 0,038714299 | ATRAB-A2C_ ATRAB11A_ ATRABA2C_ RABA2c_ RAB-A2C RAB GTPASE HOMOLOG A2C_ ARABIDOPSIS RA |
| AT2G22290.1 | 1,750618194 | 0,03887787 | RAB-H1D_ ATRABH1D_ ATRAB6_ RABH1d_ ATRAB-H1D RAB GTPASE HOMOLOG H1D_ ARABIDOPSIS RAB |
| AT1G66270.2 | 1,693787519 | 0,03998944 | BGLU21 chr1:24700110-24702995 REVERSE LENGTH=522 |
| AT3G15020.1 | 0,895735774 | 0,0401229 | mMDH2 mitochondrial malate dehydrogenase 2 chr3:5056139-5057941 FORWARD LENGTH=341 |
| AT5G01530.1 | 0,853332873 | 0,040369297 | LHCB4.1 light harvesting complex photosystem II chr5:209084-210243 FORWARD LENGTH=290 |
| AT1G10370.1 | 2,450040301 | 0,040467946 | ATGSTU17_ ERD9_ GST30B_ GST30_ GSTU17 GLUTATHIONE S-TRANSFERASE 30B_ GLUTATHIONE S-TRAN |
| AT3G01910.1 | 1,143451536 | 0,040882018 | AtSO_ AT-SO_ SOX sulfite oxidase chr3:314919-317274 REVERSE LENGTH=393 |
| AT3G52300.1 | 1,45673307 | 0,040884211 | ATPQ_ ATPd "ATP synthase D chain_ mitochondrial" chr3:19396689-19398119 FORWARD LENGTH=168 |
| AT1G03475.1 | 0,846369208 | 0,041239638 | HEMF1_ ATCPO-I_ LIN2 LESION INITIATION 2 chr1:869302-871175 REVERSE LENGTH=386 |
| AT1G21720.1 | 1,296984602 | 0,041568106 | PBC1 proteasome beta subunit C1 chr1:7626394-7628070 FORWARD LENGTH=204 |
| AT5G65010.1 | 1,349326913 | 0,041811136 | ASN2 asparagine synthetase 2 chr5:25969224-25972278 FORWARD LENGTH=578 |
| AT4G15210.3 | 1,363282621 | 0,042212535 | AT-BETA-AMY_ ATBETA-AMY_ BMY1_ RAM1_ BAM5 REDUCED BETA AMYLASE 1_ beta-amylase 5_ ARABID |
| AT1G09750.1 | 0,818210206 | 0,04227054 | no symbol available no full name available chr1:3157541-3158960 FORWARD LENGTH=449 |
| AT3G49720.1 | 0,586912268 | 0,042482924 | CGR2 chr3:18440192-18441655 REVERSE LENGTH=261 |
| AT4G24930.1 | 0,882603552 | 0,042594608 | no symbol available no full name available chr4:12821496-12822389 REVERSE LENGTH=225 |
| AT4G22930.1 | 1,572744749 | 0,042939194 | DHOASE_ PYR4 DIHYDROOROTASE_ pyrimidin 4 chr4:12019315-12021200 FORWARD LENGTH=377 |
| AT4G23670.1 | 0,815300892 | 0,042987732 | no symbol available no full name available chr4:12332846-12333656 REVERSE LENGTH=151 |
| AT4G22010.1 | 1,231785815 | 0,042988579 | sks4 SKU5 similar 4 chr4:11663429-11666463 FORWARD LENGTH=541 |
| AT4G21650.1 | 2,057002128 | 0,043007658 | SBT3.13 subtilase 3.13 chr4:11501314-11504656 REVERSE LENGTH=766 |
| AT3G54900.1 | 0,590686004 | 0,043113772 | AtGRXS14_ ATGRXCP_ CXIP1 GLUTAREDOXIN_ CAX interacting protein 1 chr3:20341850-20342371 REVERSE LE |
| AT4G33680.1 | 1,110487181 | 0,043145137 | AGD2 ABERRANT GROWTH AND DEATH 2_ ARF-GAP domain 2 chr4:16171847-16174630 REVERSE LENGTH=4[illegible] |
| AT3G22630.1 | 2,883198211 | 0,043146153 | PBD1_ PRCGB 20S proteasome beta subunit D1 chr3:8009709-8010774 REVERSE LENGTH=204 |
| AT3G53990.1 | 1,187400851 | 0,043675697 | AtUSP_ USP17 Universal stress protein chr3:19989658-19991019 REVERSE LENGTH=160 |
| AT2G32120.1 | 1,933746583 | 0,043947251 | HSP70T-2 heat-shock protein 70T-2 chr2:13651720-13653411 REVERSE LENGTH=563 |
| AT1G74910.1 | 1,344620487 | 0,044048981 | KJC1 KONJAC 1 chr1:28135770-28138456 REVERSE LENGTH=415 |
| AT5G19990.1 | 1,162273943 | 0,044064639 | RPT6A_ ATSUG1 regulatory particle triple-A ATPase 6A chr5:6752144-6754918 FORWARD LENGTH=419 |
| AT2G21590.1 | 0,570039119 | 0,044367317 | APL4 chr2:9239362-9242150 FORWARD LENGTH=523 |
| AT4G09000.1 | 1,127015579 | 0,045112728 | GRF1_ GF14 CHI GENERAL REGULATORY FACTOR1-G-BOX FACTOR 14-3-3 HOMOLOG ISOFORM CHI_ gener[illegible] |
| ATCG01110.1 | 0,678197935 | 0,045310336 | NDHH NAD(P)H dehydrogenase subunit H chrc:122011-123192 REVERSE LENGTH=393 |
| AT1G01090.1 | 0,921897069 | 0,045392582 | PDH-E1 ALPHA pyruvate dehydrogenase E1 alpha chr1:47705-49166 REVERSE LENGTH=428 |
| AT1G30120.1 | 0,515918374 | 0,045457324 | PDH-E1 BETA pyruvate dehydrogenase E1 beta chr1:10584350-10586477 REVERSE LENGTH=406 |
| AT4G38740.1 | 1,149284143 | 0,045680314 | ROC1 rotamase CYP 1 chr4:18083620-18084138 REVERSE LENGTH=172 |
| AT1G54780.1 | 0,793029582 | 0,04624225 | TLP18.3_ AtTLP18.3 thylakoid lumen protein 18.3 chr1:20439533-20440953 FORWARD LENGTH=285 |
| AT1G48420.1 | 1,532389899 | 0,04646698 | ACD1_ DCD_ ATACD1_ D-CDES_ AtDCD 1-AMINOCYCLOPROPANE-1-CARBOXYLIC ACID DEAMINASE 1_ A[illegible] |
| AT5G66120.2 | 0,670732405 | 0,046525887 | no symbol available no full name available chr5:26431516-26433649 REVERSE LENGTH=442 |
| AT1G66240.1 | 1,399435676 | 0,046726447 | AtHMP14_ ATX1_ ATATX1 HEAVY METAL ASSOCIATED PROTEIN 14_ homolog of anti-oxidant 1 chr1:24686445-[illegible] |
| AT3G14790.1 | 0,801969416 | 0,046893464 | RHM3_ ATRHM3 ARABIDOPSIS THALIANA RHAMNOSE BIOSYNTHESIS 3_ rhamnose biosynthesis 3 chr3:496479 |
| AT5G35590.1 | 0,725934671 | 0,046957781 | PAA1 proteasome alpha subunit A1 chr5:13765417-13767768 REVERSE LENGTH=246 |
| AT3G13580.1 | 1,77375979 | 0,047118707 | no symbol available no full name available chr3:4433809-4435109 FORWARD LENGTH=244 |

| | | | |
|---|---|---|---|
| AT5G47930.1 | 0,387832643 | 0,047208277 | no symbol available no full name available chr5:19406423-19407329 REVERSE LENGTH=84 |
| AT5G11880.1 | 0,791422709 | 0,047377662 | DAPDC2 meso-diaminopimelate decarboxylase 2 chr5:3827806-3829942 REVERSE LENGTH=489 |
| AT2G20260.1 | 1,109214428 | 0,048143397 | PSAE-2 photosystem I subunit E-2 chr2:8736780-8737644 FORWARD LENGTH=145 |
| AT5G66680.1 | 1,184202375 | 0,048441495 | DGL1 DEFECTIVE GLYCOSYLATION chr5:26617840-26620581 REVERSE LENGTH=437 |
| AT1G53750.1 | 1,340866078 | 0,048589455 | RPT1A regulatory particle triple-A 1A chr1:20065921-20068324 REVERSE LENGTH=426 |
| AT5G63890.1 | 1,070525955 | 0,048758712 | HISN8_ ATHDH_ HDH histidinol dehydrogenase_ HISTIDINE BIOSYNTHESIS 8 chr5:25565600-25567879 REVERSE |
| AT5G59870.1 | 1,652906217 | 0,048760262 | h2a.w.6_ HTA6 histone H2A 6 chr5:24115605-24116144 REVERSE LENGTH=150 |
| AT5G03690.1 | 1,2723894 | 0,048839468 | AtFBA4_ FBA4 fructose-bisphosphate aldolase 4 chr5:963389-964982 REVERSE LENGTH=393 |
| AT5G16990.1 | 1,281936137 | 0,049356642 | no symbol available no full name available chr5:5581831-5583849 REVERSE LENGTH=343 |
| AT3G10090.1 | 4,017716899 | 0,049918085 | no symbol available no full name available chr3:3108960-3109154 REVERSE LENGTH=64 |
| AT1G75280.1 | 1,793250542 | 0,049964426 | no symbol available no full name available chr1:28252030-28253355 FORWARD LENGTH=310 |

PROTEIN 1_ GLYCINE RICH PROTEIN 7_ "cold_ circadian rhythm_ and rna binding 2"_ GLYCINE-RICH RNA-BINDING PROTEIN 7 chr2:9265477-9266316 REVERS

SIS THALIANA PROTEIN DISULFIDE ISOMERASE 11_ UNFERTILIZED EMBRYO SAC 5_ MATERNAL EFFECT EMBRYO ARREST 30 chr2:19481503-19483683

ABIDOPSIS THALIANA CINNAMYL-ALCOHOL DEHYDROGENASE 8_ ELICITOR-ACTIVATED GENE 3 chr4:17855964-17857388 FORWARD LENGTH=359

)PSIS GLUTATHIONE S-TRANSFERASE 1_ ARABIDOPSIS THALIANA GLUATIONE S-TRANSFERASE F3_ glutathione S-transferase 6_ GLUTATHIONE S-TRANS

_ ARABIDOPSIS THALIANA PLASMA MEMBRANE INTRINSIC PROTEIN 1_ PLASMA MEMBRANE INTRINSIC PROTEIN 1;1 chr3:22733657-22735113 FORWA

NE-5'-PHOSPHOSULFATE (PAPS) REDUCTASE HOMOLOG 43_ 5'adenylylphosphosulfate reductase 2 chr1:22975794-22977465 REVERSE LENGTH=454

ST CHAPERONIN 60ALPHA_ ACCUMULATION AND REPLICATION OF CHLOROPLASTS 2_ chaperonin-60alpha chr2:11926603-11929184 FORWARD LENGTH=

GENE SILENCING 1_ MATERNAL EFFECT EMBRYO ARREST 58_ S-ADENOSYL-L-HOMOCYSTEIN HYDROLASE 1 chr4:8054931-8056676 FORWARD LENGT[illegible]

[illegible]NSIVE GENES 2_ suppressors of PIN1 overexpression 1_ ALTERED EXPRESSION OF APX2 8_ FIERY1_ ROTUNDA 1 chr5:25610002-25611802 FORWARD LENGTH

COMPLEX 33 KILODALTON PROTEIN_ OXYGEN EVOLVING ENHANCER PROTEIN 33_ PS II oxygen-evolving complex 1_ MANGANESE-STABILIZING PROTE

SFERASE 30_ GLUTATHIONE S-TRANSFERASE TAU 17_ GLUTATHIONE S-TRANSFERASE U17_ EARLY-RESPONSIVE TO DEHYDRATION 9 chr1:3397274-3

THALIANA 1-AMINOCYCLOPROPANE-1-CARBOXYLIC ACID DEAMINASE 1_ D-cysteine desulfhydrase chr1:17896767-17898803 REVERSE LENGTH=401

IN 1_ 33 KDA OXYGEN EVOLVING POLYPEPTIDE 1 chr5:26568744-26570124 FORWARD LENGTH=332
