## Supplementary material for "HY5 enhances *Arabidopsis* tolerance to combined high light and heat stress by coordinating photoprotection and hormone signaling": Table S8

**Supplemental Table S8. Differentially accumulated proteins compared to control (P < 0.05) in HY5OX leaves subjected to high light stress.**

| Protein ID | Fold Change | p-value | Protein description |
|---|---|---|---|
| AT2G21250.1 | 3,787448079 | 4,41887E-08 | no symbol available no full name available chr2:9103408-9105116 REVERSE LENGTH=309 |
| AT5G14740.2 | 0,172102015 | 7,57449E-08 | BETA CA2_ CA2_ DEG12_ CA18 BETA CARBONIC ANHYDRASE 2_ CARBONIC ANHYDRASE 18_ carbo |
| AT2G21580.2 | 4,914810141 | 8,36477E-08 | no symbol available no full name available chr2:9236629-9237510 FORWARD LENGTH=107 |
| AT3G05590.1 | 0,167371568 | 9,5009E-08 | RPL18 ribosomal protein L18 chr3:1621511-1622775 FORWARD LENGTH=187 |
| AT4G39200.2 | 5,388387706 | 2,81881E-07 | no symbol available no full name available chr4:18257464-18258464 FORWARD LENGTH=107 |
| AT3G09630.1 | 0,390739304 | 4,51739E-07 | SAC56 Suppressor of Acaulis 56 chr3:2953813-2955444 FORWARD LENGTH=406 |
| AT1G47250.1 | 0,521015851 | 5,69277E-07 | PAF2 20S proteasome alpha subunit F2 chr1:17319220-17320900 FORWARD LENGTH=277 |
| AT2G38540.1 | 29,87722379 | 5,75662E-07 | ATLTP1_ AtLtpI-4_ LTP1_ LP1 ARABIDOPSIS THALIANA LIPID TRANSFER PROTEIN 1_ lipid transfer pro |
| AT4G36130.1 | 0,056757472 | 8,1046E-07 | no symbol available no full name available chr4:17097613-17098656 FORWARD LENGTH=258 |
| AT2G18020.1 | 0,135318499 | 1,38273E-06 | EMB2296 embryo defective 2296 chr2:7837151-7838160 FORWARD LENGTH=258 |
| AT1G73600.1 | 7,323567151 | 1,50673E-06 | DEG26_ NMT_ AtPMT3_ NMT3 Phosphoethanolamine methyltransferase3 chr1:27670825-27673400 FORWARD |
| AT4G18100.1 | 0,124053082 | 2,03421E-06 | no symbol available no full name available chr4:10035715-10036475 REVERSE LENGTH=133 |
| AT5G20290.1 | 0,146508912 | 2,54037E-06 | no symbol available no full name available chr5:6851695-6853012 REVERSE LENGTH=222 |
| AT5G19140.2 | 11,71390338 | 3,13717E-06 | ATAILP1_ AILP1 chr5:6423398-6425785 FORWARD LENGTH=222 |
| AT1G11840.1 | 2,284937661 | 3,72678E-06 | AtGLYI3_ GLX1_ ATGLX1 glyoxalaseI 3_ glyoxalase I homolog chr1:3996045-3997518 FORWARD LENGTH= |
| AT1G27950.1 | 5,838894288 | 5,05201E-06 | LTPG1 glycosylphosphatidylinositol-anchored lipid protein transfer 1 chr1:9740740-9741991 FORWARD LENGT |
| AT5G40950.1 | 0,712135568 | 5,54608E-06 | PRPL27_ RPL27 ribosomal protein large subunit 27 chr5:16410866-16411845 FORWARD LENGTH=198 |
| AT2G21330.1 | 1,325624229 | 6,58568E-06 | FBA1_ AtFBA1 fructose-bisphosphate aldolase 1 chr2:9128416-9130152 REVERSE LENGTH=399 |
| AT3G10950.1 | 0,44500454 | 6,64119E-06 | no symbol available no full name available chr3:3423893-3424566 FORWARD LENGTH=92 |
| AT3G11630.1 | 1,187945155 | 8,18618E-06 | 2CPA 2-Cys peroxiredoxin A chr3:3672189-3673937 FORWARD LENGTH=266 |
| AT1G18080.1 | 1,56693433 | 9,88984E-06 | RACK1A_AT_ AtRACK1_ SAC53_ ATARCA_ RACK1A RECEPTOR FOR ACTIVATED C KINASE 1 A_ Sup |
| AT3G13470.1 | 1,787072652 | 1,01474E-05 | CPNB2_ Cpn60beta2 chaperonin-60beta2 chr3:4389685-4392624 FORWARD LENGTH=596 |
| AT5G64040.1 | 2,976112217 | 1,02811E-05 | PSAN chr5:25628724-25629409 REVERSE LENGTH=171 |
| AT5G02870.1 | 0,543790923 | 1,05484E-05 | RPL4 ribosomal large subunit 4 chr5:657830-659526 FORWARD LENGTH=407 |
| AT5G02960.1 | 0,328194172 | 1,06088E-05 | no symbol available no full name available chr5:693280-694396 REVERSE LENGTH=142 |
| AT3G49010.4 | 0,067769049 | 1,31387E-05 | BBC1_ RSU2_ ATBBC1 40S RIBOSOMAL PROTEIN_ breast basic conserved 1 chr3:18166971-18168047 REVI |
| AT2G14880.1 | 7,027660261 | 1,38969E-05 | SWIB2 chr2:6393686-6394841 REVERSE LENGTH=141 |
| AT1G74260.1 | 1,650725898 | 1,62186E-05 | PUR4 purine biosynthesis 4 chr1:27923005-27927764 REVERSE LENGTH=1407 |
| AT5G40770.1 | 1,71437848 | 1,65138E-05 | ATPHB3_ PHB3_ EER3 prohibitin 3 chr5:16315589-16316621 REVERSE LENGTH=277 |
| AT1G79550.1 | 1,848982691 | 1,66134E-05 | PGKc_ PGK_ PGK3 phosphoglycerate kinase_ phosphoglycerate kinase 3 chr1:29924347-29926295 REVERSE LI |
| AT3G46520.1 | 1,799451792 | 1,66228E-05 | ACT12 actin-12 chr3:17128567-17129981 FORWARD LENGTH=377 |
| ATCG00830.1 | 0,061360658 | 1,67671E-05 | RPL2.1 ribosomal protein L2 chrc:84337-85843 REVERSE LENGTH=274 |
| AT3G04920.2 | 0,177341803 | 1,7228E-05 | no symbol available no full name available chr3:1360989-1361719 FORWARD LENGTH=112 |
| AT1G43170.1 | 0,041516529 | 1,82057E-05 | emb2207_ RP1_ ARP1_ RPL3A embryo defective 2207_ ribosomal protein 1 chr1:16266992-16268631 FORWAR |
| AT2G39310.1 | 5,260338282 | 1,97765E-05 | JAL22 jacalin-related lectin 22 chr2:16414262-16416323 REVERSE LENGTH=458 |
| AT3G23990.1 | 1,796010618 | 2,00762E-05 | HSP60_ HSP60-3B heat shock protein 60_ HEAT SHOCK PROTEIN 60-3B chr3:8669013-8672278 FORWARD I |

| | | | |
|---|---|---|---|
| AT1G65960.2 | 1,560077995 | 2,05147E-05 | GAD2 glutamate decarboxylase 2 chr1:24552094-24557253 FORWARD LENGTH=494 |
| AT4G15802.1 | 2,492903074 | 2,08369E-05 | AtHSBP_ HSBP Arabidopsis thaliana heat shock factor binding protein_ heat shock factor binding protein chr4:898 |
| AT5G49460.1 | 1,503236889 | 2,19535E-05 | ACLB-2 ATP citrate lyase subunit B 2 chr5:20055048-20058195 FORWARD LENGTH=608 |
| AT2G38230.1 | 1,700209038 | 2,20575E-05 | ATPDX1.1_ PDX1.1 pyridoxine biosynthesis 1.1_ ARABIDOPSIS THALIANA PYRIDOXINE BIOSYNTHESIS |
| AT5G47210.1 | 4,164209889 | 2,46546E-05 | no symbol available no full name available chr5:19169222-19171012 REVERSE LENGTH=357 |
| AT2G35635.1 | 5,058536369 | 2,60861E-05 | UBQ7_ RUB2 RELATED TO UBIQUITIN 2_ ubiquitin 7 chr2:14981044-14981943 FORWARD LENGTH=154 |
| AT2G24020.1 | 2,650835409 | 2,78157E-05 | STIC2 Suppressor of TIC40 2 chr2:10217869-10219269 REVERSE LENGTH=182 |
| AT1G80600.1 | 4,553603326 | 2,83245E-05 | TUP5_ WIN1 HOPW1-1-interacting 1_ TUMOR PRONE 5 chr1:30298675-30300513 REVERSE LENGTH=457 |
| AT2G24270.1 | 1,612369774 | 2,92654E-05 | ALDH11A3 aldehyde dehydrogenase 11A3 chr2:10327325-10329601 REVERSE LENGTH=496 |
| AT4G37910.1 | 1,950862304 | 2,96517E-05 | mtHsc70-1 mitochondrial heat shock protein 70-1 chr4:17825368-17828099 REVERSE LENGTH=682 |
| AT4G22240.1 | 1,449421629 | 3,14659E-05 | FBN1b fibrillin 1b chr4:11766090-11767227 REVERSE LENGTH=310 |
| AT4G04020.1 | 2,473550043 | 3,20431E-05 | FIB_ PGL35_ FIB1a plastoglobulin 35_ fibrillin 1a_ fibrillin chr4:1932161-1933546 FORWARD LENGTH=318 |
| AT4G11820.1 | 1,876189773 | 3,36441E-05 | MVA1_ FKP1_ HMGS FLAKY POLLEN 1_ HYDROXYMETHYLGLUTARYL-COA SYNTHASE chr4:710912 |
| AT5G01410.1 | 1,778832033 | 3,44056E-05 | PDX1_ ATPDX1.3_ ATPDX1_ PDX1.3_ RSR4 REDUCED SUGAR RESPONSE 4_ PYRIDOXINE BIOSYNTH |
| AT2G43030.1 | 0,606530885 | 3,52542E-05 | PRPL3 plastid ribosomal proteins of the 50S subunit chr2:17894898-17895713 FORWARD LENGTH=271 |
| AT3G58610.1 | 1,697777932 | 3,69291E-05 | no symbol available no full name available chr3:21671561-21674639 FORWARD LENGTH=591 |
| AT1G10760.1 | 2,048665872 | 3,8961E-05 | GWD_ GWD1_ SOP1_ SOP_ SEX1 STARCH EXCESS 1 chr1:3581210-3590043 REVERSE LENGTH=1399 |
| AT4G16830.2 | 1,842799864 | 3,98323E-05 | AtRGGA chr4:9470979-9472308 FORWARD LENGTH=265 |
| AT3G13920.4 | 1,972218823 | 4,03698E-05 | TIF4A1_ EIF4A1_ RH4 eukaryotic translation initiation factor 4A1 chr3:4592635-4594128 REVERSE LENGTH= |
| AT3G03710.1 | 1,885175357 | 4,13022E-05 | PNP_ RIF10_ PDE326 PIGMENT DEFECTIVE 326_ POLYNUCLEOTIDE PHOSPHORYLASE_ resistant to inh |
| AT1G31812.1 | 6,830139317 | 4,40606E-05 | ACBP6_ ACBP_ AtACBP6 acyl-CoA-binding protein 6_ ACYL-COA-BINDING PROTEIN chr1:11411132-1141 |
| AT2G36620.1 | 0,160084748 | 4,46766E-05 | RPL24A ribosomal protein L24 chr2:15350548-15351819 REVERSE LENGTH=164 |
| AT1G41880.1 | 0,469330969 | 4,51089E-05 | no symbol available no full name available chr1:15651585-15652427 REVERSE LENGTH=111 |
| AT3G13460.2 | 2,340548118 | 4,99451E-05 | ECT2 evolutionarily conserved C-terminal region 2 chr3:4385274-4388220 REVERSE LENGTH=664 |
| AT5G20980.1 | 1,210066065 | 5,49958E-05 | MS3_ ATMS3 methionine synthase 3 chr5:7124397-7128353 REVERSE LENGTH=812 |
| AT2G05620.1 | 3,667296507 | 5,55787E-05 | PGR5_ AtPGR5 proton gradient regulation 5 chr2:2081204-2081687 REVERSE LENGTH=133 |
| AT3G19450.1 | 1,572138985 | 5,62581E-05 | CAD-C_ CAD4_ ATCAD4_ CAD CINNAMYL ALCOHOL DEHYDROGENASE 4 chr3:6744859-6747005 FOR |
| AT3G01390.1 | 8,890836429 | 5,65215E-05 | AVMA10_ VMA10 vacuolar membrane ATPase 10 chr3:150265-150922 REVERSE LENGTH=110 |
| AT5G52920.1 | 1,897374612 | 5,70787E-05 | PKP-BETA1_ PKP2_ PKP1 plastidic pyruvate kinase beta subunit 1_ PLASTIDIAL PYRUVATE KINASE 1_ PL |
| AT3G49910.1 | 0,348941458 | 5,87613E-05 | no symbol available no full name available chr3:18504311-18504751 FORWARD LENGTH=146 |
| AT5G53490.1 | 1,395201499 | 5,95588E-05 | TL17 Thylakoid lumenal 17.4 kDa protein chr5:21723488-21724621 REVERSE LENGTH=236 |
| AT4G01150.1 | 0,575243599 | 6,06439E-05 | CURT1A CURVATURE THYLAKOID 1A chr4:493692-494668 FORWARD LENGTH=164 |
| AT2G37190.1 | 2,656322191 | 6,20014E-05 | no symbol available no full name available chr2:15619559-15620059 REVERSE LENGTH=166 |
| AT3G59970.1 | 2,766337717 | 6,90526E-05 | MTHFR1 methylenetetrahydrofolate reductase 1 chr3:22151303-22153412 FORWARD LENGTH=421 |
| AT2G30200.1 | 1,668359437 | 7,23568E-05 | EMB3147_ MCAT_ MCAMT EMBRYO DEFECTIVE 3147_ malonyl CoA-ACP malonyltransferase chr2:128831 |
| AT4G09010.2 | 0,653134546 | 7,24006E-05 | APX4_ TL29 ascorbate peroxidase 4_ thylakoid lumen 29 chr4:5777502-5779064 REVERSE LENGTH=284 |
| AT3G06650.1 | 1,358258328 | 7,42096E-05 | ACLB-1 ATP-citrate lyase B-1 chr3:2079247-2082633 REVERSE LENGTH=608 |
| AT5G12860.2 | 0,766052393 | 7,48237E-05 | DiT1 dicarboxylate transporter 1 chr5:4059850-4061919 REVERSE LENGTH=556 |
| AT1G65350.1 | 3,220642949 | 7,81464E-05 | UBQ13 ubiquitin 13 chr1:24272518-24277275 REVERSE LENGTH=319 |

| | | | |
|---|---|---|---|
| AT3G12290.1 | 1,474267965 | 7,99882E-05 | MTHFD1 methylenetetrahydrofolate dehydrogenase/methenyltetrahydrofolate cyclohydrolase chr3:3919591-39213[illegible] |
| AT3G09440.1 | 1,728100572 | 8,79845E-05 | no symbol available no full name available chr3:2903434-2905632 REVERSE LENGTH=649 |
| AT3G25230.1 | 1,798702434 | 8,82514E-05 | FKBP62_ ROF1_ ATFKBP62 rotamase FKBP 1_ FK506 BINDING PROTEIN 62 chr3:9188257-9191137 FORW[illegible] |
| AT4G00100.1 | 0,345120513 | 8,83283E-05 | RPS13_ RPS13A_ PFL2_ ATRPS13A ribosomal protein S13A_ POINTED FIRST LEAF 2 chr4:37172-38123 FO[illegible] |
| AT1G20200.1 | 0,271850519 | 8,90452E-05 | HAP15_ EMB2719 HAPLESS 15_ EMBRYO DEFECTIVE 2719 chr1:7001409-7004154 REVERSE LENGTH=4[illegible] |
| AT5G19820.1 | 1,861933788 | 9,03071E-05 | IMB3_ KETCH1_ EMB2734 EMBRYO DEFECTIVE 2734_ (karyopherin enabling the transport of the cytoplasmi[illegible] |
| AT2G34480.1 | 0,321191968 | 9,23425E-05 | L18aB_ RPL18aB chr2:14532916-14534161 REVERSE LENGTH=178 |
| AT5G20890.1 | 1,893986407 | 9,25428E-05 | CCT2 Chaperonin containing T-complex polypeptide-1 subunit 2 chr5:7087020-7089906 REVERSE LENGTH=52[illegible] |
| AT3G60770.1 | 0,344431065 | 9,29071E-05 | no symbol available no full name available chr3:22460525-22461656 REVERSE LENGTH=151 |
| ATCG00900.1 | 0,464019669 | 9,3712E-05 | RPS7_ RPS7.1 CHLOROPLAST RIBOSOMAL PROTEIN S7 chrc:97478-97945 REVERSE LENGTH=155 |
| AT5G56500.1 | 0,543492345 | 9,72911E-05 | CPNB3_ Cpn60beta3 chaperonin-60beta3 chr5:22874058-22876966 FORWARD LENGTH=597 |
| AT2G40010.1 | 1,934697119 | 0,000100417 | no symbol available no full name available chr2:16708578-16710448 REVERSE LENGTH=317 |
| AT4G24280.1 | 1,41918277 | 0,000101219 | cpHsc70-1 chloroplast heat shock protein 70-1 chr4:12590094-12593437 FORWARD LENGTH=718 |
| AT1G07920.1 | 1,125709629 | 0,000101611 | ELONGATION FACTOR-TU FAMILY chr1:2455559-2457001 FORWARD LENGTH=449 |
| AT3G09790.1 | 1,914038009 | 0,000101941 | UBQ8 ubiquitin 8 chr3:3004111-3006006 REVERSE LENGTH=631 |
| AT4G17090.1 | 1,735572233 | 0,000102595 | CT-BMY_ BMY8_ AtBAM3_ BAM3 BETA-AMYLASE 8_ BETA-AMYLASE 3_ chloroplast beta-amylase chr4:[illegible] |
| AT1G72370.2 | 1,1429141 | 0,000103047 | RPSAA_ AP40_ RP40_ P40 40s ribosomal protein SA chr1:27243148-27244842 REVERSE LENGTH=294 |
| AT5G52470.1 | 2,809697188 | 0,000104215 | ATFIB1_ FBR1_ FIB1_ SKIP7_ ATFBR1 fibrillarin 1_ FIBRILLARIN 1_ SKP1/ASK1-INTERACTING PROTEI[illegible] |
| AT4G30270.1 | 0,54496125 | 0,000105816 | MERI5B_ XTH24_ MERI-5_ SEN4 xyloglucan endotransglucosylase/hydrolase 24_ SENESCENCE 4_ MERISTE[illegible] |
| AT1G70310.1 | 1,654999026 | 0,000120169 | SPDS2 spermidine synthase 2 chr1:26485497-26487352 REVERSE LENGTH=340 |
| ATCG00790.1 | 0,458115734 | 0,000122049 | RPL16 ribosomal protein L16 chrc:81189-82652 REVERSE LENGTH=135 |
| AT1G64520.1 | 1,443743887 | 0,00012458 | RPN12a regulatory particle non-ATPase 12A chr1:23956459-23958120 FORWARD LENGTH=267 |
| AT1G57720.1 | 1,648561004 | 0,000125018 | no symbol available no full name available chr1:21377873-21380114 FORWARD LENGTH=413 |
| AT3G26060.1 | 2,154585118 | 0,000131452 | ATPRX Q_ PRXQ peroxiredoxin Q chr3:9524807-9526123 FORWARD LENGTH=216 |
| AT4G09010.1 | 1,418919075 | 0,000133014 | APX4_ TL29 ascorbate peroxidase 4_ thylakoid lumen 29 chr4:5777502-5779338 REVERSE LENGTH=349 |
| AT5G02500.1 | 1,74147632 | 0,000133313 | AtHsp70-1_ HSP70-1_ HSC70-1_ HSC70_ AT-HSC70-1 ARABIDOPSIS THALIANA HEAT SHOCK COGNAT[illegible] |
| AT5G42980.1 | 1,279078702 | 0,000140648 | ATH3_ TRX3_ TRXH3_ ATTRX3_ ATTRXH3 THIOREDOXIN H3_ thioredoxin 3_ thioredoxin H-type 3 chr5:1[illegible] |
| AT3G52880.1 | 1,363585942 | 0,000143259 | ATMDAR1_ MDAR1 monodehydroascorbate reductase 1 chr3:19601477-19604366 REVERSE LENGTH=434 |
| AT5G28060.1 | 0,396027161 | 0,000143937 | RPS24B chr5:10069791-10070792 REVERSE LENGTH=133 |
| AT1G18540.1 | 0,722721433 | 0,000149204 | no symbol available no full name available chr1:6377448-6378548 REVERSE LENGTH=233 |
| AT1G74040.1 | 1,885088355 | 0,000150719 | IPMS2_ IMS1_ MAML-3 SOPROPYLMALATE SYNTHASE 2_ 2-isopropylmalate synthase 1 chr1:27842258-27[illegible] |
| AT4G29350.1 | 1,91069879 | 0,000151292 | PFN2_ PRO2_ PRF2_ AtPRF2 PROFILIN 2_ profilin 2 chr4:14450135-14451119 FORWARD LENGTH=131 |
| AT1G66580.1 | 0,249719527 | 0,000154129 | SAG24_ RPL10C senescence associated gene 24_ ribosomal protein L10 C chr1:24839208-24840439 FORWARD |
| AT3G12145.1 | 0,156460228 | 0,000154983 | FLR1_ FTM4 FLOR1_ FLORAL TRANSITION AT THE MERISTEM4 chr3:3874764-3876075 REVERSE LEN[illegible] |
| AT1G12250.2 | 0,480539905 | 0,000157782 | TL20.3 chr1:4159623-4161269 FORWARD LENGTH=206 |
| AT3G02530.1 | 1,664770062 | 0,000164769 | CCT6-2 Chaperonin containing T-complex polypeptide-1 subunit 6-2 chr3:528806-532457 REVERSE LENGTH=5[illegible] |
| AT1G03230.1 | 1,544629577 | 0,000169458 | SAP1 secreted aspartic protease 1 chr1:790110-791414 FORWARD LENGTH=434 |
| AT2G28790.1 | 0,405511429 | 0,000169474 | no symbol available no full name available chr2:12354664-12355413 REVERSE LENGTH=249 |
| AT3G24830.1 | 0,286826924 | 0,000170193 | no symbol available no full name available chr3:9064613-9065871 FORWARD LENGTH=206 |

| | | | |
|---|---|---|---|
| AT5G47890.1 | 4,970524017 | 0,000173217 | no symbol available no full name available chr5:19388806-19390409 FORWARD LENGTH=97 |
| AT5G52520.1 | 1,504805664 | 0,000176325 | ProRS-Org_ OVA6_ PRORS1_ AtProRS-Org PROLYL-TRNA SYNTHETASE 1_ OVULE ABORTION 6_ proly |
| AT5G23740.1 | 1,659527136 | 0,000180663 | RPS11-BETA ribosomal protein S11-beta chr5:8008251-8009330 REVERSE LENGTH=159 |
| AT5G22800.1 | 1,700321643 | 0,00018503 | EMB263_ EMB1030_ EMB86 EMBRYO DEFECTIVE 263_ EMBRYO DEFECTIVE 1030_ EMBRYO DEFECT |
| AT3G11130.1 | 1,461099416 | 0,000185902 | AtCHC1_ CHC1_ HAS1 clathrin heavy chain 1_ hot ABA deficiency suppressor 1 chr3:3482575-3491667 REVER |
| AT3G56340.1 | 0,178065895 | 0,000187829 | RPS26e Ribosomal Protein S26e chr3:20892309-20893343 REVERSE LENGTH=130 |
| AT3G54210.1 | 0,179355877 | 0,000188157 | PRPL17 plastid ribosomal proteins of the 50S subunit 17 chr3:20067672-20068385 REVERSE LENGTH=211 |
| AT3G01120.1 | 1,850344113 | 0,000202959 | AtCYS1_ CGS_ AtCGS1_ CGS1_ MTO1 CYSTATHIONINE GAMMA-SYNTHASE 1_ CYSTATHIONINE GA |
| AT1G52230.1 | 0,729376077 | 0,000209311 | PSI-H_ PSAH-2_ PSAH2 photosystem I subunit H2_ PHOTOSYSTEM I SUBUNIT H-2 chr1:19454902-1945550 |
| AT4G25630.1 | 0,501506622 | 0,000217843 | ATFIB2_ FIB2 fibrillarin 2 chr4:13074239-13076205 FORWARD LENGTH=320 |
| AT1G68010.1 | 1,186929225 | 0,000218124 | HPR_ ATHPR1 hydroxypyruvate reductase chr1:25493418-25495720 FORWARD LENGTH=386 |
| AT5G44500.1 | 4,78733564 | 0,000219688 | no symbol available no full name available chr5:17927505-17928269 FORWARD LENGTH=254 |
| AT1G60950.1 | 0,248485253 | 0,000223391 | ATFD2_ FD2_ FED A FERREDOXIN 2 chr1:22444565-22445011 FORWARD LENGTH=148 |
| AT3G12780.1 | 1,226735917 | 0,000225647 | PGKp1_ PGK1 phosphoglycerate kinase 1 chr3:4061127-4063140 REVERSE LENGTH=481 |
| AT3G03960.1 | 1,79495795 | 0,000233051 | CCT8 Chaperonin containing T-complex polypeptide-1 subunit 8 chr3:1024432-1027604 FORWARD LENGTH=5 |
| AT5G54770.1 | 2,500571426 | 0,000241117 | THI1_ TZ_ THI4 THIAZOLE REQUIRING_ THIAMINE4 chr5:22246634-22247891 FORWARD LENGTH=349 |
| AT3G28220.1 | 0,294530643 | 0,000253953 | no symbol available no full name available chr3:10524420-10526497 FORWARD LENGTH=370 |
| AT2G31790.1 | 1,272278368 | 0,000254044 | no symbol available no full name available chr2:13518269-13520167 FORWARD LENGTH=457 |
| AT1G31230.1 | 1,832005328 | 0,000259677 | AK-HSDH I_ AK-HSDH ASPARTATE KINASE-HOMOSERINE DEHYDROGENASE_ aspartate kinase-homos |
| AT1G16080.1 | 1,693157627 | 0,000261689 | no symbol available no full name available chr1:5514394-5515761 FORWARD LENGTH=313 |
| AT3G19710.1 | 1,547383803 | 0,000265262 | BCAT4 branched-chain aminotransferase4 chr3:6847202-6849429 REVERSE LENGTH=354 |
| AT1G66240.1 | 1,37931374 | 0,000266654 | AtHMP14_ ATX1_ ATATX1 HEAVY METAL ASSOCIATED PROTEIN 14_ homolog of anti-oxidant 1 chr1:24 |
| AT2G06050.1 | 0,485131863 | 0,000270906 | OPR3_ DDE1_ AtOPR3 DELAYED DEHISCENCE 1_ oxophytodienoate-reductase 3 chr2:2359240-2361971 RE |
| AT1G19670.1 | 0,354885158 | 0,000277066 | CLH1_ ATCLH1_ CORI1_ ATHCOR1 chlorophyllase 1_ CORONATINE-INDUCED PROTEIN 1 chr1:6803796- |
| AT1G78820.1 | 0,434931839 | 0,000278912 | no symbol available no full name available chr1:29634401-29635768 REVERSE LENGTH=455 |
| AT3G57490.1 | 0,505562902 | 0,00028093 | no symbol available no full name available chr3:21279824-21280887 REVERSE LENGTH=276 |
| AT2G43750.1 | 1,162672649 | 0,000282295 | OASB_ CPACS1_ ACS1_ ATCS-B O-acetylserine (thiol) lyase B_ ARABIDOPSIS THALIANA CYSTEIN SYNT |
| AT5G14740.5 | 1,634426724 | 0,000284224 | BETA CA2_ CA2_ DEG12_ CA18 BETA CARBONIC ANHYDRASE 2_ CARBONIC ANHYDRASE 18_ carbo |
| AT3G04870.1 | 1,45442811 | 0,000293924 | PDE181_ ZDS_ SPC1 SPONTANEOUS CELL DEATH 1_ PIGMENT DEFECTIVE EMBRYO 181_ zeta-caroten |
| AT1G55260.2 | 4,5831844 | 0,000296751 | LTPG6 glycosylphosphatidylinositol-anchored lipid protein transfer 6 chr1:20614663-20616158 FORWARD LEN |
| AT1G09590.1 | 0,281969298 | 0,000299705 | no symbol available no full name available chr1:3106549-3107606 FORWARD LENGTH=164 |
| AT3G03250.1 | 1,502756657 | 0,000303978 | AtUGP1_ UGP_ UGP1 UDP-glucose pyrophosphorylase_ UDP-GLUCOSE PYROPHOSPHORYLASE 1 chr3:74 |
| AT1G14320.2 | 0,275128265 | 0,000305192 | RPL10A_ SAC52_ RPL10 SUPPRESSOR OF ACAULIS 52_ ribosomal protein L10_ ribosomal protein L10 A chr |
| AT5G14320.1 | 5,30982018 | 0,000309853 | EMB3137 EMBRYO DEFECTIVE 3137 chr5:4617839-4618772 REVERSE LENGTH=169 |
| AT2G17360.1 | 0,628926599 | 0,000314939 | no symbol available no full name available chr2:7546598-7548138 FORWARD LENGTH=261 |
| AT1G05010.1 | 0,518498164 | 0,000326985 | EFE_ ACO4_ EAT1 ethylene forming enzyme_ ethylene-forming enzyme chr1:1431419-1432695 REVERSE LEN |
| AT1G35580.3 | 2,322987418 | 0,000340068 | CINV1_ A/N-InvG_ NIN2 cytosolic invertase 1_ alkaline/neutral invertase G_ neutral invertase 2 chr1:13123183-1 |
| AT3G19170.1 | 1,480866536 | 0,00034016 | ATZNMP_ ATPREP1_ PREP1 presequence protease 1 chr3:6625578-6631874 REVERSE LENGTH=1080 |
| AT3G08590.1 | 1,743595862 | 0,000340631 | iPGAM2 "2_3-biphosphoglycerate-independent phosphoglycerate mutase 2" chr3:2608683-2611237 REVERSE LE |

| | | | |
|---|---|---|---|
| AT5G60600.1 | 1,299035245 | 0,000341511 | ISPG_ CSB3_ HDS_ GCPE_ CLB4 CONSTITUTIVE SUBTILISIN 3_ 4-hydroxy-3-methylbut-2-enyl diphosphate |
| AT1G18210.1 | 0,47353961 | 0,0003468 | no symbol available no full name available chr1:6268273-6268785 REVERSE LENGTH=170 |
| AT3G12110.1 | 1,700379081 | 0,000351007 | ACT11 actin-11 chr3:3858116-3859609 FORWARD LENGTH=377 |
| AT5G60360.2 | 0,452352414 | 0,000356532 | AALP_ SAG2_ ALP aleurain-like protease_ SENESCENCE ASSOCIATED GENE2 chr5:24280044-24282152 FO |
| AT1G72730.1 | 1,437166445 | 0,000365335 | no symbol available no full name available chr1:27378040-27379593 REVERSE LENGTH=414 |
| AT3G57890.1 | 1,477610996 | 0,000379902 | no symbol available no full name available chr3:21438271-21441695 FORWARD LENGTH=573 |
| AT1G80380.4 | 1,279255869 | 0,000393143 | GLYK glycerate kinase chr1:30217332-30219784 FORWARD LENGTH=450 |
| AT5G13650.1 | 1,491396496 | 0,000394969 | SVR3 SUPPRESSOR OF VARIEGATION 3 chr5:4397821-4402364 FORWARD LENGTH=675 |
| AT5G44020.1 | 1,802664372 | 0,000401012 | no symbol available no full name available chr5:17712433-17714046 FORWARD LENGTH=272 |
| AT5G03630.1 | 1,406914584 | 0,000405296 | MDAR2 chr5:922378-924616 REVERSE LENGTH=435 |
| AT5G67030.2 | 1,607399528 | 0,000406966 | LOS6_ NPQ2_ ZEP_ ABA1_ ATABA1_ ATZEP_ IBS3 ABA DEFICIENT 1_ IMPAIRED IN BABA-INDUCED |
| AT2G23350.1 | 1,538963651 | 0,000413743 | PABP4_ PAB4 POLY(A) BINDING PROTEIN 4_ poly(A) binding protein 4 chr2:9943209-9946041 FORWARD |
| AT5G17170.1 | 0,77560115 | 0,000418787 | ENH1 enhancer of sos3-1 chr5:5649335-5650975 FORWARD LENGTH=271 |
| AT1G11750.1 | 1,551764414 | 0,000423131 | NCLPP1_ CLPP6_ NCLPP6 NUCLEAR-ENCODED CLPP 1_ CLP protease proteolytic subunit 6 chr1:3967609-3 |
| AT4G34200.1 | 1,781679551 | 0,000431921 | EDA9_ PGDH1 phosphoglycerate dehydrogenase 1_ embryo sac development arrest 9 chr4:16374041-16376561 R |
| AT1G76160.1 | 4,515354699 | 0,00043377 | sks5 SKU5 similar 5 chr1:28578211-28581020 REVERSE LENGTH=541 |
| AT4G39980.1 | 1,481205063 | 0,000434331 | AtDAHP1_ DHS1_ DAHP1 3-DEOXY-D-ARABINO-HEPTULOSONATE-7-PHOSPHATE 1_ 3-deoxy-D-arabin |
| AT5G04140.1 | 0,951421518 | 0,000436685 | GLUS_ FD-GOGAT_ GLS1_ GLU1 FERREDOXIN-DEPENDENT GLUTAMATE SYNTHASE 1_ glutamate syn |
| AT1G14980.1 | 7,050934161 | 0,00043855 | CPN10 chaperonin 10 chr1:5165930-5166654 REVERSE LENGTH=98 |
| AT5G06870.1 | 0,365047772 | 0,000450599 | PGIP2_ ATPGIP2 ARABIDOPSIS POLYGALACTURONASE INHIBITING PROTEIN 2_ polygalacturonase inh |
| AT2G19730.1 | 2,88254558 | 0,000450685 | no symbol available no full name available chr2:8511752-8512995 FORWARD LENGTH=143 |
| AT1G32990.1 | 0,623452574 | 0,000462792 | PRPL11 plastid ribosomal protein l11 chr1:11955827-11957139 FORWARD LENGTH=222 |
| AT3G62870.1 | 0,592430258 | 0,000471834 | no symbol available no full name available chr3:23242862-23244273 REVERSE LENGTH=256 |
| AT1G55490.1 | 1,223026385 | 0,000472935 | Cpn60beta1_ LEN1_ CPNB1_ CPN60B chaperonin-60beta1_ LESION INITIATION 1_ chaperonin 60 beta chr1:2 |
| AT3G61440.1 | 0,860799486 | 0,000477031 | ATCYSC1_ ARATH;BSAS3;1_ CAS-C1_ CYSC1 ß-cyanoalanine synthase C1_ CYSTEINE SYNTHASE C1_ BE |
| AT2G44650.1 | 1,445526278 | 0,000480161 | CHL-CPN10_ CPN10 CHLOROPLAST CHAPERONIN 10_ chloroplast chaperonin 10 chr2:18419521-18420510 |
| AT5G27850.1 | 0,570564715 | 0,000480306 | RPL18C chr5:9873169-9874297 FORWARD LENGTH=187 |
| AT3G12915.2 | 1,579779898 | 0,000483171 | no symbol available no full name available chr3:4112834-4115708 FORWARD LENGTH=788 |
| AT3G12580.1 | 2,024140892 | 0,000483759 | HSP70_ ATHSP70_ HSC70-4 ARABIDOPSIS HEAT SHOCK PROTEIN 70_ heat shock protein 70 chr3:3991487 |
| AT5G11880.1 | 1,4779839 | 0,000485423 | DAPDC2 meso-diaminopimelate decarboxylase 2 chr5:3827806-3829942 REVERSE LENGTH=489 |
| AT2G39390.1 | 0,205449695 | 0,000488915 | no symbol available no full name available chr2:16450803-16451762 REVERSE LENGTH=123 |
| AT1G62750.1 | 1,383617971 | 0,000500005 | ATSCO1/CPEF-G_ ATSCO1_ SCO1 SNOWY COTYLEDON 1 chr1:23233622-23236321 REVERSE LENGTH= |
| AT4G05180.1 | 1,375828043 | 0,000502229 | PSBQ-2_ PSBQ_ PSII-Q photosystem II subunit Q-2_ PHOTOSYSTEM II SUBUNIT Q chr4:2672093-2673170 R |
| AT4G21280.1 | 1,579677314 | 0,000504857 | PSBQ-1_ PSBQA_ PSBQ photosystem II subunit QA_ PHOTOSYSTEM II SUBUNIT Q-1_ PHOTOSYSTEM II |
| AT1G20630.1 | 1,609331266 | 0,00050498 | CAT1 catalase 1 chr1:7146812-7149609 FORWARD LENGTH=492 |
| AT4G33680.1 | 1,379654527 | 0,000540726 | AGD2 ABERRANT GROWTH AND DEATH 2_ ARF-GAP domain 2 chr4:16171847-16174630 REVERSE LEN |
| AT3G48930.1 | 0,581685131 | 0,00054785 | EMB1080 embryo defective 1080 chr3:18141017-18142189 REVERSE LENGTH=160 |
| AT1G13930.1 | 7,850006863 | 0,000548861 | no symbol available no full name available chr1:4761091-4761558 FORWARD LENGTH=155 |
| AT5G64140.1 | 1,703255581 | 0,000553793 | RPS28 ribosomal protein S28 chr5:25667529-25667723 REVERSE LENGTH=64 |

| | | | |
|---|---|---|---|
| AT5G28500.1 | 1,356708124 | 0,000561311 | no symbol available no full name available chr5:10477810-10479114 FORWARD LENGTH=434 |
| AT4G35250.1 | 1,352815181 | 0,000581934 | HCF244 high chlorophyll fluorescence phenotype 244 chr4:16771401-16773269 REVERSE LENGTH=395 |
| AT4G10320.1 | 2,185115793 | 0,000595228 | no symbol available no full name available chr4:6397526-6404509 REVERSE LENGTH=1190 |
| AT2G37270.1 | 0,777547748 | 0,000610486 | RPS5B_ ATRPS5B ribosomal protein 5B chr2:15647883-15649042 REVERSE LENGTH=207 |
| AT5G10360.1 | 0,187216089 | 0,000613472 | RPS6B_ EMB3010 Ribosomal protein small subunit 6b_ embryo defective 3010 chr5:3258734-3260142 REVERSE |
| AT5G13850.1 | 0,594808959 | 0,000620777 | NACA3 nascent polypeptide-associated complex subunit alpha-like protein 3 chr5:4471361-4472676 FORWARD L |
| AT3G15190.1 | 0,423914688 | 0,0006297 | PRPS20 plastid ribosomal protein S20 chr3:5116216-5117412 FORWARD LENGTH=202 |
| AT4G33010.1 | 1,197768737 | 0,000631562 | GLDP1_ AtGLDP1 glycine decarboxylase P-protein 1 chr4:15926852-15931150 REVERSE LENGTH=1037 |
| AT3G45140.1 | 0,457821081 | 0,000645066 | ATLOX2_ LOX2 ARABIODOPSIS THALIANA LIPOXYGENASE 2_ lipoxygenase 2 chr3:16525437-16529233 |
| AT1G79930.1 | 1,40039844 | 0,00067685 | HSP91_ AtHsp70-14 heat shock protein 91 chr1:30063781-30067067 REVERSE LENGTH=831 |
| AT1G07320.3 | 0,742767672 | 0,000677841 | RPL4_ PRPL4_ EMB2784 plastid ribosomal protein L4_ ribosomal protein L4_ EMBRYO DEFECTIVE 2784 chr |
| AT3G07110.1 | 0,322889654 | 0,000678042 | no symbol available no full name available chr3:2252092-2253332 FORWARD LENGTH=206 |
| AT1G68560.1 | 0,639466325 | 0,000684942 | XYL1_ GH31_ TRG1_ ATXYL1_ AXY3 altered xyloglucan 3_ ALPHA-XYLOSIDASE 1_ alpha-xylosidase 1_ th |
| AT1G30530.1 | 1,851434605 | 0,000688141 | UGT78D1 UDP-glucosyl transferase 78D1 chr1:10814917-10816374 FORWARD LENGTH=453 |
| AT3G11830.1 | 1,56188939 | 0,000690243 | CCT7 Chaperonin containing T-complex polypeptide-1 subunit 7 chr3:3732734-3736156 FORWARD LENGTH=5 |
| AT1G04820.1 | 1,249452966 | 0,000693423 | TUA4_ TOR2 TORTIFOLIA 2_ tubulin alpha-4 chain chr1:1356421-1358266 REVERSE LENGTH=450 |
| AT1G48860.1 | 1,682436471 | 0,000696419 | EPSPS 5-enolpyruvylshikimate-3-phosphate synthase chr1:18068892-18071331 REVERSE LENGTH=521 |
| AT3G02880.1 | 2,27925163 | 0,00069874 | KIN7 Kinase 7 chr3:634819-636982 FORWARD LENGTH=627 |
| AT1G12250.1 | 0,558425622 | 0,000703561 | TL20.3 chr1:4159287-4161269 FORWARD LENGTH=280 |
| AT1G56410.1 | 1,413294779 | 0,000715027 | HSP70T-1_ ERD2 HEAT SHOCK PROTEIN 70T-1_ EARLY-RESPONSIVE TO DEHYDRATION 2 chr1:21117 |
| AT3G25760.1 | 0,393280789 | 0,000720879 | AOC1_ ERD12 early-responsive to dehydration 12_ allene oxide cyclase 1 chr3:9403972-9405105 FORWARD LE |
| AT5G65430.2 | 0,612989143 | 0,000724033 | 14-3-3KAPPA_ GRF8_ AtMIN10_ GF14 KAPPA general regulatory factor 8_ 14-3-3 PROTEIN G-BOX FACTOR |
| AT5G66760.1 | 0,661075959 | 0,000735102 | SDH1-1 succinate dehydrogenase 1-1 chr5:26653776-26657224 FORWARD LENGTH=634 |
| AT5G07350.1 | 1,792538001 | 0,000737286 | Tudor1_ TSN1_ AtTudor1 TUDOR-SN protein 1_ Arabidopsis thaliana TUDOR-SN protein 1 chr5:2320344-2324 |
| AT1G29670.1 | 0,677637353 | 0,000744211 | GDSL1_ GGL6 chr1:10375843-10377717 FORWARD LENGTH=363 |
| AT3G11250.1 | 0,756950179 | 0,000746236 | no symbol available no full name available chr3:3521453-3522826 FORWARD LENGTH=323 |
| AT3G25860.1 | 1,493523102 | 0,000753457 | LTA2_ PLE2 PLASTID E2 SUBUNIT OF PYRUVATE DECARBOXYLASE chr3:9460632-9462585 FORWARD |
| AT2G20580.1 | 1,783381543 | 0,000763854 | ATRPN1A_ RPN1A 26S PROTEASOME REGULATORY SUBUNIT S2 1A_ 26S proteasome regulatory subunit |
| AT1G19570.1 | 0,624913055 | 0,000765897 | ATDHAR1_ DHAR1_ DHAR5 DEHYDROASCORBATE REDUCTASE 5_ dehydroascorbate reductase chr1:677 |
| AT1G23740.1 | 1,410075429 | 0,000779043 | AOR alkenal/one oxidoreductase chr1:8398245-8399656 REVERSE LENGTH=386 |
| AT5G07340.1 | 1,644990789 | 0,000800929 | no symbol available no full name available chr5:2317300-2319458 FORWARD LENGTH=532 |
| AT4G24780.1 | 0,594977833 | 0,000821632 | PLL19 chr4:12770631-12772227 REVERSE LENGTH=408 |
| AT5G14910.1 | 1,73500022 | 0,000831792 | no symbol available no full name available chr5:4823815-4825196 FORWARD LENGTH=178 |
| AT5G48300.1 | 1,422500222 | 0,000833708 | ADG1_ APS1 ADP-GLUCOSE PYROPHOSPHORYLASE SMALL SUBUNIT 1_ ADP glucose pyrophosphoryla |
| AT3G01480.1 | 1,262398135 | 0,000847454 | CYP38_ ATCYP38 cyclophilin 38_ ARABIDOPSIS CYCLOPHILIN 38 chr3:188569-190674 FORWARD LENG |
| AT1G74270.1 | 0,46133883 | 0,000857826 | no symbol available no full name available chr1:27928415-27929466 REVERSE LENGTH=112 |
| AT1G54020.1 | 0,32871548 | 0,000868534 | no symbol available no full name available chr1:20161805-20162923 REVERSE LENGTH=286 |
| AT3G46830.1 | 0,128768571 | 0,00087381 | ATRAB-A2C_ ATRAB11A_ ATRABA2C_ RABA2c_ RAB-A2C RAB GTPASE HOMOLOG A2C_ ARABIDOP |
| AT3G19760.1 | 4,136033268 | 0,000902808 | EIF4A-III_ RH2 eukaryotic initiation factor 4A-III chr3:6863790-6866242 FORWARD LENGTH=408 |

| | | | |
|---|---|---|---|
| AT5G02490.1 | 1,288720089 | 0,000914529 | AtHsp70-2_ Hsp70-2 chr5:550296-552565 REVERSE LENGTH=653 |
| AT5G59370.1 | 1,192434678 | 0,000919515 | ACT4 actin 4 chr5:23950109-23951586 FORWARD LENGTH=377 |
| AT3G10090.1 | 2,530291094 | 0,000926235 | no symbol available no full name available chr3:3108960-3109154 REVERSE LENGTH=64 |
| AT3G44300.1 | 1,566280525 | 0,00095192 | AtNIT2_ NIT2 nitrilase 2 chr3:15983351-15985172 FORWARD LENGTH=339 |
| AT1G48600.1 | 2,504786258 | 0,000964329 | AtPMT2_ AtPMEAMT_ PMEAMT phosphoethanolamine N-methyltransferase_ Phosphoethanolamine methyltrans |
| AT1G53310.1 | 0,855075148 | 0,000971153 | PPC1_ ATPEPC1_ PEPC1_ ATPPC1 phosphoenolpyruvate carboxylase 1_ PEP(PHOSPHOENOLPYRUVATE) C |
| AT3G27830.1 | 1,325902369 | 0,000981408 | RPL12-A_ RPL12 ribosomal protein L12-A_ RIBOSOMAL PROTEIN L12 chr3:10318576-10319151 FORWARD |
| AT3G25530.1 | 1,329827317 | 0,001057501 | AtGLYR1_ GLYR1_ ATGHBDH_ GHBDH_ GR1 GLYOXYLATE REDUCTASE 1_ glyoxylate reductase 1 chr3 |
| AT3G47070.1 | 3,572089947 | 0,001071654 | no symbol available no full name available chr3:17337205-17337507 REVERSE LENGTH=100 |
| AT1G55670.1 | 0,626718961 | 0,001089568 | PSAG photosystem I subunit G chr1:20802874-20803356 REVERSE LENGTH=160 |
| AT2G28000.1 | 1,702027865 | 0,001106346 | ARC2_ CH-CPN60A_ SLP_ CPN60A_ Cpn60alpha1_ CPNA1 SCHLEPPERLESS_ chaperonin-60alpha1_ CHLO |
| AT5G09660.1 | 1,278240492 | 0,001107338 | PMDH2 peroxisomal NAD-malate dehydrogenase 2 chr5:2993645-2995551 REVERSE LENGTH=354 |
| AT1G62660.1 | 0,417848536 | 0,001111657 | VI1 VACUOLAR INVERTASE 1 chr1:23199949-23203515 FORWARD LENGTH=648 |
| AT4G21650.1 | 0,484080629 | 0,001129306 | SBT3.13 subtilase 3.13 chr4:11501314-11504656 REVERSE LENGTH=766 |
| AT1G48830.1 | 0,706251457 | 0,001132691 | no symbol available no full name available chr1:18059854-18060935 REVERSE LENGTH=191 |
| AT4G18810.1 | 1,604123482 | 0,001141382 | no symbol available no full name available chr4:10322622-10325735 REVERSE LENGTH=596 |
| AT5G59890.2 | 0,560764206 | 0,001149215 | ATADF4_ ADF4 actin depolymerizing factor 4 chr5:24123107-24123596 FORWARD LENGTH=132 |
| AT1G28290.2 | 0,738303285 | 0,001152886 | AGP31 arabinogalactan protein 31 chr1:9889331-9890843 REVERSE LENGTH=315 |
| AT1G50250.1 | 0,820660516 | 0,001182177 | FTSH1 FTSH protease 1 chr1:18614398-18616930 REVERSE LENGTH=716 |
| AT1G16470.1 | 1,252275374 | 0,001192831 | PAB1 proteasome subunit PAB1 chr1:5623122-5625439 FORWARD LENGTH=235 |
| AT5G66120.2 | 1,547144447 | 0,001193871 | no symbol available no full name available chr5:26431516-26433649 REVERSE LENGTH=442 |
| AT1G07660.1 | 0,795572383 | 0,001205576 | no symbol available no full name available chr1:2369212-2369523 FORWARD LENGTH=103 |
| AT4G35100.1 | 0,142924533 | 0,001207242 | PIP3A_ SIMIP_ PIP3_ PIP2;7 plasma membrane intrinsic protein 3_ PLASMA MEMBRANE INTRINSIC PROTE |
| AT1G76180.1 | 0,404268024 | 0,001219461 | ERD14 EARLY RESPONSE TO DEHYDRATION 14 chr1:28587013-28587657 REVERSE LENGTH=185 |
| AT5G27640.1 | 1,671951598 | 0,00124174 | ATTIF3B1_ TIF3B1_ EIF3B_ ATEIF3B-1_ EIF3B-1 ARABIDOPSIS THALIANA TRANSLATION INITIATION |
| AT3G60750.1 | 1,28906683 | 0,001249954 | AtTKL1_ TKL1 transketolase 1 chr3:22454004-22456824 FORWARD LENGTH=741 |
| AT1G52000.1 | 0,491792625 | 0,001265203 | no symbol available no full name available chr1:19333352-19335700 REVERSE LENGTH=730 |
| AT1G09640.1 | 1,560859821 | 0,001266747 | no symbol available no full name available chr1:3120162-3122152 FORWARD LENGTH=414 |
| AT4G27585.1 | 0,700410247 | 0,001292508 | SLP1_ AtSLP1 stomatin-like protein 1 chr4:13766984-13769832 REVERSE LENGTH=411 |
| AT5G45280.2 | 0,530316005 | 0,001302261 | PAE11 pectin acetylesterase 11 chr5:18346862-18349488 FORWARD LENGTH=391 |
| ATCG00420.1 | 0,65417593 | 0,001321077 | NDHJ NADH dehydrogenase subunit J chrc:48677-49153 REVERSE LENGTH=158 |
| AT4G28080.1 | 2,153379937 | 0,001321735 | REC2 REDUCED CHLOROPLAST COVERAGE 2 chr4:13948993-13957840 REVERSE LENGTH=1819 |
| AT4G11600.1 | 1,483454102 | 0,001328429 | GPXL6_ PHGPX_ LSC803_ ATGPX6_ GPX6 glutathione peroxidase 6 chr4:7010021-7011330 REVERSE LENG |
| AT4G24190.1 | 1,503972265 | 0,00135587 | SHD_ HSP90_ HSP90.7_ AtHsp90-7_ AtHsp90.7 SHEPHERD_ HEAT SHOCK PROTEIN 90.7_ HEAT SHOCK |
| AT4G13940.1 | 1,420198507 | 0,001375262 | MEE58_ SAHH1_ EMB1395_ HOG1_ SAH1_ ATSAHH1 EMBRYO DEFECTIVE 1395_ HOMOLOGY-DEPEN |
| AT2G47940.2 | 1,231123574 | 0,001378712 | DEG2_ DEGP2_ EMB3117 DEGP protease 2_ EMBRYO DEFECTIVE 3117_ degradation of periplasmic proteins |
| AT1G20440.1 | 0,758599624 | 0,001382894 | AtCOR47_ RD17_ COR47 cold-regulated 47 chr1:7084722-7085664 REVERSE LENGTH=265 |
| AT4G24830.1 | 1,42188294 | 0,00139033 | no symbol available no full name available chr4:12793085-12795857 REVERSE LENGTH=494 |
| AT3G46430.1 | 1,705972555 | 0,001393788 | AtMtATP6 chr3:17087687-17088497 FORWARD LENGTH=55 |

| | | | |
|---|---|---|---|
| AT5G23860.1 | 1,627049324 | 0,001400685 | TUB8 tubulin beta 8 chr5:8042962-8044528 FORWARD LENGTH=449 |
| AT4G21990.1 | 2,827118623 | 0,001413554 | APR3_ PRH26_ PRH-26_ ATAPR3 PAPS REDUCTASE HOMOLOG 26_ APS reductase 3 chr4:11657284-1165[illegible] |
| ATCG00470.1 | 0,844755961 | 0,00142121 | ATPE ATP synthase epsilon chain chrc:52265-52663 REVERSE LENGTH=132 |
| AT2G35370.1 | 0,845875897 | 0,001427055 | GDCH glycine decarboxylase complex H chr2:14891239-14892050 FORWARD LENGTH=165 |
| AT5G50850.1 | 1,251051104 | 0,001433128 | MAB1 MACCI-BOU chr5:20689671-20692976 FORWARD LENGTH=363 |
| AT1G06000.1 | 2,357263045 | 0,001433318 | UGT89C1 chr1:1820495-1821802 REVERSE LENGTH=435 |
| AT5G23540.1 | 1,221612205 | 0,001435211 | no symbol available no full name available chr5:7937772-7939339 FORWARD LENGTH=308 |
| AT1G25490.1 | 1,315044456 | 0,001473522 | EER1_ ATB BETA BETA_ RCN1_ REGA ROOTS CURL IN NPA_ ENHANCED ETHYLENE RESPONSE 1 ch |
| AT5G10160.1 | 1,172202896 | 0,00148656 | no symbol available no full name available chr5:3185819-3187159 FORWARD LENGTH=219 |
| AT1G62180.1 | 2,80317609 | 0,001501979 | APSR_ PRH43_ PRH_ ATAPR2_ APR2 ADENOSINE-5'-PHOSPHOSULFATE REDUCTASE_ 3'-PHOSPHOAD |
| AT3G46970.1 | 1,57220116 | 0,001506871 | ATPHS2_ PHS2 alpha-glucan phosphorylase 2_ Arabidopsis thaliana alpha-glucan phosphorylase 2 chr3:17301625 |
| AT4G15440.1 | 0,219380932 | 0,001511135 | HPL1_ CYP74B2 hydroperoxide lyase 1 chr4:8835869-8838462 FORWARD LENGTH=384 |
| AT5G48810.1 | 0,679684796 | 0,001514077 | CB5-D_ ATB5-B_ ATCB5-D_ B5 #3_ CYTB5-B cytochrome B5 isoform D_ ARABIDOPSIS CYTOCHROME B |
| AT3G08580.1 | 1,199907419 | 0,001540948 | AAC1 ADP/ATP carrier 1 chr3:2605706-2607030 REVERSE LENGTH=381 |
| AT5G43330.1 | 0,789353474 | 0,001555104 | c-NAD-MDH2 cytosolic-NAD-dependent malate dehydrogenase 2 chr5:17390552-17392449 FORWARD LENGTH |
| AT5G06290.1 | 1,298744891 | 0,001557025 | 2-Cys Prx B_ 2CPB 2-cysteine peroxiredoxin B_ 2-CYS PEROXIREDOXIN B chr5:1919380-1921211 FORWARD |
| AT3G04790.1 | 1,189101951 | 0,001560032 | EMB3119 EMBRYO DEFECTIVE 3119 chr3:1313365-1314195 FORWARD LENGTH=276 |
| AT2G42520.1 | 1,851232 | 0,001564417 | RH37 RNA Helicase 37 chr2:17705382-17708744 FORWARD LENGTH=633 |
| AT2G33040.1 | 1,271288074 | 0,001581366 | ATP3 gamma subunit of Mt ATP synthase chr2:14018978-14021047 REVERSE LENGTH=325 |
| AT1G70410.1 | 0,818071003 | 0,001642146 | BCA4_ ATBCA4_ CA4 beta carbonic anhydrase 4_ BETA CARBONIC ANHYDRASE 4 chr1:26534167-2653650 |
| AT1G37130.1 | 1,59504493 | 0,001652548 | B29_ ATNR2_ NIA2_ NIA2-1_ NR2_ CHL3_ NR NITRATE REDUCTASE 2_ CHLORATE RESISTANT 3_ AR |
| AT3G62530.1 | 1,409882682 | 0,001661304 | no symbol available no full name available chr3:23132219-23133121 FORWARD LENGTH=221 |
| AT3G07390.1 | 0,792924285 | 0,001687023 | AIR12 Auxin-Induced in Root cultures 12 chr3:2365452-2366273 FORWARD LENGTH=273 |
| AT3G14415.1 | 1,313236387 | 0,00168782 | GOX2 glycolate oxidase 2 chr3:4818667-4820748 FORWARD LENGTH=367 |
| AT1G54040.2 | 0,844977738 | 0,001702272 | TASTY_ ESR_ ESP epithiospecifier protein_ EPITHIOSPECIFYING SENESCENCE REGULATOR chr1:201709 |
| AT2G07698.1 | 1,266620707 | 0,001751913 | no symbol available no full name available chr2:3361474-3364028 FORWARD LENGTH=777 |
| AT1G20950.1 | 1,641679408 | 0,001812841 | no symbol available no full name available chr1:7297467-7301336 REVERSE LENGTH=614 |
| AT5G03300.1 | 2,076632799 | 0,001838152 | ADK2 adenosine kinase 2 chr5:796573-798997 FORWARD LENGTH=345 |
| AT5G66570.1 | 0,915801059 | 0,001861584 | OEE1_ PSBO-1_ MSP-1_ OE33_ OEE33_ PSBO1 PS II OXYGEN-EVOLVING COMPLEX 1_ OXYGEN EVOL |
| AT3G58510.1 | 1,809817897 | 0,001887368 | RH11 RNA Helicase 11 chr3:21640608-21643464 FORWARD LENGTH=612 |
| AT5G41670.1 | 1,849874518 | 0,001914423 | PGD3 6-phosphogluconate dehydrogenase 3 chr5:16665647-16667110 REVERSE LENGTH=487 |
| AT1G66270.2 | 0,250959923 | 0,001944494 | BGLU21 chr1:24700110-24702995 REVERSE LENGTH=522 |
| AT4G28750.1 | 0,868782745 | 0,001978781 | PSAE-1 PSA E1 KNOCKOUT chr4:14202951-14203888 REVERSE LENGTH=143 |
| AT4G32260.1 | 1,280819089 | 0,001984358 | PDE334 PIGMENT DEFECTIVE 334 chr4:15573859-15574586 REVERSE LENGTH=219 |
| AT2G04030.1 | 1,254800925 | 0,001985869 | CR88_ EMB1956_ AtHsp90.5_ HSP90C_ AtHsp90C_ Hsp88.1_ HSP90.5 HEAT SHOCK PROTEIN 90.5_ EMB |
| AT5G14200.1 | 1,220682054 | 0,001991305 | ATIMD1_ IMD1 isopropylmalate dehydrogenase 1_ ARABIDOPSIS ISOPROPYLMALATE DEHYDROGENAS |
| AT1G04530.1 | 2,727887134 | 0,002004716 | TPR4 tetratricopeptide repeat 4 chr1:1234456-1235895 REVERSE LENGTH=310 |
| AT1G47260.1 | 0,6150894 | 0,002022488 | APFI_ GAMMA CA2 gamma carbonic anhydrase 2 chr1:17321384-17323347 REVERSE LENGTH=278 |
| AT3G08530.1 | 1,551906576 | 0,002053947 | AtCHC2_ CHC2 clathrin heavy chain 2 chr3:2587171-2595411 REVERSE LENGTH=1703 |

| | | | |
|---|---|---|---|
| AT3G58990.1 | 1,96267443 | 0,002145205 | IPMI SSU3_ IPMI1 isopropylmalate isomerase 1 chr3:21797524-21798285 REVERSE LENGTH=253 |
| AT4G27560.1 | 1,986385284 | 0,002147462 | UGT79B2 chr4:13760114-13761481 REVERSE LENGTH=455 |
| ATCG00120.1 | 1,20818624 | 0,00216915 | ATPA ATP synthase subunit alpha chrc:9938-11461 REVERSE LENGTH=507 |
| AT2G44160.1 | 1,486357734 | 0,002193838 | MTHFR2 methylenetetrahydrofolate reductase 2 chr2:18262301-18265185 FORWARD LENGTH=594 |
| AT4G12730.1 | 0,664097752 | 0,00224888 | FLA2 FASCICLIN-like arabinogalactan 2 chr4:7491598-7492809 REVERSE LENGTH=403 |
| AT2G47470.1 | 1,421157392 | 0,002310605 | UNE5_ MEE30_ ATPDI11_ PDI11_ ATPDIL2-1 PROTEIN DISULFIDE ISOMERASE 11_ PDI-LIKE 2-1_ ARA |
| AT2G29560.1 | 1,490214301 | 0,002334741 | ENOC_ ENO3 cytosolic enolase_ enolase 3 chr2:12646635-12649694 FORWARD LENGTH=475 |
| AT4G38630.1 | 2,596819948 | 0,002342238 | MBP1_ RPN10_ ATMCB1_ MCB1 MULTIUBIQUITIN CHAIN BINDING PROTEIN 1_ regulatory particle non- |
| AT3G55040.1 | 1,402337245 | 0,002356873 | GSTL2 glutathione transferase lambda 2 chr3:20398718-20400305 REVERSE LENGTH=292 |
| AT1G18500.1 | 1,300935967 | 0,00237801 | MAML-4_ IPMS1 methylthioalkylmalate synthase-like 4_ ISOPROPYLMALATE SYNTHASE 1 chr1:6369347-63 |
| AT4G35830.1 | 1,33997213 | 0,00239788 | ACO1 aconitase 1 chr4:16973007-16977949 REVERSE LENGTH=898 |
| AT3G48730.1 | 1,347905859 | 0,00243174 | GSAM_ GSA2 glutamate-l-semialdehyde aminomutase_ "glutamate-1-semialdehyde 2_1-aminomutase 2" chr3:180 |
| AT4G29010.1 | 1,267228199 | 0,002439959 | AIM1 ABNORMAL INFLORESCENCE MERISTEM chr4:14297312-14302016 REVERSE LENGTH=721 |
| AT4G33090.1 | 1,47595004 | 0,002464137 | APM1_ ATAPM1 AMINOPEPTIDASE M1_ aminopeptidase M1 chr4:15965915-15970418 REVERSE LENGTH |
| AT2G32120.1 | 0,401778878 | 0,002470657 | HSP70T-2 heat-shock protein 70T-2 chr2:13651720-13653411 REVERSE LENGTH=563 |
| AT3G63460.2 | 1,675053658 | 0,002505223 | SEC31B chr3:23431009-23437241 REVERSE LENGTH=1102 |
| AT1G03220.1 | 1,297837941 | 0,002590686 | SAP2 secreted aspartic protease 2 chr1:787143-788444 FORWARD LENGTH=433 |
| AT5G38480.1 | 1,195751557 | 0,002592946 | GRF3_ RCI1 general regulatory factor 3 chr5:15410277-15411285 FORWARD LENGTH=255 |
| AT1G47128.1 | 0,768754949 | 0,002640409 | RD21_ RD21A responsive to dehydration 21A_ responsive to dehydration 21 chr1:17283139-17285609 REVERSE |
| AT1G35720.1 | 0,837460273 | 0,002652456 | ANNAT1_ ATOXY5_ OXY5_ ANN1_ AtANN1 annexin 1 chr1:13225304-13226939 FORWARD LENGTH=317 |
| AT5G26570.1 | 1,52894268 | 0,00266351 | ATGWD3_ OK1_ PWD PHOSPHOGLUCAN WATER DIKINASE chr5:9261580-9267526 FORWARD LENGT |
| AT4G27440.1 | 0,639783811 | 0,002699899 | PORB protochlorophyllide oxidoreductase B chr4:13725648-13727107 FORWARD LENGTH=401 |
| AT1G77510.1 | 1,807340247 | 0,002721392 | ATPDI6_ PDIL1-2_ ATPDIL1-2_ PDI6 PDI-like 1-2_ PROTEIN DISULFIDE ISOMERASE 6 chr1:29126742-29 |
| AT3G50820.1 | 0,889289149 | 0,002726571 | OEC33_ PSBO-2_ PSBO2 photosystem II subunit O-2_ OXYGEN EVOLVING COMPLEX SUBUNIT 33 KDA_ |
| AT1G13440.2 | 0,860800418 | 0,002760175 | GAPC2_ GAPC-2 glyceraldehyde-3-phosphate dehydrogenase C2_ GLYCERALDEHYDE-3-PHOSPHATE DEHY |
| AT3G32980.1 | 0,450336048 | 0,002784354 | PRX32 Peroxidase 32 chr3:13526404-13529949 REVERSE LENGTH=352 |
| AT4G34450.1 | 1,691611854 | 0,002786642 | gamma2-COP gamma2 Coat Protein chr4:16471956-16476795 FORWARD LENGTH=886 |
| AT5G30510.1 | 1,299724224 | 0,002795835 | ARRPS1_ RPS1_ PRPS1 plastid ribosomal protein S1_ ribosomal protein S1 chr5:11619262-11621223 REVERSE |
| AT3G53580.1 | 1,220022216 | 0,002799158 | no symbol available no full name available chr3:19864784-19866907 FORWARD LENGTH=362 |
| AT2G20610.1 | 1,259947077 | 0,002822512 | RTY1_ RTY_ HLS3_ ALF1_ SUR1 ABERRANT LATERAL ROOT FORMATION 1_ SUPERROOT 1_ ROOTY |
| AT2G26080.1 | 1,522365297 | 0,00283422 | GLDP2_ AtGLDP2 glycine decarboxylase P-protein 2 chr2:11109330-11113786 REVERSE LENGTH=1044 |
| AT4G02770.1 | 1,223699447 | 0,002837138 | PSAD1_ PSAD-1 photosystem I subunit D-1 chr4:1229247-1229873 REVERSE LENGTH=208 |
| AT5G61410.1 | 1,323151201 | 0,002845587 | EMB2728_ RPE D-ribulose-5-phosphate-3-epimerase_ EMBRYO DEFECTIVE 2728 chr5:24684085-24685836 R |
| AT3G52500.1 | 0,684088031 | 0,002857545 | no symbol available no full name available chr3:19465644-19467053 REVERSE LENGTH=469 |
| AT3G23810.1 | 1,272563628 | 0,002876501 | ATSAHH2_ SAHH2 S-ADENOSYL-L-HOMOCYSTEINE (SAH) HYDROLASE 2_ S-adenosyl-l-homocysteine ( |
| AT1G74070.2 | 1,564447529 | 0,002884066 | no symbol available no full name available chr1:27851940-27852861 REVERSE LENGTH=280 |
| AT5G55070.1 | 0,779316832 | 0,002891332 | E2-OGDH2 chr5:22347637-22350409 FORWARD LENGTH=464 |
| AT5G58070.1 | 1,439426847 | 0,002893467 | TIL_ ATTIL TEMPERATURE-INDUCED LIPOCALIN_ temperature-induced lipocalin chr5:23500512-23501156 |
| AT5G23250.1 | 0,737485104 | 0,002896618 | no symbol available no full name available chr5:7830460-7832491 FORWARD LENGTH=341 |

| | | | |
|---|---|---|---|
| AT2G38750.2 | 0,487698337 | 0,002912688 | AtANN4_ ANNAT4 annexin 4 chr2:16196582-16198273 REVERSE LENGTH=302 |
| AT5G56030.1 | 2,040109517 | 0,002915567 | HSP81-2_ HSP90.2_ AtHsp90.2_ ERD8_ HSP81.2 EARLY-RESPONSIVE TO DEHYDRATION 8_ heat shock p[illegible] |
| AT3G47520.1 | 0,841977457 | 0,002916428 | MDH_ pNAD-MDH plastidic NAD-dependent malate dehydrogenase_ malate dehydrogenase chr3:17513657-1751[illegible] |
| AT3G60820.1 | 1,21275677 | 0,002943311 | PBF1 chr3:22472038-22473809 REVERSE LENGTH=223 |
| AT5G17920.1 | 1,182591638 | 0,002994443 | ATCIMS_ METS1_ ATMETS_ ATMS1 COBALAMIN-INDEPENDENT METHIONINE SYNTHASE_ methioni[illegible] |
| AT2G09990.1 | 2,850278193 | 0,003020917 | no symbol available no full name available chr2:3781442-3781882 FORWARD LENGTH=146 |
| AT5G12950.1 | 1,513202297 | 0,003025717 | no symbol available no full name available chr5:4093117-4096806 FORWARD LENGTH=861 |
| AT1G32080.1 | 0,616356053 | 0,003031252 | AtLrgB_ LrgB_ PLGG_ PLGG1 chr1:11537572-11539756 REVERSE LENGTH=512 |
| AT1G07890.1 | 1,192567195 | 0,003055468 | MEE6_ ATAPX01_ ATAPX1_ CS1_ APX1 ascorbate peroxidase 1_ maternal effect embryo arrest 6 chr1:2438005 |
| AT1G67430.2 | 0,558558162 | 0,003090913 | no symbol available no full name available chr1:25262209-25263627 FORWARD LENGTH=131 |
| AT1G52400.1 | 0,532196553 | 0,003091781 | BGL1_ ATBG1_ BGLU18 A. THALIANA BETA-GLUCOSIDASE 1_ BETA-GLUCOSIDASE HOMOLOG 1_ b |
| AT1G08200.1 | 1,195514026 | 0,003110404 | AXS2 UDP-D-apiose/UDP-D-xylose synthase 2 chr1:2574259-2576609 REVERSE LENGTH=389 |
| AT3G25770.1 | 0,433464993 | 0,003182034 | AOC2 allene oxide cyclase 2 chr3:9406975-9407839 FORWARD LENGTH=253 |
| AT2G21390.1 | 1,562224585 | 0,003198298 | no symbol available no full name available chr2:9152428-9156577 FORWARD LENGTH=1218 |
| AT1G42970.1 | 1,144920121 | 0,003200068 | GAPB glyceraldehyde-3-phosphate dehydrogenase B subunit chr1:16127552-16129584 FORWARD LENGTH=44[illegible] |
| AT3G49720.1 | 0,36217976 | 0,00324522 | CGR2 chr3:18440192-18441655 REVERSE LENGTH=261 |
| AT3G18780.1 | 0,63008519 | 0,003262496 | LSR2_ ACT2_ ENL2_ DER1_ FIZ2 FRIZZY AND KINKED SHOOTS 2_ LIGHT STRESS-REGULATED 2_ DE |
| AT1G03130.1 | 0,794725243 | 0,003311787 | PSAD-2 photosystem I subunit D-2 chr1:753528-754142 REVERSE LENGTH=204 |
| AT5G49910.1 | 1,685560144 | 0,003340537 | HSC70-7_ cpHsc70-2 chloroplast heat shock protein 70-2_ HEAT SHOCK PROTEIN 70-7 chr5:20303470-203062 |
| AT1G66670.1 | 1,251347262 | 0,003366827 | NCLPP3_ CLPP3 CLP protease proteolytic subunit 3 chr1:24863995-24865646 REVERSE LENGTH=309 |
| AT4G23600.1 | 0,516433467 | 0,003371578 | CORI3_ JR2 JASMONIC ACID RESPONSIVE 2_ CORONATINE INDUCED 1 chr4:12310657-12312885 FORW |
| AT3G25520.2 | 0,67363655 | 0,003379966 | RPL5A_ ATL5_ PGY3_ OLI5 ribosomal protein L5_ RIBOSOMAL PROTEIN L5 A_ PIGGYBACK3_ OLIGOC[illegible] |
| AT2G42170.1 | 2,427654135 | 0,003381716 | no symbol available no full name available chr2:17578683-17580222 FORWARD LENGTH=329 |
| AT2G30950.1 | 3,344664432 | 0,003389589 | FTSH2_ VAR2 VARIEGATED 2 chr2:13174692-13177064 FORWARD LENGTH=695 |
| AT2G42600.1 | 1,201009567 | 0,003439281 | ATPPC2_ PPC2 phosphoenolpyruvate carboxylase 2 chr2:17734541-17738679 REVERSE LENGTH=963 |
| AT4G04640.1 | 1,132618688 | 0,003506898 | ATPC1 chr4:2350761-2351882 REVERSE LENGTH=373 |
| AT1G15930.1 | 1,182744991 | 0,003512083 | no symbol available no full name available chr1:5471702-5472741 FORWARD LENGTH=144 |
| AT4G20360.1 | 1,182660241 | 0,003524616 | SVR11_ RAB8D_ ATRAB8D_ ATRABE1B_ RABE1b RAB GTPase homolog E1B_ SUPPRESSOR OF VARIEC |
| AT5G50950.1 | 0,379319989 | 0,003534539 | FUM2 FUMARASE 2 chr5:20729687-20733476 FORWARD LENGTH=510 |
| AT3G52380.1 | 1,337235489 | 0,003567902 | PDE322_ CP33 PIGMENT DEFECTIVE 322_ chloroplast RNA-binding protein 33 chr3:19421619-19422855 FOF |
| AT2G27730.1 | 2,866044387 | 0,003574692 | no symbol available no full name available chr2:11820056-11820867 REVERSE LENGTH=113 |
| AT5G56000.1 | 1,383449346 | 0,003590144 | Hsp81.4_ AtHsp90.4 HEAT SHOCK PROTEIN 90.4_ HEAT SHOCK PROTEIN 81.4 chr5:22677602-22680067 F |
| AT5G66680.1 | 1,388098318 | 0,003632126 | DGL1 DEFECTIVE GLYCOSYLATION chr5:26617840-26620581 REVERSE LENGTH=437 |
| AT3G04770.1 | 1,353518152 | 0,003684239 | RPSAb 40s ribosomal protein SA B chr3:1309465-1310846 REVERSE LENGTH=332 |
| AT5G10540.1 | 1,651480683 | 0,003717061 | TOP2 thimet metalloendopeptidase 2 chr5:3328119-3332462 FORWARD LENGTH=701 |
| AT1G07790.1 | 0,81279448 | 0,003724082 | HTB1 chr1:2413049-2413495 FORWARD LENGTH=148 |
| AT3G04840.1 | 0,86128079 | 0,003745033 | no symbol available no full name available chr3:1329751-1331418 FORWARD LENGTH=262 |
| AT2G37040.1 | 3,390800438 | 0,003775717 | PAL1_ ATPAL1 PHE ammonia lyase 1 chr2:15557602-15560237 REVERSE LENGTH=725 |
| ATCG00270.1 | 0,886928997 | 0,003869674 | PSBD photosystem II reaction center protein D chrc:32711-33772 FORWARD LENGTH=353 |

| | | | |
|---|---|---|---|
| AT5G09590.1 | 1,49191956 | 0,003870322 | MTHSC70-2_ HSC70-5 mitochondrial HSO70 2_ HEAT SHOCK COGNATE chr5:2975721-2978508 FORWARD |
| AT4G24770.1 | 1,290856349 | 0,003871395 | ATRBP33_ ATRBP31_ CP31A_ RBP31_ CP33a_ CP31 "ARABIDOPSIS THALIANA RNA BINDING PROTEIN |
| AT5G60670.1 | 0,593044671 | 0,003892361 | RPL12C Ribosomal Protein Like 12C chr5:24381066-24381566 REVERSE LENGTH=166 |
| AT3G18890.1 | 1,251349611 | 0,00392842 | Tic62_ AtTic62 translocon at the inner envelope membrane of chloroplasts 62 chr3:6511169-6514729 FORWARD |
| AT1G09310.1 | 0,650541987 | 0,003978537 | SVB2_ SVBL SVB-like chr1:3009109-3009648 FORWARD LENGTH=179 |
| AT1G21750.1 | 1,296980507 | 0,004039381 | PDIL1-1_ ATPDI5_ PDI5_ ATPDIL1-1 PDI-like 1-1_ ARABIDOPSIS THALIANA PROTEIN DISULFIDE ISOM |
| AT1G06410.1 | 1,6171437 | 0,004077838 | ATTPSA_ TPS7_ ATTPS7 TREHALOSE -6-PHOSPHATASE SYNTHASE S7_ trehalose-phosphatase/synthase 7 |
| AT1G79230.3 | 1,162160569 | 0,004155439 | ATMST1_ STR1_ MST1_ ATRDH1_ ST1 ARABIDOPSIS THALIANA RHODANESE HOMOLOGUE 1_ merca |
| AT3G54900.1 | 2,075794414 | 0,004174868 | AtGRXS14_ ATGRXCP_ CXIP1 GLUTAREDOXIN_ CAX interacting protein 1 chr3:20341850-20342371 REVE |
| ATCG00660.1 | 0,20006163 | 0,004175022 | RPL20 ribosomal protein L20 chrc:68512-68865 REVERSE LENGTH=117 |
| AT1G04420.1 | 1,952616651 | 0,00419764 | no symbol available no full name available chr1:1191634-1193699 FORWARD LENGTH=412 |
| AT1G20260.1 | 1,268545658 | 0,004229221 | AtVAB3_ VAB3 V-ATPase B subunit 3 chr1:7016971-7020290 FORWARD LENGTH=487 |
| AT3G57290.1 | 1,960119526 | 0,004247597 | EIF3E_ ATINT6_ TIF3E1_ INT6_ INT-6_ ATEIF3E-1 eukaryotic translation initiation factor 3E chr3:21196786-2 |
| AT4G29130.1 | 1,326760006 | 0,004280546 | GIN2_ HXK1_ ATHXK1 GLUCOSE INSENSITIVE 2_ hexokinase 1_ ARABIDOPSIS THALIANA HEXOKINA |
| AT5G07440.3 | 1,401954972 | 0,004282746 | GDH2 glutamate dehydrogenase 2 chr5:2356153-2357546 FORWARD LENGTH=309 |
| AT3G55610.1 | 2,460853103 | 0,004300962 | P5CS2 delta 1-pyrroline-5-carboxylate synthase 2 chr3:20624278-20628989 REVERSE LENGTH=726 |
| AT5G59880.2 | 0,619378732 | 0,004307872 | ADF3 actin depolymerizing factor 3 chr5:24120382-24121628 FORWARD LENGTH=124 |
| AT4G34120.1 | 1,247478969 | 0,004341104 | CDCP1_ CBSX2_ LEJ1 CBS domain containing protein 2_ LOSS OF THE TIMING OF ET AND JA BIOSYNTH |
| AT5G01530.1 | 0,841134984 | 0,004348487 | LHCB4.1 light harvesting complex photosystem II chr5:209084-210243 FORWARD LENGTH=290 |
| AT3G09200.2 | 1,327988789 | 0,004349013 | no symbol available no full name available chr3:2823364-2825020 REVERSE LENGTH=287 |
| AT5G12250.1 | 0,739920259 | 0,004397865 | TUB6 beta-6 tubulin chr5:3961317-3962971 REVERSE LENGTH=449 |
| AT1G72150.1 | 0,807638082 | 0,004404035 | PATL1 PATELLIN 1 chr1:27148558-27150652 FORWARD LENGTH=573 |
| AT5G20950.1 | 1,494083952 | 0,004405166 | BGLC1 chr5:7107609-7110775 REVERSE LENGTH=624 |
| AT5G48580.1 | 0,579655935 | 0,004433268 | FKBP15-2 FK506- and rapamycin-binding protein 15 kD-2 chr5:19696156-19697304 REVERSE LENGTH=163 |
| AT1G29900.1 | 1,651939116 | 0,004441963 | CARB_ VEN3 carbamoyl phosphate synthetase B_ VENOSA 3 chr1:10468164-10471976 FORWARD LENGTH= |
| AT3G15730.1 | 1,513749552 | 0,004442111 | PLD_ PLDALPHA1 phospholipase D alpha 1 chr3:5330835-5333474 FORWARD LENGTH=810 |
| AT5G26780.1 | 0,825116633 | 0,004474684 | SHM2 serine hydroxymethyltransferase 2 chr5:9418299-9421725 FORWARD LENGTH=517 |
| AT1G30580.1 | 1,268139417 | 0,004581779 | EngD-1 chr1:10831953-10835454 REVERSE LENGTH=394 |
| AT1G52040.1 | 0,322978926 | 0,004596404 | MBP1_ ATMBP myrosinase-binding protein 1 chr1:19350595-19352578 REVERSE LENGTH=462 |
| AT3G24503.1 | 1,520185255 | 0,004708326 | REF1_ ALDH2C4_ ALDH1A REDUCED EPIDERMAL FLUORESCENCE1_ aldehyde dehydrogenase 2C4_ alde |
| AT2G45810.1 | 1,424702848 | 0,004717149 | RH6 RNA Helicase 6 chr2:18859836-18862318 FORWARD LENGTH=528 |
| AT4G01690.1 | 1,317090967 | 0,004727807 | PPO1_ PPOX_ HEMG1 chr4:729929-732309 FORWARD LENGTH=537 |
| AT1G78870.4 | 1,168298228 | 0,004731765 | UBC13A_ UBC35 ubiquitin-conjugating enzyme 35_ UBIQUITIN CONJUGATING ENZYME 13A chr1:2965096 |
| AT2G46820.1 | 0,690285087 | 0,004760833 | PTAC8_ PSI-P_ PSAP_ CURT1B_ TMP14 CURVATURE THYLAKOID 1B_ PLASTID TRANSCRIPTIONALL |
| AT1G41830.1 | 0,238134299 | 0,004772105 | SKS6 SKU5-similar 6_ SKU5 SIMILAR 6 chr1:15603892-15607802 REVERSE LENGTH=542 |
| AT2G45470.1 | 0,672959117 | 0,004773771 | AGP8_ FLA8 ARABINOGALACTAN PROTEIN 8_ FASCICLIN-like arabinogalactan protein 8 chr2:18742797-1 |
| AT5G43780.1 | 0,683446027 | 0,00490675 | ATPS4_ APS4 chr5:17589631-17591480 REVERSE LENGTH=469 |
| AT1G80480.1 | 2,015799683 | 0,004932507 | PTAC17 plastid transcriptionally active 17 chr1:30258272-30260570 REVERSE LENGTH=444 |
| AT2G32920.1 | 1,561241507 | 0,004968974 | PDIL2-3_ PDI9_ ATPDIL2-3_ ATPDI9 ARABIDOPSIS THALIANA PROTEIN DISULFIDE ISOMERASE 9_ P |

| | | | |
|---|---|---|---|
| AT4G26970.1 | 1,541062605 | 0,005057618 | ACO2 aconitase 2 chr4:13543077-13548427 FORWARD LENGTH=995 |
| AT5G61170.1 | 1,418694238 | 0,005075353 | no symbol available no full name available chr5:24611158-24612202 FORWARD LENGTH=143 |
| AT1G75940.1 | 0,7242157 | 0,00508496 | ATA27_ BGLU20 BETA GLUCOSIDASE 20 chr1:28511198-28514044 FORWARD LENGTH=535 |
| AT3G11510.1 | 0,855539642 | 0,005086754 | no symbol available no full name available chr3:3623757-3624866 REVERSE LENGTH=150 |
| AT1G48920.1 | 1,546063569 | 0,005139436 | PARL1_ NUC-L1_ ATNUC-L1_ NUC1 nucleolin like 1_ PARALLEL 1_ nucleolin 1 chr1:18098186-18101422 FC |
| AT2G39330.1 | 0,647997475 | 0,005161042 | JAL23 jacalin-related lectin 23 chr2:16419787-16421573 REVERSE LENGTH=459 |
| AT5G52640.1 | 2,072207041 | 0,005214717 | HSP81-1_ ATHSP90.1_ HSP81.1_ AtHsp90-1_ HSP83_ ATHS83_ HSP90.1 HEAT SHOCK PROTEIN 90-1_ hea |
| AT3G29320.1 | 1,330290525 | 0,005272671 | PHS1 alpha-glucan phosphorylase 1 chr3:11252871-11257587 FORWARD LENGTH=962 |
| AT3G43980.1 | 1,469344498 | 0,005308697 | no symbol available no full name available chr3:15778555-15779235 REVERSE LENGTH=56 |
| AT3G04120.1 | 1,178160378 | 0,005343603 | GAPC1_ GAPC_ GAPC-1 GLYCERALDEHYDE-3-PHOSPHATE DEHYDROGENASE C SUBUNIT_ glycerald |
| AT3G53610.1 | 0,390135137 | 0,005392367 | RAB8B_ RAB8_ AtRab8B_ ATRAB8_ AtRABE1a RAB GTPase homolog 8 chr3:19876531-19878264 REVERSE |
| AT2G01290.1 | 1,383783977 | 0,005402247 | RPI2 ribose-5-phosphate isomerase 2 chr2:149192-149989 REVERSE LENGTH=265 |
| AT4G13930.1 | 1,195661796 | 0,005492801 | SHM4 serine hydroxymethyltransferase 4 chr4:8048013-8050021 REVERSE LENGTH=471 |
| AT1G69740.1 | 1,177160265 | 0,005513522 | HEMB1_ ALAD1 5-aminolevulinic acid dehydratase 1 chr1:26232197-26234713 FORWARD LENGTH=430 |
| AT3G17820.1 | 5,498506098 | 0,005513706 | GLN1.3_ ATGSKB6_ GLN1;3 glutamine synthetase 1.3_ ARABIDOPSIS THALIANA GLUTAMINE SYNTHAS |
| AT1G49630.1 | 1,407715639 | 0,005539082 | PREP2_ ATPREP2 presequence protease 2 chr1:18368405-18375336 REVERSE LENGTH=1080 |
| AT5G48180.1 | 1,436303913 | 0,005542697 | NSP5_ AtNSP5 nitrile specifier protein 5 chr5:19541283-19542358 REVERSE LENGTH=326 |
| AT3G02560.1 | 0,697391741 | 0,005571251 | no symbol available no full name available chr3:542341-543168 FORWARD LENGTH=191 |
| AT2G05840.3 | 1,39336359 | 0,005597313 | PAA2 20S proteasome subunit PAA2 chr2:2234226-2235533 FORWARD LENGTH=205 |
| ATCG00905.1 | 0,627461428 | 0,005654573 | RPS12_ RPS12C RIBOSOMAL PROTEIN S12_ ribosomal protein S12C chrc:97999-98793 REVERSE LENGTH |
| AT5G19940.1 | 0,774078239 | 0,005695052 | FBN6_ PAP8 FIBRILLIN6_ Probable Plastid-Lipid Associated Protein chr5:6739693-6740661 FORWARD LENG |
| AT1G09430.1 | 1,512895844 | 0,005722716 | ACLA-3 ATP-citrate lyase A-3 chr1:3042135-3044978 FORWARD LENGTH=424 |
| AT4G01050.1 | 0,893057588 | 0,005748336 | TROL thylakoid rhodanese-like chr4:455874-458175 FORWARD LENGTH=466 |
| AT1G74920.2 | 1,544950735 | 0,005784155 | ALDH10A8 aldehyde dehydrogenase 10A8 chr1:28139175-28142573 REVERSE LENGTH=496 |
| AT1G42960.1 | 1,772149886 | 0,005853596 | CTI1 Carboxyltransferase Interactor1 chr1:16125863-16127080 FORWARD LENGTH=168 |
| AT1G50670.1 | 1,382635933 | 0,005876874 | OTU2 ovarian tumor domain (OTU)-containing DUB (deubiquitilating enzyme) 2 chr1:18775086-18776552 REVE |
| AT3G10350.1 | 0,822004338 | 0,005951819 | AtGET3b_ GET3b Guided Entry of Tail-anchored proteins 3b chr3:3208310-3210678 FORWARD LENGTH=411 |
| AT4G26900.1 | 1,604967174 | 0,005983438 | AT-HF_ HISN4 HIS HF chr4:13515514-13519608 FORWARD LENGTH=592 |
| AT5G26000.1 | 0,80615409 | 0,005995168 | TGG1_ AtTGG1_ BGLU38 thioglucoside glucohydrolase 1_ BETA GLUCOSIDASE 38 chr5:9079678-9082347 R |
| AT1G11910.1 | 1,276023548 | 0,005996014 | ATAPA1_ AtPaspA1_ PaspA1_ APA1 aspartic proteinase A1_ putative aspartic proteinase A1 chr1:4017119-4019 |
| AT3G01280.1 | 0,839551471 | 0,005999059 | VDAC1_ ATVDAC1 ARABIDOPSIS THALIANA VOLTAGE DEPENDENT ANION CHANNEL 1_ voltage dep |
| AT1G49760.1 | 1,124357817 | 0,006018575 | PABP8_ PAB8 POLY(A) BINDING PROTEIN 8_ poly(A) binding protein 8 chr1:18416740-18419753 FORWAR |
| AT4G18440.1 | 0,680208607 | 0,006030617 | no symbol available no full name available chr4:10186385-10188832 REVERSE LENGTH=536 |
| AT5G34850.1 | 0,780274279 | 0,006047712 | PUP3_ PAP26_ ATPAP26 purple acid phosphatase 26_ phosphatase-under producer 3_ PURPLE ACID PHOSPHA |
| AT4G02530.1 | 1,216913175 | 0,006076872 | MPH2 MAINTENANCE OF PHOTOSYSTEM II UNDER HIGH LIGHT 2 chr4:1112335-1114005 REVERSE LE |
| AT1G11650.1 | 1,513000178 | 0,006099497 | RBP45B_ ATRBP45B chr1:3914895-3917301 FORWARD LENGTH=306 |
| AT5G55480.1 | 1,473938161 | 0,006135718 | GPDL1_ GDPDL4_ SVL1 SHV3-like 1_ Glycerophosphodiester phosphodiesterase (GDPD) like 4_ glycerophosph |
| AT5G09900.1 | 1,248932654 | 0,006166547 | EMB2107_ MSA_ RPN5A MARIPOSA_ EMBRYO DEFECTIVE 2107_ REGULATORY PARTICLE NON-ATP |
| AT5G20920.2 | 0,731311667 | 0,006200799 | EIF2 BETA_ EMB1401_ eIF-2bs embryo defective 1401_ eukaryotic translation initiation factor 2 beta subunit chr |

| | | | |
|---|---|---|---|
| AT1G08830.1 | 0,43013079 | 0,006263573 | CSD1_ AtSOD1_ SOD1 superoxide dismutase 1_ copper/zinc superoxide dismutase 1 chr1:2827700-2829053 FOR |
| AT5G47700.1 | 1,52023039 | 0,006323223 | RPP1C_ RPP1.3 60S acidic ribosomal protein P1-3_ RPP1 co-orthologous gene 3 chr5:19328019-19328724 REVE |
| AT1G74100.1 | 0,654681202 | 0,006370942 | ATSOT16_ ATST5A_ CORI-7_ SOT16 sulfotransferase 16_ CORONATINE INDUCED-7_ SULFOTRANSFERA |
| AT1G09180.1 | 0,902031944 | 0,006412612 | SAR1A_ ATSARA1A_ SARA1A_ ATSAR1 SECRETION-ASSOCIATED RAS 1_ secretion-associated RAS supe |
| AT1G10840.1 | 1,615296746 | 0,006431284 | TIF3H1 translation initiation factor 3 subunit H1 chr1:3607885-3610299 REVERSE LENGTH=337 |
| AT1G30120.1 | 0,649461986 | 0,006448841 | PDH-E1 BETA pyruvate dehydrogenase E1 beta chr1:10584350-10586477 REVERSE LENGTH=406 |
| AT5G36880.4 | 1,286984252 | 0,006494044 | ACS acetyl-CoA synthetase chr5:14535506-14539084 REVERSE LENGTH=610 |
| AT3G54050.1 | 1,19943025 | 0,006503041 | HCEF1_ cfbp1 high cyclic electron flow 1 chr3:20016951-20018527 FORWARD LENGTH=417 |
| AT1G53240.1 | 1,105269749 | 0,006564176 | mMDH1 mitochondrial malate dehydrogenase 1 chr1:19854966-19856802 REVERSE LENGTH=341 |
| AT2G31610.1 | 1,305665264 | 0,006638778 | no symbol available no full name available chr2:13450384-13451669 FORWARD LENGTH=250 |
| AT5G45390.1 | 1,402414849 | 0,006769489 | CLPP4_ NCLPP4 NUCLEAR-ENCODED CLP PROTEASE P4_ CLP protease P4 chr5:18396351-18397586 FOR |
| AT4G02080.1 | 0,769687038 | 0,006782484 | ATSARA1C_ SAR2_ SAR1C_ ATSAR2_ ASAR1 secretion-associated RAS super family 2 chr4:921554-922547 F |
| AT1G09750.1 | 0,749579144 | 0,006785037 | no symbol available no full name available chr1:3157541-3158960 FORWARD LENGTH=449 |
| AT4G37925.1 | 0,759415394 | 0,006794807 | NDH-M_ NdhM subunit NDH-M of NAD(P)H:plastoquinone dehydrogenase complex_ NADH dehydrogenase-like |
| AT2G12550.1 | 0,608646925 | 0,00681002 | NUB1 homolog of human NUB1 chr2:5114881-5118486 FORWARD LENGTH=562 |
| AT5G27470.1 | 1,568488778 | 0,006831845 | no symbol available no full name available chr5:9695087-9697154 FORWARD LENGTH=451 |
| AT1G48630.1 | 1,326038341 | 0,006858602 | RACK1B_ RACK1B_AT receptor for activated C kinase 1B chr1:17981977-17983268 REVERSE LENGTH=326 |
| AT4G16143.1 | 1,770785921 | 0,006882476 | IMPA-2 importin alpha isoform 2 chr4:9134450-9137134 REVERSE LENGTH=535 |
| AT5G48130.1 | 1,339774858 | 0,007006602 | no symbol available no full name available chr5:19516291-19518450 FORWARD LENGTH=625 |
| AT5G16050.1 | 1,218886324 | 0,007047155 | GRF5_ GF14 UPSILON general regulatory factor 5 chr5:5244008-5245402 REVERSE LENGTH=268 |
| AT3G26650.1 | 1,143515155 | 0,007146942 | GAPA_ GAPA1_ GAPA-1 glyceraldehyde 3-phosphate dehydrogenase A subunit_ GLYCERALDEHYDE 3-PHOS |
| AT2G36530.1 | 1,136846892 | 0,007302368 | LOS2_ ENO2 LOW EXPRESSION OF OSMOTICALLY RESPONSIVE GENES 2_ enolase 2 chr2:15321081-153 |
| AT5G08050.1 | 0,9026401 | 0,00731652 | RIQ1 chr5:2578418-2578964 FORWARD LENGTH=158 |
| AT2G36580.1 | 1,572521917 | 0,007349763 | no symbol available no full name available chr2:15339253-15342781 FORWARD LENGTH=527 |
| AT1G09795.1 | 1,476514461 | 0,007367307 | ATATP-PRT2_ HISN1B_ ATP-PRT2 ATP phosphoribosyl transferase 2 chr1:3173588-3176690 FORWARD LEN |
| AT3G14290.1 | 1,240913089 | 0,007371105 | PAE2 20S proteasome alpha subunit E2 chr3:4764364-4766381 FORWARD LENGTH=237 |
| AT5G08670.1 | 1,192532593 | 0,007518678 | no symbol available no full name available chr5:2818395-2821149 REVERSE LENGTH=556 |
| AT5G12040.1 | 1,166279796 | 0,007566102 | no symbol available no full name available chr5:3885162-3887772 FORWARD LENGTH=369 |
| AT2G21410.1 | 1,620272753 | 0,007724999 | VHA-A2 vacuolar proton ATPase A2 chr2:9162703-9168141 FORWARD LENGTH=821 |
| AT1G03680.1 | 1,26307321 | 0,007880253 | ATHM1_ TRX-M1_ ATM1_ THM1 thioredoxin M-type 1_ THIOREDOXIN M-TYPE 1_ ARABIDOPSIS THIOR |
| AT3G22460.2 | 0,780508163 | 0,007889753 | OASA2 O-acetylserine (thiol) lyase (OAS-TL) isoform A2 chr3:7963855-7964769 FORWARD LENGTH=188 |
| AT4G02930.1 | 1,189545698 | 0,007929913 | no symbol available no full name available chr4:1295751-1298354 REVERSE LENGTH=454 |
| AT4G23850.1 | 1,461942532 | 0,008013546 | LACS4 long-chain acyl-CoA synthetase 4 chr4:12403720-12408263 REVERSE LENGTH=666 |
| AT3G15060.1 | 2,744488426 | 0,008104755 | RABA1g_ AtRABA1g RAB GTPase homolog A1G chr3:5069239-5070025 FORWARD LENGTH=217 |
| AT3G22960.1 | 1,307314982 | 0,008259703 | PKP1_ PKP-ALPHA PLASTIDIAL PYRUVATE KINASE 1 chr3:8139369-8141771 FORWARD LENGTH=596 |
| AT1G23410.1 | 1,282915715 | 0,008411553 | RPS27aA chr1:8314940-8315410 FORWARD LENGTH=156 |
| AT5G46800.1 | 1,090883342 | 0,008428743 | BOU A BOUT DE SOUFFLE chr5:18988779-18989810 REVERSE LENGTH=300 |
| AT4G01800.1 | 1,603615104 | 0,008443289 | SECA1_ AGY1_ AtcpSecA Arabidopsis thaliana chloroplast SecA_ Albino or Glassy Yellow 1 chr4:770926-77613 |
| AT5G09500.1 | 0,676441584 | 0,008482881 | no symbol available no full name available chr5:2954044-2954850 REVERSE LENGTH=150 |

| | | | |
|---|---|---|---|
| AT1G19920.1 | 1,363182936 | 0,008597487 | APS2_ ATPS2_ ASA1 ATP SULFURYLASE ARABIDOPSIS 1 chr1:6914835-6916657 REVERSE LENGTH=47 |
| AT3G46230.1 | 2,376483309 | 0,00863485 | HSP17.4_ ATHSP17.4 heat shock protein 17.4_ ARABIDOPSIS THALIANA HEAT SHOCK PROTEIN 17.4 chr3 |
| AT4G37800.1 | 0,808051093 | 0,008699317 | XTH7 xyloglucan endotransglucosylase/hydrolase 7 chr4:17775703-17777372 REVERSE LENGTH=293 |
| AT1G78900.1 | 1,412937582 | 0,008784175 | VHA-A vacuolar ATP synthase subunit A chr1:29660463-29664575 FORWARD LENGTH=623 |
| AT3G15360.1 | 1,153581743 | 0,008920203 | ATM4_ TRX-M4_ ATHM4 ARABIDOPSIS THIOREDOXIN M-TYPE 4_ thioredoxin M-type 4 chr3:5188448-51 |
| AT3G53430.1 | 0,674790655 | 0,00892352 | no symbol available no full name available chr3:19809895-19810395 REVERSE LENGTH=166 |
| AT4G14130.1 | 2,633541798 | 0,008960398 | XTR7_ XTH15 xyloglucan endotransglucosylase/hydrolase 15_ xyloglucan endotransglycosylase 7 chr4:8137161-8 |
| AT1G67700.1 | 0,663383934 | 0,009002742 | HHL1 HYPERSENSITIVE TO HIGH LIGHT 1 chr1:25374295-25375716 FORWARD LENGTH=230 |
| AT3G09350.1 | 2,846444258 | 0,009013097 | Fes1A Fes1A chr3:2871216-2873109 FORWARD LENGTH=363 |
| AT4G30620.1 | 1,402863871 | 0,009188058 | STCL STIC2 Like chr4:14948724-14950035 REVERSE LENGTH=180 |
| AT5G15200.1 | 0,759981432 | 0,009277212 | no symbol available no full name available chr5:4935124-4936334 REVERSE LENGTH=198 |
| AT5G15520.1 | 1,289263992 | 0,009391457 | no symbol available no full name available chr5:5037242-5038136 REVERSE LENGTH=143 |
| AT5G65620.1 | 1,186353714 | 0,009453348 | OOP_ TOP1 thimet metalloendopeptidase 1_ organellar oligopeptidase chr5:26221951-26225784 FORWARD LEN |
| AT2G38270.1 | 1,278829135 | 0,009465554 | ATGRX2_ AtGRXS16_ CXIP2 GLUTAREDOXIN 2_ glutaredoxin 16_ CAX-interacting protein 2 chr2:16031347 |
| AT2G34430.1 | 0,662619902 | 0,009547781 | DEG11_ LHCB1.4_ LHB1B1 light-harvesting chlorophyll-protein complex II subunit B1_ LIGHT-HARVESTING |
| AT2G31810.2 | 1,28788822 | 0,00957385 | AHASS1_ AIP1 acetolactate synthase small subunit 1_ ALS-INTERACTING PROTEIN1 chr2:13524419-1352824 |
| AT4G21960.1 | 0,319646313 | 0,009723966 | PRXR1 chr4:11646613-11648312 REVERSE LENGTH=330 |
| AT5G05010.1 | 1,844271011 | 0,009797314 | no symbol available no full name available chr5:1477137-1479872 FORWARD LENGTH=527 |
| AT1G65980.1 | 1,265885832 | 0,00985979 | TPX1 thioredoxin-dependent peroxidase 1 chr1:24559524-24560753 REVERSE LENGTH=162 |
| AT3G02230.1 | 1,34984726 | 0,009930991 | RGP1_ ATRGP1 ARABIDOPSIS THALIANA REVERSIBLY GLYCOSYLATED POLYPEPTIDE 1_ reversibly |
| AT5G46290.3 | 1,31171947 | 0,010047615 | KASI_ KAS1 3-ketoacyl-acyl carrier protein synthase I_ KETOACYL-ACP SYNTHASE 1 chr5:18774439-187766 |
| AT5G08530.1 | 1,138215128 | 0,010141736 | CI51_ NDUFV1 51 kDa subunit of complex I chr5:2759848-2761726 REVERSE LENGTH=486 |
| ATCG00480.1 | 1,177432682 | 0,010159577 | PB_ ATPB_ CF1beta_ AthCF1beta ATP synthase subunit beta chrc:52660-54156 REVERSE LENGTH=498 |
| AT3G18130.1 | 1,469861343 | 0,010188562 | RACK1C_ RACK1C_AT receptor for activated C kinase 1C chr3:6211109-6212371 REVERSE LENGTH=326 |
| AT4G08390.3 | 1,268023078 | 0,010287938 | SAPX stromal ascorbate peroxidase chr4:5314999-5317071 FORWARD LENGTH=371 |
| AT1G56070.1 | 1,30291982 | 0,010369894 | LOS1 LOW EXPRESSION OF OSMOTICALLY RESPONSIVE GENES 1 chr1:20968245-20971077 REVERSE I |
| AT1G73230.1 | 2,404355619 | 0,010398138 | no symbol available no full name available chr1:27540506-27541364 REVERSE LENGTH=165 |
| AT5G11770.1 | 2,526546912 | 0,010505161 | no symbol available no full name available chr5:3791148-3792929 REVERSE LENGTH=218 |
| AT3G17810.1 | 0,917471356 | 0,01055608 | PYD1 pyrimidine 1 chr3:6094279-6096289 FORWARD LENGTH=426 |
| AT5G48570.1 | 1,912381356 | 0,010565646 | ROF2_ ATFKBP65_ FKBP65 chr5:19690746-19693656 REVERSE LENGTH=578 |
| AT1G20450.1 | 0,570506344 | 0,010601744 | LTI45_ ERD10_ LTI29 EARLY RESPONSIVE TO DEHYDRATION 10_ LOW TEMPERATURE INDUCED 45 |
| AT3G14990.1 | 0,752308919 | 0,010713938 | DJ-1a_ AtDJ1A_ DJ1A DJ-1 homolog A chr3:5047510-5049621 FORWARD LENGTH=392 |
| AT4G38510.1 | 1,184963426 | 0,01085567 | AtVAB2_ VAB2 V-ATPase B subunit 2 chr4:18011155-18014789 REVERSE LENGTH=487 |
| AT4G08900.1 | 0,5430092 | 0,010869218 | ARGAH1 arginine amidohydrolase 1 chr4:5703499-5705180 FORWARD LENGTH=342 |
| AT1G22700.3 | 0,569944068 | 0,010878081 | PYG7 chr1:8028323-8029289 REVERSE LENGTH=211 |
| AT3G22110.1 | 1,277997918 | 0,01089238 | PAC1 20S proteasome alpha subunit C1 chr3:7792819-7793571 REVERSE LENGTH=250 |
| AT5G03290.1 | 1,363643209 | 0,011001628 | IDH-V isocitrate dehydrogenase V chr5:794043-795939 FORWARD LENGTH=374 |
| AT5G41520.1 | 1,841307319 | 0,011006654 | RPS10B ribosomal protein S10e B chr5:16609377-16610583 REVERSE LENGTH=180 |
| AT3G53130.1 | 1,343422016 | 0,011073115 | LUT1_ CYP97C1 CYTOCHROME P450 97C1_ LUTEIN DEFICIENT 1 chr3:19692812-19695278 FORWARD I |

| | | | |
|---|---|---|---|
| AT1G01200.1 | 1,499366368 | 0,011190511 | RABA3_ ATRABA3_ ATRAB-A3 ARABIDOPSIS RAB GTPASE HOMOLOG A3_ RAB GTPase homolog A3 c |
| AT5G57870.1 | 1,373641398 | 0,011235817 | eIFiso4G1 eukaryotic translation Initiation Factor isoform 4G1 chr5:23439755-23443433 FORWARD LENGTH=7 |
| AT2G35040.1 | 1,247251455 | 0,011237245 | no symbol available no full name available chr2:14765347-14768269 REVERSE LENGTH=596 |
| AT1G69830.1 | 1,250035193 | 0,011361342 | ATAMY3_ AMY3 ALPHA-AMYLASE-LIKE 3_ alpha-amylase-like 3 chr1:26288518-26293003 REVERSE LEN |
| AT1G23310.1 | 1,139155491 | 0,011361532 | GGT1_ AOAT1_ GGAT1 GLUTAMATE:GLYOXYLATE AMINOTRANSFERASE 1_ glutamate:glyoxylate amin |
| AT3G23490.1 | 0,623997493 | 0,011420198 | CYN cyanase chr3:8423238-8424415 REVERSE LENGTH=168 |
| AT2G22230.1 | 1,225582431 | 0,011514769 | no symbol available no full name available chr2:9450042-9451427 FORWARD LENGTH=220 |
| AT1G58080.1 | 1,750843671 | 0,011613405 | ATATP-PRT1_ ATP-PRT1_ HISN1A ATP phosphoribosyl transferase 1 chr1:21504562-21507429 REVERSE LE |
| AT3G18190.1 | 2,279040502 | 0,011639417 | CCT4 Chaperonin containing T-complex polypeptide-1 subunit 4 chr3:6232226-6233836 FORWARD LENGTH=5 |
| AT4G34180.1 | 0,378727975 | 0,011672589 | CYCLASE1 CYCLASE1 chr4:16370060-16371383 REVERSE LENGTH=255 |
| AT1G29880.1 | 1,367815136 | 0,011899399 | no symbol available no full name available chr1:10459662-10462781 REVERSE LENGTH=729 |
| AT1G09620.1 | 1,145511406 | 0,011967507 | no symbol available no full name available chr1:3113077-3116455 REVERSE LENGTH=1091 |
| AT5G52650.1 | 1,586870598 | 0,012140721 | no symbol available no full name available chr5:21355781-21357003 REVERSE LENGTH=179 |
| AT5G28840.1 | 1,143562921 | 0,012159459 | GME GDP-D-mannose 3 chr5:10862472-10864024 REVERSE LENGTH=377 |
| AT2G34420.1 | 1,52015703 | 0,012361591 | LHCB1.5_ LHB1B2 PHOTOSYSTEM II LIGHT HARVESTING COMPLEX GENE 1.5_ photosystem II light har |
| ATCG00780.1 | 0,731362078 | 0,012419259 | RPL14 ribosomal protein L14 chrc:80696-81064 REVERSE LENGTH=122 |
| AT4G25080.1 | 1,150206696 | 0,012437407 | CHLM magnesium-protoporphyrin IX methyltransferase chr4:12877015-12878128 FORWARD LENGTH=312 |
| AT2G20550.1 | 0,43957914 | 0,012441716 | no symbol available no full name available chr2:8846051-8847113 REVERSE LENGTH=284 |
| AT1G13060.1 | 1,109558104 | 0,012459551 | PBE1 20S proteasome beta subunit E1 chr1:4452641-4454663 FORWARD LENGTH=274 |
| AT1G16030.1 | 1,199044058 | 0,012647767 | Hsp70b heat shock protein 70B chr1:5502386-5504326 REVERSE LENGTH=646 |
| AT1G56050.1 | 1,413186104 | 0,01279266 | EngD-2 chr1:20963793-20966181 FORWARD LENGTH=421 |
| AT5G26830.1 | 1,243077043 | 0,012836894 | no symbol available no full name available chr5:9437351-9441568 FORWARD LENGTH=709 |
| AT2G47400.1 | 0,793816511 | 0,012883217 | CP12_ CP12-1 CP12 DOMAIN-CONTAINING PROTEIN 1_ CP12 domain-containing protein 1 chr2:19446889-1 |
| AT1G70580.1 | 0,507095305 | 0,012914576 | GGT2_ AOAT2 GLUTAMATE:GLYOXYLATE AMINOTRANSFERASE 2_ alanine-2-oxoglutarate aminotransfe |
| AT2G29500.1 | 9,967952212 | 0,013123269 | HSP17.6B chr2:12633279-12633740 REVERSE LENGTH=153 |
| AT1G08520.1 | 1,26208768 | 0,013278647 | ALB1_ PDE166_ ALB-1V_ CHLD_ V157 PIGMENT DEFECTIVE EMBRYO 166_ ALBINA 1 chr1:2696538-27 |
| AT3G25920.1 | 0,858972554 | 0,013317403 | RPL15 ribosomal protein L15 chr3:9491268-9492558 REVERSE LENGTH=277 |
| AT1G24510.3 | 1,642693372 | 0,01333982 | CCT5 Chaperonin containing T-complex polypeptide-1 subunit 5 chr1:8685504-8687802 REVERSE LENGTH=48 |
| AT3G09840.1 | 1,158985982 | 0,013349004 | ATCDC48_ CDC48_ AtCDC48A_ CDC48A cell division cycle 48 chr3:3019494-3022832 FORWARD LENGTH= |
| AT4G38220.1 | 0,771956678 | 0,013548846 | AQI aquaporin interactor chr4:17925251-17926919 FORWARD LENGTH=430 |
| AT2G43460.1 | 1,356505302 | 0,013611362 | no symbol available no full name available chr2:18046285-18047292 REVERSE LENGTH=69 |
| AT5G51110.1 | 1,304165873 | 0,013723658 | ATP1_ SDIRIP1_ RAF2 SDIR1-INTERACTING PROTEIN1_ AtAIRP2 Target Protein 1_ Rubisco Assembly Fact |
| AT1G17220.1 | 1,537835003 | 0,013821026 | FUG1 fu-gaeri1 chr1:5885383-5890165 FORWARD LENGTH=1026 |
| AT4G25740.1 | 1,557881019 | 0,013825013 | no symbol available no full name available chr4:13107488-13108751 REVERSE LENGTH=177 |
| AT5G12140.1 | 2,613636968 | 0,013895087 | ATCYS1_ CYS1 cystatin-1 chr5:3923295-3923936 REVERSE LENGTH=101 |
| AT1G23730.1 | 0,565435746 | 0,014314556 | ATBCA3_ BCA3 beta carbonic anhydrase 3_ BETA CARBONIC ANHYDRASE 3 chr1:8395965-8398014 FORW |
| AT2G30860.1 | 0,78631634 | 0,014442205 | GSTF9_ ATGSTF9_ ATGSTF7_ GLUTTR glutathione S-transferase PHI 9 chr2:13139132-13140057 FORWARD |
| AT1G43560.1 | 2,53295351 | 0,014537788 | Aty2_ ty2 thioredoxin Y2 chr1:16398359-16399828 REVERSE LENGTH=167 |
| AT3G14310.1 | 0,899249491 | 0,014543918 | OZS2_ ATPME3_ PME3 pectin methylesterase 3_ OVERLY ZINC SENSITIVE 2 chr3:4772214-4775095 REVER |

| | | | |
|---|---|---|---|
| AT3G57260.1 | 17,62449402 | 0,014624538 | AtBG2_ PR2_ GNS2_ AtPR2_ BG2_ PR-2_ BGL2 "beta-1_3-glucanase 2"_ PATHOGENESIS-RELATED PROT[illegible] |
| AT5G65220.1 | 0,70450288 | 0,014746863 | PRPL29 plastid ribosomal proteins of the 50S subunit 29 chr5:26061301-26062506 FORWARD LENGTH=173 |
| AT1G73060.1 | 1,742503008 | 0,014791797 | LPA3 Low PSII Accumulation 3 chr1:27479027-27481258 FORWARD LENGTH=358 |
| AT1G49970.1 | 1,221925393 | 0,014935846 | ClpR1_ SVR2_ NCLPP5_ CLPR1 NUCLEAR CLPP 5_ CLP protease proteolytic subunit 1_ SUPPRESSOR OF V[illegible] |
| AT4G34670.1 | 0,914170298 | 0,014942382 | no symbol available no full name available chr4:16548724-16550222 FORWARD LENGTH=262 |
| AT4G13430.1 | 1,2531936 | 0,015012813 | IIL1_ ATLEUC1 isopropyl malate isomerase large subunit 1 chr4:7804194-7807789 REVERSE LENGTH=509 |
| AT4G09000.1 | 1,137042729 | 0,015047715 | GRF1_ GF14 CHI GENERAL REGULATORY FACTOR1-G-BOX FACTOR 14-3-3 HOMOLOG ISOFORM CH[illegible] |
| AT5G52840.1 | 1,372251141 | 0,015099071 | no symbol available no full name available chr5:21413718-21414794 FORWARD LENGTH=169 |
| AT5G59290.1 | 1,171427677 | 0,015103376 | ATUXS3_ UXS3 UDP-glucuronic acid decarboxylase 3 chr5:23915814-23917953 REVERSE LENGTH=342 |
| AT4G39260.1 | 1,346251325 | 0,015106212 | RBGA6_ CCR1_ ATGRP8_ GR-RBP8_ GRP8 "cold_ circadian rhythm_ and RNA binding 1"_ glycine-rich RNA-[illegible] |
| AT1G20620.1 | 0,824761816 | 0,015176283 | SEN2_ ROG1_ ATCAT3_ CAT3 SENESCENCE 2_ catalase 3_ REPRESSOR OF GSNOR1 chr1:7143142-71461[illegible] |
| AT1G78570.1 | 1,442835675 | 0,015250594 | RHM1_ ATRHM1_ ROL1 REPRESSOR OF LRX1 1_ rhamnose biosynthesis 1_ ARABIDOPSIS THALIANA RH |
| AT1G45000.1 | 1,305589879 | 0,015251911 | RPT4b chr1:17009220-17011607 FORWARD LENGTH=399 |
| AT5G24780.1 | 0,582735638 | 0,015338372 | VSP1_ ATVSP1 vegetative storage protein 1 chr5:8507783-8508889 REVERSE LENGTH=270 |
| AT2G28900.1 | 0,780254698 | 0,015383778 | OEP16_ OEP16-1_ ATOEP16-L_ ATOEP16-1 outer plastid envelope protein 16-1_ OUTER PLASTID ENVELO[illegible] |
| AT5G54270.1 | 0,757671034 | 0,015459966 | LHCB3_ LHCB31 light-harvesting chlorophyll B-binding protein 3 chr5:22038424-22039383 FORWARD LENGT |
| AT3G14420.1 | 1,136201377 | 0,015483964 | GOX1 glycolate oxidase 1 chr3:4821804-4823899 FORWARD LENGTH=367 |
| AT5G51070.1 | 0,725834817 | 0,015579145 | SAG15_ ERD1_ CLPD EARLY RESPONSIVE TO DEHYDRATION 1_ SENESCENCE ASSOCIATED GENE 1 |
| AT4G12800.1 | 0,765624083 | 0,015691428 | PSAL photosystem I subunit l chr4:7521469-7522493 FORWARD LENGTH=219 |
| AT1G79750.1 | 1,572774239 | 0,015787112 | NADP-ME4_ ATNADP-ME4 Arabidopsis thaliana NADP-malic enzyme 4_ NADP-malic enzyme 4 chr1:3000765[illegible] |
| AT2G21170.1 | 1,102285465 | 0,015821264 | TIM_ PDTPI triosephosphate isomerase_ PLASTID ISOFORM TRIOSE PHOSPHATE ISOMERASE chr2:90710[illegible] |
| AT4G29840.1 | 1,249320329 | 0,015954347 | MTO2_ TS THREONINE SYNTHASE_ METHIONINE OVER-ACCUMULATOR 2 chr4:14599434-14601014 R |
| AT2G21530.1 | 1,177957372 | 0,015994053 | no symbol available no full name available chr2:9219372-9220464 FORWARD LENGTH=209 |
| AT2G10940.1 | 1,336117458 | 0,01607592 | no symbol available no full name available chr2:4311160-4312035 REVERSE LENGTH=291 |
| AT5G28510.1 | 0,532640181 | 0,016150093 | BGLU24 beta glucosidase 24 chr5:10481041-10484022 REVERSE LENGTH=533 |
| AT5G42740.3 | 1,323137321 | 0,016204623 | no symbol available no full name available chr5:17136269-17140622 FORWARD LENGTH=528 |
| AT4G02510.1 | 1,669647415 | 0,016301888 | TOC86_ ATTOC159_ TOC160_ PPI2_ TOC159 translocon at the outer envelope membrane of chloroplasts 159_ [illegible] |
| AT1G75040.1 | 3,434266699 | 0,016487915 | PR-5_ PR5 pathogenesis-related gene 5 chr1:28177754-28178731 FORWARD LENGTH=239 |
| AT5G45930.1 | 1,228037068 | 0,016560641 | CHLI2_ CHL I2_ CHLI-2 magnesium chelatase i2 chr5:18628095-18629565 FORWARD LENGTH=418 |
| AT5G11560.1 | 1,397996837 | 0,016890874 | PNET5 chr5:3709734-3713994 REVERSE LENGTH=982 |
| AT5G45280.1 | 0,393391069 | 0,016895943 | PAE11 pectin acetylesterase 11 chr5:18346862-18349432 FORWARD LENGTH=370 |
| AT5G51820.1 | 0,645386108 | 0,017000732 | ATPGMP_ PGM_ PGM1_ STF1 ARABIDOPSIS THALIANA PHOSPHOGLUCOMUTASE_ STARCH-FREE 1_ |
| AT5G12020.1 | 4,482346598 | 0,017027984 | HSP17.6II 17.6 kDa class II heat shock protein chr5:3882409-3882876 REVERSE LENGTH=155 |
| AT4G26300.4 | 1,287097642 | 0,017039602 | emb1027 embryo defective 1027 chr4:13308400-13312204 REVERSE LENGTH=590 |
| AT2G22780.1 | 1,704385216 | 0,017102247 | PMDH1 peroxisomal NAD-malate dehydrogenase 1 chr2:9689995-9691923 REVERSE LENGTH=354 |
| AT1G07410.1 | 0,67716941 | 0,017177 | ATRAB-A2B_ ATRABA2B_ RABA2b_ RAB-A2B ARABIDOPSIS RAB GTPASE HOMOLOG A2B_ RAB GT[illegible] |
| AT3G48560.1 | 1,533274289 | 0,017560441 | CSR1_ IMR1_ TZP5_ ALS_ AHAS ACETOLACTATE SYNTHASE_ TRIAZOLOPYRIMIDINE RESISTANT 5_ |
| AT5G67360.1 | 0,75865567 | 0,017573549 | ARA12_ SBT1.7 Subtilisin-like Serine protease 1.7 chr5:26872192-26874465 REVERSE LENGTH=757 |
| AT4G09320.1 | 1,108750773 | 0,017664632 | ATNDK1_ NDPK1_ NDK1 nucleoside diphosphate kinase 1 chr4:5923484-5924366 FORWARD LENGTH=149 |

| | | |
|---|---|---|
| AT1G22410.1 | 1,773714076 | 0,017722241 no symbol available no full name available chr1:7912120-7914742 FORWARD LENGTH=527 |
| AT5G15450.1 | 2,409271185 | 0,017770246 APG6_ AtCLPB3_ CLPB3_ CLPB-P casein lytic proteinase B3_ CASEIN LYTIC PROTEINASE B-P_ ALBINO [illegible] |
| AT1G15820.1 | 0,88291811 | 0,017775239 CP24_ LHCB6 light harvesting complex photosystem II subunit 6 chr1:5446685-5447676 REVERSE LENGTH=2[illegible] |
| AT4G39800.1 | 1,547257367 | 0,017872676 ATMIPS1_ ATIPS1_ MIPS1_ MI-1-P SYNTHASE INOSITOL 3-PHOSPHATE SYNTHASE 1_ MYO-INOSITO[illegible] |
| AT4G39080.1 | 0,751612367 | 0,017894178 VHA-A3 vacuolar proton ATPase A3 chr4:18209513-18214752 FORWARD LENGTH=821 |
| AT5G02940.1 | 0,600320347 | 0,017942491 PEC1 PLASTID ENVELOPE ION CHANNELS 1 chr5:684671-689674 REVERSE LENGTH=813 |
| AT1G74970.1 | 1,185972574 | 0,01827877 TWN3_ SOT8_ RPS9_ PRPS9 ribosomal protein S9 chr1:28157761-28159202 REVERSE LENGTH=208 |
| AT5G07030.1 | 0,855796658 | 0,018779139 no symbol available no full name available chr5:2183600-2185717 REVERSE LENGTH=455 |
| AT1G01090.1 | 1,209326964 | 0,018873966 PDH-E1 ALPHA pyruvate dehydrogenase E1 alpha chr1:47705-49166 REVERSE LENGTH=428 |
| AT5G47930.1 | 3,932912061 | 0,018927014 no symbol available no full name available chr5:19406423-19407329 REVERSE LENGTH=84 |
| AT5G48480.1 | 2,021751985 | 0,019051254 no symbol available no full name available chr5:19644814-19645658 FORWARD LENGTH=166 |
| AT2G14610.1 | 4,493477331 | 0,019055584 PR1_ ATPR1_ AtCAPE9_ PR 1 PATHOGENESIS-RELATED GENE 1_ pathogenesis-related gene 1 chr2:624194[illegible] |
| AT2G22240.1 | 3,722227213 | 0,019164152 MIPS2_ ATIPS2_ ATMIPS2 myo-inositol-1-phosphate synthase 2_ INOSITOL 3-PHOSPHATE SYNTHASE 2_ [illegible] |
| AT5G11670.1 | 1,195871703 | 0,019212171 NADP-ME2_ ATNADP-ME2 NADP-malic enzyme 2_ Arabidopsis thaliana NADP-malic enzyme 2 chr5:3754456-[illegible] |
| AT2G19760.1 | 0,848257915 | 0,019326051 PFN1_ PRF1 profilin 1_ PROFILIN 1 chr2:8517074-8518067 REVERSE LENGTH=131 |
| AT5G26742.1 | 1,262861132 | 0,019425705 AtRH3_ RH3_ emb1138 embryo defective 1138 chr5:9285540-9288871 REVERSE LENGTH=747 |
| AT1G62780.1 | 1,421723234 | 0,019616231 no symbol available no full name available chr1:23249349-23251066 REVERSE LENGTH=237 |
| AT2G20270.1 | 1,231330884 | 0,01964563 GrxS12 chloroplast Grx 12 chr2:8738001-8739617 REVERSE LENGTH=179 |
| AT2G32060.1 | 1,179171538 | 0,020005402 no symbol available no full name available chr2:13639228-13640104 REVERSE LENGTH=144 |
| AT4G36250.1 | 1,252892354 | 0,020200749 ALDH3F1 aldehyde dehydrogenase 3F1 chr4:17151029-17153381 FORWARD LENGTH=484 |
| AT1G04710.1 | 1,897006637 | 0,020257902 KAT1_ PKT4 3-KETO-ACYL-COA THIOLASE 1_ peroxisomal 3-ketoacyl-CoA thiolase 4 chr1:1321941-132455[illegible] |
| AT4G28440.1 | 1,387927733 | 0,020284092 no symbol available no full name available chr4:14060054-14060970 FORWARD LENGTH=153 |
| AT4G05530.1 | 1,475038648 | 0,020418619 SDRA_ IBR1 indole-3-butyric acid response 1_ SHORT-CHAIN DEHYDROGENASE/REDUCTASE A chr4:281[illegible] |
| AT4G02450.2 | 1,143428973 | 0,020615398 p23-1 chr4:1073987-1075765 REVERSE LENGTH=240 |
| AT4G25050.1 | 1,198910988 | 0,020676339 ACP4_ AtACP4 acyl carrier protein 4 chr4:12870178-12871024 FORWARD LENGTH=137 |
| AT5G13630.1 | 1,973809034 | 0,020922715 ABAR_ CHLH_ GUN5_ CCH_ CCH1 ABA-BINDING PROTEIN_ H SUBUNIT OF MG-CHELATASE_ GENO[illegible] |
| AT5G50920.1 | 1,099490105 | 0,021005179 DCA1_ CLPC_ ATHSP93-V_ CLPC1_ HSP93-V HEAT SHOCK PROTEIN 93-V_ CLPC homologue 1_ DE-RE[illegible] |
| AT1G07140.1 | 0,811788256 | 0,021105096 SIRANBP chr1:2192360-2193688 REVERSE LENGTH=228 |
| AT5G13420.1 | 1,66964476 | 0,021125559 GSM2_ TRA2 Glc-hypersensitive mutant 2_ transaldolase 2 chr5:4302080-4304212 REVERSE LENGTH=438 |
| AT1G22450.1 | 0,714285714 | 0,021227254 ATCOX6B2_ COX6B cytochrome C oxidase 6B_ CYTOCHROME C OXIDASE 6B2 chr1:7925447-7926918 FO[illegible] |
| AT3G16470.1 | 0,507983068 | 0,021238454 JAL35_ AtJAC1_ JR1 jacalin-related lectin 35_ JACALIN-LECTIN LIKE 1_ JASMONATE RESPONSIVE 1 chr3[illegible] |
| AT2G42540.1 | 1,360200864 | 0,021239167 COR15_ AtCOR15A_ COR15A cold-regulated 15a chr2:17711241-17711930 REVERSE LENGTH=127 |
| AT3G48690.1 | 0,266778428 | 0,021563421 ATCXE12_ CXE12 ARABIDOPSIS THALIANA CARBOXYESTERASE 12 chr3:18037186-18038160 REVERS[illegible] |
| AT4G08870.1 | 0,560819531 | 0,02158131 ARGAH2 arginine amidohydrolase 2 chr4:5646654-5648693 REVERSE LENGTH=344 |
| AT3G18740.1 | 1,282294003 | 0,022010341 RPL30C chr3:6453437-6453870 FORWARD LENGTH=112 |
| AT1G16880.1 | 1,061997725 | 0,022250088 ACR11 ACT domain repeats 11 chr1:5773796-5776125 FORWARD LENGTH=290 |
| AT3G24170.1 | 1,464295672 | 0,022474965 ATGR1_ GR1 glutathione-disulfide reductase chr3:8729762-8734115 REVERSE LENGTH=499 |
| AT3G48000.1 | 0,885274645 | 0,02263083 ALDH2B4_ ALDH2A_ ALDH2 aldehyde dehydrogenase 2A_ aldehyde dehydrogenase 2B4_ aldehyde dehydrogen[illegible] |
| AT3G13930.1 | 1,274136763 | 0,022687825 mtE2-2 mitochondrial pyruvate dehydrogenase subunit 2-2 chr3:4596240-4600143 FORWARD LENGTH=539 |

| | | | |
|---|---|---|---|
| AT1G12900.1 | 1,097519107 | 0,023049337 | GAPA-2 glyceraldehyde 3-phosphate dehydrogenase A subunit 2 chr1:4392634-4394283 REVERSE LENGTH=39 |
| AT3G54890.4 | 0,684721417 | 0,023625851 | LHCA1 photosystem I light harvesting complex gene 1 chr3:20339881-20340922 REVERSE LENGTH=213 |
| AT1G06400.1 | 0,646868534 | 0,023655873 | ARA2_ ATRABA1A_ ATRAB11E_ ARA-2 ARABIDOPSIS THALIANA RAB GTPASE HOMOLOG A1A chr1: |
| AT4G27090.1 | 0,651123636 | 0,023871085 | RPL14B chr4:13594104-13595187 REVERSE LENGTH=134 |
| AT1G34430.1 | 0,81189886 | 0,023978238 | EMB3003 embryo defective 3003 chr1:12588027-12590084 REVERSE LENGTH=465 |
| AT3G05540.1 | 1,493824256 | 0,024091213 | TCTP2 Translationally Controlled Tumor Protein 2 chr3:1606487-1608030 REVERSE LENGTH=168 |
| AT3G11710.1 | 1,57236556 | 0,024364863 | ATKRS-1 lysyl-tRNA synthetase 1 chr3:3702359-3705613 REVERSE LENGTH=626 |
| AT3G20050.1 | 1,267042174 | 0,02476537 | ATTCP-1_ TCP-1_ CCT1 Chaperonin containing T-complex polypeptide-1 subunit 1_ T-complex protein 1 alpha s |
| AT3G56940.1 | 0,808646198 | 0,024863392 | CRD1_ ACSF_ CHL27 COPPER RESPONSE DEFECT 1 chr3:21076594-21078269 FORWARD LENGTH=409 |
| AT5G53560.1 | 0,812169251 | 0,025054701 | ATB5-A_ CB5-E_ ATCB5-E_ B5 #2 ARABIDOPSIS CYTOCHROME B5 ISOFORM E_ cytochrome B5 isoform |
| AT5G44320.1 | 1,230470009 | 0,025168172 | no symbol available no full name available chr5:17854901-17856667 REVERSE LENGTH=588 |
| AT3G44860.1 | 1,586384501 | 0,025276222 | FAMT farnesoic acid carboxyl-O-methyltransferase chr3:16379689-16380939 FORWARD LENGTH=348 |
| AT3G17390.1 | 0,868447192 | 0,025400138 | SAMS3_ AtSAMS3_ MAT4_ MTO3 METHIONINE ADENOSYLTRANSFERASE 4_ S-ADENOSYLMETHION |
| AT5G63570.1 | 1,123816363 | 0,025441378 | GSA1 "glutamate-1-semialdehyde-2_1-aminomutase" chr5:25451957-25453620 FORWARD LENGTH=474 |
| AT1G22780.1 | 0,941094114 | 0,025583718 | RPS18A_ PFL_ PFL1 POINTED FIRST LEAVES_ POINTED FIRST LEAVES 1_ 40S RIBOSOMAL PROTEIN |
| AT5G44340.1 | 1,224230659 | 0,025845587 | TUB4 tubulin beta chain 4 chr5:17859442-17860994 REVERSE LENGTH=444 |
| AT2G18040.1 | 1,069409524 | 0,025893187 | PIN1AT "peptidylprolyl cis/trans isomerase_ NIMA-interacting 1" chr2:7842346-7843537 FORWARD LENGTH= |
| ATCG00540.1 | 0,882020937 | 0,025900869 | PETA photosynthetic electron transfer A chrc:61657-62619 FORWARD LENGTH=320 |
| AT3G53230.1 | 1,234314413 | 0,025945616 | AtCDC48B cell division cycle 48B chr3:19723416-19726489 FORWARD LENGTH=815 |
| AT1G27090.1 | 1,693541874 | 0,026059969 | no symbol available no full name available chr1:9404041-9406098 REVERSE LENGTH=420 |
| AT1G56330.1 | 1,285065737 | 0,026332945 | ATSARA1B_ SAR1B_ SAR1_ ATSAR1B_ ATSAR1 SECRETION-ASSOCIATED RAS 1_ ARABIDOPSIS THA |
| AT3G27740.2 | 1,599586769 | 0,026539712 | CARA_ VEN6 carbamoyl phosphate synthetase A_ VENOSA 6 chr3:10281470-10283792 REVERSE LENGTH=3 |
| AT5G22880.1 | 1,135641015 | 0,026765513 | HTB2_ H2B histone B2_ HISTONE H2B chr5:7652130-7652567 REVERSE LENGTH=145 |
| AT3G54640.1 | 1,231175855 | 0,027015264 | TRP3_ TSA1 TRYPTOPHAN-REQUIRING 3_ tryptophan synthase alpha chain chr3:20223331-20225303 REVEF |
| AT3G04720.1 | 0,602247567 | 0,027059976 | HEL_ PR-4_ AtPR4_ PR4 HEVEIN-LIKE_ pathogenesis-related 4 chr3:1285691-1286531 REVERSE LENGTH=2 |
| AT3G47470.1 | 0,788010885 | 0,027068639 | LHCA4_ CAB4 light-harvesting chlorophyll-protein complex I subunit A4 chr3:17493622-17494773 REVERSE LI |
| AT4G31500.1 | 1,271857083 | 0,027618709 | RNT1_ RED1_ SUR2_ ATR4_ CYP83B1 RED ELONGATED 1_ SUPERROOT 2_ ALTERED TRYPTOPHAN I |
| AT3G14390.1 | 1,564068807 | 0,027973304 | DAPDC1 meso-diaminopimelate decarboxylase 1 chr3:4806771-4808954 FORWARD LENGTH=484 |
| AT1G32900.1 | 1,197146124 | 0,027991738 | GBSS1 granule bound starch synthase 1 chr1:11920582-11923506 REVERSE LENGTH=610 |
| AT3G46010.1 | 0,396315431 | 0,02799441 | atadf_ ADF1_ ATADF1 actin depolymerizing factor 1 chr3:16909679-16910678 REVERSE LENGTH=139 |
| AT3G48870.1 | 1,288726578 | 0,028046512 | ATCLPC_ HSP93-III_ ClpC2_ ATHSP93-III ClpC2 chr3:18122363-18126008 REVERSE LENGTH=952 |
| AT5G35790.1 | 1,351241677 | 0,028081641 | G6PD1 glucose-6-phosphate dehydrogenase 1 chr5:13956879-13959686 REVERSE LENGTH=576 |
| AT3G02520.1 | 1,168802327 | 0,028597647 | GRF7_ GF14 NU general regulatory factor 7 chr3:526800-527915 REVERSE LENGTH=265 |
| AT3G02630.1 | 1,133221964 | 0,028684863 | AAD5 ACYL?ACYL CARRIER PROTEIN (ACP) DESATURASE 5 chr3:562164-564524 FORWARD LENGTH= |
| AT3G61470.1 | 0,781856967 | 0,028755421 | LHCA2 photosystem I light harvesting complex gene 2 chr3:22745736-22747032 FORWARD LENGTH=257 |
| AT2G18450.1 | 1,27450723 | 0,028867542 | SDH1-2 succinate dehydrogenase 1-2 chr2:7997510-8000801 REVERSE LENGTH=632 |
| AT4G29060.1 | 1,267283433 | 0,029381327 | emb2726 embryo defective 2726 chr4:14317744-14321315 FORWARD LENGTH=953 |
| AT2G43950.1 | 1,512606867 | 0,029672204 | ATOEP37_ OEP37 chloroplast outer envelope protein 37_ ARABIDOPSIS CHLOROPLAST OUTER ENVELOPI |
| AT1G04170.1 | 1,447968006 | 0,03005688 | EIF2 GAMMA eukaryotic translation initiation factor 2 gamma subunit chr1:1097423-1099702 FORWARD LENG |

| | | | |
|---|---|---|---|
| AT1G79210.1 | 0,931005628 | 0,030072627 | no symbol available no full name available chr1:29796286-29798240 REVERSE LENGTH=235 |
| AT5G19770.1 | 1,126283889 | 0,030196784 | TUA3 tubulin alpha-3 chr5:6682761-6684474 REVERSE LENGTH=450 |
| AT4G39890.1 | 3,219054476 | 0,030346922 | AtRABH1c_ RABH1c RAB GTPase homolog H1C chr4:18506112-18507459 FORWARD LENGTH=214 |
| AT5G19990.1 | 1,281782136 | 0,030512663 | RPT6A_ ATSUG1 regulatory particle triple-A ATPase 6A chr5:6752144-6754918 FORWARD LENGTH=419 |
| AT3G29360.1 | 1,413299647 | 0,03065407 | UGD2 UDP-glucose dehydrogenase 2 chr3:11267375-11268817 REVERSE LENGTH=480 |
| AT2G20360.1 | 1,246901421 | 0,030749183 | no symbol available no full name available chr2:8786070-8789098 FORWARD LENGTH=402 |
| AT1G15140.1 | 1,588129682 | 0,031074537 | FNRL FERREDOXIN-NADP(+) OXIDOREDUCTASE -LIKE chr1:5210403-5212137 REVERSE LENGTH=295 |
| AT4G00570.1 | 1,194007645 | 0,031076293 | NAD-ME2 NAD-dependent malic enzyme 2 chr4:242817-246522 REVERSE LENGTH=607 |
| AT3G08030.2 | 0,794440036 | 0,03121612 | AthA2-1 chr3:2564517-2565819 FORWARD LENGTH=323 |
| AT1G03475.1 | 1,187154698 | 0,031360349 | HEMF1_ ATCPO-I_ LIN2 LESION INITIATION 2 chr1:869302-871175 REVERSE LENGTH=386 |
| AT2G40840.1 | 1,331502703 | 0,031564035 | DPE2 disproportionating enzyme 2 chr2:17045368-17050779 FORWARD LENGTH=955 |
| AT5G58290.1 | 0,708816686 | 0,031625075 | RPT3 regulatory particle triple-A ATPase 3 chr5:23569155-23571116 FORWARD LENGTH=408 |
| AT3G59970.3 | 1,120371428 | 0,031966276 | MTHFR1 methylenetetrahydrofolate reductase 1 chr3:22151303-22154323 FORWARD LENGTH=592 |
| AT5G63890.1 | 1,436482734 | 0,032193298 | HISN8_ ATHDH_ HDH histidinol dehydrogenase_ HISTIDINE BIOSYNTHESIS 8 chr5:25565600-25567879 RE |
| AT2G47730.1 | 1,228793476 | 0,032317931 | GST6_ GSTF8_ ATGSTF8_ ATGSTF5 glutathione S-transferase phi 8_ Arabidopsis thaliana glutathione S-transfer |
| AT1G31180.1 | 1,197112137 | 0,032333229 | ATIMD3_ IMDH3_ IMD3_ IPMDH1 ISOPROPYLMALATE DEHYDROGENASE 1_ isopropylmalate dehydrog |
| AT1G78830.1 | 1,210313833 | 0,032530887 | MNB1 chr1:29637141-29638508 REVERSE LENGTH=455 |
| AT1G22740.1 | 0,395981327 | 0,032580045 | RABG3B_ RAB7_ ATRABG3B_ RAB75 RAB GTPase homolog G3B chr1:8049247-8050494 FORWARD LENG |
| AT5G61780.1 | 1,225219042 | 0,032663285 | Tudor2_ AtTudor2_ TSN2 Arabidopsis thaliana TUDOR-SN protein 2_ TUDOR-SN protein 2 chr5:24822012-248 |
| AT5G58330.1 | 1,203347085 | 0,032714133 | NADP-MDH NADP-dependent Malate Dehydrogenase chr5:23580010-23582287 REVERSE LENGTH=443 |
| AT5G26360.1 | 1,446626208 | 0,032809118 | CCT3 Chaperonin containing T-complex polypeptide-1 subunit 3 chr5:9255561-9258891 REVERSE LENGTH=55 |
| AT1G09100.1 | 0,807678736 | 0,032840816 | RPT5B 26S proteasome AAA-ATPase subunit RPT5B chr1:2936675-2939258 REVERSE LENGTH=423 |
| AT1G09340.1 | 1,252931388 | 0,03288242 | CRB_ CSP41B_ HIP1.3 chloroplast RNA binding_ heteroglycan-interacting protein 1.3_ CHLOROPLAST STEM- |
| AT5G14780.1 | 0,695122678 | 0,032921787 | FDH_ AtFDH1 formate dehydrogenase chr5:4777043-4779190 FORWARD LENGTH=384 |
| AT1G78370.1 | 1,171046762 | 0,032925089 | ATGSTU20_ GSTU20 glutathione S-transferase TAU 20 chr1:29484428-29485204 REVERSE LENGTH=217 |
| AT2G41220.1 | 1,103881322 | 0,033046606 | GLU2 glutamate synthase 2 chr2:17177934-17188388 FORWARD LENGTH=1629 |
| AT1G75350.1 | 0,879550025 | 0,033085891 | emb2184 embryo defective 2184 chr1:28272163-28272687 FORWARD LENGTH=144 |
| AT1G59870.1 | 1,39008842 | 0,03321773 | ABCG36_ ATABCG36_ PEN3_ PDR8_ ATPDR8 Arabidopsis thaliana ATP-binding cassette G36_ PLEIOTROPI |
| AT4G28390.1 | 1,469907366 | 0,03325375 | AAC3_ ATAAC3 ADP/ATP carrier 3 chr4:14041486-14042781 REVERSE LENGTH=379 |
| AT4G12420.1 | 0,859970509 | 0,033338148 | SKU5 chr4:7349941-7352868 REVERSE LENGTH=587 |
| AT5G50950.3 | 0,647442277 | 0,033362729 | FUM2 FUMARASE 2 chr5:20731191-20733636 FORWARD LENGTH=317 |
| AT5G13410.1 | 0,863237048 | 0,033434443 | no symbol available no full name available chr5:4299830-4301706 REVERSE LENGTH=256 |
| AT5G63860.1 | 1,873417869 | 0,033536614 | AtUVR8_ UVR8 UVB-RESISTANCE 8 chr5:25554821-25558587 REVERSE LENGTH=440 |
| AT3G06050.1 | 0,843757505 | 0,033927253 | PRXIIF_ ATPRXIIF peroxiredoxin IIF_ PEROXIREDOXIN IIF chr3:1826311-1827809 REVERSE LENGTH=20 |
| AT1G09130.1 | 1,239929701 | 0,034171707 | no symbol available no full name available chr1:2940063-2942217 REVERSE LENGTH=330 |
| AT4G28706.1 | 1,314264241 | 0,034261463 | no symbol available no full name available chr4:14167805-14170619 FORWARD LENGTH=401 |
| AT2G30110.1 | 0,615706638 | 0,034447732 | MOS5_ ATUBA1_ UBA1 MODIFIER OF SNC1 5_ ubiquitin-activating enzyme 1 chr2:12852632-12857369 REV |
| AT1G14810.1 | 1,123918281 | 0,034613966 | no symbol available no full name available chr1:5102684-5104633 REVERSE LENGTH=375 |
| AT3G49870.1 | 1,219913256 | 0,034692423 | ARLA1C_ ARL8a_ ATARLA1C ADP-ribosylation factor-like A1C_ ADP-ribosylation factor-like 8a chr3:1849267 |

| | | | |
|---|---|---|---|
| AT5G02160.1 | 0,685725492 | 0,034779958 | FIP FtsH5 Interacting Protein chr5:426392-427024 FORWARD LENGTH=129 |
| AT3G56460.1 | 1,424259035 | 0,034827369 | no symbol available no full name available chr3:20933029-20934425 REVERSE LENGTH=348 |
| AT3G02360.1 | 1,313982239 | 0,03482831 | PGD2 6-phosphogluconate dehydrogenase 2 chr3:482498-483958 FORWARD LENGTH=486 |
| AT3G62120.3 | 1,191982333 | 0,035071624 | ProRS-Cyt_ AtProRS-Cyt prolyl-tRNA synthetase cytosolic chr3:23001227-23003849 REVERSE LENGTH=517 |
| AT3G54470.1 | 2,118457253 | 0,035242107 | no symbol available no full name available chr3:20168285-20170245 REVERSE LENGTH=476 |
| AT1G02930.1 | 2,632292613 | 0,035394713 | ATGSTF3_ ATGSTF6_ GST1_ ERD11_ GSTF6_ ATGST1 EARLY RESPONSIVE TO DEHYDRATION 11_ AI |
| AT3G09260.1 | 1,465426847 | 0,035805886 | LEB_ BGLU23_ PYK10_ PSR3.1 LONG ER BODY chr3:2840657-2843730 REVERSE LENGTH=524 |
| AT4G32470.1 | 1,495118627 | 0,036205339 | no symbol available no full name available chr4:15669641-15671095 REVERSE LENGTH=122 |
| AT5G42020.2 | 1,213903048 | 0,036238707 | BIP2_ BIP luminal binding protein chr5:16807697-16810480 REVERSE LENGTH=613 |
| AT3G20390.1 | 0,871541387 | 0,036281984 | RidA Reactive Intermediate Deaminase A chr3:7110227-7111695 REVERSE LENGTH=187 |
| AT4G31180.1 | 1,951395674 | 0,037110687 | IBI1 impaired in BABA-induced disease immunity 1 chr4:15156696-15159362 FORWARD LENGTH=558 |
| AT5G11170.1 | 1,169012309 | 0,037170591 | UAP56a homolog of human UAP56 a chr5:3553334-3556646 FORWARD LENGTH=427 |
| AT2G40610.1 | 0,775682172 | 0,03741598 | ATHEXP ALPHA 1.11_ EXP8_ ATEXPA8_ ATEXP8_ EXPA8 expansin A8 chr2:16949121-16950472 REVERSI |
| AT1G77940.1 | 0,565548177 | 0,037594985 | RPL30B chr1:29304116-29305288 REVERSE LENGTH=112 |
| AT3G55330.1 | 0,354044396 | 0,038494543 | PPL1 PsbP-like protein 1 chr3:20514031-20515275 REVERSE LENGTH=230 |
| ATCG00820.1 | 0,653230914 | 0,038528861 | RPS19 ribosomal protein S19 chrc:84005-84283 REVERSE LENGTH=92 |
| AT1G26230.2 | 0,753774819 | 0,039056262 | Cpn60beta4_ CPNB4 chaperonin-60beta4 chr1:9072715-9075272 REVERSE LENGTH=551 |
| AT5G51970.1 | 0,850846183 | 0,039172244 | ATSDH SORBITOL DEHYDROGENASE chr5:21111820-21113284 FORWARD LENGTH=364 |
| AT1G70730.1 | 1,142024063 | 0,039357752 | PGM2 phosphoglucomutase 2 chr1:26669020-26672726 REVERSE LENGTH=585 |
| AT2G14260.2 | 1,278646109 | 0,040134533 | PIP_ PAP1 proline iminopeptidase_ prolyl aminopeptidase 1 chr2:6041441-6043475 REVERSE LENGTH=329 |
| AT5G23010.1 | 1,329320586 | 0,040360638 | GSM1_ MAM1_ IMS3 glucosinolate metabolism 1_ 2-ISOPROPYLMALATE SYNTHASE 3_ methylthioalkylmal |
| AT3G18490.1 | 0,872198404 | 0,040524328 | ASPG1 ASPARTIC PROTEASE IN GUARD CELL 1 chr3:6349090-6350592 REVERSE LENGTH=500 |
| AT1G17650.1 | 1,234021789 | 0,040683594 | AtGLYR2_ GLYR2_ GR2 GLYOXYLATE REDUCTASE 2_ glyoxylate reductase 2 chr1:6069594-6071964 REV |
| AT3G53420.1 | 0,833536322 | 0,040725189 | PIP2;1_ AtPIP2;1_ PIP2A_ PIP2 PLASMA MEMBRANE INTRINSIC PROTEIN 2_ PLASMA MEMBRANE IN |
| AT5G45680.1 | 1,766328012 | 0,041057764 | ATFKBP13_ FKBP13 FK506 BINDING PROTEIN 13_ FK506-binding protein 13 chr5:18530894-18532128 FOR |
| AT2G17630.1 | 0,751865612 | 0,041174777 | PSAT2 phosphoserine aminotransferase 2 chr2:7666637-7667905 FORWARD LENGTH=422 |
| AT1G04270.2 | 0,770960331 | 0,041292281 | RPS15 cytosolic ribosomal protein S15 chr1:1141852-1142960 REVERSE LENGTH=151 |
| AT2G45300.3 | 2,93555533 | 0,041743004 | no symbol available no full name available chr2:18677518-18680118 FORWARD LENGTH=518 |
| AT3G52930.1 | 1,17088087 | 0,041762309 | AtFBA8_ FBA8 fructose-bisphosphate aldolase 8 chr3:19627383-19628874 REVERSE LENGTH=358 |
| AT3G63190.1 | 0,905971866 | 0,042426834 | cpRRF_ HFP108_ AtcpRRF_ RRF chloroplast ribosome recycling factor_ Arabidopsis thaliana chloroplast ribosom |
| AT1G52220.1 | 0,534499421 | 0,042734667 | CURT1C CURVATURE THYLAKOID 1C chr1:19453770-19454605 REVERSE LENGTH=156 |
| AT1G07400.1 | 2,78171583 | 0,043437928 | HSP17.8 chr1:2275148-2275621 FORWARD LENGTH=157 |
| AT2G13360.1 | 1,12760467 | 0,043511776 | SGAT_ AGT_ AGT1 ALANINE:GLYOXYLATE AMINOTRANSFERASE 1_ alanine:glyoxylate aminotransferas |
| AT5G44720.1 | 0,66291888 | 0,043945488 | no symbol available no full name available chr5:18043086-18045275 FORWARD LENGTH=308 |
| AT5G13120.1 | 0,81269783 | 0,044029092 | CYP20-2_ Pnsl5_ ATCYP20-2 cyclophilin 20-2_ Photosynthetic NDH subcomplex L 5_ ARABIDOPSIS THALIA |
| AT1G23190.1 | 1,252799109 | 0,044319553 | PGM3 phosphoglucomutase 3 chr1:8219946-8224186 FORWARD LENGTH=583 |
| AT1G09270.1 | 1,316701522 | 0,044743907 | IMPA-4 importin alpha isoform 4 chr1:2994506-2997833 FORWARD LENGTH=538 |
| AT2G31570.1 | 1,233480777 | 0,044795098 | ATGPX2_ GPX2_ GPXL2 glutathione peroxidase 2 chr2:13438211-13439775 REVERSE LENGTH=169 |
| AT4G35630.1 | 1,148417755 | 0,044854734 | PSAT1 phosphoserine aminotransferase 1 chr4:16904205-16905497 FORWARD LENGTH=430 |

| | | | |
|---|---|---|---|
| AT1G20340.1 | 0,801267143 | 0,044881768 | PETE2_ DRT112 DNA-DAMAGE-REPAIR/TOLERATION PROTEIN 112_ PLASTOCYANIN 2 chr1:7042770-[illegible] |
| AT3G56190.1 | 1,285954445 | 0,045111966 | ALPHA-SNAP2_ ASNAP alpha-soluble NSF attachment protein 2 chr3:20846119-20848356 REVERSE LENGTH |
| AT5G15970.1 | 10,57499893 | 0,045736918 | AtCor6.6_ KIN2_ COR6.6 COLD-RESPONSIVE 6.6 chr5:5211966-5212441 FORWARD LENGTH=66 |
| AT2G39730.1 | 1,073741148 | 0,045839343 | RCA rubisco activase chr2:16570951-16573345 REVERSE LENGTH=474 |
| AT3G62030.1 | 1,061792705 | 0,046169937 | ROC4_ CYP20-3 rotamase CYP 4_ cyclophilin 20-3 chr3:22973708-22975139 FORWARD LENGTH=260 |
| AT5G55220.1 | 0,55057077 | 0,046406663 | HP65b_ TIG1 chr5:22397677-22400678 FORWARD LENGTH=547 |
| AT3G28290.1 | 1,64203778 | 0,046812298 | AT14A chr3:10547873-10549030 FORWARD LENGTH=385 |
| AT5G23060.1 | 0,808222553 | 0,047805883 | CaS calcium sensing receptor chr5:7736760-7738412 REVERSE LENGTH=387 |
| AT4G12060.1 | 0,407651307 | 0,047845006 | ClpT2 chr4:7228269-7229898 REVERSE LENGTH=241 |
| AT1G35680.1 | 0,885336199 | 0,048090997 | RPL21C_ ASD chloroplast ribosomal protein L21_ ATPase-in-Seed-Development chr1:13208777-13210246 FORW |
| AT4G09650.1 | 1,05541456 | 0,048117386 | PDE332_ ATPD PIGMENT DEFECTIVE 332_ ATP synthase delta-subunit gene chr4:6100799-6101503 FORWA |
| AT2G40290.1 | 1,220709086 | 0,048391465 | no symbol available no full name available chr2:16829030-16830889 REVERSE LENGTH=344 |
| AT3G16520.3 | 1,39181458 | 0,048707677 | UGT88A1 UDP-glucosyl transferase 88A1 chr3:5619355-5620833 REVERSE LENGTH=462 |
| AT3G12260.1 | 0,773525959 | 0,048827056 | NDUFA6_ B14 chr3:3909252-3910337 REVERSE LENGTH=133 |
| AT3G23700.1 | 1,079396944 | 0,049469834 | SRRP1 S1 RNA-binding ribosomal protein 1 chr3:8531689-8533742 REVERSE LENGTH=392 |
| AT2G28815.1 | 1,261109544 | 0,049538412 | no symbol available no full name available chr2:12367001-12368064 REVERSE LENGTH=291 |
| AT3G48990.1 | 1,164800993 | 0,049861407 | AAE3 ACYL-ACTIVATING ENZYME 3 chr3:18159031-18161294 REVERSE LENGTH=514 |
| AT1G06430.1 | 0,45011708 | 0,049898697 | FTSH8 FTSH protease 8 chr1:1960214-1962525 REVERSE LENGTH=685 |

E PROTEIN 70-1_ heat shock cognate protein 70-1_ HEAT SHOCK COGNATE PROTEIN 70_ HEAT SHOCK PROTEIN 70-1 chr5:554055-556334 REVERSE LENGTH=

'HASE-B_ CHLOROPLAST O-ACETYLSERINE SULFHYDRYLASE 1_ ARABIDOPSIS CYSTEINE SYNTHASE 1 chr2:18129604-18132322 REVERSE LENGTH=392

STERILITY 3_ NON-PHOTOCHEMICAL QUENCHING 2_ ARABIDOPSIS THALIANA ZEAXANTHIN EPOXIDASE_ LOW EXPRESSION OF OSMOTIC STRESS-R

ROPLAST CHAPERONIN 60ALPHA_ ACCUMULATION AND REPLICATION OF CHLOROPLASTS 2_ chaperonin-60alpha chr2:11926603-11929184 FORWARD LE

N FACTOR 3B1_ EUKARYOTIC TRANSLATION INITIATION FACTOR 3B1_ EUKARYOTIC TRANSLATION INITIATION FACTOR 3B_ translation initiation factor

NDENT GENE SILENCING 1_ MATERNAL EFFECT EMBRYO ARREST 58_ S-ADENOSYL-L-HOMOCYSTEIN HYDROLASE 1 chr4:8054931-8056676 FORWARD I

)ENOSINE-5'-PHOSPHOSULFATE (PAPS) REDUCTASE HOMOLOG 43_ 5'adenylylphosphosulfate reductase 2 chr1:22975794-22977465 REVERSE LENGTH=454

.VING COMPLEX 33 KILODALTON PROTEIN_ OXYGEN EVOLVING ENHANCER PROTEIN 33_ PS II oxygen-evolving complex 1_ MANGANESE-STABILIZING [illegible]

ABIDOPSIS THALIANA PROTEIN DISULFIDE ISOMERASE 11_ UNFERTILIZED EMBRYO SAC 5_ MATERNAL EFFECT EMBRYO ARREST 30 chr2:19481503-19

.Y ACTIVE 8_ THYLAKOID MEMBRANE PHOSPHOPROTEIN OF 14 KDA_ photosystem I P subunit chr2:19243729-19244870 FORWARD LENGTH=174

'LASTID PROTEIN IMPORT 2_ TRANSLOCON AT THE OUTER ENVELOPE MEMBRANE OF CHLOROPLASTS 86_ TRANSLOCON AT THE OUTER ENVELOPE

_ IMIDAZOLE RESISTANT 1_ ACETOHYDROXY ACID SYNTHASE_ chlorsulfuron/imidazolinone resistant 1 chr3:18001530-18003542 REVERSE LENGTH=670

L-1-PHOSPHATE SYNTHASE 1_ myo-inositol-1-phosphate synthase 1_ D-myo-Inositol 3-Phosphate Synthase 1 chr4:18469659-18471893 REVERSE LENGTH=511

C DRUG RESISTANCE 8_ ARABIDOPSIS PLEIOTROPIC DRUG RESISTANCE 8_ PENETRATION 3_ ATP-binding cassette G36 chr1:22034661-22039844 FORWARI

RABIDOPSIS GLUTATHIONE S-TRANSFERASE 1_ ARABIDOPSIS THALIANA GLUATIONE S-TRANSFERASE F3_ glutathione S-transferase 6_ GLUTATHIONE S-

e recycling factor_ "ribosome recycling factor_ chloroplast precursor"_ HIGH CHLOROPHYLL FLUORESCENCE AND PALE GREEN MUTANT 108 chr3:23342861-233

[illegible]ESPONSIVE GENES 6_ ARABIDOPSIS THALIANA ABA DEFICIENT 1_ ZEAXANTHIN EPOXIDASE chr5:26754026-26757090 REVERSE LENGTH=610
