## Supplementary material for "HY5 enhances *Arabidopsis* tolerance to combined high light and heat stress by coordinating photoprotection and hormone signaling": Table S9

**Supplemental Table S10. Differentially accumulated proteins compared to control (P < 0.05) in HY5OX leaves subjected to combined conditions of high light and heat stress**

| Protein ID | Fold Change | p-value | Protein description |
|---|---|---|---|
| AT5G12030.1 | 17,19194702 | 3,32141E-09 | AT-HSP17.6A_ HSP17.6A_ HSP17.6 HEAT SHOCK PROTEIN 17.6_ heat shock protein 17.6A chr5:3884214-3884684 R |
| AT5G14740.2 | 0,131983079 | 2,90858E-08 | BETA CA2_ CA2_ DEG12_ CA18 BETA CARBONIC ANHYDRASE 2_ CARBONIC ANHYDRASE 18_ carbonic anhy |
| AT5G64040.1 | 2,941415217 | 3,50956E-08 | PSAN chr5:25628724-25629409 REVERSE LENGTH=171 |
| AT5G52640.1 | 17,31263125 | 3,72308E-08 | HSP81-1_ ATHSP90.1_ HSP81.1_ AtHsp90-1_ HSP83_ ATHS83_ HSP90.1 HEAT SHOCK PROTEIN 90-1_ heat shock |
| AT5G12020.1 | 23,09181512 | 5,09312E-08 | HSP17.6II 17.6 kDa class II heat shock protein chr5:3882409-3882876 REVERSE LENGTH=155 |
| AT3G05590.1 | 0,101094434 | 7,086E-08 | RPL18 ribosomal protein L18 chr3:1621511-1622775 FORWARD LENGTH=187 |
| AT2G39310.1 | 10,17361097 | 7,46683E-08 | JAL22 jacalin-related lectin 22 chr2:16414262-16416323 REVERSE LENGTH=458 |
| AT4G04020.1 | 6,890755265 | 1,00919E-07 | FIB_ PGL35_ FIB1a plastoglobulin 35_ fibrillin 1a_ fibrillin chr4:1932161-1933546 FORWARD LENGTH=318 |
| AT1G48600.1 | 2,978548576 | 1,1027E-07 | AtPMT2_ AtPMEAMT_ PMEAMT phosphoethanolamine N-methyltransferase_ Phosphoethanolamine methyltransferase2 |
| AT1G74310.1 | 22,53556359 | 1,82249E-07 | ATHSP101_ HSP101_ HOT1 heat shock protein 101 chr1:27936715-27939862 REVERSE LENGTH=911 |
| AT2G29500.1 | 61,34607251 | 1,83644E-07 | HSP17.6B chr2:12633279-12633740 REVERSE LENGTH=153 |
| AT3G09630.1 | 0,204800496 | 2,0251E-07 | SAC56 Suppressor of Acaulis 56 chr3:2953813-2955444 FORWARD LENGTH=406 |
| AT3G10950.1 | 0,237861265 | 2,81462E-07 | no symbol available no full name available chr3:3423893-3424566 FORWARD LENGTH=92 |
| AT5G40950.1 | 0,591508151 | 6,61105E-07 | PRPL27_ RPL27 ribosomal protein large subunit 27 chr5:16410866-16411845 FORWARD LENGTH=198 |
| AT4G36130.1 | 0,037572359 | 7,14223E-07 | no symbol available no full name available chr4:17097613-17098656 FORWARD LENGTH=258 |
| AT3G12580.1 | 6,347547664 | 8,13563E-07 | HSP70_ ATHSP70_ HSC70-4 ARABIDOPSIS HEAT SHOCK PROTEIN 70_ heat shock protein 70 chr3:3991487-39936 |
| AT3G01390.1 | 8,994325512 | 8,37822E-07 | AVMA10_ VMA10 vacuolar membrane ATPase 10 chr3:150265-150922 REVERSE LENGTH=110 |
| AT2G35635.1 | 20,71417375 | 8,45793E-07 | UBQ7_ RUB2 RELATED TO UBIQUITIN 2_ ubiquitin 7 chr2:14981044-14981943 FORWARD LENGTH=154 |
| AT1G79550.1 | 2,471924906 | 8,65962E-07 | PGKc_ PGK_ PGK3 phosphoglycerate kinase_ phosphoglycerate kinase 3 chr1:29924347-29926295 REVERSE LENGTH |
| AT2G18450.1 | 2,742355203 | 9,30916E-07 | SDH1-2 succinate dehydrogenase 1-2 chr2:7997510-8000801 REVERSE LENGTH=632 |
| AT1G16030.1 | 4,485044872 | 9,54772E-07 | Hsp70b heat shock protein 70B chr1:5502386-5504326 REVERSE LENGTH=646 |
| AT1G20200.1 | 1,637011477 | 1,01527E-06 | HAP15_ EMB2719 HAPLESS 15_ EMBRYO DEFECTIVE 2719 chr1:7001409-7004154 REVERSE LENGTH=488 |
| AT5G02870.1 | 0,331622571 | 1,03161E-06 | RPL4 ribosomal large subunit 4 chr5:657830-659526 FORWARD LENGTH=407 |
| AT5G48570.1 | 18,15745164 | 1,04973E-06 | ROF2_ ATFKBP65_ FKBP65 chr5:19690746-19693656 REVERSE LENGTH=578 |
| AT2G24270.1 | 1,66407234 | 1,06071E-06 | ALDH11A3 aldehyde dehydrogenase 11A3 chr2:10327325-10329601 REVERSE LENGTH=496 |
| AT2G18020.1 | 0,106362092 | 1,22084E-06 | EMB2296 embryo defective 2296 chr2:7837151-7838160 FORWARD LENGTH=258 |
| AT5G14740.5 | 1,725263423 | 1,25261E-06 | BETA CA2_ CA2_ DEG12_ CA18 BETA CARBONIC ANHYDRASE 2_ CARBONIC ANHYDRASE 18_ carbonic anhy |
| AT3G11130.1 | 2,289960897 | 1,26431E-06 | AtCHC1_ CHC1_ HAS1 clathrin heavy chain 1_ hot ABA deficiency suppressor 1 chr3:3482575-3491667 REVERSE LEN |
| AT3G09350.1 | 22,6276958 | 1,41724E-06 | Fes1A Fes1A chr3:2871216-2873109 FORWARD LENGTH=363 |
| AT1G73600.1 | 4,562565935 | 1,49098E-06 | DEG26_ NMT_ AtPMT3_ NMT3 Phosphoethanolamine methyltransferase3 chr1:27670825-27673400 FORWARD LENG |
| AT4G18100.1 | 0,072102223 | 1,57785E-06 | no symbol available no full name available chr4:10035715-10036475 REVERSE LENGTH=133 |
| AT3G13470.1 | 1,765973333 | 1,57795E-06 | CPNB2_ Cpn60beta2 chaperonin-60beta2 chr3:4389685-4392624 FORWARD LENGTH=596 |
| AT4G30270.1 | 0,369282387 | 1,89906E-06 | MERI5B_ XTH24_ MERI-5_ SEN4 xyloglucan endotransglucosylase/hydrolase 24_ SENESCENCE 4_ MERISTEM 5_ m |
| AT5G02960.1 | 0,217799392 | 1,92142E-06 | no symbol available no full name available chr5:693280-694396 REVERSE LENGTH=142 |
| AT1G13930.1 | 5,828549619 | 2,03407E-06 | no symbol available no full name available chr1:4761091-4761558 FORWARD LENGTH=155 |
| AT5G20290.1 | 0,11746146 | 2,07714E-06 | no symbol available no full name available chr5:6851695-6853012 REVERSE LENGTH=222 |

| | | | |
|---|---|---|---|
| AT4G39200.2 | 3,866765351 | 2,12593E-06 | no symbol available no full name available chr4:18257464-18258464 FORWARD LENGTH=107 |
| AT1G31812.1 | 7,743499615 | 2,17713E-06 | ACBP6_ ACBP_ AtACBP6 acyl-CoA-binding protein 6_ ACYL-COA-BINDING PROTEIN chr1:11411132-11412099 RE |
| AT1G79920.1 | 1,98588184 | 2,22562E-06 | Hsp70-15_ AtHsp70-15 heat shock protein 70-15 chr1:30059302-30062224 REVERSE LENGTH=736 |
| AT1G48630.1 | 3,393850292 | 2,28975E-06 | RACK1B_ RACK1B_AT receptor for activated C kinase 1B chr1:17981977-17983268 REVERSE LENGTH=326 |
| AT5G51440.1 | 25,75228882 | 2,31278E-06 | HSP23.5 chr5:20891242-20892013 FORWARD LENGTH=210 |
| ATCG00470.1 | 0,620542089 | 2,4153E-06 | ATPE ATP synthase epsilon chain chrc:52265-52663 REVERSE LENGTH=132 |
| AT3G09440.1 | 2,78085246 | 2,47009E-06 | no symbol available no full name available chr3:2903434-2905632 REVERSE LENGTH=649 |
| AT3G12050.2 | 2,878407019 | 2,64848E-06 | no symbol available no full name available chr3:3839289-3841303 FORWARD LENGTH=321 |
| AT1G11840.1 | 2,636934252 | 2,65497E-06 | AtGLYI3_ GLX1_ ATGLX1 glyoxalaseI 3_ glyoxalase I homolog chr1:3996045-3997518 FORWARD LENGTH=283 |
| AT5G44500.1 | 3,807866205 | 2,80101E-06 | no symbol available no full name available chr5:17927505-17928269 FORWARD LENGTH=254 |
| AT2G21580.2 | 3,992046323 | 3,0183E-06 | no symbol available no full name available chr2:9236629-9237510 FORWARD LENGTH=107 |
| AT3G13920.4 | 2,20966378 | 3,23478E-06 | TIF4A1_ EIF4A1_ RH4 eukaryotic translation initiation factor 4A1 chr3:4592635-4594128 REVERSE LENGTH=407 |
| AT5G59720.1 | 31,61254232 | 3,32733E-06 | HSP18.2 heat shock protein 18.2 chr5:24062632-24063117 FORWARD LENGTH=161 |
| AT5G06290.1 | 1,404093953 | 3,35102E-06 | 2-Cys Prx B_ 2CPB 2-cysteine peroxiredoxin B_ 2-CYS PEROXIREDOXIN B chr5:1919380-1921211 FORWARD LENG |
| AT5G27850.1 | 0,452221848 | 3,411E-06 | RPL18C chr5:9873169-9874297 FORWARD LENGTH=187 |
| AT1G18540.1 | 0,505042612 | 4,03295E-06 | no symbol available no full name available chr1:6377448-6378548 REVERSE LENGTH=233 |
| AT2G06850.1 | 1,554003543 | 4,08706E-06 | EXT_ XTH4_ EXGT-A1 ENDOXYLOGLUCAN TRANSFERASE_ endoxyloglucan transferase A1_ xyloglucan endotrans |
| AT5G57870.1 | 1,509074876 | 4,51803E-06 | eIFiso4G1 eukaryotic translation Initiation Factor isoform 4G1 chr5:23439755-23443433 FORWARD LENGTH=780 |
| AT2G38540.1 | 27,45504059 | 4,73105E-06 | ATLTP1_ AtLtpI-4_ LTP1_ LP1 ARABIDOPSIS THALIANA LIPID TRANSFER PROTEIN 1_ lipid transfer protein 1_ l |
| AT1G79930.1 | 1,937239757 | 4,91193E-06 | HSP91_ AtHsp70-14 heat shock protein 91 chr1:30063781-30067067 REVERSE LENGTH=831 |
| AT5G20010.1 | 1,199151661 | 5,0469E-06 | RAN-1_ ATRAN1_ RAN1 RAS-RELATED NUCLEAR PROTEIN_ ARABIDOPSIS THALIANA RAS-RELATED NUCI |
| AT1G07400.1 | 25,63764732 | 5,30777E-06 | HSP17.8 chr1:2275148-2275621 FORWARD LENGTH=157 |
| AT4G30690.2 | 0,135360003 | 5,54425E-06 | AtINFC-4_ SVR9L_ AtIF3- 4 SVR9-LIKE1_ Initiation factor 3-4 chr4:14960742-14962328 FORWARD LENGTH=253 |
| AT3G14210.1 | 0,412984889 | 5,63236E-06 | ESM1 epithiospecifier modifier 1 chr3:4729886-4731562 FORWARD LENGTH=392 |
| AT3G58610.1 | 1,704016872 | 5,9892E-06 | no symbol available no full name available chr3:21671561-21674639 FORWARD LENGTH=591 |
| AT5G47210.1 | 3,831945359 | 6,10749E-06 | no symbol available no full name available chr5:19169222-19171012 REVERSE LENGTH=357 |
| AT5G23740.1 | 1,605304615 | 6,13021E-06 | RPS11-BETA ribosomal protein S11-beta chr5:8008251-8009330 REVERSE LENGTH=159 |
| AT1G72730.1 | 1,562681967 | 6,94753E-06 | no symbol available no full name available chr1:27378040-27379593 REVERSE LENGTH=414 |
| AT2G43030.1 | 0,476684093 | 7,25945E-06 | PRPL3 plastid ribosomal proteins of the 50S subunit chr2:17894898-17895713 FORWARD LENGTH=271 |
| AT1G41880.1 | 0,149232883 | 7,40783E-06 | no symbol available no full name available chr1:15651585-15652427 REVERSE LENGTH=111 |
| AT2G21410.1 | 4,687160241 | 7,73463E-06 | VHA-A2 vacuolar proton ATPase A2 chr2:9162703-9168141 FORWARD LENGTH=821 |
| AT5G19820.1 | 2,331374559 | 7,96024E-06 | IMB3_ KETCH1_ EMB2734 EMBRYO DEFECTIVE 2734_ (karyopherin enabling the transport of the cytoplasmic HYL1 |
| AT5G66760.1 | 0,335409802 | 8,10942E-06 | SDH1-1 succinate dehydrogenase 1-1 chr5:26653776-26657224 FORWARD LENGTH=634 |
| AT1G65350.1 | 3,776551893 | 8,85956E-06 | UBQ13 ubiquitin 13 chr1:24272518-24277275 REVERSE LENGTH=319 |
| AT4G30190.1 | 1,82258996 | 8,87957E-06 | HA2_ PMA2_ AtHA2_ AHA2 H(+)-ATPase 2_ PLASMA MEMBRANE PROTON ATPASE 2 chr4:14770820-14775920 |
| AT2G20990.1 | 4,404852451 | 9,00002E-06 | NTMC2T1.1_ ATSYTA_ SYT1_ NTMC2TYPE1.1_ AtSYT1_ SYTA SYNAPTOTAGMIN 1_ synaptotagmin A_ ARABI |
| AT5G52470.1 | 2,381907615 | 9,27149E-06 | ATFIB1_ FBR1_ FIB1_ SKIP7_ ATFBR1 fibrillarin 1_ FIBRILLARIN 1_ SKP1/ASK1-INTERACTING PROTEIN chr5:2 |
| AT1G19670.1 | 0,390764059 | 9,32757E-06 | CLH1_ ATCLH1_ CORI1_ ATHCOR1 chlorophyllase 1_ CORONATINE-INDUCED PROTEIN 1 chr1:6803796-680492 |
| AT5G06870.1 | 0,495058366 | 9,34212E-06 | PGIP2_ ATPGIP2 ARABIDOPSIS POLYGALACTURONASE INHIBITING PROTEIN 2_ polygalacturonase inhibiting p |

| | | | |
|---|---|---|---|
| AT1G20260.1 | 1,893473428 | 9,45573E-06 | AtVAB3_ VAB3 V-ATPase B subunit 3 chr1:7016971-7020290 FORWARD LENGTH=487 |
| AT3G46520.1 | 2,127052841 | 9,6197E-06 | ACT12 actin-12 chr3:17128567-17129981 FORWARD LENGTH=377 |
| AT4G34670.1 | 0,683474142 | 1,09428E-05 | no symbol available no full name available chr4:16548724-16550222 FORWARD LENGTH=262 |
| AT3G49910.1 | 0,242186976 | 1,14383E-05 | no symbol available no full name available chr3:18504311-18504751 FORWARD LENGTH=146 |
| AT5G09590.1 | 3,935428233 | 1,21338E-05 | MTHSC70-2_ HSC70-5 mitochondrial HSO70 2_ HEAT SHOCK COGNATE chr5:2975721-2978508 FORWARD LENG |
| AT3G49010.4 | 0,07251659 | 1,40157E-05 | BBC1_ RSU2_ ATBBC1 40S RIBOSOMAL PROTEIN_ breast basic conserved 1 chr3:18166971-18168047 REVERSE L |
| AT1G07920.1 | 1,214307087 | 1,47576E-05 | ELONGATION FACTOR-TU FAMILY chr1:2455559-2457001 FORWARD LENGTH=449 |
| ATCG00830.1 | 0,033452635 | 1,48401E-05 | RPL2.1 ribosomal protein L2 chrc:84337-85843 REVERSE LENGTH=274 |
| AT3G08580.1 | 1,535724785 | 1,5098E-05 | AAC1 ADP/ATP carrier 1 chr3:2605706-2607030 REVERSE LENGTH=381 |
| AT1G59860.1 | 11,56228214 | 1,60248E-05 | HSP17.6A chr1:22031474-22031941 FORWARD LENGTH=155 |
| AT1G43170.1 | 0,044370271 | 1,6268E-05 | emb2207_ RP1_ ARP1_ RPL3A embryo defective 2207_ ribosomal protein 1 chr1:16266992-16268631 FORWARD LEN |
| AT3G09200.2 | 1,380234 | 1,68498E-05 | no symbol available no full name available chr3:2823364-2825020 REVERSE LENGTH=287 |
| AT5G15450.1 | 3,197278648 | 1,75109E-05 | APG6_ AtCLPB3_ CLPB3_ CLPB-P casein lytic proteinase B3_ CASEIN LYTIC PROTEINASE B-P_ ALBINO AND PA |
| AT3G62870.1 | 0,366179101 | 1,83839E-05 | no symbol available no full name available chr3:23242862-23244273 REVERSE LENGTH=256 |
| AT2G30200.1 | 1,711382588 | 1,84043E-05 | EMB3147_ MCAT_ MCAMT EMBRYO DEFECTIVE 3147_ malonyl CoA-ACP malonyltransferase chr2:12883162-1288 |
| AT3G04920.2 | 0,14188311 | 1,85116E-05 | no symbol available no full name available chr3:1360989-1361719 FORWARD LENGTH=112 |
| AT1G01100.1 | 1,701620179 | 1,8674E-05 | RPP1A_ RPP1.1 60S acidic ribosomal protein P1-1_ RPP1 co-orthologous gene 1 chr1:50284-50954 REVERSE LENGTH |
| AT1G62180.1 | 3,260265724 | 1,88894E-05 | APSR_ PRH43_ PRH_ ATAPR2_ APR2 ADENOSINE-5'-PHOSPHOSULFATE REDUCTASE_ 3'-PHOSPHOADENOSI |
| AT1G68560.1 | 0,612426604 | 1,93877E-05 | XYL1_ GH31_ TRG1_ ATXYL1_ AXY3 altered xyloglucan 3_ ALPHA-XYLOSIDASE 1_ alpha-xylosidase 1_ thermoinh |
| AT5G47890.1 | 4,562293913 | 1,95874E-05 | no symbol available no full name available chr5:19388806-19390409 FORWARD LENGTH=97 |
| AT1G74260.1 | 1,633004402 | 1,98072E-05 | PUR4 purine biosynthesis 4 chr1:27923005-27927764 REVERSE LENGTH=1407 |
| AT1G55060.1 | 2,289791025 | 2,02058E-05 | UBQ12 ubiquitin 12 chr1:20549533-20550225 FORWARD LENGTH=230 |
| AT1G47128.1 | 0,481484413 | 2,0288E-05 | RD21_ RD21A responsive to dehydration 21A_ responsive to dehydration 21 chr1:17283139-17285609 REVERSE LENG |
| AT3G25230.1 | 4,739370872 | 2,07191E-05 | FKBP62_ ROF1_ ATFKBP62 rotamase FKBP 1_ FK506 BINDING PROTEIN 62 chr3:9188257-9191137 FORWARD LE |
| AT4G00100.1 | 0,188314171 | 2,18275E-05 | RPS13_ RPS13A_ PFL2_ ATRPS13A ribosomal protein S13A_ POINTED FIRST LEAF 2 chr4:37172-38123 FORWARD |
| ATCG00900.1 | 0,324530841 | 2,22425E-05 | RPS7_ RPS7.1 CHLOROPLAST RIBOSOMAL PROTEIN S7 chrc:97478-97945 REVERSE LENGTH=155 |
| AT1G06000.1 | 2,095491079 | 2,26734E-05 | UGT89C1 chr1:1820495-1821802 REVERSE LENGTH=435 |
| AT4G24280.1 | 1,337022611 | 2,29255E-05 | cpHsc70-1 chloroplast heat shock protein 70-1 chr4:12590094-12593437 FORWARD LENGTH=718 |
| AT2G34480.1 | 0,046315141 | 2,29318E-05 | L18aB_ RPL18aB chr2:14532916-14534161 REVERSE LENGTH=178 |
| AT1G55490.1 | 1,334523504 | 2,30871E-05 | Cpn60beta1_ LEN1_ CPNB1_ CPN60B chaperonin-60beta1_ LESION INITIATION 1_ chaperonin 60 beta chr1:2071571 |
| AT4G11820.1 | 2,211080778 | 2,37546E-05 | MVA1_ FKP1_ HMGS FLAKY POLLEN 1_ HYDROXYMETHYLGLUTARYL-COA SYNTHASE chr4:7109124-71112 |
| AT4G16830.2 | 1,818678394 | 2,42664E-05 | AtRGGA chr4:9470979-9472308 FORWARD LENGTH=265 |
| AT5G56500.1 | 0,438891216 | 2,47212E-05 | CPNB3_ Cpn60beta3 chaperonin-60beta3 chr5:22874058-22876966 FORWARD LENGTH=597 |
| AT2G17360.1 | 0,384650588 | 2,63478E-05 | no symbol available no full name available chr2:7546598-7548138 FORWARD LENGTH=261 |
| AT3G08590.1 | 2,309119065 | 2,64782E-05 | iPGAM2 "2_3-biphosphoglycerate-independent phosphoglycerate mutase 2" chr3:2608683-2611237 REVERSE LENGTH= |
| AT5G42980.1 | 1,52350904 | 2,66291E-05 | ATH3_ TRX3_ TRXH3_ ATTRX3_ ATTRXH3 THIOREDOXIN H3_ thioredoxin 3_ thioredoxin H-type 3 chr5:1724277 |
| AT3G60770.1 | 0,202013107 | 2,6722E-05 | no symbol available no full name available chr3:22460525-22461656 REVERSE LENGTH=151 |
| ATCG00790.1 | 0,209472444 | 2,67642E-05 | RPL16 ribosomal protein L16 chrc:81189-82652 REVERSE LENGTH=135 |
| AT5G13650.1 | 0,391853916 | 2,68646E-05 | SVR3 SUPPRESSOR OF VARIEGATION 3 chr5:4397821-4402364 FORWARD LENGTH=675 |

| | | | |
|---|---|---|---|
| AT2G37190.1 | 2,596307366 | 2,72656E-05 | no symbol available no full name available chr2:15619559-15620059 REVERSE LENGTH=166 |
| AT4G22240.1 | 2,059226967 | 2,78505E-05 | FBN1b fibrillin 1b chr4:11766090-11767227 REVERSE LENGTH=310 |
| AT1G75350.1 | 0,708345812 | 2,82383E-05 | emb2184 embryo defective 2184 chr1:28272163-28272687 FORWARD LENGTH=144 |
| AT2G37620.1 | 1,930780739 | 2,87279E-05 | ACT1_ AAc1 ARABIDOPSIS ACTIN 1_ actin 1 chr2:15779761-15781241 FORWARD LENGTH=377 |
| AT4G09010.2 | 0,621899519 | 2,8853E-05 | APX4_ TL29 ascorbate peroxidase 4_ thylakoid lumen 29 chr4:5777502-5779064 REVERSE LENGTH=284 |
| AT5G01410.1 | 2,004347925 | 2,9492E-05 | PDX1_ ATPDX1.3_ ATPDX1_ PDX1.3_ RSR4 REDUCED SUGAR RESPONSE 4_ PYRIDOXINE BIOSYNTHESIS 1.3 |
| AT1G20450.1 | 1,79082209 | 3,36414E-05 | LTI45_ ERD10_ LTI29 EARLY RESPONSIVE TO DEHYDRATION 10_ LOW TEMPERATURE INDUCED 45_ LOW |
| AT3G19710.1 | 2,029126041 | 3,45186E-05 | BCAT4 branched-chain aminotransferase4 chr3:6847202-6849429 REVERSE LENGTH=354 |
| AT3G52880.1 | 1,555931501 | 3,46647E-05 | ATMDAR1_ MDAR1 monodehydroascorbate reductase 1 chr3:19601477-19604366 REVERSE LENGTH=434 |
| AT5G47700.1 | 1,692699354 | 3,6293E-05 | RPP1C_ RPP1.3 60S acidic ribosomal protein P1-3_ RPP1 co-orthologous gene 3 chr5:19328019-19328724 REVERSE LE |
| AT2G18960.1 | 1,848836682 | 3,69299E-05 | HA1_ OST2_ AHA1_ PMA H(+)-ATPase 1_ PLASMA MEMBRANE PROTON ATPASE_ OPEN STOMATA 2 chr2:82 |
| AT2G37220.1 | 0,348091129 | 3,75991E-05 | no symbol available no full name available chr2:15634980-15636331 REVERSE LENGTH=289 |
| AT5G17920.1 | 1,185406235 | 3,81297E-05 | ATCIMS_ METS1_ ATMETS_ ATMS1 COBALAMIN-INDEPENDENT METHIONINE SYNTHASE_ methionine synth |
| AT1G77590.2 | 2,525126195 | 3,94075E-05 | LACS9 long chain acyl-CoA synthetase 9 chr1:29148501-29151234 REVERSE LENGTH=545 |
| AT2G24020.1 | 2,365772287 | 3,96079E-05 | STIC2 Suppressor of TIC40 2 chr2:10217869-10219269 REVERSE LENGTH=182 |
| AT2G21330.1 | 1,193313772 | 3,98413E-05 | FBA1_ AtFBA1 fructose-bisphosphate aldolase 1 chr2:9128416-9130152 REVERSE LENGTH=399 |
| AT4G27670.1 | 72,72989355 | 4,48156E-05 | HSP21 heat shock protein 21 chr4:13819048-13819895 REVERSE LENGTH=227 |
| AT1G07320.3 | 0,501964604 | 4,55902E-05 | RPL4_ PRPL4_ EMB2784 plastid ribosomal protein L4_ ribosomal protein L4_ EMBRYO DEFECTIVE 2784 chr1:22491 |
| AT2G42600.1 | 1,292064119 | 4,5605E-05 | ATPPC2_ PPC2 phosphoenolpyruvate carboxylase 2 chr2:17734541-17738679 REVERSE LENGTH=963 |
| AT2G30490.1 | 1,638746803 | 4,65531E-05 | REF3_ CYP73A5_ C4H_ ATC4H CINNAMATE 4-HYDROXYLASE_ REDUCED EPRDERMAL FLUORESCENCE 3_ |
| AT1G23740.1 | 1,256967799 | 4,73979E-05 | AOR alkenal/one oxidoreductase chr1:8398245-8399656 REVERSE LENGTH=386 |
| AT3G26060.1 | 1,569642474 | 4,79258E-05 | ATPRX Q_ PRXQ peroxiredoxin Q chr3:9524807-9526123 FORWARD LENGTH=216 |
| AT1G13440.2 | 1,406468253 | 4,82787E-05 | GAPC2_ GAPC-2 glyceraldehyde-3-phosphate dehydrogenase C2_ GLYCERALDEHYDE-3-PHOSPHATE DEHYDROGI |
| AT4G31180.1 | 2,480764775 | 4,91529E-05 | IBI1 impaired in BABA-induced disease immunity 1 chr4:15156696-15159362 FORWARD LENGTH=558 |
| AT2G22990.2 | 0,255096359 | 4,92275E-05 | SNG1_ SCPL8 sinapoylglucose 1_ SERINE CARBOXYPEPTIDASE-LIKE 8 chr2:9787071-9789925 FORWARD LENGT |
| AT2G22990.1 | 0,255096359 | 4,92275E-05 | SNG1_ SCPL8 sinapoylglucose 1_ SERINE CARBOXYPEPTIDASE-LIKE 8 chr2:9786393-9789925 FORWARD LENGT |
| AT4G18480.1 | 0,755330672 | 4,92821E-05 | CHL11_ CH-42_ LOST1_ CH42_ CHLI-1_ CHLI1 low temperature with open-stomata 1_ CHLORINA 42 chr4:10201897- |
| AT3G09790.1 | 1,905565513 | 4,93399E-05 | UBQ8 ubiquitin 8 chr3:3004111-3006006 REVERSE LENGTH=631 |
| AT1G54050.1 | 45,76273828 | 4,94687E-05 | HSP17.4B chr1:20179558-20180122 REVERSE LENGTH=155 |
| AT5G28060.1 | 0,23369253 | 5,01355E-05 | RPS24B chr5:10069791-10070792 REVERSE LENGTH=133 |
| AT3G11250.1 | 0,757731109 | 5,04646E-05 | no symbol available no full name available chr3:3521453-3522826 FORWARD LENGTH=323 |
| AT1G09590.1 | 0,126278511 | 5,18684E-05 | no symbol available no full name available chr1:3106549-3107606 FORWARD LENGTH=164 |
| AT1G16470.1 | 1,287777679 | 5,36792E-05 | PAB1 proteasome subunit PAB1 chr1:5623122-5625439 FORWARD LENGTH=235 |
| AT3G11830.1 | 1,856211386 | 5,43286E-05 | CCT7 Chaperonin containing T-complex polypeptide-1 subunit 7 chr3:3732734-3736156 FORWARD LENGTH=557 |
| AT3G15730.1 | 2,352278294 | 5,48387E-05 | PLD_ PLDALPHA1 phospholipase D alpha 1 chr3:5330835-5333474 FORWARD LENGTH=810 |
| AT1G14980.1 | 7,36050894 | 5,6933E-05 | CPN10 chaperonin 10 chr1:5165930-5166654 REVERSE LENGTH=98 |
| AT3G02530.1 | 1,883839533 | 5,72589E-05 | CCT6-2 Chaperonin containing T-complex polypeptide-1 subunit 6-2 chr3:528806-532457 REVERSE LENGTH=535 |
| AT2G29450.1 | 1,817303462 | 5,78243E-05 | AT103-1A_ ATGSTU1_ ATGSTU5_ GSTU5 glutathione S-transferase tau 5_ ARABIDOPSIS THALIANA GLUTATHIO |
| AT5G59880.2 | 0,603911046 | 5,85185E-05 | ADF3 actin depolymerizing factor 3 chr5:24120382-24121628 FORWARD LENGTH=124 |

| | | | |
|---|---|---|---|
| AT3G03710.1 | 1,696859535 | 5,91216E-05 | PNP_ RIF10_ PDE326 PIGMENT DEFECTIVE 326_ POLYNUCLEOTIDE PHOSPHORYLASE_ resistant to inhibition [illegible] |
| AT1G74920.2 | 2,402428644 | 5,95165E-05 | ALDH10A8 aldehyde dehydrogenase 10A8 chr1:28139175-28142573 REVERSE LENGTH=496 |
| AT3G08740.1 | 0,50711495 | 5,9856E-05 | no symbol available no full name available chr3:2654788-2656154 REVERSE LENGTH=236 |
| AT3G22110.1 | 1,278476344 | 6,03468E-05 | PAC1 20S proteasome alpha subunit C1 chr3:7792819-7793571 REVERSE LENGTH=250 |
| AT1G31230.1 | 2,496237966 | 6,15092E-05 | AK-HSDH I_ AK-HSDH ASPARTATE KINASE-HOMOSERINE DEHYDROGENASE_ aspartate kinase-homoserine del |
| AT3G59970.1 | 2,744620231 | 6,29244E-05 | MTHFR1 methylenetetrahydrofolate reductase 1 chr3:22151303-22153412 FORWARD LENGTH=421 |
| AT5G66570.1 | 0,788273222 | 6,45727E-05 | OEE1_ PSBO-1_ MSP-1_ OE33_ OEE33_ PSBO1 PS II OXYGEN-EVOLVING COMPLEX 1_ OXYGEN EVOLVING [illegible] |
| AT5G26780.1 | 0,626953764 | 6,51864E-05 | SHM2 serine hydroxymethyltransferase 2 chr5:9418299-9421725 FORWARD LENGTH=517 |
| AT5G13850.1 | 0,451090432 | 6,63226E-05 | NACA3 nascent polypeptide-associated complex subunit alpha-like protein 3 chr5:4471361-4472676 FORWARD LENGTI |
| AT2G14880.1 | 5,910019199 | 6,79291E-05 | SWIB2 chr2:6393686-6394841 REVERSE LENGTH=141 |
| AT4G10320.1 | 2,39943155 | 7,39878E-05 | no symbol available no full name available chr4:6397526-6404509 REVERSE LENGTH=1190 |
| AT3G52300.1 | 1,605461688 | 7,45498E-05 | ATPQ_ ATPd "ATP synthase D chain_ mitochondrial" chr3:19396689-19398119 FORWARD LENGTH=168 |
| AT1G66580.1 | 0,107282765 | 7,54484E-05 | SAG24_ RPL10C senescence associated gene 24_ ribosomal protein L10 C chr1:24839208-24840439 FORWARD LENGT |
| AT1G56330.1 | 1,708079617 | 7,63397E-05 | ATSARA1B_ SAR1B_ SAR1_ ATSAR1B_ ATSAR1 SECRETION-ASSOCIATED RAS 1_ ARABIDOPSIS THALIANA |
| AT3G24830.1 | 0,139527272 | 7,6442E-05 | no symbol available no full name available chr3:9064613-9065871 FORWARD LENGTH=206 |
| AT5G03300.1 | 1,578916271 | 7,7425E-05 | ADK2 adenosine kinase 2 chr5:796573-798997 FORWARD LENGTH=345 |
| AT5G14040.1 | 1,507492124 | 8,27423E-05 | MPT3_ PHT3;1 phosphate transporter 3;1_ mitochondrial phosphate transporter 3 chr5:4531059-4532965 REVERSE LEN[illegible] |
| AT3G52380.1 | 0,402223687 | 8,38194E-05 | PDE322_ CP33 PIGMENT DEFECTIVE 322_ chloroplast RNA-binding protein 33 chr3:19421619-19422855 FORWARD |
| AT1G78370.1 | 1,362281824 | 8,46001E-05 | ATGSTU20_ GSTU20 glutathione S-transferase TAU 20 chr1:29484428-29485204 REVERSE LENGTH=217 |
| AT1G18210.1 | 0,375904308 | 8,65915E-05 | no symbol available no full name available chr1:6268273-6268785 REVERSE LENGTH=170 |
| AT5G02500.1 | 1,516032199 | 8,68949E-05 | AtHsp70-1_ HSP70-1_ HSC70-1_ HSC70_ AT-HSC70-1 ARABIDOPSIS THALIANA HEAT SHOCK COGNATE PROT |
| AT4G25200.1 | 7,066416544 | 8,70867E-05 | ATHSP23.6-MITO_ HSP23.6-MITO mitochondrion-localized small heat shock protein 23.6 chr4:12917089-12917858 FOI |
| AT3G48930.1 | 0,292281294 | 8,71692E-05 | EMB1080 embryo defective 1080 chr3:18141017-18142189 REVERSE LENGTH=160 |
| AT4G15802.1 | 2,467220828 | 8,77057E-05 | AtHSBP_ HSBP Arabidopsis thaliana heat shock factor binding protein_ heat shock factor binding protein chr4:8986864-8[illegible] |
| AT2G28000.1 | 1,922931832 | 8,93179E-05 | ARC2_ CH-CPN60A_ SLP_ CPN60A_ Cpn60alpha1_ CPNA1 SCHLEPPERLESS_ chaperonin-60alpha1_ CHLOROPLA |
| AT2G31610.1 | 1,325506006 | 9,00414E-05 | no symbol available no full name available chr2:13450384-13451669 FORWARD LENGTH=250 |
| AT2G23350.1 | 1,511720695 | 9,20507E-05 | PABP4_ PAB4 POLY(A) BINDING PROTEIN 4_ poly(A) binding protein 4 chr2:9943209-9946041 FORWARD LENGT |
| AT5G38480.1 | 1,480148231 | 9,25222E-05 | GRF3_ RCI1 general regulatory factor 3 chr5:15410277-15411285 FORWARD LENGTH=255 |
| AT1G15690.1 | 1,702868357 | 9,3163E-05 | AtVHP1;1_ AtAVP1_ ATAVP3_ AVP-3_ AVP1_ FUGU5_ VHP1 ARABIDOPSIS THALIANA V-PPASE 3_ FUGU 5 c[illegible] |
| AT5G20890.1 | 2,285811192 | 9,34294E-05 | CCT2 Chaperonin containing T-complex polypeptide-1 subunit 2 chr5:7087020-7089906 REVERSE LENGTH=527 |
| AT1G06410.1 | 1,953329399 | 9,4214E-05 | ATTPSA_ TPS7_ ATTPS7 TREHALOSE -6-PHOSPHATASE SYNTHASE S7_ trehalose-phosphatase/synthase 7 chr1:19 |
| AT1G18080.1 | 1,522195539 | 9,50939E-05 | RACK1A_AT_ AtRACK1_ SAC53_ ATARCA_ RACK1A RECEPTOR FOR ACTIVATED C KINASE 1 A_ Suppressor [illegible] |
| AT4G35250.1 | 1,302712824 | 9,56659E-05 | HCF244 high chlorophyll fluorescence phenotype 244 chr4:16771401-16773269 REVERSE LENGTH=395 |
| AT3G19820.1 | 3,156341951 | 9,8217E-05 | DWF1_ DIM_ DIM1_ CBB1_ EVE1 ENHANCED VERY-LOW-FLUENCE RESPONSES 1_ DIMINUTIA_ DIMINUTO |
| AT3G16950.1 | 1,473988059 | 9,85922E-05 | ptlpd1_ LPD1 lipoamide dehydrogenase 1 chr3:5786761-5790383 REVERSE LENGTH=570 |
| AT5G11560.1 | 2,357670005 | 0,000100996 | PNET5 chr5:3709734-3713994 REVERSE LENGTH=982 |
| AT5G03630.1 | 1,536564182 | 0,000102447 | MDAR2 chr5:922378-924616 REVERSE LENGTH=435 |
| AT4G27440.1 | 0,314802727 | 0,000103982 | PORB protochlorophyllide oxidoreductase B chr4:13725648-13727107 FORWARD LENGTH=401 |
| AT3G12290.1 | 1,952785608 | 0,000104859 | MTHFD1 methylenetetrahydrofolate dehydrogenase/methenyltetrahydrofolate cyclohydrolase chr3:3919591-3921326 FOR[illegible] |

| | | | |
|---|---|---|---|
| AT2G20580.1 | 2,343136277 | 0,000105942 | ATRPN1A_ RPN1A 26S PROTEASOME REGULATORY SUBUNIT S2 1A_ 26S proteasome regulatory subunit S2 1A c |
| AT5G62670.1 | 2,357712047 | 0,000106804 | HA11_ AHA11 H(+)-ATPase 11 chr5:25159495-25164957 FORWARD LENGTH=956 |
| AT3G61440.1 | 0,831284186 | 0,000107162 | ATCYSC1_ ARATH;BSAS3;1_ CAS-C1_ CYSC1 ß-cyanoalanine synthase C1_ CYSTEINE SYNTHASE C1_ BETA-SU |
| AT2G38230.1 | 1,417329903 | 0,00010804 | ATPDX1.1_ PDX1.1 pyridoxine biosynthesis 1.1_ ARABIDOPSIS THALIANA PYRIDOXINE BIOSYNTHESIS 1.1 chr2 |
| AT1G66240.1 | 1,438892054 | 0,000108705 | AtHMP14_ ATX1_ ATATX1 HEAVY METAL ASSOCIATED PROTEIN 14_ homolog of anti-oxidant 1 chr1:24686445- |
| AT2G20610.1 | 1,557248214 | 0,000109929 | RTY1_ RTY_ HLS3_ ALF1_ SUR1 ABERRANT LATERAL ROOT FORMATION 1_ SUPERROOT 1_ ROOTY 1_ RO |
| AT3G08530.1 | 2,314195225 | 0,000112745 | AtCHC2_ CHC2 clathrin heavy chain 2 chr3:2587171-2595411 REVERSE LENGTH=1703 |
| AT5G23860.1 | 2,308915519 | 0,000114182 | TUB8 tubulin beta 8 chr5:8042962-8044528 FORWARD LENGTH=449 |
| AT1G54020.1 | 0,167196545 | 0,000115773 | no symbol available no full name available chr1:20161805-20162923 REVERSE LENGTH=286 |
| AT3G55440.1 | 1,194152421 | 0,000116008 | ATCTIMC_ CYTOTPI_ TPI CYTOSOLIC ISOFORM TRIOSE PHOSPHATE ISOMERASE_ triosephosphate isomerase_ |
| AT3G54210.1 | 0,089105699 | 0,000118176 | PRPL17 plastid ribosomal proteins of the 50S subunit 17 chr3:20067672-20068385 REVERSE LENGTH=211 |
| AT2G36580.1 | 2,055463762 | 0,000121541 | no symbol available no full name available chr2:15339253-15342781 FORWARD LENGTH=527 |
| AT1G57720.1 | 1,810187551 | 0,000121834 | no symbol available no full name available chr1:21377873-21380114 FORWARD LENGTH=413 |
| AT3G56340.1 | 0,092025417 | 0,00012593 | RPS26e Ribosomal Protein S26e chr3:20892309-20893343 REVERSE LENGTH=130 |
| AT3G12145.1 | 0,235498865 | 0,000128499 | FLR1_ FTM4 FLOR1_ FLORAL TRANSITION AT THE MERISTEM4 chr3:3874764-3876075 REVERSE LENGTH=32 |
| AT4G13940.1 | 1,755834725 | 0,000129984 | MEE58_ SAHH1_ EMB1395_ HOG1_ SAH1_ ATSAHH1 EMBRYO DEFECTIVE 1395_ HOMOLOGY-DEPENDENT |
| AT5G13410.1 | 0,701978306 | 0,000130361 | no symbol available no full name available chr5:4299830-4301706 REVERSE LENGTH=256 |
| AT3G57490.1 | 0,401654588 | 0,000131224 | no symbol available no full name available chr3:21279824-21280887 REVERSE LENGTH=276 |
| AT1G32990.1 | 0,529611678 | 0,0001347 | PRPL11 plastid ribosomal protein l11 chr1:11955827-11957139 FORWARD LENGTH=222 |
| AT1G14320.2 | 0,135142907 | 0,000134865 | RPL10A_ SAC52_ RPL10 SUPPRESSOR OF ACAULIS 52_ ribosomal protein L10_ ribosomal protein L10 A chr1:48886 |
| AT1G09640.1 | 1,810704714 | 0,000136664 | no symbol available no full name available chr1:3120162-3122152 FORWARD LENGTH=414 |
| AT2G33040.1 | 1,817968988 | 0,000140457 | ATP3 gamma subunit of Mt ATP synthase chr2:14018978-14021047 REVERSE LENGTH=325 |
| AT5G09650.1 | 0,606132026 | 0,000142103 | PPa6_ AtPPa6 pyrophosphorylase 6 chr5:2991331-2993117 REVERSE LENGTH=300 |
| AT3G25520.2 | 0,501981956 | 0,000145755 | RPL5A_ ATL5_ PGY3_ OLI5 ribosomal protein L5_ RIBOSOMAL PROTEIN L5 A_ PIGGYBACK3_ OLIGOCELLULA |
| AT4G37910.1 | 1,932794254 | 0,000146195 | mtHsc70-1 mitochondrial heat shock protein 70-1 chr4:17825368-17828099 REVERSE LENGTH=682 |
| AT3G44300.1 | 1,626249089 | 0,0001499 | AtNIT2_ NIT2 nitrilase 2 chr3:15983351-15985172 FORWARD LENGTH=339 |
| AT4G37925.1 | 0,494512369 | 0,000150402 | NDH-M_ NdhM subunit NDH-M of NAD(P)H:plastoquinone dehydrogenase complex_ NADH dehydrogenase-like comple |
| AT5G59370.1 | 1,369249279 | 0,000151813 | ACT4 actin 4 chr5:23950109-23951586 FORWARD LENGTH=377 |
| AT5G17170.1 | 0,68812907 | 0,000156012 | ENH1 enhancer of sos3-1 chr5:5649335-5650975 FORWARD LENGTH=271 |
| AT1G67430.2 | 0,346391277 | 0,000159228 | no symbol available no full name available chr1:25262209-25263627 FORWARD LENGTH=131 |
| AT4G02510.1 | 2,200709435 | 0,000159676 | TOC86_ ATTOC159_ TOC160_ PPI2_ TOC159 translocon at the outer envelope membrane of chloroplasts 159_ PLASTII |
| AT1G07660.1 | 0,687441283 | 0,000161335 | no symbol available no full name available chr1:2369212-2369523 FORWARD LENGTH=103 |
| AT1G03680.1 | 0,634790419 | 0,000166293 | ATHM1_ TRX-M1_ ATM1_ THM1 thioredoxin M-type 1_ THIOREDOXIN M-TYPE 1_ ARABIDOPSIS THIOREDOXI |
| AT3G03250.1 | 1,395299582 | 0,000178442 | AtUGP1_ UGP_ UGP1 UDP-glucose pyrophosphorylase_ UDP-GLUCOSE PYROPHOSPHORYLASE 1 chr3:749761-75 |
| AT3G18130.1 | 0,745071123 | 0,000181816 | RACK1C_ RACK1C_AT receptor for activated C kinase 1C chr3:6211109-6212371 REVERSE LENGTH=326 |
| AT1G49630.1 | 0,745117745 | 0,000184775 | PREP2_ ATPREP2 presequence protease 2 chr1:18368405-18375336 REVERSE LENGTH=1080 |
| AT2G29630.1 | 0,575665185 | 0,000185385 | THIC_ PY PYRIMIDINE REQUIRING_ thiaminC chr2:12667395-12669569 FORWARD LENGTH=644 |
| AT4G29130.1 | 1,66525912 | 0,000187291 | GIN2_ HXK1_ ATHXK1 GLUCOSE INSENSITIVE 2_ hexokinase 1_ ARABIDOPSIS THALIANA HEXOKINASE 1 ch |
| AT4G09010.1 | 1,2409834 | 0,000187323 | APX4_ TL29 ascorbate peroxidase 4_ thylakoid lumen 29 chr4:5777502-5779338 REVERSE LENGTH=349 |

| | | | |
|---|---|---|---|
| AT2G28790.1 | 0,420742555 | 0,000187761 | no symbol available no full name available chr2:12354664-12355413 REVERSE LENGTH=249 |
| AT4G33090.1 | 1,571665286 | 0,000190067 | APM1_ ATAPM1 AMINOPEPTIDASE M1_ aminopeptidase M1 chr4:15965915-15970418 REVERSE LENGTH=879 |
| AT5G11420.1 | 1,739868914 | 0,000190657 | no symbol available no full name available chr5:3644655-3646991 FORWARD LENGTH=366 |
| AT5G09900.1 | 1,639936296 | 0,000192627 | EMB2107_ MSA_ RPN5A MARIPOSA_ EMBRYO DEFECTIVE 2107_ REGULATORY PARTICLE NON-ATPASE SU |
| AT2G41680.1 | 0,738965734 | 0,000196024 | NTRC NADPH-dependent thioredoxin reductase C chr2:17376349-17379028 REVERSE LENGTH=529 |
| AT3G02730.1 | 0,71765408 | 0,000196137 | TRXF1_ ATF1 thioredoxin F-type 1 chr3:588570-589591 REVERSE LENGTH=178 |
| AT2G36460.1 | 1,649744612 | 0,000199007 | FBA6 fructose-bisphosphate aldolase 6 chr2:15296929-15298387 REVERSE LENGTH=358 |
| AT1G48860.1 | 1,97486612 | 0,00020035 | EPSPS 5-enolpyruvylshikimate-3-phosphate synthase chr1:18068892-18071331 REVERSE LENGTH=521 |
| AT5G67030.2 | 1,629699297 | 0,00020264 | LOS6_ NPQ2_ ZEP_ ABA1_ ATABA1_ ATZEP_ IBS3 ABA DEFICIENT 1_ IMPAIRED IN BABA-INDUCED STERIL |
| AT5G43940.1 | 0,894133649 | 0,000205219 | ADH2_ HOT5_ ATGSNOR1_ PAR2_ GSNOR PARAQUAT RESISTANT 2_ S-NITROSOGLUTATHIONE REDUCTAS |
| AT5G07350.1 | 1,420199974 | 0,000208181 | Tudor1_ TSN1_ AtTudor1 TUDOR-SN protein 1_ Arabidopsis thaliana TUDOR-SN protein 1 chr5:2320344-2324892 REV |
| AT4G24190.1 | 1,727260238 | 0,000208453 | SHD_ HSP90_ HSP90.7_ AtHsp90-7_ AtHsp90.7 SHEPHERD_ HEAT SHOCK PROTEIN 90.7_ HEAT SHOCK PROTE |
| AT3G19760.1 | 6,351014827 | 0,000210457 | EIF4A-III_ RH2 eukaryotic initiation factor 4A-III chr3:6863790-6866242 FORWARD LENGTH=408 |
| AT3G48000.1 | 0,825123103 | 0,000213072 | ALDH2B4_ ALDH2A_ ALDH2 aldehyde dehydrogenase 2A_ aldehyde dehydrogenase 2B4_ aldehyde dehydrogenase 2 ch |
| AT1G30120.1 | 0,428246695 | 0,000218014 | PDH-E1 BETA pyruvate dehydrogenase E1 beta chr1:10584350-10586477 REVERSE LENGTH=406 |
| AT1G64520.1 | 1,661145735 | 0,000218132 | RPN12a regulatory particle non-ATPase 12A chr1:23956459-23958120 FORWARD LENGTH=267 |
| AT3G13930.1 | 1,197153608 | 0,000218609 | mtE2-2 mitochondrial pyruvate dehydrogenase subunit 2-2 chr3:4596240-4600143 FORWARD LENGTH=539 |
| AT5G27640.1 | 2,152761402 | 0,000225064 | ATTIF3B1_ TIF3B1_ EIF3B_ ATEIF3B-1_ EIF3B-1 ARABIDOPSIS THALIANA TRANSLATION INITIATION FACT |
| AT3G46430.1 | 1,845260532 | 0,000226501 | AtMtATP6 chr3:17087687-17088497 FORWARD LENGTH=55 |
| AT5G28840.1 | 1,27305943 | 0,000227711 | GME GDP-D-mannose 3 chr5:10862472-10864024 REVERSE LENGTH=377 |
| AT3G50820.1 | 0,831386135 | 0,000236255 | OEC33_ PSBO-2_ PSBO2 photosystem II subunit O-2_ OXYGEN EVOLVING COMPLEX SUBUNIT 33 KDA_ PHOTC |
| AT2G10940.1 | 1,486741301 | 0,000238541 | no symbol available no full name available chr2:4311160-4312035 REVERSE LENGTH=291 |
| AT5G07440.3 | 2,670152431 | 0,000243091 | GDH2 glutamate dehydrogenase 2 chr5:2356153-2357546 FORWARD LENGTH=309 |
| AT3G04720.1 | 0,450859507 | 0,000245936 | HEL_ PR-4_ AtPR4_ PR4 HEVEIN-LIKE_ pathogenesis-related 4 chr3:1285691-1286531 REVERSE LENGTH=212 |
| AT4G34200.1 | 2,087747538 | 0,000250753 | EDA9_ PGDH1 phosphoglycerate dehydrogenase 1_ embryo sac development arrest 9 chr4:16374041-16376561 REVERSI |
| AT1G03230.1 | 1,445210539 | 0,000252387 | SAP1 secreted aspartic protease 1 chr1:790110-791414 FORWARD LENGTH=434 |
| AT3G46230.1 | 9,43435472 | 0,000252789 | HSP17.4_ ATHSP17.4 heat shock protein 17.4_ ARABIDOPSIS THALIANA HEAT SHOCK PROTEIN 17.4 chr3:169842 |
| AT5G08530.1 | 1,258999111 | 0,00025304 | CI51_ NDUFV1 51 kDa subunit of complex I chr5:2759848-2761726 REVERSE LENGTH=486 |
| AT3G18060.1 | 1,551654318 | 0,000257399 | no symbol available no full name available chr3:6183880-6186788 FORWARD LENGTH=609 |
| AT2G21660.1 | 0,451587278 | 0,000257461 | RBGA3_ GR-RBP7_ CCR2_ ATGRP7_ GRP7_ SRBP1 RNA-binding glycine-rich protein A3_ SMALL RNA-BINDING I |
| AT2G45470.1 | 0,60697275 | 0,000262067 | AGP8_ FLA8 ARABINOGALACTAN PROTEIN 8_ FASCICLIN-like arabinogalactan protein 8 chr2:18742797-1874405 |
| AT3G63460.2 | 2,168168392 | 0,000262092 | SEC31B chr3:23431009-23437241 REVERSE LENGTH=1102 |
| AT5G40770.1 | 1,854537324 | 0,000262117 | ATPHB3_ PHB3_ EER3 prohibitin 3 chr5:16315589-16316621 REVERSE LENGTH=277 |
| AT4G33680.1 | 1,255979752 | 0,000262148 | AGD2 ABERRANT GROWTH AND DEATH 2_ ARF-GAP domain 2 chr4:16171847-16174630 REVERSE LENGTH=4 |
| AT1G78900.1 | 1,52336058 | 0,000272711 | VHA-A vacuolar ATP synthase subunit A chr1:29660463-29664575 FORWARD LENGTH=623 |
| AT4G24780.1 | 0,50065146 | 0,000273138 | PLL19 chr4:12770631-12772227 REVERSE LENGTH=408 |
| AT2G04030.1 | 1,280135073 | 0,000274311 | CR88_ EMB1956_ AtHsp90.5_ HSP90C_ AtHsp90C_ Hsp88.1_ HSP90.5 HEAT SHOCK PROTEIN 90.5_ EMBRYO DE |
| AT3G07390.1 | 0,699617102 | 0,000276965 | AIR12 Auxin-Induced in Root cultures 12 chr3:2365452-2366273 FORWARD LENGTH=273 |
| AT1G20020.1 | 0,747571663 | 0,000278196 | LFNR2_ FNR2_ ATLFNR2 leaf-type chloroplast-targeted FNR 2_ ferredoxin-NADP(+)-oxidoreductase 2_ LEAF FNR 2 cl |

| | | | |
|---|---|---|---|
| AT2G04039.1 | 0,595860263 | 0,000280764 | NdhV chr2:1333464-1334438 FORWARD LENGTH=199 |
| AT4G02520.1 | 0,701747205 | 0,000284208 | ATGSTF2_ GSTF2_ ATPM24.1_ GST2_ ATPM24 glutathione S-transferase PHI 2 chr4:1110673-1111531 REVERSE LE |
| AT3G07110.1 | 0,176786013 | 0,000292615 | no symbol available no full name available chr3:2252092-2253332 FORWARD LENGTH=206 |
| AT2G01250.1 | 0,821459619 | 0,000295108 | RPL7B chr2:132943-134264 REVERSE LENGTH=242 |
| AT4G34450.1 | 2,497862664 | 0,000299728 | gamma2-COP gamma2 Coat Protein chr4:16471956-16476795 FORWARD LENGTH=886 |
| AT3G02560.1 | 0,621894595 | 0,000308379 | no symbol available no full name available chr3:542341-543168 FORWARD LENGTH=191 |
| AT3G26650.1 | 1,208750129 | 0,00031151 | GAPA_ GAPA1_ GAPA-1 glyceraldehyde 3-phosphate dehydrogenase A subunit_ GLYCERALDEHYDE 3-PHOSPHATE |
| AT4G32260.1 | 1,282360347 | 0,000327662 | PDE334 PIGMENT DEFECTIVE 334 chr4:15573859-15574586 REVERSE LENGTH=219 |
| AT3G15356.1 | 0,23065473 | 0,000327782 | no symbol available no full name available chr3:5174603-5175418 REVERSE LENGTH=271 |
| AT1G78820.1 | 0,492650537 | 0,000331526 | no symbol available no full name available chr1:29634401-29635768 REVERSE LENGTH=455 |
| AT5G58070.1 | 2,022405242 | 0,000332476 | TIL_ ATTIL TEMPERATURE-INDUCED LIPOCALIN_ temperature-induced lipocalin chr5:23500512-23501156 REVEI |
| AT4G04640.1 | 1,158645024 | 0,000335653 | ATPC1 chr4:2350761-2351882 REVERSE LENGTH=373 |
| AT1G62660.1 | 0,595692106 | 0,000337225 | VI1 VACUOLAR INVERTASE 1 chr1:23199949-23203515 FORWARD LENGTH=648 |
| AT5G42740.3 | 1,45714022 | 0,000337931 | no symbol available no full name available chr5:17136269-17140622 FORWARD LENGTH=528 |
| AT4G25630.1 | 0,605717833 | 0,000342215 | ATFIB2_ FIB2 fibrillarin 2 chr4:13074239-13076205 FORWARD LENGTH=320 |
| AT3G42050.1 | 1,59761605 | 0,000344176 | VHA-H chr3:14228846-14232228 REVERSE LENGTH=441 |
| AT4G29010.1 | 1,500596442 | 0,000353709 | AIM1 ABNORMAL INFLORESCENCE MERISTEM chr4:14297312-14302016 REVERSE LENGTH=721 |
| AT3G17240.1 | 0,719869734 | 0,000361262 | mtLPD2 lipoamide dehydrogenase 2 chr3:5890278-5892166 REVERSE LENGTH=507 |
| AT5G28540.1 | 1,775311717 | 0,000363932 | BIP1 chr5:10540665-10543274 REVERSE LENGTH=669 |
| AT5G15200.1 | 0,481995159 | 0,000366566 | no symbol available no full name available chr5:4935124-4936334 REVERSE LENGTH=198 |
| AT3G12915.2 | 1,315298503 | 0,000379069 | no symbol available no full name available chr3:4112834-4115708 FORWARD LENGTH=788 |
| AT1G50480.1 | 1,301607588 | 0,000379801 | THFS 10-formyltetrahydrofolate synthetase chr1:18702064-18704687 FORWARD LENGTH=634 |
| AT3G18490.1 | 0,673346108 | 0,000381444 | ASPG1 ASPARTIC PROTEASE IN GUARD CELL 1 chr3:6349090-6350592 REVERSE LENGTH=500 |
| AT4G11600.1 | 1,329550537 | 0,000381731 | GPXL6_ PHGPX_ LSC803_ ATGPX6_ GPX6 glutathione peroxidase 6 chr4:7010021-7011330 REVERSE LENGTH=232 |
| AT5G19770.1 | 1,356507953 | 0,000383132 | TUA3 tubulin alpha-3 chr5:6682761-6684474 REVERSE LENGTH=450 |
| AT5G30510.1 | 1,235710022 | 0,000383244 | ARRPS1_ RPS1_ PRPS1 plastid ribosomal protein S1_ ribosomal protein S1 chr5:11619262-11621223 REVERSE LENGT |
| AT1G12410.1 | 1,592368728 | 0,000385266 | EMB3146_ CLPR2_ CLP2_ NCLPP2 EMBRYO DEFECTIVE 3146_ CLP protease proteolytic subunit 2_ NUCLEAR-EN |
| AT1G07890.1 | 1,350342961 | 0,000387223 | MEE6_ ATAPX01_ ATAPX1_ CS1_ APX1 ascorbate peroxidase 1_ maternal effect embryo arrest 6 chr1:2438005-243943 |
| AT5G10360.1 | 0,122332305 | 0,000390984 | RPS6B_ EMB3010 Ribosomal protein small subunit 6b_ embryo defective 3010 chr5:3258734-3260142 REVERSE LENG |
| AT2G30950.1 | 3,719227023 | 0,000397068 | FTSH2_ VAR2 VARIEGATED 2 chr2:13174692-13177064 FORWARD LENGTH=695 |
| AT1G25490.1 | 1,622134606 | 0,000398867 | EER1_ ATB BETA BETA_ RCN1_ REGA ROOTS CURL IN NPA_ ENHANCED ETHYLENE RESPONSE 1 chr1:8951 |
| AT1G52400.1 | 0,65302988 | 0,000399447 | BGL1_ ATBG1_ BGLU18 A. THALIANA BETA-GLUCOSIDASE 1_ BETA-GLUCOSIDASE HOMOLOG 1_ beta gluc |
| AT4G12400.2 | 9,886654897 | 0,000400067 | Hop3 Hop3 chr4:7338866-7341239 REVERSE LENGTH=558 |
| AT2G44160.1 | 1,633291406 | 0,000406679 | MTHFR2 methylenetetrahydrofolate reductase 2 chr2:18262301-18265185 FORWARD LENGTH=594 |
| AT1G75270.1 | 1,321370175 | 0,000410466 | DHAR2 dehydroascorbate reductase 2 chr1:28250255-28251237 REVERSE LENGTH=213 |
| AT1G54220.1 | 0,522058139 | 0,000410739 | mtE2-3 mitochondrial pyruvate dehydrogenase subunit 2-3 chr1:20246460-20250208 REVERSE LENGTH=539 |
| AT2G39390.1 | 0,162925582 | 0,000410874 | no symbol available no full name available chr2:16450803-16451762 REVERSE LENGTH=123 |
| AT2G38750.2 | 0,373839174 | 0,000418482 | AtANN4_ ANNAT4 annexin 4 chr2:16196582-16198273 REVERSE LENGTH=302 |
| AT4G17090.1 | 1,760850776 | 0,000422321 | CT-BMY_ BMY8_ AtBAM3_ BAM3 BETA-AMYLASE 8_ BETA-AMYLASE 3_ chloroplast beta-amylase chr4:960526 |

| | | | |
|---|---|---|---|
| AT1G68010.1 | 1,258000384 | 0,000423967 | HPR_ ATHPR1 hydroxypyruvate reductase chr1:25493418-25495720 FORWARD LENGTH=386 |
| AT3G29360.1 | 2,107640825 | 0,000424869 | UGD2 UDP-glucose dehydrogenase 2 chr3:11267375-11268817 REVERSE LENGTH=480 |
| AT1G65260.1 | 2,43080314 | 0,000429965 | VIPP1_ PTAC4_ IM30 VESICLE-INDUCING PROTEIN IN PLASTIDS 1_ plastid transcriptionally active 4 chr1:242363[illegible] |
| ATCG00780.1 | 0,619937776 | 0,000431753 | RPL14 ribosomal protein L14 chrc:80696-81064 REVERSE LENGTH=122 |
| AT5G14320.1 | 4,173550446 | 0,000435216 | EMB3137 EMBRYO DEFECTIVE 3137 chr5:4617839-4618772 REVERSE LENGTH=169 |
| AT2G36620.1 | 0,094945196 | 0,000435454 | RPL24A ribosomal protein L24 chr2:15350548-15351819 REVERSE LENGTH=164 |
| AT1G24510.3 | 2,119045363 | 0,000438607 | CCT5 Chaperonin containing T-complex polypeptide-1 subunit 5 chr1:8685504-8687802 REVERSE LENGTH=482 |
| AT1G21065.1 | 4,523654085 | 0,000445846 | no symbol available no full name available chr1:7374210-7375644 FORWARD LENGTH=217 |
| AT3G02880.1 | 2,864704505 | 0,000451005 | KIN7 Kinase 7 chr3:634819-636982 FORWARD LENGTH=627 |
| AT5G55220.1 | 0,642816475 | 0,000454593 | HP65b_ TIG1 chr5:22397677-22400678 FORWARD LENGTH=547 |
| AT3G01120.1 | 1,772871718 | 0,00046555 | AtCYS1_ CGS_ AtCGS1_ CGS1_ MTO1 CYSTATHIONINE GAMMA-SYNTHASE 1_ CYSTATHIONINE GAMMA-S |
| AT5G66530.1 | 0,695323605 | 0,000488645 | no symbol available no full name available chr5:26553821-26555575 REVERSE LENGTH=307 |
| ATCG00350.1 | 1,267723935 | 0,000491891 | PSAA chrc:39605-41857 REVERSE LENGTH=750 |
| AT5G22800.1 | 1,295183104 | 0,000497832 | EMB263_ EMB1030_ EMB86 EMBRYO DEFECTIVE 263_ EMBRYO DEFECTIVE 1030_ EMBRYO DEFECTIVE 86 |
| AT1G12840.1 | 1,495053513 | 0,000499514 | DET3_ ATVHA-C ARABIDOPSIS THALIANA VACUOLAR ATP SYNTHASE SUBUNIT C_ DE-ETIOLATED 3 chr1[illegible] |
| AT2G28190.1 | 0,435745359 | 0,00050142 | CSD2_ CZSOD2_ SOD2_ AtSOD2 COPPER/ZINC SUPEROXIDE DISMUTASE 2_ superoxide dismutase 2_ copper/zin[illegible] |
| AT3G15190.1 | 0,429037725 | 0,000506996 | PRPS20 plastid ribosomal protein S20 chr3:5116216-5117412 FORWARD LENGTH=202 |
| AT1G74270.1 | 0,476587242 | 0,000516768 | no symbol available no full name available chr1:27928415-27929466 REVERSE LENGTH=112 |
| AT2G21390.1 | 1,364788001 | 0,000517896 | no symbol available no full name available chr2:9152428-9156577 FORWARD LENGTH=1218 |
| AT5G44340.1 | 1,567652399 | 0,0005187 | TUB4 tubulin beta chain 4 chr5:17859442-17860994 REVERSE LENGTH=444 |
| AT3G22960.1 | 1,494799958 | 0,000526182 | PKP1_ PKP-ALPHA PLASTIDIAL PYRUVATE KINASE 1 chr3:8139369-8141771 FORWARD LENGTH=596 |
| AT3G53420.1 | 1,391590227 | 0,000539594 | PIP2;1_ AtPIP2;1_ PIP2A_ PIP2 PLASMA MEMBRANE INTRINSIC PROTEIN 2_ PLASMA MEMBRANE INTRINSI[illegible] |
| AT1G75780.1 | 0,745578483 | 0,000541922 | TUB1 tubulin beta-1 chain chr1:28451378-28453602 REVERSE LENGTH=447 |
| AT4G21990.1 | 3,893937105 | 0,00055966 | APR3_ PRH26_ PRH-26_ ATAPR3 PAPS REDUCTASE HOMOLOG 26_ APS reductase 3 chr4:11657284-11658973 RE |
| AT3G55040.1 | 0,790683728 | 0,000562738 | GSTL2 glutathione transferase lambda 2 chr3:20398718-20400305 REVERSE LENGTH=292 |
| AT1G12900.1 | 1,114185713 | 0,000575737 | GAPA-2 glyceraldehyde 3-phosphate dehydrogenase A subunit 2 chr1:4392634-4394283 REVERSE LENGTH=399 |
| AT2G43750.1 | 1,186352119 | 0,000577336 | OASB_ CPACS1_ ACS1_ ATCS-B O-acetylserine (thiol) lyase B_ ARABIDOPSIS THALIANA CYSTEIN SYNTHASE-[illegible] |
| AT3G09840.1 | 1,50972618 | 0,000583539 | ATCDC48_ CDC48_ AtCDC48A_ CDC48A cell division cycle 48 chr3:3019494-3022832 FORWARD LENGTH=809 |
| AT3G23990.1 | 2,476025915 | 0,000586725 | HSP60_ HSP60-3B heat shock protein 60_ HEAT SHOCK PROTEIN 60-3B chr3:8669013-8672278 FORWARD LENGT[illegible] |
| AT3G16530.1 | 0,361780165 | 0,000591254 | no symbol available no full name available chr3:5624586-5625416 REVERSE LENGTH=276 |
| AT4G28080.1 | 2,18036834 | 0,000604851 | REC2 REDUCED CHLOROPLAST COVERAGE 2 chr4:13948993-13957840 REVERSE LENGTH=1819 |
| AT4G27520.1 | 0,731898097 | 0,00060923 | ENODL2_ AtENODL2 early nodulin-like protein 2 chr4:13750668-13751819 REVERSE LENGTH=349 |
| AT3G54890.4 | 0,702616898 | 0,000614604 | LHCA1 photosystem I light harvesting complex gene 1 chr3:20339881-20340922 REVERSE LENGTH=213 |
| AT2G15620.1 | 0,721439382 | 0,000616748 | NIR_ ATHNIR_ NIR1 ARABIDOPSIS THALIANA NITRITE REDUCTASE_ nitrite reductase 1_ NITRITE REDUCTAS |
| AT5G60670.1 | 1,678144714 | 0,000619466 | RPL12C Ribosomal Protein Like 12C chr5:24381066-24381566 REVERSE LENGTH=166 |
| AT5G61790.1 | 1,293489717 | 0,000626124 | CNX1_ ATCNX1 calnexin 1 chr5:24827394-24829642 REVERSE LENGTH=530 |
| AT1G64200.1 | 0,873344682 | 0,000638208 | VHA-E3 vacuolar H+-ATPase subunit E isoform 3 chr1:23828537-23830002 REVERSE LENGTH=237 |
| AT4G24830.1 | 1,605183842 | 0,00064247 | no symbol available no full name available chr4:12793085-12795857 REVERSE LENGTH=494 |
| AT3G46060.1 | 2,384508403 | 0,000646138 | ARA3_ RAB8A_ RABE1c_ ARA-3_ ATRAB8A_ ATRABE1C RAB GTPase homolog 8A chr3:16917908-16919740 FOR |

| | | | |
|---|---|---|---|
| AT5G56030.1 | 2,481675872 | 0,000655932 | HSP81-2_ HSP90.2_ AtHsp90.2_ ERD8_ HSP81.2 EARLY-RESPONSIVE TO DEHYDRATION 8_ heat shock protein 81 |
| AT1G08200.1 | 1,123249143 | 0,000665386 | AXS2 UDP-D-apiose/UDP-D-xylose synthase 2 chr1:2574259-2576609 REVERSE LENGTH=389 |
| AT5G52840.1 | 1,900551721 | 0,000665752 | no symbol available no full name available chr5:21413718-21414794 FORWARD LENGTH=169 |
| AT4G35830.1 | 1,606557153 | 0,000669515 | ACO1 aconitase 1 chr4:16973007-16977949 REVERSE LENGTH=898 |
| AT1G29670.1 | 0,670106665 | 0,000675948 | GDSL1_ GGL6 chr1:10375843-10377717 FORWARD LENGTH=363 |
| AT5G23120.1 | 0,868564414 | 0,000683613 | HCF136 HIGH CHLOROPHYLL FLUORESCENCE 136 chr5:7778154-7780463 FORWARD LENGTH=403 |
| AT2G27730.1 | 2,920506264 | 0,000706432 | no symbol available no full name available chr2:11820056-11820867 REVERSE LENGTH=113 |
| AT1G11840.6 | 1,246000091 | 0,000708344 | AtGLYI3_ GLX1_ ATGLX1 glyoxalaseI 3_ glyoxalase I homolog chr1:3995928-3997518 FORWARD LENGTH=322 |
| AT1G09270.1 | 1,55284924 | 0,000711508 | IMPA-4 importin alpha isoform 4 chr1:2994506-2997833 FORWARD LENGTH=538 |
| AT5G56000.1 | 1,616449667 | 0,000725622 | Hsp81.4_ AtHsp90.4 HEAT SHOCK PROTEIN 90.4_ HEAT SHOCK PROTEIN 81.4 chr5:22677602-22680067 REVERS |
| AT4G20360.1 | 1,0793169 | 0,000729417 | SVR11_ RAB8D_ ATRAB8D_ ATRABE1B_ RABE1b RAB GTPase homolog E1B_ SUPPRESSOR OF VARIEGATION |
| AT1G04820.1 | 1,286232655 | 0,00073016 | TUA4_ TOR2 TORTIFOLIA 2_ tubulin alpha-4 chain chr1:1356421-1358266 REVERSE LENGTH=450 |
| AT3G16520.3 | 1,651734334 | 0,000730332 | UGT88A1 UDP-glucosyl transferase 88A1 chr3:5619355-5620833 REVERSE LENGTH=462 |
| AT1G22780.1 | 0,676439617 | 0,000741663 | RPS18A_ PFL_ PFL1 POINTED FIRST LEAVES_ POINTED FIRST LEAVES 1_ 40S RIBOSOMAL PROTEIN S18 chr |
| AT1G52230.1 | 0,801252219 | 0,000751314 | PSI-H_ PSAH-2_ PSAH2 photosystem I subunit H2_ PHOTOSYSTEM I SUBUNIT H-2 chr1:19454902-19455508 FORW |
| AT3G28220.1 | 0,429742304 | 0,000756771 | no symbol available no full name available chr3:10524420-10526497 FORWARD LENGTH=370 |
| AT3G59920.1 | 1,954002351 | 0,000761502 | GDI2_ ATGDI2 RAB GDP dissociation inhibitor 2 chr3:22135157-22138221 FORWARD LENGTH=444 |
| AT1G78380.1 | 1,515982718 | 0,00078738 | GSTU19_ GST8_ ATGSTU19 A. THALIANA GLUTATHIONE S-TRANSFERASE TAU 19_ glutathione S-transferase T |
| AT1G53310.1 | 0,705329353 | 0,000793915 | PPC1_ ATPEPC1_ PEPC1_ ATPPC1 phosphoenolpyruvate carboxylase 1_ PEP(PHOSPHOENOLPYRUVATE) CARBOX |
| AT5G54810.1 | 0,741649767 | 0,000808599 | TRP2_ ATTSB1_ TSB1_ TRPB tryptophan synthase beta-subunit 1_ TRYPTOPHAN BIOSYNTHESIS B_ TRYPTOPHA |
| AT4G05530.1 | 1,462737886 | 0,000809355 | SDRA_ IBR1 indole-3-butyric acid response 1_ SHORT-CHAIN DEHYDROGENASE/REDUCTASE A chr4:2816462-28 |
| AT3G16480.1 | 0,354763756 | 0,000814084 | MPPalpha mitochondrial processing peptidase alpha subunit chr3:5599906-5602716 FORWARD LENGTH=499 |
| AT5G25460.1 | 0,654557924 | 0,00082304 | DGR2 DUF642 L-GalL responsive gene 2 chr5:8863430-8865394 FORWARD LENGTH=369 |
| AT3G53460.1 | 0,487662027 | 0,000835558 | CP29 chloroplast RNA-binding protein 29 chr3:19819738-19821423 REVERSE LENGTH=342 |
| AT4G14880.1 | 1,160982093 | 0,000838756 | OASA1_ OLD3_ CYTACS1_ ATCYS-3A ONSET OF LEAF DEATH 3_ O-acetylserine (thiol) lyase (OAS-TL) isoform A |
| AT1G63000.1 | 0,806837866 | 0,000843668 | NRS/ER_ UER1 "UDP-4-KETO-6-DEOXY-D-GLUCOSE-3_5-EPIMERASE-4-REDUCTASE 1"_ nucleotide-rhamnose sy |
| AT5G54770.1 | 2,174864481 | 0,000843921 | THI1_ TZ_ THI4 THIAZOLE REQUIRING_ THIAMINE4 chr5:22246634-22247891 FORWARD LENGTH=349 |
| AT3G23810.1 | 1,421333444 | 0,00085365 | ATSAHH2_ SAHH2 S-ADENOSYL-L-HOMOCYSTEINE (SAH) HYDROLASE 2_ S-adenosyl-l-homocysteine (SAH) hy |
| AT1G70580.1 | 0,222556645 | 0,000855104 | GGT2_ AOAT2 GLUTAMATE:GLYOXYLATE AMINOTRANSFERASE 2_ alanine-2-oxoglutarate aminotransferase 2 c |
| AT2G26670.1 | 1,547581341 | 0,000856855 | ATHO1_ HY1_ HY6_ HO1_ GUN2_ TED4 REVERSAL OF THE DET PHENOTYPE 4_ ARABIDOPSIS THALIANA H |
| AT3G02520.1 | 1,403866197 | 0,000859117 | GRF7_ GF14 NU general regulatory factor 7 chr3:526800-527915 REVERSE LENGTH=265 |
| AT1G11650.1 | 1,509923528 | 0,000876729 | RBP45B_ ATRBP45B chr1:3914895-3917301 FORWARD LENGTH=306 |
| AT5G19760.1 | 1,189200105 | 0,000878987 | no symbol available no full name available chr5:6679591-6681845 REVERSE LENGTH=298 |
| AT1G60950.1 | 0,34684369 | 0,000879449 | ATFD2_ FD2_ FED A FERREDOXIN 2 chr1:22444565-22445011 FORWARD LENGTH=148 |
| ATCG00120.1 | 1,253985269 | 0,00088169 | ATPA ATP synthase subunit alpha chrc:9938-11461 REVERSE LENGTH=507 |
| AT2G42590.1 | 1,373128264 | 0,000894927 | GF14 MU_ GRF9_ GRF14 general regulatory factor 9 chr2:17732118-17733775 REVERSE LENGTH=263 |
| AT4G21280.1 | 1,508343943 | 0,000900233 | PSBQ-1_ PSBQA_ PSBQ photosystem II subunit QA_ PHOTOSYSTEM II SUBUNIT Q-1_ PHOTOSYSTEM II SUBUN |
| AT4G13930.1 | 1,2325258 | 0,00091315 | SHM4 serine hydroxymethyltransferase 4 chr4:8048013-8050021 REVERSE LENGTH=471 |
| AT3G57890.1 | 1,527723036 | 0,000917551 | no symbol available no full name available chr3:21438271-21441695 FORWARD LENGTH=573 |

| | | | |
|---|---|---|---|
| AT1G74470.1 | 0,672927172 | 0,000918102 | no symbol available no full name available chr1:27991248-27992845 FORWARD LENGTH=467 |
| AT1G30530.1 | 1,750370404 | 0,000930956 | UGT78D1 UDP-glucosyl transferase 78D1 chr1:10814917-10816374 FORWARD LENGTH=453 |
| AT3G54050.1 | 1,211514824 | 0,000937861 | HCEF1_ cfbp1 high cyclic electron flow 1 chr3:20016951-20018527 FORWARD LENGTH=417 |
| AT2G07698.1 | 1,293467789 | 0,000948093 | no symbol available no full name available chr2:3361474-3364028 FORWARD LENGTH=777 |
| AT3G46740.1 | 1,416293021 | 0,00095237 | MAR1_ TOC75_ TOC75-III MODIFIER OF ARG1 1_ translocon at the outer envelope membrane of chloroplasts 75-III ch |
| AT3G28270.1 | 1,789635612 | 0,001008236 | AFL1 At14a-Like1 chr3:10538725-10539849 FORWARD LENGTH=374 |
| AT1G03130.1 | 0,687322387 | 0,001009843 | PSAD-2 photosystem I subunit D-2 chr1:753528-754142 REVERSE LENGTH=204 |
| AT2G05840.3 | 1,353850226 | 0,00101931 | PAA2 20S proteasome subunit PAA2 chr2:2234226-2235533 FORWARD LENGTH=205 |
| AT5G20980.1 | 1,300386052 | 0,001020755 | MS3_ ATMS3 methionine synthase 3 chr5:7124397-7128353 REVERSE LENGTH=812 |
| AT3G57290.1 | 2,102063995 | 0,00103291 | EIF3E_ ATINT6_ TIF3E1_ INT6_ INT-6_ ATEIF3E-1 eukaryotic translation initiation factor 3E chr3:21196786-2119907 |
| AT3G25860.1 | 1,511483434 | 0,001033457 | LTA2_ PLE2 PLASTID E2 SUBUNIT OF PYRUVATE DECARBOXYLASE chr3:9460632-9462585 FORWARD LENG |
| AT3G14415.1 | 1,273861313 | 0,001043796 | GOX2 glycolate oxidase 2 chr3:4818667-4820748 FORWARD LENGTH=367 |
| AT5G26360.1 | 2,192827443 | 0,001067236 | CCT3 Chaperonin containing T-complex polypeptide-1 subunit 3 chr5:9255561-9258891 REVERSE LENGTH=555 |
| AT3G53870.1 | 0,784748223 | 0,001067361 | no symbol available no full name available chr3:19951547-19952782 FORWARD LENGTH=249 |
| AT2G35370.1 | 0,743456388 | 0,001069334 | GDCH glycine decarboxylase complex H chr2:14891239-14892050 FORWARD LENGTH=165 |
| AT2G39330.1 | 0,647467538 | 0,001075851 | JAL23 jacalin-related lectin 23 chr2:16419787-16421573 REVERSE LENGTH=459 |
| AT5G52650.1 | 1,876524963 | 0,001081145 | no symbol available no full name available chr5:21355781-21357003 REVERSE LENGTH=179 |
| AT4G26530.1 | 0,690952741 | 0,001102493 | FBA5_ AtFBA5_ DEG22 fructose-bisphosphate aldolase 5 chr4:13391566-13392937 FORWARD LENGTH=358 |
| AT1G13060.1 | 1,225712593 | 0,001103873 | PBE1 20S proteasome beta subunit E1 chr1:4452641-4454663 FORWARD LENGTH=274 |
| AT5G23540.1 | 1,240747716 | 0,001109126 | no symbol available no full name available chr5:7937772-7939339 FORWARD LENGTH=308 |
| AT3G45140.1 | 0,640093096 | 0,001109566 | ATLOX2_ LOX2 ARABIODOPSIS THALIANA LIPOXYGENASE 2_ lipoxygenase 2 chr3:16525437-16529233 FORWA |
| AT1G09620.1 | 1,345787045 | 0,001165425 | no symbol available no full name available chr1:3113077-3116455 REVERSE LENGTH=1091 |
| AT2G29560.1 | 1,936781728 | 0,001173223 | ENOC_ ENO3 cytosolic enolase_ enolase 3 chr2:12646635-12649694 FORWARD LENGTH=475 |
| AT3G48730.1 | 1,355382727 | 0,001182306 | GSAM_ GSA2 glutamate-l-semialdehyde aminomutase_ "glutamate-1-semialdehyde 2_1-aminomutase 2" chr3:18049697-1 |
| AT5G11170.1 | 1,305593569 | 0,001194316 | UAP56a homolog of human UAP56 a chr5:3553334-3556646 FORWARD LENGTH=427 |
| AT3G53430.1 | 0,455440523 | 0,001207184 | no symbol available no full name available chr3:19809895-19810395 REVERSE LENGTH=166 |
| AT4G37870.1 | 1,563088512 | 0,001216683 | PCK1_ PEPCK PHOSPHOENOLPYRUVATE CARBOXYKINASE_ phosphoenolpyruvate carboxykinase 1 chr4:1780297 |
| AT5G53490.1 | 1,259413739 | 0,001229823 | TL17 Thylakoid lumenal 17.4 kDa protein chr5:21723488-21724621 REVERSE LENGTH=236 |
| AT3G12110.1 | 0,891346852 | 0,001237782 | ACT11 actin-11 chr3:3858116-3859609 FORWARD LENGTH=377 |
| AT3G11510.1 | 0,791047156 | 0,001239924 | no symbol available no full name available chr3:3623757-3624866 REVERSE LENGTH=150 |
| AT3G15020.1 | 1,119185551 | 0,001252482 | mMDH2 mitochondrial malate dehydrogenase 2 chr3:5056139-5057941 FORWARD LENGTH=341 |
| AT1G74040.1 | 1,567936893 | 0,001259561 | IPMS2_ IMS1_ MAML-3 SOPROPYLMALATE SYNTHASE 2_ 2-isopropylmalate synthase 1 chr1:27842258-27845566 |
| AT1G21750.1 | 1,203167957 | 0,00127099 | PDIL1-1_ ATPDI5_ PDI5_ ATPDIL1-1 PDI-like 1-1_ ARABIDOPSIS THALIANA PROTEIN DISULFIDE ISOMERASI |
| AT1G09430.1 | 1,65402844 | 0,001273308 | ACLA-3 ATP-citrate lyase A-3 chr1:3042135-3044978 FORWARD LENGTH=424 |
| AT3G16390.1 | 0,768597977 | 0,001273527 | NSP3 nitrile specifier protein 3 chr3:5562602-5564356 FORWARD LENGTH=467 |
| AT3G52730.1 | 1,302438396 | 0,001282321 | no symbol available no full name available chr3:19543146-19544167 REVERSE LENGTH=72 |
| AT1G04710.1 | 2,381367537 | 0,001285185 | KAT1_ PKT4 3-KETO-ACYL-COA THIOLASE 1_ peroxisomal 3-ketoacyl-CoA thiolase 4 chr1:1321941-1324556 FORW |
| AT3G16470.1 | 0,663955388 | 0,001303638 | JAL35_ AtJAC1_ JR1 jacalin-related lectin 35_ JACALIN-LECTIN LIKE 1_ JASMONATE RESPONSIVE 1 chr3:559609 |
| AT4G24930.1 | 0,668988591 | 0,001306038 | no symbol available no full name available chr4:12821496-12822389 REVERSE LENGTH=225 |

| | | | |
|---|---|---|---|
| AT3G63410.1 | 0,83755229 | 0,00131438 | E37_ VTE3_ IEP37_ APG1 INNER ENVELOPE PROTEIN 37_ VITAMIN E DEFECTIVE 3_ ALBINO OR PALE GREI |
| AT4G23850.1 | 1,716656815 | 0,001316148 | LACS4 long-chain acyl-CoA synthetase 4 chr4:12403720-12408263 REVERSE LENGTH=666 |
| AT3G18780.1 | 0,535240915 | 0,001318341 | LSR2_ ACT2_ ENL2_ DER1_ FIZ2 FRIZZY AND KINKED SHOOTS 2_ LIGHT STRESS-REGULATED 2_ DEFORMI |
| AT5G45390.1 | 1,401029157 | 0,001339173 | CLPP4_ NCLPP4 NUCLEAR-ENCODED CLP PROTEASE P4_ CLP protease P4 chr5:18396351-18397586 FORWARD |
| AT2G05920.1 | 0,729345697 | 0,001341089 | SBT1.8 subtilase 1.8 chr2:2269831-2272207 REVERSE LENGTH=754 |
| AT3G63540.1 | 0,533368667 | 0,001341453 | no symbol available no full name available chr3:23459372-23459803 REVERSE LENGTH=143 |
| AT2G32920.1 | 2,263304748 | 0,001345375 | PDIL2-3_ PDI9_ ATPDIL2-3_ ATPDI9 ARABIDOPSIS THALIANA PROTEIN DISULFIDE ISOMERASE 9_ PDI-like [illegible] |
| ATCG00740.1 | 0,834666529 | 0,001348047 | RPOA RNA polymerase subunit alpha chrc:77901-78890 REVERSE LENGTH=329 |
| AT4G28750.1 | 0,855001046 | 0,001348348 | PSAE-1 PSA E1 KNOCKOUT chr4:14202951-14203888 REVERSE LENGTH=143 |
| AT4G11380.1 | 0,626424304 | 0,001352523 | no symbol available no full name available chr4:6920608-6925444 FORWARD LENGTH=894 |
| AT2G25140.1 | 2,84084767 | 0,001359371 | HSP98.7_ CLPB-M_ CLPB4 HEAT SHOCK PROTEIN 98.7_ CASEIN LYTIC PROTEINASE B-M_ casein lytic proteina |
| AT1G79230.3 | 1,142294017 | 0,001365325 | ATMST1_ STR1_ MST1_ ATRDH1_ ST1 ARABIDOPSIS THALIANA RHODANESE HOMOLOGUE 1_ mercaptopyru |
| AT3G25770.1 | 0,536552574 | 0,001371665 | AOC2 allene oxide cyclase 2 chr3:9406975-9407839 FORWARD LENGTH=253 |
| AT1G03220.1 | 1,329589656 | 0,00139129 | SAP2 secreted aspartic protease 2 chr1:787143-788444 FORWARD LENGTH=433 |
| AT3G23490.1 | 0,577529702 | 0,001394587 | CYN cyanase chr3:8423238-8424415 REVERSE LENGTH=168 |
| AT5G05010.1 | 2,039958065 | 0,001396864 | no symbol available no full name available chr5:1477137-1479872 FORWARD LENGTH=527 |
| AT5G12110.1 | 1,410983891 | 0,001404681 | no symbol available no full name available chr5:3914483-3915732 FORWARD LENGTH=228 |
| AT5G50850.1 | 1,248819179 | 0,001411856 | MAB1 MACCI-BOU chr5:20689671-20692976 FORWARD LENGTH=363 |
| AT5G50950.1 | 0,482613437 | 0,001414291 | FUM2 FUMARASE 2 chr5:20729687-20733476 FORWARD LENGTH=510 |
| AT5G37640.1 | 1,538833725 | 0,001432689 | UBQ9 ubiquitin 9 chr5:14952782-14953750 REVERSE LENGTH=322 |
| AT5G19510.1 | 0,623647807 | 0,001465575 | no symbol available no full name available chr5:6581854-6583137 REVERSE LENGTH=224 |
| AT5G48580.1 | 0,485159356 | 0,001495668 | FKBP15-2 FK506- and rapamycin-binding protein 15 kD-2 chr5:19696156-19697304 REVERSE LENGTH=163 |
| AT1G44575.1 | 1,167453089 | 0,001501027 | CP22_ PSBS_ NPQ4 NONPHOTOCHEMICAL QUENCHING 4_ PHOTOSYSTEM II SUBUNIT S chr1:16871768-1687[illegible] |
| AT1G19570.1 | 0,736672171 | 0,001512431 | ATDHAR1_ DHAR1_ DHAR5 DEHYDROASCORBATE REDUCTASE 5_ dehydroascorbate reductase chr1:6773462-67 |
| AT5G54190.1 | 0,706607007 | 0,001517483 | PORA protochlorophyllide oxidoreductase A chr5:21991183-21992773 REVERSE LENGTH=405 |
| AT1G08830.1 | 0,373356934 | 0,001523738 | CSD1_ AtSOD1_ SOD1 superoxide dismutase 1_ copper/zinc superoxide dismutase 1 chr1:2827700-2829053 FORWARD |
| AT3G08030.2 | 0,562444837 | 0,00152615 | AthA2-1 chr3:2564517-2565819 FORWARD LENGTH=323 |
| AT3G20820.1 | 0,798678207 | 0,001544564 | no symbol available no full name available chr3:7280930-7282027 FORWARD LENGTH=365 |
| AT1G04480.1 | 0,753654019 | 0,001544769 | no symbol available no full name available chr1:1216110-1217257 FORWARD LENGTH=140 |
| AT5G19940.1 | 0,709404052 | 0,001549209 | FBN6_ PAP8 FIBRILLIN6_ Probable Plastid-Lipid Associated Protein chr5:6739693-6740661 FORWARD LENGTH=23[illegible] |
| AT5G20920.2 | 0,678961889 | 0,001564511 | EIF2 BETA_ EMB1401_ eIF-2bs embryo defective 1401_ eukaryotic translation initiation factor 2 beta subunit chr5:7094[illegible] |
| AT5G24780.1 | 0,63773054 | 0,001571193 | VSP1_ ATVSP1 vegetative storage protein 1 chr5:8507783-8508889 REVERSE LENGTH=270 |
| AT3G12915.1 | 0,487638656 | 0,001595285 | no symbol available no full name available chr3:4112999-4115708 FORWARD LENGTH=820 |
| AT1G20440.1 | 1,428741278 | 0,001596829 | AtCOR47_ RD17_ COR47 cold-regulated 47 chr1:7084722-7085664 REVERSE LENGTH=265 |
| AT3G60820.1 | 1,258658779 | 0,001607745 | PBF1 chr3:22472038-22473809 REVERSE LENGTH=223 |
| AT5G45280.1 | 0,255668784 | 0,001618902 | PAE11 pectin acetylesterase 11 chr5:18346862-18349432 FORWARD LENGTH=370 |
| AT3G25920.1 | 0,713593921 | 0,001624244 | RPL15 ribosomal protein L15 chr3:9491268-9492558 REVERSE LENGTH=277 |
| AT2G30870.1 | 0,874943421 | 0,001625634 | GSTF10_ ATGSTF10_ ERD13_ ATGSTF4 glutathione S-transferase PHI 10_ ARABIDOPSIS THALIANA GLUTATHIC |
| AT4G09040.1 | 0,554225035 | 0,001646569 | CP33C chr4:5795075-5797315 REVERSE LENGTH=304 |

| | | | |
|---|---|---|---|
| AT4G16143.1 | 2,240690482 | 0,001671326 | IMPA-2 importin alpha isoform 2 chr4:9134450-9137134 REVERSE LENGTH=535 |
| AT1G66270.2 | 0,189164259 | 0,001677582 | BGLU21 chr1:24700110-24702995 REVERSE LENGTH=522 |
| AT5G10160.1 | 1,266181495 | 0,00167841 | no symbol available no full name available chr5:3185819-3187159 FORWARD LENGTH=219 |
| AT2G25450.1 | 0,450709939 | 0,001685541 | GSL-OH glucosinolate hydroxylase chr2:10830286-10831563 REVERSE LENGTH=359 |
| AT3G03960.1 | 2,072300165 | 0,001685584 | CCT8 Chaperonin containing T-complex polypeptide-1 subunit 8 chr3:1024432-1027604 FORWARD LENGTH=549 |
| AT4G27560.1 | 1,740992546 | 0,001695644 | UGT79B2 chr4:13760114-13761481 REVERSE LENGTH=455 |
| AT5G03340.1 | 1,465445344 | 0,001701141 | AtCDC48C cell division cycle 48C chr5:810091-813133 REVERSE LENGTH=810 |
| AT1G72370.2 | 1,177913191 | 0,001706156 | RPSAA_ AP40_ RP40_ P40 40s ribosomal protein SA chr1:27243148-27244842 REVERSE LENGTH=294 |
| AT1G76180.1 | 0,490504786 | 0,001725807 | ERD14 EARLY RESPONSE TO DEHYDRATION 14 chr1:28587013-28587657 REVERSE LENGTH=185 |
| AT4G37800.1 | 0,700206 | 0,001743279 | XTH7 xyloglucan endotransglucosylase/hydrolase 7 chr4:17775703-17777372 REVERSE LENGTH=293 |
| AT3G16400.1 | 1,425384331 | 0,00176759 | NSP1_ ATNSP1_ ATMLP-470 nitrile specifier protein 1_ NITRILE SPECIFIER PROTEIN 1_ MYROSINASE-BINDING |
| AT1G10840.1 | 1,818901356 | 0,001769398 | TIF3H1 translation initiation factor 3 subunit H1 chr1:3607885-3610299 REVERSE LENGTH=337 |
| AT5G17770.1 | 1,714285714 | 0,001773095 | CBR_ CBR1_ ATCBR NADH:cytochrome B5 reductase 1_ NADH:CYTOCHROME B5 REDUCTASE 1 chr5:5864543-5[illegible] |
| AT1G47260.1 | 0,609623895 | 0,001835745 | APFI_ GAMMA CA2 gamma carbonic anhydrase 2 chr1:17321384-17323347 REVERSE LENGTH=278 |
| AT1G07140.1 | 0,847294128 | 0,001851494 | SIRANBP chr1:2192360-2193688 REVERSE LENGTH=228 |
| AT5G52920.1 | 1,686374485 | 0,001892045 | PKP-BETA1_ PKP2_ PKP1 plastidic pyruvate kinase beta subunit 1_ PLASTIDIAL PYRUVATE KINASE 1_ PLASTIDI[illegible] |
| AT1G20620.1 | 0,804283968 | 0,001905406 | SEN2_ ROG1_ ATCAT3_ CAT3 SENESCENCE 2_ catalase 3_ REPRESSOR OF GSNOR1 chr1:7143142-7146193 FOR[illegible] |
| AT5G60600.1 | 1,228464415 | 0,001918195 | ISPG_ CSB3_ HDS_ GCPE_ CLB4 CONSTITUTIVE SUBTILISIN 3_ 4-hydroxy-3-methylbut-2-enyl diphosphate synthas |
| AT2G05620.1 | 3,338232492 | 0,001941532 | PGR5_ AtPGR5 proton gradient regulation 5 chr2:2081204-2081687 REVERSE LENGTH=133 |
| ATCG00750.1 | 0,443974474 | 0,00196356 | RPS11 ribosomal protein S11 chrc:78960-79376 REVERSE LENGTH=138 |
| AT3G12780.1 | 1,183868874 | 0,001963725 | PGKp1_ PGK1 phosphoglycerate kinase 1 chr3:4061127-4063140 REVERSE LENGTH=481 |
| AT1G76080.1 | 0,845328496 | 0,001971384 | CDSP32_ TRXL1_ ATCDSP32 ARABIDOPSIS THALIANA CHLOROPLASTIC DROUGHT-INDUCED STRESS PRO[illegible] |
| AT1G29900.1 | 1,776739843 | 0,001985834 | CARB_ VEN3 carbamoyl phosphate synthetase B_ VENOSA 3 chr1:10468164-10471976 FORWARD LENGTH=1187 |
| AT5G04140.1 | 0,909609168 | 0,001989019 | GLUS_ FD-GOGAT_ GLS1_ GLU1 FERREDOXIN-DEPENDENT GLUTAMATE SYNTHASE 1_ glutamate synthase 1[illegible] |
| AT3G17810.1 | 0,847759377 | 0,002020218 | PYD1 pyrimidine 1 chr3:6094279-6096289 FORWARD LENGTH=426 |
| AT3G58730.1 | 1,43812238 | 0,002062195 | VHA-D chr3:21718495-21719280 REVERSE LENGTH=261 |
| AT3G23700.1 | 0,751313627 | 0,002068614 | SRRP1 S1 RNA-binding ribosomal protein 1 chr3:8531689-8533742 REVERSE LENGTH=392 |
| AT1G67700.1 | 0,62735782 | 0,002074106 | HHL1 HYPERSENSITIVE TO HIGH LIGHT 1 chr1:25374295-25375716 FORWARD LENGTH=230 |
| AT3G10090.1 | 4,114586493 | 0,00209724 | no symbol available no full name available chr3:3108960-3109154 REVERSE LENGTH=64 |
| AT1G59870.1 | 1,468140532 | 0,002119832 | ABCG36_ ATABCG36_ PEN3_ PDR8_ ATPDR8 Arabidopsis thaliana ATP-binding cassette G36_ PLEIOTROPIC DRU[illegible] |
| AT5G65010.1 | 1,561552967 | 0,002122717 | ASN2 asparagine synthetase 2 chr5:25969224-25972278 FORWARD LENGTH=578 |
| AT5G45280.2 | 0,596581804 | 0,002134297 | PAE11 pectin acetylesterase 11 chr5:18346862-18349488 FORWARD LENGTH=391 |
| AT1G02920.1 | 0,419632571 | 0,002142154 | ATGSTF8_ GSTF7_ ATGSTF7_ ATGST11_ GST11 GLUTATHIONE S-TRANSFERASE 11_ glutathione S-transferase [illegible] |
| AT1G55210.1 | 0,651421074 | 0,002148627 | no symbol available no full name available chr1:20598057-20598620 REVERSE LENGTH=187 |
| AT1G35580.3 | 2,698836432 | 0,002158682 | CINV1_ A/N-InvG_ NIN2 cytosolic invertase 1_ alkaline/neutral invertase G_ neutral invertase 2 chr1:13123183-1312480[illegible] |
| AT1G41830.1 | 0,179050115 | 0,002193886 | SKS6 SKU5-similar 6_ SKU5 SIMILAR 6 chr1:15603892-15607802 REVERSE LENGTH=542 |
| AT3G63190.1 | 0,772082005 | 0,002238461 | cpRRF_ HFP108_ AtcpRRF_ RRF chloroplast ribosome recycling factor_ Arabidopsis thaliana chloroplast ribosome recycl |
| AT4G17530.1 | 1,472252749 | 0,002258304 | RAB1C_ ATRABD2C_ ATRAB1C RAB GTPase homolog 1C chr4:9773721-9775424 REVERSE LENGTH=202 |
| ATCG00660.1 | 0,090799237 | 0,002270461 | RPL20 ribosomal protein L20 chrc:68512-68865 REVERSE LENGTH=117 |

| | | | |
|---|---|---|---|
| AT2G42540.1 | 1,350091647 | 0,00229512 | COR15_ AtCOR15A_ COR15A cold-regulated 15a chr2:17711241-17711930 REVERSE LENGTH=127 |
| AT5G14780.1 | 2,21764776 | 0,002328456 | FDH_ AtFDH1 formate dehydrogenase chr5:4777043-4779190 FORWARD LENGTH=384 |
| AT3G59970.3 | 1,235034145 | 0,002366942 | MTHFR1 methylenetetrahydrofolate reductase 1 chr3:22151303-22154323 FORWARD LENGTH=592 |
| AT5G36880.4 | 1,330598499 | 0,002404395 | ACS acetyl-CoA synthetase chr5:14535506-14539084 REVERSE LENGTH=610 |
| AT2G43560.1 | 0,694076557 | 0,00241495 | no symbol available no full name available chr2:18073995-18075385 REVERSE LENGTH=223 |
| AT1G50250.1 | 0,732402338 | 0,002426887 | FTSH1 FTSH protease 1 chr1:18614398-18616930 REVERSE LENGTH=716 |
| AT4G11010.1 | 0,772405972 | 0,002441989 | NDPK3 nucleoside diphosphate kinase 3 chr4:6732780-6734298 REVERSE LENGTH=238 |
| AT1G45000.1 | 1,401440085 | 0,002489055 | RPT4b chr1:17009220-17011607 FORWARD LENGTH=399 |
| AT1G12240.1 | 0,428869002 | 0,002494801 | AtVI2_ VAC-INV_ ATBETAFRUCT4_ VI2_ FRUCT4_ AtFRUCT4_ VIN2 VACUOLAR INVERTASE_ vacuolar invert |
| AT1G18500.1 | 1,452171808 | 0,00249517 | MAML-4_ IPMS1 methylthioalkylmalate synthase-like 4_ ISOPROPYLMALATE SYNTHASE 1 chr1:6369347-6372861 I |
| AT1G74910.1 | 1,945478209 | 0,002501451 | KJC1 KONJAC 1 chr1:28135770-28138456 REVERSE LENGTH=415 |
| AT2G17340.1 | 1,741849354 | 0,00251235 | no symbol available no full name available chr2:7541615-7544089 REVERSE LENGTH=367 |
| AT2G45960.2 | 1,298074158 | 0,002518339 | TMP-A_ ATHH2_ PIP1;2_ PIP1B TRANSMEMBRANE PROTEIN A_ plasma membrane intrinsic protein 1B_ NAMED I |
| AT2G20360.1 | 1,402824296 | 0,002530908 | no symbol available no full name available chr2:8786070-8789098 FORWARD LENGTH=402 |
| AT3G49720.1 | 0,380511627 | 0,00253505 | CGR2 chr3:18440192-18441655 REVERSE LENGTH=261 |
| AT4G36250.1 | 1,690026026 | 0,00258136 | ALDH3F1 aldehyde dehydrogenase 3F1 chr4:17151029-17153381 FORWARD LENGTH=484 |
| AT5G51820.1 | 0,442311081 | 0,002611856 | ATPGMP_ PGM_ PGM1_ STF1 ARABIDOPSIS THALIANA PHOSPHOGLUCOMUTASE_ STARCH-FREE 1_ phosph |
| AT5G48180.1 | 2,430162811 | 0,002633499 | NSP5_ AtNSP5 nitrile specifier protein 5 chr5:19541283-19542358 REVERSE LENGTH=326 |
| AT5G49910.1 | 1,458571403 | 0,002640858 | HSC70-7_ cpHsc70-2 chloroplast heat shock protein 70-2_ HEAT SHOCK PROTEIN 70-7 chr5:20303470-20306295 FOR |
| AT5G16590.1 | 1,344312618 | 0,002654147 | LRR1 Leucine rich repeat protein 1 chr5:5431862-5433921 FORWARD LENGTH=625 |
| AT1G65980.1 | 1,40177062 | 0,002659483 | TPX1 thioredoxin-dependent peroxidase 1 chr1:24559524-24560753 REVERSE LENGTH=162 |
| AT1G51980.1 | 1,464972724 | 0,002670334 | no symbol available no full name available chr1:19323692-19326771 REVERSE LENGTH=503 |
| AT1G07790.1 | 0,854434902 | 0,002694941 | HTB1 chr1:2413049-2413495 FORWARD LENGTH=148 |
| AT5G12860.2 | 1,149693792 | 0,002720531 | DiT1 dicarboxylate transporter 1 chr5:4059850-4061919 REVERSE LENGTH=556 |
| AT1G12250.2 | 0,485282515 | 0,002758744 | TL20.3 chr1:4159623-4161269 FORWARD LENGTH=206 |
| AT3G14290.1 | 1,153035448 | 0,002759223 | PAE2 20S proteasome alpha subunit E2 chr3:4764364-4766381 FORWARD LENGTH=237 |
| AT4G02770.1 | 1,306111969 | 0,002780563 | PSAD1_ PSAD-1 photosystem I subunit D-1 chr4:1229247-1229873 REVERSE LENGTH=208 |
| AT1G05010.1 | 0,289409304 | 0,002783177 | EFE_ ACO4_ EAT1 ethylene forming enzyme_ ethylene-forming enzyme chr1:1431419-1432695 REVERSE LENGTH=32 |
| AT5G28500.1 | 1,246839136 | 0,002787335 | no symbol available no full name available chr5:10477810-10479114 FORWARD LENGTH=434 |
| AT5G67360.1 | 0,608570868 | 0,002831757 | ARA12_ SBT1.7 Subtilisin-like Serine protease 1.7 chr5:26872192-26874465 REVERSE LENGTH=757 |
| AT3G24170.1 | 1,759626204 | 0,002907749 | ATGR1_ GR1 glutathione-disulfide reductase chr3:8729762-8734115 REVERSE LENGTH=499 |
| AT2G36530.1 | 1,21913031 | 0,002921857 | LOS2_ ENO2 LOW EXPRESSION OF OSMOTICALLY RESPONSIVE GENES 2_ enolase 2 chr2:15321081-15323786 I |
| AT3G61050.1 | 1,375566336 | 0,002922017 | CLB1_ SYT7_ AtCLB_ NTMC2TYPE4_ NTMC2T4 calcium-dependent lipid-binding protein_ Synaptotagmin 7 chr3:225 |
| AT2G21260.1 | 0,433983766 | 0,002937165 | no symbol available no full name available chr2:9105693-9107308 REVERSE LENGTH=309 |
| AT1G21440.1 | 1,356239579 | 0,002948332 | no symbol available no full name available chr1:7502325-7504103 REVERSE LENGTH=336 |
| AT5G15970.1 | 13,92761872 | 0,002978763 | AtCor6.6_ KIN2_ COR6.6 COLD-RESPONSIVE 6.6 chr5:5211966-5212441 FORWARD LENGTH=66 |
| AT3G04840.1 | 0,897346141 | 0,002981681 | no symbol available no full name available chr3:1329751-1331418 FORWARD LENGTH=262 |
| AT3G04770.1 | 1,343771085 | 0,002985524 | RPSAb 40s ribosomal protein SA B chr3:1309465-1310846 REVERSE LENGTH=332 |
| AT5G13420.1 | 2,235684265 | 0,003000342 | GSM2_ TRA2 Glc-hypersensitive mutant 2_ transaldolase 2 chr5:4302080-4304212 REVERSE LENGTH=438 |

| | | | |
|---|---|---|---|
| AT3G46970.1 | 1,341486667 | 0,003003293 | ATPHS2_ PHS2 alpha-glucan phosphorylase 2_ Arabidopsis thaliana alpha-glucan phosphorylase 2 chr3:17301625-173061 |
| AT4G21210.1 | 0,340430715 | 0,003059602 | RP1_ ATRP1 PPDK regulatory protein chr4:11307002-11308587 FORWARD LENGTH=403 |
| AT1G65960.2 | 2,019008172 | 0,00306286 | GAD2 glutamate decarboxylase 2 chr1:24552094-24557253 FORWARD LENGTH=494 |
| AT1G42970.1 | 1,103704743 | 0,003143056 | GAPB glyceraldehyde-3-phosphate dehydrogenase B subunit chr1:16127552-16129584 FORWARD LENGTH=447 |
| AT5G02490.1 | 1,187813103 | 0,003148019 | AtHsp70-2_ Hsp70-2 chr5:550296-552565 REVERSE LENGTH=653 |
| AT5G26830.1 | 1,421414496 | 0,003151238 | no symbol available no full name available chr5:9437351-9441568 FORWARD LENGTH=709 |
| AT3G04120.1 | 1,204769213 | 0,003153705 | GAPC1_ GAPC_ GAPC-1 GLYCERALDEHYDE-3-PHOSPHATE DEHYDROGENASE C SUBUNIT_ glyceraldehyde-3 |
| AT1G52040.1 | 0,324535575 | 0,00318145 | MBP1_ ATMBP myrosinase-binding protein 1 chr1:19350595-19352578 REVERSE LENGTH=462 |
| AT2G16600.1 | 0,673655691 | 0,003188224 | AtCYP19-1_ ROC3_ CYP19 rotamase CYP 3_ cyclophilin 19 chr2:7200862-7201383 FORWARD LENGTH=173 |
| AT4G00660.1 | 2,115266488 | 0,003213097 | RH8_ ATRH8 RNAhelicase-like 8 chr4:274638-277438 FORWARD LENGTH=505 |
| AT5G16050.1 | 1,587936179 | 0,003216633 | GRF5_ GF14 UPSILON general regulatory factor 5 chr5:5244008-5245402 REVERSE LENGTH=268 |
| AT5G58260.1 | 0,69638417 | 0,003250883 | NdhN NADH dehydrogenase-like complex N chr5:23561075-23561929 REVERSE LENGTH=209 |
| AT3G13750.1 | 0,496030169 | 0,003322251 | BGAL1 beta galactosidase 1_ beta-galactosidase 1 chr3:4511192-4515756 FORWARD LENGTH=847 |
| AT2G20140.1 | 1,246981207 | 0,003349832 | RPT2b regulatory particle AAA-ATPase 2b chr2:8692736-8694837 FORWARD LENGTH=443 |
| AT5G09660.1 | 1,161620101 | 0,003365967 | PMDH2 peroxisomal NAD-malate dehydrogenase 2 chr5:2993645-2995551 REVERSE LENGTH=354 |
| AT5G19990.1 | 1,480404174 | 0,003409166 | RPT6A_ ATSUG1 regulatory particle triple-A ATPase 6A chr5:6752144-6754918 FORWARD LENGTH=419 |
| ATCG00720.1 | 1,200915062 | 0,003443146 | PETB photosynthetic electron transfer B chrc:74841-76292 FORWARD LENGTH=215 |
| AT1G27950.1 | 7,380304503 | 0,003443664 | LTPG1 glycosylphosphatidylinositol-anchored lipid protein transfer 1 chr1:9740740-9741991 FORWARD LENGTH=193 |
| AT4G16660.1 | 1,661414958 | 0,003540071 | HSP70 heat shock protein 70 chr4:9377225-9381232 FORWARD LENGTH=867 |
| AT5G17050.1 | 0,418749344 | 0,003543317 | UGT78D2 UDP-glucosyl transferase 78D2 chr5:5607828-5609392 REVERSE LENGTH=460 |
| AT4G34230.1 | 2,459892524 | 0,003586971 | CAD-5_ ATCAD5_ CAD5 cinnamyl alcohol dehydrogenase 5 chr4:16386898-16388666 REVERSE LENGTH=357 |
| AT4G03520.1 | 0,775646524 | 0,003588648 | ATHM2_ TRXm2 thioredoxin m2 chr4:1562585-1564055 REVERSE LENGTH=186 |
| AT1G30360.1 | 1,568961531 | 0,003603496 | ERD4_ OSCA3.1 early-responsive to dehydration 4 chr1:10715892-10718799 FORWARD LENGTH=724 |
| AT4G39890.1 | 3,68934025 | 0,003617832 | AtRABH1c_ RABH1c RAB GTPase homolog H1C chr4:18506112-18507459 FORWARD LENGTH=214 |
| AT2G19900.1 | 0,858726094 | 0,003643182 | ATNADP-ME1_ NADP-ME1 NADP-malic enzyme 1_ Arabidopsis thaliana NADP-malic enzyme 1 chr2:8592106-859540 |
| AT1G50670.1 | 1,339274834 | 0,003751036 | OTU2 ovarian tumor domain (OTU)-containing DUB (deubiquitilating enzyme) 2 chr1:18775086-18776552 REVERSE LE |
| AT1G43670.1 | 0,762569411 | 0,003769889 | FBP_ AtcFBP_ cyfbp_ FINS1 "fructose-1_6-bisphosphatase"_ "Arabidopsis thaliana cytosolic fructose-1_6-bisphosphatase |
| AT5G26000.1 | 0,763614883 | 0,003787206 | TGG1_ AtTGG1_ BGLU38 thioglucoside glucohydrolase 1_ BETA GLUCOSIDASE 38 chr5:9079678-9082347 REVERS |
| AT3G47070.1 | 2,841128118 | 0,003798386 | no symbol available no full name available chr3:17337205-17337507 REVERSE LENGTH=100 |
| AT2G12550.1 | 0,631319826 | 0,003813779 | NUB1 homolog of human NUB1 chr2:5114881-5118486 FORWARD LENGTH=562 |
| AT3G07770.1 | 1,494607556 | 0,003821412 | Hsp89.1_ AtHsp90.6_ AtHsp90-6 HEAT SHOCK PROTEIN 90.6_ HEAT SHOCK PROTEIN 89.1_ HEAT SHOCK PRO |
| AT5G13120.1 | 0,677752885 | 0,003866963 | CYP20-2_ Pnsl5_ ATCYP20-2 cyclophilin 20-2_ Photosynthetic NDH subcomplex L 5_ ARABIDOPSIS THALIANA CY |
| AT2G09990.1 | 2,555118045 | 0,003869205 | no symbol available no full name available chr2:3781442-3781882 FORWARD LENGTH=146 |
| AT1G11750.1 | 1,331844459 | 0,003941669 | NCLPP1_ CLPP6_ NCLPP6 NUCLEAR-ENCODED CLPP 1_ CLP protease proteolytic subunit 6 chr1:3967609-3969535 |
| AT3G55610.1 | 2,5255404 | 0,003968557 | P5CS2 delta 1-pyrroline-5-carboxylate synthase 2 chr3:20624278-20628989 REVERSE LENGTH=726 |
| AT2G47730.1 | 1,381450578 | 0,003981542 | GST6_ GSTF8_ ATGSTF8_ ATGSTF5 glutathione S-transferase phi 8_ Arabidopsis thaliana glutathione S-transferase phi |
| AT5G41670.1 | 1,5047943 | 0,003986974 | PGD3 6-phosphogluconate dehydrogenase 3 chr5:16665647-16667110 REVERSE LENGTH=487 |
| AT5G14660.1 | 0,463795573 | 0,003999751 | ATDEF2_ DEF2_ PDF1B peptide deformylase 1B chr5:4727129-4728671 REVERSE LENGTH=273 |
| AT2G24200.1 | 1,355131891 | 0,004049359 | LAP1_ atLAP1 leucyl aminopeptidase 1 chr2:10287017-10289450 REVERSE LENGTH=520 |

| | | | |
|---|---|---|---|
| AT2G26540.3 | 0,726999039 | 0,004050128 | DUF3_ ATDUF3_ HEMD_ ATUROS_ UROS ARABIDOPSIS THALIANA UROPORPHYRINOGEN III SYNTHASE_ I |
| AT3G16640.1 | 0,810038522 | 0,004091176 | AtTCTP1_ TCTP1 translationally controlled tumor protein chr3:5669709-5670729 REVERSE LENGTH=168 |
| AT2G22240.1 | 16,32293876 | 0,004145709 | MIPS2_ ATIPS2_ ATMIPS2 myo-inositol-1-phosphate synthase 2_ INOSITOL 3-PHOSPHATE SYNTHASE 2_ MYO-IN |
| AT2G31670.1 | 0,648481995 | 0,004245706 | UP3 UP3 chr2:13472699-13473490 REVERSE LENGTH=263 |
| AT3G27830.1 | 1,18063612 | 0,004288985 | RPL12-A_ RPL12 ribosomal protein L12-A_ RIBOSOMAL PROTEIN L12 chr3:10318576-10319151 FORWARD LENG |
| AT1G20010.1 | 1,170530956 | 0,004289093 | TUB5 tubulin beta-5 chain chr1:6938033-6940481 REVERSE LENGTH=449 |
| AT5G50950.3 | 0,720268123 | 0,004311075 | FUM2 FUMARASE 2 chr5:20731191-20733636 FORWARD LENGTH=317 |
| AT2G37170.1 | 1,394676858 | 0,004325892 | PIP2;2_ PIP2B plasma membrane intrinsic protein 2_ PLASMA MEMBRANE INTRINSIC PROTEIN 2;2 chr2:15613624- |
| AT3G47470.1 | 0,833723336 | 0,00434592 | LHCA4_ CAB4 light-harvesting chlorophyll-protein complex I subunit A4 chr3:17493622-17494773 REVERSE LENGTH |
| AT1G55260.2 | 5,013384169 | 0,004373789 | LTPG6 glycosylphosphatidylinositol-anchored lipid protein transfer 6 chr1:20614663-20616158 FORWARD LENGTH=22 |
| AT3G60750.1 | 1,19921279 | 0,0043792 | AtTKL1_ TKL1 transketolase 1 chr3:22454004-22456824 FORWARD LENGTH=741 |
| AT3G13870.1 | 1,981662058 | 0,004394517 | GOM8_ RHD3 GOLGI MUTANT 8_ ROOT HAIR DEFECTIVE 3 chr3:4565762-4571109 REVERSE LENGTH=802 |
| AT1G04270.2 | 0,557808066 | 0,004401968 | RPS15 cytosolic ribosomal protein S15 chr1:1141852-1142960 REVERSE LENGTH=151 |
| AT1G69740.1 | 1,185125315 | 0,004439753 | HEMB1_ ALAD1 5-aminolevulinic acid dehydratase 1 chr1:26232197-26234713 FORWARD LENGTH=430 |
| AT2G06050.1 | 0,495691513 | 0,004459535 | OPR3_ DDE1_ AtOPR3 DELAYED DEHISCENCE 1_ oxophytodienoate-reductase 3 chr2:2359240-2361971 REVERSE I |
| AT2G05710.1 | 1,485832765 | 0,00450995 | ACO3 aconitase 3 chr2:2141591-2146350 FORWARD LENGTH=990 |
| AT1G09750.1 | 0,717973132 | 0,004538349 | no symbol available no full name available chr1:3157541-3158960 FORWARD LENGTH=449 |
| AT1G10760.1 | 1,290735515 | 0,004551808 | GWD_ GWD1_ SOP1_ SOP_ SEX1 STARCH EXCESS 1 chr1:3581210-3590043 REVERSE LENGTH=1399 |
| AT1G49750.1 | 0,588200651 | 0,004564802 | no symbol available no full name available chr1:18411177-18412779 REVERSE LENGTH=494 |
| AT2G17265.1 | 0,40446157 | 0,004575515 | DRM1_ HSK_ DMR1 DOWNY MILDEW RESISTANT 1_ homoserine kinase chr2:7508606-7509718 FORWARD LENC |
| AT1G48920.1 | 1,359010814 | 0,004579025 | PARL1_ NUC-L1_ ATNUC-L1_ NUC1 nucleolin like 1_ PARALLEL 1_ nucleolin 1 chr1:18098186-18101422 FORWAR |
| AT5G43850.1 | 2,967374795 | 0,004587765 | ATARD4_ ARD4 chr5:17627364-17629122 REVERSE LENGTH=187 |
| AT1G49760.1 | 0,854376398 | 0,004606451 | PABP8_ PAB8 POLY(A) BINDING PROTEIN 8_ poly(A) binding protein 8 chr1:18416740-18419753 FORWARD LENC |
| AT3G49870.1 | 1,398328176 | 0,004614537 | ARLA1C_ ARL8a_ ATARLA1C ADP-ribosylation factor-like A1C_ ADP-ribosylation factor-like 8a chr3:18492674-1849 |
| AT2G41840.1 | 0,753693664 | 0,004618474 | no symbol available no full name available chr2:17460016-17461398 REVERSE LENGTH=285 |
| AT3G15520.2 | 0,708445238 | 0,004619553 | no symbol available no full name available chr3:5249739-5251861 REVERSE LENGTH=326 |
| AT1G37130.1 | 1,525976836 | 0,004647475 | B29_ ATNR2_ NIA2_ NIA2-1_ NR2_ CHL3_ NR NITRATE REDUCTASE 2_ CHLORATE RESISTANT 3_ ARABIDO |
| AT5G11450.1 | 0,551279319 | 0,004651051 | PPD5 PsbP domain protein 5 chr5:3654475-3656357 FORWARD LENGTH=297 |
| AT3G04790.1 | 1,159983406 | 0,004666769 | EMB3119 EMBRYO DEFECTIVE 3119 chr3:1313365-1314195 FORWARD LENGTH=276 |
| AT5G65220.1 | 0,452587653 | 0,0047409 | PRPL29 plastid ribosomal proteins of the 50S subunit 29 chr5:26061301-26062506 FORWARD LENGTH=173 |
| AT5G54640.1 | 0,741365639 | 0,004843122 | HTA1_ RAT5_ ATHTA1 histone H2A 1_ RESISTANT TO AGROBACTERIUM TRANSFORMATION 5 chr5:2219654C |
| AT1G78830.1 | 0,845767391 | 0,004926474 | MNB1 chr1:29637141-29638508 REVERSE LENGTH=455 |
| AT3G44110.1 | 1,436445654 | 0,004927319 | ATJ3_ ATJ_ J3 DNAJ homologue 3 chr3:15869115-15871059 REVERSE LENGTH=420 |
| AT5G05600.1 | 0,544537363 | 0,004998524 | JAO2_ JOX2 JASMONATE-INDUCED OXYGENASE2_ Jasmonic Acid Oxidase 2 chr5:1672266-1674602 FORWARD I |
| AT3G63140.1 | 0,844258985 | 0,005013155 | CSP41A chloroplast stem-loop binding protein of 41 kDa chr3:23327006-23328620 REVERSE LENGTH=406 |
| AT3G62030.1 | 0,847775021 | 0,005019363 | ROC4_ CYP20-3 rotamase CYP 4_ cyclophilin 20-3 chr3:22973708-22975139 FORWARD LENGTH=260 |
| AT1G35680.1 | 0,815036461 | 0,005150555 | RPL21C_ ASD chloroplast ribosomal protein L21_ ATPase-in-Seed-Development chr1:13208777-13210246 FORWARD I |
| ATCG00820.1 | 0,379975495 | 0,00519988 | RPS19 ribosomal protein S19 chrc:84005-84283 REVERSE LENGTH=92 |
| AT2G27860.1 | 1,069966217 | 0,005231121 | AXS1 UDP-D-apiose/UDP-D-xylose synthase 1 chr2:11864684-11866843 REVERSE LENGTH=389 |

| | | | |
|---|---|---|---|
| AT5G59890.2 | 0,595687207 | 0,005315395 | ATADF4_ ADF4 actin depolymerizing factor 4 chr5:24123107-24123596 FORWARD LENGTH=132 |
| AT3G61470.1 | 0,884491709 | 0,005353159 | LHCA2 photosystem I light harvesting complex gene 2 chr3:22745736-22747032 FORWARD LENGTH=257 |
| AT1G23190.1 | 1,329982736 | 0,005376113 | PGM3 phosphoglucomutase 3 chr1:8219946-8224186 FORWARD LENGTH=583 |
| AT5G66680.1 | 1,763260026 | 0,005411282 | DGL1 DEFECTIVE GLYCOSYLATION chr5:26617840-26620581 REVERSE LENGTH=437 |
| AT5G48300.1 | 1,342852999 | 0,005465486 | ADG1_ APS1 ADP-GLUCOSE PYROPHOSPHORYLASE SMALL SUBUNIT 1_ ADP glucose pyrophosphorylase 1 chr5 |
| AT1G79210.1 | 0,906445571 | 0,00549408 | no symbol available no full name available chr1:29796286-29798240 REVERSE LENGTH=235 |
| AT2G25080.1 | 0,67575888 | 0,005496183 | GPX1_ ATGPX1_ GPXL1 GLUTATHIONE PEROXIDASE 1_ glutathione peroxidase 1 chr2:10668134-10669828 FORW |
| AT4G14030.1 | 1,191986689 | 0,005530382 | SBP1_ AtSBP1 selenium-binding protein 1 chr4:8098121-8100165 REVERSE LENGTH=490 |
| AT1G56070.1 | 1,415274021 | 0,005534577 | LOS1 LOW EXPRESSION OF OSMOTICALLY RESPONSIVE GENES 1 chr1:20968245-20971077 REVERSE LENGT |
| AT4G08870.1 | 0,669309619 | 0,005558807 | ARGAH2 arginine amidohydrolase 2 chr4:5646654-5648693 REVERSE LENGTH=344 |
| AT3G52150.1 | 0,793119789 | 0,005565219 | PSRP2 plastid-speciﬁc ribosomal protein 2 chr3:19342074-19343090 FORWARD LENGTH=253 |
| AT1G70770.1 | 1,439874796 | 0,005578469 | no symbol available no full name available chr1:26688622-26691185 REVERSE LENGTH=610 |
| AT2G33070.1 | 0,280765094 | 0,00559742 | ATNSP2_ NSP2 NITRILE-SPECIFIER PROTEIN 2_ nitrile specifier protein 2 chr2:14029350-14030934 REVERSE LEN |
| AT5G39730.1 | 0,648745788 | 0,005656449 | no symbol available no full name available chr5:15901740-15902624 FORWARD LENGTH=172 |
| AT1G75940.1 | 0,638176981 | 0,005665289 | ATA27_ BGLU20 BETA GLUCOSIDASE 20 chr1:28511198-28514044 FORWARD LENGTH=535 |
| AT1G22700.3 | 0,390785988 | 0,005674264 | PYG7 chr1:8028323-8029289 REVERSE LENGTH=211 |
| AT1G72150.1 | 0,903777908 | 0,005675635 | PATL1 PATELLIN 1 chr1:27148558-27150652 FORWARD LENGTH=573 |
| AT1G06950.1 | 1,304487083 | 0,005685162 | ATTIC110_ TIC110 ARABIDOPSIS THALIANA TRANSLOCON AT THE INNER ENVELOPE MEMBRANE OF CHL |
| AT3G25760.1 | 0,656373462 | 0,005694774 | AOC1_ ERD12 early-responsive to dehydration 12_ allene oxide cyclase 1 chr3:9403972-9405105 FORWARD LENGTH= |
| AT4G15210.3 | 1,44908832 | 0,005698103 | AT-BETA-AMY_ ATBETA-AMY_ BMY1_ RAM1_ BAM5 REDUCED BETA AMYLASE 1_ beta-amylase 5_ ARABID |
| AT2G42520.1 | 1,690597899 | 0,005742104 | RH37 RNA Helicase 37 chr2:17705382-17708744 FORWARD LENGTH=633 |
| AT1G58290.1 | 0,424596176 | 0,005768047 | HEMA1_ GluTR_ AtHEMA1 glutamyl-tRNA reductase_ Arabidopsis thaliana hemA 1 chr1:21624028-21626051 REVERS |
| AT5G01530.1 | 0,88684801 | 0,005794661 | LHCB4.1 light harvesting complex photosystem II chr5:209084-210243 FORWARD LENGTH=290 |
| AT2G45740.1 | 1,814716339 | 0,00579571 | PEX11D peroxin 11D chr2:18839865-18841102 FORWARD LENGTH=236 |
| AT1G80380.4 | 1,13399146 | 0,005811368 | GLYK glycerate kinase chr1:30217332-30219784 FORWARD LENGTH=450 |
| AT1G23730.1 | 0,441034837 | 0,005814496 | ATBCA3_ BCA3 beta carbonic anhydrase 3_ BETA CARBONIC ANHYDRASE 3 chr1:8395965-8398014 FORWARD L |
| ATCG00905.1 | 0,513322273 | 0,005863906 | RPS12_ RPS12C RIBOSOMAL PROTEIN S12_ ribosomal protein S12C chrc:97999-98793 REVERSE LENGTH=85 |
| ATCG00420.1 | 0,479897904 | 0,005866082 | NDHJ NADH dehydrogenase subunit J chrc:48677-49153 REVERSE LENGTH=158 |
| AT1G22450.1 | 0,517932632 | 0,005894946 | ATCOX6B2_ COX6B cytochrome C oxidase 6B_ CYTOCHROME C OXIDASE 6B2 chr1:7925447-7926918 FORWARD |
| AT3G56460.1 | 1,626718981 | 0,00591747 | no symbol available no full name available chr3:20933029-20934425 REVERSE LENGTH=348 |
| AT1G73230.1 | 2,088350431 | 0,005938779 | no symbol available no full name available chr1:27540506-27541364 REVERSE LENGTH=165 |
| AT4G08390.3 | 1,221651085 | 0,005992975 | SAPX stromal ascorbate peroxidase chr4:5314999-5317071 FORWARD LENGTH=371 |
| AT1G74050.1 | 0,764818739 | 0,006064184 | no symbol available no full name available chr1:27847256-27848680 REVERSE LENGTH=233 |
| AT1G76160.1 | 3,533293336 | 0,006190313 | sks5 SKU5 similar 5 chr1:28578211-28581020 REVERSE LENGTH=541 |
| AT4G37000.1 | 0,713759547 | 0,006191292 | ACD2_ ATRCCR ARABIDOPSIS THALIANA RED CHLOROPHYLL CATABOLITE REDUCTASE_ ACCELERATED |
| AT3G18190.1 | 2,506794985 | 0,006227341 | CCT4 Chaperonin containing T-complex polypeptide-1 subunit 4 chr3:6232226-6233836 FORWARD LENGTH=536 |
| AT3G43980.1 | 0,58672543 | 0,006280411 | no symbol available no full name available chr3:15778555-15779235 REVERSE LENGTH=56 |
| AT2G30860.1 | 0,828354871 | 0,006479144 | GSTF9_ ATGSTF9_ ATGSTF7_ GLUTTR glutathione S-transferase PHI 9 chr2:13139132-13140057 FORWARD LENGT |
| AT4G25370.1 | 0,768967932 | 0,006518819 | ClpT1 chr4:12972747-12974580 FORWARD LENGTH=238 |

| | | | |
|---|---|---|---|
| AT3G52500.1 | 0,750264014 | 0,006544108 | no symbol available no full name available chr3:19465644-19467053 REVERSE LENGTH=469 |
| AT2G37040.1 | 2,100239665 | 0,006546307 | PAL1_ ATPAL1 PHE ammonia lyase 1 chr2:15557602-15560237 REVERSE LENGTH=725 |
| AT4G01800.1 | 1,476834385 | 0,006656795 | SECA1_ AGY1_ AtcpSecA Arabidopsis thaliana chloroplast SecA_ Albino or Glassy Yellow 1 chr4:770926-776131 REVE |
| AT1G52000.1 | 0,717238608 | 0,006698338 | no symbol available no full name available chr1:19333352-19335700 REVERSE LENGTH=730 |
| AT1G53750.1 | 1,37905184 | 0,006753558 | RPT1A regulatory particle triple-A 1A chr1:20065921-20068324 REVERSE LENGTH=426 |
| AT5G06600.2 | 2,0491881 | 0,006818014 | UBP12_ AtUBP12 ubiquitin-specific protease 12 chr5:2019545-2027834 REVERSE LENGTH=1115 |
| AT4G17560.1 | 0,500590405 | 0,006822503 | no symbol available no full name available chr4:9780343-9781752 FORWARD LENGTH=225 |
| AT5G48480.1 | 2,601917554 | 0,006874835 | no symbol available no full name available chr5:19644814-19645658 FORWARD LENGTH=166 |
| AT2G18800.1 | 1,817971585 | 0,00688303 | ATXTH21_ XTH21 xyloglucan endotransglucosylase/hydrolase 21_ XYLOGLUCAN ENDOTRANSGLUCOSYLASE/HY |
| AT5G28510.1 | 0,494492237 | 0,007028856 | BGLU24 beta glucosidase 24 chr5:10481041-10484022 REVERSE LENGTH=533 |
| AT2G37270.1 | 1,148935502 | 0,0070363 | RPS5B_ ATRPS5B ribosomal protein 5B chr2:15647883-15649042 REVERSE LENGTH=207 |
| AT2G37660.1 | 0,794222682 | 0,007088653 | no symbol available no full name available chr2:15795481-15796977 REVERSE LENGTH=325 |
| AT4G27090.1 | 0,526145823 | 0,007165777 | RPL14B chr4:13594104-13595187 REVERSE LENGTH=134 |
| AT3G48420.1 | 0,948992185 | 0,007201488 | no symbol available no full name available chr3:17929743-17931551 FORWARD LENGTH=319 |
| AT5G08670.1 | 1,213384549 | 0,007205484 | no symbol available no full name available chr5:2818395-2821149 REVERSE LENGTH=556 |
| AT3G63490.1 | 0,827762934 | 0,007208639 | EMB3126_ PRPL1 proline-rich protein-like 1_ plastid ribosomal protein L1_ EMBRYO DEFECTIVE 3126 chr3:2344426 |
| AT4G14040.1 | 1,17872501 | 0,007221654 | EDA38_ SBP2 selenium-binding protein 2_ EMBRYO SAC DEVELOPMENT ARREST 38 chr4:8100691-8102828 REVE |
| AT5G51070.1 | 0,815388848 | 0,007222346 | SAG15_ ERD1_ CLPD EARLY RESPONSIVE TO DEHYDRATION 1_ SENESCENCE ASSOCIATED GENE 15 chr5:2 |
| AT5G12040.1 | 1,431361648 | 0,007272646 | no symbol available no full name available chr5:3885162-3887772 FORWARD LENGTH=369 |
| AT2G19940.1 | 1,249911343 | 0,007358794 | no symbol available no full name available chr2:8613203-8615649 FORWARD LENGTH=389 |
| AT1G12000.1 | 1,769891675 | 0,007521245 | no symbol available no full name available chr1:4050159-4053727 REVERSE LENGTH=566 |
| AT5G10860.1 | 0,807758172 | 0,007597043 | CBSX3 CBS domain containing protein 3 chr5:3429173-3430142 REVERSE LENGTH=206 |
| AT4G02450.2 | 1,164103255 | 0,007658481 | p23-1 chr4:1073987-1075765 REVERSE LENGTH=240 |
| AT5G53850.1 | 0,680693257 | 0,007709191 | DEP1 DEHYDRATASE-ENOLASE-PHOSPHATASE-COMPLEX 1 chr5:21861617-21864817 REVERSE LENGTH=402 |
| AT2G47940.2 | 1,249720746 | 0,007731362 | DEG2_ DEGP2_ EMB3117 DEGP protease 2_ EMBRYO DEFECTIVE 3117_ degradation of periplasmic proteins 2 chr2: |
| AT4G35100.1 | 0,097581673 | 0,007732161 | PIP3A_ SIMIP_ PIP3_ PIP2;7 plasma membrane intrinsic protein 3_ PLASMA MEMBRANE INTRINSIC PROTEIN 3A_ |
| AT2G41530.1 | 3,098560481 | 0,007889 | SFGH_ ATSFGH S-formylglutathione hydrolase_ ARABIDOPSIS THALIANA S-FORMYLGLUTATHIONE HYDROLA |
| AT2G45710.1 | 1,096399288 | 0,007917832 | no symbol available no full name available chr2:18831243-18831999 FORWARD LENGTH=84 |
| AT3G20050.1 | 1,319551214 | 0,007998737 | ATTCP-1_ TCP-1_ CCT1 Chaperonin containing T-complex polypeptide-1 subunit 1_ T-complex protein 1 alpha subunit c |
| AT2G45790.1 | 0,725621335 | 0,008077549 | PMM_ ATPMM phosphomannomutase_ PHOSPHOMANNOMUTASE chr2:18855876-18857753 FORWARD LENGTH= |
| AT2G42530.1 | 1,441948728 | 0,008136162 | COR15B cold regulated 15b chr2:17709191-17709873 REVERSE LENGTH=141 |
| AT1G49240.1 | 1,374402642 | 0,008307407 | ACT8_ FIZ1 FRIZZY AND KINKED SHOOTS_ actin 8 chr1:18216539-18217947 FORWARD LENGTH=377 |
| AT1G02280.1 | 1,358890009 | 0,008393678 | TOC33_ PPI1_ ATTOC33 PLASTID PROTEIN IMPORT 1_ translocon at the outer envelope membrane of chloroplasts 3 |
| AT1G55670.1 | 0,793155521 | 0,008487708 | PSAG photosystem I subunit G chr1:20802874-20803356 REVERSE LENGTH=160 |
| AT1G05190.1 | 0,74928672 | 0,008526877 | RPL6_ EMB2394 embryo defective 2394 chr1:1502515-1503738 REVERSE LENGTH=223 |
| AT3G13120.1 | 0,730409382 | 0,008578735 | PRPS10 plastid ribosomal protein of the 30S subunit 10 chr3:4220310-4221526 REVERSE LENGTH=191 |
| AT3G10350.1 | 0,804432653 | 0,008581456 | AtGET3b_ GET3b Guided Entry of Tail-anchored proteins 3b chr3:3208310-3210678 FORWARD LENGTH=411 |
| AT3G56190.1 | 1,417606017 | 0,008656829 | ALPHA-SNAP2_ ASNAP alpha-soluble NSF attachment protein 2 chr3:20846119-20848356 REVERSE LENGTH=289 |
| AT2G44650.1 | 1,320134415 | 0,008685893 | CHL-CPN10_ CPN10 CHLOROPLAST CHAPERONIN 10_ chloroplast chaperonin 10 chr2:18419521-18420510 REVER |

| | | | |
|---|---|---|---|
| AT3G20000.1 | 1,374522915 | 0,008722253 | TOM40 translocase of the outer mitochondrial membrane 40 chr3:6967685-6970247 FORWARD LENGTH=309 |
| AT3G47800.1 | 0,71424263 | 0,008919938 | no symbol available no full name available chr3:17634971-17636998 FORWARD LENGTH=358 |
| AT3G13460.2 | 1,811424522 | 0,008962048 | ECT2 evolutionarily conserved C-terminal region 2 chr3:4385274-4388220 REVERSE LENGTH=664 |
| AT1G78570.1 | 1,56607465 | 0,009017065 | RHM1_ ATRHM1_ ROL1 REPRESSOR OF LRX1 1_ rhamnose biosynthesis 1_ ARABIDOPSIS THALIANA RHAMNO |
| AT1G29880.1 | 1,38687953 | 0,009041891 | no symbol available no full name available chr1:10459662-10462781 REVERSE LENGTH=729 |
| AT3G46780.1 | 1,223193656 | 0,009125337 | PTAC16 plastid transcriptionally active 16 chr3:17228766-17231021 FORWARD LENGTH=510 |
| AT4G26970.1 | 1,575587531 | 0,009142416 | ACO2 aconitase 2 chr4:13543077-13548427 FORWARD LENGTH=995 |
| AT5G61780.1 | 1,29428412 | 0,009408467 | Tudor2_ AtTudor2_ TSN2 Arabidopsis thaliana TUDOR-SN protein 2_ TUDOR-SN protein 2 chr5:24822012-24826641 F |
| AT4G01100.1 | 2,321642003 | 0,009711511 | ADNT1 adenine nucleotide transporter 1 chr4:477411-479590 FORWARD LENGTH=352 |
| AT2G23600.1 | 0,533722322 | 0,009790169 | ATMES2_ MES2_ ATME8_ ME8_ ACL ARABIDOPSIS THALIANA METHYL ESTERASE 2_ ARABIDOPSIS METH |
| AT1G35720.1 | 0,865763319 | 0,009790733 | ANNAT1_ ATOXY5_ OXY5_ ANN1_ AtANN1 annexin 1 chr1:13225304-13226939 FORWARD LENGTH=317 |
| AT3G58510.1 | 1,537167657 | 0,009853016 | RH11 RNA Helicase 11 chr3:21640608-21643464 FORWARD LENGTH=612 |
| AT5G63570.1 | 1,132660726 | 0,010020369 | GSA1 "glutamate-1-semialdehyde-2_1-aminomutase" chr5:25451957-25453620 FORWARD LENGTH=474 |
| AT4G12420.1 | 0,807160132 | 0,010065486 | SKU5 chr4:7349941-7352868 REVERSE LENGTH=587 |
| AT1G63660.2 | 1,973734385 | 0,010193187 | no symbol available no full name available chr1:23604874-23607080 REVERSE LENGTH=434 |
| AT5G42020.2 | 1,329087825 | 0,010207507 | BIP2_ BIP luminal binding protein chr5:16807697-16810480 REVERSE LENGTH=613 |
| AT5G15090.1 | 1,081779815 | 0,010214103 | VDAC3_ AtVDAC-3_ ATVDAC3 ARABIDOPSIS THALIANA VOLTAGE DEPENDENT ANION CHANNEL 3_ voltag |
| AT5G27770.1 | 0,767802803 | 0,010252957 | no symbol available no full name available chr5:9836166-9837113 FORWARD LENGTH=124 |
| AT4G35000.1 | 1,369901851 | 0,010481797 | APX3 ascorbate peroxidase 3 chr4:16665007-16667541 REVERSE LENGTH=287 |
| AT2G29550.1 | 1,37210841 | 0,010559978 | TUB7_ TBB7 tubulin beta-7 chain_ tubulin beta 7 chr2:12644258-12645932 REVERSE LENGTH=449 |
| AT1G22300.1 | 1,218755431 | 0,010610015 | 14-3-3EPSILON_ GRF10_ GF14 EPSILON general regulatory factor 10_ 14-3-3 PROTEIN G-BOX FACTOR14 EPSILO |
| AT1G04530.1 | 2,357297739 | 0,010747346 | TPR4 tetratricopeptide repeat 4 chr1:1234456-1235895 REVERSE LENGTH=310 |
| AT3G01280.1 | 0,844922353 | 0,010771879 | VDAC1_ ATVDAC1 ARABIDOPSIS THALIANA VOLTAGE DEPENDENT ANION CHANNEL 1_ voltage dependent |
| AT4G31500.1 | 1,61352204 | 0,010805325 | RNT1_ RED1_ SUR2_ ATR4_ CYP83B1 RED ELONGATED 1_ SUPERROOT 2_ ALTERED TRYPTOPHAN REGUL |
| AT5G55280.1 | 1,364719514 | 0,010856896 | CPFTSZ_ FTSZ1-1_ ATFTSZ1-1_ FtsZ1 CHLOROPLAST FTSZ_ homolog of bacterial cytokinesis Z-ring protein FTSZ 1 |
| AT2G04842.1 | 0,882919444 | 0,011001359 | EMB2761 EMBRYO DEFECTIVE 2761 chr2:1698466-1701271 REVERSE LENGTH=650 |
| AT2G33380.1 | 2,262377417 | 0,011073548 | AtRD20_ PXG3_ CLO-3_ CLO3_ RD20_ AtCLO3 caleosin 3_ Arabidopsis thaliana caleosin 3_ peroxygenase 3_ RESPON |
| AT2G21250.1 | 3,149493261 | 0,011087815 | no symbol available no full name available chr2:9103408-9105116 REVERSE LENGTH=309 |
| AT4G20850.1 | 0,829713192 | 0,011107774 | TPP2 tripeptidyl peptidase ii chr4:11160935-11169889 REVERSE LENGTH=1380 |
| AT3G17390.1 | 0,886417591 | 0,011400667 | SAMS3_ AtSAMS3_ MAT4_ MTO3 METHIONINE ADENOSYLTRANSFERASE 4_ S-ADENOSYLMETHIONINE SY |
| AT2G01290.1 | 1,448132462 | 0,011562324 | RPI2 ribose-5-phosphate isomerase 2 chr2:149192-149989 REVERSE LENGTH=265 |
| AT4G09000.1 | 1,165359221 | 0,011818458 | GRF1_ GF14 CHI GENERAL REGULATORY FACTOR1-G-BOX FACTOR 14-3-3 HOMOLOG ISOFORM CHI_ gener |
| AT1G69830.1 | 0,610492745 | 0,011840984 | ATAMY3_ AMY3 ALPHA-AMYLASE-LIKE 3_ alpha-amylase-like 3 chr1:26288518-26293003 REVERSE LENGTH=8 |
| AT4G05180.1 | 1,179582288 | 0,011916929 | PSBQ-2_ PSBQ_ PSII-Q photosystem II subunit Q-2_ PHOTOSYSTEM II SUBUNIT Q chr4:2672093-2673170 REVERS |
| AT2G04390.1 | 0,761606746 | 0,012111126 | dS17 chr2:1527911-1528336 FORWARD LENGTH=141 |
| AT2G13360.1 | 1,092786499 | 0,012151462 | SGAT_ AGT_ AGT1 ALANINE:GLYOXYLATE AMINOTRANSFERASE 1_ alanine:glyoxylate aminotransferase_ L-ser |
| AT5G53560.1 | 0,832361592 | 0,01220601 | ATB5-A_ CB5-E_ ATCB5-E_ B5 #2 ARABIDOPSIS CYTOCHROME B5 ISOFORM E_ cytochrome B5 isoform E chr5: |
| AT1G58080.1 | 0,86188301 | 0,012246685 | ATATP-PRT1_ ATP-PRT1_ HISN1A ATP phosphoribosyl transferase 1 chr1:21504562-21507429 REVERSE LENGTH= |
| ATCG00770.1 | 0,750712574 | 0,012269116 | RPS8 ribosomal protein S8 chrc:80068-80472 REVERSE LENGTH=134 |

| | | | |
|---|---|---|---|
| AT2G34420.1 | 1,432883534 | 0,012288843 | LHCB1.5_ LHB1B2 PHOTOSYSTEM II LIGHT HARVESTING COMPLEX GENE 1.5_ photosystem II light harvesting [illegible] |
| AT2G30050.1 | 1,592910486 | 0,012311346 | no symbol available no full name available chr2:12825540-12826448 FORWARD LENGTH=302 |
| AT5G10540.1 | 1,335792216 | 0,0123318 | TOP2 thimet metalloendopeptidase 2 chr5:3328119-3332462 FORWARD LENGTH=701 |
| AT5G62690.1 | 1,25169871 | 0,012428895 | TUB2 tubulin beta chain 2 chr5:25181560-25183501 FORWARD LENGTH=450 |
| AT3G58990.1 | 1,326934302 | 0,012488586 | IPMI SSU3_ IPMI1 isopropylmalate isomerase 1 chr3:21797524-21798285 REVERSE LENGTH=253 |
| AT1G26570.1 | 0,64948597 | 0,012552857 | UGD1_ ATUGD1 UDP-GLUCOSE DEHYDROGENASE 1_ UDP-glucose dehydrogenase 1 chr1:9182801-9184246 FOR[illegible] |
| AT4G01690.1 | 1,277885874 | 0,012662845 | PPO1_ PPOX_ HEMG1 chr4:729929-732309 FORWARD LENGTH=537 |
| AT5G63860.1 | 1,33965018 | 0,01267348 | AtUVR8_ UVR8 UVB-RESISTANCE 8 chr5:25554821-25558587 REVERSE LENGTH=440 |
| AT5G20950.1 | 1,27372739 | 0,012741323 | BGLC1 chr5:7107609-7110775 REVERSE LENGTH=624 |
| AT3G16410.1 | 0,798364859 | 0,012808946 | NSP4 nitrile specifier protein 4 chr3:5572145-5574359 FORWARD LENGTH=619 |
| AT1G22740.1 | 0,206894215 | 0,012886015 | RABG3B_ RAB7_ ATRABG3B_ RAB75 RAB GTPase homolog G3B chr1:8049247-8050494 FORWARD LENGTH=20[illegible] |
| AT3G22460.2 | 0,762342325 | 0,012890301 | OASA2 O-acetylserine (thiol) lyase (OAS-TL) isoform A2 chr3:7963855-7964769 FORWARD LENGTH=188 |
| AT1G11430.1 | 0,74375749 | 0,0128948 | RIP9_ MORF9 multiple organellar RNA editing factor 9 chr1:3847273-3848938 FORWARD LENGTH=232 |
| AT1G12250.1 | 0,598621871 | 0,013002364 | TL20.3 chr1:4159287-4161269 FORWARD LENGTH=280 |
| AT5G43330.1 | 0,915952131 | 0,013128234 | c-NAD-MDH2 cytosolic-NAD-dependent malate dehydrogenase 2 chr5:17390552-17392449 FORWARD LENGTH=332 |
| AT1G09340.1 | 1,152942819 | 0,013196855 | CRB_ CSP41B_ HIP1.3 chloroplast RNA binding_ heteroglycan-interacting protein 1.3_ CHLOROPLAST STEM-LOOP [illegible] |
| AT5G26742.1 | 1,19996043 | 0,013290953 | AtRH3_ RH3_ emb1138 embryo defective 1138 chr5:9285540-9288871 REVERSE LENGTH=747 |
| AT5G48880.1 | 0,72373684 | 0,013487092 | PKT2_ PKT1_ KAT5 3-KETO-ACYL-COENZYME A THIOLASE 5_ peroxisomal 3-keto-acyl-CoA thiolase 2_ PEROXI[illegible] |
| AT5G44320.1 | 1,219379295 | 0,01366011 | no symbol available no full name available chr5:17854901-17856667 REVERSE LENGTH=588 |
| AT2G19760.1 | 0,81979095 | 0,013675555 | PFN1_ PRF1 profilin 1_ PROFILIN 1 chr2:8517074-8518067 REVERSE LENGTH=131 |
| AT5G54940.1 | 0,454437576 | 0,013687319 | no symbol available no full name available chr5:22308420-22308758 REVERSE LENGTH=112 |
| AT1G74100.1 | 0,771679548 | 0,013712375 | ATSOT16_ ATST5A_ CORI-7_ SOT16 sulfotransferase 16_ CORONATINE INDUCED-7_ SULFOTRANSFERASE 16_ |
| AT2G42690.1 | 1,322461833 | 0,013784599 | AGAP1 ACYLATED GALACTOLIPID- ASSOCIATED PHOSPHOLIPASE 1 chr2:17776356-17777682 REVERSE LEN[illegible] |
| AT3G19450.1 | 1,090008827 | 0,013804758 | CAD-C_ CAD4_ ATCAD4_ CAD CINNAMYL ALCOHOL DEHYDROGENASE 4 chr3:6744859-6747005 FORWARD [illegible] |
| AT3G04870.1 | 1,517733955 | 0,01398162 | PDE181_ ZDS_ SPC1 SPONTANEOUS CELL DEATH 1_ PIGMENT DEFECTIVE EMBRYO 181_ zeta-carotene desatu[illegible] |
| AT3G17820.1 | 4,123708918 | 0,014124185 | GLN1.3_ ATGSKB6_ GLN1;3 glutamine synthetase 1.3_ ARABIDOPSIS THALIANA GLUTAMINE SYNTHASE CLO[illegible] |
| AT5G41520.1 | 1,672703527 | 0,01424133 | RPS10B ribosomal protein S10e B chr5:16609377-16610583 REVERSE LENGTH=180 |
| AT4G01050.1 | 1,118553492 | 0,014306054 | TROL thylakoid rhodanese-like chr4:455874-458175 FORWARD LENGTH=466 |
| AT2G45300.3 | 2,853409507 | 0,01438709 | no symbol available no full name available chr2:18677518-18680118 FORWARD LENGTH=518 |
| AT4G12730.1 | 0,794265187 | 0,01443775 | FLA2 FASCICLIN-like arabinogalactan 2 chr4:7491598-7492809 REVERSE LENGTH=403 |
| AT1G01320.2 | 1,323616744 | 0,01465765 | REC1_ FLL2 FLOURY ENDOSPERM LIKE 2_ REDUCED CHLOROPLAST COVERAGE chr1:121582-130099 REVE[illegible] |
| AT1G70310.1 | 2,213590754 | 0,014800069 | SPDS2 spermidine synthase 2 chr1:26485497-26487352 REVERSE LENGTH=340 |
| AT2G19730.1 | 2,017174523 | 0,014846338 | no symbol available no full name available chr2:8511752-8512995 FORWARD LENGTH=143 |
| AT5G44020.1 | 4,630026808 | 0,014930925 | no symbol available no full name available chr5:17712433-17714046 FORWARD LENGTH=272 |
| AT5G16710.1 | 0,842272448 | 0,014931359 | DHAR3 dehydroascorbate reductase 1 chr5:5483312-5484926 FORWARD LENGTH=258 |
| AT2G21590.1 | 0,728582195 | 0,015002623 | APL4 chr2:9239362-9242150 FORWARD LENGTH=523 |
| AT3G56090.1 | 1,119207055 | 0,015138634 | FER3_ ATFER3 ferritin 3 chr3:20814350-20815984 REVERSE LENGTH=259 |
| AT4G23900.1 | 0,805634367 | 0,015193766 | no symbol available no full name available chr4:12424505-12426318 FORWARD LENGTH=237 |
| AT2G42910.1 | 2,940617091 | 0,01519696 | AtPRS4_ PRS4 phosphoribosyl diphosphate synthase 4 chr2:17856396-17858394 FORWARD LENGTH=337 |

| | | | |
|---|---|---|---|
| AT1G01200.1 | 1,677792813 | 0,015202831 | RABA3_ ATRABA3_ ATRAB-A3 ARABIDOPSIS RAB GTPASE HOMOLOG A3_ RAB GTPase homolog A3 chr1:867 |
| AT1G17880.1 | 0,642129818 | 0,015356625 | ATBTF3_ BTF3 basic transcription factor 3 chr1:6152572-6153425 REVERSE LENGTH=165 |
| AT2G31790.1 | 1,410217133 | 0,015568708 | no symbol available no full name available chr2:13518269-13520167 FORWARD LENGTH=457 |
| AT3G44320.1 | 0,786847233 | 0,015606809 | NIT3_ AtNIT3 NITRILASE 3_ nitrilase 3 chr3:15993419-15995493 FORWARD LENGTH=346 |
| AT1G29660.1 | 0,605631973 | 0,015716698 | GGL5 chr1:10371955-10373624 FORWARD LENGTH=364 |
| ATCG00650.1 | 0,499336878 | 0,015822921 | RPS18 ribosomal protein S18 chrc:67917-68222 FORWARD LENGTH=101 |
| AT3G53230.1 | 0,736568454 | 0,015897998 | AtCDC48B cell division cycle 48B chr3:19723416-19726489 FORWARD LENGTH=815 |
| AT5G54160.1 | 1,089982306 | 0,015926128 | ATOMT1_ OMT1_ OMT3_ AtCOMT_ COMT1 caffeate O-methyltransferase 1_ O-methyltransfera |
| AT5G13630.1 | 1,46941885 | 0,016018209 | ABAR_ CHLH_ GUN5_ CCH_ CCH1 ABA-BINDING PROTEIN_ H SUBUNIT OF MG-CHELATASE_ GENOMES UN |
| AT5G38410.2 | 0,816295075 | 0,016219355 | RBCS3B Rubisco small subunit 3B chr5:15377501-15378306 REVERSE LENGTH=174 |
| AT5G08280.1 | 0,852633546 | 0,016227157 | HEMC_ RUG1 RUGOSA 1_ hydroxymethylbilane synthase chr5:2663763-2665596 REVERSE LENGTH=382 |
| AT1G34430.1 | 1,170119291 | 0,01632719 | EMB3003 embryo defective 3003 chr1:12588027-12590084 REVERSE LENGTH=465 |
| AT4G14800.1 | 2,565924215 | 0,01635415 | PBD2 20S proteasome beta subunit D2 chr4:8500398-8502058 FORWARD LENGTH=199 |
| AT5G56350.1 | 1,517185885 | 0,016725594 | no symbol available no full name available chr5:22820254-22822529 REVERSE LENGTH=498 |
| AT4G20890.1 | 0,882920286 | 0,016763028 | TUB9 tubulin beta-9 chain chr4:11182218-11183840 FORWARD LENGTH=444 |
| AT1G16080.1 | 1,228775125 | 0,016935512 | no symbol available no full name available chr1:5514394-5515761 FORWARD LENGTH=313 |
| AT1G09780.1 | 1,1295418 | 0,017045067 | iPGAM1 "2_3-biphosphoglycerate-independent phosphoglycerate mutase 1" chr1:3165550-3167812 REVERSE LENGTH= |
| AT3G25800.1 | 1,232849797 | 0,017088394 | PP2AA2_ PDF1_ PR 65 protein phosphatase 2A subunit A2 chr3:9422822-9425783 REVERSE LENGTH=587 |
| AT4G39980.1 | 0,902685309 | 0,017399851 | AtDAHP1_ DHS1_ DAHP1 3-DEOXY-D-ARABINO-HEPTULOSONATE-7-PHOSPHATE 1_ 3-deoxy-D-arabino-heptul |
| AT5G19220.1 | 0,839280441 | 0,01752389 | ADG2_ APL1 ADP glucose pyrophosphorylase large subunit 1_ ADP GLUCOSE PYROPHOSPHORYLASE 2 chr5:6463 |
| AT4G21960.1 | 0,104889605 | 0,017585853 | PRXR1 chr4:11646613-11648312 REVERSE LENGTH=330 |
| AT1G70730.1 | 1,142178416 | 0,017671427 | PGM2 phosphoglucomutase 2 chr1:26669020-26672726 REVERSE LENGTH=585 |
| AT3G19170.1 | 0,853021807 | 0,017829736 | ATZNMP_ ATPREP1_ PREP1 presequence protease 1 chr3:6625578-6631874 REVERSE LENGTH=1080 |
| AT1G77940.1 | 0,453806625 | 0,017863158 | RPL30B chr1:29304116-29305288 REVERSE LENGTH=112 |
| AT3G12260.1 | 0,688700481 | 0,017938675 | NDUFA6_ B14 chr3:3909252-3910337 REVERSE LENGTH=133 |
| AT1G65930.1 | 1,160837014 | 0,01809023 | cICDH cytosolic NADP+-dependent isocitrate dehydrogenase chr1:24539088-24541861 FORWARD LENGTH=410 |
| AT5G27670.1 | 0,837349801 | 0,018208808 | HTA7_ h2a.w.7 histone H2A 7 chr5:9792807-9793365 REVERSE LENGTH=150 |
| AT4G02530.1 | 1,179523491 | 0,018215993 | MPH2 MAINTENANCE OF PHOTOSYSTEM II UNDER HIGH LIGHT 2 chr4:1112335-1114005 REVERSE LENGTH= |
| AT5G48130.1 | 1,226842019 | 0,018248088 | no symbol available no full name available chr5:19516291-19518450 FORWARD LENGTH=625 |
| AT5G22300.1 | 0,52819908 | 0,018321406 | AtNIT4_ NIT4 nitrilase 4_ NITRILASE 4 chr5:7379401-7381764 FORWARD LENGTH=355 |
| AT1G70890.1 | 1,404144176 | 0,018421427 | MLP43 MLP-like protein 43_ major latex protein like 43 chr1:26725912-26726489 REVERSE LENGTH=158 |
| AT2G25060.1 | 0,726662791 | 0,018522005 | ENODL14_ AtENODL14 early nodulin-like protein 14 chr2:10662308-10662930 FORWARD LENGTH=182 |
| AT4G03280.1 | 0,880848485 | 0,018608116 | PGR1_ PETC PROTON GRADIENT REGULATION 1_ photosynthetic electron transfer C chr4:1440314-1441717 FORW |
| AT4G22485.1 | 1,300351086 | 0,018702481 | no symbol available no full name available chr4:11844506-11846476 REVERSE LENGTH=656 |
| AT2G21530.1 | 0,589662309 | 0,018847029 | no symbol available no full name available chr2:9219372-9220464 FORWARD LENGTH=209 |
| AT5G47190.1 | 0,852669409 | 0,018908434 | PRPL19 plastid ribosomal proteins of the 50S subunit 19 chr5:19164432-19166064 REVERSE LENGTH=229 |
| AT4G39080.1 | 0,772150891 | 0,019149157 | VHA-A3 vacuolar proton ATPase A3 chr4:18209513-18214752 FORWARD LENGTH=821 |
| AT2G21170.1 | 0,940639977 | 0,01916094 | TIM_ PDTPI triosephosphate isomerase_ PLASTID ISOFORM TRIOSE PHOSPHATE ISOMERASE chr2:9071047-9073 |
| AT3G22460.1 | 1,862625031 | 0,019513109 | OASA2 O-acetylserine (thiol) lyase (OAS-TL) isoform A2 chr3:7964204-7965751 FORWARD LENGTH=250 |

| | | | |
|---|---|---|---|
| AT1G04410.1 | 1,070236855 | 0,019534028 | c-NAD-MDH1 cytosolic-NAD-dependent malate dehydrogenase 1 chr1:1189418-1191267 REVERSE LENGTH=332 |
| AT4G01310.1 | 0,803058908 | 0,019638794 | PRPL5 plastid ribosomal proteins of the 50S subunit 5 chr4:544166-545480 REVERSE LENGTH=262 |
| AT4G34620.1 | 0,862588519 | 0,019640536 | SSR16 small subunit ribosomal protein 16 chr4:16535084-16536092 REVERSE LENGTH=113 |
| ATCG00730.1 | 1,218063227 | 0,019857955 | PETD photosynthetic electron transfer D chrc:76481-77672 FORWARD LENGTH=160 |
| AT5G49360.1 | 0,676905183 | 0,019950389 | ATBXL1_ BXL1 beta-xylosidase 1_ BETA-XYLOSIDASE 1 chr5:20012179-20016659 REVERSE LENGTH=774 |
| AT1G26480.1 | 1,409821645 | 0,020401623 | GF14 IOTA_ GRF12 general regulatory factor 12 chr1:9156573-9157845 REVERSE LENGTH=268 |
| AT4G11420.1 | 1,922600046 | 0,020462627 | ATTIF3A1_ TIF3A1_ EIF3A-1_ ATEIF3A-1_ EIF3A eukaryotic translation initiation factor 3A chr4:6947834-6952053 RI |
| AT5G27470.1 | 1,391630847 | 0,020583849 | no symbol available no full name available chr5:9695087-9697154 FORWARD LENGTH=451 |
| AT3G15840.2 | 0,867091571 | 0,020705035 | PIFI post-illumination chlorophyll fluorescence increase chr3:5356782-5358418 REVERSE LENGTH=265 |
| AT5G44720.1 | 0,533625235 | 0,02092221 | no symbol available no full name available chr5:18043086-18045275 FORWARD LENGTH=308 |
| AT5G58290.1 | 0,621287613 | 0,020946446 | RPT3 regulatory particle triple-A ATPase 3 chr5:23569155-23571116 FORWARD LENGTH=408 |
| AT3G11630.1 | 1,061240968 | 0,020983274 | 2CPA 2-Cys peroxiredoxin A chr3:3672189-3673937 FORWARD LENGTH=266 |
| AT1G47420.1 | 0,813344597 | 0,0210329 | SDH5 succinate dehydrogenase 5 chr1:17395774-17397176 REVERSE LENGTH=257 |
| AT2G42170.1 | 1,389157406 | 0,021079551 | no symbol available no full name available chr2:17578683-17580222 FORWARD LENGTH=329 |
| AT4G08900.1 | 0,617813352 | 0,021239489 | ARGAH1 arginine amidohydrolase 1 chr4:5703499-5705180 FORWARD LENGTH=342 |
| AT5G42080.3 | 1,217907555 | 0,021241802 | DRP1A_ ADL1A_ DL1_ AG68_ RSW9_ ADL1 RADIAL SWELLING 9_ DYNAMIN-RELATED PROTEIN 1A_ dynam |
| AT5G23060.1 | 0,787719555 | 0,021249796 | CaS calcium sensing receptor chr5:7736760-7738412 REVERSE LENGTH=387 |
| AT5G16240.1 | 0,862297187 | 0,02131834 | AAD1 ACYL-ACYL CARRIER PROTEIN DESATURASE1 chr5:5306981-5309639 FORWARD LENGTH=394 |
| AT1G56050.1 | 0,803176321 | 0,021496173 | EngD-2 chr1:20963793-20966181 FORWARD LENGTH=421 |
| AT2G47470.1 | 1,19260773 | 0,021503237 | UNE5_ MEE30_ ATPDI11_ PDI11_ ATPDIL2-1 PROTEIN DISULFIDE ISOMERASE 11_ PDI-LIKE 2-1_ ARABIDOP |
| AT1G78060.1 | 0,727855193 | 0,021552626 | no symbol available no full name available chr1:29349796-29352868 REVERSE LENGTH=767 |
| AT1G80410.1 | 1,284961737 | 0,021631561 | OMA_ EMB2753_ NAA15_ MUSE6 "mutant_snc1-enhancing 6"_ EMBRYO DEFECTIVE 2753_ OMISHA_ NAA15 chr |
| AT5G22580.1 | 1,163331986 | 0,021743367 | no symbol available no full name available chr5:7502709-7503137 FORWARD LENGTH=111 |
| AT1G27450.1 | 1,245955889 | 0,021860124 | APT1_ ATAPT1 ARABIDOPSIS THALIANA ADENINE PHOSPHORIBOSYLTRANSFERASE 1_ adenine phosphoribo |
| AT1G48830.1 | 0,733833596 | 0,021932438 | no symbol available no full name available chr1:18059854-18060935 REVERSE LENGTH=191 |
| AT3G61430.1 | 1,774163403 | 0,021963314 | ATPIP1_ PIP1A_ PIP1_ PIP1;1 plasma membrane intrinsic protein 1A_ PLASMA MEMBRANE INTRINSIC PROTEIN 1 |
| AT2G04400.1 | 1,242204309 | 0,021969453 | IGPS Indole-3-glycerol phosphate synthase chr2:1531208-1533578 FORWARD LENGTH=402 |
| AT3G06050.1 | 0,779308182 | 0,022085962 | PRXIIF_ ATPRXIIF peroxiredoxin IIF_ PEROXIREDOXIN IIF chr3:1826311-1827809 REVERSE LENGTH=201 |
| AT1G75280.1 | 2,676656801 | 0,022090185 | no symbol available no full name available chr1:28252030-28253355 FORWARD LENGTH=310 |
| AT1G02780.1 | 0,591700347 | 0,022291067 | emb2386 embryo defective 2386 chr1:608120-609391 REVERSE LENGTH=214 |
| AT3G09820.1 | 1,097490187 | 0,022301584 | ATADK1_ ADK1 adenosine kinase 1 chr3:3012122-3014624 FORWARD LENGTH=344 |
| AT3G20390.1 | 0,866577467 | 0,022342141 | RidA Reactive Intermediate Deaminase A chr3:7110227-7111695 REVERSE LENGTH=187 |
| AT5G35790.1 | 1,366015836 | 0,022414204 | G6PD1 glucose-6-phosphate dehydrogenase 1 chr5:13956879-13959686 REVERSE LENGTH=576 |
| AT4G02930.1 | 1,118986965 | 0,022445347 | no symbol available no full name available chr4:1295751-1298354 REVERSE LENGTH=454 |
| AT1G20340.1 | 0,769723101 | 0,022537933 | PETE2_ DRT112 DNA-DAMAGE-REPAIR/TOLERATION PROTEIN 112_ PLASTOCYANIN 2 chr1:7042770-704327 |
| AT3G11710.1 | 1,595521843 | 0,022686224 | ATKRS-1 lysyl-tRNA synthetase 1 chr3:3702359-3705613 REVERSE LENGTH=626 |
| AT3G47520.1 | 0,851256466 | 0,022971673 | MDH_ pNAD-MDH plastidic NAD-dependent malate dehydrogenase_ malate dehydrogenase chr3:17513657-17514868 FC |
| AT2G34810.1 | 0,393921627 | 0,022994266 | AtBBE16 chr2:14685292-14686914 FORWARD LENGTH=540 |
| AT2G44350.1 | 1,266824732 | 0,023155699 | ATCS_ CSY4 CITRATE SYNTHASE 4 chr2:18316673-18320524 FORWARD LENGTH=473 |

| | | | |
|---|---|---|---|
| AT5G35630.1 | 0,89399492 | 0,023163475 | GLN2_ GS2_ ATGSL1 GLUTAMINE SYNTHETASE 2_ glutamine synthetase 2_ GLUTAMINE SYNTHETASE LIKE 1 |
| AT2G26080.1 | 0,967137935 | 0,023206747 | GLDP2_ AtGLDP2 glycine decarboxylase P-protein 2 chr2:11109330-11113786 REVERSE LENGTH=1044 |
| AT1G52510.1 | 0,883984423 | 0,023265755 | no symbol available no full name available chr1:19563039-19565260 REVERSE LENGTH=380 |
| AT5G53480.1 | 2,011286362 | 0,02363486 | AtKPNB1_ KPNB1_ IMB1 homolog of human KPNB1 chr5:21714016-21716709 FORWARD LENGTH=870 |
| AT4G15440.1 | 0,528766053 | 0,023915318 | HPL1_ CYP74B2 hydroperoxide lyase 1 chr4:8835869-8838462 FORWARD LENGTH=384 |
| AT3G55330.1 | 0,225943605 | 0,023969044 | PPL1 PsbP-like protein 1 chr3:20514031-20515275 REVERSE LENGTH=230 |
| AT5G63680.1 | 1,817359632 | 0,024268269 | no symbol available no full name available chr5:25490507-25492530 FORWARD LENGTH=510 |
| AT1G09310.1 | 0,772573943 | 0,024641643 | SVB2_ SVBL SVB-like chr1:3009109-3009648 FORWARD LENGTH=179 |
| AT2G19480.3 | 0,716947279 | 0,024932801 | NFA2_ NFA02_ NAP1;2 NUCLEOSOME/CHROMATIN ASSEMBLY FACTOR GROUP A 02_ nucleosome assembly pr |
| AT3G14420.1 | 1,084267403 | 0,024957876 | GOX1 glycolate oxidase 1 chr3:4821804-4823899 FORWARD LENGTH=367 |
| AT2G44120.1 | 0,85289551 | 0,025242631 | no symbol available no full name available chr2:18249227-18250402 REVERSE LENGTH=242 |
| AT4G14130.1 | 2,003958919 | 0,025305561 | XTR7_ XTH15 xyloglucan endotransglucosylase/hydrolase 15_ xyloglucan endotransglycosylase 7 chr4:8137161-8138196 |
| AT4G14890.1 | 0,62275929 | 0,025635564 | FdC1 ferredoxin C 1 chr4:8520887-8521351 FORWARD LENGTH=154 |
| AT4G32470.1 | 1,329795566 | 0,025754425 | no symbol available no full name available chr4:15669641-15671095 REVERSE LENGTH=122 |
| AT1G63770.2 | 0,769381318 | 0,026047945 | no symbol available no full name available chr1:23658165-23664243 REVERSE LENGTH=945 |
| AT3G62410.1 | 1,388194227 | 0,026096418 | CP12_ CP12-2 CP12 DOMAIN-CONTAINING PROTEIN 1_ CP12 domain-containing protein 2 chr3:23091006-2309140 |
| AT3G63170.1 | 0,481810431 | 0,026155699 | FAP1_ AtFAP1 fatty-acid-binding protein 1 chr3:23334675-23335993 FORWARD LENGTH=279 |
| AT2G16850.1 | 0,566946945 | 0,02633388 | PIP2;8_ PIP3B PLASMA MEMBRANE INTRINSIC PROTEIN 3B_ plasma membrane intrinsic protein 2;8 chr2:7301647- |
| AT5G08380.1 | 0,752936775 | 0,026421823 | AtAGAL1_ AGAL1 alpha-galactosidase 1 chr5:2694851-2697616 REVERSE LENGTH=410 |
| AT1G56110.1 | 1,558053853 | 0,026509415 | NOP56 homolog of nucleolar protein NOP56 chr1:20984544-20986893 REVERSE LENGTH=522 |
| AT2G17630.1 | 0,708333122 | 0,026515193 | PSAT2 phosphoserine aminotransferase 2 chr2:7666637-7667905 FORWARD LENGTH=422 |
| AT3G49680.2 | 1,476302529 | 0,026566566 | BCAT3_ ATBCAT-3 branched-chain aminotransferase 3 chr3:18422768-18425473 FORWARD LENGTH=411 |
| AT3G62120.3 | 1,185573004 | 0,026618667 | ProRS-Cyt_ AtProRS-Cyt prolyl-tRNA synthetase cytosolic chr3:23001227-23003849 REVERSE LENGTH=517 |
| AT1G04170.1 | 1,588288886 | 0,026637164 | EIF2 GAMMA eukaryotic translation initiation factor 2 gamma subunit chr1:1097423-1099702 FORWARD LENGTH=465 |
| AT1G11860.1 | 0,842606468 | 0,0266981 | GLDT chr1:4001801-4003245 FORWARD LENGTH=408 |
| AT2G27720.1 | 0,733201573 | 0,026829103 | no symbol available no full name available chr2:11818696-11819370 FORWARD LENGTH=115 |
| ATCG01120.1 | 0,586291674 | 0,02682926 | RPS15 chloroplast ribosomal protein S15 chrc:123296-123562 REVERSE LENGTH=88 |
| ATCG00130.1 | 1,089080196 | 0,026970181 | ATPF chrc:11529-12798 REVERSE LENGTH=184 |
| AT2G20420.1 | 0,908391947 | 0,027059684 | no symbol available no full name available chr2:8805574-8807858 FORWARD LENGTH=421 |
| AT2G32060.1 | 1,073313927 | 0,027199938 | no symbol available no full name available chr2:13639228-13640104 REVERSE LENGTH=144 |
| AT3G10670.1 | 0,824533112 | 0,027482387 | ABCI6_ ATNAP7_ NAP7 non-intrinsic ABC protein 7_ ATP-binding cassette I6 chr3:3335325-3337304 REVERSE LENG |
| AT5G34850.1 | 0,854197567 | 0,027569958 | PUP3_ PAP26_ ATPAP26 purple acid phosphatase 26_ phosphatase-under producer 3_ PURPLE ACID PHOSPHATASE 2 |
| AT3G27925.1 | 0,845778591 | 0,027586294 | DEGP1_ DEG1 degradation of periplasmic proteins 1_ DegP protease 1 chr3:10366659-10368864 REVERSE LENGTH=4 |
| AT4G02080.1 | 0,915121918 | 0,027620716 | ATSARA1C_ SAR2_ SAR1C_ ATSAR2_ ASAR1 secretion-associated RAS super family 2 chr4:921554-922547 FORWA |
| ATCG00380.1 | 0,52175569 | 0,028000085 | RPS4 chloroplast ribosomal protein S4 chrc:45223-45828 REVERSE LENGTH=201 |
| AT2G29440.1 | 1,422542295 | 0,028031147 | GST24_ ATGSTU6_ GSTU6 glutathione S-transferase tau 6_ GLUTATHIONE S-TRANSFERASE 24 chr2:12620159-126 |
| AT5G56010.1 | 1,503029687 | 0,02824695 | AtHsp90-3_ AtHsp90.3_ Hsp81.3_ HSP81-3 HEAT SHOCK PROTEIN 90-3_ HEAT SHOCK PROTEIN 81.3_ heat shock |
| AT3G27300.1 | 1,276970429 | 0,028283348 | G6PD5 glucose-6-phosphate dehydrogenase 5 chr3:10083318-10086288 REVERSE LENGTH=516 |
| AT1G22410.1 | 0,599179583 | 0,02833352 | no symbol available no full name available chr1:7912120-7914742 FORWARD LENGTH=527 |

| | | | |
|---|---|---|---|
| AT5G03940.1 | 0,875573908 | 0,028384395 | SRP54CP_ 54CP_ FFC_ CPSRP54 54 CHLOROPLAST PROTEIN_ chloroplast signal recognition particle 54 kDa subunit |
| AT2G14260.2 | 1,30409518 | 0,028627309 | PIP_ PAP1 proline iminopeptidase_ prolyl aminopeptidase 1 chr2:6041441-6043475 REVERSE LENGTH=329 |
| AT2G40490.1 | 0,804221574 | 0,028865981 | HEME2 chr2:16912961-16914988 FORWARD LENGTH=394 |
| AT2G35840.1 | 1,875549251 | 0,029154039 | no symbol available no full name available chr2:15053952-15055776 FORWARD LENGTH=422 |
| AT3G27240.1 | 0,720680663 | 0,029445048 | Cyc1-1 chr3:10056144-10058370 REVERSE LENGTH=307 |
| AT3G26520.1 | 1,218119666 | 0,029559302 | GAMMA-TIP2_ SITIP_ TIP2_ TIP1;2 SALT-STRESS INDUCIBLE TONOPLAST INTRINSIC PROTEIN_ tonoplast int |
| AT4G09320.1 | 1,090098113 | 0,029586886 | ATNDK1_ NDPK1_ NDK1 nucleoside diphosphate kinase 1 chr4:5923484-5924366 FORWARD LENGTH=149 |
| AT5G64570.3 | 0,83041645 | 0,029587077 | ATBXL4_ XYL4 ARABIDOPSIS THALIANA BETA-D-XYLOSIDASE 4_ beta-D-xylosidase 4 chr5:25810817-2581330 |
| AT4G01150.1 | 1,066817942 | 0,029794239 | CURT1A CURVATURE THYLAKOID 1A chr4:493692-494668 FORWARD LENGTH=164 |
| AT3G56070.1 | 0,671655277 | 0,030256031 | ROC2 rotamase cyclophilin 2 chr3:20806987-20807517 REVERSE LENGTH=176 |
| AT3G05530.1 | 1,222988807 | 0,030599783 | RPT5A_ ATS6A.2 regulatory particle triple-A ATPase 5A chr3:1603540-1605993 FORWARD LENGTH=424 |
| AT4G09670.1 | 1,659324198 | 0,03073541 | no symbol available no full name available chr4:6107382-6109049 REVERSE LENGTH=362 |
| AT1G32200.1 | 0,783821945 | 0,031104791 | ACT1_ ATS1 ACYLTRANSFERASE 1 chr1:11602223-11605001 REVERSE LENGTH=459 |
| AT5G12140.1 | 3,159061696 | 0,031349107 | ATCYS1_ CYS1 cystatin-1 chr5:3923295-3923936 REVERSE LENGTH=101 |
| AT5G50920.1 | 1,148202638 | 0,031544496 | DCA1_ CLPC_ ATHSP93-V_ CLPC1_ HSP93-V HEAT SHOCK PROTEIN 93-V_ CLPC homologue 1_ DE-REGULATI |
| AT3G44860.1 | 1,627455113 | 0,031574221 | FAMT farnesoic acid carboxyl-O-methyltransferase chr3:16379689-16380939 FORWARD LENGTH=348 |
| AT5G54600.1 | 0,443321394 | 0,031615515 | RPL24_ SVR8 SUPPRESSOR OF VARIEGATION 8_ plastid ribosomal protein L24 chr5:22183046-22184403 FORWAR |
| AT4G39990.1 | 1,767550703 | 0,031638772 | ATGB3_ RABA4B_ ATRABA4B_ ATRAB11G GTP-BINDING PROTEIN 3_ RAB GTPase homolog A4B_ ARABIDOP |
| AT1G48030.1 | 0,945211611 | 0,031932161 | mtLPD1 mitochondrial lipoamide dehydrogenase 1 chr1:17717432-17719141 REVERSE LENGTH=507 |
| AT5G66510.1 | 0,630016025 | 0,032564563 | GAMMA CA3 gamma carbonic anhydrase 3 chr5:26550016-26551496 REVERSE LENGTH=258 |
| AT1G79040.1 | 0,81902935 | 0,032869553 | PSBR photosystem II subunit R chr1:29736085-29736781 FORWARD LENGTH=140 |
| AT1G32900.1 | 0,823975048 | 0,033031902 | GBSS1 granule bound starch synthase 1 chr1:11920582-11923506 REVERSE LENGTH=610 |
| AT3G18070.3 | 0,683260511 | 0,033636469 | BGLU43 beta glucosidase 43 chr3:6187294-6189947 FORWARD LENGTH=495 |
| AT3G10060.1 | 0,786817687 | 0,033732859 | no symbol available no full name available chr3:3102291-3103801 FORWARD LENGTH=230 |
| AT3G54660.1 | 0,847758384 | 0,033789844 | EMB2360_ MIAO_ ATGR2_ GR2_ GR glutathione reductase_ GRISEA 2 chr3:20230356-20233100 REVERSE LENGTH |
| AT3G16420.1 | 0,790379228 | 0,034024006 | PBPI_ JAL30_ PBP1 PYK10-binding protein 1_ JACALIN-RELATED LECTIN 30 chr3:5579560-5580674 FORWARD L |
| AT3G09260.1 | 1,414122082 | 0,034033841 | LEB_ BGLU23_ PYK10_ PSR3.1 LONG ER BODY chr3:2840657-2843730 REVERSE LENGTH=524 |
| AT1G33590.1 | 0,853526006 | 0,034355523 | no symbol available no full name available chr1:12177788-12179221 FORWARD LENGTH=477 |
| AT3G46010.1 | 0,422476192 | 0,034429837 | atadf_ ADF1_ ATADF1 actin depolymerizing factor 1 chr3:16909679-16910678 REVERSE LENGTH=139 |
| AT4G30620.1 | 0,764480226 | 0,034717873 | STCL STIC2 Like chr4:14948724-14950035 REVERSE LENGTH=180 |
| AT1G59900.1 | 1,113619116 | 0,034896508 | AT-E1 ALPHA_ E1 ALPHA_ IAR4L pyruvate dehydrogenase complex E1 alpha subunit_ IAR4-LIKE chr1:22051368-220 |
| AT5G40370.1 | 1,107631275 | 0,035033836 | AtGRXC2_ GRXC2_ GRX370 glutaredoxin C2 chr5:16147826-16149052 REVERSE LENGTH=111 |
| AT1G09130.1 | 1,103719855 | 0,03518246 | no symbol available no full name available chr1:2940063-2942217 REVERSE LENGTH=330 |
| AT1G73060.1 | 1,277859753 | 0,035286782 | LPA3 Low PSII Accumulation 3 chr1:27479027-27481258 FORWARD LENGTH=358 |
| AT4G17520.1 | 0,725812122 | 0,035515513 | HLN HYALURONAN/mRNA BINDING FAMILY PROTEIN chr4:9771496-9773313 FORWARD LENGTH=360 |
| AT2G28900.1 | 1,187143184 | 0,035798096 | OEP16_ OEP16-1_ ATOEP16-L_ ATOEP16-1 outer plastid envelope protein 16-1_ OUTER PLASTID ENVELOPE PRO |
| AT5G62390.1 | 0,661166346 | 0,036496459 | ATBAG7_ BAG7 BCL-2-associated athanogene 7 chr5:25052377-25054170 REVERSE LENGTH=446 |
| AT4G12800.1 | 0,943747294 | 0,036670524 | PSAL photosystem I subunit l chr4:7521469-7522493 FORWARD LENGTH=219 |
| AT3G02230.1 | 1,122080891 | 0,036677442 | RGP1_ ATRGP1 ARABIDOPSIS THALIANA REVERSIBLY GLYCOSYLATED POLYPEPTIDE 1_ reversibly glycosyl |

| | | | |
|---|---|---|---|
| AT3G15060.1 | 6,295671008 | 0,037622237 | RABA1g_ AtRABA1g RAB GTPase homolog A1G chr3:5069239-5070025 FORWARD LENGTH=217 |
| AT3G48110.1 | 1,498505802 | 0,037629591 | EDD1_ EDD EMBRYO-DEFECTIVE-DEVELOPMENT 1 chr3:17763111-17770964 FORWARD LENGTH=1067 |
| AT1G64740.1 | 0,873480158 | 0,037763367 | TUA1 alpha-1 tubulin chr1:24050114-24052296 FORWARD LENGTH=450 |
| AT4G39260.1 | 1,162242447 | 0,038060423 | RBGA6_ CCR1_ ATGRP8_ GR-RBP8_ GRP8 "cold_ circadian rhythm_ and RNA binding 1"_ glycine-rich RNA-binding [illegible] |
| AT4G38630.1 | 3,114422263 | 0,038112153 | MBP1_ RPN10_ ATMCB1_ MCB1 MULTIUBIQUITIN CHAIN BINDING PROTEIN 1_ regulatory particle non-ATPase |
| AT1G15820.1 | 0,928275896 | 0,038540242 | CP24_ LHCB6 light harvesting complex photosystem II subunit 6 chr1:5446685-5447676 REVERSE LENGTH=258 |
| AT5G11670.1 | 1,154107381 | 0,038870427 | NADP-ME2_ ATNADP-ME2 NADP-malic enzyme 2_ Arabidopsis thaliana NADP-malic enzyme 2 chr5:3754456-375804[illegible] |
| AT1G79850.1 | 0,594936301 | 0,039269127 | RPS17_ PDE347_ CS17_ PRPS17 PIGMENT DEFECTIVE 347_ PLASTID RIBOSOMAL SMALL SUBUNIT PROTEIN |
| ATCG00160.1 | 1,087753282 | 0,039673561 | RPS2 ribosomal protein S2 chrc:15013-15723 REVERSE LENGTH=236 |
| AT3G17020.1 | 1,245995568 | 0,040048092 | no symbol available no full name available chr3:5802728-5804063 REVERSE LENGTH=163 |
| AT5G02940.1 | 0,861791071 | 0,04024861 | PEC1 PLASTID ENVELOPE ION CHANNELS 1 chr5:684671-689674 REVERSE LENGTH=813 |
| AT3G05350.1 | 0,604002775 | 0,040636554 | no symbol available no full name available chr3:1527103-1533843 REVERSE LENGTH=710 |
| AT5G14030.5 | 1,317720172 | 0,040682007 | no symbol available no full name available chr5:4526878-4527917 FORWARD LENGTH=159 |
| AT5G47870.1 | 1,158228805 | 0,040836966 | RAD52-2_ ODB2_ RAD52-2B radiation sensitive 52-2_ Organellar DNA-Binding protein 2 chr5:19384555-19385808 RE[illegible] |
| AT5G64140.1 | 1,305473328 | 0,040891615 | RPS28 ribosomal protein S28 chr5:25667529-25667723 REVERSE LENGTH=64 |
| AT4G20260.1 | 0,934580419 | 0,040977157 | ATPCAP1_ PCAP1_ MDP25 ARABIDOPSIS THALIANA PLASMA-MEMBRANE ASSOCIATED CATION-BINDING |
| AT2G43950.1 | 1,552395169 | 0,041309684 | ATOEP37_ OEP37 chloroplast outer envelope protein 37_ ARABIDOPSIS CHLOROPLAST OUTER ENVELOPE PROT[illegible] |
| AT1G04040.1 | 3,01517957 | 0,041344836 | no symbol available no full name available chr1:1042564-1043819 REVERSE LENGTH=271 |
| AT1G13280.1 | 1,761579476 | 0,041533011 | AOC4 allene oxide cyclase 4 chr1:4547624-4548552 FORWARD LENGTH=254 |
| AT3G06860.1 | 1,472839882 | 0,041612702 | ATMFP2_ MFP2 MULTIFUNCTIONAL PROTEIN 2_ multifunctional protein 2 chr3:2161926-2166009 FORWARD LE[illegible] |
| AT4G24770.1 | 1,103497974 | 0,041701295 | ATRBP33_ ATRBP31_ CP31A_ RBP31_ CP33a_ CP31 "ARABIDOPSIS THALIANA RNA BINDING PROTEIN_ APP[illegible] |
| AT4G28440.1 | 1,161439296 | 0,041778143 | no symbol available no full name available chr4:14060054-14060970 FORWARD LENGTH=153 |
| AT3G04550.1 | 0,576334456 | 0,041839876 | RAF1 Rubisco accumulation factor 1 chr3:1225961-1227310 FORWARD LENGTH=449 |
| AT3G15360.1 | 0,934515158 | 0,042374769 | ATM4_ TRX-M4_ ATHM4 ARABIDOPSIS THIOREDOXIN M-TYPE 4_ thioredoxin M-type 4 chr3:5188448-5189457 [illegible] |
| AT5G12470.1 | 0,848028656 | 0,043089926 | RER4 RETICULATA-RELATED 4 chr5:4044950-4047290 REVERSE LENGTH=386 |
| AT4G14160.3 | 1,686512687 | 0,043446427 | AtSEC23F chr4:8167574-8172266 FORWARD LENGTH=620 |
| AT4G29410.1 | 0,752623098 | 0,043468045 | no symbol available no full name available chr4:14468439-14469964 REVERSE LENGTH=143 |
| AT1G02930.1 | 0,6044382 | 0,043667893 | ATGSTF3_ ATGSTF6_ GST1_ ERD11_ GSTF6_ ATGST1 EARLY RESPONSIVE TO DEHYDRATION 11_ ARABID[illegible] |
| AT1G09010.1 | 1,238395585 | 0,043740097 | no symbol available no full name available chr1:2895259-2899287 REVERSE LENGTH=944 |
| AT2G47400.1 | 0,899397968 | 0,043788542 | CP12_ CP12-1 CP12 DOMAIN-CONTAINING PROTEIN 1_ CP12 domain-containing protein 1 chr2:19446889-1944726[illegible] |
| AT4G00430.2 | 2,316495749 | 0,043806818 | PIP1E_ TMP-C_ PIP1;4 plasma membrane intrinsic protein 1;4_ PLASMA MEMBRANE INTRINSIC PROTEIN 1E_ TR[illegible] |
| AT4G25080.1 | 0,87621126 | 0,04386636 | CHLM magnesium-protoporphyrin IX methyltransferase chr4:12877015-12878128 FORWARD LENGTH=312 |
| AT1G10670.3 | 1,160879272 | 0,044121061 | ACLA-1 ATP-citrate lyase A-1 chr1:3535787-3538098 FORWARD LENGTH=443 |
| AT3G02780.2 | 0,734277055 | 0,044266668 | IDI2_ IPIAT1_ IPP2 isopentenyl pyrophosphate:dimethylallyl pyrophosphate isomerase 2_ ATISOPENTENYL DIPHOSP[illegible] |
| AT1G66670.1 | 1,103890174 | 0,044332837 | NCLPP3_ CLPP3 CLP protease proteolytic subunit 3 chr1:24863995-24865646 REVERSE LENGTH=309 |
| AT1G03860.1 | 1,451190497 | 0,044601276 | PHB2_ ATPHB2 prohibitin 2 chr1:979611-981157 REVERSE LENGTH=286 |
| AT4G23600.1 | 0,769102213 | 0,044851212 | CORI3_ JR2 JASMONIC ACID RESPONSIVE 2_ CORONATINE INDUCED 1 chr4:12310657-12312885 FORWARD L |
| AT2G20260.1 | 1,229265747 | 0,045335236 | PSAE-2 photosystem I subunit E-2 chr2:8736780-8737644 FORWARD LENGTH=145 |
| AT1G80600.1 | 3,909099962 | 0,045884844 | TUP5_ WIN1 HOPW1-1-interacting 1_ TUMOR PRONE 5 chr1:30298675-30300513 REVERSE LENGTH=457 |

| | | | |
|---|---|---|---|
| AT5G51970.1 | 0,869482076 | 0,04603071 | ATSDH SORBITOL DEHYDROGENASE chr5:21111820-21113284 FORWARD LENGTH=364 |
| AT5G11520.1 | 0,873034649 | 0,046045683 | YLS4_ ASP3 aspartate aminotransferase 3_ YELLOW-LEAF-SPECIFIC GENE 4 chr5:3685257-3687721 REVERSE LEN |
| AT5G47840.2 | 0,708935121 | 0,04608259 | AMK2 adenosine monophosphate kinase chr5:19375488-19378058 FORWARD LENGTH=269 |
| AT4G30610.1 | 0,622028263 | 0,046642561 | BRS1_ SCPL24 BRI1 SUPPRESSOR 1_ SERINE CARBOXYPEPTIDASE 24 PRECURSOR chr4:14944219-14948391 F |
| AT4G10480.1 | 0,83112965 | 0,047145032 | no symbol available no full name available chr4:6478089-6479079 REVERSE LENGTH=212 |
| AT2G40100.1 | 0,815798696 | 0,047209808 | LHCB8_ LHCB4.3 light harvesting complex photosystem II chr2:16745884-16747190 FORWARD LENGTH=276 |
| AT3G09810.1 | 1,244846203 | 0,047355094 | IDH-VI isocitrate dehydrogenase VI chr3:3008753-3011070 FORWARD LENGTH=374 |
| AT5G13450.1 | 1,231810767 | 0,047958078 | ATP5 delta subunit of Mt ATP synthase chr5:4310558-4311941 REVERSE LENGTH=238 |
| AT4G17170.1 | 1,505110011 | 0,048459616 | RAB2A_ AT-RAB2_ ATRAB2A_ ATRABB1C_ RAB-B1B_ ATRAB-B1B_ RABB1C ARABIDOPSIS RAB GTPASE H |
| AT5G06320.1 | 0,629112422 | 0,048523273 | NHL3 NDR1/HIN1-like 3 chr5:1931016-1931711 REVERSE LENGTH=231 |
| AT4G33510.1 | 0,883198774 | 0,048949333 | DAHP2_ AtDAHP2_ DHS2 3-deoxy-d-arabino-heptulosonate 7-phosphate synthase_ 3-DEOXY-D-ARABINO-HEPTULO |
| AT5G64050.1 | 5,678639737 | 0,048956026 | ATERS_ OVA3_ ERS glutamate tRNA synthetase_ OVULE ABORTION 3 chr5:25630196-25633099 REVERSE LENGT |
| AT5G46290.2 | 0,870979727 | 0,048965377 | KASI_ KAS1 3-ketoacyl-acyl carrier protein synthase I_ KETOACYL-ACP SYNTHASE 1 chr5:18774439-18776629 REV |
| AT4G23400.1 | 2,664941881 | 0,049767732 | PIP1D_ PIP1;5 plasma membrane intrinsic protein 1;5 chr4:12220792-12222155 FORWARD LENGTH=287 |
| AT1G04420.1 | 1,417288492 | 0,04985149 | no symbol available no full name available chr1:1191634-1193699 FORWARD LENGTH=412 |
