## Supplementary material for "HY5 enhances *Arabidopsis* tolerance to combined high light and heat stress by coordinating photoprotection and hormone signaling": Figure S1

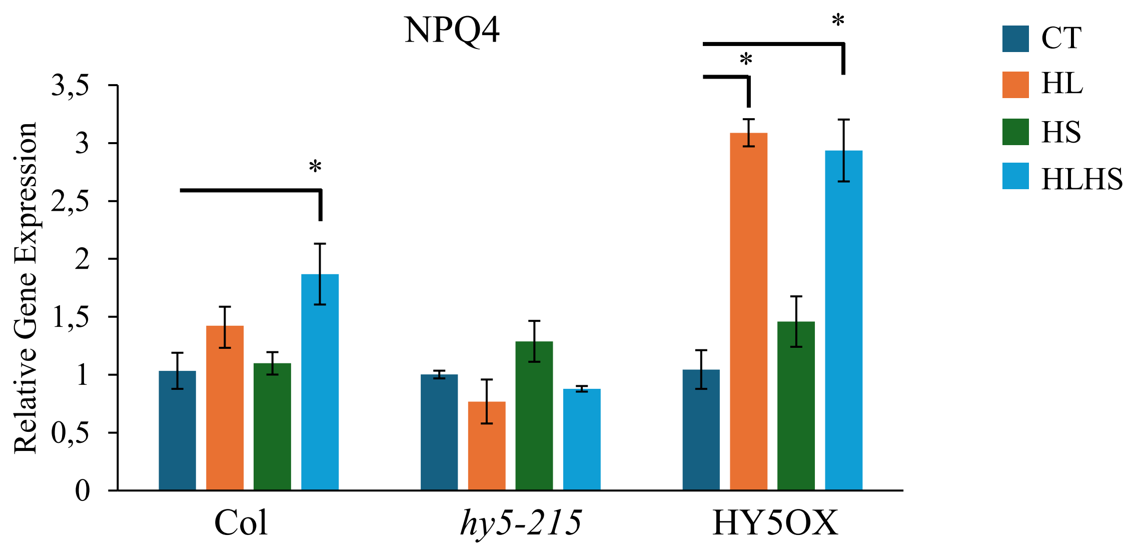


**Figure S1.** Relative expression levels of the NPQ4 gene in Col-0, *hy5-215*, and HY5OX plants under control (CT), high light (HL), heat stress (HS), and combined high light and heat stress (HLHS) conditions. Expression values were normalized to control levels for each genotype. Error bars represent mean ± standard error (SE) (n = 9).

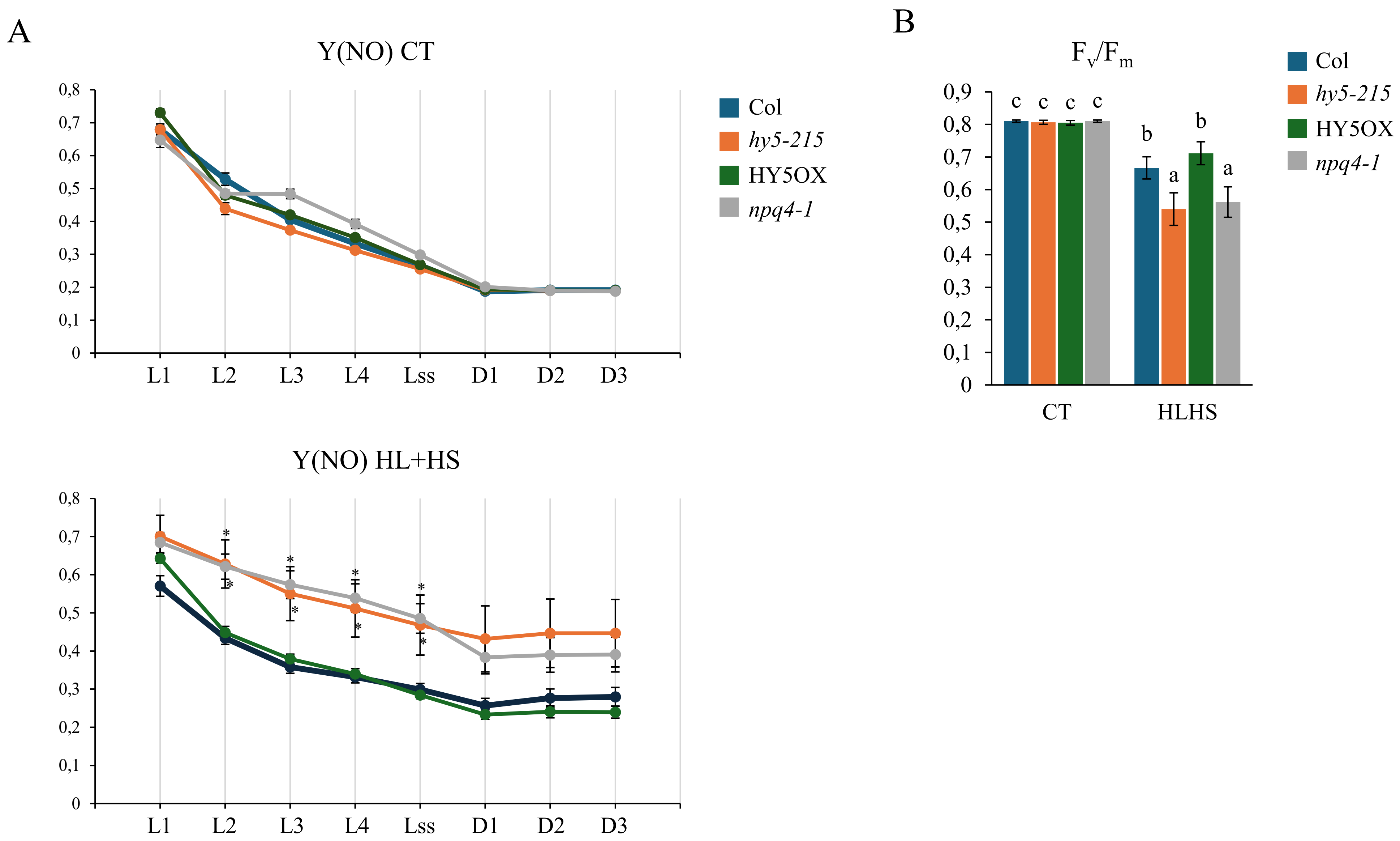


**Figure S2. Kinetics of PSII photochemical parameters in Col-0, *hy5-215*, HY5OX, and *npq4-1* plants under CT and HLHS conditions.** (A) Non-regulated energy dissipation [Y(NO)] measured over a series of actinic light pulses and dark relaxation points (L1–Lss, D1-D3) in all genotypes under CT and HLHS conditions. (B) Maximum quantum efficiency of PSII (Fv/Fm) in all genotypes under CT and after HLHS treatment. Error bars represent mean ± standard error (SE) (n = 9).
